# The human sperm basal body is a complex centrosome important for embryo pre-implantation development

**DOI:** 10.1101/2021.04.11.439346

**Authors:** Farners Amargant, Aïda Pujol, Anna Ferrer-Vaquer, Mercè Durban, Meritxell Martínez, Rita Vassena, Isabelle Vernos

## Abstract

The mechanism of conversion of the human sperm basal body to a centrosome after fertilization, and its role in supporting human early embryogenesis has not been directly addressed so far. Using proteomics and immunofluorescence studies we show here that the human zygote inherits a basal body enriched with centrosomal proteins from the sperm, establishing the first functional centrosome of the new organism. Injection of human sperm tails containing the basal body into human oocytes followed by parthenogenetically activation, showed that the centrosome contributes to the robustness of the early cell divisions, increasing the probability of parthenotes reaching the compaction stage. In the absence of the sperm-derived centrosome, pericentriolar (PCM) components stored in the oocyte can form *de novo* structures after genome activation, suggesting a tight PCM expression control in zygotes. Our results reveal that the sperm basal body is a complex organelle which converts to a centrosome after fertilization, ensuring the early steps of embryogenesis and successful compaction. However, more experiments are needed to elucidate the exact molecular mechanisms of centrosome inheritance in humans.

## Introduction

The centrosome is the main microtubule organizing center (MTOC) of the cell. It consists of two centrioles surrounded by a mass of proteins known as pericentriolar material (PCM) (Rusan and Rogers, 2009). Centrosomes are important for intracellular organization, spindle assembly, asymmetric cell division and polarity establishment as well as for the assembly of cilia and flagella. They are therefore essential for tissue architecture and function and the development of healthy organisms (Wu and Akhmanova, 2017). Defects in centrosome function and/or number are associated with conditions including cancer, ciliopathies and infertility. In cycling cells, the number of centrosomes is tightly controlled in a cell cycle dependent manner. Centrioles duplicate once (and only once) during interphase, and in mitosis the duplicated centrosomes localize each at one pole of the bipolar spindle defining spindle organization and orientation. After cell division, each daughter cell inherits one of the two centrosomes (Nigg and Stearns, 2011).

Fertilization entails the fusion of the male and the female gametes. In mammals, the regulation of the centrosome number *per* cell is established during fertilization. Indeed, human oocytes do not contain centrioles (Sathananthan, 1997) while the sperm has two centrioles (proximal and distal centrioles) in the midpiece that function as the basal body of the flagella (Familiari *et al*., 2006). After fertilization, the paternal pronucleus is associated with a centriole, suggesting that the centrosome is paternally inherited during fertilization (Sathananthan *et al*., 1991, Simerly *et al*., 1995, Van Blerkom and Davis, 1995). Studies on fertilized eggs from various species that also inherit the centrosome paternally (i.e. rhesus monkeys, drosophila, and cow) showed that the sperm basal body suffers a process known as “centrosome reduction” (Manandhar *et al*., 2000). This process consists in the elimination of the PCM and the remodeling of the centriolar microtubules generating an atypical centrosome. The current model is that upon fertilization, the sperm atypical centrioles recruit PCM components stored in the oocyte cytoplasm to assemble a functional centrosome. A very limited number of studies have addressed this process in humans, except for a few reports showing that the human sperm shares many characteristics with those of other species (Fishman *et al*., 2018, Manandhar *et al*., 2000, Sathananthan *et al*., 1996, Simerly *et al*., 1999). Indeed, the human sperm distal centriole is also remodeled, although it retains a subset of centrosomal proteins, suggesting that sperm PCM reduction is not completed in humans (Fishman *et al*., 2018). Nevertheless, the degree of conservation or loss of PCM components at the human sperm basal body, and the functional implications of these proteins for early embryonic development are still unresolved questions.

The oocyte cytoplasm provides a large store of maternal components, including proteins and mRNAs that are essential for the first embryonic divisions, that occur in the absence of transcription (Conti and Franciosi, 2018). Little is known about the storage of maternal PCM components in oocytes and how they contribute to the transition of the sperm basal body into the centrosome of the zygote. Many of these studies have been performed in mice and, although this is an excellent model to study early embryonic development, it is not ideal to study the centrosome because these phases occur in the absence of centrioles, which form *de novo* at the blastocyst stage in mice (Hiraoka *et al*., 1989). The early embryonic cell divisions in mice are instead supported by acentriolar MTOCs (aMTOCs) (Courtois *et al*., 2012, Gueth-Hallonet *et al*., 1993, Howe and FitzHarris, 2013). This suggests that although centrosomes are essential for healthy organisms, they may not be needed during the early embryo cell divisions, which may rely on spindle assembly acentrosomal pathways acting during oocyte meiosis.

In this work, we first determined the protein composition of the human sperm basal body by proteomics and super-resolution microscopy, and then devised a functional approach to study the role of the sperm derived centrosome in the early cell divisions of parthenotes. Our results suggest that the human sperm provides the zygote with centrioles and an important mass of centrosomal proteins, which may be important for the assembly of the first functional centrosome of the new organism. Furthermore, our data suggest that the sperm derived centrosome increases the robustness of the first cell divisions of the parthenotes leading to compaction. Our data also suggest that the expression of centrosomal proteins in the early parthenotes has to be tightly regulated to avoid the untimely formation of spontaneous MTOC-like aggregates in the embryonic cells. These findings not only improve our understanding of the centrosome biology during fertilization and early development, but also provide novel putative avenues to explore causes for early embryo arrests and idiopathic infertility in humans.

## Materials and methods

### Ethics

Approval to conduct this study was obtained from the ethical committee of the Parc de Recerca Biomèdica of Barcelona (PRBB), Barcelona, Spain. All procedures performed were in accordance with the ethical standards of the institutional research committees and with the 1964 Helsinki declaration of the Ethical principles for medical research involving human subjects, as revised in 2013 in Fortaleza (World Medical, 2013). Written informed consents to participate were obtained from all participants prior to their inclusions in the studies.

### Sperm thawing and swim-up

To thaw sperm samples, straws were incubated at 37°C for 5 min. To remove the sperm cryoprotectant, the sample was diluted with the same volume of Sperm Rinse (Vitrolife, Göteborg) and centrifuged for 5 min at 300 g at room temperature. Then, the pellet was washed twice with 1 ml of Sperm Rinse (Vitrolife, Göteborg) and centrifuged at 372 g 5 min at room temperature. To perform the sperm swim-up, 0.1 ml of Sperm Rinse was carefully loaded on the top of the pellets. The tubes were oriented at 45° for 10 min. During this time, the motile spermatozoa swam to the Sperm Rinse fraction. Finally, the Sperm Rinse fraction was carefully collected avoiding the aspiration of the pellet. For the tails-injected experiment, after thawing the sperm straw, the sample was diluted with 3 ml of Sperm Rinse and centrifuged once at 300 g for 5 min. Right after this wash, the sperm swim-up was performed as previously described.

### Sperm centriole enrichment

Two different methods were used to analyze sperm centrosomal composition. Only normozoospermic samples with >50% of A+B motility were used. The first approach was based on a previously published protocol (Firat-Karalar *et al*., 2014) but with some modifications. Frozen samples were thawed and washed twice with PBS and checked under the microscope for any defects caused by the freezing and/or thawing cycles (if any defect was detected, they were discarded). The total amount of cells used at each experiment was of ≈60 millions. Washed samples were pelleted at 850 g, 10 min, and resuspended with 350 μl of PBS. Then, samples were sonicated 5 times at 70% output for 15 seconds with 30 seconds intervals (Bioruptor). 1 μl of each sample was taken to check the sonication efficiency under a bright-field microscope. Sperm tails were separated from the heads through a 30% sucrose cushion (centrifugation at 200g for 10 min at 4°C). 1 μl of the tails fraction was taken and squashed in an 18×18 mm coverslip to check its purity. To sequentially extract sperm tails proteins, the tails fraction was diluted with the same volume of Buffer 1 (100 mM Tris HCl p.H. 8, 4 mM EGTA, 4 mM EDTA, 1000 mM NaCl, 2% NP40, 0.2%ß-mercaptoethanol, 2mM DTT, protease inhibitors in dH_2_O) and incubated 1 h at 4°C under movement. After this incubation, the sample was centrifuged at maximum velocity for 10 min at 4°C. The pellet and the supernatant were separated in two different tubes to be treated differentially. In the supernatant, the proteins solubilized by buffer 1 are found. 125 μl of 100% trichloroacetic acid was added to the supernatant and kept at 4°C for 10 min. To pellet the precipitated proteins, the sample was centrifuged at maximum velocity at 4°C, 5 min. The supernatant was removed, and the pellet was washed twice with cold acetone and then dried at 95°C. Finally, the pellet was diluted with 6 M urea and 200 mM of ABC (ammonium bicarbonate, Sigma, MO, USA) in ddH_2_O. The pellet fraction from the first extraction was sequentially extracted with extraction buffer 2 (50 mM Tris HCl p.H. 8, 600 mM KSCN, 2 mM DTT, protease inhibitors in dH_2_O) and extraction buffer 3 (50 mM Tris HCl p.H. 8, 4 M urea, protease inhibitors in dH_2_O). 3 different extractions diluted in 6 M urea and 200mM ABC were finally obtained for proteomic analysis.

For the second approach, sonicated tails were pelleted at 4°C at the maximum velocity and diluted with LB (Loading Buffer, 2% w/v SDS, 10% Glycerol, 50 mM Tris-HCl p.H. 6.8, 5%ß-mercaptoethanol). Samples were run on a precasted gradient gel (4 – 15% Criterion TGX, 12+2 wells, 45 μl, BioRad, CA, USA) for 45 min at 60 mA. Gels were stained for 10 min using Coomassie (Coomassie Brilliant Blue R250 (Thermo Fisher Scientific, WA, USA), 10% acetic acid, 50% methanol) and distained with 10% methanol and 10% acetic acid. Gels were cut into 9 bands to process for proteomic analysis.

### Mass spectrometry

In solution samples were reduced with 10 mM DTT, 37°C, 60 min and alkylated in the dark with iodoacetamide (IAM, 20 mM, 25 °C, 30 min). The resulting protein extract was first diluted to 2M urea for overnight digestion with LysC (Wako, USA) at 37°C and then diluted 2-fold for 8 h digestion with trypsin (Promega, USA) at 37°C.

Gels band samples were destained with 40% ACN/100mM ABC, reduced with DTT, 10 mM, 56 °C, 30 min and alkylated in the dark with iodoacetamide (IAM, 55 mM, 25°C, 30 min). Gel bands were then dehydrated with ACN and digested overnight with trypsin at 37°C.

After digestion, peptide mix was acidified with formic acid and desalted with a MicroSpin C18 column (The Nest Group, Inc, MA, USA) prior to LC-MS/MS analysis. Samples were analyzed using an LTQ-Orbitrap Velos Pro mass spectrometer (Thermo Fisher Scientific, WA, USA) coupled to an EasyLC (Thermo Fisher Scientific (Proxeon), Odense, Denmark). Peptides were loaded onto the 2-cm Nano Trap column with an inner diameter of 100 μm packed with C18 particles of 5 μm particle size (Thermo Fisher Scientific, WA, USA) and were separated by reversed-phase chromatography using a 25-cm column with an inner diameter of 75 μm, packed with 1.9 μm C18 particles (Nikkyo Technos Co., Ltd. Japan). Chromatographic gradients started at 93% buffer A and 7% buffer B with a flow rate of 250 nl/min for 5 minutes and gradually increased 65% buffer A and 35% buffer B in 60 min for the in solution samples and 120 min for the gels. After each analysis, the column was washed for 15 min with 10% buffer A and 90%buffer B. Buffer A: 0.1% formic acid in water. Buffer B: 0.1% formic acid in acetonitrile.

The mass spectrometer was operated in DDA mode and full MS scans with 1 micro scans at resolution of 60.000 were used over a mass range of m/z 350-2000 with detection in the Orbitrap. Auto gain control (AGC) was set to 1E6, dynamic exclusion (60 seconds) and charge state filtering disqualifying singly charged peptides was activated. In each cycle of DDA analysis, following each survey scan the top twenty most intense ions with multiple charged ions above a threshold ion count of 5000 were selected for fragmentation at normalized collision energy of 35%. Fragment ion spectra produced via collision-induced dissociation (CID) were acquired in the Ion Trap, AGC was set to 5e4, isolation window of 2.0 m/z, activation time of 0.1 ms and maximum injection time of 100 ms was used. All data were acquired with Xcalibur software v2.2.

For peptide identification, proteome Discoverer software suite (v1.4, Thermo Fisher Scientific, WA, USA) and the Mascot search engine (v2.5, Matrix Science (Perkins *et al*., 1999)) were used. Samples were searched against a Swiss-Prot human database plus a list of common contaminants and all the corresponding decoy entries (20797 entries). Trypsin was chosen as enzyme and a maximum of three miscleavages were allowed. Carbamidomethylation (C) was set as a fixed modification, whereas oxidation (M) and acetylation (N-terminal) were used as variable modifications. Searches were performed using a peptide tolerance of 7 ppm, a product ion tolerance of 0.5 Da. Resulting data files were filtered for FDR < 5 %.

### Oocyte warming

Oocytes were warmed according to standard procedure (Cryotop, Kitazato, BioPharma Co., Ltd; Japan). Briefly, TS medium was prewarmed at 37°C for 45 min prior to use. DS and WS media were used at room temperature. The oocyte straw cap was carefully removed and then quickly immersed in a dish with 1 ml of TS medium for 1 min. Then, oocytes were incubated in DS medium for 3 min. Finally, oocytes were placed to the WS medium for 5 min and transferred to a second WS containing plate for 1 min before incubating them 2h at 37°C, 6% CO_2_ atmosphere to let them recover from warming.

### Tails injection

In an ICSI (Intracytoplasmic Sperm Injection) dish of 60mm of diameter, several drops of G-MOPS plus containing an oocyte each (Vitrolife, Gothenburg, Sweden) surrounded a central PVP drop (polyvinylpyrrolidone; Origio, Malov, Denmark) containing the sperm swim-up sample. The plate was overlaid with mineral oil (OVOIL^TM^, Vitrolife, Gothenburg, Sweden) to prevent the evaporation of the drops and then placed on the ICSI microscope stage (Olympus IX50, Olympus, Tokio, Japan). Two different pipettes were needed to perform tails separation and injection into oocytes. A PZD (Partial Zona Dissection, Vitrolife, Gothenburg, Sweden) pipette was used for the sperm head-tail separation, and an ICSI micropipette (Vitrolife, Gothenburg, Sweden) to perform the tail injection into the oocyte. Both pipettes were located in a double needle holder. The PZD pipette was placed in the interface between the sperm head and the midpiece. With an accurate blow the separation of both parts was achieved. Immediately after the separation, the tails were collected. To confirm that the separated tails had the centrosome and not DNA, tails were aspirated with the ICSI micropipette and loaded onto a glass microscope slide (25 mm x 75 mm) and fixed with 4% PFA to immunodetect centrosomes and DNA. In all the cases, only one tail was injected per oocyte. A certain number of oocytes were sham injected as controls.

### Oocyte activation

Tail-injected and control oocytes were washed 4 times with G1-PLUS^TM^ medium (Vitrolife, Gothenburg, Sweden) and incubated during 30 min at 37°C, 6% CO_2_. The oocyte activation protocol or AOA included three 10 minutes incubations in 10μM ionomycin (calcium ionophore, MP Biomedicals, CA, USA), and three 30 minutes washes in G1-PLUS^TM^ medium (Vitrolife, Gothenburg, Sweden) at 37°C, 6% CO_2_. All the plates were covered with OVOIL^TM^. Finally, oocytes were transferred in a plate with SAGE medium (Origio, Malov, Denmark) at 37°C, 6% CO_2_ to be placed in the time-lapse system Primovision or Embryoscope (Vitrolife, Gothenburg, Sweden). These two equipments took images of each sample every 5 min, allowing for morphologic and the kinetic analyses.

### DNA cloning and transfection

DNA sequences of the 10 uncharacterized proteins were obtained from the ORFeome service (Centre for Genomic Regulation). These sequences were cloned into a pDEST vector that contained either CFP at the N-terminus (Kanamycin resistant) or GFP at the C-terminus (Ampicillin resistant).

HeLa cells were grown at 37°C in a 5% CO_2_ atmosphere in DMEM 4.5g/L Glucose, supplemented with Ultraglutamine (Lonza, Basilea, Switzerland), with 10% FBS (Fetal Bovine Serum, Invitrogen, CA, USA), 100 units/ml of penicillin and 100 μg/ml of streptomycin. HeLa cells were regularly controlled for mycoplasma contamination. DNA were transfected with the same volume of X-tremeGENE (Sigma, MO, USA) and 100 μl of Opti-MEM (Thermo Fisher Scientific, WA, USA) using 500ng of DNA per well in a 12 well plate (150,000 cells/well) of HeLa cells already attached to a glass coverslip (18 mm diameter). After 24h of protein expression (at 37°C in a 5% CO_2_ atmosphere), cells were collected and washed twice with 1 ml of PBS prior fixation with cold methanol (10 min) or 4% PFA (15 min. Sigma, MO, USA).

### Sperm immunofluorescence (IF)

Thawed sperm samples were washed twice with PBS and then loaded onto a poly-L-lysine coated glass coverslip (12 mm of diameter – from 50,000 to 100,000 cells). After 30 min, the supernatant was removed and the cells were fixed with either 4% PFA 1 hour, or methanol for 10 min. After several washes with PBS, samples were permeabilized with 0.5% Triton X-100 PBS 15 min at room temperature. Samples were loaded with 5% BSA in PBS to block unspecific interactions for 2 h. The same blocking solution was used to incubate the primary and the secondary antibody for 1 h and 45 min respectively and mounted in 10% Mowiol (Sigma, MO, USA) in 0.1M TrisHCl at pH 8.2, 25% glycerol. Confocal images were obtained using a TCS-SP5 microscope (Leica Microsystems, Wetzlar, Germany) in a 63x objective. Lasers and spectral detection bands were chosen for the optimal imaging of Alexa Fluor 488 and 568 signals. Two-channel colocalization analysis was performed using ImageJ (National Institutes of Health, MD, USA). STED images were taken on a TCS SP8 STED3X microscope (Leica Microsystems, Wetzlar, Germany).

### Oocyte and parthenotes IF

In a prewarmed plate (37°C) oocytes and parthenotes were mixed with Tyrode’s (Sigma, MO, USA) to remove the zona pellucida. Then, a quick wash with prewarmed PBS was performed. Samples were then fixed with prewarmed 4% of PFA, 15 min. Oocytes and parthenotes were permeabilized with PBS 0.2% of Triton X-100 during 15 min. After washing the samples once with PBS-T (PBS, 0.1% Tween 20) and 3 more times with PBS-TB (PBS, 0.1% Tween20, 2% BSA Fraction V) for 20 minutes each, samples were blocked with 5% normal goat serum (Vector Laboratories, CA, USA) in PBS-TB (freshly prepared) for 3 h. All primary antibodies were incubated over-night at 4°C with the blocking solution. Then, 3 washes of 20 min in PBS-TB were done while shaking to eliminate the remaining primary antibody. The secondary antibody was only incubated 1 h at room temperature in PBS-TB together with Hoechst 33342 (1 μg/ml, Invitrogen, CA, USA). Samples were mounted with Vectashield (Vector Laboratories, CA, USA) and visualized at the Zeiss 780 confocal/multiphoton microscopy at 63X with 80% glycerol (Carl Zeiss, Oberkochem, Germany).

### Cell culture IF

HeLa transfected cells were block and permeabilized at the same time with IF medium (0.1% Triton X-100, 2% BSA in PBS 1x) during 30 min. The primary and the secondary antibodies diluted in IF (0.1% Triton X-100, 2% BSA in PBS 1x) medium were placed onto the samples for 50 and 45 min, respectively. The mounted samples with 10% Mowiol (Sigma, MO, USA) were visualized at 63X or with the Leica TCS SP5 upright microscope (Leica Microsystems, Wetzlar, Germany). Two-channel colocalization analysis was performed using ImageJ (National Institutes of Health, MD, USA).

### SDS-PAGE and Western Blot

Sperm extracts (intact, heads and tails fractions) were obtained diluting the samples with 1x LB and freezing and boiling the extracts 3 times. Then, samples were run in a 4-20% Protean TGX Precast protein gels (Biorad, CA, USA). To transfer proteins, the iBlot dry system (Thermo Fisher Scientific, WA, USA) was used. PVDF membranes were blocked with TBS 5% milk 1 h. The primary and secondary antibodies were diluted with 2% milk in TBS and incubated over-night at 4°C and 1 h at room temperature, respectively. Blots were developed using the Odyssey Infrared imaging system (LI-COR Biosciences, NE, USA).

### Antibodies

The following commercial primary antibodies were used: rabbit anti-centrin (Merck Millipore, MA, USA, 20H5) at 1:100, mouse anti-acetylated tubulin (Sigma, MO, USA, T7451) at 1:1000, rabbit anti-Cep63 (Merck Millipore, MA, USA, 06-1292) at 1:100, rabbit anti-Pericentrin (Abcam, Cambridge, UK, ab448) at 1:500, mouse anti-protamine (Novus Biologicals, CO, USA, H00005619) at 1:100, rabbit anti-β-tubulin (Abcam, Cambridge, UK, ab6046) at 1:1000, mouse anti-α-tubulin (Sigma, MO, USA, DM1A T6199) at 1:1000 in sperm and at 1:100 in oocytes and parthenotes, mouse anti-γ-tubulin (Sigma, MO, USA, GTU-448) at 1:100. Secondary antibodies anti-rabbit and anti-mouse conjugated to Alexa-488 and 568 (Invitrogen, CA, USA) were used at 1:1000 in sperm and cell culture, and 1:100 in oocyte and parthenotes for IF and 680 (Invitrogen, CA, USA) or IRdye 800 CW (LI-COR Biosciences, NE, USA) at 1:10000 for WB. Hoechst 33342 (1 μg/ml, Invitrogen, CA, USA) was used to visualize DNA.

### Primo-vision/Embryoscope analysis

To analyze the kinetics of parthenotes development, the time of cell division and compaction, early pseudo-blastocyst and expanded pseudo-blastocyst was measured in minutes and transformed to hours.

### Gene Ontology enrichment and STRING analysis

The Gene Ontology analyses were performed using the “Gene Ontology Consortium” (http://geneontology.org/page/go-enrichment-analysis). Data was confirmed using Uniprot (https://www.uniprot.org) and Protein Atlas (https://www.proteinatlas.org) databases. STRING (version 11.0) was used to perform the protein-protein interaction network (https://string-db.org).

## Results

### The human sperm centrioles are associated with centrosomal proteins

The human sperm basal body consists of a proximal centriole with a conventional microtubule-based organization and a distal centriole that has been described as a degenerated centriole (Avidor-Reiss *et al*., 2015). In order to get further insights on the nature of these centrioles, we determined centrin and acetylated α-tubulin localization patterns with super-resolution microscopy (STED – Stimulated Emission Depletion). Centrin is a structural protein that localizes inside the centrioles and is often used as a centriolar marker (White *et al*., 2000). We found that centrin localizes to both centrioles **(Figure 1A)**, suggesting that the distal centriole still retains some basic intrinsic features of a conventional centriole. Acetylated α-tubulin is usually found in long-lived microtubules such as the centriole microtubules and they are considered a marker of their structural stability (Amargant *et al*., 2019). We found that both centrioles contain acetylated tubulin suggesting that they both are stable microtubule assemblies **(Figure 1B)**. Together, our results suggest that the centrioles of the human sperm basal body retain some of the basic features of conventional centrioles.

**Figure 1:**
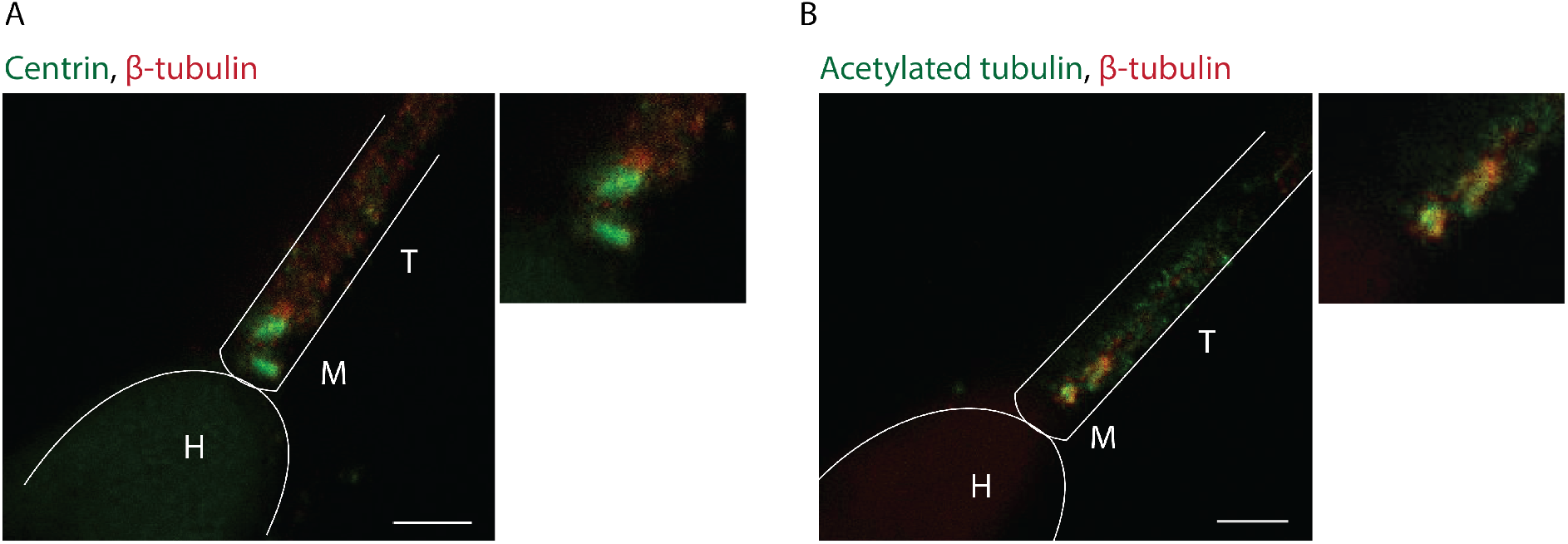
Sperm centrioles are stable structures. **A)** Super-resolution imaging (STED) of human spermatozoa stained for centrin and β-tubulin. Scale bar: 1 μm. **B)** Human sperm IF of acetylated tubulin and β-tubulin in the human sperm centrioles visualized with STED. Scale bar: 1 μm. H: Head, M: Midpiece, T: Tail. N = 2 different sperm samples.

To check whether the sperm centrioles are associated with centrosomal proteins we performed IF for Cep63, a protein involved in the centrosome duplication cycle (Brown *et al*., 2013, Watanabe *et al*., 2016). We found that Cep63 localized to both centrioles in the human sperm **(Figure 2A).** These data, in agreement with previous reports (Fishman *et al*., 2018), suggest that the human sperm basal body is associated with centrosomal proteins. Altogether we conclude that the human sperm basal body includes two centrioles that show some basic features of conventional centrosomes suggesting that it does not undergo a full process of centrosome reduction (Manandhar and Schatten, 2000, Manandhar *et al*., 2000).

**Figure 2:**
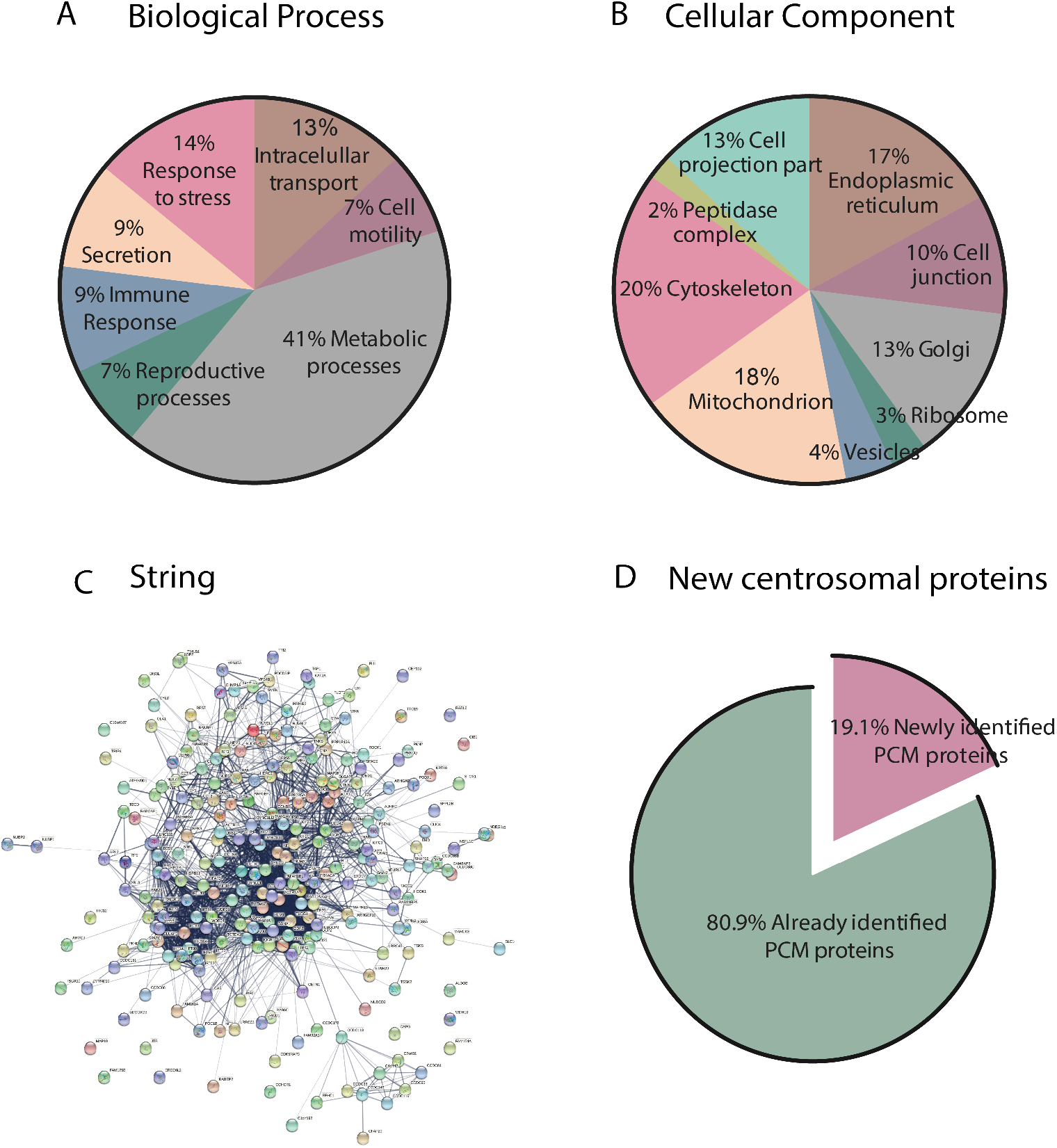
Sperm centrosome enrichment to identify centrosomal components. **A)** IF images on human sperm to visualize Cep63, α-tubulin and DNA. Scale bar: 5 μm. N = 3 different sperm samples. **B)** Schematic representation of the sonication and enrichment protocol. Only normozoospermic samples with ≥50% of A+B motility were used. N = 3 different experiments. **C)** Western Blot analysis of cell lysates from the different sperm fractions shown in B to detect heads (protamine 1) and tails (γ-tubulin). N = 3 different experiments. **D)** Representative images of intact and sperm fractions stained for DNA (blue), centrin (green) and α-tubulin (red). Scale: 10 μm. N = 3 different experiments.

### The human sperm basal body is associated with a complex set of centrosomal proteins

To define the complexity of the proteins associated with the human sperm basal body, we used mass spectrometry. Centrosomal proteins are usually in very low abundance compared with other cellular components and they may be even less represented in sperm (Bauer *et al*., 2016). We therefore aimed at reducing the sample complexity before mass spectrometry by separating the sperm heads from the tails and basal bodies (Amaral *et al*., 2013, Baker *et al*., 2013). Human normozoospermic samples were sonicated and centrifuged twice through a 30% sucrose cushion in order to obtain a fraction enriched in heads and another one enriched in tails **(Figure 2B)**. The purity of the tail fraction evaluated by bright-field microscopy showed that less than 0.01% of the samples contained a few contaminating heads **(Supplementary Figure 1A)**. To further test the purity of the tail fraction, we checked for the presence of protamines as a marker for chromatin and γ-tubulin as a marker of centrosomes in the head and tail fractions **(Figure 2C)**. Western Blot (WB) analysis showed that intact sperm lysates were positive for protamines and γ-tubulin as expected. In contrast, protamines were not detected in the tail fraction whereas γ-tubulin was present. These data confirmed that the tail fraction contains very low levels of head contaminants if any. We next examined if centrioles were present in the sperm tail fraction by IF analysis **(Figure 2D)**. Indeed, we could detect a positive signal for centrin at one of the ends of some tail fragments (10%) indicating that basal bodies were recovered in the tail fraction.

In order to identify as many centrosomal proteins as possible, we used different approaches for processing the tail fractions before mass spectrometry analysis. First, we aimed at differentially extracting the centrosomal proteins using a range of detergents as previously described (Firat-Karalar *et al*., 2014). The resulting samples derived from three successive extractions steps from three independent experiments were independently analyzed by mass spectrometry, leading to the identification of 1545 proteins with at least 2 unique peptides. In addition, as a complementary approach, the sperm tails were directly solubilized in LB and the proteins resolved by SDS-PAGE. The gel was excised in 9 fragments that were analyzed independently by mass spectrometry. This approach resulted in the identification of 3210 proteins with at least 2 unique peptides. Altogether we identified 3406 proteins with at least 2 unique peptides for the human sperm tail proteome with a significant enrichment for basal body and tail proteins **(Supplementary Table 1)**.

Gene Ontology analysis based on biological processes and cellular localization showed that many of the 3406 proteins are involved in metabolic processes, response to stress and intracellular transport, in agreement with previous reports (Amaral *et al*., 2013, Baker *et al*., 2013, Baker *et al*., 2007, Jumeau *et al*., 2015, Martinez-Heredia *et al*., 2006, Wang *et al*., 2013). Considering cellular localization, the majority had GO terms corresponding to mitochondria and cytoskeleton. Others had GO terms corresponding to the Golgi apparatus, the ER and vesicles **(Figure 3A and B)** (Amaral *et al*., 2013).

**Figure 3:**
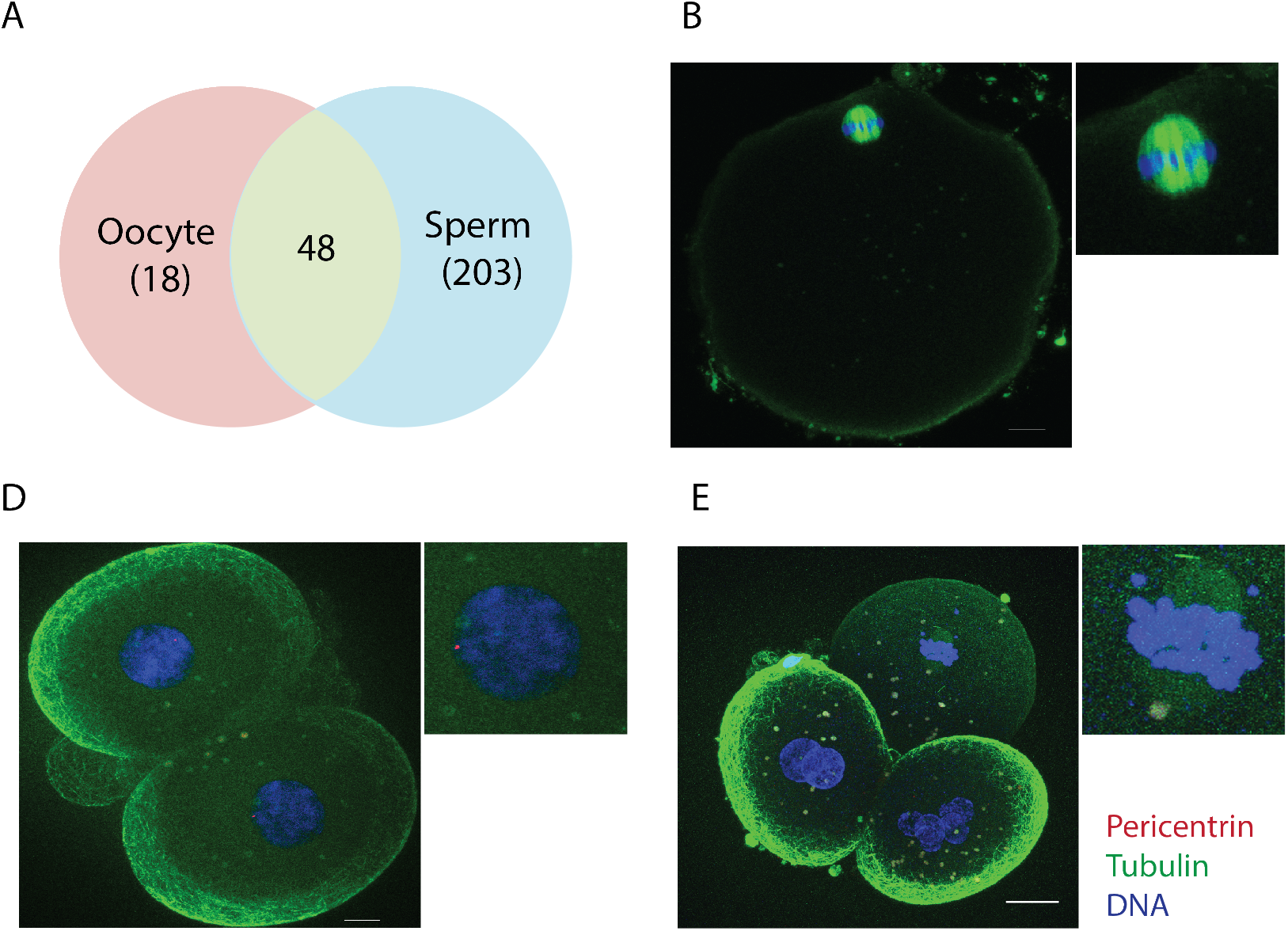
Classification of the human sperm tail proteins. **A)** Gene Ontology of the 3,406 proteins based on their biological process. **B)** Gene ontology of the same 3,406 proteins based on their subcellular localization. **C)** String network of the 251 identified centrosomal proteins. **D)** Comparison of the PCM proteins only identified in this work (19.1% - pink fraction) to all the previously published human sperm proteomic data (green fraction).

Interestingly, 170 proteins were associated to the GO terms: centrosome and/or centriole. Extending our analysis to the proteins identified with only one peptide (centrosomal proteins are in very low abundance) we found in addition 81 proteins with these GO terms **(Supplementary Table 2)**. To obtain some supporting evidence for these proteins being centrosomal, we checked them individually in the databases Uniprot and Protein Atlas. In Uniprot, 139 proteins out of the 251 total number of identified centrosomal proteins (97 identified with 2 unique peptides and 42 with 1 unique peptide) have a described centrosomal function and/or localization. In Protein Atlas, 116 were also classified as centrosomal (68 identified with 2 unique peptides and 48 with 1 unique peptide). We could not find data in these two databases for the remaining 69 proteins **(Supplementary Table 2)**. Then, a protein-protein interaction network using STRING database and the 251 proteins was established also showing a highly interconnected network **(Figure 3C)**. Taken altogether, our comprehensive analysis shows that the human sperm basal body is associated with a complex centrosomal proteome including at least 251 proteins.

Interestingly, the functions of the identified centrosomal proteins are very diverse including the regulation of centriole structure and length (centrin, POC1B), microtubule nucleation (tubulin γ-1, GCP2, GCP3 and GCP6), phosphorylation of multiple factors (Nek9 and Aurora C Kinases), centrosome cycle regulation and biogenesis (cep135, cep170) and PCM organization (ODF2, PCM1). Some of the centrosomal proteins we identified have not been reported previously as present in human sperm (12 of the 170 centrosomal proteins with 2 peptides and 36 out of the 81 centrosomal proteins with 1 peptide; in total 48 out of the 251 proteins (19,1%)) (Castillo *et al*., 2018) **(Figure 3D, Supplementary Table 2).**

The proteome of isolated centrosomes from KE37 cells contains a higher number of associated proteins (Bauer *et al*., 2016). To determine whether we only detected abundant proteins using our approach, we checked the reported abundance of the proteins we identified in the KE37 centrosome proteome database. Data were available for 20 of the proteins we identified, including some abundant ones like ODF2 and POC1B, others having intermediate abundance values such as OFD1 and Cep76, and yet others with low abundance, such as Cep170 and Cep290. These data suggest that we obtained a good representation of the centrosomal proteins associated with the human sperm basal body.

Since we successfully identified many centrosomal proteins in the human sperm samples we decided to explore whether any of the 26 proteins identified in at least two independent experiments and currently uncharacterized could be novel centrosomal components **(Supplementary Table 3)**. We obtained constructs for expression of 10 of them with a fluorescent tag **(Supplementary Table 3)** in HeLa cells. IF analysis showed that 3 of them co-localized with centrin, suggesting that they are novel human centrosomal proteins **(Supplementary Figure 2)**. Consistently, one of them C7orf31 was recently identified in bovine sperm and validated through localization studies as a novel centrosomal protein (Firat-Karalar *et al*., 2014). In summary, we have identified 251 centrosomal proteins and 3 novel centrosomal components in human sperm. Our results suggest that the human sperm basal body is associated with a complex pericentriolar protein network.

### The early embryonic centrosomal proteins are biparentally inherited

The current view is that centrosomal proteins are recruited from the oocyte cytoplasm by the sperm centrioles to assemble the first centrosome of the future organism. However, our data revealed that the sperm basal body is already associated with a complex network of centrosomal proteins. This suggested that upon fertilization the sperm not only provides half of the chromosomes to the zygote and centrioles but also a high number of centrosomal proteins suggesting that in fact, the first centrosome that assembles in the zygote has a mixed pericentriolar material derived from both the sperm basal body and the oocyte cytoplasm. To gain some insights into the centrosomal proteins stored in the human oocyte cytoplasm that may be recruited we first analyzed the 1376 proteins identified in the human oocyte (Virant-Klun *et al*., 2016). Only 66 of these proteins have the GO terms: centrosome and/or MTOC, including 48 that we identified in the human sperm **(Figure 4A, Supplementary Table 4)**. Thus, these data suggest that the first centrosome of the zygote is of biparental origin with some centrosomal proteins coming from the sperm basal body and others being recruited from the oocyte cytoplasm.

**Figure 4:**
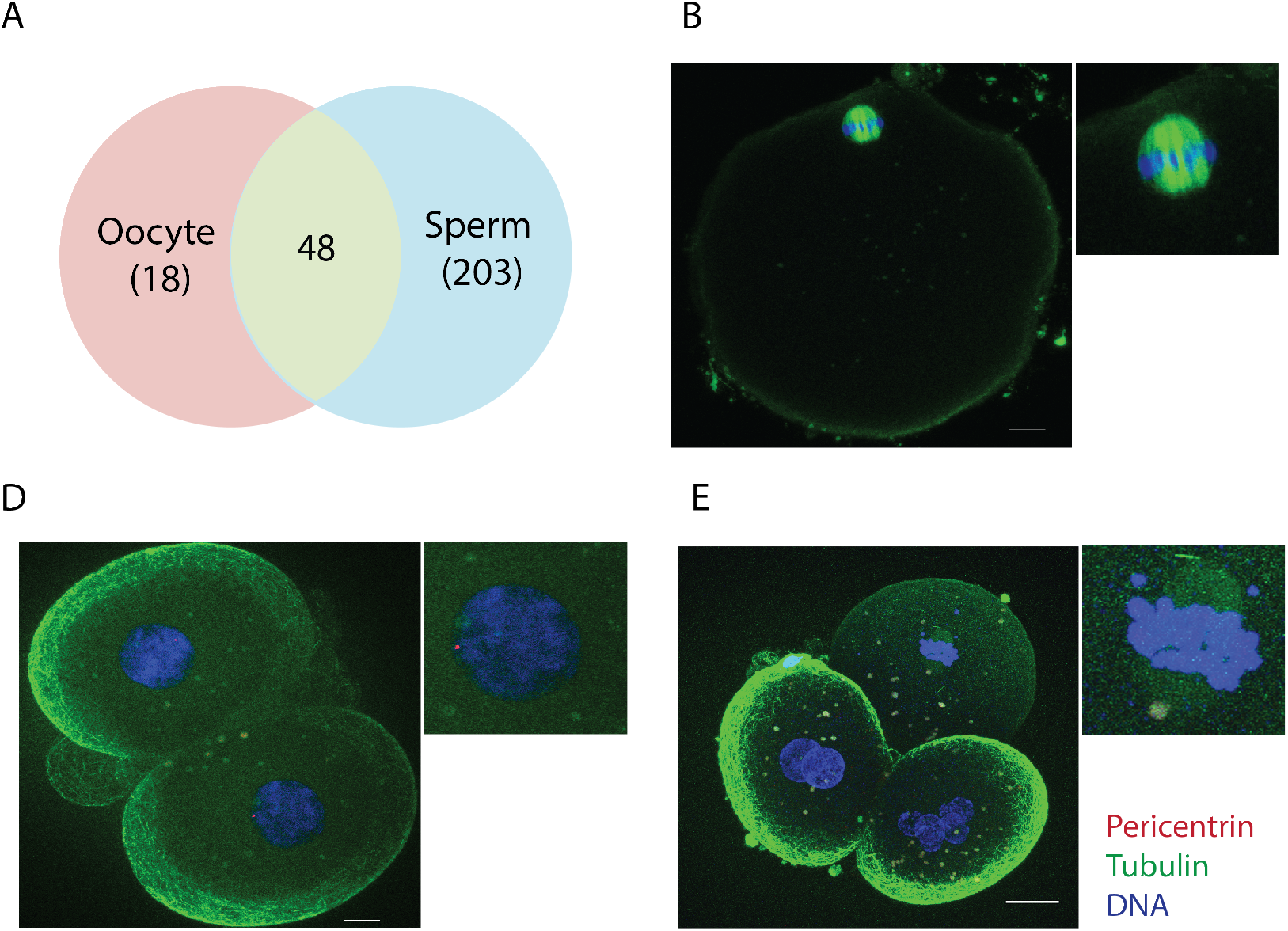
The zygote centrosomal composition is biparentally inherited during fertilization. **A)** Venn diagram showing overlap of 48 centrosomal proteins in the human oocyte and sperm. **B)** Representative IF image of an *in vitro* matured MII oocyte stained for pericentrin, tubulin and DNA. Scale: 10 μm. N = 10 oocytes. **D)** IF images of abnormally fertilized human embryos at D+2 with pericentrin, tubulin and DNA. Scale: 10 μm. N = 13 fertilized human embryos. **E)** Pericentrin, tubulin and DNA staining of a D+2 parthenogenetically activated human oocyte. Scale: 20 μm. N = 6 activated human oocytes.

To test this idea, we focused on pericentrin, a centrosomal component involved in the recruitment of other centrosomal proteins (Kim and Rhee, 2014). We identified pericentrin in our sperm proteome although with only one unique peptide. In the oocyte, pericentrin was not detected by either IF **(Figure 4B)** or proteomics (Holubcova *et al*., 2015). In *in vitro* fertilized human oocytes, 87% of embryos arrested with 1 pronucleus (PN) and 100% of embryos arrested with ≥3 PN contained pericentrin positive foci **(Figure 4C) (Supplementary Figure 3A and B)**. Interestingly, in parthenogenetically activated human oocytes, pericentrin signal was not detected **(Figure 4D)**, suggesting that pericentrin may be paternally inherited or the sperm-derived centrioles are needed to recruit pericentrin from the oocyte cytoplasm. Altogether, these data suggest that both the human sperm and the oocyte provide centrosomal components to the zygote centrosome. They further suggest that the contribution of the sperm goes beyond providing centrioles by also providing a complex set of proteins for the assembly of the first functional centrosome of the future organism.

### The sperm derived centrosome provides robustness to the early cell division cycles of the human parthenotes

To evaluate the role and contribution of the sperm derived centrosome in early embryonic development, we devised an experimental system to inject meiotically mature human oocytes with sperm basal bodies and follow their pseudo-development upon activation **(Figure 5A)**. First, we checked that 83.9% of ejaculated normozoospermic sperm showed a positive signal for centrin indicating that they do have centrioles **(Figure 5B)**. To isolate these centrioles, we microsurgically severed sperm tails at the transition between the flagella and the head **(Methods and Figure 5A)**. To confirm that the severed tails contained the centrioles we monitored the presence of centrin in these samples. 63.1% of the severed sperm tails showed either 1 or 2 centrin fluorescent dots, and no traces of DNA **(Figure 5C and D)**. Thus, the injection of these severed tails into oocytes, followed by parthenogenetic activation, should provide a model to study the role of the sperm derived centrosome in the initial phases of human development.

**Figure 5:**
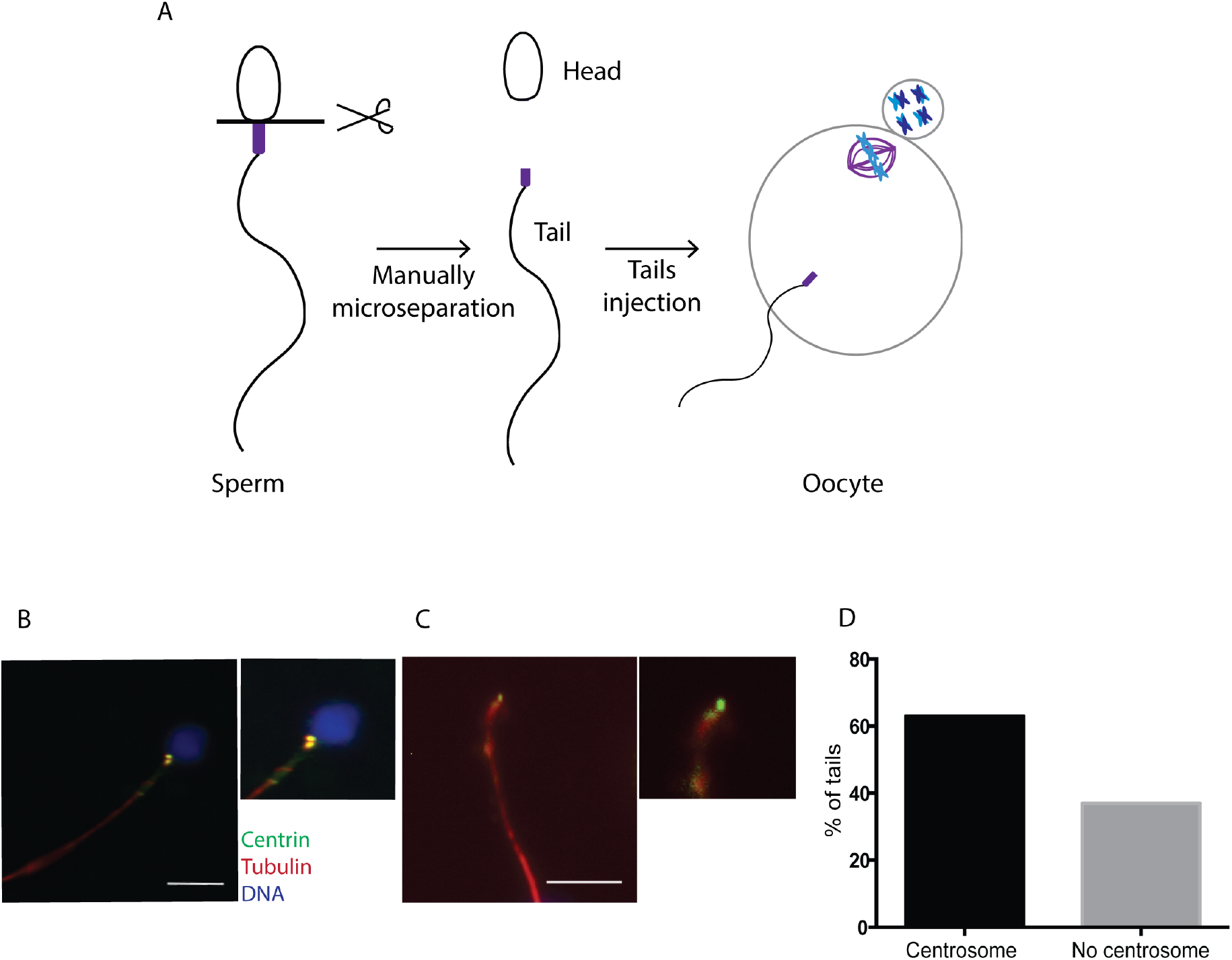
The sperm basal body localizes to the microsurgically separated tails. **A)** Schematic representation of our functional assay. **B)** IF for centrin, tubulin and DNA on intact sperm. Scale: 7.5 μm. **C)** Representative IF images of a manually separated sperm tail stained for centrin, tubulin and DNA. Scale: 7.5 μm. **D)** Graph showing the percentage of isolated tails with centrosomes. N = 2 different sperm samples.

Next, we injected severed sperm tails into human oocytes and parthenogenetically activated them **(Figure 5A)**. Parthenotes are a good experimental model to mimic the early stages of embryo development due to the similarity of most parameters used for evaluating *in vitro* development in activated or fertilized oocytes (Paffoni *et al*., 2007). We then monitored by time-lapse both control (sham-injected, n=10) and tail injected (n=15) parthenotes for up to 5 days of pseudo-development in two independent experiments. The percentage of control parthenotes that reached the pseudo-blastocyst stage by day 5 (20%) was in line with published reports (Paffoni *et al*., 2007), and it was slightly higher in tail injected parthenotes (27%) **(Figure 6A)**. However, tail injected parthenotes had a significantly higher survival rate at all the time points examined: 1-, 2-, 3-, 4, 5-cells, compaction, early pseudo-blastocyst and expanded pseudo-blastocyst stages **(Figure 6B and Supplementary table 5)**. This was particularly evident before compaction when the large-scale embryonic transcription begins in the human species **(Figure 6B and C)**. In fact, 4 of the 10 control parthenotes arrested before compaction, whereas only 4 of the 15 tail-injected parthenotes arrested before this stage **(Figure 6D).** These data suggest that tail injected parthenotes may have an increased probability to go through the earlier cell divisions successfully than control ones.

**Figure 6:**
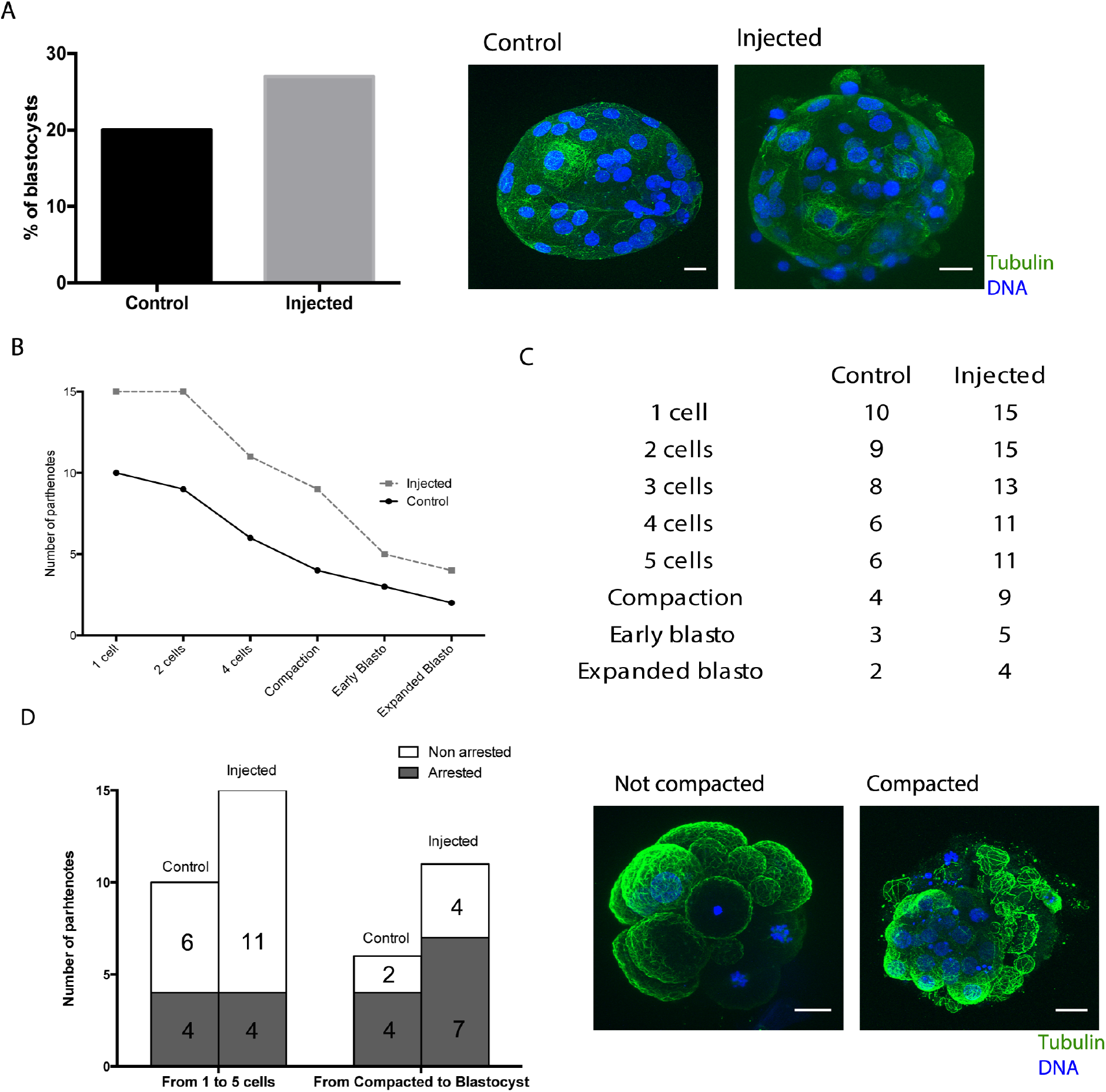
Sperm centrosome inheritance during fertilization ensures parthenotes compaction. **A)** The graph on the left shows the percentage of parthenotes that form a blastocyst-like structure at D+5 in controls and injected oocytes. The images on the right are representative pseudo-blastocysts obtained in control and injected oocytes. Scale: 20 μm. **B)** Developmental progress of control *vs* injected parthenotes. The graph represents the number of parthenotes that achieved each cellular or embryonic stage. **C)** Table with the number of control and injected parthenotes in each cellular and embryonic stage. **D)** The graph on the left show the rate of control and injected oocytes that arrested before or after compaction. On the right, representative images of non-compacted and compacted parthenotes. Scale: 20 μm. N = 10 control and N = 15 injected oocytes in 2 independent experiments.

To confirm that the tail-injected oocytes contained centrosomes, we fixed the parthenotes at day 5 and processed them for IF. We detected pericentrin in 8 out of the 15-tail injected parthenotes. Moreover, the proportion of cells with 1 or 2 centrosomes (as expected in normal cell cycles) highly resembles the ones we quantified in discarded embryos from 1 or 3 pronuclei: 66.2% in embryos and 62.8% in tail injected parthenotes **(Figure 7A and B)**. These data suggested that tail injected parthenotes do contain centrosomes that cycle correctly. In summary, our experimental approach suggests for the first time that the human sperm derived centrosome play an important role in ensuring the robustness of the early human parthenote cell divisions up to compaction when additional requirements provided by a large scale embryonic transcription come into play.

**Figure 7:**
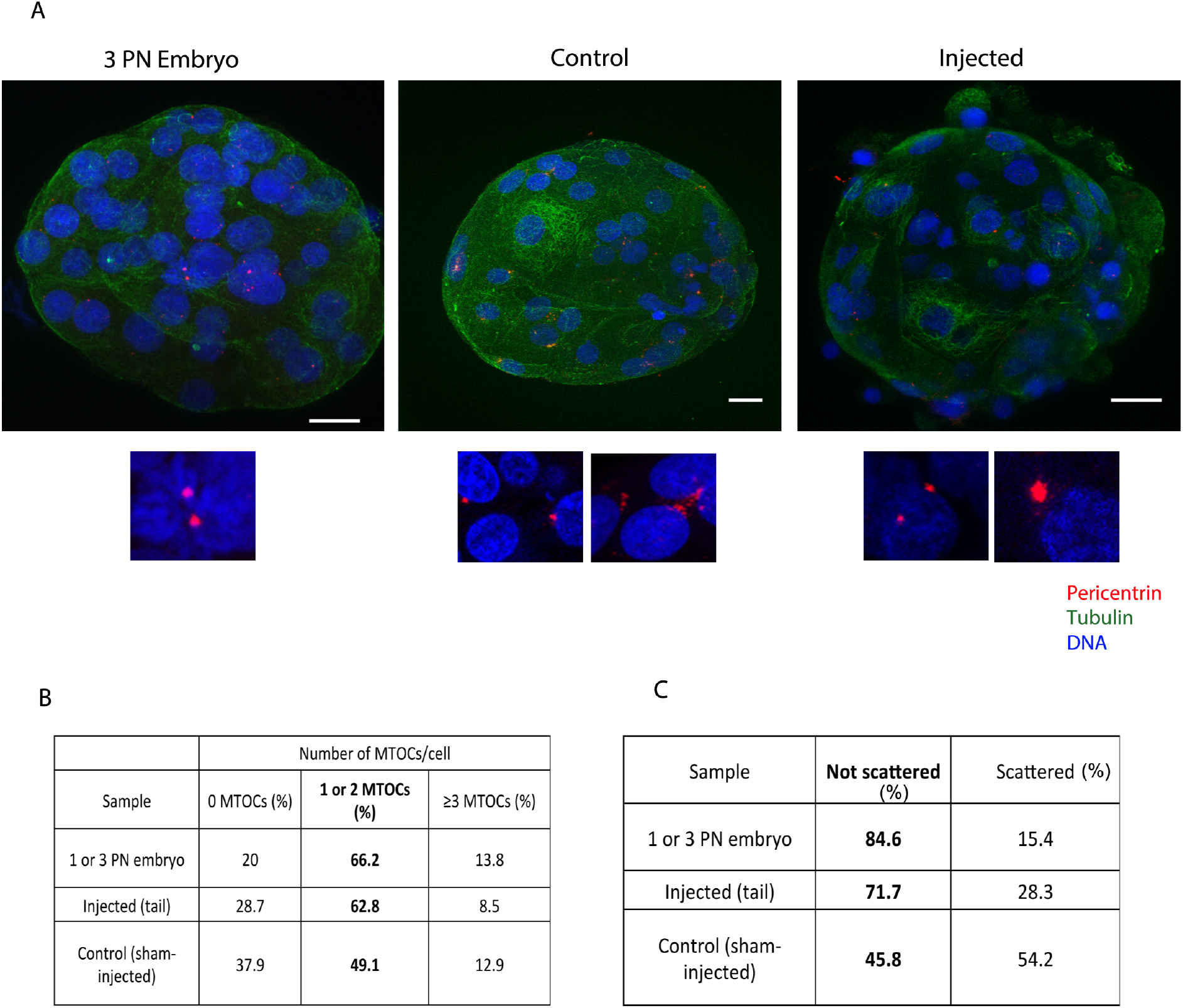
MTOCs can be formed *de novo* in control pseudo-blastocysts and after the activation of the embryonic genome. **A)** Representative IF images of blastocyst and pseudo-blastocyst of abnormal fertilized oocytes, control and injected oocytes stained for pericentrin, tubulin and DNA. The lower panels are magnifications of the MTOCs for each condition. Scale: 20 μm. **B)** Number of MTOCs *per* cell in abnormal fertilized oocytes, control and injected oocytes. **C)** Table showing the percentage of scattered MTOCs per sample. N = 5 3PN embryos, N = 10 control and N = 15 injected oocytes.

### Human parthenotes can form *de novo* MTOCs

While performing the IF studies to monitor pericentrin in parthenotes, we surprisingly found that those derived from control oocytes also contained pericentrin positive aggregates at the early pseudo-blastocyst stage after compaction **(Supplementary table 4 and Figure 7A)**. These aggregates could potentially be *de novo* MTOCs similar to those forming in mice embryos at blastocyst stage. When we quantified their number of MTOC structures, we found that the proportion of cells having pericentrin positive MTOC-like aggregates was lower (49.1%) than in the tail injected parthenotes. Moreover, the morphology of these aggregates was different than those detected in tail injected parthenotes; they had irregular shapes and some degree of scattering that was not usually observed in the tail injected parthenotes **(Figure 7A – lower panels and C)**. More than half of the pericentrin aggregates (54.2%) observed in the sham injected parthenotes showed this ‘scattered’ morphology whereas this was the case for only very few pericentrin positive MTOCs in abnormal fertilized embryos (15.4%) and injected oocytes (28.3%) **(Figure 7C)**.

We hypothesized that the embryonic genome activation could be involved in the formation of *de novo* pericentrin MTOC-like aggregates. In humans, at Day 2 after fertilization, a first burst of embryonic genome activation occurs, but it is not until Day 3 and 4 that the major activation wave happens (Vassena *et al*., 2011). We therefore monitored pericentrin by IF in another set of sham-injected and parthenogenetically activated oocytes fixed at Day 3 (before the major wave of embryonic genome activation), and at Day 5 (after embryonic genome activation). None of the oocytes fixed at Day 3 had pericentrin signal (0 out of 6). The single parthenote out of 5 parthenotes that progressed until the pseudo-blastocyst stage contained pericentrin aggregates **(Supplementary table 6)**.

Since the centrosome was previously proposed to define the kinetics of the first embryonic spindle bipolarization (Cavazza *et al*., 2016), we checked whether the presence of the sperm derived centrosome had any influence on the early embryonic development kinetics considering only the tail injected parthenotes that did have MTOCs detected by IF. Although these parthenotes reached each of the cellular and embryonic stages faster than non-injected controls, the differences were not significant **(Supplementary Figure 4)**.

Altogether, our results indicate that the paternal inheritance of the basal body is important to establish the centrosome structure and number per cell in the developing pseudoembryo.

## Discussion

The mechanism of centrosome inheritance during fertilization is an essential process to ensure the proper number of centrosomes per cell and their function in the new organism, yet this process has not been studied in humans. Our study supports the hypothesis that the human sperm basal body is a remodeled centrosome with an atypical structure that is associated with an extensive variety of centrosomal proteins. Moreover, we addressed for the first time the role of the human sperm derived centrosome in the initial phases of human pseudoembryo development. Our data suggest that the human sperm derived centrosome plays an important role during the early developmental events leading to embryo compaction.

It is currently accepted that the human sperm basal body consists of two highly remodeled centrioles and little associated PCM. After fertilization, the sperm would therefore only provide one functional centriole to the oocyte and little, if any, PCM (Avidor-Reiss *et al*., 2015, Fawcett and Phillips, 1969, Manandhar and Schatten, 2000). However, 2 functional centrosomes need to be assembled in the zygote to provide the two first cells with one centrosome each after the first division of the zygote (Palermo *et al*., 1997). Our study provides novel data that support an alternative mechanism. High-resolution microscopy has recently shown that the sperm distal centriole consists of an atypical, splayed microtubule structure that retains some centrosomal proteins (Fishman *et al*., 2018). Our super resolution analysis of centrin and acetylated tubulin localization in the human sperm basal body supports these findings. It also suggests that despite the atypical structure of the distal centriole, it is most likely a stable structure that retains some intrinsic structural characteristics shared with the proximal one. In any case, further studies are needed to comprehensively analyze the stability of both proximal and distal centrioles in the human sperm.

The identification of centrosomal proteins is usually difficult because of their low abundance in cells (Bauer *et al*., 2016). This issue was particularly challenging for this work because of the process of centrosome reduction in sperm (Avidor-Reiss *et al*., 2015, Manandhar *et al*., 2000). We could however enrich the samples by eliminating the sperm heads and identify more than 3,406 sperm tail proteins. To our knowledge, this is the most complex human sperm tail proteome. Most of the identified proteins are related to the cytoskeleton or mitochondria, in agreement with previous results (Amaral *et al*., 2013, Baker *et al*., 2013, Baker *et al*., 2007, Jumeau *et al*., 2015, Martinez-Heredia *et al*., 2006, Wang *et al*., 2013). The proteome also includes 251 centrosomal proteins with 183 having validated localizations and/or functions at the centrosome. Interestingly, these proteins have a variety of functions at various levels including structural roles, the mechanism of centriole duplication and microtubule nucleation. This is somehow surprising since these functions are in principle not required in the mature sperm. In the human oocyte, we found that some centrosomal components are stored as proteins (Virant-Klun *et al*., 2016) suggesting that the centrosomal proteins may be biparentally inherited. The sperm derived centrosomal proteins may play an important role after fertilization to promote a quick transition from the sperm basal body to a functional centrosome in the zygote cytoplasm. Interestingly, a similar mechanism was recently proposed in *Drosophila*. In flies, the sperm centrosome retains some centrosomal components that are essential to support normal embryogenesis (Khire *et al*., 2016).

The localization of a subset of human sperm centrosomal proteins has been recently described using IF (Fishman *et al*., 2018) and our proteomic analysis further identified 6 out of 18 proteins that Fishman and colleagues already described. Interestingly, we identified 3 centrosomal proteins that could not be detected by IF analysis (γ-tubulin chain 1, pericentrin and pericentriolar material 1), either because of technical limitations such as accessibility in fixed samples or their very low abundance, opening the possibility that many other centrosomal proteins may also be present in the basal body of the mature sperm.

The injection of microsurgically severed human sperm tails containing the basal body in parthenogenetically activated oocytes offered a unique system to test directly the role of the sperm basal body during preimplantation human development independently of other sperm components such as the paternal genome. This is in fact the first time that the assembly and role of the first functional centrosome in zygotes obtained using human gametes could be addressed following human parthenotes pseudo-development. Due to the use to a limited access of viable human oocytes and the technical complexity of the methodology, we had to work with a small sample size. This restricted the use of robust statistical analysis of the results and hampered the possibility of addressing mechanistically the role of the centrosome in human early development. Despite these limitations, our data suggest that the sperm derived centrosome helps parthenotes to transit successfully through the early cell divisions up to compaction. Compaction is characterized by cell internalization and embryonic reorganization (Maitre *et al*., 2016) that in mice is partially mediated by the orientation of the cell division orchestrated by the aMTOC (Korotkevich *et al*., 2017).

Surprisingly, we could detect the formation of *de novo* MTOCs in control pseudo-blastocysts. In humans, embryonic genome activation starts as early as at the 4-cell stage (D+2 of embryo development) but, the peak of gene expression occurs when the embryo is at the morula stage (compaction) (Vassena *et al*., 2011). Our results suggest that when the genome is activated, the production of centrosomal proteins increases considerably after D+3. Centrosomal proteins may cluster and form independent identities probably through a phase separation mechanism (Woodruff *et al*., 2017). Their ability to accumulate tubulin triggers microtubule nucleation. The appearance of MTOCs at the pseudo-blastocyst stage is kinetically similar to *de novo* centrosome formation in mice (Courtois *et al*., 2012, Howe and FitzHarris, 2013). Although we cannot assume that in the parthenotes they contain centrioles, it suggests that MTOCs activity is important for the development of complex organisms.

Together our results provide novel insights about the mechanism of centrosome inheritance upon fertilization in humans and its importance in supporting early embryogenesis. These results can also have an important impact in assisted reproduction technologies in which 30% of the fertilized oocytes arrest before compaction. We show here that centrosome dysfunction may not only be associated with sperm motility and morphology alterations (Amargant *et al*., 2018, Jumeau *et al*., 2017) but it could derive from an defective conversion into a fully functional centrosome in the zygote that could explain some of the unexpected early embryo arrests. Further studies using our novel tail-injection method are guaranteed to provide new strategies to address unexpected cases of human infertility.

## Data Availability statement

All data are incorporated into the article and its online supplemental material

## Acknowledgements

We want to thank all members of the Basic Laboratory from Clinica EUGIN as well as Vernos Lab for fruitful discussions. We also thank all the clinical embryology laboratory staff from Clínica EUGIN and CIRH for their assistance with sample collection and the proteomic facility at the Center for Genomic Regulation for their assistance with the proteomic analysis.

## Author’s roles

F.A. performed the experiments, analyzed data and wrote the paper. A.P. performed sperm tails separation and injection experiments. A.F.V. performed oocyte warming and provided technical support with the microscope imaging. M.D. performed AOA experiments. M.M. analyzed parthenotes development. R.V and I.V. supervised the work, designed experiments, interpreted results and critically reviewed and edited the manuscript.

