## Supplementary information for "The human sperm basal body is a complex centrosome important for embryo pre-implantation development"

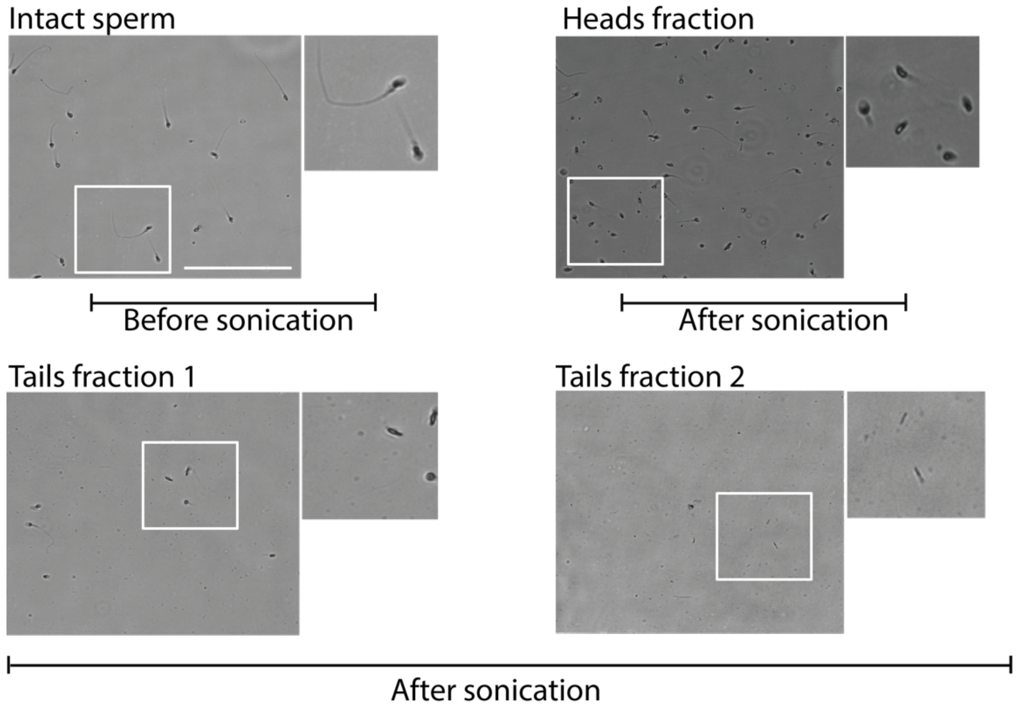


**Supplementary Figure 1: Phase-contrast images of sperm fractions.** Representative phase-contrast images of the intact human sperm, heads, and tail fraction 1 and 2 obtained before and after sonication and sucrose separation. Scale: 100 µm. N = 3 independent experiments.

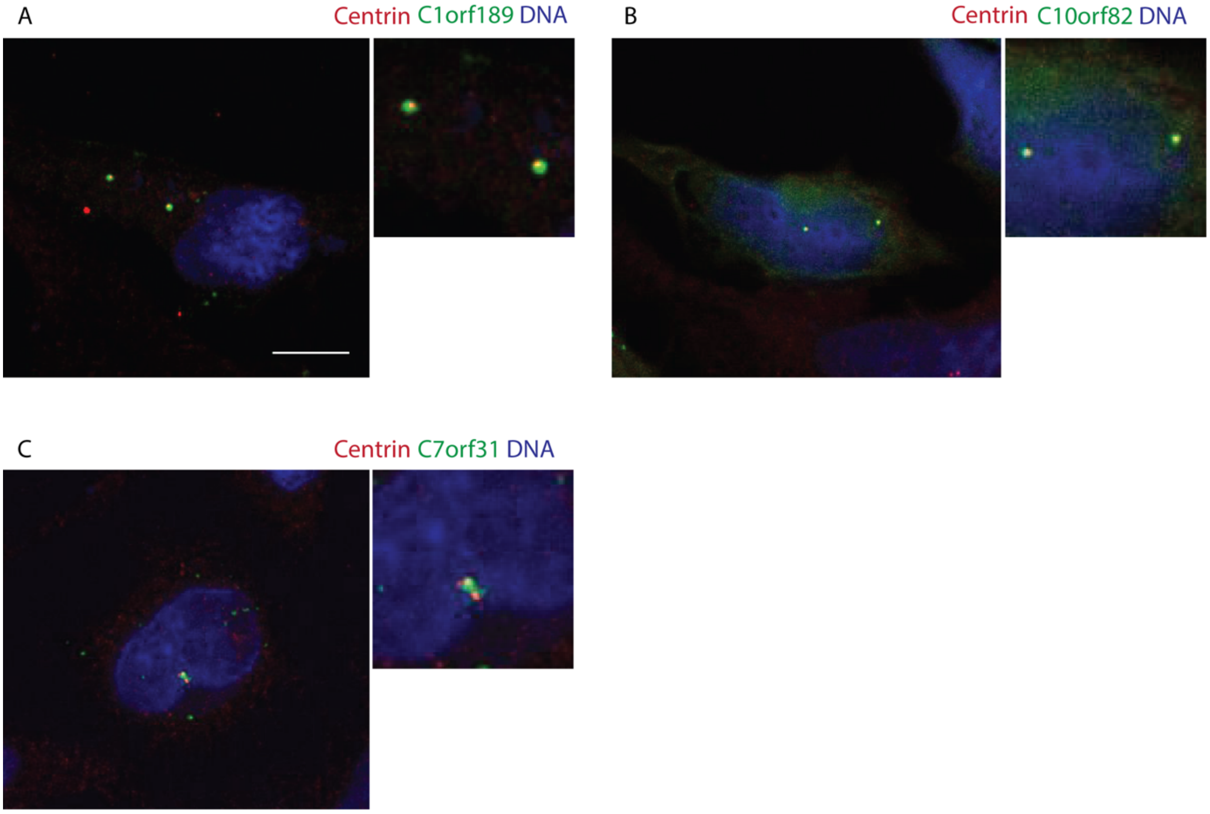


**Supplementary Figure 2: Localization of the newly identified centrosomal proteins.** IF of HeLa cells transfected for 24h and fixed with methanol for centrin, DNA and uncharacterized proteins C1orf189 (**A**), C10orf82 (**B)** and C7orf31 (**C**). Scale: 10 µm. N = 3 independent experiments.

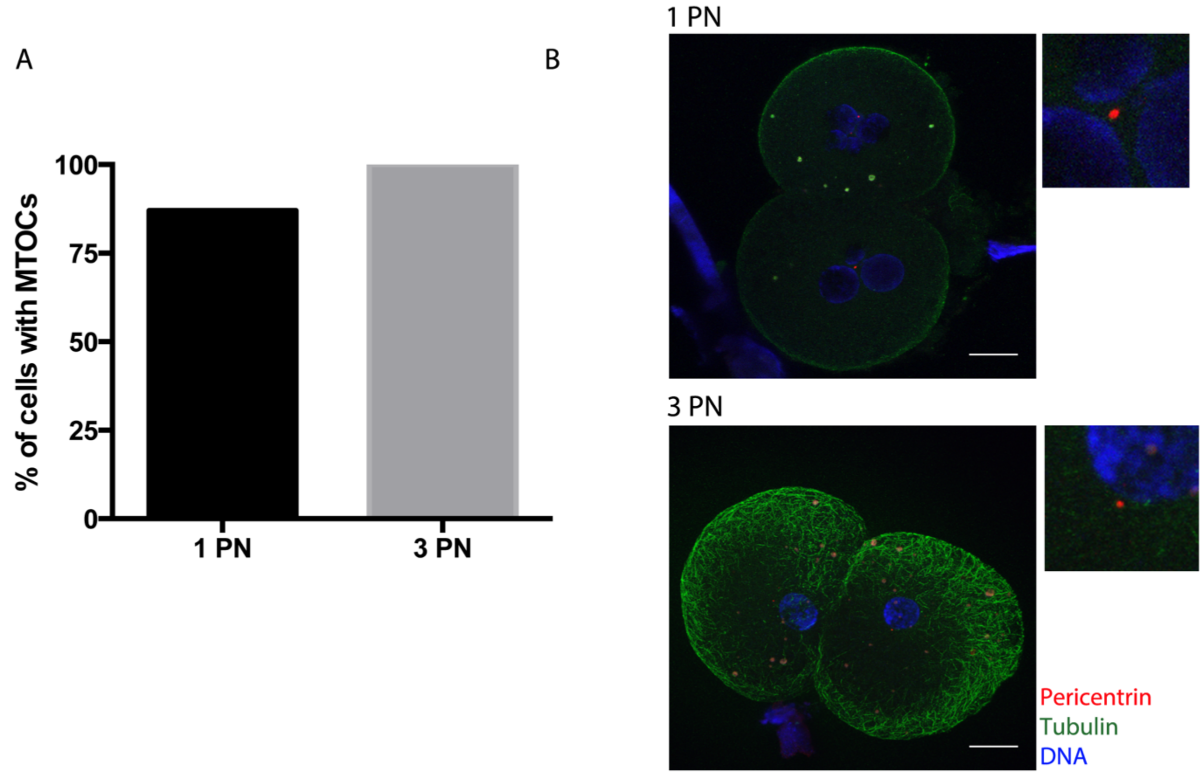


**Supplementary Figure 3: Pericentrin staining detects the centrosome in human embryos. A)** Graph showing the percentage of cells that contain at least one MTOC at 2- to 4-cell stage of 1 and ≥3-PN embryos. **B)** Representative pictures of 1 and ≥3-PN embryos stained for pericentrin, tubulin and DNA. Scale: 20 µm. N = 13 fertilized embryos.

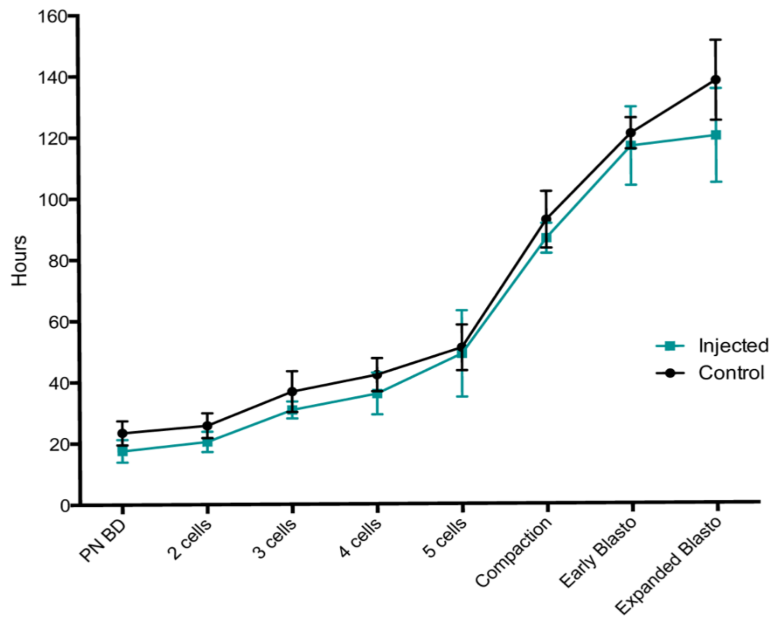


**Supplementary Figure 4: The early parthenotes development kinetics does not differ in control and injected samples.** Graph showing the time that control and injected oocytes need to reach each of the cellular and embryonic stages.

**Supplementary Table 1: Proteins identified in the sperm tail proteome.** Exp01/2/3_01: Experiment 1/2/3, fraction 1. Exp01/2/3_02: Experiment 1/2/3, fraction 2. Exp01/2/3_03: Experiment 1/2/3, fraction 3. Exp04: Experiment 4 – gel approach. In bold are proteins classified as "contaminants". These were discarded from our analyses.

**Supplementary Table 2: Sperm centrosomal proteins identified by mass spectrometry.** The first sheet contains the centrosomal proteins with more than 2 unique peptides and the second sheet the ones identified with only 1 unique peptide. In green are the new centrosomal proteins identified in this work, in yellow the proteins that have been validated as centrosomal using the Uniprot database and in blue in the Protein Atlas. In pink are the proteins identified as centrosomal in both databases. The third sheet includes the centrosomal proteins whose role in the centriole (red) and/or centrosome (green) have been validated using the Uniprot database and in the following sheet the ones validated using the Protein atlas database. The last sheet contains the proteins classified with the GO terms centrosome or centriole but their centriolar or centrosomal role could not be validated using the Uniprot and Protein atlas databases.

**Supplementary Table 3: Uncharacterized proteins identified in the sperm tail proteome.** In bold are the proteins we analyzed their localization.

**Supplementary Table 4: Sperm centrosomal proteins also identified in the single-cell oocyte proteome**

**Supplementary Table 5: Time-lapse analysis of control and injected parthenotes.** The time needed to achieve each cellular and embryonic stage per parthenotes was annotated in hours. “No data” refers to stages that could not be analyzed due to the low quality of the image. The last two columns indicate whether the sample could be analyzed by IF and the presence of MTOC signal.

**Supplementary Table 6: Time-lapse analysis of control parthenotes fixed before or after the embryonic genome activation.** The time needed to achieve each cellular and embryonic stage per sample was annotated in hours. “No data” refers to stages that could not be analyzed due to the low quality of the image. The last two columns indicate whether the sample could be analyzed by IF and the presence of MTOC signal.

#### SUPPLEMENTARY TABLE 1

| Accession | ΣCoverage | Σ# Proteins | Σ# Unique Peptides | Σ# Peptides | Σ# PSMs | Exp01_01 Coverage A4 | # Peptides A4 | # PSM A4 | Exp02_01 Coverage B4 | # Peptides B4 | # PSM B4 | Exp03_01 Coverage C4 | # Peptides C4 | # PSM C4 | Exp01_02 Coverage D4 | # Peptides D4 | # PSM D4 | Exp02_02 Coverage E4 | # Peptides E4 | # PSM E4 | Exp03_02 Coverage F4 | # Peptides F4 | # PSM F4 | Exp01_03 Coverage G4 | # Peptides G4 | # PSM G4 | Exp02_03 Coverage H4 | # Peptides H4 | # PSM H4 | Exp03_03 Coverage I4 | # Peptides I4 | # PSM I4 | Exp04 Coverage J4 | # Peptides J4 | # PSM J4 | # AAs | MW [kDa] | calc. pI |
|---|---|---|---|---|---|---|---|---|---|---|---|---|---|---|---|---|---|---|---|---|---|---|---|---|---|---|---|---|---|---|---|---|---|---|---|---|---|---|
| A0A1B0GTD5 | 29.01 | 1 | 3 | 3 | 10 | | | | | | | | | | | | | | | | | | | | | | | | | | | | 29.01 | 3 | 10 | 131 | 15.6 | 9.54 |
| A0A1B0GTY4 | 13.56 | 1 | 2 | 2 | 2 | | | | | | | | | | | | | | | | | | | | | | | | | | | | 13.56 | 2 | 2 | 177 | 20.8 | 9.25 |
| A0A1B0GTZ2 | 8.75 | 1 | 2 | 2 | 3 | | | | | | | | | | | | | | | | | | | | | | | | | | | | 8.75 | 2 | 3 | 297 | 34.5 | 8.81 |
| A0AV96 | 4.22 | 1 | 2 | 2 | 3 | | | | | | | | | | | | | | | | | | | | | | | | | 2.19 | 1 | 1 | 2.02 | 1 | 2 | 593 | 64.1 | 7.68 |
| A0AVF1 | 9.75 | 1 | 4 | 4 | 5 | | | | | | | | | | | | | | | | | | | | | | | | | | | | 9.75 | 4 | 5 | 554 | 64.1 | 6.93 |
| A0AVT1 | 44.49 | 1 | 39 | 39 | 144 | 6.08 | 6 | 6 | 8.75 | 8 | 8 | 6.18 | 6 | 6 | 3.52 | 4 | 4 | 1.90 | 2 | 2 | | | | 1.14 | 1 | 1 | 2.09 | 2 | 2 | 8.94 | 8 | 8 | 39.83 | 33 | 107 | 1052 | 117.9 | 6.14 |
| A0MZ66 | 38.51 | 1 | 26 | 26 | 77 | 2.22 | 1 | 1 | | | | 1.27 | 1 | 1 | 3.01 | 2 | 2 | | | | | | | 20.92 | 12 | 15 | 19.33 | 10 | 10 | 29.00 | 17 | 23 | 20.60 | 12 | 25 | 631 | 71.6 | 5.33 |
| A1A4V9 | 32.02 | 1 | 7 | 7 | 11 | | | | 3.63 | 1 | 1 | | | | | | | | | | | | | | | | | | | | | | 28.40 | 6 | 10 | 331 | 37.9 | 6.74 |
| A1A519 | 58.18 | 1 | 22 | 22 | 122 | 3.94 | 1 | 2 | 3.94 | 1 | 1 | 3.94 | 1 | 2 | 14.55 | 4 | 4 | 10.61 | 3 | 4 | | | | 37.58 | 10 | 13 | 31.52 | 10 | 16 | 22.12 | 7 | 8 | 56.67 | 19 | 72 | 330 | 37.1 | 5.03 |
| A1L0T0 | 20.41 | 1 | 10 | 10 | 14 | | | | 2.85 | 1 | 1 | 3.64 | 1 | 1 | | | | | | | | | | | | | | | | | | | 19.78 | 8 | 12 | 632 | 67.8 | 8.15 |
| A1XBS5 | 14.53 | 1 | 4 | 4 | 4 | | | | 2.42 | 1 | 1 | | | | | | | | | | | | | | | | | | | | | | 12.11 | 3 | 3 | 289 | 33.4 | 6.21 |
| A2RRP1 | 5.57 | 1 | 11 | 11 | 14 | | | | | | | | | | | | | | | | | | | | | | | | | | | | 5.57 | 11 | 14 | 2371 | 268.4 | 5.96 |
| A2RTX5 | 4.99 | 1 | 2 | 3 | 5 | 0.00 | | | | | | | | | | | | 0.00 | | | | | | | | | | | | | | | 4.99 | 3 | 5 | 802 | 92.6 | 6.05 |
| A2RUT3 | 26.42 | 1 | 4 | 4 | 7 | | | | | | | | | | | | | | | | | | | | | | | | | | | | 26.42 | 4 | 7 | 159 | 17.6 | 10.01 |
| A2RUU4 | 29.75 | 1 | 3 | 3 | 4 | | | | | | | | | | | | | | | | | | | | | | | | | | | | 29.75 | 3 | 4 | 121 | 14.0 | 8.54 |
| A4D126 | 4.66 | 1 | 2 | 2 | 2 | | | | | | | | | | | | | | | | | | | | | | | | | | | | 4.66 | 2 | 2 | 451 | 49.8 | 6.25 |
| A4D1F6 | 11.63 | 1 | 8 | 8 | 9 | | | | | | | | | | | | | | | | | | | | | | | | | | | | 11.63 | 8 | 9 | 860 | 98.0 | 7.46 |
| A4D1T9 | 24.68 | 1 | 3 | 3 | 9 | | | | 4.68 | 1 | 1 | | | | | | | | | | | | | | | | | | | | | | 20.00 | 2 | 8 | 235 | 26.4 | 8.87 |
| Q13885 | 77.53 | 2 | 1 | 37 | 971 | 45.39 | 18 | 44 | 52.58 | 20 | 59 | 46.07 | 19 | 41 | 53.71 | 25 | 70 | 53.48 | 27 | 93 | 6.97 | 2 | 3 | 27.42 | 11 | 20 | 29.21 | 13 | 27 | 29.89 | 14 | 28 | 70.79 | 30 | 586 | 445 | 49.9 | 4.89 |
| A4D263 | 56.16 | 1 | 17 | 17 | 40 | | | | | | | | | | 8.45 | 4 | 4 | 31.28 | 10 | 12 | | | | | | | | | | | | | 36.07 | 11 | 24 | 438 | 49.6 | 8.48 |
| A4FU69 | 27.35 | 1 | 33 | 33 | 77 | 1.33 | 1 | 1 | 2.00 | 2 | 2 | | | | 1.33 | 2 | 2 | 2.46 | 3 | 3 | | | | | | | | | | | | | 26.01 | 31 | 69 | 1503 | 173.3 | 5.82 |
| A4QMS7 | 77.55 | 1 | 11 | 11 | 29 | | | | | | | | | | 41.50 | 4 | 4 | 77.55 | 10 | 14 | | | | | | | | | | | | | 48.98 | 6 | 11 | 147 | 17.0 | 7.59 |
| A5D8V7 | 46.55 | 2 | 25 | 26 | 49 | | | | 3.03 | 2 | 2 | | | | 3.03 | 2 | 2 | 5.55 | 3 | 3 | | | | | | | | | | 2.02 | 1 | 1 | 46.55 | 26 | 41 | 595 | 69.1 | 9.07 |
| A5D8W1 | 37.62 | 1 | 28 | 28 | 50 | | | | 3.83 | 3 | 3 | | | | | | | | | | | | | | | | | | | | | | 36.45 | 27 | 47 | 941 | 105.8 | 7.17 |
| A5PLN9 | 3.84 | 1 | 2 | 2 | 6 | | | | | | | | | | | | | | | | | | | | | | | | | | | | 3.84 | 2 | 6 | 417 | 46.5 | 5.60 |
| A5YKK6 | 2.82 | 1 | 6 | 6 | 8 | | | | | | | | | | | | | | | | | | | | | | | | | | | | 2.82 | 6 | 8 | 2376 | 266.8 | 7.11 |
| P63261 | 78.93 | 2 | 1 | 35 | 758 | 69.07 | 20 | 48 | 65.60 | 20 | 45 | 73.60 | 23 | 53 | 70.93 | 23 | 53 | 72.27 | 24 | 46 | 8.27 | 3 | 3 | 54.13 | 19 | 36 | 45.87 | 16 | 29 | 63.47 | 26 | 56 | 73.33 | 25 | 389 | 375 | 41.8 | 5.48 |
| A6H8Z2 | 15.92 | 1 | 5 | 5 | 12 | 4.48 | 1 | 1 | | | | | | | | | | 4.73 | 2 | 2 | | | | | | | | | | | | | 15.92 | 5 | 9 | 402 | 45.4 | 5.60 |
| P60709 | 78.93 | 2 | 1 | 35 | 751 | 69.07 | 20 | 48 | 61.07 | 19 | 44 | 73.60 | 23 | 53 | 70.93 | 23 | 53 | 72.27 | 24 | 46 | 8.27 | 3 | 3 | 54.13 | 19 | 36 | 45.87 | 16 | 29 | 63.47 | 26 | 56 | 73.33 | 25 | 383 | 375 | 41.7 | 5.48 |
| A6NC86 | 15.20 | 1 | 2 | 2 | 3 | | | | | | | | | | | | | | | | | | | | | | | | | | | | 15.20 | 2 | 3 | 204 | 21.9 | 7.99 |
| A6NC98 | 1.96 | 3 | 2 | 3 | 3 | | | | | | | | | | | | | | | | | | | | | | | | | | | | 1.96 | 3 | 3 | 1476 | 164.7 | 5.11 |
| A6NCJ1 | 84.21 | 1 | 14 | 14 | 42 | | | | | | | | | | 20.10 | 3 | 4 | 25.36 | 6 | 7 | | | | 12.44 | 2 | 3 | 9.09 | 1 | 1 | 34.45 | 7 | 8 | 76.08 | 10 | 19 | 209 | 24.2 | 8.95 |
| A6NCL2 | 19.50 | 1 | 3 | 3 | 4 | | | | | | | | | | | | | | | | | | | | | | | | | | | | 19.50 | 3 | 4 | 159 | 17.8 | 8.13 |
| A6NCN8 | 10.16 | 1 | 2 | 2 | 3 | | | | | | | | | | | | | | | | | | | | | | | | | | | | 10.16 | 2 | 3 | 305 | 35.3 | 10.15 |
| A6NDG6 | 50.16 | 1 | 13 | 13 | 34 | 9.03 | 2 | 2 | 3.43 | 1 | 1 | 3.43 | 1 | 1 | 7.79 | 2 | 2 | | | | | | | | | | | | | 9.03 | 3 | 3 | 49.22 | 12 | 25 | 321 | 34.0 | 6.14 |
| A6NDK9 | 23.81 | 2 | 2 | 15 | 21 | | | | 1.73 | 1 | 1 | | | | | | | 10.39 | 5 | 5 | | | | | | | | | | | | | 17.60 | 10 | 15 | 693 | 79.8 | 5.39 |
| A6NE01 | 0.72 | 1 | 2 | 2 | 2 | | | | 0.00 | | | 0.00 | | | | | | | | | | | | | | | | | | | | | 0.72 | 2 | 2 | 2351 | 262.6 | 8.60 |
| A6NEK1 | 54.97 | 1 | 15 | 15 | 45 | 15.20 | 4 | 4 | 15.20 | 5 | 5 | 8.19 | 2 | 2 | | | | | | | | | | | | | | | | | | | 54.97 | 15 | 34 | 342 | 38.4 | 6.68 |
| A6NEN9 | 25.14 | 1 | 5 | 5 | 8 | | | | | | | | | | | | | | | | | | | | | | | | | | | | 25.14 | 5 | 8 | 183 | 21.3 | 10.33 |
| A6NFE3 | 75.59 | 1 | 7 | 7 | 13 | | | | | | | | | | 21.26 | 2 | 3 | | | | | | | | | | | | | | | | 75.59 | 7 | 10 | 127 | 14.7 | 5.38 |
| A6NFZ4 | 26.67 | 1 | 3 | 3 | 4 | | | | | | | | | | | | | | | | | | | | | | | | | | | | 26.67 | 3 | 4 | 105 | 11.3 | 7.84 |
| A6NGB0 | 29.97 | 2 | 8 | 8 | 21 | 3.46 | 1 | 1 | | | | | | | | | | | | | | | | 3.46 | 1 | 1 | | | | | | | 29.97 | 8 | 19 | 347 | 39.3 | 8.47 |
| A6NI87 | 14.46 | 1 | 3 | 3 | 3 | | | | | | | | | | | | | | | | | | | | | | | | | | | | 14.46 | 3 | 3 | 242 | 27.3 | 10.64 |
| A6NIV6 | 19.82 | 2 | 7 | 8 | 12 | | | | 1.96 | 1 | 1 | | | | | | | | | | | | | | | | | | | | | | 19.82 | 8 | 11 | 560 | 63.9 | 8.19 |
| A6NJI9 | 11.50 | 1 | 3 | 3 | 3 | | | | | | | | | | | | | | | | | | | | | | | | | | | | 11.50 | 3 | 3 | 287 | 33.6 | 8.76 |
| A6NJV1 | 53.23 | 1 | 12 | 12 | 33 | | | | | | | | | | 19.40 | 4 | 5 | 12.94 | 2 | 3 | | | | | | | | | | 12.44 | 2 | 2 | 44.28 | 8 | 23 | 201 | 23.4 | 9.99 |
| A6NKS8 | 9.96 | 1 | 2 | 2 | 2 | | | | | | | | | | | | | | | | | | | | | | | | | | | | 9.96 | 2 | 2 | 231 | 25.2 | 8.27 |
| A6NM11 | 22.76 | 2 | 7 | 31 | 149 | 12.18 | 15 | 18 | 14.94 | 17 | 23 | 6.71 | 9 | 11 | | | | 2.41 | 4 | 4 | | | | | | | | | | | | | 16.06 | 25 | 93 | 1700 | 188.3 | 5.50 |
| A6NNE9 | 9.70 | 1 | 3 | 3 | 3 | | | | 3.98 | 1 | 1 | | | | | | | | | | | | | | | | | | | | | | 5.72 | 2 | 2 | 402 | 43.8 | 6.92 |
| A6NNT2 | 2.54 | 1 | 3 | 3 | 3 | 0.00 | | | 0.00 | | | 0.00 | | | | | | | | | | | | | | | | | | | | | 2.54 | 3 | 3 | 1141 | 125.0 | 7.03 |
| A6NNW6 | 10.19 | 1 | 4 | 4 | 8 | | | | | | | | | | | | | 3.98 | 2 | 2 | | | | | | | | | | | | | 7.96 | 3 | 6 | 628 | 68.8 | 6.00 |
| A6PVS8 | 6.09 | 1 | 3 | 3 | 4 | | | | | | | | | | | | | | | | | | | | | | | | | | | | 6.09 | 3 | 4 | 624 | 73.6 | 9.72 |
| Q3ZCM7 | 31.08 | 3 | 1 | 15 | 465 | 24.32 | 9 | 25 | 24.32 | 9 | 26 | 24.32 | 9 | 19 | 25.00 | 11 | 38 | 27.03 | 11 | 54 | 3.38 | 1 | 1 | 10.59 | 4 | 8 | 14.64 | 6 | 12 | 22.52 | 8 | 17 | 28.38 | 12 | 265 | 444 | 49.7 | 4.89 |
| A7E2U8 | 64.08 | 1 | 15 | 15 | 51 | | | | 4.53 | 1 | 1 | | | | 27.18 | 6 | 9 | 57.93 | 13 | 19 | | | | | | | | | | | | | 56.96 | 12 | 22 | 309 | 34.4 | 9.64 |
| A8MV24 | 28.57 | 1 | 3 | 3 | 6 | | | | | | | | | | | | | | | | | | | | | | | | | | | | 28.57 | 3 | 6 | 154 | 17.6 | 9.79 |
| A8MVS5 | 16.96 | 1 | 2 | 2 | 4 | 6.52 | 1 | 1 | | | | | | | | | | | | | | | | | | | | | | | | | 16.96 | 2 | 3 | 230 | 24.7 | 7.30 |
| A8MYP8 | 80.24 | 1 | 19 | 19 | 65 | 14.62 | 3 | 3 | 17.79 | 4 | 4 | | | | 52.17 | 11 | 11 | 36.76 | 9 | 10 | | | | | | | | | | 6.32 | 1 | 1 | 73.52 | 15 | 36 | 253 | 27.3 | 10.42 |
| A8MYU2 | 5.05 | 1 | 5 | 5 | 7 | | | | | | | | | | | | | | | | | | | | | | | | | | | | 5.05 | 5 | 7 | 1149 | 129.5 | 7.83 |
| A8MYV0 | 46.76 | 1 | 19 | 19 | 69 | 14.65 | 5 | 5 | 17.46 | 6 | 7 | 6.76 | 2 | 2 | 7.61 | 3 | 3 | 6.48 | 2 | 2 | | | | 3.38 | 1 | 1 | 6.48 | 2 | 2 | 7.61 | 2 | 2 | 45.92 | 18 | 45 | 355 | 40.4 | 9.42 |
| A8MZ26 | 17.26 | 1 | 4 | 4 | 5 | | | | | | | | | | | | | | | | | | | | | | | | | | | | 17.26 | 4 | 5 | 197 | 23.9 | 8.85 |
| A8TX70 | 0.46 | 2 | 2 | 2 | 2 | | | | 0.00 | | | | | | | | | 0.23 | 1 | 1 | | | | | | | | | | | | | 0.23 | 1 | 1 | 2615 | 289.7 | 6.95 |
| B1AK53 | 9.37 | 1 | 8 | 8 | 29 | | | | | | | 1.52 | 1 | 1 | 5.15 | 4 | 4 | 3.86 | 3 | 4 | | | | | | | 1.76 | 1 | 1 | 5.74 | 4 | 5 | 9.37 | 7 | 14 | 854 | 91.7 | 6.93 |
| B1ANS9 | 24.14 | 1 | 26 | 26 | 67 | | | | | | | 0.00 | | | 0.65 | 1 | 1 | 2.68 | 3 | 3 | | | | | | | 0.65 | 1 | 1 | | | | 23.77 | 25 | 62 | 1081 | 123.6 | 8.19 |
| B2RUZ4 | 35.90 | 1 | 2 | 2 | 3 | | | | | | | | | | | | | | | | | | | | | | | | | | | | 35.90 | 2 | 3 | 78 | 8.7 | 9.03 |
| B2RV13 | 50.00 | 1 | 9 | 9 | 18 | | | | | | | | | | 12.20 | 2 | 2 | 12.20 | 2 | 2 | | | | | | | | | | | | | 49.39 | 8 | 14 | 164 | 19.5 | 10.07 |
| B3GLJ2 | 54.08 | 1 | 7 | 7 | 25 | 15.31 | 2 | 2 | 7.14 | 1 | 1 | | | | | | | | | | | | | | | | | | | | | | 54.08 | 7 | 22 | 98 | 11.7 | 8.97 |
| B3KS81 | 2.94 | 1 | 2 | 2 | 2 | | | | | | | | | | | | | | | | | | | | | | | | | | | | 2.94 | 2 | 2 | 715 | 80.3 | 12.06 |
| B7ZAQ6 | 13.63 | 1 | 4 | 4 | 5 | | | | 5.93 | 1 | 1 | 2.86 | 1 | 1 | | | | | | | | | | | | | | | | | | | 10.77 | 3 | 3 | 455 | 52.9 | 9.28 |
| **CON_00009866** | 33.41 | 1 | 2 | 18 | 39 | | | | 6.77 | 3 | 3 | 28.38 | 14 | 17 | | | | 2.62 | 1 | 1 | | | | 5.24 | 2 | 2 | 17.25 | 9 | 11 | 10.26 | 4 | 5 | | | | 458 | 49.6 | 4.96 |
| **CON_00689362** | 38.78 | 1 | 6 | 6 | 10 | | | | | | | | | | 10.20 | 1 | 1 | | | | | | | 11.56 | 2 | 2 | 28.57 | 4 | 5 | 10.20 | 1 | 2 | | | | 147 | 15.7 | 6.30 |
| **CON_00691361** | 23.94 | 2 | 2 | 8 | 24 | 16.31 | 5 | 7 | 4.03 | 2 | 2 | 10.81 | 4 | 4 | 9.11 | 4 | 4 | 4.45 | 2 | 2 | | | | 2.12 | 1 | 1 | 2.12 | 1 | 1 | 6.57 | 3 | 3 | | | | 472 | 51.2 | 7.52 |
| **CON_00701886** | 13.00 | 1 | 2 | 2 | 4 | | | | | | | | | | 13.00 | 2 | 2 | 13.00 | 2 | 2 | | | | | | | | | | | | | | | | 100 | 10.8 | 8.66 |
| **CON_00705721** | 7.24 | 2 | 2 | 5 | 18 | 1.33 | 1 | 1 | 5.52 | 3 | 3 | 3.24 | 2 | 2 | 3.62 | 2 | 2 | 1.33 | 1 | 1 | | | | 4.95 | 3 | 4 | 4.00 | 3 | 3 | 3.24 | 2 | 2 | | | | 525 | 57.4 | 6.61 |
| **CON_00706094** | 31.31 | 2 | 6 | 6 | 39 | | | | | | | | | | | | | 5.14 | 2 | 2 | | | | 3.27 | 1 | 1 | 7.01 | 1 | 1 | 7.01 | 1 | 2 | 31.31 | 6 | 29 | 214 | 24.5 | 5.02 |
| **CON_00708398** | 85.34 | 2 | 2 | 62 | 651 | 50.25 | 28 | 43 | 50.41 | 28 | 44 | 52.39 | 29 | 37 | 62.77 | 37 | 62 | 75.62 | 49 | 86 | 8.57 | 6 | 6 | 43.00 | 30 | 44 | 69.69 | 50 | 92 | 79.41 | 55 | 89 | 52.22 | 31 | 148 | 607 | 69.2 | 6.18 |
| **CON_00708398** | 85.71 | 2 | 2 | 62 | 483 | 48.44 | 26 | 40 | 48.60 | 26 | 41 | 50.57 | 27 | 35 | 60.92 | 35 | 55 | 74.88 | 48 | 84 | 8.54 | 6 | 6 | 41.22 | 28 | 41 | 67.82 | 48 | 89 | 79.80 | 55 | 88 | | | | 609 | 69.5 | 6.18 |
| **CON_00714405** | 37.10 | 1 | 6 | 10 | 24 | 24.19 | 5 | 6 | 20.56 | 5 | 6 | 10.48 | 2 | 2 | | | | 22.18 | 5 | 8 | | | | | | | | | | 10.08 | 2 | 2 | | | | 248 | 29.0 | 4.78 |
| **CON_LysC** | 30.63 | 1 | 17 | 17 | 331 | 20.83 | 7 | 35 | 26.80 | 10 | 41 | 30.63 | 13 | 73 | 25.11 | 12 | 36 | 23.12 | 12 | 28 | 15.47 | 4 | 5 | 23.12 | 14 | 66 | 15.93 | 11 | 25 | 23.12 | 12 | 22 | | | | 653 | 68.1 | 7.24 |
| **CON_P00711** | 28.87 | 1 | 3 | 3 | 5 | | | | | | | | | | | | | | | | | | | | | | | | | | | | 28.87 | 3 | 5 | 142 | 16.2 | 5.14 |
| **CON_P02663** | 22.52 | 2 | 4 | 4 | 16 | | | | | | | | | | | | | | | | | | | | | | | | | | | | 22.52 | 4 | 16 | 222 | 26.0 | 8.43 |
| **CON_P02668** | 20.53 | 2 | 3 | 3 | 29 | | | | | | | | | | | | | | | | | | | | | | | | | | | | 20.53 | 3 | 29 | 190 | 21.3 | 6.77 |
| **CON_P02754** | 55.62 | 1 | 6 | 9 | 35 | | | | | | | | | | | | | | | | | | | | | | | | | | | | 55.62 | 9 | 35 | 178 | 19.9 | 5.02 |
| **CON_Trypsin** | 58.87 | 1 | 12 | 12 | 1489 | 25.11 | 4 | 5 | 25.11 | 4 | 7 | 37.66 | 6 | 14 | 25.11 | 4 | 12 | 25.11 | 4 | 10 | 25.11 | 4 | 11 | 29.44 | 6 | 38 | 25.11 | 4 | 22 | 25.11 | 4 | 9 | 58.87 | 12 | 1361 | 231 | 24.4 | 7.18 |
| D6REC4 | 16.56 | 1 | 8 | 8 | 11 | | | | | | | | | | | | | 9.37 | 3 | 3 | | | | | | | | | | 2.40 | 1 | 1 | 13.07 | 5 | 7 | 459 | 52.3 | 10.55 |
| E9PJ23 | 24.71 | 4 | 2 | 11 | 36 | | | | 8.24 | 2 | 2 | 5.65 | 1 | 1 | 5.65 | 1 | 2 | 6.35 | 2 | 2 | | | | | | | | | | | | | 22.12 | 9 | 29 | 425 | 49.1 | 10.36 |
| H0YKK7 | 15.27 | 5 | 2 | 2 | 2 | | | | | | | | | | | | | 1.27 | 1 | 1 | | | | | | | | | | | | | 14.00 | 1 | 1 | 550 | 64.5 | 4.75 |
| H3BNL8 | 23.91 | 1 | 5 | 5 | 7 | | | | | | | | | | | | | | | | | | | | | | | | | | | | 23.91 | 5 | 7 | 230 | 26.5 | 8.90 |
| H3BRN8 | 74.38 | 1 | 7 | 7 | 17 | | | | | | | | | | 22.31 | 2 | 2 | 55.37 | 6 | 8 | | | | | | | | | | | | | 62.81 | 5 | 7 | 121 | 13.8 | 8.31 |
| H7C350 | 24.38 | 1 | 6 | 6 | 10 | | | | | | | | | | 1.74 | 1 | 1 | | | | | | | | | | | | | | | | 24.38 | 6 | 9 | 402 | 43.5 | 8.24 |
| L0R6Q1 | 15.53 | 1 | 2 | 2 | 2 | | | | | | | | | | | | | | | | | | | | | | | | | | | | 15.53 | 2 | 2 | 103 | 11.1 | 8.10 |
| O00116 | 48.63 | 1 | 24 | 24 | 44 | | | | 3.04 | 1 | 1 | | | | | | | | | | | | | | | | | | | | | | 48.63 | 24 | 43 | 658 | 72.9 | 7.34 |
| O00142 | 7.17 | 1 | 2 | 2 | 2 | | | | | | | | | | | | | | | | | | | | | | | | | | | | 7.17 | 2 | 2 | 265 | 31.0 | 8.46 |
| O00148 | 7.03 | 2 | 3 | 3 | 4 | | | | | | | | | | | | | | | | | | | | | | | | | | | | 7.03 | 3 | 4 | 427 | 49.1 | 5.68 |
| O00154 | 16.32 | 1 | 5 | 5 | 17 | 5.79 | 2 | 2 | 6.05 | 2 | 2 | | | | | | | | | | | | | | | | | | | 3.16 | 1 | 1 | 16.32 | 5 | 12 | 380 | 41.8 | 8.54 |
| O00159 | 33.77 | 1 | 30 | 30 | 78 | 1.22 | 1 | 1 | | | | 2.16 | 2 | 2 | | | | | | | | | | | | | | | | | | | 33.77 | 30 | 75 | 1063 | 121.6 | 9.41 |
| O00161 | 62.09 | 1 | 11 | 11 | 31 | 21.80 | 4 | 4 | 13.27 | 2 | 2 | 6.64 | 1 | 1 | | | | | | | | | | | | | | | | | | | 62.09 | 10 | 24 | 211 | 23.3 | 5.01 |
| **CON_00714673** | 26.90 | 1 | 1 | 49 | 283 | 21.66 | 40 | 77 | 10.68 | 19 | 24 | 24.59 | 45 | 91 | 18.10 | 33 | 43 | 8.42 | 17 | 22 | 2.09 | 4 | 4 | 2.64 | 5 | 5 | 5.03 | 9 | 11 | 3.27 | 6 | 6 | | | | 2387 | 262.3 | 5.74 |
| O00170 | 32.12 | 1 | 7 | 7 | 10 | | | | | | | | | | | | | | | | | | | | | | | | | 5.15 | 1 | 1 | 32.12 | 7 | 9 | 330 | 37.6 | 6.29 |
| O00178 | 7.62 | 1 | 4 | 4 | 6 | | | | | | | | | | | | | | | | | | | | | | | | | 1.64 | 1 | 1 | 5.98 | 3 | 5 | 669 | 72.4 | 8.34 |
| O00189 | 9.27 | 1 | 3 | 3 | 3 | | | | | | | | | | | | | | | | | | | | | | | | | | | | 9.27 | 3 | 3 | 453 | 49.9 | 7.20 |
| O00194 | 75.23 | 1 | 12 | 14 | 49 | 10.09 | 2 | 2 | 5.05 | 1 | 1 | 16.97 | 3 | 3 | 8.72 | 2 | 2 | | | | | | | 5.05 | 1 | 1 | | | | 5.05 | 1 | 1 | 75.23 | 14 | 39 | 218 | 24.6 | 5.52 |
| O00203 | 8.59 | 1 | 8 | 8 | 18 | | | | | | | | | | | | | | | | | | | | | | 1.74 | 1 | 1 | 2.01 | 2 | 2 | 7.77 | 7 | 15 | 1094 | 121.2 | 6.04 |
| O00204 | 68.22 | 1 | 18 | 18 | 67 | | | | | | | | | | 8.49 | 3 | 3 | | | | | | | | | | | | | 6.58 | 2 | 3 | 66.03 | 17 | 61 | 365 | 41.3 | 5.40 |
| O00217 | 46.67 | 1 | 8 | 8 | 11 | 4.29 | 1 | 1 | 4.29 | 1 | 1 | | | | | | | | | | | | | | | | | | | | | | 46.67 | 8 | 9 | 210 | 23.7 | 6.34 |
| O00231 | 64.45 | 1 | 28 | 28 | 103 | 23.70 | 8 | 8 | 17.77 | 6 | 7 | 11.37 | 4 | 4 | 23.22 | 8 | 10 | 2.61 | 1 | 1 | | | | 11.61 | 4 | 4 | 1.66 | 1 | 1 | 21.09 | 8 | 8 | 60.19 | 23 | 60 | 422 | 47.4 | 6.48 |
| O00232 | 57.24 | 3 | 27 | 27 | 103 | 7.02 | 2 | 2 | 18.20 | 6 | 7 | 12.06 | 5 | 5 | 7.68 | 3 | 3 | | | | | | | 2.19 | 1 | 1 | | | | 5.70 | 3 | 3 | 53.29 | 22 | 82 | 456 | 52.9 | 7.65 |
| O00233 | 8.97 | 1 | 2 | 2 | 2 | | | | 4.48 | 1 | 1 | | | | | | | | | | | | | | | | | | | | | | 4.48 | 1 | 1 | 223 | 24.7 | 6.95 |
| O00253 | 31.06 | 1 | 3 | 3 | 7 | 6.82 | 1 | 1 | | | | 6.82 | 1 | 1 | | | | | | | | | | | | | | | | | | | 31.06 | 3 | 5 | 132 | 14.4 | 7.44 |
| O00264 | 65.13 | 1 | 8 | 10 | 31 | 27.69 | 3 | 3 | 31.79 | 4 | 4 | 23.59 | 3 | 3 | 7.18 | 1 | 1 | 4.62 | 1 | 1 | | | | | | | | | | | | | 63.59 | 9 | 19 | 195 | 21.7 | 4.70 |
| O00291 | 14.18 | 2 | 13 | 13 | 18 | 2.22 | 2 | 2 | 2.60 | 2 | 2 | | | | | | | 0.87 | 1 | 1 | | | | | | | 1.64 | 1 | 1 | | | | 10.22 | 10 | 12 | 1037 | 116.1 | 5.30 |
| O00292 | 22.13 | 2 | 7 | 7 | 20 | | | | 10.38 | 3 | 3 | | | | | | | | | | | | | | | | | | | | | | 22.13 | 7 | 17 | 366 | 40.9 | 8.56 |
| O00299 | 81.33 | 2 | 18 | 19 | 107 | 22.82 | 4 | 4 | 32.37 | 5 | 5 | 38.59 | 6 | 6 | 43.15 | 8 | 8 | 22.82 | 3 | 3 | 4.98 | 1 | 1 | 29.88 | 5 | 5 | 22.82 | 4 | 4 | 48.96 | 10 | 10 | 80.50 | 18 | 61 | 241 | 26.9 | 5.17 |
| O00303 | 28.57 | 1 | 9 | 9 | 24 | | | | | | | | | | 9.52 | 3 | 3 | 3.08 | 1 | 1 | | | | | | | 3.08 | 1 | 1 | | | | 28.57 | 9 | 19 | 357 | 37.5 | 5.45 |
| O00330 | 25.15 | 1 | 11 | 11 | 45 | 13.37 | 6 | 8 | 20.16 | 8 | 10 | 14.77 | 6 | 6 | 4.79 | 2 | 2 | 2.79 | 1 | 2 | | | | | | | | | | | | | 25.15 | 11 | 17 | 501 | 54.1 | 8.66 |
| O00339 | 6.90 | 1 | 5 | 5 | 10 | | | | 1.15 | 1 | 1 | 2.30 | 2 | 2 | | | | | | | | | | | | | | | | | | | 6.90 | 5 | 7 | 956 | 106.8 | 6.27 |
| O00391 | 9.91 | 1 | 6 | 6 | 13 | 2.14 | 1 | 2 | 4.55 | 3 | 4 | | | | | | | | | | | | | | | | | | | | | | 5.35 | 3 | 7 | 747 | 82.5 | 8.92 |
| O00399 | 12.63 | 1 | 2 | 2 | 3 | | | | | | | 5.26 | 1 | 1 | | | | | | | | | | | | | 5.26 | 1 | 1 | | | | 7.37 | 1 | 1 | 190 | 20.7 | 6.32 |
| O00400 | 15.85 | 1 | 8 | 8 | 17 | | | | | | | | | | | | | | | | | | | | | | | | | | | | 15.85 | 8 | 17 | 549 | 60.9 | 7.33 |
| O00401 | 7.92 | 1 | 2 | 2 | 4 | | | | | | | | | | | | | | | | | | | | | | | | | | | | 7.92 | 2 | 4 | 505 | 54.8 | 7.93 |
| O00410 | 44.48 | 2 | 39 | 39 | 210 | 6.29 | 5 | 6 | 9.66 | 7 | 9 | 5.83 | 5 | 5 | 6.75 | 6 | 7 | 1.37 | 1 | 1 | | | | | | | | | | 1.37 | 1 | 1 | 43.85 | 38 | 181 | 1097 | 123.5 | 4.94 |
| O00429 | 28.40 | 1 | 13 | 13 | 33 | | | | | | | | | | | | | | | | | | | | | | | | | | | | 28.40 | 13 | 33 | 736 | 81.8 | 6.81 |
| O00442 | 15.30 | 1 | 5 | 5 | 7 | | | | | | | | | | | | | | | | | | | | | | | | | | | | 15.30 | 5 | 7 | 366 | 39.3 | 7.85 |
| O00443 | 2.37 | 1 | 3 | 3 | 3 | | | | | | | | | | | | | | | | | | | | | | | | | | | | 2.37 | 3 | 3 | 1686 | 190.6 | 8.02 |
| O00462 | 14.22 | 1 | 10 | 11 | 13 | | | | | | | | | | 0.80 | 1 | 1 | | | | | | | | | | | | | | | | 14.22 | 11 | 12 | 879 | 100.8 | 5.52 |
| O00468 | 36.82 | 2 | 59 | 59 | 151 | 8.71 | 12 | 13 | 4.64 | 7 | 7 | 12.19 | 18 | 20 | | | | 2.95 | 3 | 3 | | | | | | | | | | | | | 35.80 | 56 | 108 | 2067 | 217.1 | 6.39 |
| O00471 | 4.80 | 1 | 3 | 3 | 4 | | | | | | | | | | | | | | | | | | | | | | | | | | | | 4.80 | 3 | 4 | 708 | 81.8 | 6.71 |
| O00487 | 59.35 | 1 | 12 | 12 | 35 | 10.32 | 2 | 3 | 9.03 | 3 | 3 | 12.58 | 2 | 2 | 14.52 | 5 | 5 | 4.19 | 2 | 2 | | | | 6.13 | 2 | 2 | | | | 9.35 | 3 | 3 | 57.10 | 10 | 15 | 310 | 34.6 | 6.52 |
| O00505 | 42.23 | 1 | 10 | 15 | 39 | 6.14 | 3 | 3 | 13.44 | 5 | 6 | | | | | | | 1.92 | 1 | 1 | | | | | | | 1.34 | 1 | 1 | | | | 39.16 | 12 | 28 | 521 | 57.8 | 4.94 |
| O00519 | 50.09 | 1 | 19 | 19 | 47 | 6.91 | 3 | 3 | 12.95 | 5 | 5 | 6.04 | 3 | 3 | | | | | | | | | | | | | | | | | | | 44.56 | 17 | 36 | 579 | 63.0 | 7.66 |
| O00560 | 26.85 | 1 | 5 | 5 | 7 | | | | 3.69 | 1 | 1 | 3.69 | 1 | 1 | | | | | | | | | | | | | | | | | | | 23.15 | 4 | 5 | 298 | 32.4 | 7.53 |
| O00571 | 34.14 | 1 | 3 | 20 | 62 | 1.51 | 1 | 1 | | | | 6.65 | 4 | 4 | 4.83 | 3 | 3 | 1.66 | 1 | 1 | | | | 3.02 | 2 | 2 | 1.66 | 1 | 1 | 4.98 | 3 | 3 | 29.46 | 17 | 47 | 662 | 73.2 | 7.18 |
| O00602 | 26.07 | 1 | 6 | 6 | 11 | 4.91 | 2 | 2 | | | | 12.88 | 3 | 3 | 12.27 | 3 | 3 | | | | | | | | | | | | | | | | 17.48 | 3 | 3 | 326 | 35.1 | 6.86 |
| O00622 | 9.71 | 1 | 4 | 4 | 5 | | | | | | | | | | | | | | | | | | | | | | | | | | | | 9.71 | 4 | 5 | 381 | 42.0 | 8.21 |
| O00629 | 39.35 | 1 | 8 | 12 | 23 | 5.76 | 2 | 2 | | | | | | | | | | | | | | | | | | | 1.34 | 1 | 1 | | | | 37.62 | 11 | 20 | 521 | 57.9 | 4.96 |
| O00743 | 61.31 | 1 | 16 | 16 | 45 | 4.26 | 1 | 1 | | | | 3.28 | 1 | 1 | 10.16 | 3 | 3 | | | | | | | 3.28 | 1 | 1 | | | | 9.84 | 3 | 3 | 57.70 | 15 | 36 | 305 | 35.1 | 5.69 |
| O00764 | 48.72 | 1 | 11 | 11 | 44 | 5.45 | 1 | 1 | | | | | | | 3.21 | 1 | 1 | | | | | | | 6.41 | 2 | 2 | | | | 6.41 | 2 | 2 | 48.72 | 11 | 38 | 312 | 35.1 | 6.13 |
| O14494 | 20.42 | 1 | 5 | 5 | 53 | 15.85 | 3 | 4 | 10.56 | 2 | 2 | 12.32 | 2 | 2 | | | | | | | | | | | | | | | | | | | 20.42 | 5 | 45 | 284 | 32.1 | 7.97 |
| O14520 | 30.70 | 2 | 8 | 8 | 44 | | | | 7.02 | 2 | 2 | 4.68 | 2 | 2 | | | | | | | | | | | | | | | | | | | 30.70 | 8 | 40 | 342 | 37.2 | 8.95 |
| O14521 | 14.47 | 1 | 2 | 3 | 7 | | | | 3.77 | 1 | 1 | | | | | | | | | | | | | | | | | | | | | | 14.47 | 3 | 6 | 159 | 17.0 | 8.63 |
| O14556 | 96.32 | 1 | 48 | 49 | 1225 | 58.33 | 18 | 50 | 46.08 | 12 | 29 | 52.21 | 16 | 64 | 63.73 | 24 | 55 | 54.66 | 22 | 30 | 2.70 | 1 | 1 | 56.62 | 22 | 40 | 47.79 | 24 | 45 | 66.18 | 30 | 55 | 91.18 | 36 | 856 | 408 | 44.5 | 8.19 |
| O14561 | 19.23 | 1 | 3 | 3 | 4 | 19.23 | 3 | 3 | | | | | | | | | | 3.85 | 1 | 1 | | | | | | | | | | | | | | | | 156 | 17.4 | 4.93 |
| O14579 | 47.08 | 1 | 11 | 11 | 19 | 4.55 | 1 | 1 | | | | | | | | | | | | | | | | 3.90 | 1 | 1 | | | | 9.42 | 2 | 2 | 39.29 | 8 | 15 | 308 | 34.5 | 5.12 |
| O14617 | 9.71 | 1 | 8 | 8 | 23 | | | | | | | | | | | | | | | | | | | | | | | | | | | | 9.71 | 8 | 23 | 1153 | 130.1 | 8.48 |
| O14618 | 26.28 | 1 | 5 | 5 | 8 | | | | | | | | | | 2.55 | 1 | 1 | | | | | | | 3.28 | 1 | 1 | 13.14 | 3 | 3 | | | | 16.42 | 3 | 3 | 274 | 29.0 | 5.58 |
| O14638 | 21.71 | 1 | 13 | 13 | 16 | | | | | | | | | | | | | | | | | | | | | | | | | | | | 21.71 | 13 | 16 | 875 | 100.1 | 6.57 |
| O14641 | 3.13 | 2 | 2 | 2 | 2 | | | | | | | | | | | | | | | | | | | | | | | | | | | | 3.13 | 2 | 2 | 736 | 78.9 | 6.02 |
| O14645 | 76.36 | 1 | 29 | 29 | 114 | 18.22 | 4 | 4 | 25.58 | 6 | 6 | 9.69 | 2 | 2 | 36.43 | 10 | 13 | 56.20 | 21 | 33 | 10.85 | 2 | 3 | 6.59 | 1 | 1 | 5.43 | 1 | 1 | 16.28 | 3 | 3 | 63.18 | 16 | 48 | 258 | 29.6 | 8.50 |
| O14683 | 13.23 | 1 | 2 | 2 | 9 | 7.94 | 1 | 2 | 13.23 | 2 | 3 | | | | | | | | | | | | | | | | | | | | | | 13.23 | 2 | 4 | 189 | 21.0 | 9.55 |
| O14684 | 13.16 | 1 | 2 | 2 | 3 | | | | | | | | | | | | | | | | | | | | | | | | | | | | 13.16 | 2 | 3 | 152 | 17.1 | 9.50 |
| O14715 | 21.25 | 3 | 13 | 33 | 61 | | | | | | | | | | | | | | | | | | | | | | | | | | | | 21.25 | 33 | 61 | 1765 | 198.9 | 6.49 |
| O14734 | 38.56 | 1 | 9 | 9 | 15 | | | | 8.78 | 2 | 2 | | | | | | | | | | | | | | | | | | | | | | 32.60 | 8 | 13 | 319 | 35.9 | 7.56 |
| O14735 | 15.96 | 1 | 3 | 3 | 3 | | | | | | | | | | | | | | | | | | | | | | | | | | | | 15.96 | 3 | 3 | 213 | 23.5 | 8.03 |
| O14737 | 19.20 | 1 | 2 | 2 | 4 | | | | 8.80 | 1 | 1 | | | | | | | 10.40 | 1 | 1 | | | | | | | | | | | | | 10.40 | 1 | 2 | 125 | 14.3 | 6.04 |
| O14744 | 48.35 | 1 | 28 | 28 | 82 | 10.99 | 6 | 6 | 9.26 | 6 | 6 | 2.83 | 2 | 2 | 1.57 | 1 | 1 | 1.57 | 1 | 1 | | | | | | | | | | | | | 46.78 | 26 | 66 | 637 | 72.6 | 6.29 |
| O14745 | 13.97 | 1 | 4 | 4 | 6 | 7.82 | 2 | 2 | | | | 3.91 | 1 | 1 | | | | | | | | | | | | | | | | 6.70 | 2 | 2 | 3.35 | 1 | 1 | 358 | 38.8 | 5.77 |
| O14773 | 12.97 | 1 | 4 | 4 | 16 | 7.28 | 2 | 3 | 5.68 | 2 | 2 | 2.49 | 1 | 1 | | | | | | | | | | | | | | | | | | | 12.97 | 4 | 10 | 563 | 61.2 | 6.48 |
| O14786 | 5.42 | 1 | 3 | 3 | 9 | 0.00 | | | | | | | | | | | | | | | | | | | | | | | | | | | 5.42 | 3 | 9 | 923 | 103.1 | 5.88 |
| O14787 | 20.18 | 1 | 7 | 12 | 25 | | | | | | | | | | | | | | | | | | | | | | | | | | | | 20.18 | 12 | 25 | 897 | 101.3 | 5.01 |
| O14795 | 6.79 | 3 | 9 | 9 | 10 | | | | | | | | | | | | | | | | | | | | | | | | | | | | 6.79 | 9 | 10 | 1591 | 180.6 | 5.99 |
| O14818 | 56.05 | 1 | 7 | 12 | 57 | 30.65 | 6 | 8 | 41.94 | 8 | 10 | 33.47 | 6 | 7 | 4.44 | 1 | 1 | 4.44 | 1 | 1 | | | | 7.26 | 2 | 2 | | | | 4.44 | 1 | 1 | 52.42 | 11 | 27 | 248 | 27.9 | 8.46 |
| O14828 | 18.73 | 1 | 4 | 4 | 20 | 3.75 | 1 | 1 | 18.73 | 4 | 5 | 14.12 | 3 | 3 | | | | | | | | | | | | | | | | | | | 18.73 | 4 | 11 | 347 | 38.3 | 7.64 |
| O14829 | 46.55 | 1 | 28 | 28 | 110 | 11.64 | 7 | 7 | 12.71 | 8 | 10 | 11.03 | 7 | 9 | 1.68 | 1 | 1 | | | | | | | | | | | | | | | | 46.55 | 28 | 83 | 653 | 75.7 | 6.84 |
| O14841 | 46.20 | 3 | 47 | 47 | 194 | | | | | | | 3.80 | 4 | 4 | | | | 7.14 | 7 | 7 | | | | | | | 8.46 | 8 | 8 | 3.49 | 4 | 4 | 44.49 | 42 | 171 | 1288 | 137.4 | 6.58 |
| O14863 | 6.76 | 1 | 2 | 2 | 3 | | | | | | | | | | | | | | | | | | | | | | | | | | | | 6.76 | 2 | 3 | 429 | 47.5 | 6.58 |
| O14867 | 6.79 | 1 | 4 | 4 | 7 | | | | | | | | | | | | | | | | | | | | | | 1.36 | 1 | 1 | 1.36 | 1 | 1 | 6.79 | 4 | 5 | 736 | 81.9 | 5.05 |
| Q562R1 | 20.21 | 1 | 1 | 10 | 201 | 10.90 | 4 | 13 | 12.77 | 5 | 16 | 17.55 | 6 | 16 | 14.63 | 6 | 16 | 14.63 | 5 | 17 | 2.66 | 1 | 1 | 12.77 | 5 | 14 | 20.21 | 7 | 16 | 12.77 | 5 | 15 | 13.30 | 7 | 77 | 376 | 42.0 | 5.59 |
| O14880 | 18.42 | 1 | 3 | 3 | 6 | | | | | | | | | | | | | | | | | | | | | | | | | | | | 18.42 | 3 | 6 | 152 | 16.5 | 9.38 |
| O14893 | 7.50 | 1 | 2 | 2 | 2 | | | | | | | | | | | | | | | | | | | | | | | | | | | | 7.50 | 2 | 2 | 280 | 31.6 | 5.58 |
| O14896 | 20.99 | 1 | 7 | 7 | 17 | 3.43 | 1 | 1 | | | | | | | 2.57 | 1 | 1 | | | | | | | | | | | | | | | | 18.42 | 6 | 15 | 467 | 53.1 | 5.33 |
| O14910 | 45.92 | 2 | 5 | 8 | 17 | | | | | | | | | | 9.01 | 2 | 2 | | | | | | | 12.02 | 3 | 3 | 4.72 | 1 | 1 | 4.72 | 1 | 1 | 45.92 | 8 | 10 | 233 | 26.0 | 8.72 |
| O14920 | 2.78 | 1 | 2 | 2 | 2 | | | | | | | | | | | | | | | | | | | | | | | | | | | | 2.78 | 2 | 2 | 756 | 86.5 | 5.78 |
| O14925 | 18.66 | 2 | 2 | 2 | 2 | | | | | | | | | | | | | | | | | | | | | | | | | | | | 18.66 | 2 | 2 | 209 | 21.9 | 8.60 |
| O14933 | 26.14 | 1 | 3 | 3 | 3 | | | | | | | | | | | | | | | | | | | | | | | | | | | | 26.14 | 3 | 3 | 153 | 17.8 | 7.88 |
| O14949 | 47.56 | 1 | 6 | 6 | 10 | 15.85 | 1 | 1 | 15.85 | 1 | 1 | | | | | | | | | | | | | | | | | | | | | | 47.56 | 6 | 8 | 82 | 9.9 | 10.08 |
| O14950 | 69.77 | 3 | 10 | 10 | 38 | 17.44 | 3 | 3 | 5.81 | 1 | 1 | 18.02 | 2 | 2 | 18.02 | 4 | 5 | 6.40 | 2 | 2 | | | | | | | | | | 6.40 | 1 | 2 | 69.19 | 8 | 21 | 172 | 19.8 | 4.84 |
| O14964 | 5.79 | 1 | 4 | 4 | 7 | | | | | | | | | | 2.96 | 2 | 2 | | | | | | | | | | | | | | | | 4.50 | 3 | 5 | 777 | 86.1 | 6.16 |
| O14967 | 44.92 | 1 | 18 | 19 | 53 | 8.03 | 4 | 5 | 9.02 | 5 | 5 | 18.36 | 7 | 7 | 4.59 | 3 | 3 | 0.98 | 1 | 1 | | | | | | | | | | | | | 36.07 | 15 | 32 | 610 | 70.0 | 4.69 |
| O14975 | 9.52 | 1 | 5 | 5 | 9 | | | | | | | | | | | | | | | | | | | | | | | | | | | | 9.52 | 5 | 9 | 620 | 70.3 | 8.51 |
| O14980 | 45.75 | 1 | 39 | 39 | 132 | 5.42 | 4 | 4 | 7.75 | 6 | 7 | 5.79 | 4 | 4 | 0.93 | 1 | 1 | | | | | | | | | | | | | | | | 45.75 | 38 | 116 | 1071 | 123.3 | 6.06 |
| O15020 | 5.77 | 1 | 7 | 12 | 17 | 0.00 | | | 0.00 | | | 0.00 | | | | | | 0.00 | | | | | | 0.00 | | | 0.00 | | | 0.59 | 1 | 1 | 5.77 | 12 | 16 | 2390 | 271.2 | 6.11 |
| O15066 | 4.42 | 3 | 2 | 3 | 5 | 0.00 | | | 0.00 | | | | | | | | | | | | | | | | | | | | | 2.95 | 2 | 2 | 1.47 | 1 | 3 | 747 | 85.1 | 7.69 |
| O15078 | 1.53 | 4 | 4 | 5 | 6 | 0.00 | | | | | | 0.00 | | | 0.00 | | | 0.65 | 2 | 2 | | | | | | | | | | | | | 1.13 | 4 | 4 | 2479 | 290.2 | 5.95 |
| O15084 | 27.07 | 3 | 20 | 20 | 38 | 0.85 | 1 | 1 | 0.85 | 1 | 1 | 1.42 | 1 | 1 | | | | | | | | | | | | | 1.14 | 1 | 2 | | | | 25.64 | 19 | 33 | 1053 | 112.9 | 6.25 |
| O15085 | 0.85 | 1 | 2 | 2 | 2 | | | | | | | 0.85 | 2 | 2 | | | | | | | | | | | | | | | | | | | | | | 1522 | 167.6 | 5.50 |
| O15118 | 4.15 | 2 | 5 | 5 | 7 | | | | | | | | | | | | | | | | | | | | | | | | | | | | 4.15 | 5 | 7 | 1278 | 142.1 | 5.36 |
| O15120 | 13.67 | 1 | 3 | 3 | 4 | | | | | | | | | | | | | | | | | | | | | | | | | | | | 13.67 | 3 | 4 | 278 | 30.9 | 9.01 |
| O15126 | 10.65 | 1 | 2 | 2 | 5 | | | | | | | | | | | | | | | | | | | | | | | | | | | | 10.65 | 2 | 5 | 338 | 37.9 | 7.42 |
| O15127 | 10.33 | 1 | 3 | 3 | 22 | | | | 10.33 | 2 | 2 | 6.08 | 1 | 1 | | | | | | | | | | | | | | | | | | | 9.73 | 2 | 19 | 329 | 36.6 | 6.10 |
| O15143 | 30.91 | 1 | 11 | 11 | 28 | | | | | | | 8.33 | 2 | 2 | 5.11 | 1 | 2 | | | | | | | | | | | | | | | | 30.65 | 10 | 24 | 372 | 40.9 | 8.35 |
| O15144 | 49.33 | 1 | 13 | 13 | 18 | | | | | | | 7.67 | 2 | 2 | 13.33 | 4 | 4 | | | | | | | | | | | | | | | | 42.00 | 10 | 12 | 300 | 34.3 | 7.36 |
| O15145 | 39.89 | 1 | 7 | 7 | 19 | 13.48 | 2 | 2 | | | | | | | 6.18 | 1 | 1 | | | | | | | | | | | | | | | | 39.89 | 7 | 16 | 178 | 20.5 | 8.59 |
| O15155 | 24.58 | 1 | 2 | 2 | 3 | | | | | | | | | | | | | | | | | | | | | | | | | | | | 24.58 | 2 | 3 | 118 | 13.3 | 9.06 |
| O15173 | 52.91 | 1 | 10 | 12 | 30 | 18.39 | 3 | 3 | 26.91 | 4 | 4 | 10.31 | 2 | 2 | | | | 7.62 | 2 | 2 | | | | | | | | | | | | | 52.02 | 10 | 19 | 223 | 23.8 | 4.88 |
| O15228 | 38.97 | 1 | 21 | 21 | 34 | | | | 1.76 | 1 | 1 | | | | | | | | | | | | | | | | | | | | | | 37.21 | 20 | 33 | 680 | 77.1 | 6.57 |
| O15230 | 23.17 | 1 | 61 | 61 | 120 | 4.44 | 12 | 12 | 2.41 | 6 | 6 | 13.13 | 33 | 37 | | | | | | | | | | | | | | | | | | | 15.94 | 43 | 65 | 3695 | 399.5 | 7.02 |
| O15231 | 4.35 | 1 | 2 | 2 | 5 | | | | | | | | | | | | | | | | | | | | | | 4.35 | 2 | 3 | | | | 2.47 | 1 | 2 | 689 | 73.5 | 7.01 |
| O15254 | 5.00 | 1 | 3 | 3 | 3 | | | | | | | | | | | | | | | | | | | | | | | | | | | | 5.00 | 3 | 3 | 700 | 77.6 | 7.25 |
| O15258 | 24.49 | 1 | 3 | 3 | 20 | 15.31 | 2 | 2 | 15.31 | 2 | 2 | | | | | | | | | | | | | | | | | | | | | | 24.49 | 3 | 16 | 196 | 22.9 | 9.54 |
| O15260 | 21.56 | 1 | 5 | 5 | 34 | 10.41 | 2 | 4 | 3.72 | 1 | 1 | 6.69 | 1 | 1 | | | | | | | | | | | | | | | | | | | 21.56 | 5 | 28 | 269 | 30.4 | 7.78 |
| O15264 | 6.85 | 7 | 2 | 3 | 7 | | | | | | | | | | | | | 0.00 | | | | | | | | | | | | | | | 6.85 | 3 | 7 | 365 | 42.1 | 8.38 |
| O15294 | 1.82 | 1 | 2 | 2 | 2 | | | | | | | | | | | | | | | | | | | | | | | | | | | | 1.82 | 2 | 2 | 1046 | 116.8 | 6.70 |
| P53675 | 12.50 | 1 | 1 | 24 | 173 | 5.24 | 9 | 10 | 5.73 | 8 | 12 | 7.62 | 13 | 16 | 4.15 | 6 | 7 | 0.73 | 1 | 1 | | | | 1.59 | 2 | 2 | | | | 1.59 | 2 | 2 | 12.07 | 21 | 123 | 1640 | 186.9 | 5.85 |
| O15296 | 30.03 | 2 | 14 | 14 | 69 | 4.88 | 2 | 5 | | | | 17.01 | 7 | 7 | 3.85 | 2 | 2 | | | | | | | 3.25 | 2 | 2 | 3.25 | 1 | 1 | 5.33 | 2 | 2 | 26.78 | 12 | 50 | 676 | 75.8 | 6.10 |
| O15305 | 34.96 | 2 | 8 | 8 | 14 | 4.88 | 1 | 1 | | | | | | | | | | | | | | | | | | | | | | 4.88 | 1 | 1 | 34.96 | 8 | 12 | 246 | 28.1 | 6.77 |
| O15320 | 9.20 | 2 | 2 | 5 | 8 | | | | | | | | | | | | | 1.49 | 1 | 1 | | | | | | | | | | | | | 7.71 | 4 | 7 | 804 | 90.9 | 5.22 |
| O15327 | 7.47 | 1 | 4 | 4 | 9 | | | | | | | | | | | | | | | | | | | | | | | | | 2.16 | 1 | 1 | 7.47 | 4 | 8 | 924 | 104.7 | 6.27 |
| O15371 | 24.27 | 1 | 9 | 9 | 17 | | | | 2.19 | 1 | 1 | | | | | | | | | | | | | | | | | | | 5.66 | 2 | 2 | 18.61 | 7 | 14 | 548 | 63.9 | 6.05 |
| O15372 | 28.13 | 1 | 9 | 9 | 19 | 5.11 | 1 | 1 | 10.51 | 2 | 2 | | | | 5.40 | 1 | 2 | 6.53 | 2 | 2 | 6.53 | 1 | 1 | | | | 5.40 | 1 | 1 | 5.40 | 1 | 1 | 24.72 | 7 | 9 | 352 | 39.9 | 6.54 |
| O15382 | 5.61 | 1 | 2 | 2 | 2 | | | | | | | | | | | | | | | | | | | | | | | | | | | | 5.61 | 2 | 2 | 392 | 44.3 | 8.65 |
| O15393 | 40.85 | 1 | 13 | 13 | 60 | 1.63 | 1 | 1 | 1.63 | 1 | 1 | 8.54 | 3 | 3 | 7.11 | 4 | 4 | 5.49 | 3 | 3 | 2.24 | 1 | 1 | 5.49 | 3 | 3 | 3.25 | 2 | 2 | 3.66 | 2 | 2 | 40.85 | 13 | 40 | 492 | 53.8 | 7.81 |
| O15397 | 6.46 | 1 | 5 | 5 | 8 | | | | 0.00 | | | | | | | | | | | | | | | | | | | | | | | | 6.46 | 5 | 8 | 1037 | 119.9 | 5.16 |
| O15400 | 9.58 | 1 | 2 | 2 | 4 | 9.58 | 2 | 2 | 5.75 | 1 | 1 | | | | | | | | | | | | | | | | | | | | | | 5.75 | 1 | 1 | 261 | 29.8 | 5.55 |
| O15484 | 13.28 | 1 | 6 | 6 | 11 | | | | | | | | | | 0.00 | | | | | | | | | | | | | | | | | | 13.28 | 6 | 11 | 640 | 73.1 | 7.64 |
| O15498 | 10.61 | 1 | 2 | 2 | 4 | | | | | | | | | | | | | | | | | | | | | | | | | | | | 10.61 | 2 | 4 | 198 | 22.4 | 6.92 |
| O15511 | 43.71 | 1 | 4 | 4 | 8 | 11.92 | 2 | 2 | | | | | | | | | | | | | | | | | | | | | | | | | 43.71 | 4 | 6 | 151 | 16.3 | 5.67 |
| O15523 | 32.42 | 1 | 2 | 19 | 59 | 1.52 | 1 | 1 | | | | 6.67 | 4 | 4 | 4.85 | 3 | 3 | 1.67 | 1 | 1 | | | | 3.03 | 2 | 2 | 1.67 | 1 | 1 | 5.00 | 3 | 3 | 27.73 | 16 | 44 | 660 | 73.1 | 7.55 |
| O43149 | 2.06 | 1 | 4 | 5 | 10 | | | | | | | | | | | | | | | | | | | | | | | | | | | | 2.06 | 5 | 10 | 2961 | 330.9 | 5.95 |
| P50993 | 25.49 | 2 | 1 | 22 | 161 | 12.65 | 9 | 11 | 13.43 | 11 | 14 | 12.45 | 11 | 11 | 1.86 | 2 | 2 | 2.55 | 3 | 3 | | | | | | | | | | | | | 25.49 | 22 | 120 | 1020 | 112.2 | 5.66 |
| O43169 | 50.68 | 1 | 4 | 4 | 10 | 23.29 | 1 | 1 | 29.45 | 2 | 3 | | | | | | | | | | | | | | | | | | | | | | 44.52 | 3 | 6 | 146 | 16.3 | 4.97 |
| O43175 | 62.66 | 1 | 28 | 28 | 238 | 14.82 | 6 | 6 | 12.01 | 6 | 6 | 16.89 | 8 | 8 | 33.21 | 15 | 16 | 17.07 | 8 | 9 | 4.69 | 2 | 2 | 18.01 | 10 | 12 | 22.70 | 10 | 12 | 35.83 | 16 | 20 | 60.04 | 24 | 145 | 533 | 56.6 | 6.71 |
| O43181 | 17.14 | 1 | 3 | 3 | 5 | 8.57 | 1 | 1 | 8.57 | 1 | 1 | | | | | | | | | | | | | | | | | | | | | | 17.14 | 3 | 3 | 175 | 20.1 | 10.30 |
| O43182 | 2.77 | 1 | 2 | 2 | 3 | | | | | | | | | | | | | | | | | | | | | | | | | | | | 2.77 | 2 | 3 | 974 | 105.9 | 7.36 |
| O43236 | 37.87 | 1 | 17 | 18 | 65 | 1.88 | 1 | 1 | 7.74 | 3 | 3 | | | | | | | 15.27 | 8 | 10 | | | | | | | 4.60 | 2 | 2 | 1.88 | 1 | 1 | 35.15 | 13 | 48 | 478 | 55.1 | 6.11 |
| O43237 | 27.85 | 1 | 8 | 8 | 12 | | | | | | | | | | | | | | | | | | | | | | | | | | | | 27.85 | 8 | 12 | 492 | 54.1 | 6.38 |
| O43242 | 61.80 | 1 | 35 | 35 | 116 | 15.73 | 6 | 7 | 17.60 | 8 | 11 | 5.24 | 3 | 3 | 12.55 | 5 | 5 | 16.10 | 4 | 6 | | | | | | | | | | 2.81 | 1 | 1 | 55.06 | 31 | 83 | 534 | 60.9 | 8.44 |
| O43247 | 49.64 | 1 | 10 | 10 | 27 | 11.07 | 2 | 2 | 9.64 | 2 | 3 | | | | 20.71 | 4 | 4 | 41.79 | 9 | 11 | | | | | | | | | | | | | 23.21 | 5 | 7 | 280 | 30.7 | 7.99 |
| O43264 | 17.33 | 1 | 9 | 9 | 21 | | | | 1.67 | 1 | 1 | | | | | | | | | | | | | | | | | | | | | | 17.33 | 9 | 20 | 779 | 88.8 | 6.27 |
| Q14568 | 29.15 | 1 | 1 | 10 | 156 | 12.54 | 4 | 4 | 17.49 | 6 | 7 | 17.49 | 6 | 7 | 23.32 | 7 | 7 | 22.16 | 7 | 9 | 2.62 | 1 | 1 | 20.70 | 6 | 9 | 20.12 | 6 | 7 | 25.36 | 8 | 13 | 27.70 | 9 | 92 | 343 | 39.3 | 4.65 |
| O43278 | 12.29 | 1 | 7 | 7 | 9 | | | | 2.08 | 1 | 1 | 2.08 | 1 | 1 | | | | | | | | | | | | | | | | | | | 12.29 | 6 | 7 | 529 | 58.4 | 6.29 |
| O43324 | 57.47 | 1 | 8 | 8 | 23 | 24.14 | 3 | 3 | 18.39 | 3 | 3 | | | | 5.75 | 1 | 1 | | | | | | | 6.32 | 1 | 1 | 6.32 | 1 | 1 | 6.32 | 1 | 1 | 45.40 | 7 | 12 | 174 | 19.8 | 8.54 |
| O43379 | 8.04 | 1 | 9 | 9 | 20 | | | | | | | 0.79 | 1 | 1 | | | | | | | | | | | | | | | | 0.86 | 1 | 1 | 7.18 | 8 | 18 | 1518 | 165.8 | 5.91 |
| O43390 | 7.27 | 1 | 3 | 4 | 5 | | | | | | | | | | | | | | | | 1.90 | 1 | 1 | | | | | | | 1.26 | 1 | 1 | 4.11 | 2 | 3 | 633 | 70.9 | 8.13 |
| O43396 | 62.98 | 1 | 15 | 15 | 69 | 6.23 | 2 | 2 | 14.19 | 3 | 5 | 2.42 | 1 | 1 | 8.30 | 2 | 2 | 21.11 | 4 | 5 | | | | 16.61 | 4 | 4 | 22.49 | 5 | 6 | 30.10 | 6 | 8 | 50.87 | 12 | 36 | 289 | 32.2 | 4.96 |
| O43414 | 9.20 | 1 | 2 | 2 | 3 | | | | | | | | | | | | | | | | | | | | | | | | | | | | 9.20 | 2 | 3 | 337 | 37.2 | 8.07 |
| O43427 | 13.19 | 1 | 4 | 4 | 4 | | | | | | | | | | | | | | | | | | | | | | | | | | | | 13.19 | 4 | 4 | 364 | 41.9 | 6.48 |
| O43432 | 4.48 | 1 | 4 | 7 | 7 | 0.63 | 1 | 1 | | | | | | | | | | | | | | | | | | | | | | | | | 3.85 | 6 | 6 | 1585 | 176.5 | 5.38 |

| | | | | | | | | | | | | | | | | | | | | | | | | | | | | | | | | | | | | | | |
|---|---|---|---|---|---|---|---|---|---|---|---|---|---|---|---|---|---|---|---|---|---|---|---|---|---|---|---|---|---|---|---|---|---|---|---|---|---|---|
|O43451|28.70|3|44|48|127|10.07|13|16|7.75|11|12|24.50|35|48|||||||||||||||||||19.06|28|51|1857|209.7|5.50|
|Q8TAD1;Q9NY87|93.81|1|1|13|152|79.38|6|9|79.38|5|14|79.38|4|5|67.01|5|9|91.75|8|17|23.71|3|5|37.11|4|8|70.10|8|18|70.10|6|7|88.66|7|60|97|11.0|5.26|
|O43488|22.28|2|8|8|27|9.47|2|2|4.18|1|2|9.47|2|2|9.19|2|2|||||||9.19|2|2||||15.88|4|4|21.45|6|13|359|39.6|7.17|
|O43493|14.58|1|5|5|7||||3.96|1|1|2.92|1|1|||||||||||||||||||7.71|3|5|480|51.1|5.73|
|O43520|4.64|2|5|5|8||||||||||||||||||||||||||||4.64|5|8|1251|143.6|7.15|
|O43581|21.09|1|6|6|9|||||||8.68|3|3|||||||||||||||||||18.36|5|6|403|45.5|9.22|
|O43583|10.10|1|2|2|2||||||||||||||||||||||||||||10.10|2|2|198|22.1|5.30|
|O43592|28.48|1|20|20|46||||||||||||||||||||||||||||28.48|20|46|962|109.9|5.39|
|O43598|14.37|1|2|2|6||||||||||||||||||||||||||||14.37|2|6|174|19.1|5.05|
|O43615|14.60|1|6|6|8|7.52|3|3|5.09|2|2|0.00|||2.21|1|1||||||||||||||||4.65|2|2|452|51.3|8.32|
|O43617|53.89|1|12|12|49|10.00|2|2|10.00|2|2|5.56|1|1|22.78|5|5|||||||12.78|2|2||||28.89|5|6|53.89|12|31|180|20.3|4.96|
|O43633|9.91|1|3|3|6||||||||||3.15|1|1||||||||||||||||9.91|3|5|222|25.1|5.97|
|O43653|22.76|1|3|3|22|22.76|3|4|8.13|1|1|21.95|2|2|8.13|1|1||||||||||||||||22.76|3|14|123|12.9|5.29|
|O43657|13.06|1|3|3|3||||||||||||||||||||||||||||13.06|3|3|245|27.5|8.10|
|O43674|26.98|1|5|5|5||||||||||||||||||||||||||||26.98|5|5|189|21.7|9.63|
|O43676|35.71|1|4|4|5||||18.37|2|2|||||||||||||||||||||||28.57|3|3|98|11.4|9.20|
|O43678|30.30|1|2|2|5||||30.30|2|2|||||||||||||||||||||||30.30|2|3|99|10.9|9.57|
|O43681|39.08|1|9|9|16|||||||4.60|1|1||||||||||2.01|1|1||||6.61|1|1|32.47|8|13|348|38.8|4.91|
|O43684|16.16|1|3|3|5||||||||||||||||||||||||||||16.16|3|5|328|37.1|6.84|
|P11169|14.52|2|1|10|141|13.71|7|12|12.10|7|16|11.90|6|10|9.27|5|7|8.67|4|4|||||||||||||10.69|6|92|496|53.9|7.20|
|O43707|63.89|1|32|50|160|20.42|14|15|11.31|8|9|22.28|14|17|3.40|3|4|7.57|6|6||||5.16|4|4|3.95|3|3|6.26|5|7|60.26|46|95|911|104.8|5.44|
|O43708|21.76|1|3|3|11|9.26|1|2|9.26|1|2|||||||||||||||||||||||21.76|3|7|216|24.2|8.54|
|O43747|33.09|1|21|22|68||||||||||||||||||||||||||||33.09|22|68|822|91.3|6.80|
|O43759|9.44|1|2|2|4||||||||||||||||||||||||||||9.44|2|4|233|25.4|4.68|
|O43760|20.54|1|5|5|15|4.46|1|1|||||||||||||||||||||||||20.54|5|14|224|24.8|4.94|
|O43765|14.06|1|4|4|4|||||||||||||||||||||||||11.18|3|3|2.88|1|1|313|34.0|4.87|
|O43772|39.20|1|12|12|22|3.65|1|1|13.29|4|4|3.65|1|1|||||||||||||||||||39.20|11|16|301|32.9|9.41|
|O43776|32.66|1|17|17|34|1.46|1|1|||||||4.93|3|3|||||||||||||2.37|1|1|30.11|16|29|548|62.9|6.25|
|O43790|12.55|8|6|8|19|||||||||||||1.85|1|2|0.00|||1.44|1|1|1.85|2|3|1.85|1|2|12.14|7|11|486|53.5|5.66|
|O43795|16.37|1|14|15|23|0.00|||||||||||||||||||||||||||16.37|15|23|1136|131.9|9.38|
|O43805|36.97|1|5|5|12|20.17|2|2|20.17|2|2|10.92|1|1|20.17|2|2||||||||||||||||36.97|5|5|119|13.6|5.38|
|O43808|18.89|1|5|5|12||||||||||||||||||||||||||||18.89|5|12|307|34.5|10.08|
|O43813|44.86|1|14|14|30|3.01|1|1|5.51|2|2|||||||||||||||||||||||44.36|13|27|399|45.3|7.75|
|O43822|41.02|1|9|9|23|11.72|2|2|||||||17.19|3|3||||||||||||||||33.98|7|18|256|28.3|7.34|
|O43826|5.59|1|2|2|4||||||||||||||||||||||||||||5.59|2|4|429|46.3|8.62|
|O43837|50.91|1|17|17|46|20.00|5|7|11.43|3|5|14.81|4|5|||||||||||||||||||44.68|16|29|385|42.2|8.46|
|O43847|33.65|2|34|34|86|2.96|3|3|1.83|2|2|2.87|3|3|||||||||||||||||||31.83|32|78|1150|131.5|5.00|
|O43852|40.00|1|10|10|23|13.65|3|3|17.46|4|4||||2.86|1|1|14.60|4|4||||14.29|4|4|6.67|2|2|15.24|4|5||||315|37.1|4.64|
|O43854|45.00|1|18|18|31||||13.75|6|7|2.08|1|1||||2.08|1|1|||||||||||||38.54|15|22|480|53.7|7.28|
|O43865|30.38|2|6|16|43|||||||3.40|2|2||||||||||||||||1.89|1|1|30.38|14|40|530|58.9|6.89|
|O43920|57.55|1|6|6|7||||6.60|1|1|||||||||||||||||||||||57.55|6|6|106|12.5|9.14|
|O43924|28.67|1|3|3|4||||||||||||||||||||||||||||28.67|3|4|150|17.4|5.67|
|O60232|31.16|1|4|4|13||||4.02|1|2|4.02|1|1|4.02|1|1|||||||||||||12.56|2|2|31.16|4|7|199|21.5|5.24|
|O60256|30.89|1|8|9|15||||||||||||||||||||||||||||30.89|9|15|369|40.9|7.44|
|O60271|5.98|2|6|6|11||||0.61|1|1|||||||0.91|1|1||||||||||1.89|2|3|3.48|3|6|1321|146.1|5.15|
|O60282|22.78|1|7|18|47||||1.25|1|1|2.40|2|2|2.61|2|2|1.25|1|1||||3.03|2|2||||7.84|6|6|19.54|15|33|957|109.4|6.19|
|O60309|22.28|1|5|29|123|11.08|13|16|10.71|12|17|6.06|8|10||||1.90|3|3|||||||||||||17.26|25|77|1634|180.5|5.31|
|O60313|22.71|1|17|17|21|2.29|2|2|1.46|1|1|||||||||||||||||||||||21.25|16|18|960|111.6|7.87|
|O60343|7.16|1|7|7|17||||||||||2.31|2|2|||||||1.16|1|1|1.16|1|1|1.16|1|1|7.16|7|12|1298|146.5|7.01|
|O60423|12.31|3|10|12|24||||||||||||||||||||||||||||12.31|12|24|1300|146.7|7.90|
|O60493|46.30|1|7|8|21||||||||||6.79|1|1||||||||||6.79|1|1|12.35|2|2|46.30|8|17|162|18.8|8.66|
|O60494|2.24|1|5|6|7||||||||||||||||||||||||||||2.24|6|7|3623|398.5|5.35|
|O60502|5.57|1|4|4|4||||||||||||||||||||||||||||5.57|4|4|916|102.8|4.91|
|O60504|8.79|1|3|3|3||||||||||||||||||||||||||||8.79|3|3|671|75.3|9.45|
|O60506|8.51|1|3|4|6|||||||||||||||||||||||||1.28|1|1|7.22|3|5|623|69.6|8.59|
|O60547|45.43|1|14|14|26|||||||3.23|1|1||||||||||||||||3.23|1|1|45.43|14|24|372|41.9|7.31|
|O60568|7.32|1|3|3|3||||||||||||||||||||||||||||7.32|3|3|738|84.7|6.05|
|O60610|7.63|1|10|10|11||||0.00|||||||||||||||||||||||||7.63|10|11|1272|141.3|5.41|
|O60613|25.31|1|3|3|5|15.43|2|2|8.02|1|1|||||||||||||||||||||||17.28|2|2|162|17.8|5.03|
|O60645|7.54|1|5|5|5||||||||||||||||||||||||||||7.54|5|5|756|86.8|6.11|
|O60664|51.84|1|16|16|39|25.58|7|7|22.35|7|7|5.30|2|2|9.22|3|3|||||||2.76|1|1||||8.53|2|2|40.55|10|17|434|47.0|5.44|
|O60676|19.72|1|2|2|3||||||||||||||||||||||||||||19.72|2|3|142|16.3|8.81|
|O60684|33.40|1|3|12|30|1.87|1|1|||||||||||||||||||||||||33.21|11|29|536|60.0|4.98|
|O60701|60.73|1|23|23|86|23.48|9|10||||12.75|5|5|10.12|4|4|||||||13.36|5|5|2.02|1|1|8.10|3|3|58.10|21|58|494|55.0|7.12|
|O60733|32.51|2|18|18|29||||2.11|1|1|||||||||||||||||||||||32.51|18|28|806|89.8|7.27|
|O60749|20.04|1|8|10|21||||||||||2.12|1|1|||||||2.12|1|1||||4.82|2|3|19.46|9|16|519|58.4|5.12|
|O60762|15.38|1|3|3|4||||||||||||||||||||||||||||15.38|3|4|260|29.6|9.57|
|O60763|41.27|1|32|32|128|4.16|3|3|7.28|5|6|2.18|2|2|5.93|4|5|8.84|6|7||||7.38|5|5|8.52|6|7|13.41|8|9|39.71|27|84|962|107.8|4.91|
|O60784|10.57|1|4|4|13|2.64|1|1|2.64|1|1|||||||||||||||||||2.64|1|1|7.93|3|10|492|53.8|4.70|
|O60825|16.04|4|9|9|18|||||||||||||2.57|1|1||||||||||2.57|1|1|15.05|8|16|505|58.4|8.38|
|O60831|20.22|1|3|3|7|||||||6.18|1|1|||||||||||||||||||20.22|3|6|178|19.2|9.19|
|O60841|16.07|1|17|17|18|1.31|1|1|||||||0.74|1|1||||||||||||||||14.02|15|16|1220|138.7|5.49|
|O60884|68.20|1|21|21|66|28.40|5|7|30.83|6|8|8.74|3|3|7.04|3|3|12.62|5|5|||||||12.62|5|5||||50.49|18|39|412|45.7|6.48|
|O60888|17.88|1|2|2|2||||||||||||||||||||||||||||17.88|2|2|179|19.1|5.50|
|O60906|11.58|1|4|4|5||||||||||||||||||||||||||||11.58|4|5|423|47.6|6.98|
|P20020|14.47|1|1|19|117|6.76|7|7|9.38|11|13|6.76|7|10|2.23|2|2|0.87|1|1||||||||||0.87|1|1|14.39|18|83|1258|138.7|6.04|
|O75031|6.89|1|2|2|2||||||||||||||||||||||||||||6.89|2|2|334|37.6|5.53|
|O75071|21.21|1|8|8|20|2.63|1|1|8.89|3|3|||||||2.02|1|1|||||||||||||21.21|8|15|495|55.0|6.32|
|O75083|64.03|1|29|29|170|5.78|3|3|4.79|3|3|27.89|10|11|24.92|11|11|8.42|4|4||||13.20|5|5|22.77|10|11|46.04|18|26|54.13|21|96|606|66.2|6.65|
|O75110|4.68|1|5|5|6||||||||||||||||||||||||||||4.68|5|6|1047|118.5|7.77|
|O75116|9.01|2|6|10|10||||||||||||||||||||||||||||9.01|10|10|1388|160.8|6.02|
|O75131|40.04|8|18|18|31|||||||5.59|3|3||||||||||||||||2.42|1|1|36.69|15|27|537|60.1|5.85|
|O75155|5.99|1|3|6|7||||||||||||||||||||||||||||5.99|6|7|1236|135.2|5.68|
|O75165|14.76|1|28|28|49||||||||||||||||||||||||||||14.76|28|49|2243|254.3|6.74|
|O75170|12.11|1|8|8|12||||||||||||||||||||||||||||12.11|8|12|966|104.9|4.87|
|O75190|24.23|2|5|7|22|7.36|2|2|7.36|2|2|3.99|1|1|10.43|3|3|7.36|2|2|||||||||||||24.23|7|12|326|36.1|9.16|
|O75223|23.94|1|4|4|7|5.85|1|1||||12.23|2|2|||||||||||||5.32|1|1||||18.09|3|3|188|21.0|5.14|
|O75251|21.60|1|5|5|8||||4.23|1|1|||||||||||||||||||||||21.60|5|7|213|23.5|9.99|
|O75264|18.46|1|2|2|2||||||||||||||||||||||||||||18.46|2|2|130|15.0|5.29|
|O75306|27.65|1|11|11|15||||||||||||||||||||||||||||27.65|11|15|463|52.5|7.55|
|Q58FF6|10.10|1|1|8|113|8.71|4|4|10.10|6|6|10.10|6|8|9.90|5|6|7.92|6|9||||9.90|6|6|10.10|7|9|7.92|6|10|10.10|8|55|505|58.2|4.73|
|O75340|52.36|1|9|9|16|6.81|1|1|||||||||||||||||||||||||52.36|9|15|191|21.9|5.40|
|O75347|38.89|1|3|3|4|||||||||||||||||||12.04|1|1|||||||38.89|3|3|108|12.8|5.29|
|O75348|44.92|2|6|6|15|29.66|3|3|16.95|2|2|||||||7.63|1|1|||||||||||||44.92|6|9|118|13.7|8.79|
|O75352|13.77|1|3|3|7||||||||||||||||||||||||||||13.77|3|7|247|26.6|8.94|
|O75355|7.94|1|4|4|5|||||||4.54|2|2|||||||||||||||||||6.43|3|3|529|59.1|6.42|
|O75356|10.98|1|3|3|3||||||||||||||||||||||||||||10.98|3|3|428|47.5|6.33|
|O75367|7.80|2|2|2|4|2.42|1|1|||||||||||||||||||||||||7.80|2|3|372|39.6|9.79|
|O75369|54.57|3|103|108|428|3.00|5|5||||22.25|43|49|5.57|12|13|||||||9.68|20|23|9.19|18|21|20.87|37|43|48.54|92|274|2602|278.0|5.73|
|O75379|37.59|1|4|4|7|9.22|1|1|||||||||||||||||||||||||28.37|3|6|141|16.4|7.34|
|O75380|49.19|1|5|5|6|0.00|||0.00|||0.00||||||0.00||||||0.00|||0.00|||0.00|||49.19|5|6|124|13.7|8.28|
|O75381|7.96|1|2|2|4||||||||||||||||||||||||||||7.96|2|4|377|41.2|4.94|
|O75390|53.43|1|23|23|80|35.84|10|10|33.69|14|20|21.03|9|11|8.15|3|4|3.22|2|2||||||||||3.86|1|1|43.35|18|32|466|51.7|8.32|
|O75396|38.14|1|8|8|34|13.95|2|2|13.95|2|2|20.47|3|3|8.37|2|2|||||||4.65|1|1||||3.72|1|1|38.14|7|23|215|24.6|6.92|
|O75431|30.42|1|5|5|11||||8.75|2|2|||||||||||||||||||||||30.42|5|9|263|29.7|6.29|
|O75436|23.24|1|5|6|10|||||||||||||||||||||||||3.98|1|2|19.27|5|8|327|38.1|6.57|
|O75439|20.04|1|6|7|8||||5.93|1|1|||||||||||||||||||||||14.11|6|7|489|54.3|6.83|
|O75477|54.34|1|14|18|66|16.47|4|5|18.79|6|7|7.80|2|2|7.51|2|2|16.76|5|5|||||||2.02|1|1|4.05|1|1|54.05|17|43|346|38.9|7.87|
|O75487|36.87|1|14|15|23|3.24|1|1|2.70|1|1|5.40|2|2|||||||||||||||||||28.42|12|19|556|62.4|6.68|
|O75489|50.00|1|11|11|15|5.30|1|2|10.23|2|2|||||||||||||||||||||||45.08|9|11|264|30.2|7.50|
|O75521|40.61|1|15|15|38|17.51|5|5|24.37|8|10|6.60|2|2|||||||||||||||||||40.10|14|21|394|43.6|9.00|
|O75531|49.44|1|4|4|5|7.87|1|1||||||||||||||||||||||41.57|2|3|26.97|1|1|89|10.1|6.09|
|O75534|21.05|1|14|14|17||||||||||||||||||||||3.51|3|3|9.52|6|7|10.65|7|7|798|88.8|6.25|
|O75556|58.95|1|4|4|12|58.95|4|5|18.95|1|1||||21.05|2|2|||||||12.63|1|1|||||||31.58|2|3|95|10.9|5.78|
|O75569|38.66|1|11|11|38||||7.67|2|2|4.79|1|1|10.86|2|2|4.79|1|1||||4.79|1|1|4.79|1|1|4.79|1|1|38.66|11|29|313|34.4|8.41|
|O75592|2.50|1|10|10|13||||||||||||||||||||||0.00||||||2.50|10|13|4640|509.8|7.03|
|O75602|67.39|1|26|26|181|21.22|7|7|26.33|10|11|14.93|7|7|9.43|4|4|7.07|3|3||||||||||4.72|2|2|67.19|25|147|509|55.4|6.83|
|O75604|24.79|1|9|9|11||||||||||||||||||||||||||||24.79|9|11|605|68.0|8.97|
|O75608|60.43|1|8|8|34|16.52|3|3|16.52|3|3|17.39|2|2|5.65|1|1|||||||||||||12.61|2|3|49.13|7|22|230|24.7|6.77|
|O75643|1.03|1|2|2|3||||||||||||||||||||||||||||1.03|2|3|2136|244.4|6.06|
|O75663|33.09|1|7|7|7||||||||||||||||||||||||||||33.09|7|7|272|31.4|5.91|
|O75665|3.66|1|3|3|7|1.28|1|1|1.28|1|1|||||||||||||||||||||||3.66|3|5|1012|116.6|6.10|
|O75688|17.54|1|3|6|8||||2.09|1|1|||||||||||||||||||||||17.54|6|7|479|52.6|5.05|
|O75694|48.53|1|52|52|270|16.68|16|20|16.46|16|20|9.49|10|10|2.44|2|2|0.50|1|1|||||||||||||47.88|50|217|1391|155.1|6.16|
|O75695|5.14|1|2|2|3||||||||||||||||||||||||||||5.14|2|3|350|39.6|5.12|
|O75718|4.74|1|2|2|2||||0.00||||||0.00|||||||||||||||||||4.74|2|2|401|46.5|5.73|
|O75746|22.71|1|9|13|22|1.18|1|1|||||||||||||||||||||||||22.71|13|21|678|74.7|8.38|
|O75787|30.57|1|9|9|22||||||||||4.00|1|1|4.00|1|1||||8.57|3|3|6.29|2|2|8.57|3|3|27.14|7|12|350|39.0|6.10|
|O75800|25.68|1|9|9|10||||||||||||||||||||||||||||25.68|9|10|440|50.3|6.19|
|O75821|16.25|1|4|4|6||||||||||||||||||||||4.06|1|1||||12.19|3|5|320|35.6|6.13|
|O75822|7.75|1|2|2|2|||||||||||||||||||||||||4.65|1|1|3.10|1|1|258|29.0|4.83|
|A6NHL2|11.88|1|1|8|99|2.47|2|4|4.48|3|6|4.48|3|5|6.73|5|10|8.07|5|9|2.47|1|1|2.69|3|4|2.47|1|3|6.05|3|7|10.54|6|50|446|49.9|6.05|
|O75828|29.96|1|4|7|35|10.47|2|2|10.47|2|3|13.36|3|4|10.47|2|3|||||||10.47|2|3|2.89|1|1|10.47|2|2|29.96|7|17|277|30.8|6.18|
|O75832|35.40|1|6|6|12|3.54|1|1|||||||4.42|1|1|||||||4.42|1|1||||4.42|1|1|30.97|5|8|226|24.4|6.10|
|O75844|6.32|1|3|3|6|||||||2.32|1|1|||||||||||||||||||6.32|3|5|475|54.8|7.49|
|O75874|58.70|1|22|23|132|25.85|9|9|23.43|8|8|42.75|15|17|25.36|9|10|12.56|4|5||||11.59|5|6|13.53|4|5|30.92|11|14|50.97|20|58|414|46.6|7.01|
|O75882|3.64|1|4|4|5|0.70|1|1|1.12|1|1|0.91|1|1|||||||||||||||||||2.03|2|2|1429|158.4|7.31|
|O75884|37.10|1|5|5|7||||||||||||||||||||||||||||37.10|5|7|186|21.0|6.20|
|O75886|7.05|1|2|3|5|||||||||||||||||||1.90|1|1|2.48|1|1|4.38|2|2|2.67|1|1|525|58.1|5.07|
|O75891|12.20|2|7|7|8||||0.00|||0.00|||0.00|||||||0.00||||||0.00|||12.20|7|8|902|98.8|5.94|
|O75915|25.00|1|5|5|11|10.64|1|1|||||||||||||||||||||||||24.47|4|10|188|21.6|9.77|
|O75935|50.54|1|8|8|11||||10.75|2|2||||5.91|1|1||||||||||||||||45.70|7|8|186|21.1|5.47|
|P62136|58.18|1|1|16|97|35.15|9|12|12.42|3|3|22.12|6|7|10.91|3|3|4.24|1|1||||4.24|1|1|4.24|1|1|12.42|3|4|51.21|15|65|330|37.5|6.33|
|O75947|78.88|1|15|15|50|45.96|5|6|51.55|6|9|16.15|2|2|6.21|1|1|11.18|2|2|||||||||||||78.88|14|30|161|18.5|5.30|
|O75951|41.89|1|3|3|17|41.89|3|5|6.08|1|2|17.57|1|1|6.08|1|1||||||||||||||||41.89|3|8|148|16.9|6.14|
|O75952|59.03|3|32|32|435|20.08|8|13|20.08|8|13|14.81|6|7|32.05|16|25|39.15|23|35|5.88|2|2|23.73|11|16|41.99|20|34|31.44|14|20|54.56|24|270|493|52.7|4.55|
|O75955|61.12|1|22|22|50|17.33|5|5|5.15|2|2|19.91|8|8|1.87|1|1|3.28|1|1||||3.04|1|1||||13.11|5|5|57.85|20|27|427|47.3|7.49|
|O75964|46.60|1|4|4|7||||12.62|1|1|||||||||||||||||||||||46.60|4|6|103|11.4|9.64|
|O75969|79.84|1|89|89|1780|48.53|36|43|40.45|34|42|36.46|30|32|58.03|49|77|59.55|51|67|3.05|3|3|47.01|37|54|49.24|46|66|59.32|57|87|75.50|68|1309|853|94.7|6.18|
|O75976|34.86|1|40|40|112|5.65|8|9|13.99|15|17|9.35|11|13|||||||||||||||||||31.45|37|73|1380|152.8|6.05|
|O76003|40.90|1|12|12|30|7.16|2|2|10.75|3|3|3.58|1|1|15.52|4|4|||||||8.36|3|3||||23.88|6|6|27.76|8|11|335|37.4|5.39|
|O76013|5.14|10|2|3|16||||1.50|1|1|1.50|1|1||||1.50|1|1||||1.50|1|1|1.50|1|1|1.50|1|1|5.14|3|10|467|52.2|4.94|
|O76024|11.69|1|8|8|13||||||||||0.00|||||||||||||||||11.69|8|13|890|100.2|8.05|
|O76054|31.27|1|9|9|31||||0.00|||||||||2.73|1|1||||2.73|1|1||||5.71|2|2|31.27|9|27|403|46.1|7.84|
|O76062|11.24|1|4|4|5||||||||||||||||||||||||||||11.24|4|5|418|46.4|8.87|
|O76071|28.32|1|6|6|9||||||||||||||||||||||||||||28.32|6|9|339|37.8|4.97|
|O76076|22.80|1|4|4|7||||||||||||||||||||||||||||22.80|4|7|250|26.8|7.88|
|O76094|10.28|1|5|5|13|||||||||||||||||||||||||1.49|1|1|10.28|5|12|671|74.6|9.26|
|O94760|10.18|1|2|3|6|3.86|1|1||||3.86|1|1|||||||||||||||||||10.18|3|4|285|31.1|5.81|
|O94788|28.19|1|7|9|26|9.85|4|5|2.70|1|1|9.27|4|4|2.12|1|1|||||||2.12|1|1|2.12|1|1|3.86|2|2|20.46|6|11|518|56.7|6.05|
|O94813|3.92|2|5|5|6||||||||||||||||||||||||||||3.92|5|6|1529|169.8|7.06|
|O94822|3.79|1|5|5|7||||||||||||||||||||||||||||3.79|5|7|1766|200.4|6.25|
|O94829|17.13|1|12|12|19||||||||||||||||||||||||||||17.13|12|19|963|108.1|5.30|
|O94832|26.84|1|22|22|42||||||||||||||||||||||||||||26.84|22|42|1006|116.1|9.39|
|O94874|7.56|1|5|5|7||||||||||||||||||||||||||||7.56|5|7|794|89.5|6.79|
|O94886|5.33|2|5|5|11||||0.62|1|1|||||||||||||||||||||||4.71|4|10|807|92.1|7.27|
|O94903|22.91|1|5|5|7|||||||||||||||||||||||||4.73|1|1|18.18|4|6|275|30.3|7.50|
|O94905|65.49|1|19|23|99|28.32|8|10|31.27|9|11|25.66|7|9|7.08|2|2|18.58|6|7||||3.54|1|1|7.67|3|3|7.37|2|3|65.49|23|53|339|37.8|5.62|
|O94919|12.20|1|4|4|6|||||||2.40|1|1|||||||||||||||||||9.80|3|5|500|55.0|5.71|
|O94973|43.13|1|19|31|67|2.98|2|2|2.98|2|2|||||||||||||||||||||||42.07|30|63|939|103.9|6.96|
|O94979|8.52|1|11|11|13||||||||||||||||0.90|1|1|||||||1.23|2|2|7.05|8|10|1220|132.9|6.89|
|O94985|6.73|1|6|7|8|0.61|1|1|||||||||||||||||||||||||6.73|7|7|981|109.7|4.91|
|O95025|19.82|2|13|14|24|0.77|1|1|0.77|1|1|0.77|1|1||||||||||||0.77|1|1|0.77|1|1|19.82|14|19|777|89.6|7.80|
|O95070|9.22|1|2|2|2||||||||||||||||||||||||||||9.22|2|2|293|32.0|8.95|
|O95084|14.62|1|3|3|3||||||||||||||||||||||||||||14.62|3|3|383|43.0|9.42|
|O95139|19.53|1|2|2|4|14.06|1|1|||||||||||||||||||||||||19.53|2|3|128|15.5|9.63|
|O95159|11.29|1|3|3|5||||||||||||||||||||||||||||11.29|3|5|310|34.1|8.07|
|O95163|8.78|1|9|9|10||||||||||||||||||||||||||||8.78|9|10|1332|150.2|5.94|
|O95168|44.19|1|5|5|5||||||||||||||||||||||||||||44.19|5|5|129|15.2|9.85|
|O95169|25.81|1|4|4|5||||||||||||||||||||||||||||25.81|4|5|186|21.8|6.80|
|O95182|33.63|1|3|3|4||||||||||||||||||||||||||||33.63|3|4|113|12.5|10.18|
|O95197|1.84|2|2|2|14|1.07|1|1|1.07|1|1|1.07|1|2|||||||||||||||||||1.84|2|10|1032|112.5|4.96|
|O95202|35.45|1|21|21|40|5.95|3|3|11.64|6|6|1.62|1|1|1.62|1|1||||||||||||||||33.29|17|29|739|83.3|6.70|
|O95219|5.11|1|2|2|2||||||||||||||||||||||||||||5.11|2|2|450|51.9|5.99|
|O95249|9.60|1|2|2|2||||||||||||||||||||||||||||9.60|2|2|250|28.6|9.42|
|O95258|15.08|1|4|4|4||||||||||||||||||||||||||||15.08|4|4|325|36.2|9.64|
|O95292|30.86|1|5|6|10|13.17|2|3|4.94|1|1|4.94|1|1||||0.00||||||||||||||22.63|5|5|243|27.2|7.30|
|O95295|25.74|1|2|2|2||||||||||||||||||||||||||||25.74|2|2|136|14.9|9.31|
|O95298|39.50|2|5|5|5||||||||||||||||||||||||||||39.50|5|5|119|14.2|8.98|
|O95299|26.20|1|9|9|12||||||||||||||||||||||||||||26.20|9|12|355|40.7|8.48|
|O95336|54.26|1|11|11|19||||||||||3.49|1|1|||||||4.26|1|1||||7.36|2|2|50.78|10|15|258|27.5|6.05|
|O95352|3.98|1|2|2|8||||||||||||||||||||||||||||3.98|2|8|703|77.9|6.24|
|O95372|29.00|1|5|5|13|||||||||||||||||||3.46|1|1||||3.46|1|1|29.00|5|11|231|24.7|7.23|
|O95373|21.68|1|18|18|46||||||||||0.87|1|1||||||||||||||||20.81|17|45|1038|119.4|4.82|
|O95376|7.30|1|2|2|4||||||||||||||||||||||||||||7.30|2|4|493|57.8|5.63|
|O95394|49.63|1|23|23|79|8.67|3|3|1.85|1|1||||8.86|5|6|||||||2.40|1|1|4.06|2|2|9.41|6|7|49.63|22|59|542|59.8|6.25|
|O95396|18.26|1|5|5|7||||0.00|||||||||||||||||||||||||18.26|5|7|460|49.6|6.21|
|O95433|36.09|1|10|10|16|0.00|||||||||||||||||||||||||||36.09|10|16|338|38.3|5.53|
|O95456|13.19|1|3|3|5||||3.47|1|1|||||||||||||||||||||||13.19|3|4|288|32.8|7.17|
|O95473|10.26|1|3|3|15|6.41|1|2|6.41|1|2|6.41|1|2|||||||||||||||||||10.26|3|9|234|25.8|7.75|
|O95479|4.68|1|3|3|5||||||||||||||||||||||||||||4.68|3|5|791|88.8|7.30|
|O95497|10.72|2|4|4|5|||||||10.72|4|5||||||||||||||||||||||513|57.0|5.55|
|O95498|7.31|1|2|2|2|||||||7.31|2|2||||||||||||||||||||||520|58.5|6.52|
|O95563|65.35|1|8|8|25|7.87|1|1|15.75|2|2|||||||||||||||||||||||65.35|7|22|127|14.3|10.43|
|O95573|35.28|1|19|21|64|3.33|2|2|3.33|2|2|4.17|3|3|||||||||||||||||||35.28|21|57|720|80.4|8.38|
|O95671|16.26|1|7|7|17||||||||||2.74|1|1|||||||1.45|1|1||||4.19|2|2|16.26|7|13|621|68.8|6.07|
|O95685|27.09|1|6|6|7||||||||||||||||||||||||||||27.09|6|7|299|32.5|8.07|
|O95716|80.37|8|9|15|85|26.48|5|6|19.63|4|4|31.05|6|6|13.70|3|3|||||||8.68|2|2|3.65|1|1|3.65|1|1|80.37|15|62|219|24.3|4.93|
|O95721|8.53|1|2|2|2|0.00|||4.26|1|1|||||||4.26|1|1|||||||||||||0.00|||258|29.0|5.81|
|O95747|22.58|1|5|9|21|||||||2.85|1|1|5.12|2|2|||||||2.85|1|1|2.85|1|1|5.69|2|2|19.73|8|14|527|58.0|6.43|
|O95749|33.00|1|9|9|11|4.00|1|1|5.00|2|2|||||||||||||||||||||||26.67|7|8|300|34.8|6.14|
|O95757|71.16|1|45|51|211|27.18|16|17|22.29|14|16|40.52|25|32|7.75|6|6|14.78|12|13||||1.67|1|1|8.58|6|6|5.01|4|5|60.67|39|115|839|94.5|5.88|
|O95758|3.44|2|2|2|5||||||||||||||||||||||||||||3.44|2|5|552|59.7|9.04|
|O95777|23.96|1|2|2|2|||||||7.29|1|1|||||||||||||||||||16.67|1|1|96|10.4|4.48|
|O95782|37.15|1|16|28|65|2.87|2|2|2.87|2|2|||||||||||||||||||||||36.13|27|61|977|107.5|7.03|
|O95786|8.22|1|6|6|9||||||||||||||||||||||||||||8.22|6|9|925|106.5|6.40|
|Q16720|12.95|2|1|16|85|5.66|6|7|8.28|9|11|6.64|6|8|2.30|2|2|0.90|1|1||||||||||0.90|1|1|10.49|13|55|1220|134.1|5.62|
|O95816|70.14|1|15|15|47|21.33|4|4|21.80|5|5||||8.06|2|2|5.21|1|1||||4.27|1|1|18.01|4|4|37.44|6|6|63.03|11|24|211|23.8|6.70|
|O95822|12.58|1|4|5|7||||2.64|1|1|||||||1.42|1|1|||||||||||||9.94|4|5|493|55.0|8.95|
|CON_002936657|38.83|8|1|27|84|3.37|2|2|9.75|7|8|11.17|9|11|3.01|3|4|9.75|7|9|1.42|1|1|26.06|14|15|16.67|15|20|17.38|12|14||||564|60.0|8.00|
|O95825|14.61|1|3|3|3||||||||||||||||||||||||||||14.61|3|3|349|38.7|5.78|
|O95831|40.78|1|18|18|44|18.43|9|9|17.62|9|10|15.17|7|8||||1.79|1|1|||||||||||||26.75|12|16|613|66.9|8.95|
|O95833|55.08|1|11|11|29|6.78|1|1|||||||19.07|4|4|6.78|1|1|||||||8.47|2|2|19.07|4|4|51.69|9|17|236|26.6|6.43|
|O95834|20.34|1|11|11|31|||||||8.47|5|5||||||||||||||||4.47|2|3|14.02|8|23|649|70.6|6.32|
|O95861|55.52|1|13|13|28|11.36|3|3|6.17|2|2|3.57|1|1|3.57|1|1||||||||||||||||55.52|13|21|308|33.4|5.69|
|O95864|11.94|1|5|5|8||||||||||||||||||||||||||||11.94|5|8|444|52.2|8.82|
|O95865|31.23|1|7|8|15||||||||||8.07|2|2|||||||8.07|2|2|4.91|1|1|8.07|2|2|30.88|7|8|285|29.6|6.01|
|O95870|30.29|1|15|15|29||||7.35|4|4|2.87|2|2|||||||||||||||||||30.29|14|23|558|63.2|8.13|
|O95925|13.53|1|2|2|4||||||||||||||||||||||||||||13.53|2|4|133|15.3|8.13|
|O95989|29.07|1|4|4|5||||||||||||||||||||||||||||29.07|4|5|172|19.5|6.34|
|O95994|49.14|2|9|9|70|46.29|6|9|42.86|6|10|46.29|6|10|20.57|3|4|10.29|2|3||||16.57|3|4|10.29|2|2|29.14|4|7|46.86|8|21|175|20.0|9.00|
|O95995|70.92|1|49|49|139||||||||||39.54|23|28|52.72|34|44||||2.09|1|1|1.88|1|1|10.04|4|4|59.83|31|59|478|56.3|7.94|
|O96000|43.02|1|7|7|16|19.19|3|3|19.19|3|3|||||||6.40|1|1|||||||||||||43.02|7|9|172|20.8|8.48|
|O96005|13.60|1|8|8|16||||3.29|2|2|||||||||||||||||||||||13.60|8|14|669|76.0|6.30|
|O96007|13.83|1|2|2|3||||||||||||||||||||||||||||13.83|2|3|188|20.9|5.44|
|O96008|33.52|1|7|7|18||||6.93|2|2|3.05|1|1|||||||||||||||||||33.52|7|15|361|37.9|7.25|
|O96011|69.88|1|17|17|42||||12.36|3|4|4.63|1|1|||||||||||||||||||66.02|15|37|259|28.4|9.85|
|O96015|33.33|1|5|5|18|12.38|1|1|12.38|1|1|12.38|1|1|20.95|2|3|13.33|2|3|12.38|1|1|||||||12.38|1|1|33.33|5|7|105|12.0|5.49|
|O96019|16.78|2|4|4|4||||||||||||||||||||||||||||16.78|4|4|429|47.4|5.60|

| | | | | | | | | | | | | | | | | | | | | | | | | | | | | | | | | | | | | | | |
|---|---|---|---|---|---|---|---|---|---|---|---|---|---|---|---|---|---|---|---|---|---|---|---|---|---|---|---|---|---|---|---|---|---|---|---|---|---|---|
|P00156|5.00|1|2|2|3| | | | | | | | | | | | | | | | | | | | | | | | | | | |5.00|2|3|380|42.7|8.05|
|P00167|42.54|1|4|4|16| | | | | | | | | | | | | | | | | | |15.67|1|1|17.16|2|2|6.72|1|1|42.54|4|12|134|15.3|4.96|
|Q58FG1|10.77|1|1|5|79|4.78|2|4|10.77|5|8|7.42|3|5|7.42|3|4|4.78|2|3| | | |7.42|3|7|5.50|3|7|8.13|4|7|7.42|3|34|418|47.7|5.19|
|P00338|87.05|3|25|30|223|35.24|11|16|37.65|12|14|32.53|10|11|41.57|12|15|14.76|5|5| | | |14.76|5|5|6.63|2|2|15.36|6|6|85.84|28|149|332|36.7|8.27|
|P00352|47.70|1|17|20|42|12.97|5|5|11.38|4|4|19.96|8|8|2.20|1|1| | | | | | |2.20|1|1| | | |3.99|2|2|39.32|16|21|501|54.8|6.73|
|P00367|31.18|2|17|17|38|14.16|7|7|3.41|2|2|10.75|5|5| | | | | | | | | | | | | | | | | | |26.70|14|24|558|61.4|7.80|
|P00387|51.50|1|15|15|36|13.95|3|5|13.95|3|4|20.27|4|5|3.65|1|1| | | | | | | | | | | | |4.65|1|1|50.83|13|20|301|34.2|7.59|
|P00390|46.93|1|16|16|38|9.00|3|3|4.02|1|1|23.56|8|8| | | | | | | | | | | | | | | |1.34|1|1|45.98|14|25|522|56.2|8.50|
|P00395|8.77|1|2|2|7|6.04|1|2| | | | | | | | | | | | | | | | | | | | | | | | |8.77|2|5|513|57.0|6.70|
|P00403|30.84|1|7|7|45|20.70|5|8|27.75|6|9|16.30|4|7|3.08|1|1| | | | | | | | | | | | | | | |30.84|6|20|227|25.5|4.82|
|P00441|70.13|2|8|8|38|56.49|5|10|39.61|5|8|56.49|5|7|23.38|2|2|18.18|2|2| | | | | | | | | | | | |41.56|4|9|154|15.9|6.13|
|P00450|11.46|3|9|9|14|3.66|3|3|0.94|1|1|10.52|8|10| | | | | | | | | | | | | | | | | | |0.00| | |1065|122.1|5.72|
|P00491|40.83|1|8|8|13|5.88|1|1|5.88|1|1|12.11|2|2| | | | | | | | | | | | | | | | | | |40.83|8|9|289|32.1|6.95|
|P00492|69.72|1|11|11|46|49.54|9|9|44.95|8|10|32.11|6|6|11.01|2|2| | | | | | | | | |5.96|1|1| | | |59.17|9|18|218|24.6|6.68|
|P00505|53.02|1|22|22|60|19.07|6|6|32.79|11|13|23.95|9|10| | | | | | | | | |0.00| | | | | |0.00| | |44.19|19|31|430|47.5|9.01|
|P00558|87.77|1|31|37|250|38.61|13|15|31.89|11|13|49.64|18|20|42.45|14|16|19.42|7|7| | | |32.85|13|16|38.61|15|20|64.27|25|31|76.50|30|112|417|44.6|8.10|
|P00568|79.38|1|20|20|153|60.31|10|16|73.71|13|22|58.25|10|14|33.51|6|8|30.41|6|7| | | |29.90|5|6|19.59|5|9|29.38|6|9|63.92|15|62|194|21.6|8.63|
|P00738|29.56|3|11|11|16|11.82|4|4| | | |27.09|10|12| | | | | | | | | | | | | | | | | | | | | |406|45.2|6.58|
|P00747|5.80|1|4|4|5|2.35|2|2| | | | | | |1.36|1|1| | | | | | | | | | | | | | | |2.10|1|2|810|90.5|7.24|
|P00750|4.98|1|2|2|2| | | | | | | | | | | | | | | | | | | | | | | | | | | |4.98|2|2|562|62.9|7.80|
|P00846|7.08|1|2|2|9|4.42|1|1| | | |4.42|1|1| | | | | | | | | | | | | | | | | | |7.08|2|7|226|24.8|10.10|
|P00918|50.00|1|11|11|38|26.15|5|7|28.85|6|8|30.00|6|6| | | | | | | | | | | | | | | | | | |46.15|10|17|260|29.2|7.40|
|P00966|22.09|1|10|10|13| | | | | | | | | |1.70|1|1| | | | | | | | | | | | | | | |20.39|9|12|412|46.5|8.02|
|P01009|30.38|2|13|13|45|24.40|10|12|17.22|7|7|23.21|9|11|13.16|6|6|2.39|1|1| | | | | | | | | |4.31|2|2|17.22|6|6|418|46.7|5.59|
|P01011|29.79|1|9|9|15|14.42|5|6|2.36|1|1|10.17|3|3|6.86|2|2| | | | | | | | | | | | | | | |6.38|2|3|423|47.6|5.52|
|P01033|57.49|1|8|8|25|40.58|5|7|5.80|1|1|45.41|6|6|5.80|1|1|9.18|1|1| | | | | | | | | | | | |34.78|5|9|207|23.2|8.10|
|P01034|33.56|1|6|6|18|25.34|2|2|19.18|3|4|21.23|2|2| | | |7.53|1|1| | | | | | | | | | | | |33.56|6|9|146|15.8|8.75|
|P01036|57.45|3|7|7|14|46.81|5|5| | | |36.88|4|5| | | | | | | | | | | | | | | | | | |25.53|3|4|141|16.2|5.02|
|P01040|54.08|1|4|4|9| | | | | | |18.37|1|1| | | | | | | | | |41.84|3|5|30.61|2|2| | | |12.24|1|1|98|11.0|5.50|
|P35241|16.64|1|1|14|75|6.17|5|6|9.26|6|8|12.86|7|7|4.80|3|3|3.60|2|3| | | | | | | | | |2.74|2|2|12.18|10|46|583|68.5|6.37|
|P01111|25.40|3|3|3|7| | | | | | | | | |5.82|1|1| | | | | | | | | |5.82|1|1|5.82|1|1|25.40|3|4|189|21.2|5.17|
|P01133|3.98|1|3|3|4| | | | | | | | | | | | | | | | | | | | | | | | | | | |3.98|3|4|1207|133.9|5.85|
|P01833|15.31|1|9|9|14| | | |1.05|1|1|9.82|6|8| | | | | | |2.09|1|1| | | | | | | | | |5.50|3|4|764|83.2|5.74|
|CON_001360561|17.62|16|1|12|79| | | |8.47|4|4|13.96|7|8|3.20|2|2|6.41|4|4| | | |4.35|2|2|8.92|6|8|4.81|3|4|15.79|10|43|437|47.7|4.86|
|P01857|33.03|2|6|8|20|13.64|3|4| | | |28.18|7|8| | | | | | |2.12|1|1| | | |6.36|2|2| | | |14.24|4|5|330|36.1|8.19|
|P01859|18.40|1|3|5|14|9.51|2|2|5.21|1|1|18.40|5|6| | | | | | |2.15|1|1| | | |2.15|1|1| | | |9.51|3|3|326|35.9|7.59|
|P01871|12.36|2|4|4|7| | | | | | |6.40|2|2| | | | | | | | | | | | | | | | | | |9.49|3|5|453|49.4|6.77|
|P01876|45.89|1|7|12|26|2.83|1|1|9.63|3|3|45.89|12|14| | | |7.93|3|3| | | | | | | | | | | | |13.31|5|5|353|37.6|6.51|
|P01892|29.59|14|2|8|9|8.22|2|2| | | |12.33|3|3| | | | | | | | | | | | | | | | | | |15.62|4|4|365|40.9|6.99|
|P01903|11.42|2|2|2|3|11.42|2|3| | | | | | | | | | | | | | | | | | | | | | | | | | | |254|28.6|5.00|
|P01911|16.54|6|2|3|4|3.38|1|1|3.38|1|1| | | | | | | | | | | | | | | | | | | | | |13.16|2|2|266|29.9|7.74|
|P02008|32.39|1|4|4|4| | | | | | | | | | | | | | | | | | | | | | | | | | | |32.39|4|4|142|15.6|8.21|
|P02144|31.82|1|4|4|6| | | | | | | | | | | | | | | | | | | | | | | | | | | |31.82|4|6|154|17.2|7.68|
|P02511|88.00|1|15|15|79|40.57|4|5|40.57|4|4| | | |41.14|7|9| | | | | | |36.57|7|8| | | |32.57|5|8|64.57|11|41|175|20.1|7.33|
|P02533|61.44|21|6|37|206| | | |7.20|4|4|11.65|6|7|6.78|4|5|19.07|10|13| | | |31.99|15|17|48.31|24|34|15.04|7|9|43.01|22|117|472|51.5|5.16|
|P02538|56.74|7|9|37|245|3.37|2|2|9.75|7|8|11.17|9|11|3.01|3|4|9.75|7|9|1.42|1|1|23.05|13|14|14.01|13|18|16.49|11|13|45.92|29|165|564|60.0|8.00|
|P02545|19.43|1|11|11|19| | | |1.51|1|1|4.67|3|3| | | | | | | | | | | | |1.51|1|1| | | |16.57|9|14|664|74.1|7.02|
|P02647|57.30|2|17|18|48|24.34|5|6|5.24|1|1|8.61|2|2|28.84|6|7|8.61|2|2| | | |8.24|2|2| | | |19.85|4|5|55.43|16|23|267|30.8|5.76|
|P02649|48.58|2|13|13|21| | | | | | |4.73|1|1| | | | | | | | | | | | | | | | | | |48.58|13|20|317|36.1|5.73|
|P02671|4.97|1|5|5|8| | | | | | | | | | | | | | | | | | |0.92|1|1| | | | | | |4.97|5|7|866|94.9|6.01|
|P02675|9.98|1|4|4|6| | | | | | |3.05|1|1| | | | | | | | | | | | | | | | | | |9.98|4|5|491|55.9|8.27|
|P02679|27.81|1|10|10|15| | | | | | |5.74|2|2| | | | | | | | | |7.28|2|2| | | | | | |16.56|6|11|453|51.5|5.62|
|P02743|38.12|2|13|13|103|33.63|10|12|36.77|10|12|36.77|11|13|26.46|7|8|24.22|7|10| | | |14.35|4|4|7.17|2|2| | | |36.77|11|42|223|25.4|6.54|
|P02751|55.78|1|50|99|896|37.18|68|126|21.00|32|46|50.04|85|168|33.57|58|81|12.53|23|29|2.10|4|4|4.19|7|7|7.63|13|15|3.27|6|6|43.84|75|414|2386|262.5|5.71|
|P02763|35.32|1|5|7|23|35.32|7|10|27.86|5|5|22.89|4|4|19.90|3|3|4.98|1|1| | | | | | | | | | | | | | | |201|23.5|5.02|
|P02766|23.81|1|2|2|5|23.81|2|2|14.97|1|1| | | |8.84|1|1| | | | | | | | | | | | | | | |14.97|1|1|147|15.9|5.76|
|P02768|82.59|2|51|58|440|67.00|39|60|46.14|28|39|68.31|39|52|47.29|25|35|55.17|37|48|3.78|3|3|20.36|16|18|20.69|16|22|29.72|20|22|72.25|41|141|609|69.3|6.28|
|P02774|16.67|1|7|7|9|12.24|5|5| | | | | | |1.48|1|1| | | | | | | | | | | | | | | |4.43|2|3|474|52.9|5.54|
|P02786|2.89|1|2|2|2| | | | | | | | | | | | | | | | | | | | | | | | | | | |2.89|2|2|760|84.8|6.61|
|P02787|39.83|1|24|24|51|21.06|12|15|7.59|4|4|19.77|11|11| | | | | | | | | | | | | | | | | | |29.80|17|21|698|77.0|7.12|
|P02788|85.07|5|77|77|1312|71.69|50|119|70.56|48|130|71.41|53|125|41.97|27|36|64.08|47|73|10.42|5|6| | | |7.04|4|5|4.37|2|2|83.94|68|816|710|78.1|8.12|
|P02792|36.00|1|6|6|9|4.00|1|1|4.00|1|1|0.00| | | | | | | | | | | | | | | | | | | | |36.00|6|7|175|20.0|5.78|
|P62491|62.50|1|1|14|71|29.17|6|7|29.17|5|6|20.83|5|5|29.63|6|6|5.09|1|1| | | |9.26|2|2|5.09|1|1|25.00|5|6|59.72|12|37|216|24.4|6.57|
|P02794|74.86|2|15|15|54|31.15|4|5|33.33|5|8|36.07|6|6|10.93|2|2|18.03|3|5| | | | | | | | | |37.16|6|8|69.40|12|20|183|21.2|5.55|
|P03886|8.81|1|3|3|6| | | | | | | | | | | | | | | | | | | | | | | | | | | |8.81|3|6|318|35.6|6.55|
|P03905|4.14|1|2|2|2| | | | | | | | | | | | | | | | | | | | | | | | | | | |4.14|2|2|459|51.5|9.35|
|P03915|6.14|1|2|2|6| | | | | | | | | | | | | | | | | | | | | | | | | | | |6.14|2|6|603|67.0|9.03|
|P03973|34.85|1|7|7|27|27.27|4|5|9.09|1|1|26.52|3|4|9.85|1|1|34.85|6|8| | | | | | |6.06|1|1| | | |33.33|5|7|132|14.3|8.75|
|P04004|23.85|2|10|10|22|3.35|1|1| | | |10.25|3|3|2.09|1|1| | | | | | | | | | | | | | | |23.22|9|17|478|54.3|5.80|
|P04040|23.53|1|9|9|20|14.80|5|5| | | |10.63|5|6| | | | | | | | | | | | |2.47|1|1|2.47|1|1|10.82|5|7|527|59.7|7.39|
|P04066|21.46|1|8|8|24|17.60|5|9|11.16|3|5|6.01|2|2| | | | | | | | | | | | | | | | | | |9.87|4|8|466|53.7|6.84|
|P04075|96.98|2|39|46|621|49.18|13|18|34.07|11|16|60.44|18|23|74.18|25|41|62.36|26|38|3.85|1|1|62.36|22|42|69.78|29|53|69.78|31|51|92.03|34|340|364|39.4|8.09|
|P04080|85.71|2|6|6|26|48.98|3|3|24.49|2|2|24.49|2|2|12.24|1|2|12.24|1|1| | | |45.92|3|5|24.49|2|2|61.22|5|5|70.41|4|4|98|11.1|7.56|
|P04083|74.86|1|27|27|179|53.76|14|18|40.17|10|12|53.76|14|19|33.82|9|10|11.27|2|2| | | |32.95|8|9|14.74|3|3|32.08|9|11|73.70|25|95|346|38.7|7.02|
|P04156|11.86|1|3|3|11|4.74|1|1| | | | | | | | | |4.74|1|1| | | | | | | | | | | | |11.86|3|9|253|27.6|9.00|
|P04179|61.26|1|13|13|47|53.60|7|10|30.18|6|7|23.42|5|5|9.91|2|2|13.06|2|2| | | | | | | | | | | | |43.24|11|21|222|24.7|8.25|
|P04264|74.53|3|70|80|779|7.76|4|5|35.87|22|27|43.63|29|39|40.53|25|33|46.58|34|45|11.49|7|7|48.60|41|58|60.56|56|88|42.55|32|42|65.84|51|435|644|66.0|8.12|
|P04279|93.72|1|76|84|1518|76.62|40|136|73.16|36|154|65.15|32|122|75.97|36|90|67.32|39|111|56.71|26|39|66.67|32|68|66.45|41|94|67.53|42|82|81.82|54|622|462|52.1|9.29|
|P04350|79.73|1|4|35|1039|61.94|20|49|61.94|21|71|58.78|19|42|62.84|24|90|65.32|27|101|7.66|2|3|32.88|12|23|33.11|14|28|41.67|15|30|74.55|26|602|444|49.6|4.88|
|P04406|86.87|2|25|26|321|49.55|10|18|43.58|10|15|54.93|11|24|48.66|11|19|33.13|7|9|3.28|1|1|44.48|9|20|34.33|9|17|53.13|10|16|77.31|18|182|335|36.0|8.46|
|P04424|7.33|1|2|2|4| | | | | | | | | | | | | | | | | | | | | | | | | | | |7.33|2|4|464|51.6|6.48|
|P04632|68.66|2|11|11|38|13.06|3|3|8.21|2|3| | | |9.70|3|3| | | | | | | | | | | | | | | |66.42|10|29|268|28.3|5.20|
|P04792|94.15|1|24|24|218|68.29|8|15|36.10|6|7|77.07|9|13|63.90|13|18|59.51|9|14| | | |60.98|11|15|60.49|10|11|70.73|16|23|87.32|19|102|205|22.8|6.40|
|P04839|24.39|1|11|11|22|22.46|10|11| | | |14.21|7|8| | | | | | | | | | | | | | | | | | |4.74|2|3|570|65.3|8.63|
|P04843|66.39|1|38|38|172|27.35|14|14|34.10|16|19|37.56|18|24|3.79|2|2|13.51|6|8| | | | | | | | | | | | |60.30|32|105|607|68.5|6.38|
|P04844|55.15|1|26|26|152|16.32|7|9|34.71|14|16|38.83|14|16| | | | | | | | | | | | | | | |2.22|1|1|44.53|21|110|631|69.2|5.69|
|P04899|56.06|4|10|17|92|23.38|6|7| | | |34.37|9|11|20.28|6|6|3.10|1|1| | | | | | |3.10|1|1|18.59|6|8|51.27|14|58|355|40.4|5.54|
|P05023|44.09|1|28|47|239|16.62|12|14|21.11|17|20|18.96|17|19|0.88|1|1|1.56|2|2| | | | | | | | | | | | |43.01|44|183|1023|112.8|5.49|
|P05026|24.42|1|5|5|5| | | | | | |3.63|1|1| | | | | | | | | | | | | | | | | | |20.79|4|4|303|35.0|8.53|
|P05060|6.06|1|4|4|4|2.36|1|1|2.07|2|2|1.62|1|1| | | | | | | | | | | | | | | | | | | | | |677|78.2|5.07|
|P05067|4.68|1|2|3|11|1.56|1|1|1.56|1|1|1.56|1|1| | | | | | | | | | | | | | | | | | |4.68|3|8|770|86.9|4.82|
|P05090|33.33|1|7|7|11|6.35|2|2| | | |5.82|1|1| | | |5.82|1|1| | | | | | | | | | | | |32.80|6|7|189|21.3|5.15|
|P05091|42.55|1|15|18|28|20.12|6|6| | | |19.34|9|10|2.32|1|1| | | | | | | | | | | | |9.48|4|4|16.25|7|7|517|56.3|7.05|
|P05107|36.15|3|25|25|52|14.17|10|11| | | |30.69|21|25| | | | | | | | | | | | | | | | | | |17.30|13|16|769|84.7|6.95|
|P05109|86.02|1|12|12|81|86.02|11|16|54.84|8|11|77.42|11|15|53.76|7|9|33.33|4|5|7.53|1|1|26.88|3|5|11.83|1|1|52.69|7|9|47.31|5|9|93|10.8|7.03|
|P05141|51.34|1|9|20|88|22.15|6|7|34.56|10|11|18.46|6|7|2.68|1|1|2.68|1|1| | | | | | | | | | | | |48.32|18|59|298|32.8|9.69|
|P05154|49.26|2|18|19|90|31.28|12|13|27.59|10|12|30.30|12|13|2.71|1|1|2.71|1|1| | | | | | | | | | | | |42.36|16|50|406|45.6|9.26|
|P05161|18.79|1|3|3|3| | | | | | | | | | | | | | | | | | | | | | | | | | | |18.79|3|3|165|17.9|7.44|
|P05164|55.84|2|38|38|167|50.74|32|54|30.47|18|24|44.56|27|34|13.42|8|9|5.37|3|3| | | | | | | | | | | | |33.56|19|43|745|83.8|8.97|
|P05165|2.88|1|2|2|2| | | | | | | | | | | | | | | | | | | | | | | | | | | |2.88|2|2|728|80.0|7.52|
|P05186|16.98|1|6|6|6| | | | | | | | | | | | | | | | | | | | | | | | | | | |16.98|6|6|524|57.3|6.67|
|P05198|33.33|1|10|10|30|3.81|1|1|3.81|1|1|3.81|1|1|6.98|2|2|3.81|1|1| | | |3.81|1|1|3.81|1|1|10.16|2|2|33.33|10|20|315|36.1|5.08|
|P05386|89.47|1|3|4|10|14.04|1|1|14.04|1|1|14.04|1|1| | | | | | | | | | | | | | | | | | |89.47|4|7|114|11.5|4.32|
|P05387|95.65|1|8|9|32|40.87|3|3|69.57|4|5|24.35|2|3|53.04|3|4|53.91|3|3| | | | | | | | | | | | |94.78|8|14|115|11.7|4.54|
|P05388|56.47|2|16|16|66|17.67|5|5|24.61|6|7|17.67|5|5|5.99|2|2| | | | | | | | | | | | |14.51|3|3|55.52|15|44|317|34.3|5.97|
|P05452|27.72|2|4|4|6| | | | | | | | | | | | | | | | | | | | | | | | | | | |27.72|4|6|202|22.5|5.67|
|P05556|3.88|1|2|2|4| | | | | | | | | | | | | | | | | | | | | | | | | | | |3.88|2|4|798|88.4|5.39|
|P05783|43.49|13|17|19|57|9.53|2|2|5.58|3|3|12.09|6|6|6.51|3|3|3.72|2|2| | | |6.51|3|3|11.63|5|5|27.67|11|12|18.60|7|21|430|48.0|5.45|
|P05787|56.52|12|21|34|161|13.46|7|7|11.59|7|8|13.87|9|10|14.70|9|10|11.18|7|9|0.00| | |18.63|11|12|13.66|12|16|34.16|17|19|32.09|19|70|483|53.7|5.59|
|P06280|20.75|1|8|8|13|6.53|2|2|2.10|1|1|4.90|2|2| | | | | | | | | | | | | | | |5.36|2|2|11.66|5|6|429|48.7|5.60|
|P06396|38.36|2|24|24|97|18.93|10|15|9.59|5|5|20.20|11|15|3.32|2|2|1.92|1|1| | | | | | |8.06|5|6|6.27|3|4|25.19|15|49|782|85.6|6.28|
|P06576|69.94|1|27|27|365|60.68|21|37|64.27|25|53|59.17|24|44|39.32|14|16|25.14|10|11| | | |27.41|11|12|24.95|9|15|31.57|12|14|69.94|26|163|529|56.5|5.40|
|P06702|87.72|1|11|11|115|85.96|9|21|78.95|7|19|85.96|8|15|70.18|7|11|57.02|7|12|13.16|1|1|50.88|5|6|30.70|3|7|51.75|6|11|44.74|5|12|114|13.2|6.13|
|P06727|58.84|2|25|25|63|12.37|4|4| | | |9.09|3|3|19.44|7|9|6.06|2|2| | | |5.30|2|2|3.03|1|1|36.36|13|18|46.97|16|24|396|45.4|5.38|
|P06730|18.43|2|4|4|10|8.29|2|2| | | | | | | | | | | | | | | | | | | | | | | | |18.43|4|8|217|25.1|6.15|
|P06731|5.98|1|2|3|5| | | | | | |5.98|3|3| | | | | | | | | | | | | | | | | | |1.28|1|2|702|76.7|5.92|
|P06733|86.41|5|47|51|540|60.60|22|57|56.45|21|56|62.90|28|56|51.38|21|29|71.66|38|68| | | |13.82|5|5|22.12|9|13|22.58|7|8|80.41|37|248|434|47.1|7.39|
|P06737|24.56|1|12|17|48|0.94|1|1|0.00| | |1.65|2|2|2.36|2|2|0.00| | | | | |2.36|2|2|0.94|1|1|0.94|1|1|24.56|17|37|847|97.1|7.17|
|P06744|57.89|4|27|27|138|32.44|13|15|34.77|15|24|31.54|16|22|8.24|4|6|9.14|5|5| | | | | | |2.69|1|1|2.69|1|1|53.23|22|64|558|63.1|8.32|
|P06748|12.93|2|5|5|10| | | |6.80|2|3| | | | | | |12.93|4|4| | | | | | | | | | | | |6.80|2|3|294|32.6|4.78|
|P06858|39.37|1|19|20|132|17.05|6|7|20.63|8|11|24.00|9|12|13.26|5|6|3.79|2|2| | | |9.89|3|4|5.89|3|4|14.74|7|10|37.89|17|76|475|53.1|8.15|
|P06865|19.47|1|9|10|24|5.67|2|2|11.34|5|5|4.16|2|2| | | | | | | | | | | | | | | | | | |14.56|8|15|529|60.7|5.16|
|P07099|17.58|1|9|9|9| | | | | | | | | | | | | | | | | | | | | | | | | | | |17.58|9|9|455|52.9|7.25|
|P07108|74.71|2|6|6|30|52.87|4|8|52.87|4|7|52.87|3|3|41.38|3|5|63.22|4|4| | | |29.89|2|2| | | | | | |22.99|1|1|87|10.0|6.57|
|P07195|50.90|1|12|19|86|22.16|7|7|29.64|10|11|31.14|9|10|17.66|5|6|6.89|3|3| | | | | | |3.59|1|1|3.59|1|1|47.60|15|47|334|36.6|6.05|
|P07203|60.10|1|9|9|28|8.87|1|1| | | | | | |17.24|2|2| | | | | | |14.78|2|2| | | |21.67|3|3|60.10|9|18|203|22.1|6.55|
|P07205|88.97|1|35|41|429|66.43|23|30|59.71|22|30|73.14|27|39|60.67|23|30|52.28|19|23| | | |33.33|13|15|56.59|21|30|58.75|23|31|84.65|38|201|417|44.8|8.54|
|P07237|64.76|1|39|39|292|43.31|18|22|49.41|20|27|47.64|21|28|30.12|15|19|48.23|22|29|2.36|1|1|32.87|18|23|26.77|13|21|47.83|25|34|58.86|29|88|508|57.1|4.87|
|P07288|83.91|1|14|15|188|80.08|14|33|33.72|7|17|52.87|10|23|50.19|9|16|44.83|9|14|8.05|2|2|18.77|5|7|18.39|4|4|33.33|8|10|71.65|12|62|261|28.7|7.68|
|P07305|11.34|1|2|2|2| | | |4.64|1|1| | | | | | | | | | | | | | | | | | | | | |6.70|1|1|194|20.9|10.84|
|P07311|20.20|1|2|2|3|13.13|1|1| | | | | | | | | | | | | | | | | | | | | | | | |20.20|2|2|99|11.3|9.31|
|P07339|34.22|2|14|14|29|20.63|6|6|2.67|1|1|25.49|7|8| | | | | | | | | | | | | | | | | | |27.67|10|14|412|44.5|6.54|
|P07355|72.86|3|29|29|153|65.19|20|26|54.87|14|20|67.55|18|23|23.01|6|7|20.94|6|6| | | |6.19|2|2|6.49|2|2|18.29|5|6|67.26|25|61|339|38.6|7.75|
|P07358|2.54|1|2|2|3|1.35|1|1|1.18|1|1|1.18|1|1| | | | | | | | | | | | | | | | | | | | | |591|67.0|8.13|
|P07384|36.55|1|25|25|57| | | | | | |9.94|6|6| | | | | | | | | | | | | | | | | | |36.41|24|51|714|81.8|5.67|
|P07437|87.39|3|5|42|1111|61.26|22|52|57.43|21|61|56.98|21|44|71.40|30|81|61.71|31|105|6.98|2|3|34.91|14|24|37.61|18|34|50.68|20|38|79.73|31|669|444|49.6|4.89|
|P07478|12.15|3|2|2|18| | | | | | | | | | | | | | | | | | | | | | | | | | | |12.15|2|18|247|26.5|4.92|
|P07602|19.27|1|8|10|26|15.08|8|9|4.58|2|2|1.91|1|1| | | |1.72|1|1| | | | | | | | | | | | |8.02|4|13|524|58.1|5.17|
|P07686|16.55|1|8|9|13|3.60|2|2|4.86|3|3|4.14|2|2| | | | | | | | | | | | | | | | | | |9.53|5|6|556|63.1|6.76|
|P07737|85.00|2|14|14|79|53.57|5|5|24.29|3|3|35.71|4|5|48.57|6|8|23.57|3|3|12.14|2|2|23.57|3|3|34.29|6|8|54.29|7|10|81.43|12|32|140|15.0|8.27|
|P07738|8.49|1|2|2|2| | | | | | | | | | | | | | | | | | | | | | | | | | | |8.49|2|2|259|30.0|6.54|
|P07741|90.00|1|14|14|43|15.56|2|3| | | | | | |4.44|1|1| | | | | | |4.44|1|1| | | | | | |90.00|14|38|180|19.6|6.02|
|P07814|53.17|2|70|71|260|10.91|12|12|20.37|23|27|13.96|17|19|6.28|9|9|6.81|9|9| | | |1.85|3|3|3.11|4|4|5.49|8|9|46.83|59|168|1512|170.5|7.33|
|P07858|17.11|1|5|5|7|4.13|1|1|4.13|1|1|11.80|4|4| | | | | | | | | | | | | | | | | | |5.31|1|1|339|37.8|6.30|
|P07864|86.75|2|30|32|319|65.96|19|26|49.10|17|23|58.13|17|21|62.05|20|30|62.35|21|28|3.01|1|1|3.92|1|1|10.54|3|3|13.86|4|5|80.72|27|181|332|36.3|7.46|
|P07900|79.10|4|58|87|1074|56.69|38|56|57.24|44|61|54.64|42|65|49.18|42|57|47.13|46|66|1.23|1|1|47.40|41|57|48.91|45|73|52.73|52|89|70.49|68|549|732|84.6|5.02|
|P07910|25.82|4|8|8|17|6.86|2|2|6.86|2|2|7.52|2|2|3.59|1|1|3.59|1|1|3.59|1|1| | | |3.59|1|1| | | |18.95|6|7|306|33.6|5.08|
|P07911|36.72|1|20|20|179| | | |11.41|8|9|29.53|17|27| | | |9.38|6|6| | | | | | |2.19|2|2|12.03|7|8|33.13|18|127|640|69.7|5.24|
|P07919|64.84|2|5|5|10|27.47|2|3| | | | | | | | | | | | | | | | | | | | | | | | |64.84|5|7|91|10.7|4.44|
|P07951|23.24|3|2|8|18|11.27|3|3|8.10|2|2|3.17|1|1| | | |11.27|3|6| | | | | | | | | |8.80|2|2|10.56|4|4|284|32.8|4.70|
|P07954|61.96|1|26|26|140|46.08|15|23|50.39|18|28|48.82|17|30|9.22|3|3|10.59|5|6| | | | | | | | | | | | |60.39|22|50|510|54.6|8.76|
|P07996|5.73|1|6|6|7| | | | | | | | | | | | | | | | | | | | | | | | | | | |5.73|6|7|1170|129.3|4.94|
|P07998|20.51|1|2|2|3| | | | | | | | | | | | |13.46|1|1| | | | | | | | | | | | |7.05|1|2|156|17.6|8.79|
|P08107;P0DMV8|69.11|3|31|49|566|44.31|20|32|39.00|19|31|55.38|27|49|35.41|21|27|34.17|23|30|4.21|2|2|32.92|19|26|39.63|28|38|39.63|26|38|61.47|35|293|641|70.0|5.66|
|P08118|14.04|1|3|3|9|7.89|1|1| | | |14.04|3|3| | | | | | | | | | | | | | | | | | |14.04|3|5|114|12.9|5.50|
|P08133|63.15|1|44|44|136|34.62|21|22|14.41|7|7|34.03|20|23|5.20|3|3| | | | | | |9.66|5|5|1.93|1|1|17.83|9|10|44.58|27|65|673|75.8|5.60|
|P08134|62.69|1|2|11|59|10.88|2|2| | | |19.17|3|3|29.02|6|7|18.65|4|5| | | |8.81|1|1|18.65|4|5|18.65|4|4|56.48|10|32|193|22.0|6.58|
|P08174|40.42|1|14|14|37|16.01|3|4|5.25|2|2|4.46|2|2| | | |2.36|1|1| | | | | | | | | | | | |35.96|12|28|381|41.4|7.59|
|P08195|14.60|1|6|6|12|3.97|2|2|4.29|2|2|0.00| | |0.00| | | | | | | | | | | | | | |0.00| | |14.60|6|8|630|68.0|5.01|
|P08236|8.29|1|6|6|6|3.84|3|3| | | |2.00|1|1| | | | | | | | | | | | | | | | | | |2.46|2|2|651|74.7|7.02|
|P08237|62.56|1|37|40|233|7.82|4|5|6.15|3|3|17.69|11|12|6.28|4|4| | | | | | |1.15|1|1|2.56|2|2| | | |61.03|38|206|780|85.1|7.99|
|P08238|67.96|3|48|74|866|37.57|23|28|45.17|30|38|39.23|27|35|39.64|26|34|44.20|32|42|1.24|1|1|40.06|32|46|39.78|30|42|53.04|46|73|63.81|58|527|724|83.2|5.03|
|P08240|5.02|1|3|3|3| | | | | | | | | | | | | | | | | | | | | | | | | | | |5.02|3|3|638|69.8|8.95|
|P08243|8.91|1|4|4|5| | | | | | | | | | | | | | | | | | | | | | | | | | | |8.91|4|5|561|64.3|6.86|
|P08246|36.70|1|7|7|28|26.22|6|7| | | |26.22|6|6|6.37|2|2| | | | | | | | | | | | | | | |36.70|7|11|267|28.5|9.35|
|P08253|26.21|2|12|12|28|5.76|3|3|5.76|3|4|17.27|7|7| | | | | | | | | | | | | | | | | | |8.18|5|12|660|73.8|5.47|
|P08263|9.01|4|2|2|4| | | | | | | | | | | | | | | | | | | | | | | | | | | |9.01|2|4|222|25.6|8.88|
|P08294|12.50|1|2|2|2|6.67|1|1| | | | | | | | | | | | | | | | | | | | | | | | |5.83|1|1|240|25.8|6.61|
|P08311|54.90|1|16|16|59|40.39|10|12|15.29|4|4|37.25|8|10|36.86|9|13| | | | | | | | | | | | | | | |40.78|9|20|255|28.8|11.19|
|P08473|58.80|1|46|46|169|19.87|11|11|9.20|6|6|45.73|29|35| | | |2.13|1|1| | | | | | | | | |4.40|3|4|51.20|39|112|750|85.5|5.73|
|CON_007130003|9.11|3|1|6|60|6.44|4|10|6.44|4|12|6.44|4|7|6.44|5|11|9.11|4|10| | | |4.00|2|4|4.00|2|3|4.00|2|3| | | |450|50.0|5.11|
|P08559|49.74|1|13|17|47|11.28|4|4|16.92|6|10|14.87|5|6|5.90|2|2|2.31|1|1| | | |5.90|2|2| | | | | | |43.85|15|22|390|43.3|8.06|
|P08574|46.77|1|12|12|25|11.38|2|2|3.69|1|1| | | | | | | | | | | | | | | | | | | | | |46.77|12|22|325|35.4|9.00|
|P08575|16.10|1|17|17|25|4.60|5|5| | | |15.11|16|18| | | | | | | | | | | | | | | | | | |1.69|2|2|1304|147.2|6.15|
|P08670|68.03|8|38|45|137|50.64|27|33|22.10|10|11|36.70|15|15|18.03|8|8|28.97|10|11|4.51|1|1|7.73|4|4|3.86|3|4|17.81|8|9|54.29|29|41|466|53.6|5.12|
|P08697|6.11|1|2|2|2|3.46|1|1| | | | | | | | | | | | | | | | | | | | | | | | |2.65|1|1|491|54.5|6.29|
|P08727|37.75|12|7|17|76|9.00|2|2|9.00|4|4|10.25|5|5|5.75|3|3|8.00|4|4| | | |12.25|6|6|8.00|4|4|10.00|4|4|28.75|13|44|400|44.1|5.14|
|P08729|15.99|8|3|11|51|3.20|2|2|1.49|1|1|3.20|2|2|5.54|3|3|7.46|4|5|0.00| | |5.33|3|3|4.48|5|7|7.46|4|5|12.58|8|23|469|51.4|5.48|
|P08754|38.98|4|6|12|65|7.34|2|2| | | |12.43|4|5|15.25|5|5|3.11|1|1| | | | | | |3.11|1|1|14.41|5|7|38.98|12|44|354|40.5|5.69|
|P08758|84.69|1|30|32|195|47.50|13|16|35.31|11|13|45.94|14|18|32.19|11|13| | | | | | |27.50|9|13| | | |48.13|13|18|74.69|24|104|320|35.9|5.05|
|P08779|61.52|17|12|33|172| | | |7.19|4|4|11.63|6|7|6.77|4|5|12.26|7|10| | | |33.19|14|16|23.89|15|23|12.68|6|8|52.64|24|99|473|51.2|5.05|
|P08865|47.46|1|10|10|40|18.31|5|5|12.88|3|3|20.34|4|4|2.71|1|1| | | | | | | | | | | | | | | |47.46|9|27|295|32.8|4.87|
|P08962|7.56|1|2|3|11|4.20|1|1| | | | | | | | | | | | | | | | | | | | | |0.00| | |7.56|3|10|238|25.6|7.81|
|P09104|18.43|3|2|6|71|14.06|4|12|10.60|3|7|10.60|3|4|5.53|2|3|12.67|3|5| | | |0.00| | | | | | | | |18.43|6|40|434|47.2|5.03|
|P09110|41.98|1|10|10|18|4.48|1|1| | | | | | | | | | | | | | | | | | | | | | | | |41.98|10|17|424|44.3|8.44|
|P09132|11.11|1|2|2|4| | | | | | | | | | | | | | | | | | | | | | | | | | | |11.11|2|4|144|16.1|9.85|
|P09211|70.48|1|12|12|82|50.95|7|9|16.19|3|4|42.38|6|7|35.71|5|5|6.19|2|2| | | |24.76|4|4|18.57|4|4|40.00|7|8|69.52|11|39|210|23.3|5.64|
|P09382|57.04|2|6|6|13|19.26|2|2| | | | | | |19.26|2|3|13.33|1|1| | | | | | |5.93|1|1|19.26|2|2|45.19|4|4|135|14.7|5.50|
|P09417|40.57|1|10|10|17| | | |4.51|1|1| | | | | | | | | | | | | | | | | | |11.89|2|2|40.57|9|14|244|25.8|7.37|
|P09466|36.11|1|6|6|25|32.22|5|7|12.78|2|3|12.78|2|2|21.11|4|5| | | | | | | | | | | | |5.00|1|1|36.11|6|7|180|20.6|5.57|
|P09467|27.81|2|7|7|12|4.73|1|1| | | |4.73|1|1| | | | | | | | | | | | | | | | | | |27.81|7|10|338|36.8|6.99|
|P09471|27.40|1|6|8|30|3.11|1|1| | | |10.73|3|4|6.50|2|2|3.11|1|1| | | | | | |3.11|1|1|10.17|3|4|23.73|7|17|354|40.0|5.53|
|P09488|59.17|2|9|15|29| | | | | | |5.05|1|1| | | |5.05|1|1| | | | | | | | | |26.61|5|5|47.71|12|22|218|25.7|6.70|
|P09496|22.58|1|7|8|47|16.94|5|5|16.94|5|5| | | |22.58|7|8|22.58|7|11| | | |13.31|4|4| | | |11.69|3|4|13.71|4|10|248|27.1|4.51|
|P09497|10.04|1|2|2|2| | | | | | | | | | | | |10.04|2|2| | | | | | | | | | | | | | | |229|25.2|4.64|
|P09525|53.61|1|16|18|50|33.23|9|12|36.68|10|10|30.09|7|7|9.09|3|3| | | | | | |3.45|1|1| | | | | | |41.38|12|17|319|35.9|6.13|
|P09543|41.81|1|19|20|48|1.43|1|1| | | |2.61|1|1|3.33|2|2| | | | | | |1.43|1|1| | | |7.84|4|4|41.57|19|39|421|47.5|9.07|
|P09622|49.31|1|23|23|89|27.11|10|13|20.83|9|11|28.29|13|16|5.11|2|2|19.45|8|10| | | | | | | | | | | | |39.49|16|37|509|54.1|7.85|
|P09651|25.81|2|7|7|13| | | | | | | | | | | | |3.23|1|1|20.97|5|6|3.23|1|1|3.76|2|2|11.29|3|3|0.00| | |372|38.7|9.13|
|P09668|24.78|1|8|8|15|4.48|1|1| | | |7.46|2|2|3.88|1|1| | | | | | | | | | | | | | | |17.31|5|11|335|37.4|8.07|
|P09669|50.67|1|9|9|19|18.67|2|2|20.00|2|2|20.00|2|2| | | | | | | | | | | | | | | | | | |50.67|8|13|75|8.8|10.39|
|P09917|7.57|1|4|4|4| | | | | | |3.12|2|2| | | | | | | | | | | | | | | | | | |4.45|2|2|674|77.9|5.77|
|P09936|49.78|1|9|9|15|24.66|5|6|4.48|1|1|4.48|1|1| | | | | | | | | | | | |9.87|1|1| | | |34.98|5|6|223|24.8|5.48|
|P09960|25.70|1|11|11|19| | | | | | |3.93|2|2| | | | | | | | | | | | | | | |7.04|3|3|19.97|9|14|611|69.2|6.18|
|P09972|36.26|2|6|13|99|8.52|2|5|6.32|1|2|10.44|3|3|15.93|3|4|10.44|3|6| | | |1.92|1|1|13.74|5|8|13.74|5|6|35.71|11|64|364|39.4|6.87|
|P0C0S5|31.25|2|2|4|21|12.50|2|2|12.50|2|2|12.50|2|2|12.50|2|2|7.03|1|1| | | |5.47|1|1| | | | | | |31.25|4|11|128|13.5|10.58|
|Q8IW75|1.93|1|1|1|57| | | | | | |1.93|1|1| | | | | | | | | | | | | | | | | | |1.93|1|56|414|47.1|9.29|
|P0C221|12.36|1|6|6|9| | | | | | | | | | | | | | | | | | | | | | | | | | | |12.36|6|9|793|93.6|6.71|
|P0C5Z0|56.52|2|4|4|11|13.91|1|1|13.04|1|1| | | | | | | | | | | | | | | | | | | | | |56.52|4|9|115|12.7|10.67|
|P0C7A2|4.65|3|2|2|3| | | | | | | | | | | | | | | | | | | | | | | | | | | |4.65|2|3|387|43.6|4.78|
|P0C7I6|6.31|1|2|2|2| | | | | | | | | | | | | | | | | | | | | | | | |2.43|1|1|3.88|1|1|412|46.0|6.27|
|P0C870|17.72|1|4|4|4| | | | | | | | | | | | | | | | | | | | | | | | | | | |17.72|4|4|316|35.9|5.38|
|P0C875|19.94|1|6|6|9| | | | | | | | | | | | |3.12|1|1| | | | | | | | | | | | |16.82|5|8|321|37.4|9.29|
|P0C8F1|48.98|1|5|5|15|9.18|1|1|24.49|2|2|24.49|3|4|9.18|1|1|37.76|3|4| | | | | | | | | |9.18|1|1|23.47|2|2|98|11.4|8.62|
|P0CG05|74.53|7|5|5|24|47.17|3|5|14.15|1|1|74.53|5|7|14.15|1|1|23.58|2|3| | | | | | |9.43|1|1| | | |37.74|3|6|106|11.3|7.24|
|P0CG33|23.81|4|2|15|21| | | |1.73|1|1| | | | | | |10.39|5|5| | | | | | | | | | | | |17.60|10|15|693|79.8|5.45|
|P0CG38|11.35|6|3|10|177|4.28|5|8|4.93|6|12|4.93|6|9|4.93|6|9|4.93|6|8|0.93|1|1|4.93|6|8|3.63|4|4|4.93|7|12|11.26|8|106|1075|121.2|6.21|
|P0CW22;P08708|56.30|1|7|7|18|16.30|1|2|7.41|1|1| | | | | | | | | | | | | | | | | | | | | |56.30|7|15|135|15.5|9.85|
|P0CW27|13.21|1|4|4|9| | | | | | | | | | | | |5.47|1|1| | | | | | | | | | | | |13.21|4|8|439|48.7|10.59|
|P0DI81|18.57|1|2|2|2| | | | | | | | | | | | | | | | | | | | | | | | | | | |18.57|2|2|140|16.4|6.52|
|P0DJD1|19.93|3|11|31|65| | | | | | | | | | | | | | | | | | | | | | | | | | | |19.93|31|65|1756|197.2|6.20|
|P0DJG4|7.53|1|4|4|7| | | | | | | | | | | | | | | | | | | | | | | | | | | |7.53|4|7|465|53.0|9.23|
|P0DJI8|45.90|2|3|3|5| | | | | | | | | | | | | | | | | | | | | | | | |45.90|3|5| | | |122|13.5|6.79|
|P52789|5.56|2|1|5|55|4.14|4|4|4.14|4|9|4.14|4|7|1.20|1|1|1.96|2|2| | | | | | | | | |1.96|2|2|5.56|5|30|917|102.3|6.05|
|P0DKB6|68.38|1|6|7|26|14.71|1|1|36.03|3|4| | | | | | | | | | | | | | | | | | | | | |68.38|7|21|136|15.1|9.94|
|P10155|22.30|1|10|10|14| | | | | | | | | | | | | | | | | | | | | | | | | | | |22.30|10|14|538|60.6|8.03|
|P10253|24.26|2|17|17|67|8.72|6|6|5.99|3|3|2.94|2|2|4.31|3|3|2.31|2|2| | | | | | | | | |1.26|1|1|24.26|17|50|952|105.3|6.00|
|P10301|28.90|1|3|4|7| | | | | | | | | | | | | | | | | | | | | | | | | | | |28.90|4|7|218|23.5|6.93|
|P10321|24.04|7|3|6|8|4.37|1|1| | | |12.30|3|3| | | | | | | | | | | | | | | | | | |16.39|4|4|366|40.6|6.04|
|P10323|50.12|1|17|18|230|48.46|17|30|41.33|15|23|44.18|14|25|28.50|10|11|20.90|8|10| | | |2.38|1|1| | | |2.38|1|1|48.69|17|129|421|45.8|9.07|
|P10398|12.87|2|4|7|17| | | | | | | | | | | | | | | | | | | | | | | | | | | |12.87|7|17|606|67.5|9.01|
|P10515|34.00|1|21|21|134|21.64|11|13|26.58|14|21|27.51|15|20|5.26|3|3|8.50|4|4| | | |2.63|1|1|1.55|1|1| | | |33.69|18|71|647|69.0|7.84|
|P10599|51.43|2|6|6|13|12.38|1|1|12.38|1|1|12.38|1|1|12.38|1|1| | | | | | | | | |20.95|2|2| | | |51.43|6|7|105|11.7|4.92|
|P10606|51.16|1|8|8|15|23.26|3|3|29.46|4|4|8.53|1|1| | | | | | | | | | | | | | | | | | |40.31|6|7|129|13.7|8.81|
|P10620|27.10|1|3|3|3| | | | | | | | | | | | | | | | | | | | | | | | | | | |27.10|3|3|155|17.6|9.39|
|P10644|60.89|1|23|29|197|37.27|15|21|41.99|15|21|37.27|14|20|25.98|10|12|36.75|12|13| | | |23.36|9|9|21.52|9|12|32.55|11|12|58.01|23|77|381|43.0|5.35|
|P10768|63.48|1|14|14|63|21.99|3|4|4.26|1|1|21.99|5|8|14.54|4|5|11.70|3|3| | | |7.09|2|2|12.06|2|2|21.28|5|7|59.93|12|31|282|31.4|7.02|
|P10809|71.03|1|44|44|221|43.11|24|33|51.13|27|42|48.34|26|33|23.56|12|15|23.91|12|14| | | |2.09|1|1|6.46|4|4|4.89|4|4|69.28|35|75|573|61.0|5.87|
|P10909|64.59|1|43|43|456|36.75|18|28|42.76|20|50|37.86|19|27|42.32|21|29|56.57|36|53|1.78|1|1|32.96|15|22|27.17|17|22|31.63|17|23|56.35|28|201|449|52.5|6.27|

| | | | | | | | | | | | | | | | | | | | | | | | | | | | | | | | | | | | | | | |
|---|---|---|---|---|---|---|---|---|---|---|---|---|---|---|---|---|---|---|---|---|---|---|---|---|---|---|---|---|---|---|---|---|---|---|---|---|---|---|
| P11021 | 63.46 | 2 | 56 | 58 | 614 | 46.79 | 27 | 46 | 51.99 | 33 | 55 | 51.38 | 35 | 77 | 45.72 | 25 | 33 | 48.93 | 38 | 53 | 3.52 | 2 | 2 | 43.58 | 26 | 31 | 52.29 | 39 | 57 | 45.57 | 29 | 42 | 54.59 | 39 | 218 | 654 | 72.3 | 5.16 |
| P11047 | 32.07 | 1 | 41 | 41 | 114 | 7.83 | 10 | 12 | 4.47 | 5 | 5 | 23.18 | 30 | 35 | 1.68 | 2 | 2 | 0.87 | 1 | 1 | | | | | | | | | | | | | 23.18 | 28 | 59 | 1609 | 177.5 | 5.12 |
| P11142 | 62.54 | 2 | 28 | 49 | 579 | 41.18 | 22 | 31 | 41.64 | 24 | 32 | 42.72 | 25 | 37 | 38.08 | 22 | 34 | 41.18 | 26 | 36 | 8.20 | 4 | 4 | 35.91 | 21 | 30 | 42.57 | 30 | 51 | 51.24 | 38 | 57 | 53.87 | 34 | 263 | 646 | 70.9 | 5.52 |
| P11172 | 13.54 | 1 | 5 | 5 | 5 | | | | | | | | | | | | | | | | | | | | | | | | | | | | 13.54 | 5 | 5 | 480 | 52.2 | 7.24 |
| P11177 | 55.71 | 1 | 16 | 16 | 121 | 32.87 | 10 | 15 | 44.85 | 13 | 23 | 39.28 | 12 | 16 | 15.88 | 5 | 5 | 12.81 | 4 | 4 | | | | 3.06 | 1 | 1 | 19.22 | 6 | 6 | 6.13 | 2 | 2 | 55.71 | 16 | 49 | 359 | 39.2 | 6.65 |
| P11182 | 3.73 | 1 | 2 | 2 | 2 | | | | | | | | | | | | | | | | | | | | | | | | | | | | 3.73 | 2 | 2 | 482 | 53.5 | 8.51 |
| P11215 | 41.49 | 2 | 41 | 42 | 96 | 15.19 | 12 | 13 | 3.99 | 3 | 3 | 37.93 | 38 | 56 | | | | | | | | | | | | | | | | | | | 23.00 | 20 | 24 | 1152 | 127.1 | 7.23 |
| P11216 | 62.63 | 2 | 44 | 53 | 253 | 8.19 | 5 | 6 | 3.44 | 2 | 3 | 7.24 | 5 | 5 | 13.17 | 10 | 11 | | | | | | | 6.52 | 5 | 7 | 0.95 | 1 | 1 | 19.69 | 12 | 15 | 57.65 | 45 | 205 | 843 | 96.6 | 6.86 |
| P11233 | 35.44 | 1 | 7 | 9 | 34 | 27.18 | 5 | 5 | 27.18 | 5 | 5 | 27.18 | 5 | 5 | 8.74 | 2 | 2 | 5.34 | 1 | 1 | | | | 5.34 | 1 | 1 | 5.34 | 1 | 2 | 5.34 | 1 | 1 | 30.58 | 6 | 12 | 206 | 23.6 | 7.11 |
| P63096 | 25.14 | 4 | 1 | 8 | 52 | 7.06 | 2 | 2 | | | | 12.43 | 4 | 5 | 12.43 | 4 | 4 | 3.11 | 1 | 1 | | | | | | | 3.11 | 1 | 1 | 14.41 | 5 | 7 | 21.19 | 7 | 32 | 354 | 40.3 | 5.97 |
| P11234 | 33.98 | 1 | 5 | 7 | 34 | 8.74 | 2 | 2 | 8.74 | 2 | 2 | 8.74 | 2 | 2 | 13.11 | 3 | 3 | 5.34 | 1 | 1 | | | | 16.50 | 3 | 3 | 9.71 | 2 | 3 | 21.84 | 4 | 4 | 28.64 | 6 | 14 | 206 | 23.4 | 6.62 |
| P11279 | 16.79 | 1 | 9 | 9 | 63 | 7.67 | 2 | 3 | 9.83 | 3 | 4 | 14.87 | 8 | 11 | 4.80 | 2 | 2 | 2.64 | 1 | 1 | | | | | | | | | | | | | 15.35 | 7 | 42 | 417 | 44.9 | 8.75 |
| P11310 | 48.22 | 1 | 21 | 21 | 92 | 38.24 | 13 | 13 | 30.40 | 12 | 13 | 28.03 | 11 | 14 | 9.50 | 3 | 4 | 17.58 | 5 | 6 | | | | 3.09 | 1 | 1 | 12.11 | 4 | 5 | 13.06 | 4 | 4 | 32.54 | 15 | 32 | 421 | 46.6 | 8.37 |
| P11388 | 7.51 | 1 | 10 | 10 | 11 | | | | | | | | | | | | | | | | | | | | | | | | | | | | 7.51 | 10 | 11 | 1531 | 174.3 | 8.72 |
| P11413 | 34.37 | 1 | 18 | 18 | 38 | 3.88 | 2 | 2 | 1.94 | 1 | 1 | 1.36 | 1 | 1 | 4.08 | 2 | 2 | | | | | | | | | | | | | 6.80 | 3 | 3 | 26.60 | 12 | 29 | 515 | 59.2 | 6.84 |
| P11441 | 12.10 | 1 | 2 | 2 | 2 | | | | | | | | | | | | | | | | | | | | | | | | | | | | 12.10 | 2 | 2 | 157 | 17.8 | 8.66 |
| P11498 | 6.62 | 1 | 6 | 6 | 6 | | | | | | | | | | | | | | | | | | | | | | | | | | | | 6.62 | 6 | 6 | 1178 | 129.6 | 6.84 |
| P11586 | 26.10 | 1 | 24 | 24 | 39 | | | | | | | | | | 3.53 | 3 | 3 | | | | | | | | | | | | | 2.25 | 2 | 2 | 26.10 | 24 | 34 | 935 | 101.5 | 7.30 |
| P11717 | 2.21 | 1 | 5 | 5 | 8 | | | | | | | 0.44 | 1 | 1 | | | | | | | | | | | | | | | | | | | 2.21 | 5 | 7 | 2491 | 274.2 | 5.94 |
| P11766 | 40.64 | 1 | 17 | 17 | 128 | 18.45 | 5 | 5 | 8.82 | 3 | 4 | 29.41 | 9 | 10 | 13.64 | 6 | 6 | 6.95 | 2 | 2 | 2.14 | 1 | 1 | 6.95 | 2 | 2 | 6.95 | 3 | 3 | 24.87 | 11 | 13 | 38.24 | 14 | 82 | 374 | 39.7 | 7.49 |
| P11908 | 61.32 | 1 | 9 | 19 | 71 | 7.23 | 2 | 2 | | | | 14.15 | 3 | 4 | 11.95 | 3 | 3 | | | | | | | 2.52 | 1 | 1 | 5.35 | 1 | 1 | 33.65 | 7 | 10 | 57.23 | 15 | 50 | 318 | 34.7 | 6.61 |
| P11940 | 30.97 | 6 | 16 | 16 | 43 | 12.11 | 5 | 6 | 6.29 | 3 | 3 | 6.45 | 3 | 3 | 1.42 | 1 | 1 | | | | | | | 1.57 | 1 | 1 | 3.30 | 2 | 2 | 12.89 | 7 | 8 | 23.27 | 12 | 19 | 636 | 70.6 | 9.50 |
| P12036 | 25.73 | 2 | 27 | 30 | 97 | 0.88 | 1 | 1 | 3.90 | 3 | 3 | 4.00 | 4 | 4 | 2.05 | 2 | 2 | 1.95 | 2 | 2 | | | | 12.96 | 12 | 13 | 9.16 | 8 | 8 | 18.62 | 17 | 21 | 18.32 | 18 | 43 | 1026 | 112.4 | 6.18 |
| P12109 | 1.65 | 1 | 2 | 2 | 3 | 0.78 | 1 | 1 | | | | | | | | | | | | | | | | | | | | | | | | | 0.88 | 1 | 2 | 1028 | 108.5 | 5.43 |

[illegible]

| | | | | | | | | | | | | | | | | | | | | | | | | | | | | | | | | | | | | | | |
|---|---|---|---|---|---|---|---|---|---|---|---|---|---|---|---|---|---|---|---|---|---|---|---|---|---|---|---|---|---|---|---|---|---|---|---|---|---|---|
| P26436 | 21.89 | 1 | 6 | 6 | 44 | 18.49 | 4 | 10 | | | | 20.38 | 4 | 8 | 20.38 | 4 | 4 | 21.51 | 4 | 5 | | | | | | | | | | | | | 20.75 | 5 | 17 | 265 | 28.1 | 4.69 |
| P26440 | 43.50 | 1 | 15 | 15 | 31 | 12.77 | 3 | 3 | 20.09 | 6 | 7 | 10.17 | 4 | 4 | | | | | | | | | | | | | | | | 2.13 | 1 | 1 | 32.39 | 12 | 16 | 423 | 46.3 | 8.19 |
| P26639 | 43.85 | 2 | 25 | 26 | 57 | | | | 1.11 | 1 | 1 | 1.11 | 1 | 1 | | | | | | | | | | | | | | | | 4.43 | 3 | 4 | 40.53 | 24 | 51 | 723 | 83.4 | 6.67 |
| P26640 | 46.99 | 1 | 55 | 55 | 301 | 22.71 | 20 | 22 | 18.83 | 18 | 21 | 17.88 | 17 | 20 | 9.41 | 11 | 12 | 9.49 | 10 | 10 | | | | 3.16 | 3 | 3 | 3.64 | 4 | 4 | 2.14 | 2 | 2 | 42.80 | 49 | 207 | 1264 | 140.4 | 7.59 |
| P26641 | 68.42 | 1 | 30 | 30 | 274 | 37.76 | 12 | 16 | 41.42 | 16 | 20 | 42.33 | 17 | 20 | 28.15 | 12 | 17 | 28.15 | 14 | 15 | | | | 11.90 | 5 | 5 | 20.59 | 10 | 13 | 13.96 | 7 | 8 | 63.62 | 28 | 160 | 437 | 50.1 | 6.67 |
| P26885 | 63.38 | 1 | 9 | 9 | 21 | 45.77 | 4 | 5 | 46.48 | 4 | 4 | 45.77 | 4 | 4 | 8.45 | 1 | 1 | | | | | | | | | | | | | | | | 52.82 | 6 | 7 | 142 | 15.6 | 9.13 |
| P27105 | 63.89 | 2 | 15 | 15 | 80 | 46.88 | 9 | 13 | 49.31 | 10 | 14 | 42.36 | 8 | 12 | 11.11 | 3 | 3 | 4.17 | 1 | 1 | | | | 4.17 | 1 | 1 | 4.17 | 1 | 1 | 8.68 | 2 | 2 | 57.99 | 12 | 33 | 288 | 31.7 | 7.88 |
| P27144 | 8.52 | 1 | 2 | 2 | 4 | | | | | | | | | | | | | | | | | | | | | | | | | | | | 8.52 | 2 | 4 | 223 | 25.3 | 8.40 |
| P27216 | 12.34 | 1 | 3 | 3 | 6 | 3.48 | 1 | 1 | 3.48 | 1 | 1 | | | | | | | 0.00 | | | | | | | | | | | | | | | 12.34 | 3 | 4 | 316 | 35.4 | 5.60 |
| P27348 | 70.20 | 1 | 19 | 26 | 155 | 32.24 | 6 | 9 | 33.06 | 6 | 9 | 37.55 | 8 | 9 | 39.18 | 8 | 9 | 26.12 | 5 | 6 | 4.49 | 2 | 3 | 23.67 | 6 | 7 | 15.92 | 4 | 5 | 42.04 | 11 | 17 | 62.45 | 19 | 81 | 245 | 27.7 | 4.78 |
| P27361 | 31.13 | 10 | 4 | 9 | 28 | | | | | | | | | | 2.11 | 1 | 1 | | | | | | | | | | | | | | | | 31.13 | 9 | 27 | 379 | 43.1 | 6.74 |
| P27449 | 31.61 | 1 | 2 | 2 | 5 | | | | | | | | | | | | | | | | | | | | | | | | | | | | 31.61 | 2 | 5 | 155 | 15.7 | 8.44 |
| P27487 | 48.83 | 1 | 44 | 44 | 225 | 13.19 | 10 | 10 | 12.01 | 9 | 9 | 39.95 | 32 | 44 | | | | | | | | | | | | | | | | | | | 46.48 | 39 | 162 | 766 | 88.2 | 6.04 |
| P27635 | 51.40 | 1 | 7 | 12 | 40 | 15.42 | 2 | 2 | 2.80 | 1 | 1 | 6.07 | 1 | 1 | 8.88 | 2 | 2 | 4.67 | 1 | 1 | | | | | | | | | | 10.75 | 2 | 2 | 47.20 | 11 | 31 | 214 | 24.6 | 10.08 |
| P27708 | 15.87 | 1 | 25 | 26 | 42 | | | | | | | 0.58 | 1 | 1 | 0.58 | 1 | 1 | | | | | | | | | | 0.58 | 1 | 1 | 0.58 | 1 | 1 | 15.87 | 26 | 38 | 2225 | 242.8 | 6.46 |
| P27797 | 88.25 | 1 | 37 | 37 | 267 | 67.15 | 18 | 32 | 75.30 | 26 | 44 | 73.62 | 25 | 37 | 31.41 | 12 | 16 | 48.44 | 20 | 26 | 6.24 | 1 | 2 | 26.86 | 10 | 11 | 31.18 | 14 | 17 | 38.13 | 15 | 19 | 61.63 | 19 | 63 | 417 | 48.1 | 4.44 |
| P27824 | 53.89 | 1 | 34 | 35 | 221 | 40.20 | 20 | 25 | 31.25 | 18 | 23 | 33.11 | 15 | 18 | 18.92 | 11 | 12 | 16.22 | 8 | 9 | | | | 10.30 | 5 | 6 | | | | 11.66 | 5 | 6 | 44.09 | 28 | 122 | 592 | 67.5 | 4.60 |
| P28062 | 11.96 | 1 | 3 | 3 | 3 | | | | 5.07 | 1 | 1 | | | | | | | | | | | | | | | | | | | | | | 6.88 | 2 | 2 | 276 | 30.3 | 7.43 |
| P28066 | 57.26 | 1 | 14 | 14 | 88 | 26.97 | 5 | 8 | 39.00 | 7 | 8 | 39.00 | 7 | 9 | 42.74 | 7 | 8 | 16.60 | 3 | 3 | | | | 40.25 | 8 | 9 | 30.71 | 5 | 10 | 41.91 | 6 | 11 | 46.47 | 10 | 22 | 241 | 26.4 | 4.79 |
| P28070 | 37.88 | 1 | 6 | 6 | 24 | 25.00 | 3 | 4 | 34.47 | 5 | 6 | 19.70 | 3 | 4 | | | | | | | | | | | | | | | | | | | 28.79 | 5 | 10 | 264 | 29.2 | 5.97 |
| P28072 | 55.23 | 1 | 10 | 10 | 43 | 25.10 | 3 | 3 | 12.97 | 3 | 3 | 36.40 | 5 | 6 | 8.37 | 2 | 2 | 4.18 | 1 | 1 | | | | 12.97 | 3 | 3 | 17.57 | 5 | 5 | 12.97 | 3 | 3 | 48.12 | 8 | 17 | 239 | 25.3 | 4.92 |
| P28074 | 61.22 | 2 | 15 | 15 | 50 | 32.32 | 7 | 7 | 26.62 | 7 | 7 | 33.46 | 7 | 9 | 4.56 | 1 | 1 | 4.56 | 1 | 1 | | | | | | | 4.56 | 1 | 1 | | | | 60.08 | 13 | 24 | 263 | 28.5 | 6.92 |
| P28161 | 61.01 | 1 | 6 | 13 | 32 | 4.13 | 1 | 1 | 4.13 | 1 | 1 | 4.13 | 1 | 1 | 4.13 | 1 | 1 | 4.13 | 1 | 1 | | | | 4.13 | 1 | 1 | 4.13 | 1 | 1 | 27.06 | 5 | 5 | 51.83 | 10 | 20 | 218 | 25.7 | 6.37 |
| P28288 | 21.40 | 5 | 10 | 10 | 16 | | | | | | | | | | | | | | | | | | | | | | | | | | | | 21.40 | 10 | 16 | 659 | 75.4 | 9.36 |
| P28331 | 53.37 | 1 | 32 | 32 | 72 | 16.23 | 8 | 8 | 11.14 | 7 | 7 | 11.55 | 7 | 7 | | | | | | | | | | | | | | | | | | | 50.21 | 30 | 50 | 727 | 79.4 | 6.23 |
| P28482 | 35.28 | 10 | 7 | 12 | 33 | | | | | | | | | | 2.22 | 1 | 1 | | | | | | | | | | | | | | | | 35.28 | 12 | 32 | 360 | 41.4 | 6.98 |
| P28676 | 35.48 | 1 | 8 | 8 | 24 | 30.41 | 7 | 8 | | | | 7.83 | 2 | 2 | 19.82 | 4 | 5 | | | | | | | | | | | | | | | | 22.12 | 4 | 9 | 217 | 24.0 | 5.21 |
| P28838 | 66.28 | 1 | 31 | 31 | 166 | 47.21 | 18 | 22 | 59.54 | 25 | 33 | 50.48 | 19 | 27 | 8.67 | 4 | 4 | 25.05 | 11 | 11 | | | | | | | 5.20 | 2 | 2 | 7.51 | 3 | 3 | 56.07 | 25 | 64 | 519 | 56.1 | 7.93 |
| P28907 | 26.00 | 1 | 9 | 9 | 30 | 16.67 | 5 | 5 | 6.33 | 2 | 2 | 14.00 | 4 | 4 | 3.33 | 1 | 1 | 3.33 | 1 | 1 | | | | | | | | | | | | | 22.00 | 8 | 17 | 300 | 34.3 | 7.66 |
| P29144 | 61.89 | 1 | 70 | 70 | 205 | 20.10 | 17 | 18 | 16.41 | 18 | 21 | 35.39 | 34 | 43 | 1.84 | 2 | 2 | 5.44 | 5 | 5 | | | | | | | | | | | | | 53.40 | 58 | 116 | 1249 | 138.3 | 6.32 |
| P29218 | 49.46 | 1 | 14 | 14 | 50 | 43.68 | 10 | 10 | 37.18 | 8 | 9 | 44.77 | 10 | 11 | 12.64 | 3 | 3 | | | | | | | | | | | | | | | | 34.30 | 9 | 17 | 277 | 30.2 | 5.26 |
| P29350 | 9.92 | 1 | 5 | 5 | 8 | | | | | | | | | | | | | | | | | | | | | | 1.51 | 1 | 1 | 4.87 | 3 | 3 | 5.38 | 2 | 4 | 595 | 67.5 | 7.78 |
| P29373 | 58.70 | 1 | 6 | 6 | 7 | | | | | | | | | | | | | | | | | | | | | | 8.70 | 1 | 1 | | | | 50.00 | 5 | 6 | 138 | 15.7 | 5.40 |
| P29401 | 22.31 | 1 | 9 | 10 | 18 | 0.00 | | | 1.44 | 1 | 1 | 0.96 | 1 | 1 | | | | | | | | | | | | | | | | | | | 22.31 | 10 | 16 | 623 | 67.8 | 7.66 |
| P29508 | 20.51 | 2 | 7 | 7 | 8 | | | | | | | 18.46 | 6 | 6 | | | | | | | | | | | | | | | | 5.13 | 2 | 2 | | | | 390 | 44.5 | 6.81 |
| P29692 | 74.38 | 1 | 16 | 20 | 95 | 34.88 | 9 | 9 | 39.15 | 10 | 11 | 12.46 | 3 | 3 | 29.89 | 6 | 6 | 30.60 | 8 | 9 | | | | 11.74 | 2 | 2 | 34.16 | 7 | 7 | 17.44 | 4 | 4 | 37.01 | 9 | 44 | 281 | 31.1 | 5.01 |
| P29803 | 64.43 | 1 | 20 | 24 | 85 | 31.96 | 9 | 9 | 31.44 | 10 | 15 | 22.16 | 9 | 10 | 5.93 | 2 | 2 | 9.28 | 4 | 4 | | | | 5.93 | 2 | 2 | 0.00 | | | | | | 52.58 | 20 | 43 | 388 | 42.9 | 8.46 |
| P29966 | 33.73 | 2 | 6 | 6 | 23 | 20.78 | 4 | 4 | 9.64 | 2 | 2 | 15.06 | 3 | 3 | | | | | | | | | | | | | | | | | | | 29.22 | 5 | 14 | 332 | 31.5 | 4.45 |
| P29992 | 37.05 | 2 | 5 | 10 | 27 | 4.18 | 1 | 1 | | | | | | | | | | | | | | | | | | | | | | | | | 37.05 | 10 | 26 | 359 | 42.1 | 5.69 |
| P30038 | 21.85 | 1 | 8 | 8 | 12 | | | | 2.66 | 1 | 1 | | | | | | | | | | | | | | | | | | | | | | 21.85 | 8 | 11 | 563 | 61.7 | 8.07 |
| P30040 | 42.91 | 1 | 10 | 10 | 40 | 42.91 | 10 | 14 | 33.72 | 9 | 9 | 17.24 | 4 | 4 | 6.90 | 2 | 2 | 4.60 | 1 | 1 | | | | | | | | | | 4.60 | 1 | 1 | 36.78 | 7 | 9 | 261 | 29.0 | 7.31 |
| P30041 | 73.66 | 1 | 24 | 24 | 184 | 63.84 | 16 | 23 | 64.73 | 12 | 16 | 70.54 | 13 | 18 | 52.23 | 10 | 15 | 9.38 | 1 | 2 | | | | 41.07 | 9 | 11 | 9.82 | 2 | 2 | 43.75 | 6 | 9 | 71.43 | 17 | 88 | 224 | 25.0 | 6.38 |
| P30042 | 36.94 | 1 | 7 | 7 | 31 | 22.39 | 4 | 7 | 22.39 | 4 | 6 | 16.79 | 3 | 4 | 13.06 | 2 | 2 | | | | | | | | | | | | | | | | 36.94 | 7 | 12 | 268 | 28.2 | 8.27 |
| P30043 | 76.70 | 2 | 13 | 13 | 89 | 23.30 | 4 | 4 | 4.85 | 1 | 1 | 14.08 | 2 | 2 | 35.44 | 5 | 8 | 9.22 | 2 | 2 | | | | 36.89 | 5 | 9 | 33.50 | 5 | 7 | 56.31 | 7 | 11 | 65.53 | 9 | 45 | 206 | 22.1 | 7.65 |
| P30044 | 57.48 | 1 | 13 | 13 | 90 | 45.33 | 7 | 8 | 36.92 | 7 | 10 | 37.85 | 7 | 8 | 29.44 | 5 | 8 | 15.89 | 3 | 5 | | | | 29.44 | 5 | 8 | 12.15 | 2 | 6 | 21.96 | 5 | 9 | 56.54 | 12 | 28 | 214 | 22.1 | 8.70 |
| P30046 | 63.56 | 2 | 9 | 9 | 24 | 36.44 | 3 | 4 | 27.12 | 2 | 3 | | | | 33.90 | 3 | 3 | | | | | | | | | | | | | 23.73 | 2 | 2 | 63.56 | 8 | 12 | 118 | 12.7 | 7.30 |
| P30048 | 24.22 | 1 | 6 | 6 | 12 | 3.13 | 1 | 1 | | | | 9.77 | 2 | 3 | | | | | | | | | | | | | | | | 2.73 | 1 | 1 | 18.36 | 4 | 7 | 256 | 27.7 | 7.78 |
| P30049 | 17.26 | 2 | 3 | 3 | 8 | 13.69 | 2 | 2 | 13.69 | 2 | 2 | 5.36 | 1 | 1 | | | | | | | | | | | | | | | | | | | 17.26 | 3 | 3 | 168 | 17.5 | 5.49 |
| P30050 | 60.00 | 1 | 8 | 8 | 25 | 24.85 | 3 | 3 | 24.85 | 3 | 3 | 15.15 | 2 | 2 | | | | 4.24 | 1 | 1 | | | | | | | | | | | | | 60.00 | 7 | 16 | 165 | 17.8 | 9.42 |
| P30084 | 74.14 | 1 | 17 | 17 | 70 | 33.10 | 7 | 8 | 37.24 | 8 | 10 | 28.97 | 6 | 7 | 13.45 | 3 | 4 | 6.21 | 2 | 2 | | | | | | | | | | 30.69 | 6 | 9 | 68.62 | 14 | 30 | 290 | 31.4 | 8.07 |
| P30085 | 65.31 | 1 | 16 | 16 | 64 | 12.76 | 2 | 2 | 11.22 | 2 | 2 | 6.12 | 1 | 1 | 28.57 | 5 | 6 | 12.76 | 2 | 2 | | | | 21.94 | 4 | 5 | 13.27 | 3 | 4 | 56.63 | 11 | 15 | 61.22 | 11 | 27 | 196 | 22.2 | 5.57 |
| P30086 | 85.03 | 2 | 14 | 14 | 97 | 37.97 | 5 | 6 | 48.13 | 5 | 6 | 37.97 | 5 | 7 | 60.96 | 7 | 11 | 53.48 | 5 | 6 | | | | 50.80 | 9 | 12 | 36.90 | 5 | 8 | 80.75 | 9 | 12 | 73.80 | 11 | 29 | 187 | 21.0 | 7.53 |
| P30101 | 75.45 | 1 | 50 | 50 | 382 | 56.44 | 23 | 36 | 50.89 | 23 | 33 | 52.67 | 25 | 39 | 36.63 | 17 | 19 | 59.60 | 31 | 40 | | | | 28.32 | 14 | 18 | 39.80 | 21 | 29 | 56.63 | 29 | 40 | 68.51 | 35 | 128 | 505 | 56.7 | 6.35 |
| P30153 | 53.31 | 1 | 22 | 28 | 143 | 11.38 | 5 | 5 | 8.15 | 4 | 4 | 8.66 | 4 | 4 | 18.17 | 8 | 8 | | | | | | | 5.09 | 2 | 2 | 1.70 | 1 | 1 | 14.26 | 7 | 8 | 53.14 | 27 | 111 | 589 | 65.3 | 5.11 |
| P30154 | 49.25 | 1 | 17 | 23 | 77 | | | | | | | | | | | | | | | | | | | | | | | | | 4.16 | 2 | 2 | 49.25 | 23 | 79 | 601 | 66.2 | 4.94 |
| P30405 | 19.32 | 3 | 3 | 3 | 6 | 6.28 | 2 | 2 | | | | | | | | | | | | | | | | | | | | | | | | | 19.32 | 3 | 4 | 207 | 22.0 | 9.38 |
| P30419 | 4.64 | 1 | 2 | 2 | 2 | | | | | | | 2.62 | 1 | 1 | | | | | | | | | | | | | | | | | | | 2.02 | 1 | 1 | 496 | 56.8 | 7.80 |
| P30519 | 63.29 | 1 | 15 | 15 | 32 | 13.29 | 3 | 3 | 15.51 | 3 | 3 | | | | | | | 9.18 | 2 | 2 | | | | | | | | | | 5.06 | 1 | 1 | 58.86 | 13 | 23 | 316 | 36.0 | 5.41 |
| P30520 | 18.42 | 1 | 6 | 6 | 10 | | | | | | | | | | | | | | | | | | | | | | | | | | | | 18.42 | 6 | 10 | 456 | 50.1 | 6.55 |
| P30533 | 4.20 | 1 | 2 | 2 | 5 | 3.36 | 1 | 1 | | | | 3.36 | 1 | 1 | 3.36 | 1 | 1 | | | | | | | 3.36 | 1 | 1 | | | | 4.20 | 1 | 1 | | | | 357 | 41.4 | 8.78 |
| P30536 | 18.34 | 1 | 2 | 2 | 4 | 13.61 | 1 | 1 | | | | | | | | | | | | | | | | | | | | | | | | | 18.34 | 2 | 3 | 169 | 18.8 | 9.36 |
| P30622 | 5.84 | 1 | 7 | 7 | 7 | | | | | | | | | | | | | 0.00 | | | | | | | | | | | | 0.00 | | | 5.84 | 7 | 7 | 1438 | 162.1 | 5.36 |
| P30626 | 52.02 | 1 | 9 | 9 | 13 | 21.21 | 4 | 4 | 5.56 | 1 | 1 | | | | | | | | | | | | | | | | | | | | | | 42.42 | 7 | 8 | 198 | 21.7 | 5.59 |
| P30711 | 55.83 | 2 | 12 | 14 | 43 | 2.50 | 1 | 1 | 2.50 | 1 | 1 | 2.50 | 1 | 1 | | | | | | | | | | | | | | | | | | | 55.83 | 14 | 40 | 240 | 27.3 | 7.49 |
| P30740 | 16.89 | 1 | 6 | 6 | 11 | 5.28 | 2 | 2 | | | | 14.25 | 5 | 5 | 2.64 | 1 | 1 | | | | | | | | | | | | | 2.64 | 1 | 1 | 2.64 | 1 | 2 | 379 | 42.7 | 6.28 |
| P30876 | 1.53 | 1 | 2 | 2 | 5 | | | | | | | | | | | | | | | | | | | | | | | | | | | | 1.53 | 2 | 5 | 1174 | 133.8 | 6.87 |
| P31040 | 66.72 | 1 | 32 | 32 | 200 | 38.86 | 16 | 24 | 31.02 | 13 | 17 | 42.62 | 21 | 28 | | | | 17.17 | 8 | 8 | | | | | | | | | | | | | 66.42 | 31 | 123 | 664 | 72.6 | 7.39 |
| Q96HN2 | 15.22 | 2 | 1 | 11 | 34 | | | | | | | 1.31 | 1 | 1 | | | | | | | | | | | | | | | | | | | 15.22 | 10 | 33 | 611 | 66.7 | 7.36 |
| P31146 | 22.34 | 1 | 7 | 7 | 11 | 9.54 | 2 | 2 | | | | 12.80 | 5 | 5 | 5.42 | 2 | 3 | | | | | | | | | | | | | 1.74 | 1 | 1 | | | | 461 | 51.0 | 6.68 |
| P31150 | 63.09 | 2 | 15 | 24 | 105 | 37.58 | 13 | 17 | 18.79 | 7 | 9 | 22.82 | 8 | 9 | 2.24 | 1 | 1 | 1.79 | 1 | 1 | | | | 5.37 | 2 | 2 | | | | 3.80 | 2 | 2 | 45.64 | 17 | 64 | 447 | 50.6 | 5.14 |
| P31151 | 54.46 | 2 | 5 | 5 | 10 | 30.69 | 2 | 3 | 22.77 | 2 | 2 | 34.65 | 3 | 4 | | | | | | | | | | | | | 11.88 | 1 | 1 | | | | | | | 101 | 11.5 | 6.77 |
| P31153 | 37.22 | 1 | 9 | 12 | 29 | | | | | | | | | | 8.10 | 2 | 2 | | | | | | | | | | 8.10 | 2 | 2 | 6.08 | 2 | 2 | 33.92 | 11 | 23 | 395 | 43.6 | 6.48 |
| P31689 | 38.79 | 2 | 12 | 14 | 40 | 0.00 | | | 3.27 | 1 | 1 | 0.00 | | | 6.55 | 2 | 3 | 4.03 | 2 | 2 | | | | 0.00 | | | 10.33 | 4 | 4 | 8.06 | 3 | 3 | 38.54 | 13 | 27 | 397 | 44.8 | 7.08 |
| P31749 | 27.50 | 4 | 8 | 11 | 17 | | | | | | | | | | | | | | | | | | | 1.88 | 1 | 1 | | | | 2.71 | 1 | 1 | 24.79 | 10 | 15 | 480 | 55.7 | 6.07 |
| P31751 | 12.27 | 4 | 2 | 5 | 5 | | | | | | | 0.00 | | | | | | | | | | | | | | | | | | | | | 12.27 | 5 | 5 | 481 | 55.7 | 6.37 |
| P31930 | 58.54 | 1 | 20 | 21 | 100 | 25.00 | 9 | 11 | 24.17 | 10 | 13 | 17.92 | 7 | 8 | 2.50 | 1 | 1 | 2.50 | 1 | 1 | | | | | | | | | | | | | 56.46 | 19 | 66 | 480 | 52.6 | 6.37 |
| P31937 | 49.70 | 1 | 12 | 12 | 42 | 28.57 | 6 | 7 | 28.57 | 6 | 9 | 18.45 | 4 | 4 | 9.52 | 2 | 2 | 9.52 | 2 | 2 | | | | | | | | | | | | | 44.64 | 11 | 18 | 336 | 35.3 | 8.13 |
| P31939 | 57.60 | 1 | 26 | 26 | 70 | 4.90 | 3 | 3 | | | | | | | 9.63 | 5 | 5 | | | | | | | | | | 4.56 | 2 | 2 | 8.28 | 4 | 4 | 57.43 | 25 | 58 | 592 | 64.6 | 6.71 |
| P31942 | 16.76 | 3 | 5 | 5 | 9 | | | | | | | | | | | | | | | | | | | | | | 11.85 | 4 | 4 | 2.60 | 1 | 1 | 10.69 | 3 | 4 | 346 | 36.9 | 6.87 |
| P31943 | 25.17 | 2 | 7 | 9 | 28 | | | | | | | 3.79 | 1 | 1 | 4.90 | 2 | 2 | 3.56 | 1 | 1 | 2.00 | 1 | 1 | 5.79 | 2 | 2 | 9.13 | 3 | 3 | 8.69 | 3 | 4 | 17.59 | 6 | 14 | 449 | 49.2 | 6.30 |
| P31944 | 21.90 | 1 | 4 | 4 | 10 | | | | | | | | | | | | | 17.36 | 3 | 5 | | | | 5.79 | 1 | 1 | 14.88 | 3 | 4 | | | | | | | 242 | 27.7 | 5.58 |
| P31946 | 74.80 | 2 | 10 | 19 | 127 | 31.71 | 6 | 9 | 31.71 | 6 | 9 | 26.42 | 6 | 7 | 43.90 | 9 | 12 | 22.76 | 5 | 7 | 4.07 | 1 | 1 | 34.96 | 8 | 10 | 26.83 | 6 | 7 | 39.02 | 9 | 13 | 67.89 | 14 | 52 | 246 | 28.1 | 4.83 |
| P31947 | 70.97 | 1 | 17 | 22 | 156 | 34.68 | 6 | 10 | 23.79 | 4 | 7 | 37.10 | 7 | 8 | 44.76 | 10 | 13 | 20.97 | 5 | 6 | 3.63 | 1 | 1 | 31.85 | 10 | 14 | 28.23 | 6 | 9 | 50.81 | 12 | 22 | 59.27 | 17 | 66 | 248 | 27.8 | 4.74 |
| P31948 | 60.41 | 1 | 39 | 39 | 165 | 29.47 | 15 | 17 | 28.36 | 15 | 18 | 27.26 | 14 | 16 | 19.89 | 10 | 10 | 21.36 | 13 | 13 | | | | 18.23 | 10 | 10 | 13.26 | 7 | 7 | 26.89 | 15 | 16 | 51.01 | 29 | 58 | 543 | 62.6 | 6.80 |
| P31949 | 52.38 | 1 | 6 | 6 | 27 | 32.38 | 3 | 3 | 15.24 | 1 | 1 | 49.52 | 5 | 6 | 8.57 | 1 | 1 | 8.57 | 1 | 1 | | | | | | | | | | 18.10 | 2 | 2 | 52.38 | 6 | 13 | 105 | 11.7 | 7.12 |
| P31997 | 19.48 | 1 | 4 | 5 | 17 | 12.89 | 3 | 4 | | | | 19.48 | 5 | 8 | 4.58 | 1 | 1 | | | | | | | | | | | | | | | | 7.16 | 2 | 4 | 349 | 38.1 | 7.39 |
| P32119 | 72.73 | 2 | 14 | 15 | 93 | 32.32 | 7 | 7 | 32.32 | 6 | 6 | 32.32 | 8 | 8 | 23.74 | 5 | 5 | 9.09 | 2 | 2 | | | | 23.23 | 5 | 6 | 14.14 | 3 | 3 | 40.40 | 7 | 7 | 72.73 | 12 | 49 | 198 | 21.9 | 5.97 |
| P32189 | 63.15 | 1 | 2 | 27 | 96 | 21.29 | 8 | 13 | 20.04 | 10 | 12 | 13.60 | 6 | 8 | 2.33 | 1 | 1 | | | | | | | | | | | | | | | | 63.15 | 27 | 62 | 559 | 61.2 | 6.54 |
| P32856 | 30.21 | 1 | 6 | 6 | 9 | | | | | | | | | | | | | | | | | | | | | | | | | | | | 30.21 | 6 | 9 | 288 | 33.3 | 6.28 |
| P32929 | 10.62 | 1 | 4 | 4 | 4 | | | | | | | | | | | | | | | | | | | | | | | | | | | | 10.62 | 4 | 4 | 405 | 44.5 | 6.70 |
| P32942 | 5.30 | 1 | 3 | 3 | 3 | | | | | | | 5.30 | 3 | 3 | | | | | | | | | | | | | | | | | | | | | | 547 | 59.5 | 5.43 |
| P32969 | 46.35 | 1 | 7 | 7 | 18 | | | | 9.90 | 1 | 1 | | | | | | | | | | | | | | | | | | | | | | 46.35 | 7 | 17 | 192 | 21.8 | 9.95 |
| P33121 | 54.44 | 1 | 36 | 39 | 326 | 25.64 | 17 | 20 | 30.66 | 21 | 30 | 32.23 | 20 | 23 | 17.05 | 12 | 13 | 14.61 | 9 | 9 | | | | 3.15 | 2 | 3 | 1.58 | 1 | 1 | 4.15 | 4 | 4 | 54.30 | 35 | 223 | 698 | 77.9 | 7.15 |
| P33176 | 46.42 | 1 | 29 | 41 | 111 | 2.28 | 2 | 2 | 4.26 | 3 | 3 | 4.05 | 3 | 3 | 4.26 | 3 | 3 | 2.91 | 2 | 2 | | | | 3.53 | 3 | 3 | | | | 16.72 | 14 | 16 | 39.15 | 32 | 79 | 963 | 109.6 | 6.51 |
| P33316 | 36.51 | 1 | 7 | 7 | 19 | 9.92 | 2 | 2 | 28.57 | 5 | 6 | 3.97 | 1 | 1 | 3.97 | 1 | 1 | | | | | | | | | | 3.97 | 1 | 1 | | | | 25.79 | 5 | 8 | 252 | 26.5 | 9.36 |
| P34096 | 34.69 | 1 | 6 | 6 | 12 | 28.57 | 3 | 3 | 9.52 | 1 | 1 | 7.48 | 1 | 1 | | | | 7.48 | 1 | 1 | | | | | | | | | | | | | 34.01 | 5 | 6 | 147 | 16.8 | 9.03 |
| P34810 | 8.47 | 1 | 2 | 2 | 3 | | | | | | | | | | | | | | | | | | | | | | | | | | | | 8.47 | 2 | 3 | 354 | 37.4 | 8.87 |
| P34896 | 40.58 | 2 | 15 | 15 | 37 | 5.18 | 2 | 2 | 3.73 | 1 | 1 | 3.73 | 1 | 1 | 1.86 | 1 | 1 | | | | | | | 1.86 | 1 | 1 | | | | | | | 40.58 | 15 | 31 | 483 | 53.0 | 7.71 |
| P34913 | 55.14 | 1 | 21 | 21 | 98 | 3.42 | 2 | 2 | 1.26 | 1 | 1 | 12.97 | 7 | 7 | 9.73 | 5 | 5 | | | | | | | 3.24 | 2 | 2 | | | | 8.83 | 4 | 5 | 53.87 | 20 | 76 | 555 | 62.6 | 6.28 |
| P34931 | 72.39 | 1 | 25 | 44 | 526 | 49.61 | 25 | 38 | 44.15 | 23 | 40 | 55.07 | 29 | 53 | 31.36 | 18 | 22 | 31.05 | 20 | 27 | 5.46 | 3 | 3 | 28.55 | 17 | 22 | 32.76 | 22 | 34 | 30.89 | 19 | 25 | 69.89 | 38 | 264 | 641 | 70.3 | 6.02 |
| P34932 | 53.57 | 1 | 35 | 39 | 177 | 24.29 | 14 | 18 | 19.88 | 13 | 19 | 33.81 | 22 | 30 | 4.29 | 3 | 3 | 2.98 | 2 | 3 | | | | 5.60 | 3 | 3 | 0.95 | 1 | 1 | 4.64 | 3 | 4 | 47.74 | 33 | 96 | 840 | 94.3 | 5.19 |
| P34949 | 47.28 | 1 | 17 | 17 | 39 | 14.66 | 4 | 6 | 6.38 | 2 | 2 | | | | 8.04 | 2 | 2 | | | | | | | | | | 3.78 | 1 | 2 | 8.51 | 3 | 3 | 42.32 | 15 | 20 | 423 | 46.6 | 5.95 |
| P35030 | 6.58 | 1 | 2 | 2 | 10 | | | | | | | | | | | | | | | | 4.28 | 1 | 1 | | | | | | | | | | 6.58 | 2 | 9 | 304 | 32.5 | 7.49 |
| P35052 | 39.07 | 1 | 17 | 17 | 35 | 2.15 | 1 | 1 | | | | | | | | | | | | | | | | | | | | | | | | | 36.92 | 16 | 34 | 558 | 61.6 | 7.30 |
| P35080 | 54.29 | 1 | 6 | 6 | 22 | 10.00 | 1 | 1 | 10.00 | 1 | 1 | 10.00 | 1 | 1 | 15.71 | 2 | 2 | 10.00 | 1 | 1 | | | | 10.00 | 1 | 1 | 10.00 | 1 | 1 | 15.71 | 2 | 2 | 54.29 | 6 | 12 | 140 | 15.0 | 6.99 |
| P35221 | 6.29 | 2 | 4 | 4 | 6 | | | | | | | | | | | | | 1.32 | 1 | 1 | | | | | | | 0.00 | | | 1.32 | 1 | 1 | 4.97 | 3 | 4 | 906 | 100.0 | 6.29 |
| P35222 | 17.80 | 1 | 7 | 11 | 32 | | | | | | | | | | | | | 0.90 | 1 | 1 | | | | 0.90 | 1 | 1 | | | | 2.43 | 2 | 2 | 15.36 | 9 | 28 | 781 | 85.4 | 5.86 |
| P35232 | 76.10 | 1 | 16 | 16 | 36 | 19.49 | 4 | 4 | 26.84 | 6 | 6 | 18.38 | 4 | 4 | | | | | | | | | | | | | | | | | | | 72.43 | 14 | 22 | 272 | 29.8 | 5.76 |
| P35237 | 62.23 | 3 | 21 | 21 | 116 | 33.78 | 10 | 14 | 23.14 | 6 | 6 | 30.59 | 10 | 12 | 26.06 | 8 | 9 | | | | | | | 9.57 | 3 | 3 | 7.18 | 2 | 3 | 15.69 | 5 | 5 | 59.84 | 19 | 64 | 376 | 42.6 | 5.27 |
| P35268 | 57.03 | 1 | 6 | 6 | 16 | 20.31 | 1 | 1 | 39.06 | 3 | 3 | 10.16 | 1 | 1 | | | | | | | | | | | | | | | | 10.16 | 1 | 1 | 57.03 | 6 | 10 | 128 | 14.8 | 9.19 |
| P35270 | 33.33 | 1 | 6 | 6 | 20 | | | | | | | | | | 5.75 | 1 | 1 | | | | | | | | | | | | | 5.75 | 1 | 1 | 33.33 | 6 | 18 | 261 | 28.0 | 8.05 |
| P35443 | 9.89 | 1 | 6 | 6 | 11 | | | | | | | | | | | | | | | | | | | | | | | | | | | | 9.89 | 6 | 11 | 961 | 105.8 | 4.68 |
| P35527 | 69.02 | 2 | 40 | 41 | 433 | | | | 22.63 | 9 | 9 | 34.99 | 16 | 20 | 38.84 | 20 | 24 | 36.12 | 19 | 23 | 2.57 | 1 | 1 | 42.86 | 22 | 34 | 49.44 | 27 | 44 | 26.00 | 12 | 13 | 59.55 | 34 | 265 | 623 | 62.0 | 5.24 |
| P35573 | 29.57 | 1 | 36 | 36 | 79 | | | | 0.00 | | | 1.76 | 2 | 2 | 0.72 | 1 | 1 | | | | | | | 0.91 | 1 | 1 | | | | 1.50 | 2 | 2 | 27.42 | 33 | 69 | 1532 | 174.7 | 6.76 |
| P35579 | 68.01 | 8 | 140 | 164 | 918 | 32.50 | 48 | 64 | 16.79 | 25 | 31 | 41.63 | 65 | 83 | 36.84 | 61 | 73 | 17.91 | 32 | 37 | 0.66 | 1 | 1 | 0.71 | 1 | 1 | 0.61 | 1 | 1 | 12.91 | 18 | 19 | 62.76 | 139 | 608 | 1960 | 226.4 | 5.60 |
| P35580 | 52.13 | 7 | 74 | 95 | 268 | 2.94 | 5 | 5 | 2.38 | 3 | 3 | 6.58 | 10 | 10 | 6.83 | 10 | 10 | 3.09 | 5 | 5 | | | | | | | | | | 1.87 | 3 | 3 | 50.66 | 91 | 232 | 1976 | 228.9 | 5.54 |
| P35606 | 33.55 | 1 | 25 | 25 | 92 | 0.00 | | | | | | 15.67 | 10 | 11 | 0.00 | | | 1.10 | 1 | 1 | | | | | | | | | | | | | 31.57 | 24 | 80 | 906 | 102.4 | 5.27 |
| P35613 | 31.43 | 1 | 8 | 8 | 38 | 20.52 | 4 | 6 | 3.64 | 1 | 1 | 23.12 | 5 | 6 | | | | | | | | | | | | | | | | | | | 23.38 | 7 | 29 | 385 | 42.2 | 5.66 |
| P35625 | 48.82 | 1 | 9 | 9 | 18 | | | | 13.74 | 3 | 3 | 3.79 | 1 | 1 | | | | | | | | | | | | | | | | | | | 48.82 | 9 | 14 | 211 | 24.1 | 8.72 |
| P35626 | 27.33 | 1 | 12 | 17 | 23 | | | | | | | | | | | | | | | | | | | | | | | | | | | | 27.33 | 17 | 23 | 688 | 79.7 | 7.62 |
| P35658 | 6.65 | 1 | 11 | 11 | 21 | | | | 0.81 | 1 | 1 | | | | | | | | | | | | | | | | | | | | | | 6.65 | 11 | 20 | 2090 | 213.5 | 7.47 |
| P35663 | 12.90 | 1 | 6 | 6 | 8 | | | | | | | 11.06 | 5 | 6 | | | | | | | | | | | | | | | | | | | 1.84 | 1 | 2 | 651 | 74.2 | 9.67 |
| P35670 | 3.89 | 1 | 6 | 6 | 11 | | | | | | | | | | | | | | | | | | | | | | | | | | | | 3.89 | 6 | 11 | 1465 | 157.2 | 6.70 |
| P35813 | 52.62 | 1 | 14 | 16 | 29 | 2.88 | 1 | 1 | 8.38 | 3 | 3 | | | | 3.14 | 1 | 1 | 3.14 | 1 | 1 | | | | | | | | | | 3.14 | 1 | 1 | 52.62 | 16 | 22 | 382 | 42.4 | 5.36 |
| P35908 | 80.91 | 4 | 53 | 67 | 543 | 4.85 | 3 | 3 | 33.96 | 17 | 17 | 54.62 | 24 | 32 | 27.23 | 13 | 14 | 40.53 | 22 | 23 | | | | 48.36 | 30 | 34 | 67.29 | 56 | 95 | 54.62 | 30 | 35 | 70.74 | 40 | 290 | 639 | 65.4 | 8.00 |
| P35914 | 17.23 | 1 | 4 | 5 | 6 | 3.08 | 1 | 1 | | | | | | | | | | | | | | | | | | | | | | 3.08 | 1 | 1 | 14.15 | 4 | 4 | 325 | 34.3 | 8.54 |
| P35998 | 72.52 | 1 | 32 | 32 | 137 | 32.33 | 11 | 12 | 31.18 | 11 | 13 | 16.63 | 6 | 6 | 11.78 | 5 | 6 | 15.01 | 6 | 6 | 2.31 | 1 | 1 | 23.79 | 9 | 9 | 12.24 | 5 | 5 | 25.87 | 10 | 11 | 69.75 | 27 | 68 | 433 | 48.6 | 5.95 |
| P36404 | 50.00 | 1 | 7 | 7 | 10 | | | | 5.98 | 1 | 1 | | | | | | | | | | | | | | | | | | | | | | 50.00 | 7 | 9 | 184 | 20.9 | 6.34 |
| P36405 | 71.98 | 1 | 12 | 12 | 27 | 6.04 | 1 | 1 | 6.04 | 1 | 1 | | | | | | | | | | | | | | | | 4.40 | 1 | 1 | 22.53 | 3 | 3 | 71.43 | 11 | 21 | 182 | 20.4 | 7.24 |
| P36507 | 46.25 | 1 | 12 | 18 | 40 | | | | | | | 3.00 | 1 | 1 | 2.25 | 1 | 1 | | | | | | | | | | | | | 16.00 | 6 | 6 | 37.75 | 14 | 32 | 400 | 44.4 | 6.55 |
| P36542 | 45.30 | 1 | 11 | 11 | 39 | 17.45 | 5 | 6 | 32.21 | 7 | 8 | 17.11 | 5 | 5 | 7.38 | 2 | 2 | | | | | | | | | | | | | 3.69 | 1 | 1 | 33.56 | 10 | 17 | 298 | 33.0 | 9.22 |
| P36543 | 50.88 | 1 | 9 | 14 | 19 | 10.18 | 2 | 3 | 0.00 | | | | | | | | | | | | 5.31 | 1 | 1 | | | | | | | | | | 49.56 | 13 | 15 | 226 | 26.1 | 8.00 |
| P36578 | 55.50 | 1 | 26 | 26 | 74 | 4.45 | 2 | 2 | 3.51 | 2 | 2 | | | | 7.03 | 3 | 3 | 11.24 | 3 | 4 | | | | 2.58 | 1 | 1 | | | | | | | 47.54 | 22 | 62 | 427 | 47.7 | 11.06 |
| P36776 | 38.69 | 2 | 28 | 28 | 46 | 4.90 | 4 | 4 | 10.43 | 6 | 7 | 1.25 | 1 | 1 | | | | | | | | | | | | | | | | | | | 34.72 | 25 | 34 | 959 | 106.4 | 6.39 |
| P36871 | 20.46 | 1 | 11 | 11 | 12 | | | | | | | | | | | | | | | | | | | | | | | | | | | | 20.46 | 11 | 12 | 562 | 61.4 | 6.76 |
| P36873 | 67.49 | 1 | 4 | 19 | 128 | 41.80 | 11 | 14 | 18.58 | 5 | 5 | 32.51 | 9 | 10 | 21.05 | 6 | 6 | 11.15 | 3 | 3 | | | | 14.24 | 4 | 4 | 11.46 | 3 | 3 | 22.60 | 6 | 7 | 60.37 | 17 | 78 | 323 | 37.0 | 6.54 |
| P36915 | 11.37 | 1 | 5 | 5 | 9 | | | | | | | | | | | | | | | | | | | | | | | | | | | | 11.37 | 5 | 9 | 607 | 68.6 | 5.80 |
| P36957 | 27.81 | 1 | 12 | 12 | 49 | 21.19 | 8 | 8 | 18.10 | 9 | 11 | 9.05 | 4 | 5 | 7.28 | 3 | 3 | 2.87 | 1 | 2 | | | | | | | 2.87 | 1 | 1 | | | | 26.05 | 10 | 19 | 453 | 48.7 | 8.95 |
| P36959 | 46.09 | 2 | 10 | 12 | 32 | 17.39 | 3 | 4 | 2.61 | 1 | 2 | 8.99 | 2 | 2 | 3.48 | 1 | 1 | | | | | | | | | | | | | | | | 37.10 | 10 | 23 | 345 | 37.4 | 7.06 |
| P36969 | 78.68 | 1 | 20 | 20 | 217 | 53.81 | 12 | 20 | 55.33 | 12 | 23 | 55.84 | 12 | 22 | 31.47 | 7 | 8 | 38.58 | 9 | 11 | | | | 7.11 | 1 | 1 | 16.24 | 2 | 2 | 26.40 | 4 | 5 | 78.68 | 19 | 129 | 197 | 22.2 | 8.37 |
| P37198 | 23.95 | 1 | 10 | 10 | 50 | 12.07 | 4 | 4 | 11.30 | 4 | 5 | 7.28 | 3 | 3 | 1.92 | 1 | 1 | | | | | | | | | | 5.17 | 2 | 3 | 1.92 | 1 | 1 | 23.95 | 10 | 33 | 522 | 53.2 | 5.31 |
| P37235 | 11.92 | 3 | 2 | 2 | 2 | | | | | | | | | | | | | | | | | | | | | | | | | | | | 11.92 | 2 | 2 | 193 | 22.3 | 5.35 |
| P37802 | 75.38 | 3 | 12 | 12 | 24 | | | | 6.03 | 1 | 1 | | | | 10.55 | 2 | 2 | | | | | | | 15.58 | 3 | 3 | | | | | | | 71.36 | 11 | 18 | 199 | 22.4 | 8.25 |
| P37837 | 39.17 | 1 | 15 | 15 | 33 | 12.46 | 4 | 5 | 11.57 | 4 | 4 | 13.65 | 4 | 4 | | | | | | | | | | | | | | | | | | | 28.49 | 12 | 20 | 337 | 37.5 | 6.81 |
| P38117 | 59.61 | 1 | 16 | 16 | 40 | 30.59 | 8 | 8 | 37.25 | 10 | 10 | 19.22 | 5 | 5 | 4.71 | 1 | 1 | | | | | | | | | | | | | 3.92 | 1 | 1 | 41.18 | 11 | 15 | 255 | 27.8 | 8.10 |
| Q14644 | 1.08 | 1 | 1 | 1 | 31 | 1.08 | 1 | 2 | 1.08 | 1 | 1 | 1.08 | 1 | 1 | 1.08 | 1 | 2 | 1.08 | 1 | 1 | | | | 1.08 | 1 | 2 | 1.08 | 1 | 2 | 1.08 | 1 | 2 | 1.08 | 1 | 18 | 834 | 95.6 | 7.15 |
| P38432 | 6.94 | 1 | 3 | 3 | 3 | | | | | | | | | | | | | 2.43 | 1 | 1 | | | | | | | | | | 2.78 | 1 | 1 | 1.74 | 1 | 1 | 576 | 62.6 | 9.07 |
| P38567 | 28.29 | 1 | 12 | 12 | 62 | 18.47 | 7 | 8 | 9.82 | 5 | 7 | 8.45 | 4 | 4 | | | | 7.27 | 3 | 3 | | | | | | | | | | 1.77 | 1 | 1 | 28.09 | 11 | 39 | 509 | 57.8 | 7.01 |
| P38606 | 45.71 | 1 | 25 | 25 | 87 | 8.75 | 4 | 5 | 10.21 | 4 | 6 | 16.05 | 7 | 7 | 5.02 | 2 | 2 | 2.92 | 2 | 2 | | | | 3.89 | 2 | 2 | | | | 6.32 | 3 | 3 | 44.41 | 22 | 60 | 617 | 68.3 | 5.52 |
| P38646 | 53.61 | 1 | 40 | 41 | 141 | 27.39 | 15 | 15 | 30.19 | 17 | 18 | 32.70 | 20 | 23 | 5.60 | 3 | 3 | 13.11 | 7 | 7 | | | | 3.98 | 2 | 2 | 21.80 | 12 | 12 | 29.16 | 14 | 16 | 42.12 | 25 | 49 | 679 | 73.6 | 6.16 |
| P38919 | 13.38 | 1 | 3 | 7 | 19 | | | | | | | | | | 3.89 | 1 | 1 | 3.89 | 1 | 1 | | | | | | | 3.89 | 1 | 1 | 6.08 | 2 | 3 | 13.14 | 6 | 13 | 411 | 46.8 | 6.73 |
| P39019 | 48.28 | 1 | 9 | 9 | 23 | 32.41 | 5 | 5 | 33.10 | 5 | 5 | 6.21 | 1 | 1 | 6.21 | 1 | 1 | | | | | | | | | | | | | | | | 48.28 | 9 | 11 | 145 | 16.1 | 10.32 |
| P39023 | 41.94 | 2 | 18 | 18 | 100 | 3.97 | 2 | 2 | 17.12 | 4 | 4 | 7.94 | 3 | 3 | 7.20 | 3 | 3 | 2.98 | 1 | 1 | | | | 2.98 | 1 | 1 | | | | 9.93 | 3 | 3 | 37.97 | 17 | 83 | 403 | 46.1 | 10.18 |
| P39060 | 14.37 | 2 | 21 | 21 | 78 | 7.30 | 9 | 9 | 5.82 | 7 | 7 | 3.36 | 4 | 4 | 1.77 | 2 | 2 | | | | 0.51 | 1 | 1 | | | | | | | | | | 10.66 | 16 | 55 | 1754 | 178.1 | 6.01 |
| P39210 | 17.61 | 1 | 2 | 2 | 6 | 13.07 | 1 | 1 | 13.07 | 1 | 1 | | | | | | | | | | | | | | | | | | | | | | 17.61 | 2 | 4 | 176 | 19.7 | 9.47 |
| P39656 | 56.80 | 1 | 18 | 18 | 71 | 21.05 | 6 | 7 | 29.82 | 8 | 11 | 29.61 | 9 | 9 | 1.97 | 1 | 1 | | | | | | | | | | | | | | | | 55.04 | 16 | 43 | 456 | 50.8 | 6.55 |
| P40121 | 37.36 | 1 | 13 | 13 | 38 | 6.90 | 2 | 2 | | | | 8.33 | 3 | 3 | 17.24 | 5 | 6 | | | | | | | 6.90 | 2 | 2 | 17.53 | 5 | 7 | 14.66 | 4 | 5 | 26.15 | 9 | 13 | 348 | 38.5 | 6.19 |
| P40199 | 9.88 | 1 | 2 | 2 | 5 | 6.10 | 1 | 1 | | | | 9.88 | 2 | 4 | | | | | | | | | | | | | | | | | | | | | | 344 | 37.2 | 5.82 |
| P40227 | 66.10 | 1 | 27 | 32 | 182 | 21.09 | 8 | 10 | 26.74 | 12 | 15 | 17.33 | 8 | 9 | 23.54 | 12 | 12 | 20.53 | 10 | 10 | | | | 12.81 | 7 | 8 | 12.05 | 5 | 5 | 23.92 | 12 | 16 | 55.18 | 26 | 97 | 531 | 58.0 | 6.68 |
| P40429 | 40.39 | 2 | 14 | 14 | 31 | | | | | | | | | | 8.37 | 2 | 2 | | | | | | | 9.85 | 2 | 2 | | | | 14.78 | 3 | 3 | 38.92 | 12 | 24 | 203 | 23.6 | 10.93 |
| P40616 | 77.35 | 2 | 12 | 12 | 34 | | | | | | | 9.39 | 1 | 1 | 12.71 | 2 | 2 | | | | | | | 4.97 | 1 | 1 | | | | | | | 77.35 | 12 | 30 | 181 | 20.4 | 5.72 |
| P40617 | 22.00 | 2 | 3 | 4 | 6 | | | | | | | | | | | | | | | | | | | | | | | | | | | | 22.00 | 4 | 6 | 200 | 22.6 | 9.17 |
| P40763 | 50.52 | 2 | 27 | 27 | 92 | | | | | | | | | | | | | | | | | | | | | | | | | 4.81 | 2 | 2 | 50.52 | 27 | 90 | 770 | 88.0 | 6.30 |
| P40879 | 5.24 | 1 | 4 | 4 | 4 | | | | | | | 1.44 | 1 | 1 | | | | | | | | | | | | | | | | | | | 3.80 | 3 | 3 | 764 | 84.5 | 8.69 |
| P40925 | 35.93 | 1 | 11 | 11 | 39 | 17.07 | 5 | 5 | 17.37 | 5 | 5 | 15.87 | 5 | 5 | 11.68 | 3 | 3 | 5.69 | 2 | 2 | | | | | | | | | | 2.99 | 1 | 1 | 35.93 | 9 | 18 | 334 | 36.4 | 7.36 |
| P40926 | 67.75 | 1 | 20 | 20 | 157 | 54.14 | 15 | 22 | 58.88 | 17 | 25 | 63.61 | 17 | 25 | 34.32 | 9 | 10 | 40.83 | 10 | 11 | | | | | | | 2.66 | 1 | 1 | 4.73 | 1 | 1 | 61.24 | 16 | 62 | 338 | 35.5 | 8.68 |
| P40939 | 66.84 | 1 | 44 | 44 | 318 | 47.71 | 25 | 35 | 50.72 | 29 | 49 | 48.23 | 27 | 41 | 16.51 | 11 | 11 | 17.43 | 11 | 13 | | | | 2.88 | 2 | 2 | 2.75 | 2 | 2 | 11.80 | 7 | 7 | 64.09 | 40 | 158 | 763 | 82.9 | 9.04 |
| P41091 | 48.52 | 2 | 17 | 17 | 41 | | | | | | | 3.39 | 1 | 1 | 3.39 | 1 | 1 | | | | | | | | | | | | | | | | 48.52 | 17 | 39 | 472 | 51.1 | 8.40 |
| P41218 | 25.80 | 1 | 9 | 9 | 17 | 10.57 | 4 | 5 | | | | | | | | | | | | | | | | | | | | | | 4.67 | 2 | 2 | 20.39 | 6 | 10 | 407 | 45.8 | 9.76 |
| P41222 | 46.84 | 2 | 7 | 7 | 30 | 17.37 | 2 | 3 | 17.37 | 2 | 3 | 27.89 | 5 | 6 | 21.05 | 3 | 3 | 19.47 | 4 | 5 | | | | 8.42 | 1 | 1 | 15.79 | 3 | 4 | | | | 36.32 | 4 | 5 | 190 | 21.0 | 7.80 |
| P41240 | 7.78 | 1 | 3 | 3 | 4 | | | | | | | | | | | | | | | | | | | | | | | | | | | | 7.78 | 3 | 4 | 450 | 50.7 | 7.06 |
| P41250 | 39.24 | 1 | 29 | 29 | 94 | 7.31 | 4 | 5 | 5.14 | 4 | 4 | 7.71 | 5 | 5 | 2.30 | 2 | 2 | | | | | | | 2.84 | 2 | 2 | 1.08 | 1 | 1 | 4.87 | 4 | 5 | 36.81 | 25 | 70 | 739 | 83.1 | 7.03 |
| P41252 | 38.75 | 1 | 40 | 40 | 95 | | | | 1.43 | 2 | 2 | 1.51 | 2 | 2 | 1.51 | 2 | 2 | | | | | | | | | | | | | 0.95 | 1 | 1 | 38.75 | 40 | 88 | 1262 | 144.4 | 6.15 |
| P42025 | 48.94 | 1 | 6 | 14 | 33 | 11.70 | 4 | 4 | 9.04 | 3 | 3 | 11.70 | 3 | 3 | 6.65 | 2 | 2 | | | | | | | | | | | | | | | | 43.09 | 12 | 21 | 376 | 42.3 | 6.40 |
| P42126 | 32.12 | 1 | 9 | 9 | 31 | 18.87 | 4 | 5 | 19.87 | 5 | 8 | 15.23 | 4 | 6 | 4.97 | 1 | 1 | | | | | | | | | | | | | | | | 32.12 | 9 | 11 | 302 | 32.8 | 8.54 |
| P42224 | 33.60 | 1 | 22 | 22 | 53 | | | | | | | | | | | | | | | | | | | | | | 0.93 | 1 | 1 | 11.60 | 6 | 6 | 31.33 | 20 | 46 | 750 | 87.3 | 6.05 |
| P42226 | 11.33 | 1 | 7 | 8 | 9 | | | | | | | | | | | | | | | | | | | | | | | | | | | | 11.33 | 8 | 9 | 847 | 94.1 | 6.23 |
| P42338 | 2.90 | 1 | 2 | 2 | 7 | | | | 0.00 | | | | | | | | | | | | | | | | | | | | | | | | 2.90 | 2 | 7 | 1070 | 122.7 | 7.09 |
| P42345 | 4.39 | 2 | 9 | 10 | 16 | | | | | | | | | | | | | | | | | | | | | | | | | | | | 4.39 | 10 | 16 | 2549 | 288.7 | 7.17 |
| P42356 | 1.33 | 1 | 2 | 2 | 3 | | | | | | | | | | | | | | | | | | | | | | | | | | | | 1.33 | 2 | 3 | 2102 | 236.7 | 7.06 |
| P42566 | 6.92 | 1 | 4 | 4 | 7 | 0.00 | | | 0.00 | | | | | | | | | | | | | | | 3.35 | 2 | 2 | 1.56 | 1 | 1 | | | | 5.13 | 3 | 4 | 896 | 98.6 | 4.64 |
| P42685 | 4.16 | 1 | 2 | 2 | 2 | | | | | | | | | | | | | | | | | | | | | | | | | | | | 4.16 | 2 | 2 | 505 | 58.2 | 6.67 |
| P42704 | 22.81 | 1 | 25 | 25 | 33 | | | | 0.00 | | | | | | | | | | | | | | | | | | | | | | | | 22.81 | 25 | 33 | 1394 | 157.8 | 6.13 |
| P42765 | 56.17 | 1 | 18 | 18 | 36 | 17.88 | 5 | 5 | 24.18 | 7 | 8 | 13.85 | 5 | 5 | | | | | | | | | | | | | | | | | | | 41.31 | 11 | 18 | 397 | 41.9 | 8.09 |
| P42766 | 18.70 | 1 | 2 | 2 | 4 | | | | | | | | | | | | | | | | | | | | | | | | | | | | 18.70 | 2 | 4 | 123 | 14.5 | 11.05 |
| P42785 | 29.64 | 1 | 11 | 11 | 23 | 18.15 | 5 | 5 | 9.68 | 4 | 6 | 12.70 | 5 | 5 | 2.22 | 1 | 1 | | | | | | | | | | | | | | | | 7.26 | 3 | 6 | 496 | 55.8 | 7.21 |
| P42858 | 1.34 | 1 | 3 | 3 | 4 | | | | | | | 0.00 | | | | | | | | | | | | | | | | | | | | | 1.34 | 3 | 4 | 3142 | 347.4 | 6.20 |
| P42898 | 10.67 | 1 | 6 | 6 | 7 | | | | | | | | | | | | | | | | | | | | | | | | | | | | 10.67 | 6 | 7 | 656 | 74.5 | 5.34 |
| P43034 | 41.95 | 1 | 16 | 17 | 69 | 1.71 | 1 | 1 | 10.98 | 4 | 4 | 24.88 | 8 | 8 | 11.95 | 4 | 4 | 9.27 | 3 | 3 | | | | 3.90 | 1 | 1 | 17.07 | 5 | 5 | 18.29 | 5 | 5 | 36.83 | 15 | 38 | 410 | 46.6 | 7.37 |
| P43155 | 37.38 | 1 | 23 | 23 | 63 | 12.46 | 7 | 7 | 19.33 | 11 | 11 | 26.52 | 15 | 17 | 1.92 | 1 | 1 | | | | | | | | | | | | | | | | 28.27 | 18 | 27 | 626 | 70.8 | 8.44 |
| P43251 | 4.60 | 1 | 2 | 2 | 7 | | | | | | | | | | | | | | | | | | | | | | | | | | | | 4.60 | 2 | 7 | 543 | 61.1 | 6.25 |
| P43304 | 20.77 | 1 | 11 | 11 | 20 | 3.03 | 1 | 1 | 3.03 | 1 | 1 | 1.24 | 1 | 1 | | | | | | | | | | | | | | | | | | | 20.77 | 11 | 17 | 727 | 80.8 | 7.69 |
| P43307 | 19.93 | 1 | 4 | 4 | 8 | | | | 10.84 | 2 | 2 | 2.80 | 1 | 1 | | | | | | | | | | | | | | | | | | | 11.89 | 3 | 5 | 286 | 32.2 | 4.49 |
| P43358 | 10.09 | 10 | 3 | 3 | 4 | | | | | | | | | | | | | | | | | | | | | | | | | | | | 10.09 | 3 | 4 | 317 | 34.9 | 4.69 |
| P43487 | 43.28 | 2 | 7 | 7 | 22 | 5.47 | 1 | 1 | 5.47 | 1 | 1 | 8.96 | 2 | 2 | | | | 9.95 | 2 | 2 | | | | | | | 4.48 | 1 | 1 | 15.42 | 3 | 3 | 37.81 | 6 | 12 | 201 | 23.3 | 5.29 |
| P43490 | 48.68 | 1 | 19 | 19 | 112 | 16.90 | 5 | 5 | 14.05 | 4 | 5 | 13.65 | 4 | 5 | 10.39 | 4 | 4 | | | | | | | 1.43 | 1 | 1 | | | | 15.89 | 7 | 10 | 46.84 | 18 | 82 | 491 | 55.5 | 7.15 |
| P43686 | 63.64 | 1 | 24 | 25 | 106 | 27.27 | 8 | 9 | 21.53 | 7 | 7 | 6.70 | 2 | 2 | 10.29 | 4 | 4 | 4.78 | 2 | 2 | | | | 16.03 | 8 | 8 | 6.94 | 3 | 3 | 26.56 | 9 | 11 | 48.56 | 16 | 60 | 418 | 47.3 | 5.21 |
| P43897 | 30.15 | 1 | 6 | 6 | 7 | | | | 9.23 | 2 | 2 | | | | | | | | | | | | | | | | | | | | | | 24.92 | 5 | 5 | 325 | 35.4 | 8.38 |
| P45877 | 17.45 | 1 | 4 | 4 | 7 | 3.77 | 1 | 1 | 3.77 | 1 | 1 | | | | | | | | | | | | | | | | | | | | | | 13.68 | 3 | 5 | 212 | 22.7 | 8.40 |
| P45880 | 70.07 | 1 | 15 | 15 | 192 | 53.74 | 12 | 18 | 46.26 | 11 | 17 | 57.48 | 12 | 17 | 25.17 | 5 | 7 | 31.29 | 8 | 10 | | | | | | | 3.40 | 1 | 1 | | | | 67.35 | 14 | 122 | 294 | 31.5 | 7.56 |
| P45954 | 12.50 | 1 | 4 | 4 | 8 | 3.01 | 1 | 2 | 3.01 | 1 | 1 | 3.01 | 1 | 1 | | | | | | | | | | | | | | | | | | | 12.50 | 4 | 4 | 432 | 47.5 | 6.99 |
| P45974 | 43.82 | 2 | 27 | 28 | 128 | 8.62 | 4 | 5 | 10.49 | 6 | 6 | 2.45 | 2 | 2 | 5.13 | 3 | 3 | | | | | | | 6.64 | 5 | 5 | 6.41 | 3 | 4 | 16.32 | 11 | 13 | 38.81 | 23 | 90 | 858 | 95.7 | 5.03 |
| P45984 | 6.84 | 8 | 2 | 3 | 7 | | | | | | | | | | | | | | | | | | | | | | | | | | | | 6.84 | 3 | 7 | 424 | 48.1 | 5.63 |
| P45985 | 18.80 | 1 | 6 | 6 | 11 | | | | | | | | | | | | | | | | | | | | | | | | | | | | 18.80 | 6 | 11 | 399 | 44.3 | 8.07 |
| P46019 | 1.94 | 1 | 2 | 3 | 4 | | | | | | | | | | | | | | | | | | | | | | | | | | | | 1.94 | 3 | 4 | 1235 | 138.3 | 6.44 |
| P46020 | 4.01 | 1 | 3 | 4 | 5 | | | | | | | | | | | | | | | | | | | | | | | | | | | | 4.01 | 4 | 5 | 1223 | 137.2 | 6.19 |
| P46060 | 8.69 | 1 | 3 | 3 | 5 | | | | | | | | | | | | | | | | | | | | | | | | | | | | 8.69 | 3 | 5 | 587 | 63.5 | 4.68 |
| P46108 | 20.72 | 1 | 5 | 5 | 8 | | | | | | | | | | | | | | | | | | | | | | | | | 9.87 | 2 | 2 | 14.47 | 3 | 6 | 304 | 33.8 | 5.55 |
| P46379 | 22.35 | 1 | 20 | 21 | 50 | 2.65 | 3 | 3 | 1.86 | 2 | 2 | | | | 1.50 | 2 | 2 | 0.88 | 1 | 1 | | | | 1.59 | 2 | 3 | 1.86 | 2 | 2 | 2.56 | 3 | 3 | 22.26 | 20 | 34 | 1132 | 119.3 | 5.60 |
| P46459 | 61.29 | 2 | 52 | 52 | 230 | 25.27 | 18 | 19 | 23.52 | 20 | 22 | 20.03 | 15 | 17 | 13.31 | 9 | 11 | 4.70 | 3 | 3 | | | | 9.27 | 7 | 8 | 7.39 | 5 | 6 | 24.87 | 18 | 20 | 54.03 | 37 | 124 | 744 | 82.5 | 6.95 |
| P46736 | 15.82 | 1 | 3 | 3 | 3 | | | | | | | 0.00 | | | | | | | | | | | | | | | | | | | | | 15.82 | 3 | 3 | 316 | 36.0 | 5.92 |
| P46776 | 37.84 | 1 | 6 | 6 | 16 | 14.19 | 2 | 2 | 14.19 | 2 | 2 | | | | | | | 7.43 | 1 | 1 | | | | | | | | | | 7.43 | 1 | 1 | 37.84 | 6 | 10 | 148 | 16.6 | 11.00 |
| P46777 | 44.44 | 1 | 12 | 12 | 33 | 8.75 | 2 | 2 | 4.04 | 1 | 2 | 6.06 | 2 | 2 | 4.04 | 1 | 2 | 8.08 | 2 | 2 | | | | | | | | | | | | | 42.42 | 11 | 23 | 297 | 34.3 | 9.72 |
| P46778 | 41.88 | 1 | 7 | 8 | 14 | | | | | | | | | | | | | | | | | | | | | | | | | | | | 41.88 | 8 | 14 | 160 | 18.6 | 10.49 |
| P46779 | 45.26 | 1 | 8 | 8 | 20 | | | | 12.41 | 2 | 2 | | | | | | | | | | | | | | | | | | | | | | 45.26 | 8 | 18 | 137 | 15.7 | 12.02 |
| P46781 | 60.31 | 1 | 18 | 18 | 35 | | | | 4.12 | 1 | 1 | | | | | | | | | | | | | | | | | | | | | | 60.31 | 18 | 34 | 194 | 22.6 | 10.65 |
| P46782 | 45.10 | 1 | 9 | 9 | 25 | 7.35 | 1 | 2 | | | | 7.35 | 1 | 2 | | | | | | | | | | | | | | | | | | | 45.10 | 9 | 21 | 204 | 22.9 | 9.72 |
| P46783 | 27.88 | 1 | 4 | 4 | 10 | 5.45 | 1 | 1 | 5.45 | 1 | 1 | 5.45 | 1 | 1 | | | | | | | | | | | | | | | | | | | 27.88 | 4 | 7 | 165 | 18.9 | 10.15 |
| CON_007605243 | 25.45 | 1 | 1 | 10 | 28 | 9.88 | 4 | 4 | 14.37 | 6 | 6 | 21.86 | 7 | 7 | 6.89 | 2 | 2 | 9.88 | 4 | 4 | | | | 2.99 | 1 | 1 | 3.59 | 1 | 1 | 6.89 | 3 | 3 | | | | 334 | 36.7 | 6.44 |
| P46926 | 25.61 | 1 | 2 | 6 | 11 | 4.15 | 1 | 1 | 11.76 | 3 | 3 | 4.15 | 1 | 1 | | | | | | | | | | | | | | | | | | | 22.15 | 5 | 6 | 289 | 32.6 | 6.92 |
| P46939 | 0.47 | 1 | 2 | 2 | 2 | 0.20 | 1 | 1 | | | | | | | | | | | | | | | | | | | | | | | | | 0.26 | 1 | 1 | 3433 | 394.2 | 5.33 |
| P46940 | 51.78 | 2 | 63 | 66 | 207 | 4.10 | 4 | 5 | | | | 3.44 | 4 | 4 | | | | | | | | | | 6.16 | 8 | 8 | | | | 4.04 | 6 | 6 | 50.75 | 63 | 184 | 1657 | 189.1 | 6.48 |
| P46976 | 14.86 | 1 | 5 | 5 | 15 | | | | | | | 2.86 | 1 | 1 | 2.86 | 1 | 1 | | | | | | | 2.86 | 1 | 1 | | | | 5.71 | 2 | 2 | 14.86 | 5 | 10 | 350 | 39.4 | 5.53 |
| P46977 | 17.30 | 1 | 12 | 14 | 45 | 4.11 | 3 | 3 | 4.11 | 4 | 4 | 6.81 | 4 | 4 | | | | | | | | | | | | | | | | | | | 14.18 | 11 | 34 | 705 | 80.5 | 8.07 |
| P47224 | 21.95 | 1 | 2 | 2 | 2 | | | | | | | | | | | | | | | | | | | | | | | | | | | | 21.95 | 2 | 2 | 123 | 13.8 | 5.52 |
| P47712 | 8.54 | 1 | 4 | 4 | 8 | | | | | | | | | | | | | | | | | | | | | | | | | | | | 8.54 | 4 | 8 | 749 | 85.2 | 5.38 |
| P47755 | 63.99 | 1 | 10 | 13 | 21 | 6.29 | 1 | 1 | 3.50 | 1 | 1 | 5.24 | 1 | 1 | 3.50 | 1 | 1 | | | | | | | | | | | | | | | | 54.20 | 11 | 17 | 286 | 32.9 | 5.85 |
| P47756 | 58.12 | 1 | 17 | 17 | 71 | 22.74 | 5 | 6 | 19.49 | 4 | 4 | 25.63 | 6 | 6 | 19.49 | 4 | 5 | 15.52 | 5 | 5 | | | | | | | | | | 2.89 | 1 | 1 | 55.23 | 14 | 44 | 277 | 31.3 | 5.59 |
| P47813 | 40.28 | 2 | 6 | 6 | 8 | | | | | | | | | | | | | | | | | | | | | | | | | | | | 40.28 | 6 | 8 | 144 | 16.5 | 5.24 |
| P47895 | 61.52 | 2 | 25 | 27 | 110 | 3.91 | 2 | 2 | | | | 4.88 | 2 | 3 | 16.80 | 7 | 8 | | | | | | | 16.21 | 7 | 9 | 8.40 | 4 | 4 | 27.15 | 15 | 17 | 58.40 | 23 | 67 | 512 | 56.1 | 7.25 |
| P47897 | 44.77 | 1 | 29 | 30 | 96 | 4.77 | 3 | 3 | 4.39 | 3 | 3 | 2.06 | 1 | 1 | 5.42 | 3 | 3 | | | | | | | | | | | | | 5.42 | 3 | 3 | 42.84 | 29 | 83 | 775 | 87.7 | 7.15 |
| P47914 | 25.16 | 1 | 3 | 3 | 7 | 9.43 | 1 | 1 | 9.43 | 1 | 1 | | | | | | | | | | | | | | | | | | | | | | 25.16 | 3 | 5 | 159 | 17.7 | 11.66 |
| P47929 | 16.91 | 1 | 2 | 2 | 4 | | | | | | | | | | | | | | | | | | | | | | 16.91 | 2 | 2 | 16.91 | 2 | 2 | | | | 136 | 15.1 | 7.62 |
| P47985 | 24.82 | 2 | 8 | 8 | 32 | 13.87 | 3 | 3 | 13.87 | 3 | 5 | | | | | | | 3.28 | 1 | 1 | | | | | | | | | | | | | 24.82 | 8 | 23 | 274 | 29.6 | 8.32 |
| P48047 | 70.89 | 1 | 13 | 13 | 63 | 37.56 | 6 | 7 | 49.77 | 8 | 9 | 22.07 | 4 | 4 | 11.74 | 2 | 2 | | | | | | | | | | | | | | | | 70.89 | 13 | 41 | 213 | 23.3 | 9.96 |
| Q15155 | 15.71 | 1 | 1 | 15 | 28 | | | | | | | 6.06 | 6 | 6 | | | | | | | | | | | | | | | | | | | 14.57 | 14 | 22 | 1222 | 134.2 | 5.81 |
| P69849 | 16.12 | 2 | 1 | 15 | 28 | | | | | | | 6.06 | 6 | 6 | | | | | | | | | | | | | | | | | | | 14.98 | 14 | 22 | 1222 | 134.0 | 5.67 |
| P48147 | 23.80 | 1 | 12 | 12 | 30 | | | | | | | | | | | | | | | | | | | 1.83 | 1 | 1 | | | | 1.83 | 1 | 1 | 23.80 | 12 | 28 | 710 | 80.6 | 5.86 |
| P48163 | 5.07 | 1 | 2 | 2 | 6 | | | | | | | 1.40 | 1 | 1 | | | | | | | | | | | | | | | | | | | 3.67 | 1 | 5 | 572 | 64.1 | 6.13 |

| | | | | | | | | | | | | | | | | | | | | | | | | | | | | | | | | | | | | | | |
|---|---|---|---|---|---|---|---|---|---|---|---|---|---|---|---|---|---|---|---|---|---|---|---|---|---|---|---|---|---|---|---|---|---|---|---|---|---|---|
| P48444 | 27.79 | 1 | 14 | 14 | 38 | 2.15 | 1 | 1 | 2.15 | 1 | 1 | 4.31 | 2 | 2 | 4.31 | 2 | 2 | | | | | | | 2.15 | 1 | 1 | 3.91 | 2 | 2 | 8.81 | 4 | 4 | 27.59 | 13 | 23 | 511 | 57.2 | 6.21 |
| P48449 | 27.73 | 1 | 17 | 17 | 22 | | | | | | | 4.37 | 3 | 3 | | | | | | | | | | | | | | | | | | | 24.59 | 15 | 19 | 732 | 83.3 | 6.61 |
| O00507 | 5.75 | 5 | 1 | 14 | 28 | | | | | | | | | | | | | | | | | | | | | | | | | | | | 5.75 | 14 | 28 | 2555 | 290.9 | 5.86 |
| CON_P02666 | 8.48 | 1 | 1 | 1 | 28 | | | | | | | | | | | | | | | | | | | | | | | | | | | | 8.48 | 1 | 28 | 224 | 25.1 | 5.35 |
| P48454 | 24.02 | 1 | 8 | 11 | 35 | 5.66 | 3 | 3 | 4.30 | 2 | 2 | | | | | | | 0.00 | | | | | | 1.17 | 1 | 1 | 0.00 | | | 1.17 | 1 | 1 | 19.92 | 9 | 28 | 512 | 58.1 | 6.98 |
| P48506 | 17.58 | 1 | 9 | 10 | 16 | | | | | | | | | | | | | | | | | | | | | | | | | | | | 17.58 | 10 | 16 | 637 | 72.7 | 6.09 |
| P48507 | 18.98 | 1 | 4 | 4 | 6 | | | | | | | | | | | | | | | | | | | | | | | | | 8.39 | 2 | 2 | 14.23 | 3 | 4 | 274 | 30.7 | 6.02 |
| P48556 | 36.86 | 1 | 9 | 9 | 23 | 13.71 | 3 | 3 | 11.14 | 3 | 3 | 11.71 | 4 | 4 | 6.57 | 3 | 3 | | | | | | | | | | | | | | | | 27.43 | 7 | 10 | 350 | 39.6 | 9.70 |
| P48637 | 11.60 | 1 | 6 | 6 | 11 | 3.80 | 2 | 2 | 3.80 | 2 | 2 | 1.90 | 1 | 1 | | | | | | | | | | | | | | | | | | | 11.60 | 6 | 6 | 474 | 52.4 | 5.92 |
| P48643 | 71.72 | 1 | 42 | 43 | 346 | 53.42 | 23 | 31 | 53.23 | 26 | 46 | 40.11 | 19 | 25 | 44.73 | 23 | 30 | 31.79 | 18 | 23 | | | | 15.53 | 9 | 9 | 16.45 | 9 | 11 | 32.53 | 17 | 23 | 62.85 | 32 | 148 | 541 | 59.6 | 5.66 |
| P48723 | 17.41 | 1 | 7 | 7 | 14 | 2.55 | 1 | 1 | 4.67 | 2 | 2 | 8.28 | 3 | 3 | | | | | | | | | | | | | | | | 3.18 | 1 | 1 | 8.07 | 4 | 7 | 471 | 51.9 | 5.76 |
| P48729 | 36.80 | 1 | 4 | 10 | 19 | 2.97 | 1 | 1 | 5.34 | 2 | 2 | | | | | | | | | | | | | | | | | | | | | | 34.42 | 9 | 16 | 337 | 38.9 | 9.57 |
| P31321 | 8.40 | 1 | 1 | 7 | 27 | 6.04 | 3 | 3 | 5.77 | 3 | 4 | 3.67 | 2 | 2 | 3.67 | 2 | 2 | 2.10 | 1 | 1 | | | | | | | 2.10 | 1 | 1 | 2.10 | 1 | 1 | 8.14 | 6 | 13 | 381 | 43.0 | 5.71 |
| P48735 | 47.57 | 1 | 21 | 22 | 57 | 23.01 | 9 | 9 | 5.53 | 3 | 3 | 26.77 | 10 | 11 | 4.42 | 2 | 2 | | | | | | | | | | | | | 11.95 | 4 | 4 | 46.46 | 18 | 28 | 452 | 50.9 | 8.69 |
| P48739 | 22.14 | 1 | 2 | 4 | 6 | | | | | | | | | | | | | | | | | | | | | | | | | | | | 22.14 | 4 | 6 | 271 | 31.5 | 6.87 |
| P49069 | 30.07 | 1 | 4 | 4 | 6 | | | | | | | | | | | | | | | | | | | | | | | | | | | | 30.07 | 4 | 6 | 296 | 32.9 | 8.05 |
| P49184 | 15.23 | 1 | 2 | 2 | 4 | | | | | | | | | | | | | | | | | | | | | | | | | | | | 15.23 | 2 | 4 | 302 | 33.9 | 5.74 |
| P49189 | 51.42 | 1 | 23 | 23 | 106 | 30.57 | 12 | 12 | 15.99 | 7 | 8 | 27.13 | 12 | 15 | 18.83 | 8 | 9 | 8.91 | 5 | 5 | | | | 6.88 | 3 | 3 | 2.83 | 1 | 1 | 20.24 | 9 | 9 | 40.69 | 18 | 44 | 494 | 53.8 | 5.87 |
| P49207 | 23.08 | 1 | 4 | 4 | 4 | | | | | | | | | | | | | | | | | | | | | | | | | | | | 23.08 | 4 | 4 | 117 | 13.3 | 11.47 |
| P49221 | 78.22 | 1 | 47 | 48 | 657 | 40.35 | 21 | 26 | 38.60 | 19 | 25 | 52.63 | 26 | 34 | 46.49 | 28 | 36 | 37.43 | 22 | 28 | | | | 31.87 | 18 | 23 | 36.84 | 22 | 29 | 40.94 | 23 | 28 | 73.68 | 45 | 428 | 684 | 77.1 | 6.76 |
| P49223 | 29.21 | 1 | 3 | 3 | 19 | 10.11 | 1 | 1 | 24.72 | 2 | 3 | 24.72 | 2 | 2 | 24.72 | 2 | 3 | 24.72 | 2 | 4 | | | | | | | 10.11 | 1 | 2 | 10.11 | 1 | 1 | 29.21 | 3 | 3 | 89 | 10.2 | 6.44 |
| P49257 | 29.02 | 1 | 13 | 13 | 39 | 4.51 | 2 | 2 | 6.67 | 3 | 4 | 13.14 | 5 | 7 | 3.92 | 2 | 2 | 3.92 | 2 | 2 | | | | | | | | | | 8.24 | 3 | 3 | 28.63 | 10 | 19 | 510 | 57.5 | 6.77 |
| P49327 | 59.06 | 2 | 118 | 119 | 1058 | 11.79 | 19 | 22 | 5.14 | 10 | 12 | 24.53 | 46 | 53 | 21.43 | 43 | 47 | 3.90 | 7 | 7 | | | | 11.31 | 24 | 26 | 5.02 | 9 | 9 | 26.48 | 49 | 63 | 56.83 | 108 | 819 | 2511 | 273.3 | 6.44 |
| P49354 | 25.86 | 1 | 8 | 8 | 24 | | | | | | | | | | | | | 3.43 | 1 | 1 | | | | | | | | | | 3.43 | 1 | 1 | 25.86 | 8 | 22 | 379 | 44.4 | 5.08 |
| P49356 | 12.13 | 1 | 4 | 4 | 5 | | | | | | | | | | | | | | | | | | | | | | | | | | | | 12.13 | 4 | 5 | 437 | 48.7 | 5.82 |
| P49368 | 88.62 | 2 | 48 | 48 | 350 | 56.15 | 21 | 23 | 62.39 | 27 | 39 | 42.20 | 18 | 21 | 44.22 | 21 | 24 | 44.77 | 20 | 23 | | | | 20.18 | 7 | 9 | 24.77 | 11 | 11 | 33.21 | 17 | 20 | 84.95 | 41 | 180 | 545 | 60.5 | 6.49 |
| P49411 | 60.40 | 1 | 26 | 26 | 69 | 18.58 | 7 | 7 | 16.59 | 6 | 6 | 15.93 | 6 | 6 | 0.00 | | | 3.54 | 1 | 1 | | | | 7.96 | 3 | 3 | 5.75 | 2 | 2 | 26.77 | 9 | 11 | 51.99 | 22 | 33 | 452 | 49.5 | 7.61 |
| P49419 | 40.63 | 1 | 14 | 14 | 23 | | | | | | | 2.60 | 1 | 1 | | | | | | | | | | | | | | | | 2.78 | 1 | 1 | 40.63 | 14 | 21 | 539 | 58.5 | 7.99 |
| P49441 | 7.02 | 1 | 3 | 3 | 3 | | | | | | | | | | | | | | | | | | | | | | | | | | | | 7.02 | 3 | 3 | 399 | 44.0 | 5.26 |
| P49447 | 14.34 | 1 | 3 | 3 | 10 | | | | | | | 4.38 | 1 | 1 | | | | | | | | | | | | | | | | | | | 14.34 | 3 | 9 | 251 | 27.5 | 8.56 |
| P49458 | 41.86 | 1 | 3 | 3 | 4 | | | | | | | | | | 12.79 | 1 | 1 | | | | | | | | | | | | | | | | 29.07 | 2 | 3 | 86 | 10.1 | 7.97 |
| P49585 | 40.87 | 2 | 12 | 12 | 19 | | | | | | | | | | 6.27 | 2 | 2 | | | | | | | | | | | | | | | | 40.87 | 12 | 17 | 367 | 41.7 | 7.25 |
| P49588 | 48.97 | 1 | 41 | 41 | 238 | 5.17 | 4 | 4 | 1.14 | 1 | 1 | 3.93 | 3 | 3 | 9.61 | 7 | 8 | 1.14 | 1 | 1 | 0.00 | | | 10.12 | 8 | 8 | 9.61 | 8 | 9 | 20.87 | 18 | 20 | 45.66 | 35 | 182 | 968 | 106.7 | 5.53 |
| P49589 | 22.19 | 1 | 13 | 13 | 26 | | | | 8.16 | 4 | 5 | 2.67 | 2 | 2 | | | | | | | | | | | | | | | | | | | 18.32 | 11 | 19 | 748 | 85.4 | 6.76 |
| P49591 | 33.85 | 1 | 14 | 14 | 38 | 1.56 | 1 | 1 | 3.70 | 2 | 2 | | | | | | | | | | | | | 1.75 | 1 | 1 | | | | 2.53 | 1 | 2 | 32.30 | 12 | 32 | 514 | 58.7 | 6.43 |
| CON_P22629 | 10.38 | 1 | 1 | 1 | 27 | | | | | | | | | | | | | | | | | | | | | | | | | | | | 10.38 | 1 | 27 | 183 | 18.8 | 8.35 |
| P49641 | 5.74 | 1 | 5 | 6 | 7 | | | | | | | | | | | | | | | | | | | | | | | | | | | | 5.74 | 6 | 7 | 1150 | 130.5 | 6.84 |
| P49720 | 41.95 | 1 | 8 | 8 | 37 | 26.34 | 4 | 4 | 28.29 | 4 | 7 | 28.29 | 4 | 7 | | | | | | | | | | | | | | | | | | | 40.98 | 7 | 19 | 205 | 22.9 | 6.55 |
| P49721 | 55.72 | 2 | 12 | 13 | 26 | 8.46 | 2 | 2 | 20.40 | 3 | 3 | 4.98 | 1 | 1 | | | | | | | | | | | | | | | | | | | 45.77 | 11 | 20 | 201 | 22.8 | 7.02 |
| P49748 | 60.31 | 1 | 32 | 32 | 73 | 9.92 | 5 | 5 | 16.95 | 8 | 8 | 42.29 | 21 | 21 | | | | | | | | | | | | | | | | 1.83 | 1 | 1 | 52.37 | 26 | 38 | 655 | 70.3 | 8.75 |
| P49754 | 10.77 | 1 | 7 | 7 | 8 | | | | | | | | | | | | | | | | | | | | | | | | | | | | 10.77 | 7 | 8 | 854 | 98.5 | 5.85 |
| P49755 | 51.60 | 1 | 13 | 13 | 63 | 31.51 | 5 | 7 | 24.20 | 5 | 10 | 31.51 | 6 | 9 | 16.44 | 3 | 4 | 21.00 | 4 | 4 | | | | | | | | | | | | | 42.47 | 11 | 29 | 219 | 25.0 | 7.44 |
| P49768 | 18.42 | 2 | 6 | 6 | 12 | | | | | | | | | | | | | | | | | | | | | | | | | | | | 18.42 | 6 | 12 | 467 | 52.6 | 5.31 |
| P49770 | 31.05 | 1 | 9 | 9 | 20 | | | | 5.13 | 1 | 1 | 3.42 | 1 | 1 | | | | | | | | | | | | | | | | | | | 31.05 | 9 | 18 | 351 | 39.0 | 6.16 |
| P49773 | 89.68 | 1 | 13 | 13 | 53 | 16.67 | 2 | 3 | 11.11 | 1 | 1 | 11.11 | 1 | 1 | 71.43 | 4 | 6 | 79.37 | 6 | 6 | 15.87 | 1 | 1 | 30.16 | 2 | 3 | 50.79 | 3 | 4 | 84.92 | 9 | 15 | 78.57 | 7 | 13 | 126 | 13.8 | 6.95 |
| P49788 | 16.67 | 1 | 5 | 5 | 13 | 10.88 | 3 | 3 | 3.06 | 1 | 1 | 13.95 | 4 | 5 | 3.06 | 1 | 1 | | | | | | | | | | | | | | | | 7.14 | 2 | 3 | 294 | 33.3 | 8.51 |
| P49821 | 34.91 | 1 | 15 | 15 | 26 | 1.29 | 1 | 1 | 6.03 | 2 | 2 | 3.45 | 2 | 2 | | | | | | | | | | | | | | | | | | | 29.53 | 12 | 21 | 464 | 50.8 | 8.21 |
| P49840 | 15.94 | 1 | 3 | 7 | 10 | | | | | | | | | | | | | | | | | | | | | | | | | | | | 15.94 | 7 | 10 | 483 | 50.9 | 8.75 |
| P49841 | 32.38 | 1 | 6 | 10 | 13 | | | | | | | | | | | | | | | | | | | | | | | | | | | | 32.38 | 10 | 13 | 420 | 46.7 | 8.78 |
| P49901 | 81.03 | 1 | 6 | 6 | 29 | 50.86 | 4 | 6 | 16.38 | 1 | 1 | 30.17 | 2 | 3 | | | | 53.45 | 3 | 5 | | | | | | | | | | | | | 81.03 | 5 | 14 | 116 | 12.8 | 8.07 |
| P49902 | 40.11 | 1 | 16 | 16 | 53 | 8.20 | 3 | 5 | 4.99 | 2 | 2 | 11.23 | 5 | 7 | | | | | | | | | | | | | | | | | | | 37.43 | 15 | 39 | 561 | 64.9 | 6.14 |
| P49908 | 8.40 | 1 | 2 | 2 | 4 | 5.77 | 1 | 2 | 5.77 | 1 | 1 | | | | | | | | | | | | | | | | | | | | | | 2.62 | 1 | 1 | 381 | 43.2 | 7.87 |
| P49913 | 65.29 | 2 | 19 | 19 | 131 | 41.18 | 9 | 14 | 41.18 | 9 | 14 | 45.88 | 9 | 13 | 19.41 | 4 | 6 | 53.53 | 9 | 13 | 7.65 | 1 | 1 | | | | 12.35 | 2 | 2 | 7.65 | 1 | 1 | 62.35 | 16 | 67 | 170 | 19.3 | 9.41 |
| P50135 | 20.55 | 1 | 5 | 5 | 12 | | | | | | | | | | | | | | | | | | | | | | | | | | | | 20.55 | 5 | 12 | 292 | 33.3 | 5.34 |
| P50148 | 40.67 | 2 | 7 | 13 | 31 | 4.18 | 1 | 1 | | | | | | | | | | | | | | | | | | | | | | | | | 40.67 | 13 | 30 | 359 | 42.1 | 5.68 |
| P50213 | 44.26 | 1 | 16 | 16 | 53 | 20.49 | 7 | 7 | 24.04 | 8 | 9 | 20.77 | 6 | 6 | 2.73 | 1 | 1 | | | | | | | | | | | | | | | | 38.80 | 12 | 30 | 366 | 39.6 | 6.92 |
| P50225 | 27.80 | 3 | 7 | 7 | 9 | | | | | | | | | | | | | | | | | | | | | | | | | 2.37 | 1 | 1 | 27.80 | 7 | 8 | 295 | 34.1 | 6.62 |
| P50336 | 12.58 | 1 | 3 | 4 | 5 | | | | | | | | | | | | | | | | | | | | | | | | | 0.00 | | | 12.58 | 4 | 5 | 477 | 50.7 | 8.16 |
| P50395 | 72.36 | 1 | 20 | 29 | 151 | 37.53 | 13 | 17 | 36.85 | 13 | 15 | 43.37 | 14 | 16 | 8.76 | 3 | 4 | 8.09 | 3 | 3 | | | | 4.49 | 2 | 2 | 6.29 | 2 | 2 | 13.48 | 5 | 5 | 67.87 | 25 | 87 | 445 | 50.6 | 6.47 |
| P50416 | 4.40 | 1 | 3 | 3 | 3 | | | | | | | | | | | | | | | | | | | | | | | | | 0.00 | | | 4.40 | 3 | 3 | 773 | 88.3 | 8.65 |
| P50443 | 7.31 | 1 | 4 | 4 | 8 | | | | | | | | | | | | | | | | | | | | | | | | | | | | 7.31 | 4 | 8 | 739 | 81.6 | 8.38 |
| P50454 | 5.26 | 1 | 2 | 2 | 2 | | | | | | | | | | | | | | | | | | | | | | | | | | | | 5.26 | 2 | 2 | 418 | 46.4 | 8.69 |
| P50502 | 46.88 | 3 | 14 | 14 | 98 | 36.31 | 11 | 12 | 17.89 | 7 | 8 | 17.07 | 5 | 5 | 21.68 | 8 | 10 | 22.49 | 7 | 9 | | | | 24.93 | 8 | 8 | 8.67 | 4 | 4 | 17.34 | 7 | 8 | 24.93 | 8 | 34 | 369 | 41.3 | 5.27 |
| P50552 | 20.79 | 1 | 7 | 7 | 12 | 3.16 | 1 | 1 | 3.16 | 1 | 1 | 3.16 | 1 | 1 | 3.16 | 1 | 1 | | | | | | | 3.16 | 1 | 1 | | | | 10.00 | 3 | 3 | 10.53 | 3 | 4 | 380 | 39.8 | 8.94 |
| P50570 | 23.45 | 3 | 16 | 16 | 31 | | | | | | | | | | 2.53 | 2 | 2 | | | | | | | 2.76 | 2 | 3 | | | | 6.55 | 5 | 5 | 19.66 | 13 | 21 | 870 | 98.0 | 7.44 |
| P50583 | 72.11 | 1 | 8 | 8 | 13 | | | | | | | | | | | | | | | | | | | | | | | | | 4.76 | 1 | 1 | 67.35 | 7 | 12 | 147 | 16.8 | 5.35 |
| P50591 | 25.62 | 1 | 6 | 6 | 8 | | | | | | | | | | | | | | | | | | | | | | 0.00 | | | 2.85 | 1 | 1 | 25.62 | 6 | 7 | 281 | 32.5 | 7.42 |
| P50851 | 9.57 | 2 | 24 | 24 | 48 | | | | | | | 0.00 | | | | | | | | | | | | 0.28 | 1 | 1 | | | | | | | 9.57 | 24 | 47 | 2863 | 318.9 | 5.60 |
| P50897 | 23.20 | 1 | 4 | 4 | 7 | | | | | | | | | | | | | | | | | | | | | | | | | | | | 23.20 | 4 | 7 | 306 | 34.2 | 6.52 |
| P50914 | 27.44 | 1 | 7 | 7 | 28 | 5.58 | 1 | 1 | 14.88 | 3 | 3 | 5.58 | 1 | 1 | 14.88 | 3 | 4 | | | | | | | | | | | | | 5.58 | 1 | 1 | 26.51 | 6 | 18 | 215 | 23.4 | 10.93 |
| P50990 | 72.26 | 1 | 45 | 45 | 322 | 45.26 | 21 | 24 | 51.46 | 26 | 32 | 49.27 | 26 | 32 | 61.86 | 32 | 38 | 42.70 | 24 | 26 | | | | 13.69 | 6 | 7 | 12.41 | 6 | 7 | 32.66 | 16 | 19 | 66.24 | 38 | 137 | 548 | 59.6 | 5.60 |
| P50991 | 73.65 | 1 | 38 | 39 | 263 | 31.91 | 12 | 15 | 39.33 | 18 | 22 | 30.98 | 15 | 17 | 25.60 | 11 | 14 | 26.16 | 12 | 15 | 5.19 | 3 | 3 | 6.31 | 3 | 3 | 3.90 | 2 | 3 | 15.03 | 7 | 8 | 66.05 | 33 | 163 | 539 | 57.9 | 7.83 |
| P50995 | 22.57 | 1 | 10 | 10 | 28 | 15.05 | 6 | 6 | 4.55 | 2 | 2 | 10.10 | 4 | 4 | 2.38 | 1 | 1 | | | | | | | | | | | | | | | | 17.03 | 7 | 15 | 505 | 54.4 | 7.65 |
| P51114 | 15.62 | 2 | 7 | 9 | 16 | | | | 2.74 | 1 | 1 | 4.03 | 2 | 2 | 2.58 | 2 | 2 | | | | | | | 2.09 | 1 | 1 | | | | 1.29 | 1 | 1 | 14.33 | 7 | 9 | 621 | 69.7 | 6.15 |
| P51148 | 58.80 | 1 | 6 | 9 | 28 | 11.11 | 2 | 2 | 18.06 | 3 | 4 | 11.57 | 2 | 2 | | | | | | | | | | | | | | | | | | | 54.17 | 8 | 20 | 216 | 23.5 | 8.41 |
| P51149 | 83.09 | 2 | 16 | 17 | 62 | 36.71 | 6 | 6 | 16.43 | 3 | 4 | 31.88 | 6 | 6 | 4.83 | 1 | 1 | | | | | | | 13.04 | 3 | 3 | 4.83 | 1 | 1 | 14.49 | 3 | 3 | 74.40 | 16 | 38 | 207 | 23.5 | 6.70 |
| P51153 | 38.92 | 2 | 6 | 8 | 29 | 5.42 | 1 | 1 | 5.42 | 1 | 1 | 9.36 | 2 | 2 | 3.94 | 1 | 1 | | | | | | | 3.94 | 1 | 1 | 9.36 | 2 | 2 | 9.36 | 2 | 2 | 38.92 | 8 | 19 | 203 | 22.8 | 9.19 |
| P51157 | 56.56 | 1 | 11 | 11 | 24 | | | | | | | 0.00 | | | | | | | | | | | | | | | 9.95 | 2 | 2 | 10.86 | 2 | 2 | 50.68 | 10 | 20 | 221 | 24.8 | 5.97 |
| P51159 | 59.28 | 1 | 10 | 12 | 45 | 9.05 | 2 | 2 | 9.95 | 2 | 2 | 8.60 | 2 | 2 | 21.72 | 4 | 5 | | | | | | | 4.98 | 1 | 1 | | | | 9.95 | 2 | 2 | 59.28 | 11 | 31 | 221 | 24.9 | 5.22 |
| P51452 | 48.11 | 1 | 7 | 7 | 10 | | | | | | | | | | | | | | | | | | | | | | | | | 7.03 | 1 | 1 | 48.11 | 7 | 9 | 185 | 20.5 | 7.80 |
| P51553 | 21.37 | 1 | 4 | 4 | 6 | | | | 13.74 | 2 | 3 | | | | | | | | | | | | | | | | | | | | | | 13.49 | 3 | 3 | 393 | 42.8 | 8.50 |
| P51570 | 16.58 | 1 | 6 | 6 | 8 | | | | | | | | | | | | | | | | | | | | | | | | | | | | 16.58 | 6 | 8 | 392 | 42.2 | 6.46 |
| P51571 | 36.42 | 1 | 5 | 5 | 17 | 6.36 | 1 | 1 | 12.14 | 2 | 3 | | | | | | | | | | | | | | | | | | | | | | 36.42 | 5 | 13 | 173 | 19.0 | 6.15 |
| P51572 | 30.49 | 1 | 11 | 11 | 24 | 8.54 | 3 | 3 | 3.66 | 1 | 1 | 10.98 | 3 | 3 | | | | | | | 0.00 | | | | | | | | | | | | 26.83 | 8 | 17 | 246 | 28.0 | 8.44 |
| P51580 | 27.35 | 1 | 7 | 7 | 10 | | | | | | | | | | | | | | | | | | | | | | | | | | | | 27.35 | 7 | 10 | 245 | 28.2 | 6.23 |
| P51648 | 10.93 | 2 | 5 | 5 | 6 | | | | | | | | | | | | | | | | | | | | | | | | | | | | 10.93 | 5 | 6 | 485 | 54.8 | 7.88 |
| P51649 | 10.84 | 1 | 5 | 5 | 7 | | | | | | | | | | | | | | | | | | | | | | | | | | | | 10.84 | 5 | 7 | 535 | 57.2 | 8.28 |
| P51654 | 3.79 | 1 | 2 | 2 | 2 | | | | | | | | | | | | | | | | | | | | | | | | | | | | 3.79 | 2 | 2 | 580 | 65.5 | 6.37 |
| P51659 | 50.41 | 1 | 27 | 27 | 74 | 1.77 | 1 | 1 | 8.97 | 4 | 4 | 16.58 | 8 | 10 | 1.77 | 1 | 1 | | | | | | | | | | | | | 1.77 | 1 | 1 | 46.06 | 24 | 57 | 736 | 79.6 | 8.84 |
| P51665 | 54.94 | 1 | 15 | 15 | 55 | 7.72 | 2 | 2 | 27.78 | 6 | 8 | | | | 33.02 | 7 | 10 | 16.05 | 5 | 6 | | | | | | | | | | 11.42 | 3 | 4 | 39.51 | 11 | 25 | 324 | 37.0 | 6.77 |
| P51668 | 24.49 | 2 | 2 | 3 | 3 | | | | | | | | | | | | | | | | | | | | | | | | | | | | 24.49 | 3 | 3 | 147 | 16.6 | 7.42 |
| P51692 | 24.78 | 2 | 16 | 17 | 28 | | | | | | | | | | 1.40 | 1 | 1 | | | | | | | 1.40 | 1 | 2 | | | | | | | 24.78 | 16 | 23 | 787 | 89.8 | 6.05 |
| P51808 | 49.14 | 1 | 4 | 4 | 4 | | | | | | | | | | | | | | | | | | | | | | | | | | | | 49.14 | 4 | 4 | 116 | 13.1 | 5.66 |
| P51809 | 9.55 | 1 | 2 | 2 | 2 | | | | | | | | | | | | | | | | | | | | | | | | | | | | 9.55 | 2 | 2 | 220 | 24.9 | 8.60 |
| P51812 | 22.03 | 10 | 8 | 15 | 51 | | | | | | | | | | | | | | | | | | | 1.22 | 1 | 1 | | | | 2.70 | 2 | 2 | 20.81 | 14 | 48 | 740 | 83.7 | 6.89 |
| P51854 | 39.93 | 1 | 17 | 18 | 46 | | | | 3.36 | 1 | 2 | 4.87 | 2 | 2 | | | | 1.68 | 1 | 1 | | | | | | | 1.68 | 1 | 1 | 1.68 | 1 | 1 | 39.93 | 18 | 39 | 596 | 65.3 | 5.86 |
| P51970 | 30.23 | 1 | 6 | 6 | 9 | 12.21 | 2 | 2 | 15.12 | 2 | 2 | | | | | | | | | | | | | | | | | | | | | | 30.23 | 5 | 5 | 172 | 20.1 | 7.65 |
| P51991 | 43.92 | 1 | 16 | 16 | 30 | | | | 8.20 | 2 | 3 | 4.23 | 1 | 1 | 6.08 | 1 | 1 | 10.05 | 2 | 2 | | | | 8.99 | 2 | 2 | 23.54 | 8 | 10 | 17.72 | 4 | 4 | 16.40 | 4 | 7 | 378 | 39.6 | 9.01 |
| P52198 | 37.44 | 1 | 8 | 8 | 8 | | | | | | | | | | | | | | | | | | | | | | | | | | | | 37.44 | 8 | 8 | 227 | 25.4 | 8.35 |
| P52209 | 49.69 | 1 | 20 | 20 | 124 | 27.33 | 10 | 13 | 21.53 | 8 | 10 | 28.36 | 10 | 12 | 24.22 | 10 | 10 | 7.66 | 3 | 3 | | | | 7.87 | 3 | 3 | 4.14 | 2 | 2 | 12.01 | 4 | 4 | 45.55 | 18 | 67 | 483 | 53.1 | 7.23 |
| P52272 | 34.38 | 1 | 23 | 23 | 52 | 1.51 | 1 | 1 | 1.37 | 1 | 1 | | | | 3.42 | 2 | 2 | 8.36 | 5 | 5 | 1.10 | 1 | 1 | 2.19 | 1 | 1 | 13.29 | 8 | 8 | 10.27 | 6 | 6 | 27.40 | 17 | 27 | 730 | 77.5 | 8.70 |
| P52292 | 65.22 | 2 | 21 | 21 | 116 | 28.36 | 9 | 11 | 25.14 | 10 | 11 | 10.02 | 4 | 4 | 11.34 | 5 | 5 | 10.40 | 5 | 5 | | | | | | | 5.29 | 2 | 2 | 5.10 | 2 | 2 | 54.63 | 18 | 76 | 529 | 57.8 | 5.40 |
| P52294 | 15.80 | 1 | 2 | 8 | 22 | 1.86 | 1 | 1 | | | | | | | | | | | | | | | | | | | | | | | | | 15.61 | 7 | 21 | 538 | 60.2 | 5.01 |
| P52306 | 27.18 | 1 | 11 | 12 | 16 | | | | | | | | | | | | | | | | | | | | | | | | | | | | 27.18 | 12 | 16 | 607 | 66.3 | 5.31 |
| P52434 | 18.00 | 1 | 2 | 2 | 2 | | | | | | | | | | | | | | | | | | | | | | | | | | | | 18.00 | 2 | 2 | 150 | 17.1 | 4.68 |
| P52565 | 80.88 | 1 | 13 | 13 | 67 | 36.76 | 4 | 4 | 33.33 | 4 | 5 | 21.08 | 3 | 3 | 22.55 | 4 | 5 | 8.33 | 1 | 1 | | | | 29.90 | 4 | 5 | 31.37 | 3 | 3 | 29.90 | 5 | 6 | 65.20 | 9 | 35 | 204 | 23.2 | 5.11 |
| P52566 | 24.38 | 1 | 3 | 3 | 4 | | | | | | | 6.47 | 1 | 1 | | | | | | | | | | | | | | | | 4.98 | 1 | 1 | 12.94 | 1 | 1 | 201 | 23.0 | 5.21 |
| P52597 | 21.20 | 1 | 4 | 6 | 12 | | | | | | | 2.89 | 1 | 1 | | | | | | | | | | 2.41 | 1 | 1 | 3.86 | 1 | 2 | | | | 17.35 | 5 | 8 | 415 | 45.6 | 5.58 |
| P52630 | 4.00 | 1 | 2 | 2 | 6 | | | | | | | | | | | | | | | | | | | | | | | | | | | | 4.00 | 2 | 6 | 851 | 97.9 | 5.49 |
| P52758 | 33.58 | 1 | 3 | 3 | 4 | | | | | | | | | | | | | | | | | | | | | | | | | | | | 33.58 | 3 | 3 | 137 | 14.5 | 8.68 |
| P52788 | 51.37 | 1 | 19 | 19 | 46 | | | | | | | 3.01 | 1 | 1 | 16.67 | 4 | 4 | | | | 6.28 | 1 | 1 | 2.19 | 1 | 1 | 5.46 | 2 | 2 | 20.49 | 8 | 8 | 46.72 | 16 | 29 | 366 | 41.2 | 5.02 |
| P52888 | 8.85 | 1 | 5 | 5 | 7 | 0.00 | | | | | | | | | | | | | | | | | | | | | | | | | | | 8.85 | 5 | 7 | 689 | 78.8 | 6.05 |
| P52907 | 76.22 | 1 | 12 | 15 | 58 | 9.44 | 2 | 2 | 30.07 | 6 | 7 | 27.62 | 5 | 5 | 18.53 | 4 | 4 | 9.44 | 2 | 2 | | | | | | | 15.03 | 3 | 3 | 15.03 | 3 | 3 | 72.73 | 13 | 32 | 286 | 32.9 | 5.69 |
| P52943 | 32.21 | 1 | 3 | 3 | 9 | | | | | | | | | | | | | | | | | | | | | | | | | | | | 32.21 | 3 | 9 | 208 | 22.5 | 8.72 |
| P52948 | 17.34 | 3 | 29 | 29 | 55 | 1.05 | 2 | 2 | 1.71 | 3 | 3 | 1.21 | 2 | 2 | | | | 1.27 | 2 | 2 | | | | | | | 1.10 | 1 | 1 | | | | 16.84 | 27 | 45 | 1817 | 197.5 | 6.40 |
| P53004 | 24.66 | 1 | 6 | 6 | 10 | | | | | | | | | | | | | | | | | | | | | | | | | 4.73 | 1 | 1 | 24.66 | 6 | 9 | 296 | 33.4 | 6.44 |
| P53007 | 40.19 | 1 | 11 | 11 | 23 | 6.11 | 2 | 2 | 20.26 | 6 | 7 | 3.54 | 1 | 1 | | | | | | | | | | | | | | | | | | | 40.19 | 11 | 13 | 311 | 34.0 | 9.89 |
| P53041 | 13.23 | 1 | 6 | 6 | 7 | | | | | | | 0.00 | | | | | | | | | | | | | | | | | | | | | 13.23 | 6 | 7 | 499 | 56.8 | 6.28 |
| P53367 | 24.40 | 1 | 8 | 8 | 17 | 4.83 | 1 | 1 | 6.17 | 2 | 2 | | | | | | | 0.00 | | | | | | 0.00 | | | 0.00 | | | 0.00 | | | 24.40 | 7 | 14 | 373 | 41.7 | 6.70 |
| P63027 | 34.48 | 2 | 1 | 4 | 24 | 20.69 | 2 | 3 | 20.69 | 2 | 3 | 20.69 | 2 | 2 | 20.69 | 1 | 1 | | | | | | | | | | | | | | | | 34.48 | 4 | 15 | 116 | 12.7 | 8.13 |
| P53384 | 24.69 | 1 | 5 | 5 | 11 | | | | | | | | | | 2.81 | 1 | 1 | | | | | | | | | | | | | 7.81 | 2 | 2 | 19.06 | 4 | 8 | 320 | 34.5 | 5.33 |
| P53396 | 52.59 | 1 | 51 | 51 | 293 | 14.62 | 12 | 13 | 15.53 | 14 | 15 | 24.70 | 24 | 26 | 14.17 | 12 | 12 | 3.91 | 4 | 4 | | | | 1.54 | 2 | 2 | 1.63 | 2 | 3 | 11.08 | 10 | 10 | 47.77 | 46 | 208 | 1101 | 120.8 | 7.33 |
| P53582 | 5.96 | 1 | 2 | 2 | 6 | | | | | | | | | | | | | | | | | | | | | | | | | | | | 5.96 | 2 | 6 | 386 | 43.2 | 7.17 |
| P53597 | 32.66 | 1 | 10 | 10 | 35 | 8.09 | 2 | 3 | 10.98 | 4 | 7 | 21.97 | 5 | 9 | | | | 11.27 | 3 | 4 | | | | | | | | | | 6.07 | 2 | 2 | 24.57 | 6 | 10 | 346 | 36.2 | 8.79 |
| P53602 | 11.50 | 1 | 5 | 5 | 8 | | | | 2.50 | 1 | 1 | 2.50 | 1 | 1 | | | | | | | | | | | | | | | | | | | 11.50 | 5 | 6 | 400 | 43.4 | 7.23 |
| P53609 | 13.00 | 1 | 5 | 5 | 6 | | | | | | | | | | | | | | | | | | | | | | | | | | | | 13.00 | 5 | 6 | 377 | 42.3 | 6.83 |
| P53611 | 9.06 | 1 | 3 | 3 | 3 | | | | | | | | | | | | | | | | | | | | | | | | | | | | 9.06 | 3 | 3 | 331 | 36.9 | 5.03 |
| P53618 | 42.71 | 1 | 31 | 31 | 94 | 1.15 | 1 | 1 | 1.15 | 1 | 1 | 1.15 | 1 | 1 | 1.15 | 1 | 1 | | | | | | | | | | | | | 3.15 | 2 | 2 | 42.71 | 31 | 88 | 953 | 107.1 | 6.05 |
| P53621 | 44.53 | 1 | 47 | 47 | 160 | 0.82 | 1 | 1 | 0.82 | 1 | 1 | 6.70 | 7 | 7 | 1.06 | 1 | 1 | 1.06 | 1 | 1 | | | | 1.06 | 1 | 1 | | | | 1.06 | 1 | 1 | 43.55 | 46 | 147 | 1224 | 138.3 | 7.66 |
| P53680 | 30.28 | 1 | 4 | 4 | 12 | | | | | | | | | | 4.93 | 1 | 1 | | | | | | | | | | | | | | | | 30.28 | 4 | 11 | 142 | 17.0 | 6.18 |
| P53985 | 17.80 | 1 | 8 | 8 | 37 | 2.40 | 1 | 1 | 11.00 | 6 | 9 | 9.00 | 3 | 3 | | | | | | | | | | | | | | | | | | | 15.80 | 6 | 24 | 500 | 53.9 | 8.66 |
| P53990 | 18.41 | 1 | 7 | 7 | 20 | 1.92 | 1 | 1 | | | | | | | | | | | | | | | | | | | | | | | | | 18.41 | 7 | 19 | 364 | 39.7 | 5.35 |
| P54107 | 58.23 | 1 | 12 | 12 | 92 | 34.94 | 7 | 8 | 35.74 | 8 | 12 | 24.50 | 4 | 8 | 14.86 | 3 | 3 | 36.55 | 8 | 11 | | | | 4.82 | 1 | 1 | 9.64 | 2 | 2 | 6.02 | 1 | 1 | 54.62 | 9 | 46 | 249 | 28.5 | 5.91 |
| P54108 | 6.94 | 1 | 2 | 2 | 3 | 3.67 | 1 | 1 | | | | 6.94 | 2 | 2 | | | | | | | | | | | | | | | | | | | | | | 245 | 27.6 | 7.80 |
| P54136 | 46.21 | 1 | 30 | 30 | 87 | 2.73 | 1 | 1 | 8.79 | 5 | 5 | 10.30 | 6 | 6 | 5.30 | 3 | 3 | 2.88 | 2 | 2 | | | | 1.36 | 1 | 1 | | | | 9.39 | 5 | 5 | 44.24 | 26 | 64 | 660 | 75.3 | 6.68 |
| P54289 | 4.71 | 1 | 4 | 4 | 4 | | | | | | | 2.36 | 2 | 2 | | | | 0.00 | | | | | | | | | | | | | | | 2.36 | 2 | 2 | 1103 | 124.5 | 5.27 |
| P54577 | 57.77 | 1 | 28 | 28 | 66 | | | | 9.28 | 5 | 5 | 3.79 | 1 | 1 | 3.41 | 2 | 2 | | | | | | | 7.95 | 4 | 4 | 4.55 | 2 | 2 | 17.23 | 9 | 9 | 50.38 | 22 | 43 | 528 | 59.1 | 7.05 |
| P54578 | 50.20 | 1 | 24 | 24 | 97 | 7.09 | 3 | 3 | 10.73 | 5 | 5 | 8.10 | 4 | 4 | 9.11 | 4 | 5 | 2.23 | 1 | 1 | | | | 7.09 | 3 | 3 | 14.98 | 5 | 5 | 18.62 | 7 | 9 | 49.19 | 21 | 62 | 494 | 56.0 | 5.30 |
| P54619 | 32.33 | 2 | 9 | 9 | 15 | | | | | | | | | | 3.02 | 1 | 1 | | | | | | | | | | | | | | | | 29.31 | 8 | 14 | 331 | 37.6 | 6.92 |
| P54652 | 82.94 | 1 | 50 | 72 | 952 | 56.81 | 29 | 65 | 66.82 | 35 | 80 | 62.28 | 37 | 83 | 50.39 | 28 | 47 | 62.44 | 43 | 61 | 13.77 | 6 | 6 | 40.06 | 24 | 35 | 58.69 | 46 | 82 | 57.90 | 41 | 65 | 69.17 | 43 | 428 | 639 | 70.0 | 5.74 |
| P54709 | 47.67 | 1 | 10 | 10 | 39 | 7.53 | 2 | 2 | 23.30 | 6 | 7 | 7.53 | 2 | 2 | | | | | | | | | | | | | | | | | | | 42.29 | 8 | 28 | 279 | 31.5 | 8.35 |
| P54727 | 27.87 | 1 | 8 | 9 | 16 | | | | 9.29 | 3 | 4 | 4.16 | 2 | 2 | | | | 2.69 | 1 | 1 | | | | | | | | | | | | | 21.03 | 6 | 9 | 409 | 43.1 | 4.84 |
| P54802 | 10.77 | 1 | 5 | 5 | 8 | | | | | | | | | | | | | | | | | | | | | | | | | | | | 10.77 | 5 | 8 | 743 | 82.2 | 6.65 |
| P54803 | 4.09 | 1 | 2 | 2 | 2 | | | | | | | | | | | | | | | | | | | | | | | | | | | | 4.09 | 2 | 2 | 685 | 77.0 | 6.64 |
| P54819 | 37.66 | 1 | 7 | 7 | 11 | 23.43 | 3 | 3 | 10.04 | 2 | 2 | | | | | | | | | | | | | | | | | | | | | | 20.08 | 3 | 6 | 239 | 26.5 | 7.81 |
| P54868 | 7.48 | 1 | 3 | 3 | 3 | | | | | | | | | | | | | | | | | | | | | | | | | | | | 7.48 | 3 | 3 | 508 | 56.6 | 8.16 |
| P54886 | 10.06 | 1 | 7 | 7 | 7 | | | | | | | | | | | | | | | | | | | | | | | | | | | | 10.06 | 7 | 7 | 795 | 87.2 | 7.12 |
| P54920 | 78.64 | 2 | 21 | 21 | 70 | 17.29 | 4 | 4 | 18.98 | 4 | 4 | 7.80 | 2 | 2 | 20.34 | 5 | 6 | 6.10 | 1 | 1 | 3.39 | 1 | 1 | | | | | | | 26.78 | 6 | 7 | 73.90 | 20 | 45 | 295 | 33.2 | 5.36 |
| P55010 | 7.19 | 1 | 3 | 3 | 6 | | | | 2.09 | 1 | 1 | | | | | | | | | | | | | | | | | | | | | | 7.19 | 3 | 5 | 431 | 49.2 | 5.58 |
| P55036 | 42.44 | 1 | 12 | 12 | 40 | 9.81 | 3 | 3 | 10.88 | 3 | 3 | 8.22 | 2 | 2 | 8.22 | 2 | 2 | 21.75 | 5 | 5 | | | | | | | 3.71 | 1 | 1 | 4.51 | 1 | 1 | 35.54 | 10 | 23 | 377 | 40.7 | 4.79 |
| P55039 | 25.00 | 1 | 8 | 8 | 17 | | | | | | | | | | | | | | | | | | | | | | | | | | | | 25.00 | 8 | 17 | 364 | 40.7 | 8.88 |
| P55060 | 44.28 | 1 | 39 | 39 | 107 | 5.66 | 4 | 4 | 6.90 | 5 | 6 | 3.50 | 2 | 2 | 3.19 | 3 | 3 | 0.00 | | | | | | | | | | | | | | | 43.36 | 38 | 92 | 971 | 110.3 | 5.77 |
| P55061 | 3.80 | 1 | 2 | 2 | 3 | 3.38 | 1 | 1 | | | | | | | | | | | | | | | | | | | | | | | | | 3.80 | 2 | 2 | 237 | 26.5 | 8.54 |
| P55064 | 15.85 | 1 | 5 | 5 | 40 | 9.81 | 2 | 2 | 6.04 | 2 | 3 | 6.04 | 1 | 2 | 6.04 | 1 | 2 | | | | | | | | | | | | | | | | 15.85 | 5 | 31 | 265 | 28.3 | 8.62 |
| P55072 | 72.21 | 2 | 65 | 65 | 465 | 42.93 | 25 | 32 | 43.67 | 35 | 46 | 37.10 | 25 | 30 | 34.12 | 22 | 27 | 30.40 | 20 | 22 | 2.36 | 2 | 2 | 16.75 | 12 | 14 | 16.50 | 12 | 14 | 22.83 | 14 | 16 | 69.98 | 51 | 262 | 806 | 89.3 | 5.26 |
| P55083 | 19.61 | 1 | 4 | 4 | 12 | 9.41 | 2 | 2 | 9.41 | 2 | 2 | 7.06 | 1 | 3 | | | | | | | | | | | | | | | | | | | 19.61 | 4 | 5 | 255 | 28.6 | 5.63 |
| P55084 | 73.00 | 1 | 34 | 34 | 165 | 35.65 | 16 | 19 | 43.25 | 20 | 27 | 42.62 | 18 | 25 | 6.12 | 2 | 2 | 9.92 | 2 | 2 | | | | 4.43 | 2 | 2 | | | | 5.70 | 1 | 1 | 69.83 | 30 | 87 | 474 | 51.3 | 9.41 |
| P55145 | 54.95 | 1 | 13 | 13 | 34 | 40.11 | 7 | 9 | 13.19 | 3 | 3 | 34.07 | 6 | 6 | | | | 16.48 | 3 | 3 | | | | | | | | | | | | | 38.46 | 9 | 13 | 182 | 20.7 | 8.69 |
| P55157 | 5.03 | 1 | 4 | 4 | 4 | | | | | | | 1.23 | 1 | 1 | | | | | | | | | | | | | | | | | | | 3.80 | 3 | 3 | 894 | 99.3 | 8.41 |
| P55196 | 1.92 | 1 | 2 | 2 | 2 | | | | | | | | | | | | | | | | | | | | | | | | | 0.93 | 1 | 1 | 0.99 | 1 | 1 | 1824 | 206.7 | 6.47 |
| P55209 | 15.09 | 1 | 5 | 5 | 10 | 2.81 | 1 | 1 | 2.81 | 1 | 1 | | | | | | | | | | | | | | | | | | | 7.42 | 2 | 2 | 15.09 | 4 | 6 | 391 | 45.3 | 4.46 |
| P55210 | 10.23 | 1 | 2 | 2 | 4 | | | | | | | | | | | | | | | | | | | | | | | | | | | | 10.23 | 2 | 4 | 303 | 34.3 | 6.07 |
| P55212 | 10.24 | 1 | 2 | 3 | 5 | | | | | | | | | | | | | | | | | | | | | | | | | | | | 10.24 | 3 | 5 | 293 | 33.3 | 6.93 |
| P55259 | 13.97 | 4 | 7 | 8 | 11 | | | | 2.05 | 2 | 2 | 13.97 | 8 | 8 | 1.12 | 1 | 1 | | | | | | | | | | | | | | | | 0.00 | | | 537 | 59.4 | 5.24 |
| P55263 | 18.23 | 1 | 5 | 5 | 6 | | | | | | | | | | | | | | | | | | | | | | | | | | | | 18.23 | 5 | 6 | 362 | 40.5 | 6.70 |
| P55268 | 38.38 | 3 | 53 | 54 | 184 | 17.35 | 23 | 24 | 6.51 | 11 | 11 | 31.09 | 40 | 47 | 4.12 | 5 | 5 | 2.22 | 2 | 2 | | | | | | | | | | | | | 27.25 | 37 | 95 | 1798 | 195.9 | 6.52 |
| P55327 | 24.11 | 1 | 3 | 3 | 4 | | | | | | | | | | | | | | | | | | | | | | | | | | | | 24.11 | 3 | 3 | 224 | 24.3 | 4.83 |
| P55735 | 40.37 | 1 | 9 | 9 | 31 | 8.07 | 1 | 2 | 8.07 | 1 | 1 | 8.07 | 1 | 1 | 8.07 | 1 | 1 | | | | | | | 8.07 | 1 | 1 | 3.73 | 1 | 1 | 11.80 | 2 | 2 | 40.06 | 8 | 22 | 322 | 35.5 | 5.48 |
| P55769 | 18.75 | 1 | 2 | 2 | 2 | | | | | | | | | | | | | | | | | | | | | | | | | | | | 18.75 | 2 | 2 | 128 | 14.2 | 8.46 |
| P55786 | 46.25 | 2 | 38 | 38 | 126 | 7.94 | 7 | 7 | 12.95 | 12 | 14 | 13.49 | 13 | 13 | 5.11 | 5 | 5 | 1.52 | 1 | 1 | | | | 1.41 | 1 | 1 | | | | 4.35 | 3 | 3 | 44.94 | 35 | 82 | 919 | 103.2 | 5.72 |
| P55789 | 32.68 | 1 | 7 | 7 | 19 | 5.85 | 1 | 1 | 18.05 | 2 | 2 | | | | | | | | | | | | | | | | | | | | | | 32.68 | 7 | 16 | 205 | 23.4 | 7.62 |
| P55809 | 45.77 | 1 | 17 | 18 | 87 | 30.19 | 10 | 13 | 33.27 | 10 | 18 | 28.85 | 11 | 16 | 2.12 | 1 | 1 | 11.54 | 4 | 5 | | | | | | | | | | | | | 37.31 | 15 | 34 | 520 | 56.1 | 7.46 |
| P55884 | 26.17 | 1 | 18 | 18 | 53 | 1.47 | 1 | 1 | 3.19 | 2 | 2 | 1.72 | 1 | 1 | 1.72 | 1 | 1 | 1.72 | 1 | 1 | | | | | | | | | | | | | 26.17 | 17 | 47 | 814 | 92.4 | 5.00 |
| P56134 | 41.49 | 1 | 4 | 4 | 17 | 11.70 | 1 | 1 | 25.53 | 2 | 2 | 11.70 | 1 | 1 | | | | | | | | | | | | | | | | | | | 41.49 | 4 | 13 | 94 | 10.9 | 9.67 |
| Q9ULC5 | 4.83 | 1 | 1 | 3 | 23 | 1.61 | 1 | 1 | 2.93 | 2 | 2 | 2.93 | 2 | 2 | 1.61 | 1 | 1 | 1.61 | 1 | 1 | | | | 1.61 | 1 | 2 | | | | 1.61 | 1 | 1 | 4.83 | 3 | 13 | 683 | 75.9 | 6.92 |
| P56180 | 13.43 | 2 | 7 | 7 | 17 | 8.35 | 4 | 4 | 3.63 | 2 | 2 | 4.72 | 2 | 3 | | | | | | | | | | | | | | | | | | | 11.80 | 6 | 9 | 551 | 64.3 | 8.68 |
| P56192 | 42.89 | 1 | 27 | 27 | 100 | 7.11 | 5 | 5 | 11.89 | 8 | 8 | 5.33 | 4 | 4 | | | | | | | | | | | | | | | | | | | 40.22 | 25 | 83 | 900 | 101.1 | 6.16 |
| P56385 | 40.58 | 1 | 3 | 3 | 7 | 30.43 | 2 | 2 | 14.49 | 1 | 1 | | | | | | | | | | | | | | | | | | | | | | 40.58 | 3 | 4 | 69 | 7.9 | 9.35 |
| P56537 | 56.73 | 1 | 8 | 8 | 21 | 4.08 | 1 | 1 | 5.71 | 1 | 1 | | | | | | | | | | | | | 5.71 | 1 | 1 | 13.06 | 2 | 2 | 9.80 | 2 | 2 | 52.65 | 7 | 14 | 245 | 26.6 | 4.68 |
| P56556 | 12.34 | 1 | 2 | 2 | 4 | | | | 5.19 | 1 | 1 | | | | | | | | | | | | | | | | | | | | | | 12.34 | 2 | 3 | 154 | 17.9 | 10.14 |
| P56557 | 12.03 | 1 | 2 | 2 | 4 | | | | | | | | | | | | | | | | | | | | | | | | | | | | 12.03 | 2 | 4 | 158 | 17.9 | 5.39 |
| P56589 | 16.09 | 1 | 5 | 5 | 9 | | | | 2.68 | 1 | 1 | | | | | | | | | | | | | | | | | | | | | | 16.09 | 5 | 8 | 373 | 42.1 | 8.15 |
| P56597 | 79.72 | 1 | 11 | 11 | 41 | 17.92 | 2 | 2 | 12.26 | 2 | 2 | | | | 25.94 | 5 | 6 | 5.19 | 1 | 1 | | | | | | | | | | | | | 79.72 | 11 | 30 | 212 | 24.2 | 6.29 |
| P56730 | 3.89 | 1 | 3 | 3 | 4 | | | | | | | | | | | | | | | | | | | | | | | | | | | | 3.89 | 3 | 4 | 875 | 97.0 | 8.03 |
| P56851 | 54.42 | 1 | 7 | 8 | 26 | 7.48 | 1 | 1 | 7.48 | 1 | 1 | 22.45 | 2 | 2 | 7.48 | 1 | 1 | 18.37 | 2 | 3 | | | | 7.48 | 1 | 1 | | | | | | | 53.06 | 7 | 17 | 147 | 17.6 | 6.99 |
| P57088 | 13.36 | 1 | 3 | 3 | 6 | | | | | | | 0.00 | | | | | | | | | | | | | | | | | | | | | 13.36 | 3 | 6 | 247 | 28.0 | 9.70 |
| P57105 | 51.03 | 1 | 8 | 8 | 21 | | | | 6.21 | 1 | 1 | | | | | | | | | | | | | | | | | | | | | | 51.03 | 8 | 20 | 145 | 15.9 | 6.30 |
| P57678 | 41.59 | 1 | 33 | 34 | 81 | | | | | | | | | | | | | | | | | | | | | | | | | | | | 41.59 | 34 | 81 | 1058 | 120.0 | 6.04 |
| P57735 | 26.29 | 1 | 4 | 5 | 12 | 3.76 | 1 | 1 | 0.00 | | | | | | | | | | | | | | | | | | | | | | | | 26.29 | 5 | 11 | 213 | 23.5 | 5.96 |
| P57740 | 33.73 | 1 | 21 | 21 | 33 | | | | | | | | | | | | | | | | | | | | | | | | | | | | 33.73 | 21 | 33 | 925 | 106.3 | 5.43 |
| P57764 | 4.75 | 1 | 2 | 2 | 5 | | | | | | | | | | | | | | | | | | | | | | | | | | | | 4.75 | 2 | 5 | 484 | 52.8 | 5.08 |
| P57772 | 11.58 | 1 | 5 | 5 | 8 | | | | | | | | | | | | | | | | | | | | | | | | | | | | 11.58 | 5 | 8 | 596 | 65.3 | 8.35 |
| P58340 | 83.96 | 1 | 29 | 29 | 156 | 14.55 | 3 | 3 | 30.60 | 6 | 9 | 33.96 | 7 | 10 | 51.12 | 11 | 16 | 66.79 | 17 | 26 | 4.10 | 1 | 1 | 19.40 | 5 | 6 | 33.96 | 11 | 17 | 47.76 | 14 | 19 | 60.07 | 13 | 49 | 268 | 30.6 | 9.45 |
| P58499 | 11.49 | 1 | 2 | 2 | 3 | | | | 3.83 | 1 | 1 | | | | | | | | | | | | | | | | | | | | | | 7.66 | 1 | 2 | 235 | 26.0 | 8.75 |
| B2RC85 | 20.00 | 1 | 1 | 13 | 23 | | | | | | | | | | | | | | | | | | | | | | | | | | | | 20.00 | 13 | 23 | 870 | 100.5 | 7.49 |
| P0C881 | 20.00 | 1 | 1 | 13 | 23 | | | | | | | | | | | | | | | | | | | | | | | | | | | | 20.00 | 13 | 23 | 870 | 100.6 | 7.49 |
| P58546 | 49.15 | 1 | 7 | 7 | 18 | 20.34 | 2 | 2 | 14.41 | 1 | 1 | | | | 14.41 | 1 | 1 | | | | | | | 22.03 | 2 | 2 | 33.90 | 4 | 4 | 26.27 | 3 | 3 | 41.53 | 4 | 5 | 118 | 12.9 | 5.52 |
| P58876 | 57.14 | 9 | 2 | 8 | 62 | 33.33 | 4 | 8 | 40.48 | 5 | 10 | 40.48 | 5 | 7 | 22.22 | 4 | 5 | 12.70 | 2 | 2 | 5.56 | 1 | 1 | 8.73 | 1 | 1 | | | | 8.73 | 1 | 1 | 57.14 | 8 | 27 | 126 | 13.9 | 10.32 |
| P59282 | 57.06 | 1 | 9 | 9 | 24 | 48.24 | 6 | 6 | 38.24 | 4 | 4 | 43.53 | 4 | 4 | | | | 12.35 | 2 | 2 | | | | | | | 11.18 | 1 | 1 | | | | 42.94 | 5 | 7 | 170 | 18.5 | 9.00 |
| P59665 | 26.60 | 2 | 5 | 5 | 42 | 26.60 | 5 | 6 | | | | 20.21 | 4 | 5 | 20.21 | 4 | 6 | 20.21 | 4 | 5 | 9.57 | 1 | 1 | | | | 20.21 | 2 | 2 | 20.21 | 4 | 6 | 19.15 | 2 | 11 | 94 | 10.2 | 6.99 |
| P59780 | 16.06 | 1 | 2 | 2 | 2 | | | | | | | | | | | | | | | | | | | | | | | | | | | | 16.06 | 2 | 2 | 193 | 22.0 | 5.22 |
| P59910 | 67.41 | 1 | 20 | 20 | 81 | 32.28 | 6 | 7 | 18.99 | 4 | 5 | 2.22 | 1 | 1 | 26.90 | 8 | 8 | 48.42 | 12 | 14 | | | | | | | 14.24 | 4 | 4 | 3.80 | 1 | 1 | 52.85 | 15 | 41 | 316 | 36.1 | 7.87 |
| P59998 | 51.19 | 1 | 7 | 7 | 13 | 14.88 | 2 | 2 | | | | 7.14 | 1 | 1 | 13.69 | 2 | 2 | | | | | | | | | | | | | | | | 35.71 | 5 | 8 | 168 | 19.7 | 8.43 |
| P60033 | 15.25 | 1 | 2 | 2 | 4 | 6.78 | 1 | 1 | | | | 8.47 | 1 | 1 | | | | | | | | | | | | | | | | | | | 15.25 | 2 | 2 | 236 | 25.8 | 5.29 |
| P60174 | 94.76 | 2 | 32 | 32 | 503 | 67.48 | 15 | 20 | 61.89 | 14 | 22 | 70.28 | 17 | 22 | 71.68 | 19 | 28 | 66.08 | 18 | 28 | 8.74 | 2 | 2 | 67.13 | 19 | 38 | 76.57 | 24 | 59 | 75.52 | 23 | 41 | 94.06 | 27 | 243 | 286 | 30.8 | 5.92 |
| P60228 | 51.91 | 1 | 20 | 20 | 44 | 4.27 | 1 | 1 | | | | | | | | | | | | | | | | | | | | | | | | | 51.91 | 20 | 43 | 445 | 52.2 | 6.04 |
| P60468 | 37.50 | 1 | 3 | 3 | 6 | 27.08 | 2 | 2 | 26.04 | 2 | 2 | | | | | | | | | | | | | | | | | | | | | | 27.08 | 2 | 2 | 96 | 10.0 | 11.56 |
| P60484 | 4.47 | 1 | 2 | 2 | 3 | | | | | | | | | | | | | | | | | | | | | | | | | | | | 4.47 | 2 | 3 | 403 | 47.1 | 6.37 |
| P60510 | 31.27 | 1 | 6 | 8 | 17 | | | | | | | | | | 4.89 | 1 | 1 | | | | | | | | | | | | | | | | 31.27 | 8 | 16 | 307 | 35.1 | 5.06 |
| P60660 | 69.54 | 4 | 11 | 13 | 73 | 46.36 | 6 | 7 | 46.36 | 6 | 6 | 37.75 | 5 | 5 | 39.74 | 7 | 9 | 41.06 | 5 | 5 | | | | 23.84 | 4 | 4 | 41.06 | 6 | 7 | 58.28 | 8 | 10 | 66.23 | 9 | 20 | 151 | 16.9 | 4.65 |
| P60673 | 34.31 | 1 | 6 | 6 | 13 | 5.84 | 1 | 1 | 5.84 | 1 | 1 | | | | | | | | | | | | | | | | | | | | | | 34.31 | 6 | 11 | 137 | 14.6 | 9.26 |
| P60842 | 64.29 | 1 | 15 | 29 | 132 | 10.34 | 4 | 4 | 10.59 | 4 | 4 | 8.62 | 3 | 3 | 16.50 | 5 | 5 | 6.40 | 2 | 2 | | | | 15.52 | 5 | 5 | 9.85 | 3 | 3 | 23.15 | 8 | 9 | 63.79 | 25 | 97 | 406 | 46.1 | 5.48 |
| P60866 | 28.57 | 1 | 4 | 4 | 14 | 15.13 | 2 | 2 | 9.24 | 1 | 1 | 9.24 | 1 | 1 | 9.24 | 1 | 1 | | | | | | | 9.24 | 1 | 1 | | | | | | | 28.57 | 4 | 8 | 119 | 13.4 | 9.94 |
| P60891 | 50.94 | 1 | 6 | 16 | 56 | 7.23 | 2 | 2 | | | | 5.35 | 1 | 1 | 11.32 | 3 | 3 | | | | | | | 2.52 | 1 | 1 | | | | 15.09 | 3 | 5 | 49.37 | 13 | 44 | 318 | 34.8 | 6.98 |
| P60900 | 53.66 | 1 | 12 | 12 | 53 | 41.06 | 8 | 8 | 37.40 | 7 | 8 | 44.72 | 9 | 11 | 17.48 | 3 | 3 | | | | | | | | | | 9.35 | 2 | 2 | 14.23 | 3 | 4 | 43.09 | 10 | 17 | 246 | 27.4 | 6.76 |
| P60903 | 24.74 | 1 | 2 | 2 | 3 | 17.53 | 1 | 1 | | | | 7.22 | 1 | 1 | | | | | | | | | | | | | | | | 17.53 | 1 | 1 | | | | 97 | 11.2 | 7.37 |
| P60953 | 62.83 | 3 | 8 | 9 | 37 | 14.14 | 2 | 3 | | | | 36.65 | 5 | 5 | 16.75 | 3 | 3 | 10.99 | 2 | 2 | | | | 5.76 | 1 | 1 | 5.24 | 1 | 1 | 33.51 | 5 | 5 | 56.02 | 8 | 17 | 191 | 21.2 | 6.55 |
| P60981 | 63.64 | 1 | 14 | 15 | 55 | 12.73 | 2 | 2 | 15.76 | 3 | 3 | 22.42 | 5 | 5 | 21.21 | 3 | 4 | 18.79 | 3 | 3 | | | | 20.00 | 4 | 4 | 19.39 | 3 | 3 | 32.12 | 7 | 8 | 55.76 | 11 | 23 | 165 | 18.5 | 7.85 |
| P61006 | 48.79 | 7 | 4 | 12 | 88 | 24.64 | 4 | 5 | 30.43 | 5 | 6 | 27.54 | 5 | 5 | 26.09 | 5 | 5 | | | | | | | 18.36 | 4 | 4 | 13.53 | 4 | 4 | 26.57 | 5 | 5 | 47.83 | 11 | 52 | 207 | 23.7 | 9.07 |
| P61009 | 30.56 | 1 | 6 | 6 | 17 | 11.67 | 2 | 2 | 5.00 | 1 | 1 | 5.00 | 1 | 1 | | | | | | | | | | | | | | | | | | | 30.56 | 6 | 13 | 180 | 20.3 | 8.62 |
| P61018 | 60.09 | 6 | 7 | 10 | 41 | 28.17 | 4 | 5 | 15.96 | 3 | 3 | 5.16 | 1 | 1 | 5.16 | 1 | 1 | | | | | | | 5.16 | 1 | 1 | | | | | | | 60.09 | 10 | 30 | 213 | 23.6 | 6.06 |
| P61019 | 76.89 | 1 | 11 | 20 | 172 | 61.32 | 14 | 17 | 65.57 | 15 | 21 | 61.32 | 13 | 16 | 29.72 | 5 | 6 | 6.60 | 1 | 1 | | | | 9.91 | 2 | 2 | 6.60 | 1 | 1 | 6.60 | 1 | 1 | 76.42 | 19 | 107 | 212 | 23.5 | 6.54 |
| P61020 | 60.93 | 1 | 6 | 9 | 22 | 11.16 | 2 | 2 | 11.63 | 2 | 2 | 5.12 | 1 | 1 | | | | | | | | | | | | | | | | | | | 56.28 | 8 | 17 | 215 | 23.7 | 8.13 |
| P61026 | 46.50 | 7 | 8 | 11 | 61 | 26.50 | 6 | 9 | 23.00 | 4 | 6 | 23.00 | 4 | 4 | 11.50 | 2 | 2 | | | | | | | 5.50 | 1 | 1 | 5.50 | 1 | 1 | 5.50 | 1 | 1 | 46.50 | 11 | 37 | 200 | 22.5 | 8.38 |
| P61077 | 36.05 | 2 | 2 | 3 | 4 | | | | | | | | | | | | | | | | | | | | | | | | | | | | 36.05 | 3 | 4 | 147 | 16.7 | 7.80 |
| P61081 | 32.79 | 1 | 6 | 6 | 14 | 12.02 | 2 | 3 | | | | 4.37 | 1 | 1 | | | | | | | | | | | | | | | | 4.37 | 1 | 1 | 32.79 | 6 | 9 | 183 | 20.9 | 7.69 |
| P61086 | 13.50 | 1 | 2 | 2 | 6 | | | | | | | | | | | | | | | | | | | | | | | | | | | | 13.50 | 2 | 6 | 200 | 22.4 | 5.44 |
| P61088 | 46.71 | 3 | 6 | 6 | 19 | 4.61 | 1 | 1 | 7.24 | 1 | 1 | 7.24 | 1 | 1 | 18.42 | 3 | 3 | | | | | | | 7.24 | 1 | 1 | 7.24 | 1 | 1 | 7.24 | 1 | 1 | 46.71 | 6 | 10 | 152 | 17.1 | 6.57 |
| P61106 | 91.16 | 6 | 19 | 20 | 88 | 30.23 | 6 | 8 | 43.72 | 8 | 10 | 16.74 | 3 | 3 | 5.12 | 1 | 1 | 6.98 | 1 | 1 | | | | 5.12 | 1 | 1 | 6.98 | 1 | 1 | 5.12 | 1 | 1 | 91.16 | 18 | 60 | 215 | 23.9 | 6.21 |
| P61158 | 48.56 | 3 | 15 | 15 | 41 | 8.85 | 4 | 4 | 2.15 | 1 | 1 | 18.66 | 4 | 4 | 12.44 | 4 | 4 | 4.07 | 1 | 1 | | | | | | | | | | | | | 42.34 | 12 | 27 | 418 | 47.3 | 5.88 |
| P61160 | 28.93 | 1 | 10 | 10 | 37 | 3.05 | 1 | 1 | 5.84 | 2 | 2 | 7.36 | 3 | 3 | 2.79 | 1 | 1 | | | | | | | | | | | | | 9.14 | 3 | 4 | 28.93 | 10 | 26 | 394 | 44.7 | 6.74 |
| P61163 | 61.17 | 1 | 10 | 18 | 69 | 34.31 | 9 | 11 | 31.65 | 8 | 11 | 34.31 | 7 | 8 | 12.50 | 3 | 3 | | | | | | | | | | | | | | | | 47.34 | 14 | 36 | 376 | 42.6 | 6.64 |
| P61201 | 23.93 | 1 | 9 | 9 | 14 | | | | | | | | | | | | | | | | | | | | | | | | | | | | 23.93 | 9 | 14 | 443 | 51.6 | 5.53 |
| P61204 | 70.72 | 4 | 7 | 14 | 78 | 35.36 | 5 | 7 | 41.44 | 8 | 10 | 43.09 | 6 | 7 | 31.49 | 4 | 4 | | | | | | | 13.81 | 2 | 2 | | | | 7.73 | 1 | 1 | 57.46 | 12 | 45 | 181 | 20.6 | 7.43 |
| P61221 | 20.03 | 1 | 10 | 10 | 18 | | | | | | | 1.00 | 1 | 1 | | | | | | | | | | | | | | | | 2.00 | 2 | 2 | 19.03 | 8 | 15 | 599 | 67.3 | 8.34 |
| P61224 | 53.26 | 2 | 2 | 11 | 32 | 12.50 | 2 | 2 | 12.50 | 2 | 2 | 25.54 | 4 | 4 | | | | | | | | | | 5.98 | 1 | 1 | | | | 5.98 | 1 | 1 | 53.26 | 11 | 22 | 184 | 20.8 | 5.78 |
| P61225 | 17.49 | 2 | 2 | 3 | 3 | | | | | | | | | | | | | | | | | | | | | | | | | | | | 17.49 | 3 | 3 | 183 | 20.5 | 4.81 |
| P61247 | 62.12 | 1 | 18 | 18 | 58 | 17.42 | 4 | 4 | 15.15 | 4 | 4 | | | | 16.67 | 4 | 4 | 3.79 | 1 | 1 | | | | 3.41 | 1 | 1 | | | | 21.59 | 5 | 5 | 56.82 | 15 | 37 | 264 | 29.9 | 9.73 |
| P61254 | 51.03 | 1 | 2 | 12 | 23 | | | | 5.52 | 1 | 1 | | | | | | | | | | | | | | | | | | | | | | 51.03 | 12 | 22 | 145 | 17.2 | 10.55 |

| | | | | | | | | | | | | | | | | | | | | | | | | | | | | | | | | | | | | | | |
|---|---|---|---|---|---|---|---|---|---|---|---|---|---|---|---|---|---|---|---|---|---|---|---|---|---|---|---|---|---|---|---|---|---|---|---|---|---|---|
| P61313 | 42.16 | 1 | 9 | 9 | 24 | 12.75 | 2 | 2 | 20.59 | 3 | 4 | | | | | | | | | | | | | | | | | | | 6.86 | 1 | 1 | 36.27 | 8 | 17 | 204 | 24.1 | 11.62 |
| P61353 | 43.38 | 1 | 5 | 5 | 12 | | | | 6.62 | 1 | 1 | | | | | | | | | | | | | 5.88 | 1 | 1 | | | | | | | 43.38 | 5 | 10 | 136 | 15.8 | 10.56 |
| P61421 | 43.59 | 2 | 11 | 11 | 24 | 13.11 | 3 | 3 | | | | 3.70 | 1 | 1 | 5.41 | 2 | 2 | | | | | | | | | | | | | | | | 43.59 | 11 | 18 | 351 | 40.3 | 5.00 |
| P61513 | 50.00 | 1 | 4 | 4 | 5 | | | | | | | | | | | | | | | | | | | | | | | | | | | | 50.00 | 4 | 5 | 92 | 10.3 | 10.43 |
| P61586 | 79.79 | 1 | 5 | 14 | 75 | 35.75 | 5 | 5 | 8.81 | 1 | 1 | 25.39 | 4 | 4 | 37.82 | 8 | 9 | 18.65 | 4 | 5 | | | | 17.62 | 2 | 2 | 33.68 | 6 | 9 | 24.87 | 5 | 5 | 56.48 | 10 | 35 | 193 | 21.8 | 6.10 |
| P61604 | 75.49 | 1 | 11 | 11 | 40 | 50.98 | 6 | 6 | 57.84 | 7 | 9 | 60.78 | 7 | 8 | 32.35 | 4 | 4 | 39.22 | 5 | 5 | | | | | | | 6.86 | 1 | 1 | | | | 58.82 | 7 | 7 | 102 | 10.9 | 8.92 |
| P61626 | 62.16 | 2 | 9 | 9 | 25 | 56.76 | 7 | 8 | 10.81 | 2 | 2 | 56.76 | 7 | 8 | 12.16 | 2 | 2 | | | | | | | 8.11 | 1 | 1 | | | | | | | 35.81 | 4 | 4 | 148 | 16.5 | 9.16 |
| P61758 | 25.89 | 1 | 2 | 3 | 6 | 22.34 | 2 | 2 | | | | | | | | | | 3.55 | 1 | 1 | | | | | | | | | | | | | 3.55 | 1 | 3 | 197 | 22.6 | 7.11 |
| P61769 | 57.98 | 2 | 8 | 8 | 66 | 35.29 | 3 | 6 | 40.34 | 4 | 6 | 35.29 | 3 | 7 | 21.85 | 3 | 5 | 49.58 | 6 | 11 | 8.40 | 1 | 1 | 21.85 | 3 | 5 | 26.05 | 4 | 7 | 39.50 | 6 | 9 | 40.34 | 4 | 9 | 119 | 13.7 | 6.52 |
| P61803 | 35.40 | 1 | 4 | 4 | 8 | | | | 0.00 | | | | | | | | | | | | | | | | | | | | | | | | 35.40 | 4 | 8 | 113 | 12.5 | 7.08 |
| P61916 | 50.33 | 1 | 7 | 7 | 33 | 32.45 | 3 | 3 | 18.54 | 3 | 3 | 44.37 | 5 | 8 | 15.23 | 2 | 2 | 21.85 | 3 | 3 | | | | | | | | | | | | | 43.05 | 5 | 14 | 151 | 16.6 | 7.65 |
| P61923 | 41.81 | 1 | 6 | 6 | 13 | | | | | | | | | | | | | | | | | | | | | | | | | | | | 41.81 | 6 | 13 | 177 | 20.2 | 4.81 |
| P61962 | 18.42 | 1 | 6 | 6 | 7 | 2.63 | 1 | 1 | | | | | | | | | | | | | | | | | | | | | | | | | 15.79 | 5 | 6 | 342 | 38.9 | 5.52 |
| P61966 | 41.77 | 2 | 4 | 6 | 17 | | | | | | | | | | | | | | | | | | | | | | | | | | | | 41.77 | 6 | 17 | 158 | 18.7 | 5.73 |
| P61970 | 53.54 | 1 | 5 | 5 | 23 | 35.43 | 4 | 5 | 24.41 | 3 | 3 | 18.11 | 2 | 2 | 5.51 | 1 | 1 | 5.51 | 1 | 1 | | | | 5.51 | 1 | 1 | | | | 22.83 | 3 | 3 | 47.24 | 4 | 7 | 127 | 14.5 | 5.38 |
| P61978 | 29.59 | 1 | 11 | 11 | 34 | 2.38 | 1 | 1 | 6.70 | 2 | 2 | 2.59 | 1 | 1 | 11.02 | 4 | 4 | 9.50 | 3 | 3 | | | | 3.67 | 1 | 2 | 9.50 | 3 | 3 | 9.50 | 3 | 3 | 20.52 | 7 | 15 | 463 | 50.9 | 5.54 |
| P61981 | 68.42 | 1 | 11 | 18 | 126 | 34.82 | 7 | 10 | 34.82 | 7 | 9 | 23.89 | 6 | 7 | 29.15 | 7 | 9 | 25.10 | 7 | 9 | 3.64 | 1 | 1 | 34.41 | 8 | 11 | 20.65 | 5 | 7 | 38.06 | 10 | 16 | 57.09 | 13 | 47 | 247 | 28.3 | 4.89 |
| P62070 | 22.06 | 1 | 3 | 4 | 9 | | | | | | | | | | | | | | | | | | | | | | | | | | | | 22.06 | 4 | 9 | 204 | 23.4 | 6.01 |
| P62081 | 60.82 | 1 | 10 | 10 | 24 | 5.67 | 1 | 1 | 10.31 | 2 | 2 | | | | | | | | | | | | | | | | | | | | | | 60.82 | 10 | 21 | 194 | 22.1 | 10.10 |
| P62140 | 66.67 | 1 | 7 | 19 | 114 | 31.19 | 8 | 10 | 15.60 | 4 | 4 | 22.32 | 6 | 7 | 11.01 | 4 | 5 | 4.28 | 1 | 1 | | | | 7.34 | 2 | 2 | 4.28 | 1 | 1 | 16.82 | 5 | 6 | 65.44 | 17 | 78 | 327 | 37.2 | 6.19 |
| P62158 | 67.11 | 3 | 12 | 12 | 80 | 43.62 | 7 | 13 | 44.97 | 8 | 16 | 38.93 | 6 | 10 | 34.90 | 6 | 10 | 30.20 | 4 | 5 | | | | 5.37 | 1 | 1 | 11.41 | 1 | 1 | | | | 46.31 | 8 | 24 | 149 | 16.8 | 4.22 |
| P62191 | 53.41 | 1 | 19 | 21 | 110 | 30.23 | 11 | 12 | 32.95 | 11 | 14 | 9.55 | 3 | 3 | 10.00 | 4 | 5 | 5.91 | 2 | 2 | | | | 16.82 | 7 | 8 | 10.68 | 4 | 4 | 12.73 | 5 | 5 | 45.68 | 17 | 57 | 440 | 49.2 | 6.21 |
| A6NHN6 | 20.77 | 2 | 1 | 10 | 21 | | | | 2.48 | 1 | 1 | | | | | | | 2.03 | 1 | 1 | | | | | | | | | | | | | 18.28 | 8 | 19 | 443 | 51.2 | 10.49 |
| P62195 | 71.92 | 1 | 26 | 28 | 131 | 26.60 | 7 | 10 | 41.13 | 13 | 16 | 20.44 | 6 | 6 | 7.14 | 3 | 3 | 12.56 | 4 | 4 | | | | 21.43 | 8 | 9 | 14.53 | 5 | 7 | 39.90 | 15 | 15 | 61.82 | 20 | 61 | 406 | 45.6 | 7.55 |
| P62241 | 58.17 | 1 | 10 | 10 | 41 | 11.54 | 2 | 2 | 18.75 | 3 | 3 | 18.75 | 3 | 3 | 18.27 | 3 | 3 | 18.27 | 3 | 3 | | | | 4.33 | 1 | 1 | | | | 14.42 | 3 | 3 | 47.60 | 8 | 23 | 208 | 24.2 | 10.32 |
| P62244 | 66.15 | 1 | 9 | 9 | 24 | 16.15 | 3 | 3 | 17.69 | 3 | 3 | | | | 17.69 | 3 | 3 | | | | | | | | | | | | | 4.62 | 1 | 1 | 66.15 | 9 | 14 | 130 | 14.8 | 10.13 |
| P62249 | 56.85 | 1 | 10 | 10 | 15 | | | | 13.01 | 2 | 2 | 7.53 | 1 | 1 | | | | | | | | | | | | | | | | | | | 55.48 | 9 | 12 | 146 | 16.4 | 10.21 |
| P62258 | 80.39 | 1 | 26 | 29 | 228 | 42.35 | 9 | 10 | 34.90 | 7 | 10 | 51.76 | 12 | 14 | 67.84 | 17 | 21 | 36.08 | 8 | 10 | 4.71 | 1 | 1 | 58.43 | 17 | 22 | 32.55 | 8 | 10 | 58.43 | 16 | 27 | 80.39 | 27 | 103 | 255 | 29.2 | 4.74 |
| P62263 | 41.06 | 1 | 9 | 9 | 23 | | | | 13.91 | 1 | 1 | 7.28 | 1 | 1 | 13.91 | 2 | 2 | | | | | | | 7.28 | 1 | 1 | | | | | | | 41.06 | 9 | 18 | 151 | 16.3 | 10.05 |
| P62266 | 48.95 | 1 | 7 | 7 | 11 | 7.69 | 1 | 1 | 7.69 | 1 | 1 | | | | | | | | | | | | | | | | | | | 7.69 | 1 | 1 | 48.95 | 6 | 8 | 143 | 15.8 | 10.49 |
| P62269 | 57.24 | 1 | 11 | 11 | 28 | 11.18 | 2 | 2 | 17.11 | 3 | 3 | | | | 13.16 | 3 | 3 | | | | | | | | | | | | | | | | 57.24 | 11 | 20 | 152 | 17.7 | 10.99 |
| P62277 | 43.71 | 1 | 7 | 7 | 17 | | | | 7.95 | 1 | 1 | | | | | | | | | | | | | | | | | | | | | | 43.71 | 7 | 16 | 151 | 17.2 | 10.54 |
| P62280 | 61.39 | 1 | 12 | 12 | 31 | 6.33 | 1 | 1 | 10.76 | 2 | 2 | 6.33 | 1 | 2 | 4.43 | 1 | 1 | | | | | | | 5.06 | 1 | 1 | | | | | | | 61.39 | 12 | 24 | 158 | 18.4 | 10.30 |
| P62316 | 23.73 | 1 | 2 | 2 | 2 | | | | | | | | | | | | | | | | | | | | | | | | | | | | 23.73 | 2 | 2 | 118 | 13.5 | 9.91 |
| P62318 | 31.75 | 1 | 3 | 3 | 5 | | | | | | | | | | | | | | | | | | | 7.94 | 1 | 1 | | | | | | | 31.75 | 3 | 4 | 126 | 13.9 | 10.32 |
| P62330 | 78.29 | 2 | 7 | 10 | 30 | 6.29 | 1 | 1 | 6.29 | 1 | 1 | | | | 17.14 | 2 | 2 | 0.00 | | | | | | | | | | | | | | | 72.57 | 9 | 26 | 175 | 20.1 | 8.95 |
| P62333 | 64.52 | 1 | 26 | 26 | 81 | 11.31 | 3 | 4 | 31.11 | 8 | 11 | 17.99 | 4 | 4 | 7.97 | 3 | 3 | 4.88 | 2 | 2 | | | | 10.54 | 4 | 5 | 2.83 | 1 | 1 | 20.05 | 7 | 7 | 56.04 | 20 | 44 | 389 | 44.1 | 7.49 |
| P62424 | 47.37 | 1 | 15 | 16 | 35 | 4.14 | 1 | 1 | 14.29 | 3 | 3 | | | | | | | | | | | | | | | | | | | | | | 47.37 | 15 | 31 | 266 | 30.0 | 10.61 |
| P62495 | 21.74 | 1 | 8 | 8 | 22 | | | | 2.97 | 1 | 1 | 5.03 | 2 | 2 | 2.29 | 1 | 1 | | | | | | | | | | | | | 4.58 | 2 | 2 | 21.74 | 7 | 16 | 437 | 49.0 | 5.71 |
| P62633 | 19.77 | 1 | 3 | 3 | 3 | | | | | | | | | | | | | | | | | | | | | | | | | | | | 19.77 | 3 | 3 | 177 | 19.4 | 7.71 |
| P62699 | 44.63 | 1 | 4 | 4 | 6 | | | | | | | | | | | | | | | | | | | | | | | | | | | | 44.63 | 4 | 6 | 121 | 13.8 | 7.31 |
| P62701 | 56.27 | 1 | 11 | 18 | 58 | 3.42 | 1 | 1 | | | | 9.89 | 2 | 2 | 14.07 | 3 | 3 | 3.42 | 1 | 1 | | | | | | | | | | 17.49 | 4 | 4 | 56.27 | 18 | 47 | 263 | 29.6 | 10.15 |
| P16403 | 23.94 | 3 | 1 | 6 | 20 | 17.84 | 4 | 5 | 18.31 | 3 | 3 | 15.96 | 3 | 3 | 12.68 | 2 | 2 | | | | | | | | | | | | | | | | 15.49 | 4 | 7 | 213 | 21.4 | 10.93 |
| P62714 | 59.22 | 1 | 2 | 15 | 39 | 7.77 | 2 | 2 | 3.56 | 1 | 1 | 3.56 | 1 | 1 | 8.41 | 2 | 2 | | | | | | | 3.56 | 1 | 1 | | | | 8.41 | 2 | 2 | 55.66 | 14 | 30 | 309 | 35.6 | 5.43 |
| P62750 | 34.62 | 1 | 7 | 7 | 12 | | | | 14.74 | 2 | 2 | 8.33 | 1 | 1 | | | | | | | | | | | | | | | | | | | 28.21 | 6 | 9 | 156 | 17.7 | 10.45 |
| P62753 | 28.92 | 1 | 10 | 10 | 26 | 6.43 | 1 | 2 | 14.86 | 4 | 5 | | | | | | | 8.43 | 2 | 2 | | | | | | | | | | 11.24 | 2 | 3 | 24.10 | 6 | 14 | 249 | 28.7 | 10.84 |
| P62805 | 59.22 | 1 | 12 | 12 | 73 | 53.40 | 8 | 10 | 52.43 | 8 | 12 | 39.81 | 6 | 8 | 57.28 | 9 | 11 | | | | | | | 17.48 | 2 | 2 | | | | 11.65 | 1 | 1 | 54.37 | 9 | 29 | 103 | 11.4 | 11.36 |
| P62820 | 81.95 | 7 | 7 | 18 | 113 | 22.93 | 4 | 7 | 34.63 | 6 | 6 | 48.29 | 8 | 9 | 13.17 | 2 | 2 | 9.27 | 3 | 3 | | | | 8.78 | 2 | 2 | 17.07 | 5 | 5 | 31.71 | 5 | 6 | 80.49 | 15 | 73 | 205 | 22.7 | 6.21 |
| P62826 | 51.39 | 1 | 12 | 12 | 72 | 9.26 | 2 | 2 | 18.98 | 4 | 4 | 18.98 | 4 | 4 | 14.81 | 3 | 3 | 13.89 | 3 | 3 | | | | 4.63 | 1 | 1 | 10.19 | 2 | 2 | 15.74 | 3 | 3 | 50.93 | 11 | 50 | 216 | 24.4 | 7.49 |
| P62829 | 46.43 | 1 | 6 | 6 | 10 | | | | | | | | | | | | | 7.14 | 1 | 1 | | | | | | | 7.14 | 1 | 1 | | | | 46.43 | 6 | 8 | 140 | 14.9 | 10.51 |
| P62834 | 53.26 | 1 | 2 | 11 | 33 | 12.50 | 2 | 2 | 12.50 | 2 | 2 | 18.48 | 3 | 3 | | | | | | | | | | 5.98 | 1 | 1 | | | | 5.98 | 1 | 1 | 53.26 | 11 | 24 | 184 | 21.0 | 6.67 |
| P62841 | 13.10 | 1 | 3 | 3 | 5 | | | | | | | | | | | | | | | | | | | | | | | | | | | | 13.10 | 3 | 5 | 145 | 17.0 | 10.39 |
| P62847 | 35.34 | 1 | 5 | 5 | 8 | | | | | | | | | | | | | | | | | | | | | | | | | | | | 35.34 | 5 | 8 | 133 | 15.4 | 10.78 |
| P62851 | 42.40 | 1 | 7 | 7 | 17 | 14.40 | 2 | 2 | 21.60 | 3 | 3 | 7.20 | 1 | 1 | 7.20 | 1 | 1 | 11.20 | 1 | 1 | | | | | | | | | | | | | 29.60 | 5 | 9 | 125 | 13.7 | 10.11 |
| P62854 | 31.30 | 2 | 3 | 3 | 12 | | | | 20.87 | 2 | 2 | 7.83 | 1 | 1 | 7.83 | 1 | 1 | | | | | | | 7.83 | 1 | 1 | | | | | | | 31.30 | 3 | 7 | 115 | 13.0 | 11.00 |
| P62857 | 36.23 | 1 | 3 | 3 | 3 | 30.43 | 2 | 2 | 23.19 | 1 | 1 | | | | | | | | | | | | | | | | | | | | | | | | | 69 | 7.8 | 10.70 |
| P62873 | 61.18 | 2 | 10 | 16 | 63 | 23.24 | 5 | 5 | 10.29 | 3 | 3 | 45.88 | 11 | 16 | | | | 3.53 | 1 | 1 | | | | | | | 3.53 | 1 | 1 | 7.65 | 2 | 2 | 60.29 | 15 | 35 | 340 | 37.4 | 6.00 |
| P62877 | 25.00 | 1 | 3 | 3 | 5 | 17.59 | 1 | 1 | | | | | | | | | | | | | | | | | | | | | | 17.59 | 1 | 1 | 24.07 | 2 | 3 | 108 | 12.3 | 6.96 |
| P62879 | 70.29 | 3 | 11 | 17 | 79 | 25.00 | 5 | 6 | 10.29 | 3 | 3 | 47.65 | 11 | 15 | 7.65 | 2 | 2 | 3.53 | 1 | 1 | | | | | | | 3.53 | 1 | 1 | 3.53 | 1 | 1 | 70.29 | 17 | 50 | 340 | 37.3 | 6.00 |
| P62888 | 69.57 | 1 | 6 | 7 | 12 | 13.91 | 1 | 1 | | | | | | | | | | | | | | | | | | | | | | | | | 69.57 | 7 | 11 | 115 | 12.8 | 9.63 |
| P62899 | 37.60 | 1 | 5 | 5 | 8 | | | | | | | | | | | | | | | | | | | | | | | | | | | | 37.60 | 5 | 8 | 125 | 14.5 | 10.54 |
| P62906 | 27.19 | 1 | 5 | 5 | 10 | 9.68 | 2 | 2 | 9.68 | 2 | 2 | | | | | | | | | | | | | | | | | | | | | | 23.50 | 4 | 6 | 217 | 24.8 | 9.94 |
| P62910 | 41.48 | 1 | 6 | 6 | 12 | | | | | | | | | | | | | | | | | | | | | | | | | | | | 41.48 | 6 | 12 | 135 | 15.8 | 11.33 |
| P62913 | 32.58 | 1 | 7 | 7 | 26 | 12.92 | 2 | 2 | 12.92 | 2 | 2 | 5.06 | 1 | 1 | 9.55 | 2 | 2 | | | | | | | 7.87 | 1 | 1 | 4.49 | 1 | 1 | 12.36 | 2 | 2 | 32.02 | 6 | 15 | 178 | 20.2 | 9.60 |
| P62917 | 34.24 | 1 | 8 | 8 | 22 | 4.28 | 1 | 1 | 4.28 | 1 | 1 | 4.28 | 1 | 1 | 4.28 | 1 | 1 | | | | | | | 2.72 | 1 | 1 | | | | | | | 34.24 | 8 | 17 | 257 | 28.0 | 11.03 |
| P62937 | 82.42 | 7 | 14 | 14 | 133 | 41.21 | 6 | 7 | 26.06 | 4 | 6 | 43.64 | 6 | 7 | 42.42 | 8 | 11 | 43.64 | 9 | 14 | 16.97 | 2 | 2 | 33.94 | 7 | 10 | 43.64 | 9 | 17 | 51.52 | 11 | 19 | 74.55 | 13 | 40 | 165 | 18.0 | 7.81 |
| P62942 | 40.74 | 2 | 3 | 3 | 8 | | | | | | | | | | 15.74 | 1 | 1 | | | | | | | | | | 27.78 | 2 | 2 | 27.78 | 2 | 2 | 25.00 | 2 | 3 | 108 | 11.9 | 8.16 |
| P62979 | 65.38 | 5 | 11 | 11 | 150 | 35.26 | 6 | 8 | 35.26 | 6 | 8 | 35.26 | 6 | 8 | 15.38 | 3 | 4 | 28.21 | 4 | 5 | | | | 25.64 | 4 | 6 | 39.10 | 5 | 6 | 37.82 | 5 | 7 | 60.26 | 9 | 98 | 156 | 18.0 | 9.64 |
| Q9UNX3 | 44.83 | 1 | 1 | 11 | 20 | | | | 5.52 | 1 | 1 | | | | | | | | | | | | | | | | | | | | | | 44.83 | 11 | 19 | 145 | 17.2 | 10.55 |
| P62993 | 8.76 | 1 | 2 | 2 | 2 | | | | | | | | | | | | | | | | | | | | | | | | | | | | 8.76 | 2 | 2 | 217 | 25.2 | 6.32 |
| P63000 | 44.27 | 4 | 5 | 9 | 23 | 4.17 | 1 | 1 | | | | | | | 17.19 | 3 | 3 | 5.73 | 1 | 1 | | | | | | | | | | | | | 41.15 | 8 | 18 | 192 | 21.4 | 8.50 |
| P63010 | 31.06 | 1 | 7 | 25 | 78 | 1.07 | 1 | 1 | | | | 2.99 | 2 | 2 | 1.17 | 1 | 1 | | | | | | | | | | | | | 2.77 | 2 | 3 | 29.88 | 23 | 71 | 937 | 104.5 | 5.38 |
| P63092 | 38.07 | 3 | 12 | 13 | 41 | 6.85 | 2 | 3 | | | | 9.39 | 3 | 3 | 9.14 | 3 | 3 | 2.79 | 1 | 1 | | | | | | | 2.79 | 1 | 1 | 6.85 | 2 | 2 | 34.01 | 12 | 28 | 394 | 45.6 | 5.82 |
| P63104 | 79.59 | 1 | 21 | 27 | 215 | 44.49 | 9 | 17 | 52.24 | 11 | 15 | 49.39 | 10 | 13 | 54.29 | 13 | 18 | 60.41 | 14 | 18 | 4.49 | 2 | 3 | 33.88 | 10 | 15 | 46.53 | 10 | 12 | 45.31 | 12 | 20 | 73.06 | 20 | 84 | 245 | 27.7 | 4.79 |
| P63151 | 16.33 | 4 | 5 | 5 | 14 | 3.13 | 1 | 1 | | | | | | | | | | | | | | | | | | | | | | | | | 16.33 | 5 | 13 | 447 | 51.7 | 6.20 |
| P63167 | 65.17 | 1 | 4 | 7 | 108 | 59.55 | 6 | 8 | 59.55 | 4 | 6 | 58.43 | 5 | 6 | 59.55 | 6 | 11 | 59.55 | 5 | 10 | | | | 46.07 | 4 | 8 | 59.55 | 4 | 7 | 59.55 | 4 | 10 | 65.17 | 7 | 42 | 89 | 10.4 | 7.40 |
| P63172 | 79.65 | 1 | 5 | 5 | 30 | 30.09 | 2 | 3 | 30.09 | 2 | 4 | 30.09 | 2 | 3 | 14.16 | 1 | 2 | 14.16 | 1 | 3 | | | | | | | 14.16 | 1 | 1 | 14.16 | 1 | 2 | 79.65 | 5 | 12 | 113 | 12.4 | 5.08 |
| P63173 | 35.71 | 1 | 4 | 4 | 6 | 10.00 | 1 | 1 | | | | | | | 10.00 | 1 | 1 | | | | | | | | | | | | | | | | 35.71 | 3 | 4 | 70 | 8.2 | 10.10 |
| P63208 | 54.60 | 1 | 9 | 10 | 26 | 38.04 | 4 | 5 | 23.93 | 3 | 4 | | | | | | | | | | | | | | | | | | | 8.59 | 1 | 1 | 30.06 | 7 | 16 | 163 | 18.6 | 4.54 |
| P63220 | 22.89 | 1 | 2 | 2 | 2 | 12.05 | 1 | 1 | | | | | | | | | | | | | | | | | | | | | | | | | 10.84 | 1 | 1 | 83 | 9.1 | 8.50 |
| P63241 | 88.31 | 3 | 14 | 14 | 58 | | | | 7.79 | 1 | 2 | 7.79 | 1 | 1 | 36.36 | 3 | 4 | 19.48 | 2 | 2 | | | | 15.58 | 3 | 4 | 46.10 | 5 | 9 | 46.10 | 5 | 8 | 82.47 | 12 | 28 | 154 | 16.8 | 5.24 |
| P63244 | 79.81 | 1 | 20 | 20 | 90 | 25.87 | 7 | 8 | | | | 29.34 | 8 | 9 | 8.83 | 3 | 3 | 5.05 | 1 | 1 | | | | 2.21 | 1 | 1 | 5.05 | 1 | 1 | 31.86 | 8 | 9 | 78.55 | 19 | 58 | 317 | 35.1 | 7.69 |
| P67809 | 12.96 | 1 | 3 | 3 | 4 | 8.33 | 2 | 2 | | | | 9.57 | 2 | 2 | | | | | | | | | | | | | | | | | | | | | | 324 | 35.9 | 9.88 |
| P67812 | 40.78 | 2 | 8 | 8 | 34 | 6.70 | 1 | 1 | 18.44 | 4 | 7 | 6.70 | 1 | 1 | | | | | | | | | | | | | | | | | | | 33.52 | 7 | 25 | 179 | 20.6 | 9.48 |
| P67870 | 57.21 | 1 | 10 | 10 | 29 | 23.26 | 3 | 3 | 11.63 | 2 | 2 | 7.44 | 1 | 1 | 6.51 | 1 | 1 | 10.70 | 2 | 3 | | | | | | | | | | | | | 57.21 | 10 | 19 | 215 | 24.9 | 5.55 |
| P67936 | 33.06 | 3 | 2 | 8 | 18 | 12.90 | 3 | 3 | 9.27 | 2 | 2 | 3.63 | 1 | 1 | | | | 12.90 | 3 | 6 | | | | | | | | | | 10.08 | 2 | 2 | 18.55 | 4 | 4 | 248 | 28.5 | 4.69 |
| P68032 | 38.73 | 5 | 4 | 21 | 425 | 29.71 | 11 | 23 | 33.16 | 12 | 30 | 33.16 | 13 | 28 | 33.16 | 13 | 28 | 33.16 | 17 | 35 | 8.22 | 3 | 3 | 33.16 | 13 | 25 | 33.95 | 12 | 24 | 37.93 | 17 | 36 | 31.83 | 13 | 193 | 377 | 42.0 | 5.39 |
| P68036 | 53.25 | 1 | 7 | 7 | 11 | | | | | | | | | | 5.84 | 1 | 1 | | | | | | | | | | | | | 5.84 | 1 | 1 | 53.25 | 7 | 9 | 154 | 17.9 | 8.51 |
| P68104 | 78.79 | 2 | 17 | 33 | 502 | 28.79 | 11 | 18 | 31.17 | 12 | 20 | 40.26 | 15 | 24 | 35.93 | 15 | 24 | 35.06 | 16 | 23 | 6.71 | 3 | 3 | 29.87 | 14 | 21 | 42.42 | 18 | 27 | 43.51 | 21 | 42 | 73.81 | 26 | 300 | 462 | 50.1 | 9.01 |
| P68366 | 69.64 | 2 | 7 | 31 | 756 | 41.96 | 16 | 27 | 39.96 | 16 | 32 | 39.73 | 14 | 23 | 43.75 | 22 | 62 | 39.96 | 15 | 34 | 6.47 | 2 | 2 | 19.87 | 9 | 14 | 28.79 | 10 | 16 | 43.97 | 14 | 24 | 68.30 | 28 | 522 | 448 | 49.9 | 5.06 |
| P68371 | 87.19 | 3 | 2 | 43 | 1314 | 69.44 | 24 | 55 | 73.26 | 25 | 78 | 70.11 | 25 | 54 | 75.96 | 32 | 109 | 78.43 | 37 | 133 | 13.93 | 4 | 5 | 44.27 | 16 | 31 | 40.90 | 19 | 39 | 54.83 | 22 | 45 | 83.82 | 34 | 765 | 445 | 49.8 | 4.89 |
| P68400 | 54.99 | 2 | 16 | 18 | 64 | 16.88 | 5 | 5 | 25.32 | 8 | 12 | 13.30 | 4 | 4 | 3.84 | 1 | 1 | | | | | | | | | | | | | | | | 51.15 | 16 | 42 | 391 | 45.1 | 7.74 |
| Q9NVA2 | 15.38 | 2 | 1 | 7 | 19 | 2.10 | 1 | 1 | 2.10 | 1 | 1 | 2.10 | 1 | 1 | 2.10 | 1 | 1 | 6.99 | 2 | 3 | | | | | | | | | | | | | 14.92 | 6 | 12 | 429 | 49.4 | 6.81 |
| P68402 | 37.55 | 1 | 7 | 8 | 24 | 3.93 | 1 | 1 | 3.93 | 1 | 1 | 3.93 | 1 | 1 | 3.93 | 1 | 1 | | | | | | | 3.93 | 1 | 1 | | | | 6.99 | 2 | 2 | 37.55 | 8 | 17 | 229 | 25.6 | 5.92 |
| P68431 | 52.21 | 5 | 9 | 9 | 40 | 46.32 | 6 | 13 | 17.65 | 4 | 4 | 12.50 | 3 | 3 | 11.76 | 2 | 2 | | | | | | | 5.15 | 1 | 1 | | | | 5.15 | 1 | 1 | 51.47 | 7 | 16 | 136 | 15.4 | 11.12 |
| P68871 | 42.86 | 8 | 5 | 5 | 7 | | | | | | | | | | 21.77 | 3 | 3 | | | | | | | | | | | | | | | | 36.05 | 4 | 4 | 147 | 16.0 | 7.28 |
| P69905 | 14.79 | 2 | 2 | 2 | 3 | | | | | | | | | | 14.79 | 2 | 3 | | | | | | | | | | | | | | | | | | | 142 | 15.2 | 8.68 |
| P78318 | 18.58 | 1 | 7 | 7 | 12 | 4.13 | 1 | 1 | 6.78 | 2 | 2 | | | | | | | 4.13 | 1 | 1 | | | | | | | | | | | | | 18.58 | 7 | 8 | 339 | 39.2 | 5.38 |
| P78324 | 9.92 | 4 | 4 | 4 | 5 | 1.98 | 1 | 1 | | | | 6.75 | 3 | 3 | | | | | | | | | | | | | | | | | | | 3.17 | 1 | 1 | 504 | 54.9 | 6.98 |
| P78344 | 6.28 | 1 | 5 | 5 | 7 | | | | | | | | | | | | | | | | | | | | | | | | | | | | 6.28 | 5 | 7 | 907 | 102.3 | 7.14 |
| P78368 | 23.37 | 2 | 2 | 9 | 18 | 5.78 | 2 | 2 | 13.49 | 5 | 5 | | | | | | | | | | | | | | | | | | | | | | 17.59 | 7 | 11 | 415 | 47.4 | 9.06 |
| P78371 | 84.49 | 1 | 46 | 46 | 404 | 51.59 | 20 | 24 | 65.05 | 26 | 41 | 59.07 | 24 | 32 | 51.96 | 21 | 31 | 46.36 | 21 | 27 | | | | 31.78 | 14 | 19 | 33.64 | 17 | 25 | 46.17 | 22 | 30 | 80.19 | 40 | 175 | 535 | 57.5 | 6.46 |
| P78382 | 6.53 | 1 | 2 | 2 | 2 | | | | | | | | | | | | | | | | | | | | | | | | | | | | 6.53 | 2 | 2 | 337 | 36.8 | 8.95 |
| P78406 | 62.50 | 1 | 20 | 21 | 64 | 7.07 | 3 | 3 | 12.77 | 4 | 4 | 14.67 | 4 | 4 | 10.87 | 4 | 4 | 26.09 | 8 | 9 | | | | | | | 4.08 | 2 | 2 | 1.36 | 1 | 1 | 62.50 | 20 | 37 | 368 | 40.9 | 7.83 |
| P78417 | 36.10 | 1 | 7 | 7 | 17 | | | | | | | | | | 7.47 | 2 | 2 | | | | | | | | | | | | | 4.15 | 1 | 1 | 36.10 | 7 | 14 | 241 | 27.5 | 6.60 |
| P78527 | 2.23 | 1 | 7 | 7 | 10 | | | | | | | | | | | | | | | | | | | | | | | | | | | | 2.23 | 7 | 10 | 4128 | 468.8 | 7.12 |
| P78537 | 20.92 | 1 | 3 | 3 | 3 | | | | | | | | | | | | | | | | | | | | | | | | | | | | 20.92 | 3 | 3 | 153 | 17.3 | 9.33 |
| P78540 | 12.43 | 1 | 3 | 3 | 3 | | | | | | | | | | | | | | | | | | | | | | | | | | | | 12.43 | 3 | 3 | 354 | 38.6 | 6.47 |
| P80188 | 60.10 | 1 | 10 | 10 | 26 | 38.89 | 5 | 6 | 7.58 | 1 | 1 | 37.88 | 5 | 6 | 7.58 | 1 | 1 | | | | | | | | | | | | | | | | 54.55 | 9 | 12 | 198 | 22.6 | 8.91 |
| P80303 | 61.43 | 1 | 25 | 26 | 141 | 41.67 | 16 | 22 | 42.38 | 17 | 28 | 47.14 | 14 | 20 | 13.81 | 5 | 7 | 40.71 | 14 | 19 | | | | 12.86 | 5 | 5 | 21.19 | 8 | 9 | 40.00 | 11 | 14 | 26.90 | 10 | 17 | 420 | 50.2 | 5.12 |
| P80404 | 6.60 | 1 | 2 | 2 | 3 | | | | | | | | | | | | | | | | | | | | | | | | | | | | 6.60 | 2 | 3 | 500 | 56.4 | 7.96 |
| P80511 | 43.48 | 1 | 6 | 6 | 15 | 21.74 | 2 | 3 | | | | 30.43 | 3 | 4 | 31.52 | 4 | 4 | 30.43 | 2 | 2 | | | | | | | | | | 35.87 | 2 | 2 | | | | 92 | 10.6 | 6.25 |
| P80723 | 92.51 | 1 | 20 | 20 | 136 | 56.83 | 10 | 10 | 63.44 | 9 | 14 | 63.00 | 11 | 16 | 63.44 | 8 | 8 | 66.96 | 11 | 12 | | | | | | | 12.33 | 2 | 2 | 21.59 | 3 | 3 | 89.87 | 14 | 71 | 227 | 22.7 | 4.63 |
| P81605 | 34.55 | 1 | 5 | 5 | 29 | 10.00 | 1 | 1 | 10.00 | 1 | 1 | | | | 10.00 | 1 | 1 | 22.73 | 3 | 4 | 31.82 | 4 | 5 | 20.00 | 2 | 2 | 22.73 | 3 | 4 | 22.73 | 3 | 3 | 22.73 | 3 | 8 | 110 | 11.3 | 6.54 |
| P82094 | 1.92 | 1 | 2 | 2 | 3 | 0.00 | | | | | | | | | | | | | | | | | | | | | 0.91 | 1 | 1 | | | | 1.92 | 2 | 2 | 1093 | 122.8 | 4.92 |
| P82909 | 69.90 | 1 | 6 | 6 | 13 | 29.13 | 2 | 2 | 44.66 | 3 | 4 | 11.65 | 1 | 1 | | | | | | | | | | | | | | | | | | | 69.90 | 5 | 6 | 103 | 11.5 | 9.99 |
| P83436 | 22.99 | 1 | 12 | 12 | 14 | | | | | | | | | | | | | | | | | | | | | | | | | | | | 22.99 | 12 | 14 | 770 | 86.3 | 5.47 |
| P83731 | 31.85 | 1 | 6 | 6 | 15 | | | | 13.38 | 2 | 2 | | | | | | | | | | | | | | | | | | | | | | 31.85 | 6 | 13 | 157 | 17.8 | 11.25 |
| P84085 | 71.11 | 2 | 5 | 12 | 45 | 15.56 | 2 | 2 | 17.22 | 3 | 3 | 11.67 | 1 | 1 | 5.56 | 1 | 1 | | | | | | | 5.56 | 1 | 1 | | | | | | | 71.11 | 12 | 37 | 180 | 20.5 | 6.79 |
| P84090 | 16.35 | 1 | 2 | 2 | 2 | | | | | | | | | | | | | | | | | | | | | | | | | | | | 16.35 | 2 | 2 | 104 | 12.3 | 5.92 |
| P84095 | 46.60 | 3 | 6 | 7 | 11 | | | | | | | 8.90 | 1 | 1 | 6.81 | 1 | 1 | 5.76 | 1 | 1 | | | | | | | | | | | | | 46.60 | 6 | 8 | 191 | 21.3 | 8.12 |
| P84098 | 26.53 | 1 | 6 | 6 | 19 | 8.67 | 1 | 1 | 13.27 | 2 | 2 | 4.59 | 1 | 1 | 4.59 | 1 | 1 | 9.69 | 2 | 2 | | | | | | | | | | | | | 21.43 | 5 | 12 | 196 | 23.5 | 11.47 |
| P84103 | 14.02 | 2 | 2 | 2 | 2 | | | | | | | | | | | | | | | | | | | | | | | | | | | | 14.02 | 2 | 2 | 164 | 19.3 | 11.65 |
| P86790 | 4.15 | 1 | 2 | 2 | 3 | | | | | | | | | | | | | | | | | | | | | | | | | | | | 4.15 | 2 | 3 | 482 | 55.8 | 6.48 |
| P98088 | 5.89 | 1 | 12 | 13 | 15 | | | | | | | 5.89 | 13 | 15 | | | | | | | | | | | | | | | | | | | | | | 5654 | 585.2 | 7.02 |
| P98095 | 8.53 | 1 | 10 | 10 | 35 | 2.96 | 3 | 3 | | | | 2.53 | 2 | 2 | | | | | | | | | | | | | | | | | | | 7.52 | 8 | 30 | 1184 | 126.5 | 4.82 |
| P98160 | 8.34 | 1 | 28 | 28 | 38 | | | | | | | 0.25 | 1 | 1 | | | | | | | | | | | | | | | | | | | 8.34 | 28 | 37 | 4391 | 468.5 | 6.51 |
| P98164 | 0.69 | 1 | 3 | 3 | 4 | | | | | | | | | | | | | | | | | | | | | | | | | | | | 0.69 | 3 | 4 | 4655 | 521.6 | 5.08 |
| P98194 | 2.72 | 1 | 2 | 2 | 3 | | | | | | | | | | | | | | | | | | | | | | | | | | | | 2.72 | 2 | 3 | 919 | 100.5 | 6.74 |
| P99999 | 65.71 | 2 | 11 | 11 | 46 | 64.76 | 9 | 14 | 41.90 | 5 | 8 | 40.95 | 4 | 8 | 24.76 | 3 | 3 | 25.71 | 3 | 3 | | | | | | | | | | | | | 56.19 | 7 | 10 | 105 | 11.7 | 9.57 |
| Q00169 | 23.70 | 1 | 3 | 5 | 9 | 8.52 | 1 | 1 | | | | | | | | | | | | | | | | | | | | | | | | | 23.70 | 5 | 8 | 270 | 31.8 | 6.55 |
| Q00266 | 26.84 | 1 | 6 | 9 | 17 | | | | | | | | | | 3.80 | 1 | 1 | 2.03 | 1 | 1 | | | | | | | | | | | | | 26.84 | 9 | 15 | 395 | 43.6 | 6.29 |
| Q00325 | 38.67 | 1 | 16 | 16 | 100 | 20.72 | 7 | 7 | 20.72 | 7 | 7 | 20.17 | 7 | 7 | 5.52 | 2 | 2 | | | | | | | | | | | | | | | | 38.67 | 16 | 77 | 362 | 40.1 | 9.38 |
| Q00341 | 22.63 | 2 | 25 | 25 | 46 | | | | | | | | | | 0.95 | 1 | 1 | 0.71 | 1 | 1 | | | | | | | | | | 11.12 | 11 | 11 | 17.35 | 20 | 33 | 1268 | 141.4 | 6.87 |
| Q00535 | 15.75 | 17 | 3 | 4 | 9 | 0.00 | | | 0.00 | | | 2.74 | 1 | 1 | | | | | | | | | | | | | | | | | | | 15.75 | 4 | 8 | 292 | 33.3 | 7.66 |
| Q00577 | 34.47 | 3 | 8 | 10 | 25 | | | | | | | | | | | | | | | | | | | | | | | | | 2.80 | 1 | 1 | 34.47 | 10 | 24 | 322 | 34.9 | 6.44 |
| Q00610 | 62.09 | 1 | 70 | 93 | 666 | 30.63 | 40 | 47 | 26.51 | 34 | 40 | 37.43 | 50 | 63 | 16.18 | 24 | 26 | 6.03 | 8 | 9 | | | | 3.64 | 7 | 7 | 2.75 | 4 | 4 | 10.51 | 15 | 15 | 54.57 | 81 | 455 | 1675 | 191.5 | 5.69 |
| Q00688 | 15.18 | 1 | 3 | 3 | 6 | | | | | | | | | | | | | 6.70 | 1 | 1 | | | | | | | | | | | | | 15.18 | 3 | 5 | 224 | 25.2 | 9.28 |
| Q00765 | 36.51 | 1 | 9 | 9 | 17 | 14.81 | 3 | 3 | 10.05 | 2 | 2 | | | | | | | | | | | | | | | | | | | | | | 30.69 | 7 | 12 | 189 | 21.5 | 8.10 |
| Q00796 | 78.99 | 2 | 25 | 25 | 258 | 36.97 | 10 | 13 | 22.97 | 7 | 11 | 47.06 | 12 | 22 | 53.22 | 17 | 26 | 44.26 | 11 | 15 | | | | 19.61 | 6 | 8 | 8.40 | 3 | 4 | 37.54 | 9 | 14 | 71.15 | 21 | 145 | 357 | 38.3 | 7.97 |
| Q00839 | 13.94 | 1 | 9 | 9 | 18 | 1.45 | 1 | 1 | 1.45 | 1 | 1 | 7.39 | 4 | 4 | | | | | | | 0.73 | 1 | 1 | | | | | | | | | | 9.94 | 6 | 11 | 825 | 90.5 | 6.00 |
| P16402 | 19.91 | 3 | 1 | 6 | 18 | 9.95 | 3 | 4 | 10.41 | 2 | 2 | 15.38 | 3 | 3 | 4.98 | 1 | 1 | | | | | | | | | | | | | | | | 19.00 | 5 | 8 | 221 | 22.3 | 11.02 |
| Q01082 | 26.78 | 3 | 48 | 53 | 73 | | | | | | | | | | | | | | | | | | | | | | | | | 0.47 | 1 | 1 | 26.78 | 53 | 72 | 2364 | 274.4 | 5.57 |
| Q01415 | 10.92 | 1 | 3 | 3 | 4 | | | | | | | | | | | | | | | | | | | | | | | | | | | | 10.92 | 3 | 4 | 458 | 50.3 | 6.61 |
| P41219 | 9.79 | 4 | 1 | 6 | 18 | 3.83 | 2 | 2 | 2.13 | 1 | 1 | 3.83 | 2 | 2 | 3.83 | 2 | 2 | 6.38 | 3 | 3 | | | | | | | | | | 3.83 | 2 | 2 | 7.45 | 5 | 6 | 470 | 53.6 | 5.47 |
| Q01469 | 85.19 | 2 | 16 | 16 | 89 | 49.63 | 5 | 8 | 37.78 | 4 | 5 | 45.19 | 5 | 9 | 51.85 | 5 | 5 | 60.74 | 6 | 9 | | | | 14.81 | 2 | 2 | 80.74 | 10 | 13 | 67.41 | 7 | 10 | 76.30 | 14 | 28 | 135 | 15.2 | 7.01 |
| Q01518 | 54.53 | 2 | 18 | 24 | 103 | 27.16 | 9 | 11 | 6.74 | 3 | 3 | 22.53 | 10 | 10 | 12.42 | 5 | 5 | 6.53 | 3 | 3 | | | | 2.11 | 1 | 1 | 6.53 | 3 | 3 | 18.53 | 8 | 8 | 42.95 | 20 | 59 | 475 | 51.9 | 8.06 |
| Q01581 | 8.27 | 1 | 3 | 3 | 3 | | | | | | | | | | | | | | | | | | | | | | | | | | | | 8.27 | 3 | 3 | 520 | 57.3 | 5.41 |
| Q01813 | 62.88 | 1 | 43 | 48 | 440 | 39.29 | 22 | 28 | 36.22 | 21 | 31 | 37.76 | 22 | 26 | 32.53 | 19 | 23 | 14.92 | 9 | 13 | | | | 2.42 | 2 | 2 | 3.44 | 3 | 3 | 7.14 | 4 | 4 | 59.18 | 42 | 310 | 784 | 85.5 | 7.55 |
| Q02127 | 15.70 | 1 | 6 | 6 | 6 | | | | | | | | | | | | | | | | | | | | | | | | | | | | 15.70 | 6 | 6 | 395 | 42.8 | 9.67 |
| O15131 | 23.13 | 1 | 1 | 8 | 18 | 1.87 | 1 | 1 | | | | | | | | | | | | | | | | | | | | | | | | | 22.95 | 7 | 17 | 536 | 60.3 | 5.16 |
| Q02218 | 56.01 | 2 | 52 | 52 | 170 | 16.81 | 12 | 15 | 20.23 | 16 | 19 | 20.72 | 17 | 20 | | | | 2.25 | 2 | 2 | | | | | | | | | | | | | 49.36 | 44 | 114 | 1023 | 115.9 | 6.86 |
| Q02224 | 0.89 | 3 | 2 | 3 | 4 | | | | | | | 0.00 | | | 0.00 | | | 0.00 | | | | | | | | | | | | 0.30 | 1 | 1 | 0.89 | 3 | 3 | 2701 | 316.2 | 5.64 |
| Q02241 | 2.71 | 1 | 2 | 2 | 3 | | | | | | | | | | | | | | | | | | | | | | | | | | | | 2.71 | 2 | 3 | 960 | 110.0 | 8.51 |
| Q02252 | 18.13 | 1 | 8 | 8 | 9 | | | | | | | | | | | | | | | | | | | | | | | | | 2.24 | 1 | 1 | 15.89 | 7 | 8 | 535 | 57.8 | 8.50 |
| Q02338 | 15.16 | 1 | 4 | 4 | 4 | | | | | | | | | | | | | | | | | | | | | | | | | | | | 15.16 | 4 | 4 | 343 | 38.1 | 8.95 |
| Q02383 | 90.89 | 1 | 69 | 77 | 1977 | 62.54 | 34 | 184 | 63.06 | 33 | 157 | 64.26 | 35 | 140 | 62.71 | 35 | 83 | 75.95 | 47 | 111 | 47.25 | 26 | 40 | 63.06 | 34 | 63 | 68.21 | 41 | 96 | 76.80 | 46 | 87 | 79.04 | 57 | 1016 | 582 | 65.4 | 9.07 |
| Q02413 | 10.77 | 1 | 8 | 8 | 17 | | | | | | | | | | 1.33 | 1 | 1 | 1.33 | 1 | 1 | | | | | | | 8.87 | 6 | 10 | | | | 3.72 | 3 | 5 | 1049 | 113.7 | 5.03 |
| Q02543 | 43.75 | 1 | 10 | 10 | 27 | | | | | | | | | | | | | | | | | | | | | | | | | | | | 43.75 | 10 | 27 | 176 | 20.7 | 10.71 |
| Q02750 | 34.35 | 1 | 7 | 13 | 30 | | | | | | | 3.05 | 1 | 1 | | | | | | | | | | | | | | | | 10.69 | 4 | 4 | 28.75 | 11 | 25 | 393 | 43.4 | 6.62 |
| Q02790 | 76.91 | 1 | 37 | 37 | 174 | 19.83 | 7 | 8 | 12.42 | 6 | 6 | 17.21 | 7 | 7 | 19.83 | 7 | 7 | 14.60 | 6 | 6 | | | | 27.02 | 11 | 11 | 39.00 | 13 | 19 | 51.42 | 20 | 30 | 72.98 | 27 | 80 | 459 | 51.8 | 5.43 |
| Q02809 | 20.36 | 1 | 10 | 10 | 23 | 8.39 | 5 | 5 | 5.64 | 3 | 3 | 3.58 | 2 | 2 | | | | | | | | | | | | | | | | 2.61 | 1 | 1 | 16.23 | 8 | 12 | 727 | 83.5 | 6.95 |
| Q02817 | 1.74 | 1 | 9 | 9 | 10 | | | | | | | 1.74 | 9 | 10 | | | | | | | | | | | | | | | | | | | | | | 5179 | 540.0 | 5.80 |
| Q02818 | 9.76 | 2 | 3 | 4 | 7 | | | | | | | | | | | | | 5.42 | 2 | 2 | | | | 2.39 | 1 | 1 | 1.52 | 1 | 1 | 5.42 | 2 | 2 | 2.82 | 1 | 1 | 461 | 53.8 | 5.25 |
| Q02878 | 42.71 | 1 | 16 | 16 | 66 | 6.94 | 2 | 2 | 11.46 | 3 | 3 | 6.25 | 2 | 2 | 6.94 | 2 | 2 | | | | | | | | | | | | | 6.25 | 2 | 2 | 42.71 | 16 | 55 | 288 | 32.7 | 10.58 |
| Q02952 | 4.99 | 1 | 5 | 5 | 5 | 0.73 | 1 | 1 | | | | | | | | | | | | | | | | | | | | | | | | | 4.26 | 4 | 4 | 1782 | 191.4 | 4.41 |
| Q02978 | 63.38 | 1 | 14 | 14 | 37 | 7.64 | 2 | 2 | 21.66 | 6 | 6 | 5.10 | 1 | 1 | | | | | | | | | | | | | | | | | | | 63.38 | 14 | 28 | 314 | 34.0 | 9.91 |
| Q03154 | 6.86 | 1 | 2 | 2 | 3 | | | | | | | | | | | | | | | | | | | | | | | | | | | | 6.86 | 2 | 3 | 408 | 45.9 | 6.18 |
| Q03167 | 13.28 | 2 | 9 | 9 | 14 | 0.00 | | | | | | 1.41 | 1 | 1 | | | | 0.00 | | | | | | | | | | | | | | | 13.28 | 8 | 13 | 851 | 93.4 | 5.71 |
| Q03252 | 6.77 | 1 | 2 | 4 | 6 | 2.58 | 2 | 2 | 2.26 | 1 | 1 | 4.52 | 3 | 3 | | | | | | | | | | | | | | | | | | | 0.00 | | | 620 | 69.9 | 5.59 |
| Q03426 | 27.53 | 1 | 6 | 6 | 15 | | | | | | | | | | | | | | | | | | | | | | | | | | | | 27.53 | 6 | 15 | 396 | 42.4 | 6.46 |
| Q04206 | 19.06 | 1 | 10 | 10 | 22 | | | | | | | 2.54 | 1 | 1 | | | | | | | | | | 4.36 | 2 | 2 | | | | 2.00 | 1 | 1 | 18.69 | 9 | 18 | 551 | 60.2 | 5.68 |
| Q04446 | 17.09 | 1 | 8 | 8 | 14 | | | | | | | | | | | | | | | | | | | | | | | | | | | | 17.09 | 8 | 14 | 702 | 80.4 | 6.32 |
| Q04609 | 48.13 | 2 | 30 | 32 | 109 | 7.07 | 5 | 5 | 4.67 | 3 | 3 | 33.60 | 20 | 26 | | | | | | | | | | | | | | | | | | | 47.20 | 31 | 75 | 750 | 84.3 | 6.98 |
| Q04637 | 16.76 | 1 | 18 | 20 | 51 | 1.31 | 2 | 2 | 0.69 | 1 | 1 | 0.69 | 1 | 1 | 1.44 | 2 | 2 | | | | | | | | | | | | | | | | 15.38 | 18 | 45 | 1599 | 175.4 | 5.33 |
| Q04695 | 41.44 | 15 | 6 | 25 | 104 | | | | 7.87 | 4 | 4 | 7.87 | 4 | 4 | 5.32 | 3 | 3 | 9.26 | 5 | 5 | | | | 15.05 | 8 | 9 | 19.91 | 11 | 13 | 6.94 | 3 | 3 | 34.49 | 18 | 63 | 432 | 48.1 | 5.02 |
| Q04724 | 7.53 | 4 | 4 | 4 | 7 | 1.43 | 1 | 1 | | | | 2.86 | 2 | 2 | | | | | | | | | | | | | | | | | | | 6.10 | 3 | 4 | 770 | 83.1 | 7.24 |
| Q04760 | 70.65 | 1 | 13 | 13 | 56 | 28.26 | 5 | 5 | 22.83 | 4 | 5 | 27.72 | 5 | 5 | 22.28 | 5 | 5 | | | | | | | 9.78 | 2 | 2 | | | | 17.93 | 5 | 5 | 70.65 | 12 | 29 | 184 | 20.8 | 5.31 |
| Q04837 | 69.59 | 1 | 11 | 11 | 69 | 55.41 | 7 | 9 | 60.14 | 9 | 18 | 59.46 | 8 | 13 | 16.22 | 2 | 2 | 26.35 | 4 | 4 | | | | | | | 10.14 | 1 | 1 | 10.14 | 1 | 1 | 62.84 | 10 | 21 | 148 | 17.2 | 9.60 |
| Q04917 | 53.25 | 1 | 12 | 17 | 83 | 10.98 | 3 | 4 | 12.20 | 3 | 5 | 10.57 | 3 | 4 | 14.23 | 4 | 4 | 9.76 | 3 | 4 | | | | 9.35 | 3 | 4 | 13.01 | 3 | 4 | 34.55 | 8 | 10 | 50.41 | 13 | 43 | 246 | 28.2 | 4.84 |
| Q04941 | 18.42 | 1 | 2 | 2 | 5 | | | | | | | 8.55 | 1 | 1 | | | | | | | | | | | | | | | | | | | 18.42 | 2 | 4 | 152 | 16.7 | 7.24 |
| Q05086 | 3.09 | 1 | 2 | 2 | 5 | | | | | | | | | | | | | | | | | | | | | | | | | 1.49 | 1 | 1 | 3.09 | 2 | 4 | 875 | 100.6 | 5.22 |
| Q05469 | 13.75 | 1 | 9 | 9 | 32 | | | | | | | | | | | | | | | | | | | | | | | | | | | | 13.75 | 9 | 32 | 1076 | 116.5 | 6.70 |
| Q05639 | 50.76 | 1 | 5 | 21 | 240 | 12.53 | 6 | 8 | 14.04 | 7 | 9 | 20.30 | 8 | 10 | 15.77 | 7 | 9 | 16.20 | 8 | 11 | 4.54 | 2 | 2 | 15.77 | 7 | 9 | 22.25 | 10 | 14 | 23.11 | 12 | 20 | 48.16 | 17 | 148 | 463 | 50.4 | 9.03 |
| Q05655 | 29.73 | 5 | 15 | 16 | 35 | | | | | | | | | | | | | | | | | | | | | | | | | 6.80 | 4 | 4 | 28.25 | 14 | 31 | 676 | 77.5 | 7.75 |
| Q06210 | 69.96 | 3 | 47 | 47 | 260 | 7.87 | 4 | 5 | 9.30 | 5 | 6 | 22.03 | 12 | 13 | 18.88 | 11 | 11 | 2.00 | 1 | 1 | | | | 9.59 | 5 | 5 | 3.43 | 2 | 2 | 23.03 | 11 | 12 | 69.67 | 41 | 209 | 699 | 78.8 | 7.11 |
| Q06278 | 7.85 | 1 | 9 | 9 | 13 | | | | | | | 0.82 | 1 | 1 | | | | | | | | | | | | | | | | | | | 7.85 | 9 | 12 | 1338 | 147.8 | 7.17 |
| Q06323 | 48.59 | 1 | 13 | 14 | 34 | 18.47 | 4 | 4 | 4.82 | 1 | 1 | 4.82 | 1 | 1 | 16.47 | 3 | 3 | | | | | | | | | | | | | 8.03 | 1 | 1 | 44.18 | 11 | 24 | 249 | 28.7 | 6.02 |
| Q06481 | 3.93 | 1 | 2 | 3 | 3 | 0.79 | 1 | 1 | | | | | | | | | | | | | | | | | | | | | | | | | 3.15 | 2 | 2 | 763 | 86.9 | 4.79 |
| Q06730 | 1.60 | 1 | 2 | 2 | 2 | | | | | | | | | | | | | | | | | | | | | | | | | | | | 1.60 | 2 | 2 | 810 | 94.3 | 8.07 |
| Q06828 | 7.45 | 1 | 2 | 2 | 4 | | | | | | | | | | | | | | | | | | | | | | | | | | | | 7.45 | 2 | 4 | 376 | 43.2 | 6.04 |
| Q06830 | 75.88 | 1 | 13 | 17 | 155 | 36.18 | 8 | 9 | 41.71 | 8 | 9 | 49.75 | 10 | 10 | 58.29 | 11 | 11 | 41.21 | 8 | 9 | | | | 33.67 | 8 | 11 | 57.79 | 11 | 15 | 54.27 | 11 | 13 | 72.36 | 16 | 68 | 199 | 22.1 | 8.13 |
| Q07020 | 43.09 | 1 | 9 | 9 | 32 | 5.85 | 1 | 1 | 6.91 | 1 | 1 | 6.91 | 1 | 1 | 27.13 | 4 | 4 | | | | | | | | | | | | | | | | 43.09 | 9 | 25 | 188 | 21.6 | 11.72 |
| Q07021 | 31.91 | 1 | 7 | 7 | 23 | 21.63 | 4 | 6 | 10.99 | 2 | 2 | 21.28 | 3 | 3 | | | | 5.32 | 1 | 2 | | | | | | | 5.32 | 1 | 1 | | | | 31.56 | 5 | 9 | 282 | 31.3 | 4.84 |
| Q07065 | 39.53 | 5 | 18 | 20 | 42 | 9.14 | 4 | 4 | 6.48 | 3 | 3 | 6.98 | 3 | 3 | | | | 2.49 | 1 | 1 | | | | | | | | | | 7.48 | 3 | 3 | 37.71 | 19 | 28 | 602 | 66.0 | 5.92 |
| P62745 | 19.39 | 1 | 1 | 3 | 17 | 4.59 | 1 | 1 | | | | 13.27 | 2 | 2 | 4.59 | 1 | 1 | 4.59 | 1 | 1 | | | | 8.67 | 1 | 1 | 4.59 | 1 | 1 | 4.59 | 1 | 1 | 19.39 | 3 | 9 | 196 | 22.1 | 5.24 |
| Q07522 | 31.06 | 1 | 5 | 5 | 27 | 29.55 | 3 | 3 | 16.67 | 2 | 2 | 16.67 | 2 | 2 | | | | 17.42 | 3 | 3 | | | | | | | | | | | | | 31.06 | 5 | 16 | 132 | 15.7 | 7.17 |
| Q07812 | 42.19 | 1 | 7 | 7 | 9 | | | | | | | | | | | | | | | | | | | | | | | | | | | | 42.19 | 7 | 9 | 192 | 21.2 | 5.22 |
| Q96BM9 | 58.60 | 1 | 1 | 10 | 17 | 8.06 | 1 | 1 | 12.90 | 2 | 3 | 8.06 | 1 | 1 | | | | | | | | | | | | | | | | | | | 50.54 | 9 | 12 | 186 | 21.4 | 7.77 |
| Q07960 | 16.17 | 1 | 6 | 6 | 18 | | | | | | | | | | 2.51 | 1 | 1 | | | | | | | 2.51 | 1 | 1 | | | | 2.51 | 1 | 1 | 16.17 | 6 | 15 | 439 | 50.4 | 6.29 |
| Q08209 | 24.76 | 2 | 8 | 11 | 34 | 1.15 | 1 | 1 | | | | | | | | | | | | | | | | 1.15 | 1 | 1 | | | | 1.15 | 1 | 1 | 24.76 | 11 | 31 | 521 | 58.7 | 5.86 |
| Q08211 | 8.35 | 1 | 8 | 8 | 19 | | | | | | | | | | | | | | | | | | | | | | | | | | | | 8.35 | 8 | 19 | 1270 | 140.9 | 6.84 |
| Q08257 | 61.09 | 1 | 13 | 13 | 35 | 3.34 | 1 | 1 | | | | 10.03 | 3 | 3 | | | | | | | | | | 5.17 | 1 | 1 | 5.17 | 1 | 1 | 8.81 | 2 | 3 | 61.09 | 13 | 26 | 329 | 35.2 | 8.44 |
| Q08378 | 8.08 | 1 | 9 | 10 | 17 | | | | | | | | | | | | | | | | | | | | | | | | | | | | 8.08 | 10 | 17 | 1498 | 167.3 | 5.44 |
| Q08379 | 2.00 | 1 | 2 | 2 | 2 | | | | | | | | | | | | | | | | | | | | | | | | | | | | 2.00 | 2 | 2 | 1002 | 113.0 | 5.02 |
| Q08380 | 34.53 | 1 | 16 | 16 | 71 | 22.56 | 9 | 11 | 13.16 | 6 | 9 | 31.62 | 14 | 19 | | | | | | | | | | | | | | | | | | | 26.15 | 12 | 32 | 585 | 65.3 | 5.27 |
| Q08431 | 64.60 | 1 | 19 | 19 | 39 | 6.72 | 2 | 2 | 13.95 | 4 | 5 | 23.00 | 7 | 8 | | | | | | | | | | | | | | | | | | | 58.91 | 17 | 24 | 387 | 43.1 | 8.15 |
| Q08554 | 9.73 | 1 | 6 | 6 | 11 | | | | | | | | | | | | | | | | 1.45 | 1 | 1 | 1.68 | 1 | 1 | 6.71 | 4 | 6 | | | | 3.24 | 2 | 3 | 894 | 99.9 | 5.43 |
| Q08629 | 2.28 | 1 | 2 | 2 | 2 | | | | | | | 2.28 | 1 | 1 | | | | | | | | | | | | | | | | | | | 2.05 | 1 | 1 | 439 | 49.1 | 6.10 |
| Q08722 | 5.88 | 1 | 2 | 2 | 19 | 2.48 | 1 | 1 | 5.88 | 2 | 2 | 5.88 | 2 | 2 | | | | 2.48 | 1 | 1 | | | | | | | | | | | | | 5.88 | 2 | 13 | 323 | 35.2 | 7.21 |
| Q08752 | 12.97 | 1 | 4 | 4 | 7 | | | | | | | | | | 4.32 | 1 | 1 | | | | | | | | | | 2.43 | 1 | 1 | 12.97 | 4 | 4 | 2.43 | 1 | 1 | 370 | 40.7 | 7.21 |
| Q08830 | 9.94 | 1 | 3 | 3 | 5 | | | | 7.05 | 2 | 2 | 2.56 | 1 | 1 | | | | | | | | | | | | | | | | | | | 5.45 | 2 | 2 | 312 | 36.4 | 5.87 |
| Q08AM6 | 8.57 | 1 | 6 | 6 | 7 | | | | | | | | | | | | | | | | | | | | | | | | | | | | 8.57 | 6 | 7 | 782 | 87.9 | 6.13 |
| Q09666 | 10.14 | 1 | 14 | 14 | 16 | 3.45 | 2 | 3 | | | | | | | | | | | | | | | | 0.44 | 2 | 2 | 0.19 | 1 | 1 | 0.75 | 3 | 3 | 5.76 | 7 | 7 | 5890 | 628.7 | 6.15 |
| Q0IIM8 | 5.27 | 1 | 3 | 4 | 10 | 0.63 | 1 | 1 | 0.63 | 1 | 1 | | | | 0.63 | 1 | 1 | 0.63 | 1 | 1 | | | | 0.63 | 1 | 1 | | | | | | | 5.27 | 4 | 5 | 1120 | 128.6 | 5.95 |
| Q0PNE2 | 10.15 | 1 | 2 | 2 | 6 | | | | | | | | | | | | | | | | | | | | | | | | | | | | 10.15 | 2 | 6 | 266 | 29.8 | 6.84 |
| Q0VAA2 | 19.88 | 1 | 8 | 8 | 14 | | | | | | | | | | 1.84 | 1 | 1 | | | | | | | | | | | | | | | | 19.88 | 8 | 13 | 488 | 54.5 | 5.34 |
| Q0VDD8 | 2.57 | 1 | 6 | 7 | 20 | 0.26 | 1 | 1 | 0.26 | 1 | 1 | 0.26 | 1 | 1 | 0.26 | 1 | 1 | | | | | | | | | | | | | | | | 2.57 | 7 | 16 | 3507 | 399.6 | 6.93 |
| Q0VDF9 | 6.68 | 1 | 3 | 3 | 4 | | | | | | | | | | | | | | | | | | | | | | | | | | | | 6.68 | 3 | 4 | 509 | 54.8 | 5.59 |
| Q0VF26 | 60.87 | 1 | 44 | 44 | 149 | 2.36 | 1 | 1 | 4.17 | 2 | 2 | 2.36 | 1 | 1 | 20.83 | 10 | 12 | 44.57 | 22 | 28 | | | | 19.20 | 10 | 10 | 2.36 | 1 | 1 | 22.10 | 11 | 13 | 51.09 | 33 | 81 | 552 | 66.4 | 8.87 |
| Q10567 | 40.57 | 1 | 14 | 32 | 120 | 3.06 | 2 | 2 | 2.85 | 2 | 2 | 2.95 | 2 | 2 | 3.37 | 3 | 3 | | | | | | | | | | | | | 2.63 | 2 | 3 | 37.41 | 29 | 108 | 949 | 104.6 | 5.06 |
| Q10713 | 8.57 | 1 | 5 | 5 | 6 | 1.90 | 1 | 1 | 1.90 | 1 | 1 | | | | | | | | | | | | | | | | | | | | | | 6.67 | 4 | 4 | 525 | 58.2 | 6.92 |
| Q12756 | 3.31 | 3 | 4 | 4 | 12 | 0.00 | | | | | | | | | 0.71 | 1 | 1 | | | | | | | 0.71 | 1 | 1 | | | | | | | 3.31 | 4 | 10 | 1690 | 190.9 | 6.21 |
| Q12765 | 25.60 | 1 | 10 | 10 | 22 | | | | | | | | | | 3.14 | 1 | 1 | | | | | | | 2.90 | 1 | 1 | 7.49 | 3 | 4 | 6.52 | 3 | 3 | 16.18 | 5 | 13 | 414 | 46.4 | 4.75 |
| Q12768 | 19.41 | 1 | 17 | 18 | 24 | | | | | | | | | | | | | | | | | | | | | | | | | | | | 19.41 | 18 | 24 | 1159 | 134.2 | 6.98 |
| Q12769 | 18.66 | 1 | 21 | 21 | 36 | | | | | | | | | | | | | | | | | | | | | | | | | | | | 18.66 | 21 | 36 | 1436 | 162.0 | 5.50 |
| Q12792 | 29.71 | 1 | 7 | 10 | 21 | | | | | | | | | | 3.71 | 1 | 1 | | | | | | | 9.43 | 3 | 3 | | | | 11.14 | 3 | 3 | 27.71 | 9 | 14 | 350 | 40.3 | 6.96 |
| Q12798 | 52.33 | 2 | 10 | 10 | 21 | 5.23 | 1 | 1 | | | | | | | 25.58 | 4 | 5 | 42.44 | 8 | 10 | | | | | | | | | | | | | 38.95 | 5 | 5 | 172 | 19.6 | 4.88 |
| Q12800 | 5.98 | 2 | 2 | 2 | 5 | | | | | | | | | | | | | | | | | | | | | | | | | 3.59 | 1 | 1 | 5.98 | 2 | 4 | 502 | 57.2 | 5.80 |
| Q12805 | 19.27 | 1 | 7 | 7 | 15 | | | | | | | | | | | | | | | | | | | | | | | | | | | | 19.27 | 7 | 15 | 493 | 54.6 | 5.07 |
| Q12841 | 9.09 | 1 | 2 | 2 | 6 | 2.60 | 1 | 1 | 2.60 | 1 | 1 | 2.60 | 1 | 1 | 2.60 | 1 | 1 | | | | | | | | | | | | | | | | 9.09 | 2 | 2 | 308 | 35.0 | 5.52 |
| Q12894 | 8.50 | 1 | 2 | 2 | 3 | | | | | | | | | | | | | | | | | | | | | | | | | | | | 8.50 | 2 | 3 | 506 | 54.8 | 7.94 |
| Q12904 | 53.21 | 1 | 12 | 12 | 23 | | | | 7.69 | 2 | 2 | | | | | | | 4.81 | 2 | 2 | | | | | | | | | | 9.62 | 3 | 3 | 47.12 | 9 | 16 | 312 | 34.3 | 8.43 |
| Q12907 | 62.36 | 1 | 20 | 20 | 97 | 33.99 | 10 | 13 | 43.54 | 11 | 17 | 45.51 | 14 | 21 | 14.61 | 4 | 4 | 4.49 | 1 | 1 | | | | | | | | | | | | | 55.90 | 15 | 41 | 356 | 40.2 | 6.95 |
| Q12913 | 12.79 | 1 | 11 | 11 | 21 | 0.90 | 1 | 1 | | | | 11.97 | 10 | 12 | | | | | | | | | | | | | | | | | | | 5.09 | 4 | 8 | 1337 | 145.9 | 5.58 |
| Q12931 | 24.43 | 1 | 12 | 13 | 28 | 5.97 | 3 | 3 | 8.38 | 4 | 4 | 4.12 | 2 | 2 | 1.99 | 1 | 1 | 1.99 | 1 | 1 | | | | 1.99 | 1 | 1 | 1.99 | 1 | 1 | 1.99 | 1 | 1 | 22.30 | 12 | 14 | 704 | 80.1 | 8.21 |
| Q12965 | 9.21 | 3 | 8 | 9 | 22 | | | | | | | | | | | | | | | | | | | | | | | | | | | | 9.21 | 9 | 22 | 1108 | 127.0 | 8.92 |
| Q13011 | 52.74 | 1 | 16 | 16 | 48 | 25.00 | 7 | 9 | 24.09 | 8 | 10 | 21.34 | 7 | 8 | 2.13 | 1 | 1 | | | | | | | | | | | | | | | | 41.16 | 11 | 20 | 328 | 35.8 | 8.00 |
| Q13029 | 1.16 | 1 | 2 | 2 | 2 | | | | 1.16 | 2 | 2 | | | | | | | | | | | | | | | | | | | 0.00 | | | 0.00 | | | 1718 | 188.8 | 7.31 |

| | | | | | | | | | | | | | | | | | | | | | | | | | | | | | | | | | | | | | | |
|---|---|---|---|---|---|---|---|---|---|---|---|---|---|---|---|---|---|---|---|---|---|---|---|---|---|---|---|---|---|---|---|---|---|---|---|---|---|---|
| Q13043 | 8.83 | 1 | 3 | 3 | 3 |  |  |  |  |  |  |  |  |  |  |  |  |  |  |  |  |  |  |  |  |  |  |  |  | 5.54 | 2 | 2 | 3.29 | 1 | 1 | 487 | 55.6 | 5.07 |
| Q13045 | 14.58 | 1 | 15 | 15 | 22 |  |  |  |  |  |  | 0.95 | 1 | 1 |  |  |  | 0.00 |  |  |  |  |  |  |  |  |  |  |  | 0.63 | 1 | 1 | 13.95 | 14 | 20 | 1269 | 144.7 | 6.05 |
| Q13057 | 46.63 | 1 | 18 | 18 | 44 | 2.13 | 1 | 1 | 2.66 | 1 | 1 |  |  |  | 2.66 | 1 | 1 |  |  |  |  |  |  |  |  |  |  |  |  | 5.85 | 2 | 2 | 45.39 | 16 | 39 | 564 | 62.3 | 6.99 |
| Q13085 | 19.78 | 1 | 37 | 41 | 72 | 0.00 |  |  |  |  |  |  |  |  | 0.30 | 1 | 1 |  |  |  |  |  |  |  |  |  |  |  |  |  |  |  | 19.78 | 41 | 71 | 2346 | 265.4 | 6.37 |
| Q13093 | 32.43 | 1 | 11 | 11 | 28 | 6.35 | 2 | 2 | 1.59 | 1 | 1 | 3.63 | 1 | 1 |  |  |  |  |  |  |  |  |  |  |  |  |  |  |  |  |  |  | 30.84 | 10 | 24 | 441 | 50.0 | 7.56 |
| Q13098 | 27.70 | 1 | 10 | 10 | 18 |  |  |  |  |  |  |  |  |  |  |  |  |  |  |  |  |  |  |  |  |  |  |  |  | 1.63 | 1 | 1 | 26.07 | 9 | 17 | 491 | 55.5 | 6.74 |
| Q13099 | 2.04 | 1 | 2 | 2 | 2 |  |  |  |  |  |  |  |  |  |  |  |  |  |  |  |  |  |  |  |  |  |  |  |  |  |  |  | 2.04 | 2 | 2 | 833 | 94.2 | 6.54 |
| Q13107 | 18.07 | 3 | 13 | 13 | 18 | 3.12 | 1 | 1 |  |  |  |  |  |  |  |  |  |  |  |  |  |  |  |  |  |  |  |  |  |  |  |  | 14.95 | 12 | 17 | 963 | 108.5 | 5.71 |
| Q13126 | 36.75 | 1 | 7 | 7 | 10 |  |  |  |  |  |  |  |  |  |  |  |  |  |  |  |  |  |  |  |  |  |  |  |  |  |  |  | 36.75 | 7 | 10 | 283 | 31.2 | 7.18 |
| P28340 | 1.26 | 1 | 1 | 2 | 16 | 0.63 | 1 | 1 |  |  |  | 0.63 | 1 | 1 | 0.63 | 1 | 1 | 0.63 | 1 | 1 |  |  |  | 0.63 | 1 | 1 |  |  |  | 0.63 | 1 | 1 | 1.26 | 2 | 10 | 1107 | 123.6 | 7.03 |
| Q13131 | 14.13 | 2 | 6 | 6 | 14 |  |  |  |  |  |  |  |  |  |  |  |  |  |  |  |  |  |  |  |  |  |  |  |  |  |  |  | 14.13 | 6 | 14 | 559 | 64.0 | 8.12 |
| Q13137 | 4.71 | 1 | 2 | 2 | 3 |  |  |  |  |  |  |  |  |  |  |  |  |  |  |  |  |  |  |  |  |  |  |  |  | 2.24 | 1 | 1 | 2.47 | 1 | 2 | 446 | 52.2 | 5.02 |
| Q13144 | 12.62 | 1 | 8 | 8 | 17 |  |  |  |  |  |  |  |  |  |  |  |  |  |  |  |  |  |  |  |  |  |  |  |  |  |  |  | 12.62 | 8 | 17 | 721 | 80.3 | 5.08 |
| Q13148 | 17.39 | 1 | 8 | 8 | 19 |  |  |  |  |  |  | 3.14 | 1 | 1 |  |  |  | 3.14 | 1 | 1 |  |  |  |  |  |  | 2.17 | 1 | 1 | 8.21 | 3 | 3 | 17.15 | 7 | 13 | 414 | 44.7 | 6.19 |
| Q9H082 | 9.61 | 7 | 1 | 2 | 16 | 4.80 | 1 | 2 | 4.80 | 1 | 1 | 4.80 | 1 | 1 | 4.80 | 1 | 1 |  |  |  |  |  |  | 4.80 | 1 | 1 |  |  |  |  |  |  | 9.61 | 2 | 10 | 229 | 25.7 | 7.18 |
| Q13155 | 48.75 | 1 | 10 | 10 | 34 | 6.88 | 2 | 2 | 2.81 | 1 | 1 |  |  |  | 7.81 | 1 | 1 |  |  |  |  |  |  |  |  |  |  |  |  |  |  |  | 48.75 | 10 | 30 | 320 | 35.3 | 8.22 |
| Q13162 | 67.16 | 1 | 12 | 15 | 80 | 29.89 | 6 | 8 | 28.41 | 6 | 8 | 32.10 | 7 | 8 | 18.45 | 5 | 6 | 21.03 | 5 | 5 |  |  |  | 6.64 | 2 | 2 | 15.87 | 5 | 6 | 20.30 | 5 | 5 | 64.58 | 14 | 32 | 271 | 30.5 | 6.29 |
| Q13188 | 4.28 | 1 | 2 | 2 | 2 |  |  |  |  |  |  |  |  |  |  |  |  |  |  |  |  |  |  |  |  |  |  |  |  |  |  |  | 4.28 | 2 | 2 | 491 | 56.3 | 5.24 |
| Q13200 | 69.16 | 1 | 54 | 54 | 256 | 16.41 | 8 | 9 | 23.35 | 15 | 17 | 16.74 | 10 | 11 | 12.89 | 10 | 12 | 9.14 | 5 | 6 |  |  |  | 10.35 | 6 | 6 | 8.04 | 5 | 6 | 15.53 | 11 | 14 | 60.13 | 44 | 175 | 908 | 100.1 | 5.20 |
| Q13217 | 47.02 | 1 | 17 | 17 | 33 | 11.31 | 3 | 3 | 9.92 | 3 | 3 |  |  |  |  |  |  |  |  |  |  |  |  |  |  |  |  |  |  |  |  |  | 47.02 | 17 | 27 | 504 | 57.5 | 6.15 |
| Q13228 | 20.34 | 1 | 6 | 6 | 7 |  |  |  |  |  |  | 6.78 | 2 | 2 |  |  |  |  |  |  |  |  |  |  |  |  |  |  |  |  |  |  | 13.56 | 4 | 5 | 472 | 52.4 | 6.37 |
| Q13231 | 7.08 | 1 | 3 | 3 | 5 | 7.08 | 2 | 2 |  |  |  |  |  |  |  |  |  |  |  |  |  |  |  |  |  |  |  |  |  |  |  |  | 4.51 | 1 | 3 | 466 | 51.6 | 6.96 |
| Q13232 | 46.15 | 1 | 6 | 6 | 7 |  |  |  |  |  |  |  |  |  |  |  |  |  |  |  |  |  |  |  |  |  |  |  |  |  |  |  | 46.15 | 6 | 7 | 169 | 19.0 | 7.84 |
| Q13275 | 4.59 | 1 | 3 | 3 | 5 |  |  |  |  |  |  |  |  |  |  |  |  |  |  |  |  |  |  |  |  |  |  |  |  |  |  |  | 4.59 | 3 | 5 | 785 | 88.3 | 8.27 |
| Q13277 | 9.00 | 1 | 3 | 3 | 6 | 3.11 | 1 | 1 | 3.11 | 1 | 1 |  |  |  |  |  |  |  |  |  |  |  |  |  |  |  |  |  |  |  |  |  | 9.00 | 3 | 4 | 289 | 33.1 | 5.44 |
| Q13325 | 4.56 | 1 | 2 | 2 | 3 |  |  |  |  |  |  |  |  |  |  |  |  |  |  |  |  |  |  |  |  |  |  |  |  |  |  |  | 4.56 | 2 | 3 | 482 | 55.8 | 7.40 |
| Q13332 | 1.59 | 2 | 2 | 2 | 2 |  |  |  | 0.00 |  |  | 1.59 | 2 | 2 |  |  |  |  |  |  |  |  |  |  |  |  |  |  |  |  |  |  |  |  |  | 1948 | 216.9 | 6.46 |
| Q13347 | 59.38 | 1 | 14 | 14 | 40 | 3.69 | 1 | 1 |  |  |  | 3.69 | 1 | 1 | 15.08 | 4 | 5 |  |  |  |  |  |  | 4.92 | 1 | 1 | 22.77 | 4 | 4 | 9.23 | 3 | 3 | 50.77 | 11 | 25 | 325 | 36.5 | 5.64 |
| Q13363 | 4.32 | 1 | 2 | 2 | 2 |  |  |  |  |  |  |  |  |  |  |  |  |  |  |  |  |  |  |  |  |  |  |  |  |  |  |  | 4.32 | 2 | 2 | 440 | 47.5 | 6.77 |
| Q13404 | 22.45 | 2 | 3 | 3 | 5 |  |  |  | 6.80 | 1 | 1 |  |  |  |  |  |  |  |  |  |  |  |  |  |  |  |  |  |  |  |  |  | 22.45 | 3 | 4 | 147 | 16.5 | 7.93 |
| Q13409 | 19.91 | 1 | 8 | 8 | 16 | 2.19 | 1 | 1 | 0.00 |  |  | 5.49 | 2 | 2 | 3.29 | 1 | 1 | 5.49 | 2 | 2 |  |  |  |  |  |  |  |  |  |  |  |  | 19.91 | 7 | 10 | 638 | 71.4 | 5.20 |
| Q13421 | 9.21 | 1 | 5 | 5 | 6 |  |  |  |  |  |  |  |  |  |  |  |  |  |  |  |  |  |  |  |  |  |  |  |  |  |  |  | 9.21 | 5 | 6 | 630 | 68.9 | 6.38 |
| Q13423 | 7.64 | 1 | 6 | 6 | 12 |  |  |  |  |  |  |  |  |  |  |  |  |  |  |  |  |  |  |  |  |  |  |  |  |  |  |  | 7.64 | 6 | 12 | 1086 | 113.8 | 8.09 |
| P01834 | 79.44 | 1 | 1 | 5 | 16 | 34.58 | 2 | 2 | 18.69 | 1 | 1 | 47.66 | 3 | 4 |  |  |  | 15.89 | 1 | 1 |  |  |  |  |  |  | 15.89 | 1 | 1 |  |  |  | 66.36 | 4 | 7 | 107 | 11.8 | 6.52 |
| P0DOX7 | 34.11 | 16 | 1 | 5 | 16 | 17.29 | 2 | 2 | 9.35 | 1 | 1 | 23.83 | 3 | 4 |  |  |  | 7.94 | 1 | 1 |  |  |  |  |  |  | 7.94 | 1 | 1 |  |  |  | 27.57 | 4 | 7 | 214 | 23.4 | 7.17 |
| Q13438 | 14.09 | 1 | 5 | 6 | 11 | 6.60 | 3 | 4 | 1.65 | 1 | 1 | 5.40 | 2 | 2 | 3.00 | 2 | 2 | 2.10 | 1 | 1 |  |  |  |  |  |  |  |  |  | 2.10 | 1 | 1 |  |  |  | 667 | 75.5 | 4.87 |
| Q13439 | 0.54 | 1 | 2 | 2 | 2 |  |  |  | 0.27 | 1 | 1 |  |  |  |  |  |  |  |  |  |  |  |  |  |  |  |  |  |  |  |  |  | 0.27 | 1 | 1 | 2230 | 261.0 | 5.39 |
| Q13443 | 3.17 | 1 | 2 | 2 | 8 |  |  |  |  |  |  | 1.71 | 1 | 1 |  |  |  |  |  |  |  |  |  |  |  |  |  |  |  |  |  |  | 3.17 | 2 | 7 | 819 | 90.5 | 7.52 |
| Q13445 | 18.50 | 1 | 4 | 4 | 5 | 3.96 | 1 | 1 | 3.96 | 1 | 1 |  |  |  |  |  |  |  |  |  |  |  |  |  |  |  |  |  |  |  |  |  | 14.54 | 3 | 3 | 227 | 25.2 | 4.48 |
| Q13449 | 11.83 | 1 | 3 | 3 | 3 | 2.66 | 1 | 1 |  |  |  | 9.17 | 2 | 2 |  |  |  |  |  |  |  |  |  |  |  |  |  |  |  |  |  |  |  |  |  | 338 | 37.4 | 6.98 |
| Q13451 | 30.42 | 1 | 12 | 12 | 31 |  |  |  |  |  |  |  |  |  |  |  |  |  |  |  |  |  |  | 3.94 | 2 | 2 | 4.38 | 2 | 2 | 18.82 | 7 | 7 | 20.13 | 8 | 20 | 457 | 51.2 | 5.90 |
| Q13454 | 5.75 | 1 | 2 | 2 | 2 |  |  |  |  |  |  |  |  |  |  |  |  |  |  |  |  |  |  |  |  |  |  |  |  |  |  |  | 5.75 | 2 | 2 | 348 | 39.7 | 9.91 |
| Q13459 | 1.21 | 1 | 2 | 2 | 3 |  |  |  |  |  |  |  |  |  |  |  |  |  |  |  |  |  |  |  |  |  |  |  |  |  |  |  | 1.21 | 2 | 3 | 2157 | 243.2 | 8.75 |
| Q13464 | 12.33 | 1 | 14 | 17 | 19 |  |  |  |  |  |  |  |  |  |  |  |  |  |  |  |  |  |  |  |  |  |  |  |  | 2.14 | 2 | 2 | 11.82 | 16 | 17 | 1354 | 158.1 | 5.90 |
| Q13485 | 5.43 | 1 | 2 | 2 | 2 |  |  |  |  |  |  |  |  |  |  |  |  |  |  |  |  |  |  |  |  |  |  |  |  |  |  |  | 5.43 | 2 | 2 | 552 | 60.4 | 6.99 |
| Q13501 | 16.36 | 1 | 4 | 4 | 8 |  |  |  |  |  |  |  |  |  | 8.64 | 2 | 2 | 2.73 | 1 | 1 |  |  |  |  |  |  |  |  |  | 2.73 | 1 | 1 | 7.73 | 2 | 4 | 440 | 47.7 | 5.22 |
| Q13505 | 22.32 | 1 | 7 | 8 | 13 |  |  |  | 3.22 | 1 | 1 |  |  |  |  |  |  |  |  |  |  |  |  |  |  |  |  |  |  |  |  |  | 22.32 | 8 | 12 | 466 | 51.4 | 9.79 |
| Q13508 | 35.99 | 4 | 10 | 11 | 68 | 23.39 | 6 | 12 | 23.91 | 7 | 10 | 26.99 | 6 | 10 | 6.17 | 2 | 2 | 9.77 | 3 | 5 |  |  |  |  |  |  |  |  |  |  |  |  | 33.42 | 10 | 29 | 389 | 43.9 | 6.06 |
| Q13509 | 43.33 | 1 | 2 | 21 | 594 | 30.00 | 14 | 29 | 30.00 | 14 | 40 | 26.89 | 13 | 26 | 30.22 | 17 | 43 | 30.89 | 17 | 46 | 4.22 | 1 | 2 | 25.56 | 10 | 20 | 24.44 | 10 | 19 | 24.44 | 10 | 21 | 39.11 | 17 | 348 | 450 | 50.4 | 4.93 |
| Q13510 | 29.11 | 1 | 10 | 10 | 25 | 11.14 | 3 | 3 | 9.37 | 3 | 3 | 14.18 | 6 | 6 |  |  |  |  |  |  |  |  |  |  |  |  |  |  |  |  |  |  | 27.09 | 9 | 13 | 395 | 44.6 | 7.62 |
| Q13526 | 63.80 | 2 | 7 | 7 | 14 |  |  |  |  |  |  |  |  |  |  |  |  |  |  |  |  |  |  |  |  |  | 9.20 | 1 | 2 | 15.34 | 2 | 4 | 57.67 | 6 | 8 | 163 | 18.2 | 8.82 |
| Q13546 | 4.92 | 1 | 2 | 2 | 3 |  |  |  |  |  |  |  |  |  |  |  |  |  |  |  |  |  |  |  |  |  |  |  |  |  |  |  | 4.92 | 2 | 3 | 671 | 75.9 | 6.33 |
| Q13555 | 11.83 | 1 | 2 | 5 | 19 |  |  |  |  |  |  | 2.15 | 1 | 1 | 2.15 | 1 | 2 |  |  |  |  |  |  | 3.41 | 2 | 2 |  |  |  | 5.56 | 2 | 2 | 11.83 | 5 | 12 | 558 | 62.6 | 7.83 |
| Q13557 | 41.28 | 3 | 12 | 15 | 67 | 11.22 | 4 | 4 |  |  |  | 7.01 | 3 | 3 | 5.61 | 2 | 3 |  |  |  |  |  |  | 13.23 | 6 | 8 |  |  |  | 20.84 | 7 | 8 | 30.46 | 11 | 41 | 499 | 56.3 | 7.25 |
| Q13561 | 46.63 | 1 | 17 | 17 | 68 | 27.43 | 8 | 8 | 31.42 | 9 | 11 | 17.46 | 5 | 5 | 19.20 | 6 | 6 | 20.20 | 7 | 10 |  |  |  |  |  |  | 6.23 | 2 | 2 | 17.96 | 5 | 5 | 35.66 | 11 | 21 | 401 | 44.2 | 5.21 |
| Q13564 | 18.16 | 1 | 7 | 7 | 11 |  |  |  |  |  |  |  |  |  |  |  |  |  |  |  |  |  |  |  |  |  |  |  |  |  |  |  | 18.16 | 7 | 11 | 534 | 60.2 | 5.40 |
| Q13572 | 10.39 | 1 | 3 | 3 | 4 |  |  |  |  |  |  |  |  |  |  |  |  |  |  |  |  |  |  |  |  |  |  |  |  |  |  |  | 10.39 | 3 | 4 | 414 | 45.6 | 6.16 |
| Q13576 | 39.17 | 2 | 49 | 53 | 117 | 0.83 | 1 | 1 |  |  |  | 1.65 | 2 | 2 | 0.83 | 1 | 2 |  |  |  |  |  |  |  |  |  |  |  |  | 0.76 | 1 | 1 | 38.35 | 52 | 111 | 1575 | 180.5 | 5.64 |
| Q13596 | 18.77 | 1 | 6 | 8 | 21 |  |  |  |  |  |  |  |  |  | 4.02 | 2 | 2 |  |  |  |  |  |  | 2.11 | 1 | 1 |  |  |  | 16.86 | 7 | 7 | 13.79 | 6 | 11 | 522 | 59.0 | 5.15 |
| Q13616 | 8.25 | 2 | 6 | 7 | 7 |  |  |  |  |  |  |  |  |  |  |  |  |  |  |  |  |  |  |  |  |  |  |  |  |  |  |  | 8.25 | 7 | 7 | 776 | 89.6 | 8.00 |
| Q13617 | 18.39 | 1 | 12 | 12 | 16 |  |  |  | 2.68 | 2 | 2 | 3.09 | 2 | 2 |  |  |  |  |  |  |  |  |  |  |  |  |  |  |  |  |  |  | 15.44 | 10 | 12 | 745 | 86.9 | 6.92 |
| Q13618 | 58.33 | 2 | 41 | 42 | 135 | 13.41 | 10 | 10 | 18.49 | 13 | 13 | 15.10 | 12 | 12 | 1.69 | 1 | 1 | 2.73 | 2 | 2 |  |  |  |  |  |  |  |  |  |  |  |  | 57.42 | 40 | 97 | 768 | 88.9 | 8.48 |
| Q13619 | 19.10 | 1 | 8 | 13 | 19 | 3.03 | 2 | 2 |  |  |  |  |  |  |  |  |  |  |  |  |  |  |  |  |  |  |  |  |  |  |  |  | 19.10 | 12 | 17 | 759 | 87.6 | 8.13 |
| P61956 | 12.63 | 3 | 1 | 1 | 15 | 12.63 | 1 | 1 | 12.63 | 1 | 1 | 12.63 | 1 | 1 | 12.63 | 1 | 2 | 12.63 | 1 | 1 |  |  |  |  |  |  | 12.63 | 1 | 1 | 12.63 | 1 | 1 | 12.63 | 1 | 7 | 95 | 10.9 | 5.50 |
| Q13620 | 8.11 | 1 | 3 | 8 | 12 | 1.10 | 1 | 1 |  |  |  |  |  |  |  |  |  |  |  |  |  |  |  |  |  |  |  |  |  |  |  |  | 8.11 | 7 | 11 | 913 | 103.9 | 7.37 |
| Q13630 | 40.19 | 1 | 9 | 9 | 25 |  |  |  | 3.43 | 1 | 1 |  |  |  | 13.08 | 3 | 3 |  |  |  |  |  |  |  |  |  | 3.74 | 1 | 1 | 7.17 | 2 | 2 | 40.19 | 9 | 18 | 321 | 35.9 | 6.60 |
| Q13637 | 15.56 | 3 | 3 | 3 | 3 |  |  |  |  |  |  |  |  |  |  |  |  |  |  |  |  |  |  |  |  |  |  |  |  |  |  |  | 15.56 | 3 | 3 | 225 | 25.0 | 6.54 |
| Q13642 | 40.25 | 1 | 11 | 11 | 32 | 6.19 | 2 | 2 |  |  |  | 15.48 | 4 | 5 |  |  |  | 6.50 | 2 | 2 |  |  |  |  |  |  | 13.00 | 4 | 4 | 3.72 | 1 | 1 | 38.39 | 10 | 18 | 323 | 36.2 | 8.97 |
| Q13685 | 5.53 | 1 | 2 | 2 | 2 |  |  |  |  |  |  |  |  |  |  |  |  |  |  |  |  |  |  |  |  |  | 3.00 | 1 | 1 |  |  |  | 2.53 | 1 | 1 | 434 | 46.7 | 4.42 |
| Q13724 | 31.90 | 1 | 19 | 19 | 37 | 3.94 | 2 | 2 | 9.20 | 5 | 6 | 3.35 | 2 | 2 |  |  |  |  |  |  |  |  |  |  |  |  |  |  |  |  |  |  | 25.93 | 15 | 27 | 837 | 91.9 | 8.90 |
| Q13733 | 47.04 | 1 | 31 | 46 | 345 | 23.91 | 19 | 22 | 28.67 | 26 | 33 | 27.31 | 23 | 26 | 1.85 | 2 | 2 | 5.73 | 5 | 5 |  |  |  |  |  |  |  |  |  |  |  |  | 46.94 | 45 | 257 | 1029 | 114.1 | 6.64 |
| Q13748 | 85.33 | 2 | 11 | 46 | 1082 | 60.00 | 22 | 42 | 67.33 | 25 | 61 | 60.00 | 22 | 35 | 58.22 | 28 | 82 | 54.44 | 29 | 60 | 9.11 | 4 | 5 | 31.78 | 13 | 18 | 44.67 | 16 | 22 | 54.67 | 19 | 34 | 82.89 | 36 | 723 | 450 | 49.9 | 5.10 |
| Q13765 | 13.49 | 2 | 2 | 2 | 5 | 6.51 | 1 | 1 |  |  |  |  |  |  |  |  |  |  |  |  |  |  |  |  |  |  |  |  |  |  |  |  | 13.49 | 2 | 4 | 215 | 23.4 | 4.56 |
| Q13813 | 27.39 | 1 | 53 | 53 | 93 |  |  |  |  |  |  |  |  |  |  |  |  |  |  |  |  |  |  |  |  |  |  |  |  |  |  |  | 27.39 | 53 | 93 | 2472 | 284.4 | 5.35 |
| Q13867 | 7.25 | 1 | 3 | 3 | 7 | 2.42 | 1 | 1 | 2.42 | 1 | 1 | 2.42 | 1 | 1 |  |  |  |  |  |  |  |  |  |  |  |  |  |  |  |  |  |  | 7.25 | 3 | 4 | 455 | 52.5 | 6.27 |
| Q13907 | 29.96 | 3 | 6 | 6 | 10 |  |  |  |  |  |  |  |  |  |  |  |  |  |  |  |  |  |  |  |  |  |  |  |  |  |  |  | 29.96 | 6 | 10 | 227 | 26.3 | 6.34 |
| Q13938 | 39.15 | 1 | 7 | 7 | 7 |  |  |  |  |  |  |  |  |  |  |  |  |  |  |  |  |  |  |  |  |  |  |  |  |  |  |  | 39.15 | 7 | 7 | 189 | 21.0 | 4.89 |
| Q13939 | 38.27 | 1 | 21 | 21 | 45 |  |  |  | 4.08 | 2 | 2 | 4.08 | 2 | 2 |  |  |  |  |  |  |  |  |  |  |  |  |  |  |  |  |  |  | 38.27 | 21 | 41 | 588 | 66.5 | 8.18 |
| Q14008 | 1.87 | 1 | 3 | 3 | 5 |  |  |  |  |  |  |  |  |  |  |  |  |  |  |  |  |  |  |  |  |  |  |  |  |  |  |  | 1.87 | 3 | 5 | 2032 | 225.4 | 7.80 |
| Q14012 | 30.00 | 2 | 7 | 7 | 9 |  |  |  |  |  |  |  |  |  |  |  |  |  |  |  |  |  |  | 2.70 | 1 | 1 |  |  |  |  |  |  | 27.30 | 6 | 8 | 370 | 41.3 | 5.29 |
| Q14525 | 4.70 | 10 | 1 | 2 | 15 |  |  |  | 1.73 | 1 | 1 | 1.73 | 1 | 1 |  |  |  | 1.73 | 1 | 1 |  |  |  | 1.73 | 1 | 1 | 1.73 | 1 | 1 | 1.73 | 1 | 1 | 4.70 | 2 | 9 | 404 | 46.2 | 4.84 |
| Q14019 | 64.79 | 2 | 10 | 10 | 13 |  |  |  |  |  |  |  |  |  |  |  |  |  |  |  |  |  |  |  |  |  |  |  |  | 16.90 | 3 | 3 | 59.15 | 9 | 10 | 142 | 15.9 | 5.67 |
| Q14093 | 11.21 | 1 | 3 | 3 | 6 |  |  |  | 3.45 | 1 | 1 | 2.87 | 1 | 1 | 3.45 | 1 | 1 |  |  |  |  |  |  |  |  |  |  |  |  |  |  |  | 8.33 | 2 | 3 | 348 | 39.1 | 9.74 |
| Q14103 | 5.92 | 1 | 2 | 2 | 2 |  |  |  |  |  |  |  |  |  |  |  |  |  |  |  |  |  |  |  |  |  |  |  |  |  |  |  | 5.92 | 2 | 2 | 355 | 38.4 | 7.81 |
| Q14108 | 25.31 | 1 | 8 | 8 | 15 |  |  |  |  |  |  | 14.44 | 4 | 4 |  |  |  |  |  |  |  |  |  |  |  |  |  |  |  |  |  |  | 16.11 | 6 | 11 | 478 | 54.3 | 5.14 |
| Q14139 | 12.57 | 2 | 11 | 11 | 12 |  |  |  | 0.00 |  |  |  |  |  |  |  |  |  |  |  |  |  |  |  |  |  |  |  |  |  |  |  | 12.57 | 11 | 12 | 1066 | 122.5 | 5.24 |
| Q14141 | 29.26 | 2 | 5 | 10 | 24 | 2.07 | 1 | 1 | 2.07 | 1 | 1 | 2.07 | 1 | 1 | 2.07 | 1 | 1 | 3.46 | 2 | 2 |  |  |  |  |  |  |  |  |  |  |  |  | 29.26 | 10 | 18 | 434 | 49.7 | 6.67 |
| Q14145 | 3.04 | 1 | 2 | 2 | 2 |  |  |  |  |  |  |  |  |  |  |  |  |  |  |  |  |  |  |  |  |  |  |  |  |  |  |  | 3.04 | 2 | 2 | 624 | 69.6 | 6.44 |
| Q14152 | 22.43 | 2 | 30 | 30 | 70 | 0.65 | 1 | 1 | 2.24 | 3 | 3 | 0.87 | 1 | 1 |  |  |  |  |  |  |  |  |  |  |  |  | 0.51 | 1 | 1 |  |  |  | 20.55 | 27 | 64 | 1382 | 166.5 | 6.79 |
| Q14156 | 19.24 | 2 | 10 | 10 | 25 |  |  |  |  |  |  |  |  |  |  |  |  |  |  |  |  |  |  |  |  |  |  |  |  |  |  |  | 19.24 | 10 | 25 | 821 | 92.9 | 6.70 |
| Q14165 | 42.81 | 1 | 10 | 10 | 49 | 19.18 | 5 | 6 | 21.58 | 6 | 7 | 16.78 | 4 | 5 | 5.82 | 1 | 1 |  |  |  |  |  |  | 4.11 | 1 | 1 |  |  |  |  |  |  | 42.81 | 10 | 29 | 292 | 32.2 | 5.41 |
| Q14166 | 44.41 | 1 | 19 | 19 | 80 | 3.11 | 1 | 2 | 3.11 | 1 | 2 | 3.11 | 1 | 1 | 2.33 | 1 | 1 |  |  |  |  |  |  | 2.33 | 1 | 1 | 2.33 | 1 | 1 | 8.54 | 4 | 5 | 42.55 | 18 | 67 | 644 | 74.4 | 5.53 |
| Q14185 | 4.93 | 1 | 7 | 8 | 9 |  |  |  |  |  |  |  |  |  |  |  |  |  |  |  |  |  |  |  |  |  |  |  |  |  |  |  | 4.93 | 8 | 9 | 1865 | 215.2 | 7.56 |
| Q14195 | 7.37 | 3 | 3 | 3 | 8 |  |  |  |  |  |  |  |  |  |  |  |  |  |  |  |  |  |  |  |  |  | 1.75 | 1 | 1 | 4.56 | 2 | 3 | 5.61 | 2 | 4 | 570 | 61.9 | 6.49 |
| Q14203 | 50.47 | 2 | 57 | 58 | 161 | 10.09 | 10 | 12 | 10.72 | 11 | 12 | 5.01 | 5 | 5 | 4.46 | 5 | 6 | 4.30 | 5 | 5 |  |  |  | 3.60 | 4 | 4 | 6.26 | 7 | 9 | 12.52 | 13 | 14 | 45.54 | 47 | 94 | 1278 | 141.6 | 5.81 |
| Q14204 | 47.27 | 3 | 200 | 201 | 618 | 0.65 | 3 | 3 |  |  |  | 5.47 | 21 | 21 | 4.20 | 18 | 18 | 0.24 | 1 | 1 |  |  |  | 0.60 | 3 | 3 | 0.30 | 1 | 1 | 4.52 | 18 | 20 | 45.16 | 190 | 551 | 4646 | 532.1 | 6.40 |
| Q9Y4G6 | 2.79 | 1 | 1 | 7 | 15 |  |  |  |  |  |  |  |  |  |  |  |  |  |  |  |  |  |  |  |  |  |  |  |  | 0.87 | 2 | 2 | 2.79 | 7 | 13 | 2542 | 271.4 | 5.57 |
| Q14232 | 37.70 | 1 | 10 | 10 | 15 |  |  |  |  |  |  |  |  |  |  |  |  |  |  |  |  |  |  |  |  |  |  |  |  |  |  |  | 37.70 | 10 | 15 | 305 | 33.7 | 7.33 |
| Q14240 | 51.35 | 1 | 8 | 22 | 86 | 10.81 | 4 | 4 | 7.62 | 3 | 3 | 5.90 | 2 | 2 | 16.22 | 5 | 5 | 6.39 | 2 | 2 |  |  |  | 15.48 | 5 | 5 | 9.83 | 3 | 3 | 12.04 | 4 | 5 | 50.86 | 20 | 57 | 407 | 46.4 | 5.48 |
| Q14247 | 5.64 | 1 | 2 | 2 | 2 |  |  |  |  |  |  |  |  |  |  |  |  |  |  |  |  |  |  |  |  |  | 3.27 | 1 | 1 |  |  |  | 2.36 | 1 | 1 | 550 | 61.5 | 5.40 |
| Q14254 | 40.89 | 1 | 17 | 17 | 27 | 7.94 | 3 | 3 |  |  |  | 19.86 | 7 | 7 | 2.80 | 1 | 1 |  |  |  |  |  |  |  |  |  |  |  |  |  |  |  | 31.78 | 13 | 16 | 428 | 47.0 | 5.25 |
| Q14257 | 47.63 | 1 | 12 | 12 | 63 | 18.93 | 4 | 5 | 45.43 | 9 | 14 | 17.03 | 4 | 4 | 29.02 | 6 | 7 | 22.08 | 5 | 7 |  |  |  | 29.02 | 7 | 10 | 8.83 | 3 | 4 | 5.36 | 2 | 2 | 17.03 | 3 | 10 | 317 | 36.9 | 4.40 |
| Q14258 | 21.27 | 1 | 12 | 12 | 26 |  |  |  |  |  |  |  |  |  |  |  |  |  |  |  |  |  |  | 2.06 | 1 | 1 |  |  |  | 4.13 | 2 | 3 | 21.11 | 11 | 22 | 630 | 70.9 | 8.09 |
| Q14289 | 2.38 | 2 | 3 | 3 | 5 |  |  |  | 0.59 | 1 | 1 | 1.59 | 2 | 2 |  |  |  |  |  |  |  |  |  |  |  |  |  |  |  |  |  |  | 1.39 | 2 | 2 | 1009 | 115.8 | 6.25 |
| Q14344 | 23.87 | 3 | 6 | 8 | 23 | 6.10 | 2 | 2 |  |  |  | 2.92 | 1 | 1 | 2.92 | 1 | 1 | 2.92 | 1 | 1 |  |  |  |  |  |  | 2.92 | 1 | 1 | 2.92 | 1 | 1 | 23.87 | 8 | 16 | 377 | 44.0 | 8.00 |
| Q14353 | 43.64 | 1 | 5 | 5 | 7 |  |  |  |  |  |  |  |  |  |  |  |  |  |  |  |  |  |  |  |  |  |  |  |  |  |  |  | 43.64 | 5 | 7 | 236 | 26.3 | 6.14 |
| Q14376 | 16.95 | 1 | 5 | 5 | 14 |  |  |  |  |  |  |  |  |  |  |  |  |  |  |  |  |  |  |  |  |  |  |  |  |  |  |  | 16.95 | 5 | 14 | 348 | 38.3 | 6.73 |
| Q14393 | 33.01 | 2 | 18 | 18 | 37 | 8.18 | 4 | 4 | 3.05 | 2 | 2 | 7.35 | 4 | 4 |  |  |  |  |  |  |  |  |  |  |  |  |  |  |  |  |  |  | 27.74 | 15 | 27 | 721 | 79.6 | 6.21 |
| Q14409 | 63.83 | 1 | 3 | 27 | 93 | 24.95 | 10 | 15 | 20.25 | 10 | 12 | 12.48 | 5 | 7 | 2.35 | 1 | 1 |  |  |  |  |  |  |  |  |  |  |  |  |  |  |  | 61.66 | 26 | 58 | 553 | 60.6 | 6.39 |
| Q14410 | 59.67 | 1 | 21 | 28 | 116 | 27.12 | 11 | 16 | 28.39 | 15 | 20 | 17.72 | 9 | 10 | 2.35 | 1 | 1 | 1.81 | 1 | 1 |  |  |  |  |  |  |  |  |  |  |  |  | 58.59 | 25 | 68 | 553 | 60.6 | 5.77 |
| Q14435 | 15.80 | 2 | 10 | 10 | 12 |  |  |  | 3.63 | 2 | 2 | 2.05 | 1 | 1 |  |  |  |  |  |  |  |  |  |  |  |  |  |  |  |  |  |  | 14.22 | 9 | 9 | 633 | 72.6 | 7.99 |
| Q14444 | 3.24 | 1 | 2 | 2 | 3 |  |  |  |  |  |  |  |  |  |  |  |  |  |  |  |  |  |  |  |  |  | 1.69 | 1 | 1 | 3.24 | 2 | 2 |  |  |  | 709 | 78.3 | 5.25 |
| Q14507 | 37.41 | 1 | 4 | 5 | 10 |  |  |  | 7.48 | 1 | 1 |  |  |  |  |  |  |  |  |  |  |  |  |  |  |  |  |  |  |  |  |  | 37.41 | 5 | 9 | 147 | 17.6 | 8.35 |
| CON_009038861 | 40.23 | 1 | 1 | 7 | 14 | 30.08 | 5 | 6 | 19.55 | 4 | 4 | 22.18 | 4 | 4 |  |  |  |  |  |  |  |  |  |  |  |  |  |  |  |  |  |  |  |  |  | 266 | 30.1 | 9.04 |
| Q14508 | 38.71 | 1 | 3 | 3 | 8 | 38.71 | 3 | 4 |  |  |  | 11.29 | 1 | 1 | 11.29 | 1 | 1 | 11.29 | 1 | 1 |  |  |  |  |  |  |  |  |  |  |  |  | 11.29 | 1 | 1 | 124 | 13.0 | 4.84 |
| Q14558 | 38.20 | 1 | 8 | 9 | 14 |  |  |  |  |  |  |  |  |  |  |  |  |  |  |  |  |  |  |  |  |  |  |  |  |  |  |  | 38.20 | 9 | 14 | 356 | 39.4 | 7.20 |
| Q14642 | 19.42 | 1 | 7 | 7 | 9 | 0.00 |  |  |  |  |  |  |  |  |  |  |  |  |  |  |  |  |  |  |  |  |  |  |  |  |  |  | 19.42 | 7 | 9 | 412 | 47.8 | 7.03 |
| Q14651 | 10.97 | 1 | 2 | 7 | 27 | 5.72 | 3 | 3 | 3.50 | 2 | 2 | 5.72 | 3 | 4 | 3.97 | 2 | 2 |  |  |  |  |  |  |  |  |  |  |  |  |  |  |  | 8.74 | 6 | 16 | 629 | 70.2 | 5.41 |
| Q14693 | 3.48 | 1 | 2 | 2 | 4 |  |  |  |  |  |  |  |  |  |  |  |  |  |  |  |  |  |  |  |  |  |  |  |  |  |  |  | 3.48 | 2 | 4 | 890 | 98.6 | 6.58 |
| Q14696 | 19.66 | 1 | 5 | 5 | 5 | 5.56 | 1 | 1 | 11.11 | 3 | 3 |  |  |  |  |  |  |  |  |  |  |  |  |  |  |  |  |  |  |  |  |  | 8.55 | 1 | 1 | 234 | 26.1 | 7.78 |
| Q14697 | 52.97 | 1 | 44 | 44 | 278 | 26.69 | 18 | 23 | 29.45 | 23 | 31 | 44.07 | 31 | 44 | 12.82 | 10 | 10 | 6.14 | 5 | 5 |  |  |  | 9.00 | 7 | 7 | 4.98 | 3 | 4 | 11.97 | 9 | 9 | 41.63 | 34 | 145 | 944 | 106.8 | 6.14 |
| Q14738 | 14.78 | 1 | 3 | 7 | 19 |  |  |  |  |  |  |  |  |  |  |  |  |  |  |  |  |  |  |  |  |  |  |  |  |  |  |  | 14.78 | 7 | 19 | 602 | 69.9 | 8.13 |
| Q14739 | 5.53 | 1 | 3 | 3 | 4 |  |  |  |  |  |  | 1.30 | 1 | 1 |  |  |  |  |  |  |  |  |  |  |  |  |  |  |  |  |  |  | 5.53 | 3 | 3 | 615 | 70.7 | 9.36 |
| Q14746 | 34.42 | 1 | 16 | 17 | 20 |  |  |  |  |  |  |  |  |  |  |  |  |  |  |  |  |  |  |  |  |  |  |  |  |  |  |  | 34.42 | 17 | 20 | 738 | 83.2 | 6.62 |
| Q14749 | 23.73 | 1 | 5 | 5 | 12 | 10.17 | 2 | 2 |  |  |  |  |  |  | 3.39 | 1 | 1 |  |  |  |  |  |  | 6.78 | 1 | 1 |  |  |  |  |  |  | 12.88 | 3 | 8 | 295 | 32.7 | 7.02 |
| Q14764 | 35.50 | 1 | 22 | 22 | 29 | 1.46 | 1 | 1 |  |  |  | 4.48 | 3 | 3 |  |  |  |  |  |  |  |  |  |  |  |  |  |  |  |  |  |  | 32.81 | 20 | 25 | 893 | 99.3 | 5.48 |
| Q14847 | 9.20 | 1 | 2 | 2 | 3 |  |  |  |  |  |  |  |  |  |  |  |  |  |  |  |  |  |  |  |  |  |  |  |  | 3.83 | 1 | 1 | 9.20 | 2 | 2 | 261 | 29.7 | 7.05 |
| Q14894 | 44.27 | 1 | 12 | 12 | 47 |  |  |  |  |  |  | 3.18 | 1 | 1 | 11.46 | 3 | 3 |  |  |  |  |  |  | 25.80 | 7 | 7 | 12.10 | 3 | 4 | 20.70 | 5 | 5 | 43.63 | 11 | 27 | 314 | 33.8 | 5.14 |
| Q14914 | 43.47 | 1 | 13 | 13 | 53 |  |  |  |  |  |  | 3.65 | 1 | 2 | 7.90 | 2 | 2 |  |  |  |  |  |  |  |  |  |  |  |  | 9.12 | 3 | 5 | 43.47 | 13 | 44 | 329 | 35.8 | 8.29 |
| Q14974 | 49.89 | 1 | 33 | 33 | 229 | 11.30 | 8 | 8 | 16.44 | 12 | 13 | 16.67 | 11 | 11 | 8.90 | 6 | 7 |  |  |  |  |  |  | 1.37 | 1 | 1 |  |  |  | 4.57 | 4 | 5 | 47.26 | 32 | 184 | 876 | 97.1 | 4.78 |
| Q14980 | 2.17 | 1 | 4 | 4 | 4 |  |  |  |  |  |  |  |  |  |  |  |  | 0.00 |  |  |  |  |  |  |  |  |  |  |  |  |  |  | 2.17 | 4 | 4 | 2115 | 238.1 | 5.78 |
| Q14990 | 70.00 | 1 | 23 | 23 | 253 | 18.80 | 4 | 4 | 8.40 | 3 | 3 | 37.20 | 10 | 10 | 38.40 | 12 | 14 | 38.80 | 8 | 10 |  |  |  | 9.60 | 2 | 2 | 36.40 | 11 | 16 | 55.60 | 14 | 18 | 61.20 | 16 | 176 | 250 | 28.3 | 8.03 |
| Q14997 | 34.89 | 4 | 57 | 58 | 158 | 6.19 | 8 | 10 | 8.30 | 10 | 11 | 3.36 | 4 | 4 |  |  |  | 0.54 | 1 | 1 |  |  |  |  |  |  |  |  |  |  |  |  | 33.32 | 55 | 132 | 1843 | 211.2 | 6.90 |
| Q149M9 | 4.67 | 1 | 7 | 7 | 10 |  |  |  |  |  |  |  |  |  |  |  |  |  |  |  |  |  |  |  |  |  |  |  |  |  |  |  | 4.67 | 7 | 10 | 1564 | 174.4 | 6.74 |
| Q14CX7 | 2.47 | 1 | 2 | 2 | 2 |  |  |  |  |  |  |  |  |  |  |  |  |  |  |  |  |  |  |  |  |  |  |  |  |  |  |  | 2.47 | 2 | 2 | 972 | 112.2 | 6.64 |
| Q15005 | 61.50 | 1 | 19 | 19 | 73 | 27.43 | 6 | 8 | 31.42 | 7 | 12 | 24.34 | 4 | 5 |  |  |  | 8.41 | 2 | 2 |  |  |  |  |  |  |  |  |  |  |  |  | 61.06 | 17 | 46 | 226 | 25.0 | 8.47 |
| Q15006 | 14.81 | 1 | 3 | 3 | 3 |  |  |  |  |  |  |  |  |  |  |  |  |  |  |  |  |  |  |  |  |  |  |  |  |  |  |  | 14.81 | 3 | 3 | 297 | 34.8 | 6.57 |
| Q15008 | 67.35 | 1 | 29 | 29 | 83 | 9.51 | 3 | 3 | 14.65 | 5 | 6 | 9.77 | 3 | 3 | 21.08 | 8 | 11 | 19.28 | 8 | 8 |  |  |  |  |  |  |  |  |  | 5.91 | 2 | 2 | 63.50 | 25 | 50 | 389 | 45.5 | 5.62 |
| Q15018 | 16.63 | 1 | 6 | 6 | 10 | 2.65 | 1 | 1 |  |  |  | 2.65 | 1 | 1 | 2.65 | 1 | 1 |  |  |  |  |  |  |  |  |  |  |  |  |  |  |  | 16.63 | 6 | 7 | 415 | 46.9 | 6.21 |
| Q15019 | 41.83 | 1 | 10 | 10 | 19 |  |  |  |  |  |  |  |  |  |  |  |  |  |  |  |  |  |  | 3.88 | 1 | 1 |  |  |  | 5.82 | 2 | 2 | 39.89 | 9 | 16 | 361 | 41.5 | 6.60 |
| P01877 | 23.53 | 2 | 1 | 6 | 14 | 2.94 | 1 | 1 | 5.00 | 2 | 2 | 23.53 | 6 | 7 |  |  |  | 2.06 | 1 | 1 |  |  |  |  |  |  |  |  |  |  |  |  | 7.65 | 3 | 3 | 340 | 36.6 | 6.27 |
| Q15041 | 16.75 | 1 | 4 | 4 | 4 |  |  |  |  |  |  |  |  |  |  |  |  |  |  |  |  |  |  |  |  |  |  |  |  |  |  |  | 16.75 | 4 | 4 | 203 | 23.3 | 9.32 |
| Q15046 | 45.73 | 1 | 25 | 25 | 92 | 19.93 | 10 | 10 | 18.26 | 9 | 9 | 13.07 | 6 | 7 | 7.37 | 4 | 4 | 11.06 | 5 | 5 |  |  |  |  |  |  |  |  |  | 1.51 | 1 | 1 | 43.05 | 22 | 56 | 597 | 68.0 | 6.35 |
| Q15056 | 25.81 | 1 | 7 | 7 | 11 |  |  |  |  |  |  |  |  |  |  |  |  |  |  |  |  |  |  | 5.24 | 1 | 1 | 16.53 | 4 | 4 | 14.11 | 3 | 4 | 6.45 | 2 | 2 | 248 | 27.4 | 7.23 |
| Q15067 | 5.76 | 1 | 2 | 2 | 4 |  |  |  |  |  |  |  |  |  |  |  |  |  |  |  |  |  |  |  |  |  |  |  |  |  |  |  | 5.76 | 2 | 4 | 660 | 74.4 | 8.16 |
| Q15075 | 20.62 | 1 | 23 | 23 | 34 | 0.00 |  |  | 0.00 |  |  | 0.78 | 1 | 1 | 0.00 |  |  | 0.00 |  |  |  |  |  | 1.13 | 1 | 1 | 1.84 | 2 | 3 | 1.13 | 1 | 1 | 19.84 | 22 | 28 | 1411 | 162.4 | 5.68 |
| Q15084 | 48.41 | 1 | 19 | 19 | 85 | 25.45 | 8 | 12 | 29.32 | 9 | 12 | 23.64 | 8 | 9 | 8.64 | 3 | 3 | 10.68 | 4 | 4 |  |  |  |  |  |  |  |  |  |  |  |  | 44.32 | 15 | 45 | 440 | 48.1 | 5.08 |
| Q15102 | 41.99 | 1 | 9 | 9 | 21 | 14.29 | 2 | 2 | 3.90 | 1 | 1 |  |  |  | 7.79 | 2 | 2 |  |  |  |  |  |  | 16.88 | 3 | 3 |  |  |  | 13.85 | 3 | 3 | 28.57 | 6 | 10 | 231 | 25.7 | 6.84 |
| Q15120 | 16.50 | 1 | 5 | 5 | 5 |  |  |  | 2.46 | 1 | 1 |  |  |  |  |  |  |  |  |  |  |  |  |  |  |  |  |  |  |  |  |  | 14.04 | 4 | 4 | 406 | 46.9 | 8.37 |
| Q15121 | 63.08 | 1 | 8 | 8 | 20 | 10.77 | 1 | 1 |  |  |  |  |  |  | 7.69 | 1 | 1 |  |  |  |  |  |  | 26.92 | 3 | 4 |  |  |  | 52.31 | 5 | 7 | 43.85 | 5 | 7 | 130 | 15.0 | 5.02 |
| Q15125 | 12.17 | 1 | 2 | 2 | 13 | 5.22 | 1 | 1 | 5.22 | 1 | 1 | 5.22 | 1 | 1 | 5.22 | 1 | 1 |  |  |  |  |  |  |  |  |  |  |  |  |  |  |  | 12.17 | 2 | 9 | 230 | 26.3 | 7.90 |
| Q15126 | 61.46 | 1 | 14 | 14 | 26 |  |  |  |  |  |  |  |  |  | 5.73 | 1 | 1 |  |  |  |  |  |  | 5.73 | 1 | 1 |  |  |  | 18.75 | 3 | 3 | 60.94 | 13 | 21 | 192 | 22.0 | 5.73 |
| Q15139 | 4.71 | 8 | 4 | 4 | 7 |  |  |  |  |  |  |  |  |  |  |  |  |  |  |  |  |  |  |  |  |  |  |  |  |  |  |  | 4.71 | 4 | 7 | 912 | 101.6 | 6.62 |
| Q15149 | 5.36 | 5 | 23 | 24 | 28 | 0.62 | 3 | 3 |  |  |  | 0.47 | 2 | 2 |  |  |  |  |  |  |  |  |  |  |  |  |  |  |  |  |  |  | 4.53 | 20 | 23 | 4684 | 531.5 | 5.96 |
| Q15154 | 3.90 | 1 | 6 | 6 | 6 |  |  |  | 0.49 | 1 | 1 |  |  |  |  |  |  |  |  |  |  |  |  |  |  |  |  |  |  |  |  |  | 3.41 | 5 | 5 | 2024 | 228.4 | 5.02 |
| Q15172 | 25.93 | 1 | 9 | 11 | 20 |  |  |  |  |  |  |  |  |  |  |  |  |  |  |  |  |  |  |  |  |  |  |  |  |  |  |  | 25.93 | 11 | 20 | 486 | 56.2 | 6.71 |
| Q15181 | 70.59 | 1 | 18 | 19 | 45 | 16.26 | 4 | 4 |  |  |  |  |  |  | 15.57 | 4 | 5 | 12.80 | 2 | 2 |  |  |  |  |  |  | 6.57 | 2 | 2 | 37.02 | 9 | 13 | 53.63 | 12 | 19 | 289 | 32.6 | 5.86 |
| Q15185 | 56.88 | 1 | 10 | 10 | 52 | 13.13 | 2 | 2 | 8.13 | 1 | 1 | 23.75 | 3 | 3 | 29.38 | 4 | 4 | 35.63 | 6 | 7 |  |  |  | 5.63 | 1 | 1 | 31.25 | 6 | 6 | 35.00 | 6 | 9 | 52.50 | 8 | 19 | 160 | 18.7 | 4.54 |
| Q15233 | 7.86 | 1 | 3 | 3 | 4 | 2.97 | 1 | 1 |  |  |  |  |  |  |  |  |  |  |  |  |  |  |  |  |  |  |  |  |  | 2.55 | 1 | 1 | 2.34 | 1 | 2 | 471 | 54.2 | 8.95 |
| Q15257 | 53.63 | 1 | 16 | 16 | 39 | 18.16 | 4 | 6 | 4.75 | 2 | 2 | 2.79 | 1 | 1 | 5.87 | 2 | 2 |  |  |  |  |  |  |  |  |  | 3.35 | 1 | 1 |  |  |  | 45.25 | 14 | 23 | 358 | 40.6 | 5.94 |
| Q15276 | 3.25 | 1 | 2 | 2 | 3 |  |  |  |  |  |  |  |  |  |  |  |  |  |  |  |  |  |  |  |  |  | 1.04 | 1 | 1 |  |  |  | 2.20 | 1 | 2 | 862 | 99.2 | 5.01 |
| Q15286 | 35.82 | 7 | 5 | 7 | 39 | 10.95 | 2 | 3 | 10.95 | 2 | 2 | 15.42 | 3 | 3 | 5.47 | 1 | 1 |  |  |  |  |  |  | 5.47 | 1 | 1 | 5.47 | 1 | 1 | 5.47 | 1 | 1 | 35.82 | 7 | 23 | 201 | 23.0 | 8.29 |
| Q15293 | 23.26 | 1 | 7 | 7 | 20 | 4.83 | 2 | 2 | 5.74 | 1 | 1 | 2.72 | 1 | 1 |  |  |  | 2.72 | 1 | 1 |  |  |  | 9.06 | 2 | 2 | 14.80 | 4 | 4 | 16.62 | 4 | 7 | 3.63 | 1 | 2 | 331 | 38.9 | 5.00 |
| Q15311 | 4.73 | 1 | 2 | 2 | 2 |  |  |  |  |  |  |  |  |  |  |  |  |  |  |  |  |  |  |  |  |  |  |  |  |  |  |  | 4.73 | 2 | 2 | 655 | 76.0 | 5.88 |
| Q15363 | 36.82 | 1 | 7 | 7 | 39 | 18.41 | 2 | 2 | 27.86 | 4 | 6 | 25.87 | 5 | 7 |  |  |  |  |  |  |  |  |  |  |  |  |  |  |  |  |  |  | 27.36 | 4 | 20 | 201 | 22.7 | 5.17 |
| Q15365 | 62.36 | 1 | 11 | 16 | 114 | 15.17 | 4 | 4 | 5.34 | 2 | 2 | 11.80 | 4 | 4 | 18.26 | 5 | 5 | 9.55 | 3 | 3 |  |  |  | 6.74 | 2 | 2 | 26.40 | 7 | 7 | 36.52 | 9 | 9 | 59.27 | 12 | 78 | 356 | 37.5 | 7.09 |
| Q15366 | 52.60 | 2 | 8 | 13 | 60 | 6.85 | 2 | 2 | 8.22 | 3 | 3 | 5.21 | 2 | 2 | 9.59 | 3 | 3 | 3.84 | 1 | 1 |  |  |  |  |  |  | 5.21 | 2 | 2 | 13.15 | 3 | 3 | 49.59 | 10 | 44 | 365 | 38.6 | 6.79 |
| Q15369 | 66.07 | 1 | 8 | 8 | 55 | 50.89 | 4 | 5 | 50.89 | 4 | 4 | 23.21 | 2 | 2 | 66.07 | 6 | 7 | 33.04 | 3 | 4 |  |  |  | 58.93 | 5 | 7 | 33.04 | 3 | 5 | 58.93 | 4 | 7 | 66.07 | 7 | 14 | 112 | 12.5 | 4.78 |
| Q15370 | 97.46 | 1 | 11 | 11 | 54 | 45.76 | 4 | 4 | 45.76 | 4 | 4 | 6.78 | 1 | 1 | 45.76 | 4 | 5 | 38.14 | 3 | 4 |  |  |  | 31.36 | 4 | 5 | 52.54 | 5 | 6 | 51.69 | 5 | 6 | 91.53 | 10 | 19 | 118 | 13.1 | 4.88 |
| Q15382 | 28.80 | 1 | 5 | 5 | 9 |  |  |  |  |  |  |  |  |  |  |  |  |  |  |  |  |  |  |  |  |  |  |  |  |  |  |  | 28.80 | 5 | 9 | 184 | 20.5 | 5.92 |
| Q15386 | 8.96 | 1 | 8 | 8 | 14 |  |  |  |  |  |  |  |  |  |  |  |  |  |  |  |  |  |  |  |  |  |  |  |  |  |  |  | 8.96 | 8 | 14 | 1083 | 123.8 | 6.71 |
| Q15393 | 3.62 | 1 | 3 | 3 | 3 |  |  |  |  |  |  |  |  |  |  |  |  |  |  |  |  |  |  |  |  |  |  |  |  |  |  |  | 3.62 | 3 | 3 | 1217 | 135.5 | 5.26 |
| Q15418 | 20.68 | 9 | 6 | 13 | 40 |  |  |  |  |  |  |  |  |  |  |  |  |  |  |  |  |  |  |  |  |  |  |  |  | 3.67 | 2 | 2 | 18.50 | 12 | 38 | 735 | 82.7 | 7.83 |
| Q15435 | 66.11 | 1 | 22 | 22 | 113 | 44.44 | 13 | 15 | 33.33 | 11 | 15 | 31.39 | 9 | 9 | 15.56 | 6 | 6 |  |  |  |  |  |  | 20.83 | 7 | 7 | 16.94 | 7 | 7 | 25.00 | 8 | 8 | 55.00 | 17 | 46 | 360 | 41.5 | 4.91 |
| Q15437 | 17.08 | 1 | 8 | 9 | 12 |  |  |  |  |  |  |  |  |  |  |  |  |  |  |  |  |  |  |  |  |  |  |  |  |  |  |  | 17.08 | 9 | 12 | 767 | 86.4 | 6.89 |
| Q15477 | 12.04 | 1 | 11 | 11 | 14 |  |  |  |  |  |  |  |  |  |  |  |  |  |  |  |  |  |  |  |  |  |  |  |  |  |  |  | 12.04 | 11 | 14 | 1246 | 137.7 | 6.06 |
| Q15506 | 92.72 | 1 | 23 | 23 | 135 | 57.62 | 6 | 8 | 64.90 | 8 | 11 | 49.01 | 5 | 5 | 76.82 | 10 | 13 | 91.39 | 17 | 24 |  |  |  | 44.37 | 5 | 7 | 39.07 | 8 | 11 | 41.72 | 6 | 7 | 87.42 | 14 | 49 | 151 | 17.4 | 4.78 |
| Q15628 | 17.31 | 1 | 4 | 4 | 7 |  |  |  |  |  |  |  |  |  |  |  |  |  |  |  |  |  |  |  |  |  |  |  |  |  |  |  | 17.31 | 4 | 7 | 312 | 34.2 | 6.27 |
| Q15631 | 47.81 | 1 | 11 | 11 | 52 | 41.67 | 6 | 10 | 41.67 | 6 | 9 | 31.14 | 4 | 5 | 18.42 | 3 | 4 |  |  |  |  |  |  | 4.39 | 1 | 1 |  |  |  | 11.40 | 2 | 2 | 33.33 | 9 | 21 | 228 | 26.2 | 6.44 |
| Q15645 | 41.20 | 1 | 17 | 17 | 21 |  |  |  |  |  |  |  |  |  |  |  |  |  |  |  |  |  |  |  |  |  | 3.01 | 1 | 1 |  |  |  | 41.20 | 17 | 20 | 432 | 48.5 | 6.09 |
| Q15691 | 23.13 | 1 | 4 | 4 | 4 |  |  |  |  |  |  |  |  |  |  |  |  |  |  |  |  |  |  |  |  |  |  |  |  |  |  |  | 23.13 | 4 | 4 | 268 | 30.0 | 5.14 |
| Q15738 | 13.94 | 1 | 4 | 4 | 9 |  |  |  |  |  |  | 5.09 | 1 | 1 |  |  |  |  |  |  |  |  |  |  |  |  |  |  |  |  |  |  | 13.94 | 4 | 8 | 373 | 41.9 | 8.06 |
| Q15751 | 0.58 | 1 | 2 | 2 | 2 |  |  |  |  |  |  |  |  |  |  |  |  |  |  |  |  |  |  |  |  |  |  |  |  |  |  |  | 0.58 | 2 | 2 | 4861 | 531.9 | 6.04 |
| Q15785 | 63.43 | 1 | 21 | 21 | 87 | 23.30 | 5 | 5 | 34.30 | 8 | 11 | 10.03 | 2 | 2 | 25.24 | 6 | 6 | 46.60 | 13 | 14 |  |  |  | 2.91 | 1 | 1 | 7.44 | 2 | 2 | 33.66 | 10 | 10 | 60.52 | 17 | 36 | 309 | 34.5 | 8.98 |
| Q15813 | 22.39 | 2 | 8 | 8 | 10 |  |  |  | 5.50 | 2 | 2 |  |  |  |  |  |  | 2.09 | 1 | 1 |  |  |  |  |  |  |  |  |  |  |  |  | 17.08 | 6 | 7 | 527 | 59.3 | 6.76 |
| Q15818 | 8.10 | 3 | 3 | 3 | 4 |  |  |  |  |  |  |  |  |  |  |  |  |  |  |  |  |  |  |  |  |  |  |  |  |  |  |  | 8.10 | 3 | 4 | 432 | 47.1 | 6.55 |
| P06241 | 8.38 | 19 | 1 | 5 | 13 | 0.00 |  |  | 0.00 |  |  | 1.49 | 1 | 1 |  |  |  |  |  |  |  |  |  |  |  |  |  |  |  | 1.86 | 1 | 1 | 8.38 | 5 | 11 | 537 | 60.7 | 6.67 |
| P05062 | 3.85 | 2 | 1 | 2 | 13 | 1.92 | 1 | 1 | 0.00 |  |  | 1.92 | 1 | 1 |  |  |  | 1.92 | 1 | 1 |  |  |  | 1.92 | 1 | 1 | 1.92 | 1 | 1 | 1.92 | 1 | 1 | 1.92 | 1 | 7 | 364 | 39.4 | 7.87 |
| Q15831 | 11.78 | 1 | 4 | 4 | 8 |  |  |  |  |  |  |  |  |  |  |  |  |  |  |  |  |  |  |  |  |  |  |  |  |  |  |  | 11.78 | 4 | 8 | 433 | 48.6 | 7.43 |
| Q15833 | 44.35 | 1 | 21 | 22 | 53 | 7.76 | 3 | 4 | 9.44 | 4 | 5 | 1.85 | 1 | 1 |  |  |  | 0.00 |  |  |  |  |  | 2.02 | 1 | 1 | 0.00 |  |  | 4.72 | 2 | 2 | 40.81 | 20 | 40 | 593 | 66.4 | 6.55 |
| Q15836 | 53.00 | 2 | 3 | 6 | 39 | 41.00 | 4 | 7 | 41.00 | 4 | 8 | 24.00 | 2 | 2 | 24.00 | 1 | 1 |  |  |  |  |  |  |  |  |  |  |  |  |  |  |  | 53.00 | 6 | 21 | 100 | 11.3 | 8.79 |
| Q15843 | 34.57 | 1 | 2 | 2 | 3 |  |  |  | 17.28 | 1 | 2 |  |  |  | 17.28 | 1 | 1 |  |  |  |  |  |  |  |  |  |  |  |  |  |  |  |  |  |  | 81 | 9.1 | 8.43 |
| Q15904 | 7.23 | 1 | 4 | 4 | 5 |  |  |  |  |  |  |  |  |  |  |  |  |  |  |  |  |  |  |  |  |  |  |  |  |  |  |  | 7.23 | 4 | 5 | 470 | 52.0 | 6.14 |
| Q15907 | 64.22 | 1 | 2 | 15 | 78 | 42.20 | 8 | 10 | 42.20 | 7 | 8 | 24.31 | 6 | 6 | 29.36 | 6 | 6 | 5.05 | 1 | 1 |  |  |  | 9.17 | 2 | 2 | 5.05 | 1 | 1 | 24.77 | 5 | 6 | 61.47 | 13 | 38 | 218 | 24.5 | 5.94 |
| Q16134 | 20.75 | 1 | 9 | 9 | 13 |  |  |  | 1.94 | 1 | 1 |  |  |  |  |  |  |  |  |  |  |  |  |  |  |  |  |  |  |  |  |  | 18.80 | 8 | 12 | 617 | 68.5 | 7.55 |
| Q16181 | 35.24 | 2 | 15 | 16 | 68 | 5.26 | 3 | 3 | 5.95 | 2 | 2 | 2.06 | 1 | 1 | 2.06 | 1 | 1 | 21.51 | 9 | 11 |  |  |  |  |  |  | 2.29 | 1 | 1 |  |  |  | 34.78 | 13 | 47 | 437 | 50.6 | 8.63 |
| Q16186 | 17.94 | 1 | 6 | 6 | 21 |  |  |  | 3.93 | 1 | 1 | 3.93 | 1 | 1 | 3.93 | 1 | 1 | 3.93 | 1 | 1 |  |  |  | 3.93 | 1 | 1 | 3.93 | 1 | 1 | 3.93 | 1 | 1 | 17.94 | 6 | 14 | 407 | 42.1 | 5.07 |
| Q16222 | 14.94 | 1 | 5 | 6 | 10 | 1.92 | 1 | 1 |  |  |  | 1.92 | 1 | 1 |  |  |  |  |  |  |  |  |  |  |  |  |  |  |  | 2.49 | 1 | 1 | 12.45 | 5 | 7 | 522 | 58.7 | 6.33 |
| Q16348 | 7.13 | 1 | 5 | 5 | 10 |  |  |  | 1.10 | 1 | 1 | 2.19 | 2 | 2 |  |  |  |  |  |  |  |  |  |  |  |  |  |  |  |  |  |  | 6.04 | 4 | 7 | 729 | 81.7 | 8.16 |
| Q16401 | 41.07 | 1 | 16 | 16 | 37 | 8.93 | 3 | 3 | 4.37 | 2 | 2 | 1.79 | 1 | 1 | 4.37 | 2 | 2 |  |  |  |  |  |  |  |  |  |  |  |  |  |  |  | 35.12 | 14 | 29 | 504 | 56.2 | 5.48 |
| Q16531 | 32.19 | 1 | 31 | 31 | 57 |  |  |  | 2.63 | 2 | 2 | 15.09 | 15 | 15 |  |  |  |  |  |  |  |  |  |  |  |  |  |  |  |  |  |  | 24.65 | 23 | 40 | 1140 | 126.9 | 5.26 |
| Q16537 | 7.71 | 1 | 2 | 4 | 6 |  |  |  |  |  |  |  |  |  |  |  |  |  |  |  |  |  |  |  |  |  |  |  |  |  |  |  | 7.71 | 4 | 6 | 467 | 54.7 | 6.95 |
| Q16543 | 25.66 | 1 | 10 | 10 | 28 | 6.61 | 2 | 2 | 5.82 | 2 | 2 | 6.08 | 2 | 2 | 3.17 | 1 | 1 |  |  |  |  |  |  | 8.20 | 4 | 4 |  |  |  | 8.47 | 3 | 3 | 19.31 | 7 | 14 | 378 | 44.4 | 5.25 |
| Q16563 | 38.22 | 1 | 9 | 9 | 58 | 15.83 | 4 | 4 | 14.67 | 4 | 4 | 5.79 | 2 | 2 | 7.34 | 2 | 2 | 3.09 | 1 | 1 |  |  |  |  |  |  |  |  |  |  |  |  | 38.22 | 9 | 45 | 259 | 28.5 | 8.43 |
| Q16566 | 28.12 | 2 | 12 | 12 | 19 | 5.50 | 2 | 3 |  |  |  | 2.33 | 1 | 1 |  |  |  |  |  |  |  |  |  |  |  |  |  |  |  |  |  |  | 23.47 | 10 | 15 | 473 | 51.9 | 5.82 |
| Q16576 | 8.94 | 2 | 3 | 3 | 3 |  |  |  |  |  |  |  |  |  |  |  |  |  |  |  |  |  |  |  |  |  | 8.94 | 3 | 3 |  |  |  |  |  |  | 425 | 47.8 | 5.05 |
| Q16610 | 48.52 | 2 | 20 | 20 | 76 | 35.00 | 14 | 16 | 20.00 | 10 | 11 | 10.56 | 5 | 5 | 13.52 | 6 | 6 | 16.11 | 9 | 10 |  |  |  | 2.41 | 1 | 1 |  |  |  |  |  |  | 27.04 | 10 | 27 | 540 | 60.6 | 6.71 |
| Q16651 | 20.41 | 1 | 4 | 4 | 39 | 18.37 | 3 | 6 | 18.37 | 3 | 5 | 18.37 | 3 | 5 | 4.08 | 1 | 2 | 4.08 | 1 | 2 |  |  |  |  |  |  | 4.08 | 1 | 2 | 4.08 | 1 | 2 | 11.66 | 3 | 11 | 343 | 36.4 | 5.85 |
| Q16698 | 66.87 | 1 | 24 | 24 | 119 | 46.57 | 13 | 18 | 50.15 | 14 | 20 | 39.40 | 11 | 14 | 17.01 | 6 | 6 | 22.39 | 7 | 7 |  |  |  | 3.88 | 1 | 1 | 3.88 | 1 | 1 | 3.88 | 1 | 1 | 60.90 | 19 | 51 | 335 | 36.0 | 9.28 |
| Q16706 | 2.88 | 1 | 2 | 3 | 3 |  |  |  |  |  |  |  |  |  |  |  |  |  |  |  |  |  |  |  |  |  |  |  |  |  |  |  | 2.88 | 3 | 3 | 1144 | 131.1 | 7.58 |
| Q16718 | 66.38 | 1 | 7 | 7 | 17 | 31.03 | 2 | 2 | 38.79 | 3 | 3 |  |  |  | 7.76 | 1 | 1 |  |  |  |  |  |  |  |  |  |  |  |  | 7.76 | 1 | 1 | 66.38 | 7 | 10 | 116 | 13.5 | 5.99 |
| Q16774 | 19.29 | 1 | 4 | 4 | 7 |  |  |  |  |  |  |  |  |  |  |  |  |  |  |  |  |  |  | 5.58 | 1 | 1 |  |  |  | 5.08 | 1 | 1 | 19.29 | 3 | 5 | 197 | 21.7 | 6.55 |
| Q16775 | 45.78 | 1 | 11 | 11 | 45 | 37.01 | 8 | 12 | 34.09 | 7 | 11 | 11.69 | 4 | 6 | 9.09 | 2 | 2 | 12.99 | 3 | 3 |  |  |  |  |  |  |  |  |  | 3.90 | 1 | 1 | 31.82 | 7 | 10 | 308 | 33.8 | 8.12 |
| Q16795 | 41.11 | 1 | 13 | 13 | 23 | 5.84 | 2 | 2 | 11.14 | 4 | 4 |  |  |  |  |  |  |  |  |  |  |  |  |  |  |  |  |  |  |  |  |  | 38.46 | 12 | 17 | 377 | 42.5 | 9.80 |
| Q16822 | 4.06 | 1 | 2 | 2 | 2 | 1.88 | 1 | 1 |  |  |  |  |  |  |  |  |  |  |  |  |  |  |  |  |  |  |  |  |  |  |  |  | 2.19 | 1 | 1 | 640 | 70.7 | 7.62 |
| Q16836 | 53.18 | 2 | 13 | 13 | 41 | 37.58 | 6 | 8 | 38.85 | 7 | 11 | 19.43 | 5 | 6 |  |  |  | 2.55 | 1 | 1 |  |  |  |  |  |  |  |  |  | 2.55 | 1 | 1 | 30.57 | 9 | 14 | 314 | 34.3 | 8.85 |
| Q93070 | 2.23 | 1 | 1 | 1 | 13 |  |  |  |  |  |  |  |  |  | 2.23 | 1 | 1 | 2.23 | 1 | 3 |  |  |  | 2.23 | 1 | 3 | 2.23 | 1 | 3 | 2.23 | 1 | 3 |  |  |  | 314 | 35.9 | 9.22 |
| Q16851 | 48.43 | 1 | 21 | 21 | 55 | 12.99 | 4 | 4 | 25.79 | 10 | 12 | 25.39 | 10 | 11 |  |  |  | 2.76 | 1 | 1 |  |  |  | 2.76 | 1 | 1 | 2.76 | 1 | 1 | 2.76 | 1 | 1 | 34.45 | 14 | 24 | 508 | 56.9 | 8.15 |
| Q16854 | 23.47 | 1 | 5 | 5 | 15 | 16.25 | 3 | 3 | 16.97 | 3 | 5 | 11.55 | 2 | 2 |  |  |  |  |  |  |  |  |  |  |  |  |  |  |  |  |  |  | 19.49 | 4 | 5 | 277 | 32.0 | 8.66 |
| Q16864 | 28.57 | 1 | 2 | 2 | 2 | 28.57 | 2 | 2 |  |  |  |  |  |  |  |  |  |  |  |  |  |  |  |  |  |  |  |  |  |  |  |  |  |  |  | 119 | 13.4 | 5.52 |
| Q16881 | 27.58 | 1 | 9 | 10 | 19 | 6.32 | 3 | 3 | 1.69 | 1 | 1 | 3.70 | 2 | 2 |  |  |  |  |  |  |  |  |  |  |  |  |  |  |  |  |  |  | 23.88 | 8 | 13 | 649 | 70.9 | 7.39 |
| Q16891 | 68.87 | 1 | 46 | 46 | 135 | 6.99 | 4 | 4 | 15.70 | 10 | 10 | 10.29 | 7 | 7 |  |  |  | 1.45 | 1 | 1 |  |  |  |  |  |  |  |  |  |  |  |  | 66.23 | 43 | 113 | 758 | 83.6 | 6.48 |
| Q17RY6 | 24.85 | 1 | 4 | 4 | 13 | 18.79 | 3 | 3 | 11.52 | 2 | 2 |  |  |  |  |  |  | 10.91 | 2 | 2 |  |  |  |  |  |  |  |  |  |  |  |  | 24.85 | 4 | 6 | 165 | 18.7 | 7.43 |
| Q1ZYL8 | 50.00 | 1 | 12 | 12 | 135 | 38.79 | 7 | 12 | 49.57 | 10 | 25 | 38.36 | 7 | 9 | 14.22 | 2 | 4 | 16.81 | 4 | 4 |  |  |  |  |  |  |  |  |  |  |  |  | 43.10 | 10 | 81 | 232 | 26.5 | 7.46 |
| Q2KHT3 | 3.61 | 1 | 3 | 3 | 3 | 1.14 | 1 | 1 |  |  |  |  |  |  |  |  |  |  |  |  |  |  |  |  |  |  |  |  |  |  |  |  | 2.47 | 2 | 2 | 1053 | 117.6 | 5.86 |
| Q2M243 | 29.42 | 1 | 19 | 19 | 63 | 2.59 | 1 | 1 | 7.62 | 4 | 4 | 1.52 | 1 | 1 | 1.52 | 1 | 1 | 5.79 | 3 | 3 |  |  |  |  |  |  | 2.59 | 1 | 1 | 1.52 | 1 | 1 | 27.90 | 18 | 51 | 656 | 75.3 | 5.68 |
| Q2M2I8 | 9.68 | 2 | 5 | 5 | 9 |  |  |  |  |  |  |  |  |  |  |  |  |  |  |  |  |  |  |  |  |  |  |  |  |  |  |  | 9.68 | 5 | 9 | 961 | 103.8 | 6.60 |
| Q2M329 | 38.38 | 1 | 25 | 25 | 38 |  |  |  | 3.24 | 1 | 1 |  |  |  | 5.05 | 2 | 2 | 28.47 | 12 | 13 |  |  |  |  |  |  |  |  |  |  |  |  | 24.14 | 13 | 22 | 555 | 62.7 | 4.94 |
| Q2M389 | 11.00 | 3 | 10 | 10 | 19 |  |  |  |  |  |  |  |  |  |  |  |  |  |  |  |  |  |  |  |  |  |  |  |  |  |  |  | 11.00 | 10 | 19 | 1173 | 136.3 | 7.44 |
| Q2PPJ7 | 2.24 | 1 | 3 | 4 | 8 | 0.00 |  |  | 0.00 |  |  |  |  |  |  |  |  |  |  |  |  |  |  |  |  |  |  |  |  |  |  |  | 2.24 | 4 | 8 | 1873 | 210.6 | 6.07 |
| Q2TAA2 | 14.52 | 1 | 2 | 2 | 2 |  |  |  |  |  |  |  |  |  |  |  |  |  |  |  |  |  |  |  |  |  |  |  |  |  |  |  | 14.52 | 2 | 2 | 248 | 27.6 | 5.30 |
| Q2TAA8 | 22.04 | 11 | 13 | 13 | 17 |  |  |  |  |  |  |  |  |  |  |  |  |  |  |  |  |  |  |  |  |  |  |  |  |  |  |  | 22.04 | 13 | 17 | 658 | 76.7 | 5.06 |
| Q32MB4 | 33.20 | 2 | 17 | 17 | 40 | 1.38 | 1 | 1 |  |  |  |  |  |  |  |  |  |  |  |  |  |  |  |  |  |  |  |  |  |  |  |  | 33.20 | 17 | 39 | 506 | 58.4 | 9.20 |
| Q32MZ4 | 10.27 | 1 | 5 | 6 | 10 | 1.73 | 1 | 1 |  |  |  | 4.95 | 3 | 3 |  |  |  |  |  |  |  |  |  | 1.49 | 1 | 1 | 1.49 | 1 | 1 |  |  |  | 7.05 | 4 | 4 | 808 | 89.2 | 4.65 |
| Q330K2 | 7.81 | 1 | 2 | 2 | 2 |  |  |  |  |  |  |  |  |  |  |  |  |  |  |  |  |  |  |  |  |  |  |  |  |  |  |  | 7.81 | 2 | 2 | 333 | 38.2 | 9.36 |
| Q3KQV9 | 10.85 | 1 | 4 | 5 | 9 |  |  |  |  |  |  |  |  |  |  |  |  |  |  |  |  |  |  | 1.78 | 1 | 1 |  |  |  | 2.56 | 1 | 1 | 8.28 | 4 | 7 | 507 | 57.0 | 6.32 |
| Q3KQZ1 | 36.67 | 1 | 7 | 7 | 11 |  |  |  | 3.00 | 1 | 1 |  |  |  |  |  |  |  |  |  |  |  |  |  |  |  |  |  |  |  |  |  | 36.67 | 7 | 10 | 300 | 32.4 | 9.11 |
| Q3LXA3 | 52.70 | 1 | 22 | 22 | 69 | 24.17 | 9 | 10 | 23.65 | 9 | 15 | 8.70 | 3 | 3 | 5.04 | 2 | 2 | 3.65 | 1 | 1 |  |  |  |  |  |  |  |  |  |  |  |  | 45.91 | 18 | 38 | 575 | 58.9 | 7.49 |
| Q3MIX3 | 6.72 | 1 | 3 | 3 | 3 |  |  |  | 3.28 | 1 | 1 |  |  |  |  |  |  |  |  |  |  |  |  |  |  |  |  |  |  |  |  |  | 3.45 | 2 | 2 | 580 | 65.9 | 8.97 |

| | | | | | | | | | | | | | | | | | | | | | | | | | | | | | | | | | | | | | | |
|---|---|---|---|---|---|---|---|---|---|---|---|---|---|---|---|---|---|---|---|---|---|---|---|---|---|---|---|---|---|---|---|---|---|---|---|---|---|---|
| Q3SXM5 | 6.36 | 1 | 2 | 2 | 2 | | | | | | | | | | | | | | | | | | | | | | | | | | | | 6.36 | 2 | 2 | 330 | 37.0 | 8.72 |
| Q3SYG4 | 4.06 | 1 | 2 | 2 | 3 | | | | | | | | | | | | | | | | | | | | | | | | | | | | 4.06 | 2 | 3 | 887 | 99.2 | 5.55 |
| Q3ZCQ8 | 18.70 | 1 | 5 | 5 | 11 | | | | 4.82 | 1 | 1 | | | | | | | | | | | | | | | | | | | | | | 18.70 | 5 | 10 | 353 | 39.6 | 8.37 |
| Q3ZCV2 | 26.79 | 1 | 10 | 10 | 11 | | | | | | | | | | | | | | | | | | | | | | | | | | | | 26.79 | 10 | 11 | 418 | 47.6 | 9.82 |
| Q3ZCW2 | 23.84 | 1 | 4 | 4 | 6 | 6.40 | 1 | 1 | | | | | | | | | | | | | | | | | | | | | | | | | 23.84 | 4 | 5 | 172 | 19.0 | 5.35 |
| Q494V2 | 24.55 | 2 | 17 | 17 | 41 | | | | | | | | | | 5.89 | 4 | 5 | 9.33 | 6 | 6 | | | | | | | | | | | | | 22.26 | 15 | 30 | 611 | 71.1 | 7.11 |
| Q499Z3 | 25.31 | 1 | 9 | 9 | 24 | 3.93 | 2 | 2 | 5.65 | 2 | 2 | | | | | | | | | | | | | 2.95 | 1 | 1 | | | | | | | 23.59 | 8 | 19 | 407 | 45.6 | 5.90 |
| Q4G0F5 | 17.26 | 1 | 4 | 5 | 7 | | | | | | | | | | | | | | | | | | | | | | | | | | | | 17.26 | 5 | 7 | 336 | 39.1 | 7.36 |
| Q4G0N4 | 14.93 | 1 | 5 | 5 | 6 | | | | 3.17 | 1 | 1 | 2.26 | 1 | 1 | | | | | | | | | | | | | | | | | | | 12.67 | 4 | 4 | 442 | 49.4 | 8.18 |
| Q4G0P3 | 20.25 | 1 | 87 | 87 | 166 | 0.21 | 1 | 1 | 0.41 | 2 | 2 | 0.20 | 1 | 1 | | | | | | | | | | | | | | | | | | | 20.05 | 86 | 162 | 5121 | 575.5 | 6.06 |
| Q4G0U5 | 10.83 | 1 | 7 | 7 | 10 | | | | | | | | | | | | | 1.19 | 1 | 1 | | | | | | | | | | | | | 10.83 | 7 | 9 | 840 | 96.8 | 8.62 |
| Q4G0X9 | 50.96 | 2 | 68 | 68 | 179 | 2.10 | 2 | 2 | 2.45 | 2 | 2 | | | | 12.00 | 12 | 14 | 31.35 | 31 | 37 | | | | | | | | | | | | | 47.81 | 57 | 124 | 1142 | 130.0 | 5.29 |
| Q4G1C9 | 14.24 | 1 | 4 | 4 | 6 | 2.91 | 1 | 1 | 2.91 | 1 | 1 | | | | | | | | | | | | | | | | | | | | | | 11.34 | 3 | 4 | 344 | 40.2 | 4.91 |
| Q4KMQ2 | 2.20 | 1 | 2 | 2 | 3 | | | | | | | | | | | | | | | | | | | | | | | | | | | | 2.20 | 2 | 3 | 910 | 106.1 | 7.77 |
| Q4LDG9 | 52.63 | 1 | 10 | 10 | 24 | 29.47 | 4 | 4 | 24.21 | 4 | 4 | | | | | | | | | | | | | | | | | | | | | | 52.63 | 10 | 16 | 190 | 21.5 | 6.10 |
| Q4VNC1 | 2.93 | 1 | 3 | 3 | 4 | | | | | | | | | | | | | | | | | | | | | | | | | | | | 2.93 | 3 | 4 | 1196 | 133.9 | 6.70 |
| Q502W7 | 11.55 | 1 | 5 | 6 | 7 | | | | | | | | | | | | | | | | | | | | | | | | | | | | 11.55 | 6 | 7 | 563 | 65.3 | 8.75 |
| Q537H7 | 48.98 | 1 | 5 | 5 | 10 | | | | | | | | | | | | | 8.16 | 1 | 1 | | | | | | | | | | | | | 48.98 | 5 | 9 | 98 | 11.3 | 10.07 |
| Q53EL6 | 10.66 | 1 | 4 | 4 | 5 | | | | | | | | | | | | | | | | | | | | | | | | | | | | 10.66 | 4 | 5 | 469 | 51.7 | 5.21 |
| Q53EV4 | 75.22 | 1 | 27 | 27 | 66 | 7.29 | 1 | 1 | 12.54 | 3 | 3 | | | | 35.86 | 10 | 11 | 38.78 | 13 | 16 | | | | | | | | | | | | | 64.14 | 20 | 35 | 343 | 39.7 | 4.65 |
| Q53GD3 | 7.18 | 1 | 6 | 6 | 28 | 3.94 | 2 | 2 | 1.97 | 1 | 1 | 3.94 | 2 | 3 | | | | | | | | | | | | | | | | | | | 7.18 | 6 | 22 | 710 | 79.2 | 8.59 |
| Q53GQ0 | 47.44 | 1 | 11 | 11 | 21 | 3.53 | 1 | 1 | 3.53 | 1 | 1 | 3.53 | 1 | 1 | | | | | | | | | | | | | | | | | | | 47.44 | 10 | 18 | 312 | 34.3 | 9.32 |
| Q53H12 | 9.72 | 1 | 3 | 3 | 5 | | | | | | | | | | | | | | | | | | | | | | | | | | | | 9.72 | 3 | 5 | 422 | 47.1 | 8.09 |
| Q53H76 | 16.01 | 1 | 5 | 5 | 26 | 4.82 | 1 | 1 | 4.82 | 1 | 1 | | | | | | | | | | | | | | | | | | | | | | 16.01 | 5 | 24 | 456 | 49.7 | 7.37 |
| Q53H82 | 66.67 | 1 | 17 | 17 | 68 | 16.32 | 4 | 4 | 16.32 | 4 | 4 | 13.19 | 4 | 4 | 3.13 | 1 | 1 | 7.99 | 2 | 2 | | | | | | | 21.53 | 5 | 6 | 10.07 | 3 | 4 | 61.11 | 15 | 43 | 288 | 32.8 | 6.80 |
| Q8N7S2 | 27.30 | 1 | 1 | 7 | 12 | 2.97 | 1 | 1 | 2.97 | 1 | 1 | | | | | | | | | | | | | | | | | | | | | | 27.30 | 7 | 10 | 337 | 39.1 | 9.45 |
| Q53H96 | 51.46 | 1 | 11 | 11 | 20 | | | | 4.01 | 1 | 1 | | | | 4.01 | 1 | 1 | | | | | | | 4.01 | 1 | 1 | | | | 4.74 | 1 | 1 | 51.46 | 11 | 16 | 274 | 28.6 | 7.72 |
| Q53HC9 | 6.98 | 1 | 2 | 3 | 8 | | | | | | | | | | | | | 1.29 | 1 | 1 | | | | 3.36 | 1 | 1 | 3.36 | 1 | 1 | 3.62 | 2 | 2 | 6.98 | 3 | 3 | 387 | 43.6 | 5.06 |
| Q53QW1 | 19.24 | 1 | 5 | 5 | 21 | 3.80 | 1 | 1 | 2.03 | 1 | 1 | | | | 2.28 | 1 | 1 | | | | | | | | | | 8.10 | 3 | 3 | 2.03 | 1 | 1 | 19.24 | 5 | 14 | 395 | 41.6 | 5.12 |
| Q53T59 | 23.47 | 1 | 8 | 8 | 10 | | | | | | | | | | | | | | | | | | | 2.55 | 1 | 1 | | | | 2.81 | 1 | 1 | 20.66 | 6 | 8 | 392 | 42.8 | 5.01 |
| Q53TS8 | 14.13 | 2 | 7 | 7 | 9 | | | | 2.41 | 1 | 1 | | | | | | | | | | | | | | | | | | | | | | 11.72 | 6 | 8 | 623 | 71.1 | 7.02 |
| Q5BJE1 | 6.81 | 1 | 5 | 5 | 6 | | | | | | | | | | | | | 0.00 | | | | | | | | | | | | | | | 6.81 | 5 | 6 | 867 | 101.9 | 6.77 |
| Q5BJF6 | 65.86 | 3 | 97 | 98 | 1166 | 34.50 | 26 | 38 | 36.79 | 28 | 41 | 34.14 | 27 | 38 | 45.48 | 39 | 71 | 55.37 | 65 | 134 | 3.38 | 3 | 3 | 36.43 | 34 | 56 | 48.01 | 55 | 105 | 55.01 | 57 | 110 | 55.97 | 65 | 570 | 829 | 95.3 | 7.62 |
| Q5BN46 | 85.29 | 1 | 12 | 12 | 27 | | | | | | | | | | 32.35 | 4 | 4 | 36.03 | 5 | 5 | | | | | | | | | | | | | 66.18 | 9 | 18 | 136 | 15.3 | 8.81 |
| Q5CZC0 | 41.49 | 2 | 247 | 247 | 1019 | 1.72 | 10 | 10 | 0.52 | 3 | 3 | 0.29 | 2 | 2 | 4.99 | 29 | 30 | 2.55 | 17 | 18 | | | | 1.78 | 12 | 13 | 0.52 | 3 | 3 | 2.42 | 17 | 18 | 41.12 | 242 | 922 | 6907 | 780.1 | 6.71 |
| Q5D862 | 10.29 | 1 | 11 | 11 | 36 | | | | 0.79 | 1 | 1 | | | | 2.89 | 2 | 2 | 0.79 | 1 | 1 | | | | 4.48 | 4 | 4 | 8.28 | 8 | 12 | 2.09 | 1 | 2 | 5.23 | 6 | 14 | 2391 | 247.9 | 8.31 |
| Q5EBM0 | 19.82 | 1 | 4 | 4 | 5 | | | | | | | | | | | | | | | | | | | | | | | | | | | | 19.82 | 4 | 5 | 449 | 49.4 | 7.01 |
| Q5FVE4 | 50.45 | 2 | 25 | 25 | 74 | 8.26 | 4 | 4 | 7.06 | 4 | 4 | 6.46 | 4 | 4 | 1.95 | 1 | 1 | | | | | | | | | | | | | 2.10 | 1 | 1 | 47.15 | 23 | 60 | 666 | 74.3 | 8.46 |
| Q5GAN3 | 35.26 | 1 | 5 | 5 | 23 | 17.95 | 3 | 3 | 28.21 | 4 | 4 | 7.05 | 1 | 1 | | | | 10.90 | 2 | 2 | | | | | | | | | | | | | 35.26 | 5 | 13 | 156 | 17.8 | 8.59 |
| P61619 | 5.67 | 1 | 1 | 3 | 12 | | | | 1.89 | 1 | 1 | | | | | | | | | | | | | | | | | | | | | | 5.67 | 3 | 11 | 476 | 52.2 | 8.06 |
| Q5GLZ8 | 8.33 | 1 | 6 | 6 | 11 | | | | | | | | | | | | | | | | | | | | | | | | | | | | 8.33 | 6 | 11 | 1057 | 118.5 | 6.19 |
| Q5H943 | 19.47 | 1 | 2 | 2 | 4 | | | | | | | | | | | | | | | | | | | | | | | | | 9.73 | 1 | 1 | 19.47 | 2 | 3 | 113 | 12.8 | 10.20 |
| Q5H9R7 | 10.08 | 1 | 7 | 7 | 12 | | | | | | | | | | | | | | | | | | | | | | | | | | | | 10.08 | 7 | 12 | 873 | 97.6 | 4.60 |
| Q5HYA8 | 2.31 | 1 | 2 | 2 | 2 | | | | | | | | | | | | | | | | | | | | | | | | | | | | 2.31 | 2 | 2 | 995 | 111.7 | 6.77 |
| Q5HYI8 | 35.17 | 1 | 6 | 6 | 13 | 0.00 | | | 4.66 | 1 | 1 | 4.66 | 1 | 1 | | | | | | | | | | 4.66 | 1 | 1 | | | | 4.66 | 1 | 1 | 35.17 | 6 | 9 | 236 | 26.4 | 7.11 |
| Q5I0G3 | 20.85 | 2 | 11 | 11 | 17 | | | | | | | | | | 7.14 | 3 | 4 | 3.28 | 2 | 2 | | | | | | | | | | | | | 19.11 | 10 | 11 | 518 | 58.6 | 6.23 |
| Q5JPF3 | 0.84 | 3 | 2 | 2 | 8 | | | | | | | | | | | | | | | | | | | | | | | | | | | | 0.84 | 2 | 8 | 1778 | 199.6 | 7.83 |
| Q5JQC4 | 17.01 | 2 | 4 | 4 | 20 | 4.86 | 1 | 1 | 11.46 | 2 | 3 | | | | 4.17 | 1 | 1 | 7.99 | 1 | 2 | | | | | | | 7.99 | 1 | 3 | 7.99 | 1 | 1 | 9.03 | 2 | 9 | 288 | 30.1 | 4.25 |
| Q5JQC9 | 95.32 | 2 | 129 | 129 | 4627 | 65.22 | 45 | 107 | 54.68 | 41 | 103 | 59.25 | 43 | 84 | 70.14 | 56 | 150 | 75.53 | 70 | 158 | 7.49 | 7 | 9 | 52.93 | 44 | 77 | 64.87 | 58 | 123 | 72.01 | 69 | 143 | 93.56 | 114 | 3673 | 854 | 94.4 | 6.96 |
| Q5JRA6 | 16.41 | 1 | 25 | 25 | 51 | 0.94 | 1 | 1 | 4.51 | 6 | 6 | | | | | | | 4.82 | 7 | 7 | | | | | | | | | | | | | 14.00 | 19 | 37 | 1907 | 213.6 | 4.84 |
| Q5JRC9 | 14.29 | 1 | 8 | 8 | 9 | | | | | | | | | | | | | | | | | | | | | | | | | | | | 14.29 | 8 | 9 | 791 | 90.5 | 9.11 |
| Q5JRX3 | 49.57 | 1 | 39 | 39 | 87 | 7.71 | 5 | 5 | 13.31 | 11 | 12 | 5.21 | 4 | 4 | 0.96 | 1 | 1 | 0.96 | 1 | 1 | | | | | | | | | | | | | 46.87 | 36 | 64 | 1037 | 117.3 | 6.92 |
| Q5JST6 | 68.62 | 1 | 55 | 55 | 272 | 11.62 | 8 | 8 | 13.62 | 8 | 8 | 2.40 | 1 | 1 | 36.58 | 24 | 30 | 37.78 | 21 | 31 | | | | | | | | | | | | | 62.75 | 41 | 194 | 749 | 87.3 | 7.37 |
| Q5JTV8 | 18.52 | 1 | 8 | 8 | 14 | 2.06 | 1 | 1 | | | | | | | | | | 1.54 | 1 | 1 | | | | | | | 8.23 | 3 | 3 | | | | 10.29 | 5 | 9 | 583 | 66.2 | 8.18 |
| Q5JU00 | 26.95 | 1 | 8 | 8 | 17 | | | | | | | | | | 5.59 | 2 | 2 | | | | | | | | | | | | | | | | 25.35 | 7 | 15 | 501 | 55.6 | 6.80 |
| Q5JU67 | 34.04 | 1 | 16 | 16 | 30 | | | | | | | | | | 7.88 | 4 | 4 | 11.54 | 6 | 6 | | | | | | | | | | | | | 28.65 | 12 | 20 | 520 | 60.5 | 8.81 |
| Q5JU69 | 9.03 | 1 | 2 | 2 | 3 | 4.05 | 1 | 1 | | | | | | | | | | | | | | | | | | | | | | | | | 9.03 | 2 | 2 | 321 | 35.7 | 7.97 |
| Q5JUQ0 | 23.32 | 1 | 4 | 4 | 5 | | | | | | | | | | | | | | | | | | | | | | | | | | | | 23.32 | 4 | 5 | 283 | 31.9 | 8.06 |
| Q5JUR7 | 11.89 | 1 | 2 | 2 | 3 | | | | | | | | | | | | | | | | | | | | | | | | | | | | 11.89 | 2 | 3 | 227 | 25.6 | 8.66 |
| Q5JVL4 | 70.16 | 1 | 50 | 50 | 267 | 15.78 | 8 | 10 | 14.06 | 8 | 10 | 13.28 | 7 | 7 | 40.63 | 25 | 30 | 51.56 | 35 | 40 | | | | | | | 8.13 | 4 | 4 | | | | 65.16 | 39 | 166 | 640 | 73.9 | 6.16 |
| Q5JWF8 | 36.73 | 1 | 6 | 6 | 21 | | | | | | | | | | | | | | | | | | | | | | | | | | | | 36.73 | 6 | 21 | 245 | 26.7 | 9.32 |
| Q5JX69 | 41.52 | 1 | 4 | 7 | 13 | 14.62 | 2 | 2 | | | | | | | | | | | | | | | | | | | | | | | | | 41.52 | 7 | 11 | 171 | 19.5 | 8.84 |
| Q5JX71 | 25.15 | 1 | 2 | 5 | 10 | | | | | | | | | | | | | | | | | | | | | | | | | | | | 25.15 | 5 | 10 | 171 | 19.6 | 9.42 |
| Q5M9N0 | 1.26 | 1 | 2 | 2 | 2 | | | | | | | | | | | | | | | | | | | | | | | | | | | | 1.26 | 2 | 2 | 1113 | 127.1 | 6.46 |
| Q5QJ74 | 16.75 | 1 | 5 | 5 | 5 | | | | | | | | | | | | | | | | | | | | | | | | | 4.72 | 1 | 1 | 15.09 | 4 | 4 | 424 | 48.2 | 5.38 |
| Q5R3I4 | 33.05 | 1 | 13 | 13 | 23 | | | | | | | | | | | | | | | | | | | | | | | | | 1.49 | 1 | 1 | 31.56 | 12 | 22 | 469 | 52.8 | 5.99 |
| Q5RI15 | 22.88 | 1 | 3 | 3 | 4 | | | | 11.02 | 1 | 1 | | | | | | | | | | | | | | | | | | | | | | 22.88 | 3 | 3 | 118 | 13.3 | 8.76 |
| Q5SQI0 | 5.70 | 1 | 2 | 2 | 2 | | | | | | | | | | | | | | | | | | | | | | | | | | | | 5.70 | 2 | 2 | 421 | 46.8 | 9.95 |
| Q5SQS8 | 31.64 | 1 | 11 | 11 | 19 | | | | | | | | | | | | | 8.36 | 3 | 3 | | | | | | | | | | | | | 26.27 | 9 | 16 | 335 | 39.2 | 9.44 |
| Q5SRE5 | 33.68 | 2 | 50 | 50 | 118 | 3.49 | 4 | 4 | 3.54 | 5 | 5 | 0.91 | 1 | 1 | | | | | | | | | | | | | | | | | | | 32.93 | 48 | 108 | 1749 | 195.9 | 6.73 |
| Q5SRH9 | 15.17 | 1 | 8 | 8 | 10 | | | | | | | | | | | | | | | | | | | | | | | | | | | | 15.17 | 8 | 10 | 613 | 69.7 | 7.47 |
| Q5SRN2 | 16.52 | 1 | 7 | 7 | 13 | | | | | | | | | | | | | | | | | | | | | | | | | | | | 16.52 | 7 | 13 | 563 | 61.6 | 9.25 |
| Q969Z0 | 1.11 | 1 | 1 | 1 | 12 | | | | | | | | | | | | | | | | | | | | | | | | | | | | 1.11 | 1 | 12 | 631 | 70.7 | 7.42 |
| Q5SVJ3 | 38.10 | 1 | 5 | 5 | 9 | | | | | | | | | | | | | | | | | | | | | | | | | 8.16 | 1 | 1 | 38.10 | 5 | 8 | 147 | 17.6 | 9.61 |
| Q5SW79 | 1.77 | 2 | 3 | 3 | 3 | | | | | | | | | | | | | | | | | | | | | | | | | | | | 1.77 | 3 | 3 | 1584 | 175.2 | 7.11 |
| Q5SY80 | 3.58 | 1 | 3 | 3 | 4 | | | | | | | | | | | | | | | | | | | | | | | | | | | | 3.58 | 3 | 4 | 951 | 109.6 | 7.27 |
| Q5T0J7 | 12.45 | 1 | 3 | 3 | 4 | | | | | | | | | | 3.86 | 1 | 1 | | | | | | | | | | | | | | | | 8.58 | 2 | 3 | 233 | 26.5 | 9.11 |
| Q5T0N1 | 31.13 | 1 | 31 | 31 | 99 | 3.93 | 2 | 2 | | | | 1.25 | 1 | 1 | 2.41 | 2 | 2 | 6.24 | 7 | 7 | | | | | | | 1.16 | 1 | 1 | | | | 30.51 | 29 | 86 | 1121 | 125.6 | 5.81 |
| Q5T124 | 25.00 | 1 | 11 | 11 | 26 | | | | 2.88 | 1 | 1 | | | | 2.31 | 1 | 1 | 8.08 | 4 | 4 | | | | | | | 1.73 | 1 | 1 | | | | 19.62 | 8 | 19 | 520 | 57.3 | 5.19 |
| Q5T1B0 | 12.65 | 1 | 11 | 11 | 15 | | | | | | | | | | | | | 2.57 | 2 | 2 | | | | | | | | | | | | | 11.36 | 10 | 13 | 1012 | 118.0 | 5.68 |
| Q96PE2 | 0.73 | 1 | 1 | 2 | 12 | | | | | | | | | | | | | | | | | | | | | | | | | | | | 0.73 | 2 | 12 | 2063 | 221.5 | 6.29 |
| Q5T2S8 | 38.70 | 1 | 33 | 33 | 113 | 6.03 | 5 | 6 | 4.98 | 5 | 5 | | | | 1.05 | 1 | 1 | 4.60 | 4 | 4 | | | | | | | | | | | | | 35.73 | 30 | 97 | 1044 | 115.6 | 7.77 |
| Q5T440 | 19.10 | 1 | 3 | 3 | 4 | | | | | | | | | | | | | | | | | | | | | | | | | | | | 19.10 | 3 | 4 | 356 | 38.1 | 9.83 |
| Q5T447 | 10.45 | 1 | 7 | 7 | 9 | | | | | | | | | | | | | | | | | | | | | | | | | | | | 10.45 | 7 | 9 | 861 | 97.1 | 5.64 |
| Q5T4S7 | 20.66 | 3 | 79 | 80 | 204 | 0.68 | 2 | 2 | 0.89 | 3 | 3 | 1.22 | 5 | 5 | 0.37 | 1 | 1 | | | | | | | | | | | | | | | | 20.30 | 79 | 193 | 5183 | 573.5 | 6.04 |
| Q5T5A4 | 85.80 | 1 | 15 | 15 | 72 | 8.28 | 1 | 1 | 27.81 | 3 | 4 | | | | 52.07 | 7 | 14 | 45.56 | 7 | 12 | | | | | | | | | | | | | 83.43 | 12 | 41 | 169 | 19.3 | 9.28 |
| Q5T5S1 | 58.80 | 4 | 38 | 39 | 110 | 1.31 | 1 | 1 | 7.30 | 4 | 4 | 0.00 | | | 18.73 | 9 | 10 | 27.90 | 16 | 21 | | | | | | | | | | 4.31 | 2 | 2 | 55.43 | 33 | 72 | 534 | 62.6 | 8.79 |
| Q5T655 | 70.30 | 2 | 82 | 83 | 245 | 1.26 | 1 | 1 | 10.09 | 7 | 7 | | | | 40.48 | 32 | 39 | 58.49 | 53 | 62 | | | | | | | | | | 5.05 | 3 | 3 | 57.00 | 58 | 133 | 872 | 103.4 | 8.41 |
| Q5T681 | 34.53 | 1 | 7 | 7 | 15 | | | | 3.59 | 1 | 1 | | | | | | | 4.04 | 1 | 1 | | | | 7.62 | 2 | 2 | | | | | | | 30.49 | 6 | 11 | 223 | 25.1 | 7.59 |
| Q5T6J7 | 14.97 | 1 | 2 | 2 | 4 | | | | | | | | | | | | | | | | | | | | | | | | | | | | 14.97 | 2 | 4 | 187 | 20.6 | 6.21 |
| Q5T6V5 | 26.69 | 1 | 8 | 8 | 12 | | | | | | | | | | | | | | | | | | | | | | | | | | | | 26.69 | 8 | 12 | 341 | 39.0 | 5.88 |
| Q5T749 | 3.45 | 1 | 2 | 2 | 4 | | | | | | | | | | | | | | | | | | | | | | | | | | | | 3.45 | 2 | 4 | 579 | 64.1 | 8.27 |
| Q5T750 | 8.00 | 1 | 2 | 2 | 4 | | | | | | | | | | | | | | | | | | | | | | | | | | | | 8.00 | 2 | 4 | 250 | 26.2 | 7.97 |
| Q5T871 | 54.08 | 1 | 4 | 4 | 21 | 17.35 | 1 | 1 | | | | | | | | | | 24.49 | 1 | 1 | | | | | | | | | | | | | 54.08 | 4 | 19 | 98 | 10.7 | 8.03 |
| Q5T9G4 | 50.00 | 1 | 13 | 13 | 59 | 7.06 | 2 | 2 | 3.53 | 1 | 1 | 3.53 | 1 | 1 | | | | | | | | | | | | | | | | | | | 50.00 | 13 | 55 | 340 | 38.6 | 7.88 |
| Q5TA50 | 48.13 | 1 | 8 | 9 | 25 | | | | | | | | | | | | | | | | | | | | | | | | | | | | 48.13 | 9 | 25 | 214 | 24.3 | 7.21 |
| Q5TCS8 | 38.36 | 1 | 65 | 65 | 138 | | | | 0.00 | | | 0.00 | | | 6.23 | 10 | 10 | 10.52 | 16 | 19 | | | | | | | | | | | | | 35.27 | 58 | 109 | 1911 | 221.3 | 5.01 |
| Q5TD94 | 10.47 | 1 | 5 | 5 | 11 | | | | | | | | | | | | | | | | | | | | | | | | | | | | 10.47 | 5 | 11 | 716 | 80.7 | 4.44 |
| O60488 | 5.49 | 1 | 1 | 3 | 11 | 2.53 | 1 | 1 | | | | | | | | | | | | | | | | | | | | | | | | | 5.49 | 3 | 10 | 711 | 79.1 | 8.38 |
| Q5TD97 | 53.52 | 1 | 12 | 12 | 17 | | | | | | | | | | | | | | | | | | | | | | | | | | | | 53.52 | 12 | 17 | 284 | 32.7 | 7.52 |
| Q5TEA6 | 6.25 | 1 | 4 | 4 | 4 | | | | | | | | | | | | | | | | | | | | | | | | | | | | 6.25 | 4 | 4 | 688 | 77.9 | 9.23 |
| Q5TEZ5 | 66.87 | 1 | 23 | 23 | 66 | 3.34 | 1 | 1 | 6.69 | 2 | 2 | | | | 34.04 | 9 | 11 | 48.33 | 15 | 21 | | | | | | | | | | 6.99 | 2 | 2 | 55.02 | 18 | 29 | 329 | 38.5 | 6.96 |
| Q5TFE4 | 13.85 | 1 | 5 | 5 | 9 | | | | | | | | | | | | | | | | | | | | | | | | | | | | 13.85 | 5 | 9 | 455 | 51.8 | 6.35 |
| Q5TH69 | 4.36 | 1 | 6 | 6 | 7 | | | | | | | | | | | | | | | | | | | | | | | | | | | | 4.36 | 6 | 7 | 2177 | 240.5 | 5.82 |
| Q5TH74 | 42.22 | 1 | 9 | 9 | 21 | | | | | | | | | | 5.39 | 1 | 1 | 5.39 | 1 | 1 | | | | | | | | | | | | | 42.22 | 9 | 19 | 334 | 36.8 | 9.76 |
| Q5THR3 | 53.36 | 1 | 78 | 78 | 280 | 4.80 | 6 | 6 | 7.86 | 10 | 11 | 1.47 | 2 | 2 | 12.46 | 15 | 16 | 17.26 | 21 | 25 | | | | | | | | | | | | | 49.37 | 71 | 220 | 1501 | 172.8 | 8.40 |
| Q5TZF3 | 18.09 | 1 | 4 | 4 | 7 | 6.03 | 1 | 1 | 6.03 | 1 | 2 | | | | | | | | | | | | | | | | | | | | | | 18.09 | 4 | 4 | 282 | 31.8 | 4.64 |
| Q5U5X0 | 54.81 | 1 | 5 | 5 | 6 | | | | 9.62 | 1 | 1 | | | | | | | | | | | | | | | | | | | | | | 45.19 | 4 | 5 | 104 | 11.9 | 9.66 |
| Q5VIR6 | 4.43 | 1 | 2 | 2 | 6 | | | | | | | | | | | | | | | | | | | | | | | | | | | | 4.43 | 2 | 6 | 699 | 79.6 | 6.02 |
| Q5VST6 | 30.56 | 1 | 6 | 6 | 7 | | | | | | | | | | | | | | | | | | | | | | | | | | | | 30.56 | 6 | 7 | 288 | 32.2 | 6.29 |
| Q5VTH2 | 59.89 | 1 | 10 | 10 | 48 | | | | | | | | | | 33.33 | 5 | 7 | 45.76 | 7 | 9 | 9.04 | 2 | 2 | | | | | | | | | | 57.06 | 8 | 30 | 177 | 19.3 | 10.08 |
| Q5VTH9 | 30.42 | 1 | 23 | 23 | 49 | | | | | | | 0.83 | 1 | 1 | 5.54 | 4 | 5 | 8.73 | 7 | 8 | | | | | | | | | | | | | 28.54 | 20 | 35 | 848 | 94.5 | 5.76 |
| Q5VTT2 | 67.69 | 1 | 14 | 14 | 39 | | | | 6.55 | 1 | 1 | | | | 39.30 | 6 | 6 | 50.22 | 9 | 13 | | | | | | | 6.55 | 1 | 1 | | | | 57.21 | 10 | 18 | 229 | 26.4 | 7.21 |
| Q5VU65 | 40.68 | 1 | 73 | 73 | 327 | 12.34 | 17 | 19 | 25.79 | 39 | 47 | 27.97 | 46 | 60 | 1.59 | 3 | 3 | 4.34 | 7 | 8 | | | | | | | | | | | | | 35.65 | 63 | 190 | 1888 | 210.5 | 7.50 |
| Q5VU69 | 65.35 | 1 | 10 | 10 | 23 | 12.87 | 1 | 1 | 6.93 | 1 | 1 | | | | 26.73 | 2 | 2 | 34.65 | 3 | 5 | | | | | | | | | | 20.79 | 2 | 3 | 63.37 | 8 | 11 | 101 | 12.1 | 9.70 |
| Q5VUJ9 | 33.46 | 1 | 7 | 7 | 11 | | | | | | | | | | 10.41 | 2 | 3 | 7.43 | 2 | 2 | | | | | | | | | | | | | 26.02 | 4 | 6 | 269 | 29.7 | 8.79 |
| Q5VW32 | 16.30 | 1 | 6 | 6 | 8 | | | | | | | 2.68 | 1 | 1 | | | | | | | | | | | | | | | | | | | 16.30 | 6 | 7 | 411 | 46.4 | 7.65 |
| Q5VW36 | 2.11 | 1 | 3 | 4 | 7 | | | | | | | | | | | | | | | | | | | | | | | | | | | | 2.11 | 4 | 7 | 1801 | 199.9 | 6.62 |
| Q5VWP2 | 7.93 | 1 | 2 | 2 | 3 | | | | | | | | | | | | | | | | | | | | | | | | | | | | 7.93 | 2 | 3 | 391 | 44.9 | 5.69 |
| Q5VWZ2 | 24.05 | 1 | 5 | 5 | 6 | | | | | | | | | | | | | | | | | | | | | | | | | | | | 24.05 | 5 | 6 | 237 | 26.3 | 7.84 |
| Q5VYK3 | 21.46 | 1 | 32 | 32 | 48 | 0.65 | 1 | 1 | 0.65 | 1 | 1 | | | | | | | | | | | | | | | | | | | 0.00 | | | 20.81 | 31 | 46 | 1845 | 204.2 | 7.12 |
| Q5VZ52 | 57.14 | 1 | 7 | 7 | 19 | | | | | | | | | | 21.74 | 3 | 4 | 26.09 | 4 | 4 | | | | | | | 16.77 | 2 | 3 | | | | 45.34 | 5 | 8 | 161 | 18.7 | 5.88 |
| Q5VZ72 | 26.78 | 1 | 6 | 6 | 12 | | | | | | | | | | | | | | | | | | | | | | | | | | | | 26.78 | 6 | 12 | 239 | 27.7 | 8.29 |
| Q5VZQ5 | 34.95 | 1 | 6 | 6 | 7 | | | | | | | | | | | | | | | | | | | | | | | | | | | | 34.95 | 6 | 7 | 186 | 21.5 | 9.76 |
| Q5W041 | 50.11 | 1 | 35 | 35 | 93 | 4.47 | 3 | 3 | 3.10 | 2 | 2 | | | | | | | 4.47 | 3 | 3 | | | | | | | | | | | | | 49.66 | 34 | 85 | 872 | 96.3 | 6.21 |
| Q5W0Z9 | 8.22 | 1 | 2 | 2 | 2 | | | | | | | | | | | | | | | | | | | | | | | | | | | | 8.22 | 2 | 2 | 365 | 42.2 | 7.71 |
| Q5W111 | 44.39 | 1 | 9 | 9 | 19 | | | | 12.76 | 2 | 2 | 5.10 | 1 | 1 | | | | | | | | | | | | | | | | | | | 36.73 | 8 | 16 | 196 | 21.7 | 6.70 |
| Q5XKP0 | 46.61 | 1 | 4 | 4 | 6 | | | | | | | | | | | | | | | | | | | | | | | | | | | | 46.61 | 4 | 6 | 118 | 13.1 | 9.42 |
| Q5XPI4 | 2.21 | 1 | 2 | 2 | 3 | | | | | | | | | | | | | | | | | | | | | | | | | | | | 2.21 | 2 | 3 | 1314 | 148.4 | 6.74 |
| Q5XUX1 | 14.75 | 1 | 6 | 6 | 7 | | | | | | | | | | | | | | | | | | | | | | | | | | | | 14.75 | 6 | 7 | 488 | 54.1 | 6.48 |
| Q5XX13 | 9.22 | 2 | 9 | 9 | 15 | | | | | | | | | | | | | | | | | | | | | | | | | 0.00 | | | 9.22 | 9 | 15 | 1052 | 119.8 | 9.39 |
| Q63HN8 | 5.26 | 1 | 25 | 25 | 30 | | | | | | | | | | | | | | | | | | | | | | | | | | | | 5.26 | 25 | 30 | 5207 | 591.0 | 6.48 |
| Q658Y4 | 7.40 | 4 | 3 | 4 | 5 | | | | 0.72 | 1 | 1 | | | | | | | | | | | | | | | | | | | | | | 7.40 | 4 | 4 | 838 | 93.8 | 6.39 |
| Q66GS9 | 6.93 | 4 | 6 | 8 | 11 | 0.00 | | | 0.00 | | | 0.00 | | | 0.00 | | | 0.00 | | | | | | 0.70 | 1 | 1 | | | | | | | 6.93 | 8 | 10 | 1140 | 133.4 | 6.21 |
| Q66K14 | 2.88 | 1 | 2 | 2 | 5 | | | | | | | | | | | | | | | | | | | | | | | | | | | | 2.88 | 2 | 5 | 1250 | 140.4 | 5.25 |
| Q66K79 | 48.62 | 1 | 25 | 25 | 120 | 16.41 | 9 | 11 | 18.87 | 8 | 11 | 15.34 | 8 | 11 | 6.90 | 4 | 4 | | | | | | | | | | | | | 2.91 | 2 | 2 | 43.25 | 22 | 81 | 652 | 73.6 | 7.97 |
| Q676U5 | 7.74 | 1 | 3 | 3 | 4 | | | | | | | | | | | | | | | | | | | 3.46 | 2 | 2 | | | | 1.81 | 1 | 1 | 4.28 | 1 | 1 | 607 | 68.2 | 6.64 |
| Q687X5 | 45.10 | 1 | 16 | 16 | 76 | 9.59 | 3 | 3 | | | | 11.11 | 4 | 4 | | | | | | | | | | | | | | | | 4.79 | 1 | 1 | 45.10 | 15 | 68 | 459 | 51.9 | 9.29 |
| Q68CQ1 | 26.38 | 1 | 27 | 27 | 73 | 1.97 | 2 | 2 | 1.81 | 2 | 2 | 0.83 | 1 | 1 | | | | | | | | | | | | | | | | | | | 25.40 | 26 | 68 | 1323 | 145.6 | 6.96 |
| Q68CQ7 | 11.32 | 1 | 3 | 3 | 3 | | | | | | | | | | | | | | | | | | | | | | | | | | | | 11.32 | 3 | 3 | 371 | 41.9 | 9.35 |
| Q68DQ2 | 4.68 | 1 | 9 | 9 | 14 | | | | | | | 0.44 | 1 | 1 | | | | | | | | | | 0.71 | 1 | 1 | 2.19 | 4 | 4 | | | | 3.20 | 6 | 8 | 2970 | 330.4 | 5.20 |
| Q6BCY4 | 77.17 | 1 | 18 | 18 | 70 | 44.20 | 9 | 11 | 56.16 | 12 | 15 | 36.23 | 8 | 9 | 9.06 | 3 | 3 | 15.22 | 3 | 3 | | | | 0.00 | | | 2.54 | 1 | 1 | 6.88 | 2 | 2 | 75.72 | 16 | 26 | 276 | 31.4 | 8.50 |
| Q6DD88 | 36.60 | 1 | 14 | 14 | 41 | 7.39 | 4 | 4 | 7.21 | 4 | 4 | 7.39 | 3 | 3 | 2.03 | 1 | 1 | | | | | | | | | | | | | | | | 36.60 | 12 | 29 | 541 | 60.5 | 5.66 |
| Q6DKJ4 | 13.79 | 1 | 4 | 4 | 10 | | | | | | | | | | | | | | | | | | | | | | | | | | | | 13.79 | 4 | 10 | 435 | 48.4 | 4.97 |
| Q6FI81 | 25.00 | 1 | 6 | 6 | 10 | 9.29 | 1 | 1 | 3.85 | 1 | 1 | | | | 3.85 | 1 | 1 | | | | | | | | | | | | | | | | 15.71 | 5 | 7 | 312 | 33.6 | 5.62 |
| Q6GMR7 | 1.50 | 1 | 2 | 2 | 2 | | | | 0.00 | | | 0.00 | | | | | | | | | | | | | | | | | | | | | 1.50 | 2 | 2 | 532 | 58.3 | 9.07 |
| Q6GMV3 | 30.71 | 1 | 4 | 4 | 11 | | | | 15.00 | 2 | 2 | 8.57 | 1 | 1 | | | | | | | | | | 18.57 | 2 | 2 | 8.57 | 1 | 1 | 14.29 | 2 | 2 | 24.29 | 3 | 3 | 140 | 15.8 | 9.10 |
| Q8TF72 | 0.45 | 1 | 1 | 1 | 11 | | | | | | | | | | | | | | | | | | | | | | 0.45 | 1 | 1 | | | | 0.45 | 1 | 10 | 1996 | 216.7 | 7.80 |
| Q6GYQ0 | 2.60 | 1 | 4 | 4 | 6 | | | | | | | | | | | | | | | | | | | | | | | | | | | | 2.60 | 4 | 6 | 2036 | 229.7 | 6.19 |
| Q6IA69 | 16.71 | 1 | 10 | 10 | 14 | | | | 1.70 | 1 | 1 | | | | | | | | | | | | | | | | 0.00 | | | | | | 16.71 | 10 | 13 | 706 | 79.2 | 6.44 |
| Q6IA86 | 5.69 | 1 | 3 | 3 | 5 | | | | | | | | | | | | | | | | | | | | | | | | | | | | 5.69 | 3 | 5 | 826 | 92.4 | 5.96 |
| Q6IAA8 | 31.06 | 1 | 3 | 3 | 3 | 8.07 | 1 | 1 | | | | | | | | | | | | | | | | | | | | | | | | | 22.98 | 2 | 2 | 161 | 17.7 | 5.15 |
| Q13362 | 11.45 | 1 | 1 | 5 | 11 | | | | | | | | | | | | | | | | | | | | | | | | | | | | 11.45 | 5 | 11 | 524 | 61.0 | 6.87 |
| Q6IAN0 | 39.69 | 1 | 11 | 11 | 17 | | | | 10.77 | 3 | 3 | | | | | | | | | | | | | | | | | | | | | | 37.23 | 10 | 14 | 325 | 35.1 | 9.55 |
| Q6IBS0 | 20.06 | 1 | 3 | 6 | 12 | | | | | | | | | | | | | | | | | | | 2.01 | 1 | 1 | | | | 3.72 | 1 | 2 | 14.33 | 4 | 9 | 349 | 39.5 | 6.84 |
| Q6ICG8 | 16.18 | 1 | 5 | 5 | 11 | | | | 5.83 | 1 | 1 | | | | | | | 3.24 | 1 | 1 | | | | | | | | | | | | | 16.18 | 5 | 9 | 309 | 31.9 | 7.84 |
| Q6IPR1 | 24.44 | 1 | 3 | 3 | 4 | | | | 17.78 | 2 | 2 | | | | | | | | | | | | | | | | | | | | | | 14.44 | 2 | 2 | 90 | 10.9 | 9.92 |
| Q6IQ20 | 11.45 | 1 | 3 | 3 | 5 | 3.56 | 1 | 1 | | | | | | | | | | | | | | | | | | | | | | | | | 7.89 | 2 | 4 | 393 | 45.6 | 6.06 |
| Q6IWH7 | 6.54 | 1 | 4 | 4 | 7 | | | | | | | | | | | | | | | | | | | | | | | | | | | | 6.54 | 4 | 7 | 933 | 105.5 | 7.87 |
| Q6J272 | 79.50 | 1 | 27 | 27 | 200 | 41.32 | 10 | 12 | 35.02 | 10 | 12 | 17.67 | 4 | 4 | 58.36 | 18 | 23 | 66.25 | 20 | 34 | | | | | | | 6.94 | 2 | 2 | 7.89 | 2 | 2 | 73.50 | 20 | 112 | 317 | 36.1 | 7.81 |
| Q6JEL2 | 53.62 | 1 | 27 | 27 | 79 | 1.81 | 1 | 1 | 11.18 | 6 | 6 | 22.20 | 10 | 10 | 8.06 | 4 | 4 | 3.95 | 2 | 2 | | | | | | | | | | 1.81 | 1 | 1 | 48.68 | 24 | 55 | 608 | 68.9 | 5.68 |
| Q6KB66 | 7.30 | 4 | 2 | 4 | 7 | | | | | | | | | | 2.43 | 1 | 1 | 2.43 | 1 | 1 | | | | 2.43 | 1 | 1 | 3.54 | 2 | 2 | 2.43 | 1 | 1 | 1.33 | 1 | 1 | 452 | 50.5 | 5.67 |
| Q6KC79 | 0.46 | 1 | 2 | 2 | 2 | | | | | | | | | | | | | | | | | | | | | | | | | | | | 0.46 | 2 | 2 | 2804 | 315.9 | 7.91 |
| Q6N063 | 8.86 | 1 | 3 | 3 | 6 | | | | | | | | | | | | | | | | | | | | | | | | | | | | 8.86 | 3 | 6 | 350 | 39.0 | 5.68 |
| Q6NT32 | 7.48 | 1 | 4 | 4 | 9 | | | | 1.57 | 1 | 1 | | | | | | | | | | | | | | | | | | | | | | 7.48 | 4 | 8 | 575 | 63.9 | 6.44 |
| Q6NUK1 | 17.19 | 1 | 9 | 9 | 10 | 2.73 | 1 | 1 | 1.68 | 1 | 1 | 1.89 | 1 | 1 | | | | | | | | | | | | | | | | | | | 13.63 | 7 | 7 | 477 | 53.3 | 6.33 |
| Q6NUT2 | 17.28 | 3 | 12 | 12 | 56 | 1.45 | 1 | 1 | 2.51 | 2 | 2 | 2.51 | 2 | 2 | | | | | | | | | | | | | | | | | | | 17.28 | 12 | 51 | 758 | 87.3 | 9.10 |
| Q6NVY1 | 26.94 | 1 | 10 | 10 | 16 | 8.29 | 3 | 3 | 8.81 | 3 | 3 | 8.55 | 3 | 3 | | | | | | | | | | | | | | | | | | | 16.32 | 6 | 7 | 386 | 43.5 | 8.19 |
| Q6NXE6 | 37.52 | 1 | 15 | 15 | 23 | | | | | | | | | | | | | | | | | | | | | | | | | 3.59 | 1 | 1 | 37.52 | 15 | 22 | 501 | 54.1 | 6.24 |
| Q6NXR0 | 54.64 | 1 | 28 | 28 | 122 | | | | | | | | | | 4.32 | 2 | 2 | | | | | | | 14.69 | 6 | 7 | 20.73 | 8 | 10 | 12.31 | 5 | 5 | 52.48 | 25 | 98 | 463 | 50.3 | 5.35 |
| Q6NXR4 | 5.31 | 1 | 2 | 2 | 2 | | | | 1.18 | 1 | 1 | 0.00 | | | | | | | | | | | | | | | | | | | | | 4.13 | 1 | 1 | 508 | 56.9 | 7.09 |
| Q6P047 | 22.79 | 1 | 6 | 6 | 6 | | | | 0.00 | | | | | | | | | | | | | | | | | | | | | | | | 22.79 | 6 | 6 | 294 | 33.7 | 7.88 |
| Q6P1A2 | 16.02 | 1 | 7 | 7 | 13 | | | | | | | | | | | | | | | | | | | | | | | | | | | | 16.02 | 7 | 13 | 487 | 56.0 | 8.69 |
| Q6P1M0 | 18.51 | 1 | 10 | 10 | 15 | | | | | | | | | | | | | | | | | | | | | | | | | | | | 18.51 | 10 | 15 | 643 | 72.0 | 8.47 |
| Q6P1X6 | 25.00 | 1 | 4 | 4 | 4 | | | | | | | | | | | | | | | | | | | | | | | | | | | | 25.00 | 4 | 4 | 216 | 23.9 | 9.14 |
| Q6P2D8 | 4.42 | 1 | 3 | 3 | 5 | | | | | | | | | | | | | | | | | | | | | | | | | | | | 4.42 | 3 | 5 | 792 | 89.8 | 8.92 |
| Q6P2H8 | 19.13 | 1 | 5 | 5 | 7 | | | | | | | | | | | | | | | | | | | | | | | | | | | | 19.13 | 5 | 7 | 277 | 31.6 | 8.53 |
| Q6P2I3 | 45.86 | 1 | 3 | 12 | 30 | 24.20 | 5 | 7 | 21.34 | 4 | 4 | 19.11 | 5 | 6 | | | | | | | | | | | | | | | | | | | 42.99 | 10 | 13 | 314 | 34.6 | 7.75 |
| Q6P3X3 | 24.56 | 1 | 15 | 15 | 29 | | | | 1.07 | 1 | 1 | | | | | | | | | | | | | | | | | | | | | | 23.49 | 14 | 28 | 843 | 96.6 | 5.59 |
| Q6P4A7 | 9.79 | 1 | 2 | 2 | 2 | | | | | | | | | | | | | | | | | | | | | | | | | | | | 9.79 | 2 | 2 | 337 | 38.0 | 9.19 |
| Q6P4A8 | 7.78 | 1 | 4 | 4 | 7 | 7.78 | 4 | 5 | | | | 3.62 | 2 | 2 | | | | | | | | | | | | | | | | | | | | | | 553 | 63.2 | 9.06 |
| Q6P656 | 62.79 | 1 | 17 | 17 | 74 | 11.30 | 2 | 2 | 15.61 | 3 | 3 | | | | 32.56 | 8 | 10 | 29.90 | 9 | 11 | | | | | | | | | | | | | 61.46 | 14 | 48 | 301 | 34.3 | 6.28 |
| P07948 | 5.08 | 18 | 1 | 3 | 10 | 0.00 | | | 0.00 | | | 1.56 | 1 | 1 | | | | | | | | | | | | | | | | | | | 5.08 | 3 | 9 | 512 | 58.5 | 7.11 |
| Q6P996 | 12.44 | 2 | 7 | 7 | 14 | 0.00 | | | | | | | | | | | | | | | | | | | | | | | | 1.52 | 1 | 1 | 10.91 | 6 | 13 | 788 | 86.7 | 5.38 |
| Q6P9G0 | 42.54 | 1 | 8 | 8 | 17 | | | | 10.96 | 2 | 2 | | | | 25.44 | 4 | 4 | 6.14 | 2 | 2 | | | | | | | | | | | | | 36.84 | 5 | 9 | 228 | 26.7 | 5.60 |
| Q6PCB0 | 34.38 | 1 | 8 | 8 | 12 | 11.24 | 2 | 2 | 7.64 | 2 | 2 | | | | | | | | | | | | | | | | | | | | | | 25.39 | 6 | 8 | 445 | 46.8 | 7.68 |
| Q6PEW0 | 22.53 | 1 | 5 | 5 | 18 | 20.00 | 4 | 8 | 11.65 | 3 | 4 | 9.37 | 2 | 2 | | | | | | | | | | | | | | | | | | | 7.09 | 2 | 4 | 395 | 43.8 | 6.70 |
| Q6PF18 | 36.25 | 1 | 8 | 8 | 14 | | | | | | | | | | 10.83 | 2 | 2 | 4.58 | 1 | 1 | | | | | | | | | | | | | 31.25 | 6 | 11 | 240 | 27.6 | 8.44 |
| Q6PGP7 | 21.04 | 1 | 25 | 25 | 50 | | | | | | | | | | | | | | | | | | | | | | | | | 2.37 | 3 | 3 | 21.04 | 25 | 47 | 1564 | 175.4 | 7.53 |
| Q6PGQ1 | 17.47 | 1 | 3 | 3 | 17 | 9.61 | 2 | 2 | 17.47 | 3 | 4 | | | | | | | | | | | | | | | | | | | | | | 17.47 | 3 | 11 | 229 | 25.0 | 4.11 |
| Q6PI48 | 3.88 | 1 | 2 | 2 | 2 | | | | | | | | | | | | | | | | | | | | | | | | | | | | 3.88 | 2 | 2 | 645 | 73.5 | 8.02 |
| Q6PIY5 | 47.50 | 1 | 15 | 15 | 28 | | | | | | | | | | | | | 2.27 | 1 | 1 | | | | | | | | | | | | | 47.50 | 15 | 27 | 440 | 48.8 | 5.74 |
| Q6PJG6 | 9.01 | 1 | 4 | 5 | 5 | | | | | | | | | | | | | | | | | | | | | | | | | | | | 9.01 | 5 | 5 | 821 | 88.1 | 5.27 |
| Q6Q759 | 23.53 | 1 | 36 | 36 | 93 | | | | 2.16 | 4 | 4 | 0.40 | 1 | 1 | 0.00 | | | 0.67 | 1 | 1 | | | | 0.00 | | | 0.67 | 1 | 1 | 0.00 | | | 22.76 | 34 | 86 | 2223 | 251.6 | 6.24 |
| Q6QHF9 | 10.17 | 1 | 3 | 3 | 6 | | | | | | | | | | | | | | | | | | | | | | | | | | | | 10.17 | 3 | 6 | 649 | 70.2 | 5.53 |
| Q13214 | 2.67 | 3 | 1 | 3 | 10 | 0.80 | 1 | 1 | 0.80 | 1 | 1 | 0.80 | 1 | 1 | | | | | | | | | | | | | 0.80 | 1 | 1 | 1.47 | 2 | 3 | 2.00 | 2 | 3 | 749 | 83.1 | 8.91 |
| Q6RW13 | 29.56 | 1 | 3 | 3 | 4 | | | | | | | | | | | | | | | | | | | | | | | | | | | | 29.56 | 3 | 4 | 159 | 17.4 | 6.14 |
| Q6SA08 | 27.44 | 1 | 9 | 9 | 13 | | | | | | | | | | | | | | | | | | | | | | | | | | | | 27.44 | 9 | 13 | 328 | 37.4 | 8.57 |
| Q6TDU7 | 38.13 | 1 | 23 | 24 | 52 | 1.68 | 1 | 1 | 4.19 | 3 | 3 | | | | 4.61 | 3 | 3 | 12.99 | 7 | 7 | | | | | | | | | | | | | 38.13 | 23 | 38 | 716 | 83.1 | 5.29 |
| Q6UDR6 | 41.41 | 1 | 3 | 3 | 4 | | | | | | | | | | | | | | | | | | | | | | | | | | | | 41.41 | 3 | 4 | 99 | 11.4 | 9.23 |
| Q6URK8 | 66.42 | 1 | 16 | 16 | 70 | 8.86 | 2 | 2 | 6.64 | 2 | 2 | | | | 33.58 | 9 | 12 | 40.96 | 10 | 12 | | | | | | | | | | 9.96 | 3 | 3 | 59.78 | 13 | 39 | 271 | 30.7 | 8.82 |
| Q9H0N0 | 11.02 | 1 | 1 | 2 | 10 | | | | 2.36 | 1 | 1 | | | | | | | | | | | | | | | | | | | | | | 11.02 | 2 | 9 | 254 | 28.3 | 7.71 |
| Q6UW02 | 17.10 | 1 | 6 | 6 | 7 | | | | | | | | | | | | | | | | | | | | | | | | | | | | 17.10 | 6 | 7 | 462 | 52.4 | 6.21 |
| Q6UW15 | 41.14 | 1 | 5 | 5 | 21 | 14.29 | 2 | 2 | 27.43 | 3 | 4 | | | | 17.14 | 2 | 2 | 30.86 | 4 | 5 | | | | | | | | | | | | | 33.71 | 4 | 8 | 175 | 19.3 | 7.09 |
| Q6UW49 | 45.14 | 1 | 15 | 15 | 66 | 41.71 | 11 | 12 | 40.57 | 10 | 12 | 16.29 | 4 | 4 | 14.29 | 3 | 5 | 9.71 | 3 | 3 | | | | | | | | | | | | | 34.57 | 8 | 30 | 350 | 38.9 | 5.73 |
| Q6UW68 | 33.33 | 1 | 7 | 7 | 43 | 16.93 | 3 | 4 | 16.93 | 3 | 6 | | | | | | | | | | | | | | | | | | | | | | 32.28 | 6 | 33 | 189 | 21.2 | 8.62 |
| Q6UWB4 | 35.80 | 1 | 9 | 9 | 19 | 11.36 | 3 | 3 | 2.56 | 1 | 1 | | | | | | | | | | | | | | | | | | | | | | 35.80 | 9 | 15 | 352 | 38.8 | 7.53 |
| Q6UWM5 | 31.82 | 1 | 8 | 8 | 27 | 11.16 | 2 | 2 | 2.48 | 1 | 2 | 6.20 | 2 | 3 | | | | | | | | | | | | | | | | | | | 23.14 | 7 | 20 | 242 | 27.1 | 8.21 |
| Q6UWN0 | 11.38 | 1 | 2 | 2 | 7 | | | | 4.07 | 1 | 1 | 4.07 | 1 | 1 | 4.07 | 1 | 1 | | | | | | | | | | | | | | | | 11.38 | 2 | 4 | 246 | 26.7 | 8.38 |
| Q6UWP2 | 5.77 | 1 | 2 | 2 | 3 | | | | 5.77 | 2 | 2 | | | | | | | | | | | | | | | | | | | | | | 4.23 | 1 | 1 | 260 | 28.3 | 6.64 |
| Q6UWP8 | 35.08 | 1 | 10 | 10 | 21 | | | | 9.15 | 1 | 1 | | | | | | | 11.53 | 2 | 2 | | | | 23.05 | 5 | 6 | 27.29 | 7 | 9 | 9.15 | 1 | 1 | 9.15 | 1 | 2 | 590 | 60.5 | 7.01 |
| Q6UWQ5 | 13.51 | 1 | 2 | 4 | 11 | 8.78 | 2 | 4 | 8.11 | 1 | 2 | 4.73 | 1 | 1 | | | | | | | | | | | | | | | | | | | 12.84 | 3 | 4 | 148 | 16.6 | 8.05 |
| Q6UWU2 | 30.12 | 1 | 16 | 16 | 48 | 27.37 | 14 | 14 | 19.42 | 10 | 11 | 9.48 | 4 | 4 | | | | | | | | | | | | | | | | | | | 21.25 | 10 | 19 | 654 | 74.1 | 8.92 |
| Q6UWW0 | 27.72 | 1 | 4 | 4 | 5 | | | | | | | | | | | | | | | | | | | | | | | | | | | | 27.72 | 4 | 5 | 184 | 20.4 | 4.94 |
| Q6UX06 | 52.94 | 2 | 28 | 28 | 107 | 16.08 | 8 | 9 | 28.63 | 14 | 19 | 50.59 | 25 | 56 | | | | 10.59 | 6 | 7 | | | | | | | | | | | | | 23.73 | 12 | 16 | 510 | 57.2 | 5.69 |
| Q6UXG2 | 20.24 | 1 | 17 | 17 | 47 | 1.09 | 1 | 1 | 1.09 | 1 | 1 | 10.46 | 9 | 9 | | | | | | | | | | | | | | | | | | | 17.18 | 15 | 36 | 1013 | 111.3 | 6.55 |
| Q6UXH1 | 12.75 | 1 | 4 | 4 | 9 | 2.83 | 1 | 1 | 3.40 | 1 | 1 | | | | | | | 4.53 | 1 | 1 | | | | | | | | | | | | | 12.75 | 4 | 6 | 353 | 38.2 | 4.59 |
| Q6UXI9 | 24.96 | 1 | 11 | 11 | 26 | 2.30 | 1 | 2 | | | | 2.12 | 1 | 1 | | | | | | | | | | | | | | | | | | | 24.96 | 11 | 23 | 565 | 61.9 | 8.34 |
| Q6UXV1 | 26.24 | 1 | 5 | 5 | 8 | | | | | | | | | | | | | | | | | | | | | | | | | | | | 26.24 | 5 | 8 | 221 | 24.8 | 8.79 |
| Q6UY27 | 76.11 | 1 | 8 | 8 | 25 | 20.35 | 2 | 2 | 8.85 | 1 | 1 | 52.21 | 4 | 5 | | | | | | | | | | | | | | | | | | | 67.26 | 7 | 17 | 113 | 13.0 | 7.25 |
| Q6V702 | 26.61 | 1 | 7 | 7 | 11 | | | | | | | | | | 6.44 | 2 | 2 | 8.58 | 2 | 2 | | | | | | | | | | | | | 21.03 | 6 | 7 | 233 | 26.9 | 5.44 |
| Q6W4X9 | 15.99 | 1 | 30 | 30 | 74 | 3.65 | 6 | 6 | 1.52 | 4 | 5 | 8.00 | 15 | 19 | | | | 0.45 | 1 | 1 | | | | | | | 1.72 | 2 | 2 | | | | 10.78 | 21 | 41 | 2439 | 256.9 | 7.39 |
| Q6X4U4 | 12.14 | 1 | 2 | 2 | 4 | | | | | | | | | | | | | | | | | | | | | | | | | | | | 12.14 | 2 | 4 | 206 | 23.3 | 9.74 |
| Q6X784 | 31.07 | 1 | 10 | 10 | 46 | 18.64 | 6 | 7 | 10.65 | 4 | 5 | 8.58 | 3 | 3 | 2.37 | 1 | 1 | 3.85 | 1 | 1 | | | | | | | | | | | | | 30.18 | 9 | 29 | 338 | 38.6 | 7.78 |
| O95263 | 2.26 | 2 | 1 | 2 | 10 | | | | | | | | | | | | | | | | | | | | | | 0.68 | 1 | 1 | | | | 2.26 | 2 | 9 | 885 | 98.9 | 6.83 |
| Q6XZB0 | 26.96 | 1 | 13 | 13 | 43 | 4.78 | 2 | 2 | 8.04 | 4 | 4 | 6.09 | 3 | 3 | | | | | | | | | | | | | | | | | | | 26.96 | 13 | 34 | 460 | 53.0 | 9.03 |
| Q6Y7W6 | 2.54 | 1 | 2 | 2 | 4 | | | | | | | | | | | | | | | | | | | | | | | | | | | | 2.54 | 2 | 4 | 1299 | 150.0 | 5.54 |
| Q6YN16 | 33.73 | 1 | 11 | 11 | 23 | 4.31 | 1 | 1 | 4.31 | 1 | 2 | 4.31 | 1 | 1 | | | | | | | | | | | | | | | | | | | 33.73 | 11 | 19 | 418 | 45.4 | 7.99 |
| Q6ZMP0 | 11.89 | 1 | 9 | 9 | 19 | | | | | | | 2.55 | 2 | 2 | | | | | | | | | | | | | | | | | | | 11.89 | 9 | 17 | 1018 | 112.4 | 7.65 |
| Q6ZMW3 | 1.48 | 2 | 3 | 3 | 7 | 0.00 | | | | | | | | | | | | | | | | | | | | | | | | | | | 1.48 | 3 | 7 | 1958 | 217.8 | 7.44 |
| Q6ZN84 | 50.61 | 1 | 37 | 37 | 59 | | | | | | | | | | 11.04 | 6 | 8 | 36.35 | 18 | 19 | | | | | | | 0.00 | | | | | | 33.74 | 24 | 32 | 652 | 76.0 | 9.14 |
| Q6ZNM6 | 72.39 | 1 | 11 | 11 | 40 | | | | | | | | | | 30.60 | 4 | 7 | 35.07 | 5 | 7 | | | | | | | | | | | | | 63.43 | 9 | 26 | 134 | 15.4 | 9.16 |
| Q6ZNW5 | 15.32 | 1 | 3 | 3 | 5 | | | | | | | | | | | | | | | | | | | | | | | | | | | | 15.32 | 3 | 5 | 385 | 42.3 | 6.47 |
| Q6ZQQ6 | 1.88 | 2 | 4 | 6 | 9 | 0.00 | | | | | | | | | | | | | | | | | | | | | | | | | | | 1.88 | 6 | 9 | 2873 | 333.0 | 7.28 |
| Q6ZQR2 | 56.88 | 1 | 20 | 21 | 58 | 6.88 | 2 | 2 | 7.19 | 2 | 2 | | | | 23.13 | 7 | 7 | 27.50 | 10 | 12 | | | | 14.37 | 5 | 5 | 4.06 | 1 | 1 | 19.06 | 5 | 6 | 35.94 | 11 | 21 | 320 | 36.5 | 10.26 |
| Q6ZR08 | 37.55 | 2 | 99 | 107 | 241 | 1.00 | 3 | 3 | 0.61 | 2 | 2 | 0.71 | 2 | 2 | 4.08 | 11 | 12 | 0.97 | 1 | 1 | 0.39 | 1 | 1 | | | | | | | | | | 36.48 | 104 | 220 | 3092 | 356.7 | 6.19 |
| Q6ZRH7 | 5.78 | 1 | 4 | 4 | 6 | 3.19 | 1 | 1 | | | | | | | | | | | | | | | | | | | | | | | | | 2.59 | 3 | 5 | 1159 | 132.9 | 6.37 |
| Q6ZS30 | 1.26 | 1 | 3 | 3 | 4 | | | | | | | | | | | | | | | | | | | | | | | | | | | | 1.26 | 3 | 4 | 2694 | 307.0 | 6.44 |
| Q6ZT12 | 1.38 | 1 | 2 | 2 | 2 | | | | | | | | | | | | | | | | | | | | | | | | | | | | 1.38 | 2 | 2 | 1888 | 212.3 | 6.10 |

| | | | | | | | | | | | | | | | | | | | | | | | | | | | | | | | | | | | | | | |
|---|---|---|---|---|---|---|---|---|---|---|---|---|---|---|---|---|---|---|---|---|---|---|---|---|---|---|---|---|---|---|---|---|---|---|---|---|---|---|
| Q6ZTR5 | 25.18 | 1 | 68 | 69 | 107 | 0.74 | 2 | 2 | | | | | | | | | | | | | | | | | | | | | | | | | 25.18 | 68 | 109 | 3102 | 351.7 | 7.40 |
| Q6ZU64 | 21.82 | 1 | 37 | 37 | 72 | | | | | | | | | | | | | 0.99 | 2 | 2 | | | | | | | | | | | | | 21.30 | 36 | 70 | 1925 | 217.1 | 6.51 |
| Q6ZUG5 | 7.52 | 1 | 4 | 4 | 9 | | | | | | | | | | | | | | | | | | | | | | | | | | | | 7.52 | 4 | 9 | 572 | 66.0 | 7.30 |
| Q6ZVM7 | 18.54 | 1 | 7 | 7 | 21 | 2.56 | 1 | 1 | | | | | | | 1.38 | 1 | 1 | | | | | | | 1.38 | 1 | 1 | 1.38 | 1 | 1 | | | | 18.54 | 7 | 17 | 507 | 55.5 | 4.79 |
| Q6ZVS7 | 77.78 | 1 | 9 | 10 | 33 | | | | | | | | | | 50.37 | 4 | 4 | 77.78 | 8 | 14 | | | | | | | | | | | | | 65.19 | 6 | 15 | 135 | 16.2 | 8.43 |
| Q709C8 | 8.34 | 1 | 25 | 25 | 34 | | | | 0.00 | | | 0.32 | 1 | 1 | | | | 0.00 | | | | | | 0.00 | | | | | | | | | 8.02 | 24 | 33 | 3753 | 422.1 | 6.83 |
| Q709F0 | 11.28 | 1 | 7 | 7 | 8 | | | | | | | | | | | | | | | | | | | | | | | | | | | | 11.28 | 7 | 8 | 780 | 87.2 | 8.12 |
| Q70HW3 | 12.77 | 1 | 2 | 2 | 4 | | | | | | | | | | | | | | | | | | | | | | | | | | | | 12.77 | 2 | 4 | 274 | 29.4 | 9.35 |
| Q70Z35 | 1.25 | 1 | 2 | 2 | 2 | 0.00 | | | | | | | | | | | | | | | | | | | | | | | | | | | 1.25 | 2 | 2 | 1606 | 182.5 | 7.44 |
| Q75WM6 | 17.65 | 1 | 4 | 4 | 10 | | | | 4.31 | 1 | 1 | | | | | | | | | | | | | | | | | | | | | | 17.65 | 4 | 9 | 255 | 28.1 | 11.78 |
| Q76KD6 | 8.46 | 1 | 4 | 4 | 5 | | | | | | | | | | | | | | | | | | | | | | | | | | | | 8.46 | 4 | 5 | 591 | 62.4 | 8.19 |
| Q7KYR7 | 7.02 | 3 | 3 | 4 | 7 | | | | | | | | | | | | | | | | | | | | | | | | | | | | 7.02 | 4 | 7 | 527 | 59.6 | 6.48 |
| Q7KZF4 | 43.19 | 1 | 36 | 36 | 117 | | | | | | | | | | 1.43 | 1 | 1 | 1.21 | 1 | 1 | | | | 6.26 | 5 | 5 | 3.52 | 3 | 3 | 13.19 | 10 | 14 | 38.35 | 30 | 93 | 910 | 101.9 | 7.17 |
| Q7L1Q6 | 52.03 | 2 | 18 | 18 | 38 | | | | | | | | | | | | | | | | | | | 1.91 | 1 | 1 | | | | 1.91 | 1 | 1 | 50.12 | 17 | 36 | 419 | 48.0 | 5.92 |
| Q7L266 | 64.29 | 1 | 14 | 14 | 98 | 58.44 | 11 | 13 | 56.17 | 11 | 15 | 40.58 | 8 | 10 | 35.39 | 7 | 8 | 27.60 | 5 | 5 | | | | 4.87 | 1 | 1 | 30.19 | 6 | 7 | 32.14 | 6 | 6 | 61.36 | 13 | 33 | 308 | 32.0 | 6.24 |
| Q7L2E3 | 8.21 | 1 | 7 | 7 | 10 | | | | | | | | | | | | | | | | | | | | | | | | | | | | 8.21 | 7 | 10 | 1194 | 133.9 | 8.78 |
| Q7L2H7 | 42.25 | 1 | 12 | 12 | 24 | | | | | | | | | | | | | | | | | | | | | | | | | | | | 42.25 | 12 | 24 | 374 | 42.5 | 5.63 |
| Q7L3T8 | 12.84 | 1 | 4 | 4 | 4 | | | | | | | | | | | | | | | | | | | | | | | | | | | | 12.84 | 4 | 4 | 475 | 53.2 | 8.10 |
| Q7L523 | 11.50 | 2 | 3 | 3 | 4 | | | | | | | | | | | | | | | | | | | | | | | | | | | | 11.50 | 3 | 4 | 313 | 36.5 | 7.72 |
| Q7L576 | 16.44 | 1 | 8 | 19 | 31 | 0.96 | 1 | 1 | | | | | | | | | | | | | | | | | | | | | | | | | 16.44 | 19 | 30 | 1253 | 145.1 | 6.90 |
| Q7L590 | 1.49 | 2 | 2 | 2 | 3 | | | | | | | | | | | | | | | | | | | | | | | | | | | | 1.49 | 2 | 3 | 875 | 98.1 | 8.75 |
| Q7L5D6 | 32.42 | 1 | 8 | 8 | 12 | | | | | | | | | | | | | | | | | | | | | | | | | | | | 32.42 | 8 | 12 | 327 | 36.5 | 5.41 |
| Q7L5N1 | 44.34 | 1 | 10 | 10 | 16 | | | | | | | | | | 3.36 | 1 | 1 | | | | | | | | | | | | | | | | 44.34 | 10 | 15 | 327 | 36.1 | 5.73 |
| Q7L5N7 | 5.33 | 1 | 3 | 3 | 4 | | | | | | | | | | | | | | | | | | | | | | | | | | | | 5.33 | 3 | 4 | 544 | 60.2 | 6.55 |
| Q7L5Y1 | 12.87 | 1 | 5 | 5 | 10 | | | | | | | | | | | | | | | | | | | | | | | | | | | | 12.87 | 5 | 10 | 443 | 49.8 | 6.48 |
| Q7L5Y9 | 5.05 | 1 | 2 | 2 | 4 | | | | | | | | | | | | | | | | | | | | | | | | | | | | 5.05 | 2 | 4 | 396 | 45.3 | 8.69 |
| Q7L7V1 | 3.90 | 1 | 2 | 2 | 4 | 1.62 | 1 | 1 | | | | | | | | | | | | | | | | | | | | | | | | | 3.90 | 2 | 3 | 743 | 84.4 | 4.97 |
| Q7L8S5 | 20.14 | 1 | 4 | 4 | 5 | | | | | | | | | | | | | | | | | | | | | | | | | | | | 20.14 | 4 | 5 | 288 | 33.3 | 6.74 |
| Q7L9L4 | 8.33 | 2 | 2 | 2 | 3 | | | | | | | | | | | | | | | | | | | | | | | | | | | | 8.33 | 2 | 3 | 216 | 25.1 | 6.73 |
| Q7LBR1 | 16.58 | 1 | 5 | 5 | 6 | | | | | | | | | | 4.02 | 1 | 1 | | | | | | | | | | | | | | | | 16.58 | 4 | 5 | 199 | 22.1 | 8.10 |
| P12532 | 5.28 | 2 | 1 | 2 | 9 | 1.44 | 1 | 1 | 1.44 | 1 | 2 | | | | | | | | | | | | | | | | | | | | | | 5.28 | 2 | 6 | 417 | 47.0 | 8.34 |
| Q7LG56 | 13.96 | 1 | 4 | 4 | 8 | | | | | | | | | | | | | | | | | | | | | | | | | | | | 13.96 | 4 | 8 | 351 | 40.7 | 4.97 |
| Q7RTN6 | 12.53 | 1 | 4 | 4 | 10 | | | | | | | | | | | | | | | | | | | | | | | | | | | | 12.53 | 4 | 10 | 431 | 48.3 | 6.48 |
| Q7RTX7 | 6.57 | 1 | 3 | 3 | 3 | | | | 2.33 | 1 | 1 | | | | | | | | | | | | | | | | | | | | | | 4.24 | 2 | 2 | 472 | 54.1 | 5.26 |
| Q7RTZ1 | 9.04 | 1 | 4 | 4 | 5 | | | | | | | | | | | | | | | | | | | | | | | | | | | | 9.04 | 4 | 5 | 564 | 62.6 | 7.43 |
| Q7Z2H8 | 6.93 | 1 | 2 | 2 | 3 | 3.78 | 1 | 1 | | | | | | | | | | | | | | | | | | | | | | | | | 3.15 | 1 | 2 | 476 | 53.0 | 6.99 |
| Q7Z2K6 | 5.75 | 1 | 5 | 5 | 8 | 0.00 | | | | | | 0.00 | | | | | | | | | | | | | | | | | | | | | 5.75 | 5 | 8 | 904 | 100.2 | 7.52 |
| Q7Z2W7 | 5.89 | 1 | 6 | 6 | 8 | | | | | | | | | | | | | | | | | | | | | | | | | | | | 5.89 | 6 | 8 | 1104 | 127.6 | 7.24 |
| Q7Z2Z2 | 6.16 | 1 | 5 | 6 | 6 | | | | | | | | | | | | | | | | | | | | | | | | | | | | 6.16 | 6 | 6 | 1120 | 125.3 | 5.91 |
| Q7Z304 | 10.35 | 1 | 6 | 6 | 9 | | | | | | | | | | | | | | | | | | | | | | | | | | | | 10.35 | 6 | 9 | 686 | 77.5 | 5.16 |
| Q7Z3B4 | 49.11 | 1 | 20 | 20 | 53 | 4.34 | 2 | 2 | 23.67 | 9 | 9 | 8.48 | 4 | 4 | | | | | | | | | | | | | | | | | | | 48.72 | 19 | 38 | 507 | 55.4 | 7.02 |
| Q7Z3C6 | 7.99 | 1 | 5 | 5 | 9 | | | | 2.15 | 1 | 1 | | | | | | | | | | | | | | | | | | | | | | 7.99 | 5 | 8 | 839 | 94.4 | 6.67 |
| Q7Z3S9 | 13.56 | 2 | 2 | 2 | 3 | | | | | | | | | | | | | | | | | | | | | | | | | | | | 13.56 | 2 | 3 | 236 | 25.8 | 5.71 |
| Q7Z3U7 | 7.92 | 1 | 13 | 13 | 20 | 1.28 | 2 | 2 | | | | | | | | | | | | | | | | | | | | | | | | | 7.22 | 12 | 18 | 1717 | 190.2 | 6.06 |
| Q99719 | 4.88 | 1 | 1 | 2 | 9 | 2.44 | 1 | 1 | 2.44 | 1 | 1 | | | | | | | 2.44 | 1 | 1 | | | | | | | | | | | | | 4.88 | 2 | 6 | 369 | 42.7 | 6.67 |
| Q7Z406 | 34.99 | 7 | 52 | 63 | 180 | 1.30 | 2 | 3 | 0.55 | 1 | 1 | 3.81 | 6 | 6 | 1.05 | 2 | 2 | 0.50 | 1 | 1 | | | | 0.50 | 1 | 1 | | | | 1.75 | 4 | 4 | 34.84 | 60 | 162 | 1995 | 227.7 | 5.60 |
| Q4KWH8 | 0.47 | 1 | 1 | 1 | 9 | 0.47 | 1 | 2 | 0.47 | 1 | 1 | 0.47 | 1 | 2 | | | | 0.47 | 1 | 1 | | | | | | | 0.47 | 1 | 2 | 0.47 | 1 | 1 | | | | 1693 | 189.1 | 7.74 |
| Q7Z4H3 | 63.24 | 1 | 9 | 9 | 33 | 63.24 | 9 | 11 | 51.96 | 8 | 8 | 13.24 | 2 | 2 | 12.25 | 2 | 2 | | | | | | | 5.39 | 1 | 1 | | | | | | | 46.08 | 7 | 9 | 204 | 23.4 | 5.49 |
| Q7Z4L5 | 15.20 | 1 | 14 | 15 | 20 | | | | | | | | | | | | | | | | | | | | | | | | | | | | 15.20 | 15 | 20 | 1316 | 150.8 | 6.96 |
| Q7Z4S6 | 5.32 | 2 | 6 | 6 | 9 | | | | | | | | | | | | | | | | | | | | | | | | | | | | 5.32 | 6 | 9 | 1674 | 187.1 | 6.42 |
| Q7Z4T9 | 50.08 | 1 | 26 | 26 | 87 | | | | | | | | | | 19.57 | 10 | 11 | 27.20 | 14 | 16 | | | | | | | | | | | | | 39.14 | 19 | 60 | 603 | 71.0 | 7.74 |
| Q7Z4W1 | 61.48 | 1 | 15 | 15 | 95 | 27.87 | 7 | 8 | 22.13 | 6 | 6 | 29.10 | 7 | 9 | 20.49 | 5 | 7 | 4.92 | 1 | 1 | | | | 3.28 | 1 | 1 | | | | 4.10 | 1 | 1 | 61.48 | 14 | 62 | 244 | 25.9 | 8.10 |
| Q7Z4W2 | 31.76 | 1 | 4 | 6 | 16 | 30.41 | 5 | 7 | 8.11 | 1 | 2 | | | | | | | | | | | | | | | | | | | | | | 30.41 | 4 | 7 | 148 | 16.6 | 7.84 |
| Q7Z5G4 | 27.74 | 2 | 4 | 4 | 4 | | | | | | | | | | | | | | | | | | | | | | | | | | | | 27.74 | 4 | 4 | 137 | 15.8 | 7.05 |
| Q7Z5J8 | 11.16 | 1 | 13 | 13 | 17 | | | | | | | | | | | | | | | | | | | | | | | | | | | | 11.16 | 13 | 17 | 1434 | 161.9 | 8.10 |
| Q7Z5L4 | 50.30 | 1 | 7 | 7 | 27 | 23.95 | 3 | 3 | 15.57 | 2 | 3 | 8.98 | 1 | 1 | | | | | | | | | | 8.98 | 1 | 2 | | | | 8.98 | 1 | 1 | 50.30 | 7 | 17 | 167 | 19.2 | 6.96 |
| Q7Z5V6 | 81.18 | 1 | 27 | 27 | 69 | 3.53 | 1 | 1 | 5.65 | 2 | 2 | | | | 16.47 | 6 | 6 | 57.88 | 17 | 21 | | | | | | | | | | | | | 72.24 | 21 | 39 | 425 | 47.3 | 8.54 |
| Q7Z6K5 | 9.73 | 1 | 2 | 2 | 3 | | | | | | | | | | | | | | | | | | | | | | | | | 4.42 | 1 | 1 | 5.31 | 1 | 2 | 226 | 24.9 | 5.83 |
| Q7Z6L1 | 3.69 | 1 | 2 | 2 | 2 | | | | | | | | | | | | | | | | | | | | | | | | | | | | 3.69 | 2 | 2 | 1165 | 129.6 | 6.24 |
| Q7Z6W1 | 26.37 | 1 | 3 | 3 | 6 | | | | | | | | | | | | | | | | | | | | | | | | | | | | 26.37 | 3 | 6 | 182 | 20.1 | 8.68 |
| Q7Z6Z7 | 10.43 | 1 | 38 | 38 | 97 | | | | | | | 0.25 | 1 | 1 | 0.91 | 2 | 2 | | | | | | | | | | | | | | | | 10.43 | 38 | 94 | 4374 | 481.6 | 5.22 |
| Q7Z745 | 32.74 | 1 | 45 | 45 | 73 | 0.82 | 1 | 1 | | | | | | | | | | | | | | | | | | | | | | | | | 32.74 | 45 | 72 | 1585 | 180.7 | 6.28 |
| Q7Z7B7 | 28.42 | 1 | 2 | 2 | 3 | | | | | | | | | | | | | | | | | | | | | | | | | | | | 28.42 | 2 | 3 | 95 | 10.6 | 9.60 |
| Q7Z7H3 | 28.17 | 1 | 10 | 10 | 14 | | | | | | | | | | 4.39 | 2 | 2 | | | | | | | | | | | | | | | | 25.84 | 9 | 12 | 387 | 43.9 | 5.41 |
| Q7Z7H5 | 22.91 | 1 | 6 | 7 | 33 | 16.74 | 5 | 6 | 7.93 | 2 | 2 | 20.26 | 6 | 7 | 4.85 | 1 | 1 | | | | | | | | | | | | | | | | 22.47 | 6 | 17 | 227 | 25.9 | 8.28 |
| Q86SX6 | 31.85 | 1 | 4 | 4 | 5 | | | | 12.74 | 2 | 2 | | | | | | | | | | | | | | | | | | | | | | 28.03 | 3 | 3 | 157 | 16.6 | 6.79 |
| Q86SZ2 | 43.04 | 1 | 6 | 6 | 8 | | | | | | | 0.00 | | | | | | | | | | | | | | | | | | | | | 43.04 | 6 | 8 | 158 | 18.0 | 8.68 |
| Q86T03 | 11.19 | 1 | 3 | 3 | 4 | | | | | | | | | | | | | | | | | | | | | | | | | | | | 11.19 | 3 | 4 | 277 | 29.4 | 8.91 |
| Q86TM6 | 1.62 | 1 | 2 | 2 | 4 | | | | | | | | | | | | | | | | | | | | | | | | | | | | 1.62 | 2 | 4 | 617 | 67.6 | 6.95 |
| Q86TX2 | 5.23 | 2 | 2 | 2 | 2 | | | | | | | | | | | | | | | | | | | | | | | | | | | | 5.23 | 2 | 2 | 421 | 46.2 | 7.34 |
| Q86U28 | 12.99 | 1 | 2 | 2 | 2 | | | | | | | 7.79 | 1 | 1 | | | | | | | | | | | | | | | | | | | 5.19 | 1 | 1 | 154 | 16.5 | 5.25 |
| Q86UB9 | 4.15 | 1 | 2 | 2 | 2 | | | | | | | | | | | | | | | | | | | | | | | | | | | | 4.15 | 2 | 2 | 458 | 52.3 | 9.45 |
| Q86UC2 | 44.29 | 1 | 23 | 23 | 89 | 7.32 | 3 | 5 | 7.32 | 3 | 4 | | | | 22.14 | 12 | 12 | 25.54 | 14 | 20 | | | | | | | | | | 1.61 | 1 | 1 | 31.43 | 13 | 43 | 560 | 63.6 | 5.69 |
| Q86UE4 | 4.64 | 1 | 2 | 2 | 2 | | | | | | | | | | | | | | | | | | | | | | | | | | | | 4.64 | 2 | 2 | 582 | 63.8 | 9.32 |
| Q86UG4 | 11.68 | 1 | 7 | 7 | 18 | 1.53 | 1 | 1 | | | | 1.53 | 1 | 1 | | | | | | | | | | | | | | | | | | | 11.68 | 7 | 16 | 719 | 79.2 | 8.59 |
| Q86UK7 | 2.32 | 1 | 2 | 2 | 2 | | | | | | | | | | | | | | | | | | | | | | | | | | | | 2.32 | 2 | 2 | 904 | 98.6 | 8.40 |
| Q86UN6 | 16.75 | 1 | 4 | 4 | 8 | | | | 7.11 | 1 | 1 | | | | 6.09 | 1 | 1 | 6.09 | 1 | 1 | | | | | | | 6.09 | 1 | 1 | 6.09 | 1 | 1 | 16.75 | 3 | 7 | 197 | 22.8 | 6.80 |
| Q86UP2 | 24.83 | 1 | 30 | 30 | 52 | 5.75 | 6 | 6 | 1.11 | 1 | 2 | 4.79 | 5 | 5 | 1.11 | 1 | 2 | 0.00 | | | | | | 1.11 | 1 | 1 | 0.88 | 1 | 1 | 1.11 | 1 | 1 | 24.61 | 28 | 34 | 1357 | 156.2 | 5.64 |
| Q86UT6 | 7.28 | 1 | 6 | 6 | 7 | | | | | | | | | | | | | | | | | | | | | | | | | | | | 7.28 | 6 | 7 | 975 | 107.5 | 7.37 |
| Q86UW7 | 1.54 | 2 | 2 | 2 | 2 | | | | | | | | | | | | | | | | | | | | | | | | | | | | 1.54 | 2 | 2 | 1296 | 147.6 | 6.19 |
| Q86UX2 | 7.75 | 1 | 6 | 6 | 7 | | | | | | | | | | | | | | | | | | | | | | | | | | | | 7.75 | 6 | 7 | 942 | 104.5 | 8.28 |
| Q86V21 | 11.90 | 1 | 8 | 8 | 12 | | | | | | | | | | | | | | | | | | | | | | | | | | | | 11.90 | 8 | 12 | 672 | 75.1 | 6.24 |
| Q86VD7 | 12.26 | 1 | 2 | 2 | 2 | | | | | | | | | | | | | | | | | | | | | | | | | | | | 12.26 | 2 | 2 | 318 | 35.4 | 10.07 |
| Q86VP6 | 49.76 | 1 | 52 | 55 | 225 | 26.91 | 24 | 30 | 20.00 | 18 | 19 | 18.54 | 18 | 19 | 3.41 | 3 | 3 | | | | | | | | | | | | | | | | 46.99 | 49 | 154 | 1230 | 136.3 | 5.78 |
| Q86VQ3 | 26.40 | 1 | 10 | 10 | 26 | 15.73 | 5 | 5 | 6.51 | 2 | 2 | 9.76 | 2 | 2 | 4.88 | 1 | 1 | 10.85 | 3 | 3 | | | | 4.88 | 1 | 1 | 3.25 | 1 | 1 | | | | 20.98 | 7 | 11 | 553 | 60.4 | 4.87 |
| Q86VQ6 | 13.22 | 1 | 4 | 4 | 5 | | | | | | | | | | | | | | | | | | | | | | | | | | | | 13.22 | 4 | 5 | 643 | 70.6 | 7.87 |
| Q86VS3 | 7.89 | 1 | 5 | 5 | 5 | | | | | | | | | | | | | | | | | | | | | | | | | | | | 7.89 | 5 | 5 | 1027 | 117.3 | 9.10 |
| Q86VY4 | 6.00 | 1 | 2 | 2 | 3 | | | | | | | | | | | | | | | | | | | | | | | | | | | | 6.00 | 2 | 3 | 417 | 45.1 | 9.54 |
| Q86W54 | 16.59 | 1 | 3 | 3 | 4 | | | | | | | | | | | | | | | | | | | | | | | | | | | | 16.59 | 3 | 4 | 205 | 23.6 | 7.99 |
| Q86WA6 | 11.34 | 1 | 2 | 2 | 2 | | | | | | | | | | | | | | | | | | | | | | | | | | | | 11.34 | 2 | 2 | 291 | 32.5 | 9.14 |
| Q86WQ0 | 28.06 | 1 | 2 | 2 | 3 | | | | | | | | | | | | | | | | | | | | | | | | | | | | 28.06 | 2 | 3 | 139 | 15.9 | 6.16 |
| Q86WS5 | 29.31 | 1 | 7 | 7 | 12 | | | | 6.61 | 1 | 2 | | | | | | | | | | | | | | | | | | | | | | 22.70 | 6 | 10 | 348 | 38.6 | 8.62 |
| Q86WT1 | 32.03 | 2 | 17 | 17 | 20 | | | | | | | | | | | | | | | | | | | | | | | | | | | | 32.03 | 17 | 20 | 665 | 76.1 | 5.20 |
| Q86WT6 | 25.40 | 1 | 10 | 11 | 17 | | | | 2.20 | 1 | 1 | | | | | | | | | | | | | | | | | | | | | | 25.40 | 10 | 16 | 500 | 57.4 | 6.48 |
| Q86WZ0 | 2.14 | 1 | 2 | 2 | 2 | | | | | | | | | | | | | | | | | | | | | | | | | | | | 2.14 | 2 | 2 | 1026 | 117.1 | 9.11 |
| Q86X10 | 8.50 | 1 | 11 | 11 | 14 | | | | | | | | | | | | | | | | | | | | | | | | | | | | 8.50 | 11 | 14 | 1494 | 166.7 | 6.79 |
| Q86X29 | 7.86 | 1 | 4 | 4 | 8 | 1.69 | 1 | 1 | 1.69 | 1 | 1 | 1.69 | 1 | 1 | | | | | | | | | | | | | | | | | | | 6.16 | 3 | 3 | 649 | 71.4 | 7.97 |
| Q86X55 | 9.21 | 1 | 5 | 5 | 7 | | | | | | | | | | | | | | | | | | | | | | | | | | | | 9.21 | 5 | 7 | 608 | 65.8 | 6.73 |
| Q86X76 | 35.78 | 1 | 10 | 10 | 25 | 8.56 | 3 | 3 | 5.20 | 2 | 2 | 5.20 | 2 | 2 | 3.06 | 1 | 1 | | | | | | | 2.75 | 1 | 1 | | | | | | | 35.78 | 10 | 16 | 327 | 35.9 | 7.74 |
| Q86XH1 | 31.02 | 1 | 18 | 19 | 29 | | | | | | | | | | 4.26 | 3 | 3 | | | | | | | | | | | | | | | | 31.02 | 19 | 26 | 822 | 95.3 | 9.47 |
| Q86XM0 | 10.03 | 1 | 6 | 6 | 8 | | | | | | | | | | | | | | | | | | | | | | | | | | | | 10.03 | 6 | 8 | 798 | 90.4 | 7.37 |
| Q86XQ3 | 7.79 | 1 | 3 | 3 | 4 | | | | | | | | | | | | | | | | | | | | | | | | | | | | 7.79 | 3 | 4 | 398 | 46.4 | 6.30 |
| O75152 | 3.46 | 1 | 1 | 2 | 9 | | | | | | | | | | | | | | | | | | | | | | | | | | | | 3.46 | 2 | 9 | 810 | 89.1 | 8.37 |
| Q86Y39 | 57.45 | 1 | 5 | 5 | 8 | | | | | | | | | | | | | | | | | | | | | | | | | | | | 57.45 | 5 | 8 | 141 | 14.8 | 8.72 |
| Q86Y56 | 7.95 | 1 | 6 | 6 | 8 | | | | | | | | | | | | | | | | | | | | | | | | | | | | 7.95 | 6 | 8 | 855 | 93.5 | 6.42 |
| Q86Y82 | 42.03 | 1 | 7 | 7 | 30 | 30.43 | 5 | 5 | 21.74 | 3 | 4 | 10.87 | 2 | 2 | | | | | | | | | | | | | | | | | | | 33.33 | 5 | 19 | 276 | 31.6 | 5.59 |
| Q86YB8 | 15.42 | 1 | 5 | 6 | 14 | 8.57 | 3 | 3 | 10.92 | 4 | 4 | 9.42 | 4 | 4 | | | | | | | | | | | | | | | | | | | 7.28 | 3 | 3 | 467 | 53.5 | 7.99 |
| Q86YN1 | 8.40 | 1 | 2 | 2 | 9 | | | | | | | | | | | | | | | | | | | | | | | | | | | | 8.40 | 2 | 9 | 238 | 27.0 | 9.44 |
| Q86YS6 | 35.85 | 6 | 3 | 5 | 20 | 5.19 | 1 | 2 | 5.19 | 1 | 1 | 5.19 | 1 | 1 | 5.19 | 1 | 1 | | | | | | | 5.19 | 1 | 1 | | | | | | | 35.85 | 5 | 14 | 212 | 23.3 | 5.64 |
| Q86YZ3 | 12.56 | 1 | 10 | 10 | 49 | | | | 2.74 | 1 | 1 | | | | 7.89 | 4 | 4 | 3.47 | 2 | 2 | | | | 8.70 | 5 | 5 | 8.35 | 7 | 9 | 6.21 | 3 | 3 | 7.82 | 6 | 25 | 2850 | 282.2 | 10.04 |
| Q8IUA0 | 59.34 | 1 | 17 | 17 | 58 | 28.22 | 5 | 7 | 2.49 | 1 | 1 | 23.24 | 5 | 5 | | | | 29.46 | 6 | 7 | | | | | | | | | | | | | 58.51 | 15 | 38 | 241 | 27.8 | 8.02 |
| Q8IUC4 | 9.77 | 2 | 5 | 5 | 5 | | | | | | | | | | | | | | | | | | | | | | | | | | | | 9.77 | 5 | 5 | 686 | 76.9 | 6.80 |
| Q8IUE6 | 35.38 | 1 | 3 | 4 | 22 | 30.00 | 3 | 5 | 24.62 | 2 | 3 | 6.92 | 1 | 1 | 6.92 | 1 | 1 | 6.92 | 1 | 1 | | | | | | | | | | | | | 30.00 | 3 | 11 | 130 | 14.0 | 10.89 |
| Q8IUH8 | 4.24 | 1 | 2 | 2 | 2 | | | | | | | | | | | | | | | | | | | | | | | | | | | | 4.24 | 2 | 2 | 684 | 74.5 | 6.76 |
| Q8IUI8 | 15.84 | 1 | 5 | 5 | 7 | | | | | | | | | | | | | | | | | | | | | | | | | | | | 15.84 | 5 | 7 | 442 | 49.7 | 5.14 |
| Q8IUR0 | 30.32 | 1 | 4 | 4 | 7 | | | | 0.00 | | | | | | | | | | | | | | | | | | 0.00 | | | 0.00 | | | 30.32 | 4 | 7 | 188 | 20.8 | 9.66 |
| Q8IUR7 | 17.09 | 1 | 9 | 9 | 11 | | | | | | | | | | | | | | | | | | | | | | | | | | | | 17.09 | 9 | 11 | 673 | 75.5 | 6.73 |
| Q8IVD9 | 6.93 | 1 | 2 | 2 | 3 | | | | | | | | | | | | | | | | | | | | | | | | | | | | 6.93 | 2 | 3 | 361 | 40.8 | 5.25 |
| Q8IVF4 | 42.47 | 1 | 185 | 187 | 415 | 1.21 | 4 | 4 | 0.20 | 1 | 1 | 0.45 | 2 | 2 | 7.20 | 28 | 29 | 3.47 | 14 | 14 | | | | | | | | | | | | | 41.20 | 177 | 365 | 4471 | 514.5 | 5.88 |
| Q8IVH4 | 6.22 | 1 | 2 | 2 | 2 | 3.83 | 1 | 1 | | | | | | | | | | | | | | | | | | | | | | | | | 2.39 | 1 | 1 | 418 | 46.5 | 9.29 |
| Q8IVL8 | 6.68 | 1 | 2 | 2 | 3 | | | | 4.28 | 1 | 1 | | | | | | | | | | | | | | | | | | | | | | 6.68 | 2 | 2 | 374 | 42.5 | 7.06 |
| Q8IVU9 | 26.44 | 1 | 5 | 5 | 6 | | | | | | | | | | | | | | | | | | | | | | | | | | | | 26.44 | 5 | 6 | 208 | 23.9 | 4.45 |
| Q8IW40 | 22.31 | 1 | 4 | 4 | 4 | | | | | | | | | | | | | | | | | | | | | | | | | | | | 22.31 | 4 | 4 | 242 | 27.1 | 5.97 |
| Q8IW45 | 22.48 | 1 | 6 | 6 | 6 | | | | | | | | | | | | | | | | | | | | | | | | | | | | 22.48 | 6 | 6 | 347 | 36.6 | 8.06 |
| Q8IWA4 | 6.21 | 1 | 3 | 3 | 3 | | | | | | | | | | | | | | | | | | | | | | | | | | | | 6.21 | 3 | 3 | 741 | 84.0 | 6.25 |
| Q8IWB7 | 5.61 | 1 | 2 | 2 | 3 | | | | 0.00 | | | | | | | | | | | | | | | | | | | | | | | | 5.61 | 2 | 3 | 410 | 46.3 | 7.33 |
| Q8IWE2 | 12.79 | 1 | 5 | 5 | 9 | | | | | | | | | | | | | | | | | | | 6.75 | 3 | 3 | | | | 8.70 | 3 | 3 | 6.04 | 2 | 3 | 563 | 60.7 | 4.68 |
| Q8IWF9 | 10.65 | 1 | 3 | 3 | 4 | | | | | | | | | | | | | | | | | | | | | | | | | | | | 10.65 | 3 | 4 | 413 | 48.8 | 5.45 |
| Q8IWG1 | 33.33 | 1 | 32 | 32 | 68 | | | | | | | | | | 3.70 | 3 | 3 | 10.10 | 8 | 8 | | | | | | | | | | 1.35 | 1 | 1 | 33.11 | 30 | 56 | 891 | 102.9 | 5.74 |
| Q8IWT0 | 16.17 | 1 | 2 | 2 | 3 | | | | | | | | | | | | | | | | | | | | | | | | | | | | 16.17 | 2 | 3 | 167 | 19.5 | 4.49 |
| Q8IWV7 | 3.54 | 1 | 6 | 6 | 7 | | | | | | | | | | | | | | | | | | | | | | | | | | | | 3.54 | 6 | 7 | 1749 | 200.1 | 6.01 |
| Q8IWZ5 | 7.75 | 1 | 4 | 4 | 5 | | | | | | | 1.80 | 1 | 1 | | | | | | | | | | | | | | | | | | | 5.95 | 3 | 4 | 723 | 82.7 | 7.93 |
| Q8IWZ6 | 14.13 | 1 | 8 | 8 | 8 | | | | | | | | | | | | | | | | | | | | | | | | | | | | 14.13 | 8 | 8 | 715 | 80.3 | 6.05 |
| Q8IX01 | 4.71 | 1 | 3 | 3 | 3 | | | | | | | | | | | | | | | | | | | | | | | | | | | | 4.71 | 3 | 3 | 1082 | 120.1 | 7.28 |
| Q8IXA5 | 33.49 | 1 | 7 | 7 | 41 | 28.37 | 5 | 10 | 15.81 | 3 | 5 | 21.40 | 4 | 5 | 21.40 | 4 | 5 | 15.81 | 3 | 3 | | | | | | | | | | 3.72 | 1 | 1 | 33.49 | 7 | 12 | 215 | 23.4 | 7.94 |
| Q8IXB1 | 12.23 | 1 | 8 | 8 | 10 | 1.26 | 1 | 1 | | | | | | | | | | | | | | | | | | | | | | | | | 12.23 | 8 | 9 | 793 | 91.0 | 7.18 |
| Q8IXI2 | 6.15 | 1 | 3 | 3 | 3 | | | | | | | | | | | | | | | | | | | | | | | | | | | | 6.15 | 3 | 3 | 618 | 70.7 | 6.27 |
| Q8IXK2 | 2.75 | 1 | 2 | 2 | 3 | | | | | | | | | | | | | | | | | | | | | | | | | | | | 2.75 | 2 | 2 | 581 | 66.9 | 6.80 |
| Q8IXM7 | 58.76 | 1 | 12 | 12 | 31 | | | | | | | | | | 27.74 | 6 | 6 | 41.61 | 8 | 8 | | | | 8.39 | 1 | 1 | | | | | | | 42.70 | 8 | 16 | 274 | 31.0 | 9.54 |
| Q8IXQ6 | 2.93 | 1 | 2 | 2 | 2 | | | | 0.00 | | | 0.00 | | | | | | | | | | | | | | | | | | | | | 2.93 | 2 | 2 | 854 | 96.3 | 7.91 |
| Q8IXS2 | 47.52 | 1 | 27 | 27 | 58 | | | | | | | | | | 14.88 | 7 | 8 | 30.99 | 15 | 19 | | | | | | | 1.24 | 1 | 1 | | | | 43.39 | 24 | 30 | 484 | 57.3 | 7.43 |
| Q8IXU6 | 9.63 | 1 | 2 | 2 | 3 | | | | | | | | | | | | | | | | | | | | | | | | | | | | 9.63 | 2 | 3 | 374 | 41.2 | 6.21 |
| Q8IXW0 | 7.57 | 1 | 3 | 3 | 3 | | | | | | | | | | | | | | | | | | | | | | | | | | | | 7.57 | 3 | 3 | 634 | 70.3 | 8.84 |
| P16188 | 21.64 | 12 | 1 | 6 | 8 | 7.40 | 2 | 2 | | | | 12.33 | 3 | 3 | | | | | | | | | | | | | | | | | | | 12.05 | 3 | 3 | 365 | 40.9 | 6.10 |
| Q8IXY8 | 51.77 | 1 | 15 | 15 | 32 | | | | 4.82 | 1 | 1 | | | | 4.50 | 1 | 1 | 18.65 | 5 | 6 | | | | | | | | | | | | | 47.27 | 13 | 24 | 311 | 35.2 | 7.02 |
| Q8IY17 | 14.49 | 1 | 15 | 15 | 26 | 0.88 | 1 | 1 | 1.83 | 2 | 2 | | | | | | | | | | | | | | | | | | | | | | 14.49 | 15 | 23 | 1366 | 149.9 | 7.81 |
| Q8IY31 | 40.91 | 1 | 4 | 4 | 6 | 10.61 | 1 | 1 | | | | | | | | | | | | | | | | | | | | | | | | | 40.91 | 4 | 5 | 132 | 15.3 | 5.12 |
| Q8IY45 | 13.95 | 1 | 2 | 2 | 3 | | | | | | | | | | | | | | | | | | | | | | | | | | | | 13.95 | 2 | 3 | 258 | 28.4 | 7.85 |
| Q9C037 | 2.20 | 1 | 1 | 1 | 8 | 2.20 | 1 | 1 | 2.20 | 1 | 1 | 2.20 | 1 | 1 | | | | | | | | | | | | | | | | 2.20 | 1 | 1 | 2.20 | 1 | 4 | 500 | 57.4 | 8.10 |
| Q8IY82 | 45.31 | 1 | 41 | 41 | 77 | | | | | | | | | | 15.79 | 13 | 14 | 19.68 | 14 | 15 | | | | | | | | | | | | | 34.10 | 29 | 48 | 874 | 103.4 | 5.67 |
| Q8IYD1 | 28.34 | 1 | 2 | 14 | 33 | | | | 0.00 | | | 1.75 | 1 | 1 | | | | | | | | | | | | | 1.59 | 1 | 1 | 12.26 | 6 | 6 | 18.95 | 10 | 25 | 628 | 68.8 | 5.43 |
| Q8IYE0 | 30.99 | 1 | 29 | 29 | 51 | | | | | | | 0.00 | | | 3.04 | 3 | 3 | 6.81 | 7 | 7 | | | | | | | 0.00 | | | | | | 28.06 | 26 | 41 | 955 | 112.7 | 8.48 |
| Q8IYF3 | 28.30 | 1 | 22 | 22 | 29 | | | | | | | | | | | | | | | | | | | | | | | | | | | | 28.30 | 22 | 29 | 940 | 107.8 | 5.16 |
| Q8IYI6 | 4.69 | 1 | 4 | 4 | 4 | | | | | | | | | | | | | | | | | | | | | | | | | 1.79 | 1 | 1 | 4.69 | 3 | 3 | 725 | 81.7 | 5.49 |
| Q8IYJ3 | 16.37 | 1 | 8 | 8 | 21 | 1.78 | 1 | 1 | | | | 1.96 | 1 | 1 | 5.16 | 2 | 2 | | | | | | | 8.36 | 4 | 4 | | | | 10.85 | 5 | 6 | 6.05 | 3 | 7 | 562 | 61.8 | 5.48 |
| O60635 | 5.39 | 1 | 1 | 1 | 8 | 5.39 | 1 | 1 | | | | 5.39 | 1 | 1 | | | | | | | | | | | | | | | | | | | 5.39 | 1 | 6 | 241 | 26.3 | 5.25 |
| Q8IYK2 | 74.35 | 1 | 48 | 48 | 218 | | | | 8.62 | 4 | 4 | | | | 47.49 | 25 | 32 | 58.52 | 32 | 44 | | | | 17.23 | 8 | 9 | 4.61 | 2 | 2 | 36.67 | 16 | 21 | 61.72 | 31 | 106 | 499 | 56.9 | 9.86 |
| Q9H3S1 | 2.63 | 2 | 1 | 2 | 8 | 0.79 | 1 | 1 | 0.79 | 1 | 1 | 0.79 | 1 | 1 | | | | | | | | | | | | | 0.79 | 1 | 1 | 0.79 | 1 | 1 | 2.63 | 2 | 3 | 761 | 83.5 | 6.84 |
| Q8IYM1 | 30.45 | 1 | 9 | 9 | 26 | 3.35 | 1 | 1 | 11.17 | 3 | 4 | | | | | | | 8.66 | 2 | 5 | | | | | | | | | | | | | 30.45 | 9 | 16 | 358 | 40.7 | 7.11 |
| Q8IYP2 | 14.11 | 1 | 3 | 3 | 3 | | | | | | | | | | | | | | | | | | | | | | | | | | | | 14.11 | 3 | 3 | 241 | 27.1 | 6.84 |
| Q8IYQ7 | 22.48 | 1 | 12 | 12 | 15 | 2.02 | 1 | 1 | 3.77 | 2 | 2 | 1.75 | 1 | 1 | | | | | | | | | | | | | | | | | | | 18.71 | 10 | 11 | 743 | 83.0 | 7.12 |
| Q8IYR0 | 43.25 | 1 | 25 | 25 | 45 | | | | | | | | | | 6.11 | 4 | 4 | 3.54 | 2 | 2 | | | | | | | | | | | | | 41.48 | 22 | 39 | 622 | 71.1 | 6.84 |
| Q8IYS1 | 14.22 | 1 | 4 | 5 | 6 | | | | | | | | | | | | | | | | | | | | | | | | | | | | 14.22 | 5 | 6 | 436 | 47.7 | 5.85 |
| Q8IYS2 | 14.20 | 1 | 6 | 6 | 12 | | | | | | | | | | | | | | | | | | | | | | | | | | | | 14.20 | 6 | 12 | 634 | 69.1 | 8.19 |
| Q8IYS8 | 62.79 | 2 | 9 | 9 | 29 | | | | 9.88 | 1 | 1 | | | | 25.00 | 3 | 5 | 25.58 | 3 | 3 | | | | | | | 18.60 | 2 | 3 | 8.72 | 1 | 1 | 55.81 | 7 | 16 | 172 | 18.1 | 5.33 |
| Q8IYT1 | 14.48 | 2 | 8 | 9 | 29 | 2.53 | 1 | 1 | 5.39 | 3 | 3 | 4.21 | 2 | 2 | 2.53 | 1 | 1 | 2.53 | 1 | 1 | | | | | | | 2.53 | 1 | 1 | | | | 14.48 | 9 | 20 | 594 | 63.1 | 9.64 |
| Q8IYU2 | 3.30 | 1 | 3 | 3 | 3 | | | | | | | | | | | | | | | | | | | | | | | | | | | | 3.30 | 3 | 3 | 909 | 102.3 | 5.92 |
| Q8IYV9 | 30.57 | 1 | 9 | 9 | 14 | 2.00 | 1 | 1 | 5.43 | 2 | 3 | | | | | | | | | | | | | | | | | | | | | | 28.00 | 8 | 10 | 350 | 38.9 | 6.54 |
| Q8IYW2 | 13.15 | 1 | 27 | 27 | 58 | | | | | | | | | | | | | 0.59 | 1 | 1 | | | | | | | | | | | | | 13.15 | 27 | 57 | 2715 | 303.3 | 7.36 |
| Q8IYX1 | 34.23 | 1 | 10 | 10 | 15 | | | | | | | | | | | | | | | | | | | | | | | | | | | | 34.23 | 10 | 15 | 336 | 39.2 | 6.19 |
| Q8IYX3 | 30.68 | 1 | 12 | 12 | 47 | | | | 1.75 | 1 | 1 | | | | 1.75 | 1 | 1 | 1.75 | 1 | 1 | | | | | | | | | | | | | 30.68 | 12 | 44 | 515 | 56.9 | 9.64 |
| Q8IYX7 | 87.34 | 1 | 44 | 44 | 279 | 20.46 | 7 | 8 | 19.83 | 7 | 8 | | | | 60.76 | 20 | 28 | 85.23 | 38 | 59 | | | | | | | 25.74 | 10 | 11 | | | | 73.00 | 30 | 165 | 474 | 54.6 | 8.37 |
| Q8IZ02 | 40.33 | 1 | 13 | 13 | 21 | | | | | | | | | | 9.79 | 3 | 3 | | | | | | | | | | | | | | | | 40.33 | 13 | 18 | 419 | 46.4 | 6.52 |
| Q8IZ16 | 23.79 | 1 | 4 | 4 | 12 | 7.28 | 1 | 1 | 4.37 | 1 | 1 | | | | | | | | | | | | | | | | | | | | | | 23.79 | 4 | 10 | 206 | 23.8 | 10.40 |
| Q8IZ81 | 11.60 | 1 | 3 | 3 | 3 | | | | | | | | | | | | | | | | | | | | | | | | | | | | 11.60 | 3 | 3 | 293 | 34.9 | 8.05 |
| Q8IZ83 | 7.61 | 1 | 4 | 4 | 5 | 1.62 | 1 | 1 | | | | | | | 0.00 | | | | | | | | | | | | | | | | | | 5.99 | 3 | 4 | 802 | 85.1 | 6.79 |
| Q8IZ96 | 24.85 | 1 | 3 | 3 | 11 | 8.88 | 1 | 1 | 8.88 | 1 | 1 | | | | | | | | | | | | | | | | | | | | | | 24.85 | 3 | 9 | 169 | 18.6 | 6.99 |
| Q8IZH2 | 1.23 | 1 | 2 | 2 | 2 | | | | | | | | | | | | | | | | | | | | | | | | | | | | 1.23 | 2 | 2 | 1706 | 194.0 | 7.21 |
| Q8IZP9 | 13.77 | 1 | 12 | 12 | 34 | 4.42 | 3 | 3 | 1.87 | 2 | 2 | 1.87 | 2 | 2 | 1.18 | 1 | 1 | 2.65 | 2 | 3 | | | | | | | 1.18 | 1 | 1 | | | | 13.77 | 12 | 22 | 1017 | 111.5 | 7.55 |
| Q8IZS6 | 47.47 | 1 | 9 | 10 | 30 | 14.65 | 2 | 2 | 18.69 | 3 | 3 | 5.05 | 1 | 1 | | | | 9.09 | 2 | 2 | | | | | | | | | | | | | 46.97 | 9 | 22 | 198 | 23.2 | 9.33 |
| Q8IZT6 | 0.40 | 1 | 3 | 3 | 3 | | | | | | | | | | | | | | | | | | | | | | | | | 0.00 | | | 0.40 | 3 | 3 | 3477 | 409.5 | 10.45 |
| Q8N0U8 | 25.57 | 1 | 5 | 5 | 11 | 5.68 | 1 | 1 | | | | | | | | | | | | | | | | | | | | | | | | | 25.57 | 5 | 10 | 176 | 19.8 | 9.13 |
| Q8N0W3 | 3.87 | 1 | 3 | 3 | 3 | | | | | | | | | | | | | | | | | | | | | | | | | | | | 3.87 | 3 | 3 | 1084 | 117.5 | 6.25 |
| P20936 | 1.62 | 2 | 1 | 2 | 8 | | | | | | | | | | 0.00 | | | 0.57 | 1 | 1 | | | | 0.57 | 1 | 1 | 0.57 | 1 | 1 | 0.00 | | | 1.62 | 2 | 5 | 1047 | 116.3 | 6.54 |
| Q8N0W5 | 20.91 | 1 | 4 | 4 | 9 | | | | | | | | | | | | | | | | | | | | | | | | | | | | 20.91 | 4 | 9 | 287 | 33.3 | 8.21 |
| Q8N0W7 | 20.78 | 2 | 5 | 5 | 14 | 1.96 | 1 | 1 | 1.96 | 1 | 1 | | | | | | | | | | | | | | | | | | | | | | 20.78 | 5 | 12 | 255 | 29.2 | 8.94 |
| Q8N0X2 | 50.55 | 1 | 25 | 26 | 70 | | | | | | | 2.06 | 1 | 1 | 3.33 | 2 | 2 | 12.68 | 7 | 7 | | | | | | | 1.74 | 1 | 1 | | | | 47.54 | 22 | 59 | 631 | 70.8 | 6.33 |
| Q8N0X4 | 8.82 | 1 | 2 | 2 | 2 | | | | 2.94 | 1 | 1 | | | | | | | | | | | | | | | | | | | | | | 5.88 | 1 | 1 | 340 | 37.3 | 8.73 |
| Q8N158 | 5.35 | 1 | 3 | 3 | 3 | | | | 2.07 | 2 | 2 | | | | | | | | | | | | | | | | | | | | | | 3.28 | 1 | 1 | 579 | 62.8 | 8.00 |
| Q8N183 | 34.91 | 1 | 4 | 4 | 4 | | | | | | | | | | | | | | | | | | | | | | | | | | | | 34.91 | 4 | 4 | 169 | 19.8 | 8.97 |
| Q8N1B4 | 2.77 | 1 | 2 | 2 | 2 | | | | | | | | | | | | | | | | | | | | | | | | | | | | 2.77 | 2 | 2 | 723 | 82.2 | 5.99 |
| Q8N1D5 | 64.95 | 1 | 13 | 13 | 38 | 6.19 | 1 | 1 | 9.79 | 2 | 2 | | | | 25.77 | 5 | 5 | 26.80 | 5 | 9 | | | | 6.19 | 1 | 1 | | | | 16.49 | 3 | 3 | 62.37 | 10 | 21 | 194 | 23.1 | 9.73 |
| Q8N1F7 | 64.35 | 1 | 52 | 52 | 175 | 14.90 | 10 | 10 | 29.06 | 19 | 23 | 5.86 | 4 | 4 | | | | | | | | | | | | | | | | | | | 60.20 | 44 | 138 | 819 | 93.4 | 5.72 |
| Q8N1G4 | 9.43 | 1 | 5 | 5 | 8 | | | | | | | | | | 2.57 | 1 | 1 | | | | | | | | | | | | | 2.74 | 2 | 2 | 9.43 | 4 | 5 | 583 | 63.4 | 8.28 |
| Q8N1N4 | 9.42 | 5 | 2 | 6 | 32 | | | | 3.46 | 2 | 2 | 3.46 | 2 | 2 | | | | 3.27 | 2 | 3 | 0.00 | | | 1.35 | 1 | 1 | 5.58 | 4 | 5 | 5.19 | 3 | 4 | 6.73 | 4 | 15 | 520 | 56.8 | 6.02 |
| Q8N1V2 | 60.81 | 1 | 35 | 36 | 210 | 6.61 | 4 | 6 | 6.61 | 4 | 5 | 18.55 | 10 | 13 | 30.65 | 15 | 19 | 24.84 | 12 | 14 | | | | | | | | | | | | | 58.71 | 30 | 153 | 620 | 68.3 | 6.95 |
| Q8N2K0 | 22.11 | 1 | 7 | 7 | 7 | | | | | | | | | | | | | | | | | | | | | | | | | | | | 22.11 | 7 | 7 | 398 | 45.1 | 8.65 |
| Q8N2U9 | 8.86 | 1 | 3 | 3 | 3 | | | | | | | | | | | | | | | | | | | | | | | | | | | | 8.86 | 3 | 3 | 271 | 30.5 | 8.18 |
| Q8N309 | 16.16 | 1 | 6 | 6 | 16 | | | | | | | | | | | | | | | | | | | | | | | | | | | | 16.16 | 6 | 16 | 656 | 73.0 | 5.24 |
| Q8N335 | 67.52 | 1 | 23 | 24 | 97 | 3.42 | 1 | 1 | 2.28 | 1 | 1 | 15.10 | 5 | 5 | 21.65 | 6 | 6 | | | | | | | 15.10 | 5 | 5 | | | | 27.92 | 10 | 11 | 66.10 | 21 | 68 | 351 | 38.4 | 7.02 |
| O00763 | 2.44 | 1 | 1 | 5 | 8 | | | | | | | | | | | | | | | | | | | | | | | | | | | | 2.44 | 5 | 8 | 2458 | 276.4 | 6.49 |
| Q8N3C0 | 3.59 | 1 | 7 | 7 | 7 | | | | | | | | | | | | | | | | | | | | | | | | | | | | 3.59 | 7 | 7 | 2202 | 251.3 | 7.09 |
| Q71UM5 | 38.10 | 1 | 1 | 3 | 8 | | | | | | | | | | | | | | | | | | | | | | | | | | | | 38.10 | 3 | 8 | 84 | 9.5 | 9.45 |
| Q8N3I7 | 6.16 | 1 | 2 | 2 | 2 | | | | | | | | | | | | | | | | | | | | | | | | | | | | 6.16 | 2 | 2 | 341 | 38.7 | 5.60 |
| Q8N3P4 | 3.71 | 1 | 5 | 5 | 7 | | | | | | | | | | | | | | | | | | | | | | | | | | | | 3.71 | 5 | 7 | 1428 | 161.7 | 5.64 |
| Q8N3R9 | 5.48 | 1 | 3 | 3 | 5 | 1.78 | 1 | 1 | | | | | | | | | | | | | | | | | | | | | | | | | 5.48 | 3 | 4 | 675 | 77.2 | 6.14 |
| Q8N427 | 60.54 | 2 | 36 | 36 | 181 | 13.78 | 8 | 8 | 16.16 | 8 | 8 | 4.08 | 3 | 3 | 16.67 | 9 | 10 | 31.12 | 19 | 24 | 1.02 | 1 | 1 | 1.87 | 1 | 1 | 5.44 | 4 | 5 | 11.05 | 6 | 6 | 57.14 | 32 | 115 | 588 | 67.2 | 4.97 |
| Q8N443 | 74.14 | 1 | 25 | 25 | 71 | 2.37 | 1 | 1 | | | | | | | 38.52 | 10 | 15 | 48.55 | 16 | 24 | 2.90 | 1 | 1 | | | | | | | 8.44 | 2 | 2 | 60.69 | 16 | 28 | 379 | 44.0 | 8.76 |
| Q8N4S6 | 33.72 | 1 | 6 | 6 | 10 | | | | | | | | | | | | | | | | | | | | | | | | | | | | 33.72 | 6 | 10 | 261 | 29.7 | 9.80 |
| Q8N474 | 43.63 | 1 | 9 | 10 | 23 | 2.23 | 1 | 1 | 3.18 | 1 | 1 | | | | | | | | | | | | | | | | | | | | | | 43.63 | 10 | 21 | 314 | 35.4 | 8.85 |
| Q8N4E7 | 59.50 | 1 | 12 | 12 | 71 | 42.56 | 6 | 10 | 49.59 | 8 | 19 | 52.48 | 9 | 15 | 6.20 | 1 | 1 | 10.74 | 2 | 2 | | | | | | | | | | | | | 44.21 | 8 | 24 | 242 | 27.5 | 7.27 |
| Q8N4P6 | 25.04 | 1 | 11 | 11 | 21 | | | | 2.15 | 1 | 1 | 2.33 | 1 | 1 | | | | | | | | | | | | | | | | | | | 22.90 | 10 | 19 | 559 | 61.8 | 9.22 |
| Q8N4T8 | 13.08 | 1 | 2 | 2 | 2 | | | | | | | | | | | | | | | | | | | | | | | | | | | | 13.08 | 2 | 2 | 237 | 25.3 | 9.33 |
| Q8N4V1 | 33.59 | 1 | 3 | 3 | 3 | | | | | | | | | | | | | | | | | | | | | | | | | | | | 33.59 | 3 | 3 | 131 | 14.7 | 9.16 |
| Q8N573 | 5.26 | 1 | 4 | 5 | 9 | | | | 1.37 | 1 | 1 | 2.17 | 2 | 2 | | | | | | | | | | | | | 1.72 | 1 | 1 | 1.72 | 1 | 1 | 3.09 | 2 | 4 | 874 | 97.9 | 5.47 |
| Q8N5K1 | 46.67 | 1 | 6 | 6 | 7 | | | | | | | | | | | | | | | | | | | | | | | | | | | | 46.67 | 6 | 7 | 135 | 15.3 | 9.61 |
| P15056 | 4.96 | 2 | 1 | 4 | 8 | | | | | | | | | | | | | | | | | | | | | | | | | | | | 4.96 | 4 | 8 | 766 | 84.4 | 7.53 |
| Q8N5M4 | 19.88 | 3 | 3 | 3 | 4 | | | | 0.00 | | | | | | 0.00 | | | 0.00 | | | | | | | | | | | | | | | 19.88 | 3 | 4 | 171 | 20.0 | 8.92 |
| Q8N5Q1 | 8.57 | 1 | 5 | 5 | 9 | | | | 1.08 | 1 | 1 | | | | | | | | | | | | | | | | | | | | | | 7.48 | 4 | 8 | 922 | 99.9 | 9.39 |
| Q8N5S3 | 11.50 | 1 | 3 | 3 | 3 | | | | | | | | | | | | | | | | | | | | | | | | | | | | 11.50 | 3 | 3 | 287 | 32.1 | 8.97 |
| Q8N6G2 | 6.23 | 1 | 2 | 2 | 2 | | | | | | | | | | | | | 6.23 | 2 | 2 | | | | | | | | | | | | | | | | 289 | 33.6 | 9.35 |
| Q8N6M5 | 8.29 | 1 | 2 | 2 | 3 | | | | | | | | | | | | | | | | | | | | | | | | | | | | 8.29 | 2 | 2 | 410 | 45.7 | 6.18 |
| Q8N6M8 | 15.61 | 2 | 2 | 3 | 4 | | | | | | | | | | | | | | | | | | | | | | | | | | | | 15.61 | 3 | 4 | 205 | 23.7 | 9.61 |
| Q8N6Q1 | 44.10 | 1 | 10 | 10 | 25 | 14.24 | 3 | 3 | | | | | | | 5.90 | 1 | 1 | | | | | | | | | | | | | | | | 44.10 | 10 | 21 | 288 | 34.2 | 6.05 |
| Q8N6Q3 | 25.63 | 1 | 10 | 10 | 60 | 12.59 | 6 | 7 | 2.52 | 2 | 2 | 25.40 | 9 | 14 | 4.12 | 2 | 2 | 5.03 | 3 | 4 | | | | | | | 6.64 | 3 | 3 | 6.64 | 4 | 6 | 18.54 | 7 | 22 | 437 | 46.3 | 6.29 |
| Q8N729 | 10.91 | 1 | 2 | 2 | 2 | | | | | | | | | | | | | | | | | | | | | | | | | | | | 10.91 | 2 | 2 | 165 | 18.0 | 11.62 |
| Q8N766 | 3.83 | 1 | 3 | 3 | 7 | | | | | | | | | | 0.00 | | | 0.00 | | | | | | | | | | | | | | | 3.83 | 3 | 7 | 993 | 111.7 | 7.66 |
| Q8N7B9 | 57.99 | 1 | 23 | 24 | 48 | | | | | | | | | | 11.42 | 4 | 4 | 11.87 | 5 | 5 | | | | | | | | | | | | | 55.71 | 21 | 39 | 438 | 50.1 | 9.22 |
| Q8N7C0 | 23.96 | 1 | 6 | 6 | 9 | | | | | | | | | | | | | | | | | | | | | | | | | | | | 23.96 | 6 | 9 | 313 | 35.1 | 5.74 |
| Q8N7F7 | 10.34 | 1 | 2 | 2 | 3 | 4.02 | 1 | 1 | 4.02 | 1 | 1 | | | | | | | | | | | | | | | | | | | | | | 6.32 | 1 | 1 | 174 | 19.9 | 5.95 |
| Q8N7U6 | 57.26 | 1 | 44 | 44 | 159 | 1.56 | 1 | 1 | | | | | | | 20.17 | 13 | 15 | 33.49 | 23 | 29 | | | | | | | | | | | | | 51.14 | 35 | 110 | 833 | 93.7 | 7.58 |
| Q8N7X0 | 21.18 | 1 | 26 | 26 | 51 | 0.72 | 1 | 1 | 0.00 | | | | | | | | | 0.00 | | | | | | | | | | | | | | | 20.46 | 25 | 50 | 1667 | 189.6 | 8.31 |
| Q8N7X2 | 19.54 | 1 | 6 | 6 | 8 | | | | | | | | | | | | | | | | | | | | | | | | | | | | 19.54 | 6 | 8 | 389 | 42.8 | 9.99 |
| Q8N7Z2 | 17.55 | 3 | 8 | 8 | 16 | | | | 2.25 | 1 | 2 | | | | | | | 13.20 | 5 | 6 | | | | | | | | | | | | | 12.24 | 5 | 8 | 621 | 77.0 | 5.36 |
| Q8N801 | 54.44 | 1 | 10 | 10 | 18 | | | | | | | | | | 15.73 | 3 | 3 | 4.44 | 1 | 1 | | | | | | | | | | | | | 49.19 | 9 | 14 | 248 | 27.8 | 9.58 |
| Q8N807 | 9.76 | 1 | 6 | 6 | 11 | | | | 8.56 | 5 | 5 | 7.19 | 4 | 4 | | | | | | | | | | | | | | | | | | | 2.91 | 2 | 2 | 584 | 66.6 | 6.86 |
| Q8N865 | 61.36 | 3 | 31 | 31 | 118 | | | | | | | | | | 23.05 | 11 | 13 | 38.14 | 18 | 24 | | | | 4.58 | 3 | 3 | 3.56 | 2 | 2 | 3.05 | 2 | 2 | 49.32 | 24 | 74 | 590 | 68.4 | 7.31 |
| Q8N8N7 | 56.70 | 1 | 14 | 14 | 34 | 13.11 | 2 | 4 | 4.27 | 1 | 1 | 2.85 | 1 | 1 | 13.11 | 4 | 4 | | | | | | | 5.98 | 2 | 2 | 1.99 | 1 | 1 | 4.84 | 2 | 2 | 36.75 | 10 | 19 | 351 | 38.5 | 5.41 |
| Q8N9A8 | 43.20 | 1 | 5 | 5 | 7 | | | | | | | | | | | | | | | | | | | | | | | | | | | | 43.20 | 5 | 7 | 125 | 14.3 | 8.38 |

| | | | | | | | | | | | | | | | | | | | | | | | | | | | | | | | | | | | | | | |
|---|---|---|---|---|---|---|---|---|---|---|---|---|---|---|---|---|---|---|---|---|---|---|---|---|---|---|---|---|---|---|---|---|---|---|---|---|---|---|
|Q8N9F7|33.76|1|8|8|22| 4.14|1|2| 9.24|2|3| 4.14|1|1| | | | | | | | | | | | | | | | | | | 29.62|7|16|314|36.1|8.47|
|Q8N9N7|12.97|1|3|3|3| | | | | | | | | | | | | | | | | | | | | | | | | | | | 12.97|3|3|239|26.7|8.43|
|Q8N9U0|10.82|1|6|6|8| | | | | | | | | | | | | | | | | | | | | | | | | | | | 10.82|6|8|490|55.2|9.19|
|Q8N9V2|28.85|1|12|12|28| 4.91|2|2| 1.92|1|1| | | | 6.20|2|2| 2.14|1|1| | | | 3.21|1|1| 6.62|3|3| 3.21|1|1| 27.14|10|17|468|53.0|5.49|
|Q8N9Z9|6.70|1|2|2|2| | | | | | | | | | | | | | | | | | | | | | | | | | | | 6.70|2|2|388|43.4|9.07|
|Q8NA47|68.38|1|52|52|177| 13.50|6|7| 23.45|12|13| 4.62|2|2| 29.48|15|17| 48.13|29|36| | | | 1.60|1|1| | | | 5.51|3|3| 60.21|39|98|563|66.2|9.06|
|Q56NI9|1.00|1|1|1|7| 0.00| | | | | | | | | 0.00| | | | | | | | | | | | | | | | | | 1.00|1|7|601|68.3|9.39|
|Q8NA54|33.12|1|25|25|65| | | | | | | | | | 5.94|4|5| 4.68|3|4| | | | | | | | | | | | | 31.73|24|58|791|92.5|6.67|
|Q8NA56|28.42|1|12|12|23| | | | | | | | | | 5.26|2|2| 16.21|6|6| | | | | | | | | | | | | 23.16|10|15|475|55.0|5.71|
|Q8NA61|52.90|1|27|27|80| 13.62|5|6| 5.58|2|2| 0.00| | | 13.62|4|4| 28.13|14|15| | | | 8.04|3|3| 15.63|7|8| 16.07|7|8| 28.79|15|34|448|51.5|7.15|
|Q8NA69|29.31|1|13|13|23| | | | | | | | | | 2.57|1|1| 13.86|6|6| | | | | | | | | | | | | 23.96|10|16|505|57.3|8.81|
|Q8NA82|6.68|1|6|6|14| 1.11|1|1| | | | | | | | | | 1.24|1|1| | | | | | | 4.70|4|4| | | | 4.33|4|8|808|90.5|6.71|
|Q14004|1.19|17|1|2|7| 0.00| | | 0.00| | | 1.19|2|2| | | | | | | | | | | | | | | | | | | 1.19|2|5|1512|164.8|9.69|
|Q00536|4.64|18|1|3|7| 0.00| | | 0.00| | | 1.61|1|1| 1.81|1|1| | | | | | | | | | | | | | | | 2.82|2|5|496|55.7|7.62|
|Q8NB37|41.36|1|5|5|9| | | | | | | | | | | | | | | | | | | | | | | | | 5.91|1|1| 41.36|5|8|220|23.3|6.61|
|Q8NB90|6.94|1|4|5|18| 1.34|1|2| 1.34|1|2| 1.34|1|1| | | | | | | | | | 1.34|1|2| 1.34|1|1| 1.34|1|1| 6.94|5|9|893|97.8|5.66|
|Q8NBF2|30.17|1|16|16|37| | | | | | | | | | 1.24|1|1| | | | | | | 1.52|1|1| | | | 8.13|5|6| 25.62|13|29|726|79.4|5.55|
|Q8NBJ7|20.27|1|5|5|11| 4.98|1|1| 11.96|3|3| | | | | | | | | | | | | | | | | | | | | | 20.27|5|7|301|33.8|8.00|
|Q8NBM4|11.63|1|3|3|3| | | | | | | | | | | | | | | | | | | | | | | | | | | | 11.63|3|3|344|38.9|9.01|
|Q8NBN3|7.93|1|5|5|9| | | | | | | | | | | | | | | | | | | | | | | | | | | | 7.93|5|9|555|63.4|6.74|
|Q8NBQ5|7.67|1|2|2|2| 7.67|2|2| | | | | | | | | | | | | | | | | | | | | | | | | | | |300|32.9|9.07|
|Q8NBS9|6.02|1|2|2|7| 3.70|1|1| | | | 2.31|1|1| | | | 3.70|1|1| | | | | | | 3.70|1|1| | | | 3.70|1|3|432|47.6|5.97|
|Q8NBU5|17.73|1|5|5|9| | | | | | | | | | | | | | | | | | | | | | | | | | | | 17.73|5|9|361|40.7|6.90|
|Q8NBX0|78.09|1|25|25|201| 40.56|13|17| 49.18|16|26| 40.79|14|18| 11.42|5|5| 7.93|3|3| | | | | | | 1.86|1|1| | | | 73.89|20|131|429|47.1|9.14|
|Q8NBZ7|12.14|1|3|3|8| | | | | | | | | | | | | | | | | | | | | | | | | 4.29|1|1| 7.86|2|7|420|47.5|8.94|
|Q8NCM8|16.23|1|59|61|85| | | | | | | | | | | | | | | | | | | | | | | | | | | | 16.23|61|85|4307|492.3|6.54|
|Q8NCN5|8.99|1|5|5|6| | | | | | | | | | | | | | | | | | | | | | | | | | | | 8.99|5|6|879|99.3|6.35|
|Q8NCQ7|8.52|1|3|3|4| | | | | | | | | | 2.20|1|1| | | | | | | 2.20|1|1| | | | | | | 6.32|2|2|364|40.5|7.81|
|Q8NCR3|6.07|1|2|2|2| | | | | | | | | | | | | | | | | | | | | | | | | | | | 6.07|2|2|313|36.7|7.72|
|Q8NCR6|95.04|1|33|33|231| 30.15|7|8| 23.28|6|6| 7.25|2|2| 61.07|15|21| 67.18|22|33| 0.00| | | 8.40|2|2| 26.34|6|7| 35.88|9|10| 86.26|25|142|262|30.1|8.40|
|Q8NCS7|25.03|1|16|16|33| | | | | | | | | | | | | | | | | | | | | | | | | | | | 25.03|16|33|719|81.6|8.18|
|Q8NCW5|10.76|1|2|2|5| 6.60|1|1| 10.76|2|2| | | | | | | | | | | | | | | | | | | | | | 10.76|2|2|288|31.7|7.66|
|Q8ND07|30.81|1|15|15|18| | | | | | | | | | 1.89|1|1| 8.32|4|4| | | | | | | | | | | | | 25.52|12|13|529|61.9|9.04|
|Q8ND76|19.65|1|3|5|7| | | | | | | | | | | | | | | | | | | | | | | | | | | | 19.65|5|7|341|39.3|7.20|
|Q8NDA8|23.16|1|36|36|48| | | | | | | | | | | | | | | | | | | | | | | | | | | | 23.16|36|48|1641|181.1|6.89|
|Q8NDM7|38.38|1|55|55|158| | | | 0.78|1|1| | | | 10.21|15|16| 12.13|18|20| | | | | | | | | | | | | 33.09|45|121|1665|191.9|5.99|
|Q8NDV3|1.54|1|2|2|2| | | | | | | | | | | | | | | | | | | | | | | | | | | | 1.54|2|2|1235|143.8|7.66|
|Q8NDW8|23.18|1|27|28|36| | | | 0.91|1|1| | | | | | | | | | | | | | | | | | | | | | 23.18|28|35|1320|150.8|7.24|
|Q8NE09|39.87|1|42|42|132| | | | 1.42|2|2| | | | | | | 2.93|3|3| | | | | | | | | | | | | 39.87|42|127|1264|147.1|7.90|
|Q8NE22|29.43|1|6|6|7| | | | 4.35|1|1| | | | | | | | | | | | | | | | | | | | | | 29.43|6|6|299|34.1|8.76|
|Q8NE28|14.41|1|9|9|18| | | | | | | | | | | | | | | | | | | | | | | | | | | | 14.41|9|18|680|75.6|5.36|
|Q8NE62|20.20|1|9|9|12| | | | 3.03|2|2| | | | | | | | | | | | | | | | | | | | | | 18.69|8|10|594|65.3|8.28|
|Q8NE71|13.73|1|8|8|24| | | | | | | | | | 2.84|2|2| | | | | | | 1.30|1|1| | | | 1.30|1|1| 12.19|7|20|845|95.9|6.80|
|Q8NEA4|25.00|1|4|4|4| | | | | | | | | | | | | | | | | | | | | | | | | | | | 25.00|4|4|188|22.1|8.43|
|Q8NEB7|71.82|1|28|28|209| 57.27|19|37| 45.67|18|28| 31.31|12|17| 21.55|9|14| 28.36|15|20| | | | 2.03|1|1| 5.89|3|3| 8.66|4|5| 44.38|18|86|543|61.3|5.16|
|Q8NEC7|17.06|1|9|9|26| 2.37|1|1| 2.37|1|1| | | | 2.37|1|1| 2.37|1|1| | | | | | | | | | | | | 17.06|9|22|633|71.0|7.78|
|Q8NEH6|64.04|1|50|50|132| | | | 4.65|2|2| | | | 40.40|22|29| 59.60|37|58| | | | 2.42|1|1| 2.42|1|1| 2.42|1|1| 45.05|27|40|495|60.5|7.12|
|Q8NEL9|1.89|1|2|2|2| 0.78|1|1| | | | | | | 0.00| | | | | | | | | | | | | | | | | | 1.11|1|1|900|100.4|5.67|
|Q8NEP4|33.86|1|15|15|36| 1.93|1|1| 4.74|2|2| | | | | | | 18.42|7|7| | | | | | | 7.54|3|3| | | | 27.02|11|23|570|63.1|9.20|
|Q8NEU8|21.54|1|10|11|17| | | | | | | | | | 2.11|1|1| | | | | | | | | | | | | 3.92|2|2| 18.07|9|14|664|74.4|4.94|
|Q8NEW0|12.50|1|4|4|4| | | | | | | | | | | | | | | | | | | | | | | | | | | | 12.50|4|4|376|41.6|6.95|
|Q8NEX5|22.47|1|2|2|5| | | | 8.99|1|1| | | | | | | | | | | | | | | | | | | | | | 22.47|2|4|89|10.5|7.55|
|Q8NEY3|46.23|1|11|11|19| 3.93|1|1| | | | | | | | | | | | | | | | | | | | | | | | | 46.23|11|18|305|34.7|9.85|
|Q8NEZ3|8.64|1|10|10|13| | | | | | | | | | | | | | | | | | | | | | | | | | | | 8.64|10|13|1342|151.5|6.35|
|Q8NF91|0.52|3|5|6|19| 0.22|2|3| 0.22|2|3| 0.32|3|3| 0.00| | | 0.15|1|1| | | | | | | | | | | | | 0.42|5|9|8797|1010.5|5.53|
|Q8NFF5|18.57|1|8|8|15| | | | | | | | | | | | | | | | | | | | | | | | | | | | 18.57|8|15|587|65.2|6.93|
|P56377|15.29|2|1|3|7| | | | 5.73|1|1| | | | | | | | | | | | | | | | | | | | | | 15.29|3|6|157|18.6|5.47|
|Q8NFH3|31.84|1|7|8|12| | | | | | | | | | | | | | | | | | | | | | 2.37|1|1| | | | 29.47|7|11|380|42.1|5.63|
|Q8NFH4|35.89|1|7|7|18| | | | | | | | | | | | | | | | | | | | | | | | | | | | 35.89|7|18|326|36.7|5.92|
|Q8NFH5|30.67|1|7|7|25| 19.02|4|4| 19.02|4|4| 3.99|1|1| 3.99|1|1| | | | | | | | | | | | | | | | 30.67|7|15|326|34.8|9.09|
|Q8NFT2|14.08|1|5|5|9| | | | | | | | | | | | | | | | | | | | | | | | | | | | 14.08|5|9|490|56.0|9.20|
|Q8NFU3|26.09|1|4|4|9| | | | | | | | | | 19.13|2|2| 8.70|1|1| | | | 10.43|1|1| | | | 19.13|2|2| 15.65|3|3|115|12.5|6.07|
|Q8NFV4|12.70|1|3|3|4| | | | | | | | | | | | | | | | | | | | | | | | | | | | 12.70|3|4|315|34.7|9.48|
|Q8NFX7|13.33|1|3|3|4| | | | | | | | | | | | | | | | | | | | | | | | | | | | 13.33|3|4|210|23.5|9.04|
|Q8NG35|25.64|1|2|2|2| 25.64|2|2| | | | | | | | | | | | | | | | | | | | | | | | | | | |78|8.9|8.32|
|Q8NGA1|4.79|1|2|2|7| 2.24|1|1| 2.24|1|1| 2.24|1|1| 2.24|1|1| | | | | | | 2.24|1|1| | | | 2.24|1|1| 2.56|1|1|313|34.8|8.91|
|Q8NHG7|42.86|1|3|3|6| | | | 32.47|2|2| | | | | | | | | | | | | | | | | | | | | | 42.86|3|4|77|8.4|8.91|
|Q8NHR9|46.51|1|5|5|7| | | | | | | | | | | | | | | | | | | | | | | | | | | | 46.51|5|7|129|14.3|8.57|
|Q8NHS0|51.72|1|12|14|79| 5.60|1|1| 14.66|3|4| 29.31|4|4| 36.64|7|9| 26.72|6|7| | | | 23.28|4|5| 33.19|8|11| 21.12|4|6| 49.14|10|28|232|25.7|6.42|
|Q8NHS1|5.99|1|2|2|2| | | | | | | | | | | | | | | | | | | | | | | | | | | | 5.99|2|2|167|18.0|7.62|
|Q8NHS2|17.10|1|6|6|7| | | | | | | | | | | | | | | | | | | | | | | | | | | | 17.10|6|7|421|47.3|7.01|
|Q8NHU2|21.42|1|26|26|72| | | | | | | | | | 3.64|5|5| 4.77|6|6| | | | | | | | | | | | | 19.32|22|61|1237|141.3|6.16|
|Q8NHX4|20.31|1|4|4|6| | | | | | | | | | | | | | | | | | | | | | | | | | | | 20.31|4|6|192|20.9|10.02|
|Q8NI22|28.08|1|2|2|5| 19.18|1|1| 28.08|2|4| | | | | | | | | | | | | | | | | | | | | | | | |146|16.4|4.63|
|Q8TAA3|50.78|1|6|11|58| 32.81|6|8| 37.11|7|9| 37.11|7|8| 4.30|1|1| 4.30|1|1| | | | 14.45|3|3| | | | 14.45|3|3| 50.78|10|25|256|28.5|8.98|
|Q8TAG9|7.09|1|4|4|6| | | | | | | | | | | | | | | | | | | | | | | | | | | | 7.09|4|6|804|93.7|6.20|
|Q8TAP6|8.50|1|5|5|10| | | | | | | | | | | | | | | | | | | | | | | | | | | | 8.50|5|10|659|74.4|6.74|
|P04198|1.72|1|1|1|7| | | | | | | | | | | | | 1.72|1|1| | | | 1.72|1|1| | | | 0.00| | | 1.72|1|5|464|49.5|5.66|
|O60739|48.67|1|1|3|7| | | | | | | | | | | | | | | | | | | 12.39|1|1| 12.39|1|1| 12.39|1|1| 48.67|3|4|113|12.8|7.37|
|Q8TAQ9|39.22|1|11|11|17| 4.48|1|1| 1.96|1|1| | | | | | | | | | | | | | | | | | | | | | 39.22|11|15|357|40.5|7.83|
|Q8TAT6|9.38|1|4|4|8| 2.96|1|1| 2.96|1|1| 0.00| | | | | | | | | | | | | | | | | | | | | 9.38|4|6|608|68.1|6.38|
|Q8TB22|18.32|1|12|12|15| | | | | | | | | | | | | | | | | | | | | | | | | | | | 18.32|12|15|786|87.8|7.43|
|Q8TB36|12.57|1|3|3|3| | | | | | | | | | | | | | | | | | | | | | | | | | | | 12.57|3|3|358|41.3|8.34|
|Q8TB61|5.09|1|2|2|3| | | | | | | | | | | | | | | | | | | | | | | | | | | | 5.09|2|3|432|47.5|9.16|
|Q8TBC4|14.90|1|4|4|7| | | | | | | | | | | | | | | | | | | | | | | | | | | | 14.90|4|7|463|51.8|5.45|
|Q8TBF2|23.74|1|4|4|5| | | | | | | | | | | | | | | | | | | | | | | | | | | | 23.74|4|5|198|21.2|6.67|
|Q8TBP6|15.38|1|3|3|3| | | | | | | | | | | | | | | | | | | | | | | | | | | | 15.38|3|3|338|38.1|9.35|
|Q8TBX8|30.17|1|12|12|48| 13.06|5|5| 6.18|2|2| 3.56|1|1| 5.94|2|2| 4.04|1|1| | | | 1.66|1|1| 4.04|1|1| 7.60|3|3| 22.33|9|32|421|47.3|6.84|
|Q8TBY8|14.97|1|13|13|17| | | | | | | | | | | | | | | | | | | | | | | | | 3.62|3|3| 13.80|12|14|1022|119.0|6.29|
|Q8TBY9|22.45|1|28|28|60| | | | 0.78|1|1| | | | 1.83|3|3| 2.00|1|1| | | | | | | | | | | | | 22.11|25|55|1149|129.9|5.08|
|Q8TBZ2|19.85|1|20|20|32| 1.06|1|1| | | | | | | 1.16|1|1| | | | | | | | | | | | | | | | 17.63|18|30|947|108.1|7.49|
|Q8TBZ9|41.50|1|8|8|12| | | | | | | | | | 3.95|1|1| | | | | | | | | | | | | | | | 41.50|8|11|253|29.5|7.69|
|Q8TC07|12.45|1|7|8|14| | | | | | | | | | | | | | | | | | | | | | | | | 4.20|2|2| 8.25|6|12|691|79.4|5.67|
|Q8TC12|55.35|2|15|15|64| | | | 7.86|2|2| 8.18|2|2| 4.40|1|2| | | | | | | | | | | | | 4.40|1|1| 55.35|15|57|318|35.4|8.82|
|Q8TC27|12.71|2|9|9|44| 8.89|5|9| | | | 11.82|7|9| | | | 2.41|1|2| | | | | | | | | | | | | 11.18|7|24|787|87.9|5.55|
|Q8TC29|80.47|2|31|31|120| | | | 17.97|4|4| | | | 57.03|15|20| 70.31|25|43| | | | | | | | | | | | | 63.67|17|53|256|29.4|9.31|
|P42677|40.48|1|1|3|7| | | | | | | | | | | | | | | | | | | | | | | | | | | | 40.48|3|7|84|9.5|9.45|
|Q8TC36|13.72|1|4|4|5| | | | | | | | | | | | | | | | | | | | | | | | | | | | 13.72|4|5|379|43.1|8.31|
|Q8TC44|12.76|1|5|5|6| | | | | | | | | | | | | | | | | | | | | | | | | | | | 12.76|5|6|478|53.6|7.24|
|Q8TC56|40.50|1|21|22|92| 13.88|6|6| 25.45|12|13| 17.36|8|8| | | | 3.64|2|2| | | | | | | 7.93|4|4| | | | 23.64|14|59|605|64.7|9.48|
|Q8TC71|47.96|1|26|26|119| 28.44|13|17| 22.86|11|12| 6.13|2|2| 4.46|3|3| 18.22|7|7| | | | | | | 1.67|1|1| 1.67|1|1| 46.47|21|76|538|61.1|8.63|
|Q8TC94|32.93|1|7|7|11| | | | | | | | | | | | | | | | | | | | | | | | | | | | 32.93|7|11|416|45.6|7.08|
|Q8TC99|29.94|1|7|7|16| 5.25|1|1| | | | | | | | | | | | | | | | | | | 5.56|1|1| | | | 29.94|6|14|324|35.9|5.08|
|Q8TCD5|27.86|1|3|3|8| 8.96|1|1| | | | | | | | | | | | | | | | 6.47|1|1| | | | | | | 27.86|3|6|201|23.4|6.64|
|Q8TCG1|3.76|1|4|4|4| | | | | | | | | | | | | | | | | | | 0.99|1|1| 1.10|1|1| 0.00| | | 1.66|2|2|905|102.1|6.23|
|Q8TCI5|26.18|1|4|4|10| | | | | | | | | | | | | | | | | | | | | | | | | | | | 26.18|4|10|191|22.0|9.89|
|Q8TCJ2|15.74|1|11|13|32| 4.12|3|3| 3.51|3|3| 1.09|1|1| | | | | | | | | | | | | | | | | | | 15.74|12|25|826|93.6|8.91|
|O75351|8.78|2|1|3|7| | | | | | | | | | | | | | | | | | | | | | | | | | | | 8.78|3|7|444|49.3|7.23|
|Q8TCS8|18.26|1|11|11|13| | | | | | | 4.34|3|3| | | | | | | | | | | | | | | | | | | 13.92|8|10|783|85.9|7.77|
|Q8TCT0|10.99|1|4|4|5| | | | 2.98|1|1| 1.86|1|1| | | | | | | | | | | | | | | | | | | 6.15|2|3|537|59.9|8.19|
|Q8TCT7|8.28|1|4|4|8| | | | 1.86|1|1| | | | | | | | | | | | | | | | | | | | | | 8.28|4|7|592|64.6|8.32|
|Q8TCT9|20.69|1|8|8|30| 3.18|1|1| | | | 0.00| | | | | | | | | | | | | | | | | | | | | 20.69|8|29|377|41.5|6.43|
|P45983|4.92|7|1|2|7| | | | | | | | | | | | | | | | | | | | | | | | | | | | 4.92|2|7|427|48.3|6.89|
|Q8TCU4|1.30|1|3|3|6| | | | | | | | | | | | | | | | | | | | | | | | | | | | 1.30|3|6|4167|460.7|6.28|
|Q8TD19|5.62|1|4|4|7| | | | | | | | | | | | | | | | | | | | | | | | | | | | 5.62|4|7|979|107.1|5.74|
|Q8TD57|40.21|2|148|159|373| 0.95|3|4| 0.97|4|4| 0.22|1|1| 6.85|25|25| 4.08|14|14| | | | | | | | | | | | | 37.59|149|325|4116|470.5|6.43|
|Q8TDB8|22.69|2|7|16|197| 13.08|7|12| 14.42|9|18| 11.92|7|14| 11.92|7|9| 8.27|4|4| | | | | | | | | | | | | 20.00|12|140|520|56.3|7.83|
|Q8TDL5|14.26|1|4|4|4| | | | 14.26|4|4| | | | | | | | | | | | | | | | | | | | | | | | |484|52.4|7.23|
|Q8TDM5|21.77|1|3|3|13| 21.77|2|2| 10.48|1|1| 11.29|1|1| | | | 10.48|1|1| | | | | | | | | | | | | 21.77|3|8|124|13.0|5.80|
|Q8TDQ7|27.90|1|2|6|9| 4.35|1|1| 7.61|2|2| | | | | | | | | | | | | | | | | | | | | | 24.64|5|6|276|31.1|6.95|
|Q8TDR0|2.75|1|2|2|2| | | | | | | | | | | | | | | | | | | | | | | | | | | | 2.75|2|2|691|78.6|7.93|
|Q8TDY3|49.60|1|14|14|28| 10.88|3|3| 10.88|3|3| 5.57|2|2| 5.84|2|2| 2.92|1|1| | | | | | | | | | 5.84|2|2| 38.46|11|15|377|41.7|5.47|
|Q8TDZ2|19.03|1|14|14|41| | | | 1.12|1|1| | | | | | | 0.00| | | | | | | | | 0.00| | | 1.31|1|1| 18.46|13|39|1067|117.8|6.40|
|Q8TE73|4.54|1|3|26|87| 0.89|5|5| 0.89|5|5| 0.54|3|3| 0.69|4|4| 0.50|3|3| | | | | | | | | | | | | 4.28|23|67|4624|528.7|6.10|
|Q8TE77|7.44|1|3|3|4| | | | | | | | | | | | | | | | | | | | | | | | | 1.97|1|1| 5.46|2|3|659|73.0|5.30|
|Q8TEM1|22.58|1|34|34|66| 2.49|4|4| 4.35|7|7| 7.95|12|12| | | | | | | | | | | | | | | | | | | 21.04|31|43|1887|205.0|6.81|
|Q8TEV9|2.99|1|3|3|4| | | | | | | | | | | | | | | | | | | | | | | | | | | | 2.99|3|4|937|105.0|5.58|
|Q8TEX9|50.69|1|40|40|259| 2.68|3|3| 3.24|2|2| 1.02|1|1| 1.85|2|2| 0.00| | | | | | | | | | | | | | | 50.69|40|251|1081|118.6|4.96|
|Q8TF05|27.37|1|19|19|61| 3.26|3|4| 6.32|5|6| 3.47|2|2| 1.89|2|2| | | | | | | | | | | | | | | | 25.68|17|47|950|106.9|4.77|
|Q8TF09|75.00|2|8|8|33| 29.17|2|2| 41.67|4|5| 29.17|2|2| 57.29|5|7| 36.46|3|6| | | | | | | | | | 16.67|1|1| 69.79|6|10|96|10.8|7.50|
|Q8TF46|1.80|1|2|2|2| | | | 0.00| | | 0.00| | | | | | | | | | | | | | | | | | | | | 1.80|2|2|1054|120.7|6.54|
|Q8WTQ1|25.00|1|2|2|3| | | | | | | | | | | | | | | | | | | | | | | | | | | | 25.00|2|3|72|8.5|9.19|
|Q8WTQ4|8.30|1|3|3|3| | | | 0.00| | | | | | | | | 6.04|1|1| | | | | | | | | | | | | 6.42|2|2|265|30.8|9.79|
|Q8WTS6|7.65|1|2|2|4| | | | | | | | | | | | | | | | | | | 2.73|1|1| 2.73|1|1| 2.73|1|1| 4.92|1|1|366|40.7|4.63|
|Q8WTU0|32.83|4|8|9|21| | | | | | | | | | | | | | | | | | | | | | | | | | | | 32.83|9|21|396|44.1|5.73|
|Q8WTW3|7.14|1|6|6|9| | | | | | | | | | | | | | | | | | | | | | | | | | | | 7.14|6|9|980|108.9|7.31|
|Q8WU76|21.78|1|10|10|11| 0.00| | | 0.00| | | 0.00| | | | | | 0.00| | | | | | 0.00| | | 0.00| | | | | | 21.78|10|11|684|75.1|6.68|
|Q8WUD1|82.41|1|9|18|111| 34.26|7|8| 52.31|10|16| 38.43|8|10| 16.20|3|4| | | | | | | | | | | | | | | | 82.41|18|73|216|24.2|7.83|
|Q8WUK0|64.18|1|11|11|21| 12.44|2|2| 30.85|5|5| | | | | | | | | | | | | | | | | | | | | | 51.74|9|14|201|22.8|9.77|
|Q8WUM0|16.09|1|12|12|21| | | | | | | 0.00| | | | | | | | | | | | | | | | | | | | | 16.09|12|21|1156|128.9|5.10|
|Q8WUM4|56.91|1|48|48|208| 13.36|9|9| 8.06|5|5| 12.44|10|10| 12.10|9|9| 1.27|1|1| | | | 12.21|10|11| 3.57|3|3| 25.23|18|22| 54.84|43|138|868|96.0|6.52|
|Q8WUN7|31.62|2|10|10|19| 20.94|4|4| 19.66|5|6| 5.56|1|1| | | | | | | | | | | | | | | | | | | 28.21|7|8|234|26.2|5.67|
|Q8WUW1|66.67|1|5|5|9| 17.33|1|1| | | | | | | 22.67|2|2| | | | | | | | | | | | | 22.67|2|2| 56.00|4|4|75|8.7|5.45|
|Q8WUX2|12.50|1|2|2|3| | | | | | | | | | | | | | | | | | | | | | | | | | | | 12.50|2|3|184|20.9|5.43|
|Q8WUY3|1.26|1|3|4|7| | | | | | | | | | | | | | | | | | | | | | | | | | | | 1.26|4|7|3088|340.4|4.45|
|Q8WV93|7.28|1|3|3|3| | | | | | | | | | | | | | | | | | | | | | | | | | | | 7.28|3|3|481|54.8|7.27|
|Q8WVC6|9.96|1|2|2|2| | | | | | | | | | | | | | | | | | | | | | | | | | | | 9.96|2|2|231|26.5|9.58|
|Q8WVD3|14.29|1|2|2|3| | | | | | | | | | | | | | | | | | | | | | | | | | | | 14.29|2|3|245|28.2|6.93|
|Q8WVF1|10.54|1|2|2|4| | | | | | | | | | | | | | | | | | | | | | | | | | | | 10.54|2|4|389|44.6|5.83|
|Q8WVJ2|52.87|1|6|6|16| | | | | | | | | | 7.01|1|1| 7.01|1|1| | | | 11.46|1|1| | | | 22.93|2|3| 52.87|6|10|157|17.7|5.07|
|Q8WVM8|26.32|1|11|11|17| 0.00| | | 1.40|1|1| 0.00| | | 0.00| | | | | | | | | | | | | | | | | | 26.32|11|16|642|72.3|6.27|
|Q8WVP5|23.12|1|3|3|4| | | | | | | | | | | | | | | | | | | | | | | | | | | | 23.12|3|4|186|20.8|9.50|
|Q8WVZ1|6.80|1|2|2|3| | | | | | | | | | | | | | | | | | | | | | | | | | | | 6.80|2|3|309|34.3|8.18|
|Q8WW14|75.65|1|16|16|67| 18.26|3|3| 7.83|2|2| | | | 40.87|9|11| 51.74|13|18| | | | | | | | | | 14.35|3|3| 63.04|11|30|230|25.9|8.47|
|Q8WW22|58.69|1|22|22|90| 17.13|5|5| 27.71|9|10| 15.11|4|5| 12.09|4|4| 19.65|7|8| | | | 6.80|2|2| 12.34|4|4| 11.08|4|4| 54.41|17|48|397|44.8|7.59|
|Q8WW24|88.28|1|54|54|366| 28.97|10|11| 36.55|13|16| 8.51|3|3| 60.23|27|34| 65.98|30|48| | | | 40.23|18|29| 42.99|21|30| 59.08|28|45| 82.07|39|150|435|50.6|6.44|
|Q8WW35|38.73|1|3|3|4| | | | | | | | | | | | | | | | | | | | | | | | | | | | 38.73|3|4|142|16.1|4.93|
|Q8WW59|9.66|1|2|2|2| | | | | | | | | | | | | | | | | | | | | | | | | | | | 9.66|2|2|207|23.1|6.93|
|Q8WWB3|62.15|1|12|12|42| 18.08|2|2| 9.04|1|1| | | | 51.98|8|8| 27.12|4|5| 4.52|1|1| 3.39|1|1| 5.08|1|1| 16.95|3|3| 44.07|8|20|177|20.9|4.67|
|Q8WWC4|11.34|1|3|3|3| | | | | | | | | | | | | | | | | | | | | | | | | | | | 11.34|3|3|291|32.5|9.17|
|Q8WWU5|64.21|1|27|27|174| 2.78|1|1| | | | | | | 10.14|5|5| 2.78|1|1| | | | 4.17|1|4| | | | 6.36|2|4| 64.21|27|159|503|56.1|5.17|
|Q8WWZ7|1.83|1|2|3|3| | | | | | | | | | | | | 0.00| | | | | | | | | | | | | | | 1.83|3|3|1642|186.4|6.95|
|Q8WXA2|65.08|1|9|9|40| 38.89|5|6| 7.14|1|1| 35.71|5|7| 11.90|1|1| 13.49|2|2| | | | | | | | | | | | | 62.70|7|23|126|14.3|7.93|
|Q8WXF7|9.68|1|4|4|6| 2.69|1|1| | | | | | | | | | | | | | | | | | | | | | | | | 9.68|4|5|558|63.5|6.18|
|Q8WXH0|0.52|1|3|4|4| 0.00| | | 0.00| | | | | | | | | 0.00| | | | | | | | | | | | 0.00| | | 0.52|4|4|6885|795.9|5.36|
|Q8WXR4|13.80|2|13|15|27| | | | | | | | | | | | | | | | | | | | | | | | | | | | 13.80|15|27|1341|151.7|8.15|
|Q8WXX0|50.22|1|173|186|557| 1.19|4|5| 1.32|5|5| 2.46|8|8| 15.06|53|55| 8.70|32|34| | | | | | | | | | 0.57|2|2| 48.06|175|448|4024|460.9|6.00|
|Q8WY22|22.71|1|4|4|7| 6.77|1|1| 6.77|1|1| | | | | | | | | | | | | | | | | | | | | | 22.71|4|5|251|27.8|9.44|
|Q8WYA0|15.68|1|11|11|11| | | | | | | | | | | | | 1.33|1|1| | | | | | | | | | | | | 15.53|10|10|676|79.7|8.82|
|Q8WYR4|54.37|1|15|15|84| 8.74|2|2| 11.97|3|3| 14.56|3|3| 38.19|9|16| 35.92|8|12| | | | 7.77|2|2| 12.30|3|5| 6.80|2|3| 46.60|12|38|309|35.1|4.63|
|Q8WZ42|0.20|1|6|8|16| 0.02|1|1| 0.05|2|2| 0.02|1|1| 0.09|3|3| 0.00| | | | | | | | | 0.00| | | 0.00| | | 0.15|6|9|34350|3813.7|6.35|
|Q8WZ59|39.55|1|7|7|29| 17.51|2|2| 7.91|2|2| | | | | | | | | | | | | | | | | | | | | | 39.55|7|25|177|19.4|5.24|
|Q8WZ82|6.61|1|2|2|2| | | | | | | | | | | | | | | | | | | | | | | | | | | | 6.61|2|2|227|24.4|6.89|
|Q92485|16.26|1|6|6|7| | | | | | | | | | | | | | | | | | | | | | | | | | | | 16.26|6|7|455|50.8|5.64|
|Q92499|13.51|1|8|8|15| 2.84|1|1| | | | 0.00| | | 1.89|1|1| 1.89|1|1| | | | | | | | | | 4.05|2|2| 11.62|7|10|740|82.4|7.23|
|Q92503|15.94|1|8|8|12| | | | | | | | | | | | | | | | | | | | | | | | | | | | 15.94|8|12|715|81.2|6.43|
|Q92506|15.71|1|3|3|3| | | | 4.60|1|1| | | | | | | | | | | | | | | | | | | | | | 11.11|2|2|261|27.0|6.54|
|Q92520|15.86|1|3|3|3| | | | | | | | | | | | | | | | | | | | | | | | | | | | 15.86|3|3|227|24.7|8.29|
|Q92523|45.08|1|32|32|145| 18.78|11|13| 20.98|14|18| 9.97|7|7| | | | | | | | | | | | | | | | | | | 41.32|29|107|772|87.7|8.62|
|Q92526|54.34|1|16|21|141| 29.06|10|11| 31.70|13|14| 17.17|8|8| 13.21|6|7| 13.40|6|7| | | | 10.19|5|5| 12.08|5|5| 7.36|4|5| 49.81|19|79|530|57.8|7.24|
|Q92530|19.93|1|4|4|13| | | | | | | | | | | | | | | | | | | | | | | | | 3.69|1|1| 16.24|3|12|271|29.8|5.74|
|Q92538|6.83|1|11|13|30| 0.32|1|1| 0.32|1|1| 0.32|1|1| | | | 0.32|1|1| | | | 0.32|1|1| 0.32|1|1| 0.32|1|1| 6.83|13|23|1859|206.3|5.73|
|Q92542|17.77|1|10|10|39| 5.92|3|3| 5.78|3|3| 7.62|5|5| | | | | | | | | | | | | | | | | | | 15.80|9|28|709|78.4|5.99|
|Q92544|11.99|1|7|7|12| | | | 1.25|1|1| | | | | | | | | | | | | | | | | | | | | | 11.99|7|11|642|74.5|6.54|
|Q92572|29.02|1|4|4|10| | | | | | | | | | | | | | | | | | | | | | | | | 5.70|1|1| 29.02|4|9|193|21.7|5.39|
|Q92574|3.95|1|3|3|3| | | | | | | | | | | | | | | | | | | | | | | | | | | | 3.95|3|3|1164|129.7|6.47|
|Q92597|45.18|1|12|12|160| 19.80|5|6| 16.50|4|4| 27.92|7|9| 27.41|7|12| 24.37|6|7| | | | 23.60|6|11| 31.22|9|14| 28.43|9|16| 44.16|10|81|394|42.8|5.82|
|Q92598|42.77|2|25|30|86| 4.43|3|3| 7.81|5|5| 15.50|10|10| | | | 1.05|1|2| | | | 1.63|1|1| 2.56|2|2| 1.86|2|3| 34.50|24|60|858|96.8|5.39|
|Q92599|24.84|2|5|10|23| 1.86|1|1| 1.86|1|1| 1.86|1|1| 1.86|1|1| 6.21|2|3| | | | | | | | | | | | | 24.43|9|16|483|55.7|6.28|
|P06493|6.06|17|1|2|6| 0.00| | | 0.00| | | 2.69|1|1| | | | | | | | | | | | | | | | | | | 6.06|2|5|297|34.1|8.40|
|Q92600|7.02|1|2|2|2| | | | | | | | | | | | | | | | | | | | | | | | | | | | 7.02|2|2|299|33.6|8.03|
|Q92616|30.66|1|66|66|194| | | | 0.00| | | 0.00| | | 0.49|1|1| 0.00| | | | | | | | | | | | | | | 30.66|66|193|2671|292.6|7.47|
|Q92621|38.62|2|68|68|231| 3.38|5|5| 6.46|10|12| 1.24|2|2| | | | 0.75|1|1| | | | | | | | | | | | | 37.87|67|211|2012|227.8|6.19|
|P21802|1.95|17|1|2|6| 0.00| | | 0.00| | | 0.97|1|1| | | | | | | | | | | | | | | | | | | 1.95|2|5|821|92.0|5.88|
|Q92643|8.10|1|3|3|3| | | | | | | | | | | | | | | | | | | | | | | | | | | | 8.10|3|3|395|45.2|6.16|
|Q92667|3.32|1|2|2|2| | | | | | | | | | | | | | | | | | | | | | | | | | | | 3.32|2|2|903|97.3|4.94|
|Q92696|5.29|1|3|3|3| | | | | | | | | | | | | | | | | | | | | | | | | | | | 5.29|3|3|567|65.0|5.67|
|Q92734|12.00|1|4|4|5| | | | | | | | | | | | | | | | | | | | | | | | | | | | 12.00|4|5|400|43.4|5.10|
|Q92743|35.42|1|14|14|36| 4.58|2|2| 6.67|3|3| 2.92|1|1| 2.92|1|1| 13.54|5|5| | | | | | | | | | 2.92|1|1| 32.29|11|23|480|51.3|7.83|
|Q92747|27.30|1|9|9|20| | | | | | | 9.19|2|3| | | | | | | | | | | | | | | | | | | 27.30|9|17|370|41.5|8.18|
|Q92765|5.23|1|2|2|2| | | | | | | | | | | | | | | | | | | | | | | | | | | | 5.23|2|2|325|36.2|8.46|
|Q92783|11.30|1|4|5|8| | | | | | | | | | | | | | | | | | | 1.85|1|1| | | | 4.44|2|2| 9.44|4|5|540|59.1|4.82|
|Q92820|47.48|2|14|14|52| 22.64|8|13| 27.99|10|12| 27.99|9|11| | | | 4.09|1|1| | | | | | | | | | | | | 33.96|8|15|318|35.9|7.11|
|Q92834|7.35|1|5|5|5| | | | | | | | | | | | | | | | | | | | | | | | | | | | 7.35|5|5|1020|113.3|4.86|
|Q92878|11.74|2|14|15|16| | | | | | | | | | | | | | | | | | | | | | | | | | | | 11.74|15|16|1312|153.8|6.89|
|Q92882|34.11|1|5|5|12| 5.61|1|1| | | | | | | 6.54|1|1| | | | | | | 6.54|1|1| | | | 22.90|3|3| 17.76|3|6|214|23.8|5.68|
|Q92890|8.79|1|2|2|3| | | | | | | | | | | | | | | | | | | | | | | | | | | | 8.79|2|3|307|34.5|6.70|
|Q92665|2.78|1|1|1|6| 2.78|1|2| 2.78|1|2| 2.78|1|2| | | | | | | | | | | | | | | | | | | | | |395|45.3|9.29|
|Q92896|8.65|1|9|9|12| 3.22|3|3| | | | 1.10|1|1| | | | | | | | | | | | | | | | | | | 7.38|8|8|1179|134.5|6.90|
|Q53FT3|6.60|1|1|1|6| 6.60|1|1| 6.60|1|1| | | | 6.60|1|1| | | | | | | | | | 6.60|1|1| 6.60|1|1| 6.60|1|1|197|21.6|5.45|
|Q92900|12.40|2|12|12|20| | | | | | | 0.97|1|1| | | | | | | | | | | | | | | | | | | 12.40|12|19|1129|124.3|6.61|
|Q92905|49.40|1|12|12|23| | | | 5.09|2|2| 2.99|1|1| 9.58|3|3| | | | 2.99|1|1| 2.99|1|1| | | | | | | 41.92|10|15|334|37.6|6.54|
|Q8IWA5|2.55|1|1|2|6| 1.70|1|1| | | | 1.70|1|1| | | | | | | | | | | | | | | | | | | 2.55|2|4|706|80.1|8.57|
|Q92930|48.31|7|4|11|65| 23.19|4|5| 16.43|3|4| 20.29|4|4| 13.04|3|3| | | | | | | 12.56|3|3| 13.53|4|4| 13.53|3|3| 47.34|10|39|207|23.6|9.07|
|Q92947|4.11|1|2|2|2| | | | | | | | | | | | | | | | | | | | | | | | | | | | 4.11|2|2|438|48.1|8.06|
|Q92968|19.11|1|4|4|5| | | | | | | | | | | | | | | | | | | | | | | | | | | | 19.11|4|5|403|44.1|8.05|
|Q92973|20.04|1|8|13|38| | | | | | | 1.34|1|1| 1.34|1|1| | | | | | | | | | | | | | | | 20.04|13|38|898|102.3|4.98|
|Q92990|6.73|1|3|3|4| | | | | | | | | | | | | | | | | | | | | | | | | | | | 6.73|3|4|594|68.2|5.33|
|Q93008|11.98|8|13|26|57| | | | | | | | | | | | | | | | | | | | | | | | | | | | 11.98|26|57|2570|292.1|5.80|
|Q93009|39.56|1|38|38|131| 7.44|5|6| 7.62|7|8| 9.62|8|9| | | | 0.91|1|1| | | | | | | 3.09|3|3| | | | 38.75|37|104|1102|128.2|5.55|
|Q93034|14.74|1|11|11|12| | | | | | | | | | | | | | | | | | | | | | | | | | | | 14.74|11|12|780|90.9|7.94|
|Q6N069|5.90|1|1|5|6| | | | 0.00| | | | | | | | | | | | | | | 0.00| | | | | | | | | 5.90|5|6|864|101.4|7.87|
|Q93050|19.12|1|13|14|33| 1.43|1|1| 1.43|1|1| 4.78|3|3| 2.15|1|1| | | | | | | | | | | | | | | | 19.12|13|27|837|96.4|6.43|
|Q93052|15.20|1|6|6|13| | | | | | | | | | | | | | | | | | | 2.29|1|1| 7.03|3|4| | | | 11.93|4|8|612|65.7|7.37|
|Q93084|17.16|1|8|15|42| 4.03|3|3| 2.21|2|2| 6.81|6|7| 2.68|2|2| 1.25|1|1| | | | | | | | | | 1.44|1|1| 13.04|11|26|1043|113.9|5.57|
|Q93096|36.42|1|2|6|7| | | | | | | | | | | | | | | | | | | | | | | | | | | | 36.42|6|7|173|19.8|8.97|
|Q93099|40.90|1|13|13|25| 4.94|2|2| 2.47|1|1| | | | | | | | | | | | | | | | | | | | | | 38.43|12|22|445|49.9|6.96|
|Q93100|13.54|1|14|14|24| | | | | | | | | | | | | | | | | | | | | | | | | | | | 13.54|14|24|1093|124.8|6.95|
|Q969E2|10.04|1|2|2|15| 4.80|1|1| 4.80|1|1| 4.80|1|1| | | | | | | | | | | | | | | | | | | 10.04|2|12|229|25.7|8.82|
|Q969H8|30.06|1|6|6|21| 16.18|3|3| 27.17|5|5| 16.18|3|3| 6.36|1|1| 6.36|1|1| | | | | | | 11.56|1|1| | | | 27.17|4|7|173|18.8|6.68|
|Q969N2|4.84|1|2|2|3| | | | | | | | | | | | | | | | | | | | | | | | | | | | 4.84|2|3|578|65.7|8.38|
|Q969Q0|17.92|2|2|2|3| | | | | | | | | | | | | | | | | | | | | | | | | | | | 17.92|2|3|106|12.5|10.65|
|P51116|5.94|2|1|3|6| | | | | | | 1.19|1|1| 2.38|2|2| | | | | | | | | | | | | 1.19|1|1| 4.75|2|2|673|74.2|6.23|
|Q969R2|6.66|1|4|6|13| 2.95|2|2| | | | | | | | | | | | | | | | | | | | | | | | | 4.91|4|11|916|101.2|6.54|
|Q969U7|11.36|1|3|3|3| | | | | | | | | | | | | | | | | | | | | | | | | | | | 11.36|3|3|264|29.4|6.98|
|Q969V3|12.08|1|5|5|6| | | | | | | | | | | | | | | | | | | | | | | | | | | | 12.08|5|6|563|62.9|6.89|
|Q969V4|76.56|1|39|39|199| 9.09|3|3| 23.68|9|10| 2.39|1|1| 28.23|11|13| 43.54|17|22| | | | 34.45|17|21| 31.58|14|19| 48.56|22|30| 67.22|28|80|418|48.3|6.37|
|Q969X1|4.82|1|2|2|5| | | | | | | 4.82|1|2| | | | | | | | | | | | | | | | | | | 4.50|1|3|311|34.6|7.72|
|Q969X5|43.45|1|10|10|22| 3.45|1|1| 3.45|1|1| 11.38|2|2| | | | | | | | | | | | | | | | | | | 43.45|10|18|290|32.6|7.06|

| | | | | | | | | | | | | | | | | | | | | | | | | | | | | | | | | | | | | | | |
|---|---|---|---|---|---|---|---|---|---|---|---|---|---|---|---|---|---|---|---|---|---|---|---|---|---|---|---|---|---|---|---|---|---|---|---|---|---|---|
| Q96A05 | 56.64 | 1 | 9 | 14 | 20 | 7.96 | 2 | 2 | 9.73 | 2 | 2 | | | | | | | | | | | | | | | | | | | | | | 56.64 | 14 | 16 | 226 | 26.1 | 8.76 |
| Q96A08 | 66.14 | 1 | 4 | 9 | 69 | 33.07 | 4 | 6 | 40.16 | 5 | 12 | 33.07 | 4 | 5 | 14.17 | 2 | 3 | 22.05 | 3 | 3 | 5.51 | 1 | 1 | 8.66 | 1 | 1 | | | | 8.66 | 1 | 1 | 66.14 | 9 | 37 | 127 | 14.2 | 10.32 |
| P36888 | 1.71 | 9 | 1 | 2 | 6 | | | | | | | | | | 0.81 | 1 | 1 | | | | | | | | | | | | | | | | 1.71 | 2 | 5 | 993 | 112.8 | 5.68 |
| A6NL82 | 20.15 | 1 | 1 | 2 | 6 | | | | | | | | | | 11.94 | 1 | 1 | 11.94 | 1 | 2 | | | | | | | | | | | | | 20.15 | 2 | 3 | 134 | 16.0 | 8.65 |
| Q96A26 | 33.12 | 1 | 8 | 8 | 12 | | | | 16.23 | 2 | 2 | | | | | | | | | | | | | | | | | | | | | | 33.12 | 7 | 10 | 154 | 17.3 | 9.77 |
| Q96A65 | 12.94 | 1 | 10 | 10 | 18 | | | | | | | | | | | | | | | | | | | | | | | | | 2.98 | 2 | 2 | 12.94 | 10 | 16 | 974 | 110.4 | 6.49 |
| Q96AA3 | 4.07 | 1 | 3 | 3 | 4 | | | | | | | | | | | | | | | | | | | | | | | | | | | | 4.07 | 3 | 4 | 541 | 60.3 | 8.85 |
| Q96AB3 | 56.59 | 1 | 7 | 7 | 32 | 38.05 | 4 | 7 | 38.05 | 4 | 6 | 28.29 | 3 | 3 | 4.39 | 1 | 1 | | | | | | | | | | | | | | | | 52.20 | 6 | 15 | 205 | 22.3 | 7.77 |
| Q96AB6 | 7.42 | 1 | 3 | 3 | 3 | | | | | | | | | | | | | | | | | | | | | | | | | | | | 7.42 | 3 | 3 | 310 | 34.7 | 6.27 |
| P41567 | 48.67 | 1 | 1 | 3 | 6 | | | | | | | | | | | | | | | | | | | 12.39 | 1 | 1 | 12.39 | 1 | 1 | 12.39 | 1 | 1 | 48.67 | 3 | 3 | 113 | 12.7 | 7.44 |
| Q96AG4 | 19.54 | 1 | 4 | 4 | 6 | | | | | | | | | | | | | | | | | | | | | | | | | | | | 19.54 | 4 | 6 | 307 | 34.9 | 9.57 |
| Q96AJ1 | 20.58 | 2 | 8 | 9 | 15 | | | | | | | | | | | | | | | | | | | | | | | | | | | | 20.58 | 9 | 15 | 413 | 48.1 | 4.77 |
| Q96AQ6 | 2.74 | 1 | 2 | 2 | 2 | 1.37 | 1 | 1 | | | | | | | | | | | | | | | | | | | | | | | | | 1.37 | 1 | 1 | 731 | 80.6 | 5.33 |
| Q96BH3 | 63.68 | 1 | 13 | 13 | 81 | 35.87 | 7 | 8 | 39.46 | 8 | 9 | 39.01 | 7 | 9 | 12.56 | 3 | 3 | 26.91 | 7 | 9 | | | | | | | | | | | | | 63.68 | 12 | 43 | 223 | 26.1 | 6.62 |
| Q9P289 | 9.38 | 2 | 1 | 3 | 6 | | | | | | | | | | | | | | | | | | | | | | | | | 1.68 | 1 | 1 | 9.38 | 3 | 5 | 416 | 46.5 | 5.29 |
| Q96BJ3 | 4.58 | 1 | 2 | 2 | 2 | | | | | | | | | | | | | | | | | | | | | | | | | | | | 4.58 | 2 | 2 | 306 | 35.0 | 6.55 |
| Q96BJ8 | 2.50 | 1 | 2 | 2 | 2 | | | | | | | | | | | | | | | | | | | | | | | | | | | | 2.50 | 2 | 2 | 720 | 81.4 | 6.30 |
| Q96BN8 | 5.11 | 1 | 2 | 2 | 4 | | | | | | | | | | | | | | | | | | | | | | | | | | | | 5.11 | 2 | 4 | 352 | 40.2 | 5.47 |
| Q96BS2 | 10.75 | 1 | 2 | 2 | 2 | | | | | | | | | | | | | | | | | | | | | | | | | | | | 10.75 | 2 | 2 | 214 | 24.7 | 4.93 |
| Q96BW5 | 10.60 | 1 | 2 | 2 | 3 | | | | | | | | | | | | | | | | | | | | | | | | | | | | 10.60 | 2 | 3 | 349 | 39.0 | 6.52 |
| Q96BX8 | 11.98 | 1 | 2 | 2 | 2 | | | | | | | | | | | | | | | | | | | | | | | | | | | | 11.98 | 2 | 2 | 217 | 25.4 | 8.63 |
| Q96BY7 | 1.92 | 1 | 3 | 3 | 3 | | | | | | | | | | | | | | | | | | | | | | | | | | | | 1.92 | 3 | 3 | 2078 | 232.6 | 5.76 |
| Q96C19 | 8.75 | 1 | 2 | 2 | 4 | | | | | | | | | | | | | | | | | | | | | | | | | | | | 8.75 | 2 | 4 | 240 | 26.7 | 5.20 |
| Q96C23 | 13.45 | 1 | 3 | 3 | 12 | | | | | | | | | | | | | | | | | | | 4.39 | 1 | 1 | 4.09 | 1 | 1 | 8.48 | 2 | 2 | 13.45 | 3 | 8 | 342 | 37.7 | 6.65 |
| Q12974 | 30.54 | 1 | 1 | 5 | 6 | | | | | | | | | | | | | | | | | | | | | | | | | | | | 30.54 | 5 | 6 | 167 | 19.1 | 8.37 |
| Q96C24 | 5.51 | 1 | 4 | 4 | 4 | | | | | | | | | | | | | | | | | | | | | | | | | 1.94 | 1 | 1 | 3.58 | 3 | 3 | 671 | 76.0 | 8.98 |
| Q96C74 | 78.26 | 1 | 18 | 18 | 66 | 15.22 | 3 | 3 | 23.04 | 5 | 5 | | | | 27.83 | 6 | 6 | 23.04 | 4 | 5 | | | | | | | | | | | | | 72.17 | 15 | 47 | 230 | 26.1 | 7.72 |
| Q96C92 | 9.89 | 1 | 4 | 4 | 5 | | | | | | | | | | | | | | | | | | | | | | | | | 3.45 | 1 | 1 | 8.97 | 3 | 4 | 435 | 47.9 | 5.14 |
| Q96CD0 | 22.99 | 1 | 6 | 6 | 11 | | | | | | | | | | | | | | | | | | | | | | | | | | | | 22.99 | 6 | 11 | 374 | 40.5 | 7.31 |
| Q96CN7 | 33.22 | 1 | 7 | 7 | 7 | | | | 5.03 | 1 | 1 | | | | | | | | | | | | | | | | | | | | | | 28.19 | 6 | 6 | 298 | 32.2 | 7.39 |
| Q96CS3 | 25.62 | 1 | 8 | 8 | 14 | 4.04 | 1 | 1 | 3.37 | 1 | 1 | 3.37 | 1 | 1 | | | | 3.37 | 1 | 1 | | | | | | | | | | 3.37 | 1 | 1 | 20.22 | 6 | 9 | 445 | 52.6 | 5.62 |
| Q96CV9 | 8.67 | 1 | 4 | 4 | 8 | 1.91 | 1 | 1 | 1.91 | 1 | 1 | | | | | | | | | | | | | 2.25 | 1 | 1 | 4.85 | 2 | 2 | | | | 6.07 | 3 | 3 | 577 | 65.9 | 5.24 |
| Q96CW1 | 13.10 | 1 | 5 | 5 | 11 | | | | | | | | | | 2.07 | 1 | 1 | | | | | | | | | | | | | | | | 13.10 | 5 | 10 | 435 | 49.6 | 9.54 |
| Q96D96 | 36.63 | 1 | 9 | 9 | 42 | 10.62 | 3 | 4 | 18.32 | 4 | 4 | 9.52 | 2 | 2 | | | | | | | | | | | | | | | | | | | 30.40 | 8 | 32 | 273 | 31.7 | 6.80 |
| P41227 | 20.43 | 1 | 1 | 4 | 6 | | | | | | | | | | | | | | | | | | | | | | | | | | | | 20.43 | 4 | 6 | 235 | 26.4 | 5.64 |
| Q96DA0 | 33.65 | 1 | 6 | 6 | 26 | 4.33 | 1 | 1 | | | | 22.12 | 3 | 3 | 4.33 | 1 | 1 | | | | | | | 4.33 | 1 | 1 | | | | | | | 33.65 | 6 | 20 | 208 | 22.7 | 7.39 |
| Q96DA6 | 18.97 | 1 | 2 | 2 | 2 | | | | | | | | | | | | | | | | | | | | | | | | | | | | 18.97 | 2 | 2 | 116 | 12.5 | 10.10 |
| Q96DB2 | 38.04 | 2 | 15 | 16 | 24 | | | | 3.17 | 1 | 1 | | | | 0.00 | | | | | | | | | | | | 0.00 | | | 0.00 | | | 38.04 | 16 | 23 | 347 | 39.2 | 7.65 |
| Q96DC9 | 50.00 | 1 | 12 | 12 | 29 | 5.56 | 1 | 1 | 8.97 | 2 | 2 | | | | | | | | | | | | | | | | | | | | | | 50.00 | 12 | 22 | 234 | 27.2 | 6.23 |
| Q96DE0 | 21.03 | 1 | 2 | 2 | 3 | | | | | | | | | | | | | | | | | | | | | | | | | | | | 21.03 | 2 | 3 | 195 | 21.3 | 6.89 |
| Q96DG6 | 54.69 | 1 | 12 | 12 | 32 | | | | | | | | | | 6.53 | 2 | 2 | | | | | | | 10.20 | 3 | 3 | | | | 8.98 | 2 | 2 | 54.69 | 12 | 25 | 245 | 28.0 | 7.18 |
| Q96DH6 | 9.45 | 1 | 2 | 2 | 3 | | | | | | | | | | | | | | | | | | | | | | | | | 4.88 | 1 | 1 | 4.57 | 1 | 2 | 328 | 35.2 | 8.48 |
| Q96DM3 | 4.11 | 1 | 2 | 2 | 4 | | | | | | | | | | | | | | | | | | | | | | | | | | | | 4.11 | 2 | 4 | 657 | 74.9 | 7.83 |
| Q96DY2 | 64.37 | 1 | 25 | 25 | 68 | | | | | | | | | | 10.24 | 5 | 5 | 26.50 | 10 | 10 | | | | | | | | | | | | | 54.79 | 22 | 53 | 449 | 52.3 | 9.25 |
| Q96DZ1 | 17.39 | 1 | 7 | 7 | 12 | 2.07 | 1 | 1 | 2.69 | 1 | 1 | | | | | | | | | | | | | | | | | | | | | | 17.39 | 7 | 10 | 483 | 54.8 | 6.28 |
| Q96E40 | 91.44 | 1 | 20 | 20 | 144 | 48.20 | 7 | 11 | 37.84 | 6 | 10 | 15.32 | 2 | 2 | 58.56 | 9 | 13 | 72.52 | 12 | 14 | | | | 5.86 | 1 | 1 | 5.86 | 1 | 1 | 9.91 | 2 | 2 | 68.92 | 14 | 90 | 222 | 25.1 | 9.13 |
| Q96E66 | 37.50 | 1 | 5 | 5 | 7 | | | | | | | | | | | | | | | | | | | | | | | | | | | | 37.50 | 5 | 7 | 192 | 22.2 | 9.35 |
| Q96EE3 | 23.89 | 1 | 7 | 7 | 13 | | | | | | | | | | | | | | | | | | | | | | | | | | | | 23.89 | 7 | 13 | 360 | 39.6 | 8.09 |
| Q96EE4 | 9.29 | 1 | 2 | 2 | 5 | 9.29 | 1 | 1 | 9.29 | 1 | 2 | | | | | | | | | | | | | 0.00 | | | | | | | | | 8.57 | 1 | 2 | 140 | 15.7 | 9.88 |
| Q96EK5 | 6.44 | 1 | 2 | 2 | 2 | | | | | | | | | | | | | | | | | | | | | | | | | | | | 6.44 | 2 | 2 | 621 | 71.8 | 5.49 |
| Q96EK6 | 40.22 | 1 | 4 | 4 | 8 | | | | | | | | | | | | | | | | | | | | | | | | | | | | 40.22 | 4 | 8 | 184 | 20.7 | 7.99 |
| Q96EP5 | 34.64 | 1 | 9 | 9 | 35 | 10.57 | 2 | 2 | 7.62 | 2 | 2 | 14.25 | 3 | 3 | 7.62 | 2 | 2 | 17.69 | 4 | 4 | | | | 3.69 | 1 | 1 | 11.55 | 3 | 3 | 3.69 | 1 | 1 | 24.32 | 6 | 17 | 407 | 43.4 | 8.56 |
| Q96ER3 | 22.57 | 1 | 6 | 6 | 9 | | | | | | | | | | | | | | | | | | | | | | | | | | | | 22.57 | 6 | 9 | 474 | 53.5 | 4.50 |
| Q96EY1 | 15.63 | 1 | 6 | 6 | 8 | | | | | | | 3.33 | 1 | 1 | | | | | | | | | | | | | | | | | | | 15.63 | 6 | 7 | 480 | 52.5 | 9.26 |
| Q96EY8 | 10.40 | 1 | 2 | 2 | 5 | | | | 10.40 | 2 | 2 | | | | | | | | | | | | | | | | | | | | | | 10.40 | 2 | 3 | 250 | 27.4 | 8.60 |
| Q96F07 | 12.99 | 1 | 4 | 15 | 21 | 0.94 | 1 | 1 | | | | | | | | | | | | | | | | | | | | | | | | | 12.99 | 15 | 20 | 1278 | 148.3 | 7.31 |
| Q96F10 | 36.47 | 1 | 5 | 5 | 7 | | | | | | | 0.00 | | | 0.00 | | | 0.00 | | | | | | 0.00 | | | 0.00 | | | 0.00 | | | 36.47 | 5 | 7 | 170 | 19.1 | 6.04 |
| Q96FC7 | 7.98 | 1 | 3 | 3 | 4 | | | | | | | | | | | | | | | | | | | | | | | | | | | | 7.98 | 3 | 4 | 376 | 42.5 | 6.42 |
| Q96FJ2 | 65.17 | 1 | 5 | 8 | 113 | 59.55 | 6 | 7 | 58.43 | 4 | 5 | 59.55 | 6 | 8 | 59.55 | 6 | 14 | 65.17 | 7 | 13 | | | | 51.69 | 5 | 10 | 59.55 | 5 | 8 | 59.55 | 5 | 12 | 64.04 | 6 | 36 | 89 | 10.3 | 7.37 |
| Q96FV2 | 19.53 | 1 | 5 | 5 | 6 | | | | | | | | | | | | | | | | | | | | | | | | | | | | 19.53 | 5 | 6 | 425 | 46.6 | 5.67 |
| Q96FW1 | 52.40 | 2 | 13 | 13 | 67 | 9.96 | 2 | 2 | 9.96 | 2 | 3 | | | | 16.97 | 4 | 4 | 4.43 | 1 | 1 | | | | 25.46 | 6 | 7 | 7.75 | 2 | 2 | 25.09 | 6 | 8 | 46.86 | 12 | 40 | 271 | 31.3 | 4.94 |
| Q96FZ7 | 17.41 | 1 | 3 | 3 | 5 | | | | | | | | | | 6.47 | 1 | 1 | | | | | | | | | | | | | | | | 17.41 | 3 | 4 | 201 | 23.5 | 5.31 |
| Q96CB9 | 2.08 | 1 | 1 | 1 | 6 | | | | | | | | | | | | | | | | | | | | | | | | | | | | 2.08 | 1 | 6 | 384 | 43.1 | 8.18 |
| Q96G75 | 4.83 | 1 | 2 | 2 | 2 | | | | | | | | | | | | | | | | | | | | | | | | | | | | 4.83 | 2 | 2 | 393 | 44.4 | 6.61 |
| Q96G97 | 4.02 | 1 | 2 | 2 | 3 | | | | | | | | | | | | | | | | | | | | | | | | | | | | 4.02 | 2 | 3 | 398 | 44.4 | 5.17 |
| Q96GA7 | 13.07 | 1 | 3 | 3 | 5 | | | | | | | | | | | | | | | | | | | | | | | | | | | | 13.07 | 3 | 5 | 329 | 34.7 | 6.89 |
| Q96GG9 | 21.62 | 1 | 5 | 5 | 9 | 13.90 | 3 | 4 | 2.70 | 1 | 1 | | | | | | | | | | | | | | | | | | | | | | 15.06 | 4 | 4 | 259 | 30.1 | 5.34 |
| Q96GK7 | 45.54 | 1 | 3 | 12 | 30 | 27.71 | 6 | 8 | 13.06 | 3 | 3 | 23.57 | 6 | 7 | | | | | | | | | | | | | | | | | | | 39.49 | 10 | 12 | 314 | 34.6 | 8.24 |
| Q96GQ5 | 13.46 | 1 | 5 | 5 | 7 | | | | | | | | | | | | | | | | | | | | | | | | | | | | 13.46 | 5 | 7 | 468 | 51.0 | 6.93 |
| Q96GX9 | 19.83 | 1 | 3 | 3 | 4 | | | | | | | | | | | | | | | | | | | | | | | | | | | | 19.83 | 3 | 4 | 242 | 27.1 | 7.12 |
| Q96H20 | 8.14 | 1 | 2 | 2 | 6 | | | | | | | | | | | | | | | | | | | | | | | | | 8.14 | 2 | 2 | 8.14 | 2 | 4 | 258 | 28.8 | 6.65 |
| P20023 | 0.68 | 1 | 1 | 1 | 6 | | | | | | | | | | | | | | | | | | | | | | | | | | | | 0.68 | 1 | 6 | 1033 | 112.8 | 7.52 |
| Q96HA9 | 44.81 | 1 | 9 | 9 | 19 | | | | | | | | | | | | | | | | | | | | | | | | | | | | 44.81 | 9 | 19 | 241 | 26.6 | 9.13 |
| Q96HC4 | 12.92 | 1 | 8 | 8 | 22 | 1.51 | 1 | 1 | | | | | | | 3.52 | 2 | 2 | | | | | | | 3.52 | 2 | 2 | | | | 7.55 | 4 | 4 | 10.74 | 7 | 13 | 596 | 63.9 | 8.21 |
| Q96HD1 | 11.43 | 1 | 4 | 4 | 4 | | | | | | | | | | | | | | | | | | | | | | | | | | | | 11.43 | 4 | 4 | 420 | 45.4 | 4.87 |
| Q96HE7 | 9.40 | 1 | 3 | 4 | 8 | 7.05 | 3 | 3 | 2.35 | 1 | 1 | 5.34 | 2 | 2 | | | | | | | | | | | | | | | | | | | 5.34 | 2 | 2 | 468 | 54.4 | 5.68 |
| Q96HR9 | 23.22 | 1 | 6 | 6 | 27 | 19.43 | 4 | 6 | 15.64 | 3 | 4 | | | | | | | | | | | | | | | | | | | | | | 23.22 | 6 | 17 | 211 | 23.4 | 8.56 |
| Q96HS1 | 29.41 | 2 | 9 | 9 | 12 | 3.46 | 1 | 1 | 3.46 | 1 | 1 | 4.50 | 1 | 1 | | | | | | | | | | | | | | | | | | | 29.07 | 8 | 9 | 289 | 32.0 | 8.68 |
| Q96HV5 | 8.33 | 1 | 2 | 2 | 2 | | | | | | | | | | | | | | | | | | | | | | | | | | | | 8.33 | 2 | 2 | 264 | 29.6 | 9.03 |
| Q96HY6 | 14.97 | 1 | 3 | 3 | 5 | | | | | | | | | | | | | | | | | | | | | | | | | 4.78 | 1 | 1 | 10.19 | 2 | 4 | 314 | 35.6 | 5.12 |
| Q96I99 | 25.46 | 1 | 11 | 11 | 14 | 4.40 | 2 | 2 | 6.94 | 3 | 3 | 5.09 | 2 | 2 | | | | | | | | | | | | | | | | 2.08 | 1 | 1 | 14.12 | 6 | 6 | 432 | 46.5 | 6.39 |
| Q96IG2 | 5.73 | 1 | 2 | 2 | 2 | | | | | | | | | | | | | | | | | | | | | | | | | | | | 5.73 | 2 | 2 | 436 | 48.4 | 7.49 |
| Q96IJ6 | 31.90 | 1 | 10 | 10 | 23 | 3.33 | 1 | 1 | 3.33 | 1 | 1 | 3.33 | 1 | 1 | 3.33 | 1 | 1 | | | | | | | | | | | | | 6.90 | 2 | 2 | 31.90 | 10 | 17 | 420 | 46.3 | 7.21 |
| Q96IU4 | 46.19 | 1 | 8 | 8 | 43 | | | | | | | 4.76 | 1 | 1 | 24.76 | 4 | 4 | | | | 3.33 | 1 | 1 | 15.24 | 2 | 2 | 15.24 | 2 | 4 | 15.24 | 2 | 2 | 46.19 | 8 | 29 | 210 | 22.3 | 6.40 |
| Q96IV0 | 15.44 | 1 | 10 | 10 | 19 | 3.36 | 2 | 2 | 1.68 | 1 | 1 | | | | | | | | | | | | | | | | | | | | | | 15.44 | 10 | 16 | 654 | 74.3 | 6.89 |
| Q96IX5 | 46.55 | 1 | 4 | 4 | 9 | 25.86 | 1 | 1 | 25.86 | 1 | 1 | | | | | | | 27.59 | 1 | 1 | | | | | | | | | | | | | 46.55 | 4 | 6 | 58 | 6.5 | 9.76 |
| Q96J02 | 3.32 | 2 | 2 | 2 | 2 | | | | | | | | | | | | | | | | | | | | | | | | | | | | 3.32 | 2 | 2 | 903 | 102.7 | 6.30 |
| Q96J94 | 5.23 | 1 | 3 | 3 | 5 | | | | | | | | | | | | | | | | | | | | | | | | | | | | 5.23 | 3 | 5 | 861 | 98.5 | 9.44 |
| Q96JB1 | 55.23 | 1 | 240 | 263 | 1136 | 10.76 | 40 | 41 | 9.49 | 36 | 38 | 12.54 | 45 | 47 | 8.06 | 33 | 35 | 7.64 | 33 | 36 | | | | | | | | | | 0.45 | 2 | 2 | 53.21 | 253 | 937 | 4490 | 514.3 | 6.32 |
| Q9P2P6 | 0.51 | 3 | 1 | 3 | 5 | 0.00 | | | 0.00 | | | 0.00 | | | | | | | | | | | | | | | | | | | | | 0.51 | 3 | 5 | 4700 | 516.0 | 6.32 |
| Q96JB2 | 12.56 | 1 | 6 | 6 | 15 | | | | | | | | | | | | | | | | | | | | | | | | | | | | 12.56 | 6 | 15 | 828 | 94.0 | 5.57 |
| Q96JC1 | 1.92 | 1 | 2 | 2 | 2 | | | | | | | | | | | | | | | | | | | | | | | | | | | | 1.92 | 2 | 2 | 886 | 101.7 | 6.99 |
| Q96JN2 | 50.43 | 1 | 50 | 50 | 206 | 9.10 | 6 | 8 | 14.04 | 11 | 12 | 14.47 | 12 | 15 | 2.08 | 2 | 2 | 9.36 | 8 | 10 | | | | 2.43 | 2 | 2 | 3.12 | 3 | 3 | 1.21 | 1 | 1 | 47.40 | 47 | 153 | 1154 | 134.0 | 4.65 |
| Q96JQ2 | 35.03 | 1 | 28 | 28 | 76 | 5.79 | 4 | 4 | 12.08 | 9 | 9 | | | | | | | 2.89 | 2 | 2 | | | | | | | 4.89 | 4 | 4 | | | | 26.25 | 19 | 57 | 1002 | 111.6 | 4.94 |
| Q96JY0 | 5.99 | 1 | 2 | 2 | 3 | | | | | | | | | | | | | | | | | | | 2.07 | 1 | 1 | | | | | | | 5.99 | 2 | 2 | 434 | 49.2 | 8.69 |
| Q96K19 | 11.24 | 1 | 2 | 2 | 2 | | | | | | | | | | | | | | | | | | | | | | | | | | | | 11.24 | 2 | 2 | 258 | 29.8 | 5.58 |
| Q96K76 | 6.11 | 1 | 7 | 7 | 9 | | | | | | | | | | | | | | | | | | | | | | | | | 0.00 | | | 6.11 | 7 | 9 | 1375 | 157.2 | 5.08 |
| Q96KC9 | 16.96 | 1 | 7 | 7 | 24 | 3.54 | 1 | 1 | 6.33 | 2 | 3 | | | | 3.54 | 1 | 1 | 10.63 | 3 | 3 | | | | 6.33 | 2 | 2 | 16.96 | 7 | 11 | 7.34 | 2 | 2 | 3.54 | 1 | 1 | 395 | 43.0 | 4.21 |
| O75923 | 2.12 | 1 | 1 | 4 | 5 | 0.77 | 1 | 1 | | | | | | | | | | | | | | | | | | | | | | | | | 1.35 | 3 | 4 | 2080 | 237.1 | 5.64 |
| Q96KK5 | 35.94 | 10 | 3 | 5 | 43 | 32.81 | 4 | 5 | 32.81 | 4 | 7 | 32.81 | 4 | 6 | 17.97 | 3 | 4 | 17.97 | 2 | 2 | | | | 10.94 | 2 | 2 | | | | 10.94 | 1 | 2 | 35.94 | 4 | 15 | 128 | 13.9 | 10.89 |
| Q96KN8 | 16.13 | 1 | 3 | 3 | 5 | | | | | | | | | | | | | | | | | | | | | | | | | | | | 16.13 | 3 | 5 | 279 | 30.3 | 8.24 |
| Q96KP1 | 7.36 | 1 | 6 | 6 | 7 | | | | | | | | | | | | | | | | | | | | | | | | | | | | 7.36 | 6 | 7 | 924 | 104.0 | 6.90 |
| Q96KP4 | 76.00 | 1 | 34 | 34 | 239 | 33.89 | 12 | 14 | 24.63 | 9 | 9 | 33.26 | 12 | 13 | 36.21 | 16 | 22 | 20.63 | 8 | 11 | | | | 19.37 | 8 | 8 | 22.11 | 8 | 12 | 32.63 | 13 | 18 | 73.89 | 28 | 132 | 475 | 52.8 | 5.97 |
| Q96KW9 | 17.44 | 1 | 2 | 2 | 7 | 9.74 | 1 | 3 | 17.44 | 2 | 3 | | | | 7.69 | 1 | 1 | | | | | | | | | | | | | | | | | | | 195 | 21.5 | 4.78 |
| Q96KX0 | 47.26 | 1 | 6 | 6 | 21 | 40.41 | 5 | 5 | 23.29 | 3 | 4 | 22.60 | 3 | 3 | | | | | | | | | | | | | | | | | | | 47.26 | 6 | 9 | 146 | 16.4 | 8.28 |
| Q96KX2 | 52.51 | 1 | 16 | 16 | 24 | | | | | | | | | | | | | 2.68 | 1 | 1 | | | | | | | | | | | | | 52.17 | 15 | 23 | 299 | 35.0 | 7.69 |
| P20702 | 1.72 | 2 | 1 | 2 | 5 | 0.95 | 1 | 1 | | | | 0.95 | 1 | 1 | | | | | | | | | | | | | | | | | | | 1.72 | 2 | 3 | 1163 | 127.7 | 6.64 |
| Q96L03 | 26.04 | 1 | 9 | 10 | 13 | | | | | | | | | | | | | | | | | | | | | | | | | | | | 26.04 | 10 | 13 | 361 | 43.5 | 9.86 |
| Q96L21 | 43.46 | 1 | 3 | 8 | 23 | 2.80 | 1 | 1 | 2.80 | 1 | 1 | | | | 2.80 | 1 | 1 | | | | | | | | | | | | | | | | 43.46 | 8 | 20 | 214 | 24.5 | 10.01 |
| Q96LB3 | 15.33 | 1 | 8 | 8 | 15 | | | | | | | | | | | | | 3.67 | 2 | 2 | | | | 4.17 | 2 | 2 | 2.00 | 1 | 1 | 7.33 | 4 | 4 | 6.33 | 3 | 6 | 600 | 69.2 | 6.00 |
| Q96LD8 | 27.83 | 1 | 4 | 4 | 8 | | | | | | | | | | | | | | | | | | | | | | | | | | | | 27.83 | 4 | 8 | 212 | 24.1 | 5.97 |
| Q96LI6 | 20.45 | 1 | 8 | 8 | 11 | | | | | | | | | | | | | | | | | | | | | | | | | | | | 20.45 | 8 | 11 | 401 | 45.1 | 7.18 |
| Q96LJ7 | 13.10 | 1 | 3 | 3 | 4 | | | | | | | | | | | | | | | | | | | | | | | | | | | | 13.10 | 3 | 4 | 313 | 33.9 | 7.83 |
| Q96LK8 | 35.94 | 1 | 11 | 11 | 47 | 2.34 | 1 | 1 | 2.34 | 1 | 1 | | | | 9.11 | 3 | 4 | 15.10 | 4 | 5 | | | | 2.34 | 1 | 1 | 9.90 | 4 | 5 | 6.25 | 2 | 3 | 32.03 | 9 | 27 | 384 | 42.3 | 4.77 |
| Q96LM5 | 39.25 | 1 | 6 | 6 | 32 | | | | | | | | | | 36.02 | 5 | 6 | 25.27 | 4 | 5 | 8.06 | 1 | 1 | 13.98 | 2 | 2 | 8.06 | 1 | 1 | 8.06 | 1 | 1 | 39.25 | 6 | 16 | 186 | 21.7 | 9.96 |
| Q96LM6 | 68.33 | 1 | 14 | 14 | 42 | | | | 5.00 | 1 | 1 | | | | 32.22 | 5 | 6 | 58.89 | 10 | 16 | 16.11 | 1 | 1 | | | | | | | | | | 57.22 | 9 | 18 | 180 | 20.6 | 7.55 |
| Q96LP2 | 19.69 | 1 | 7 | 7 | 14 | | | | | | | | | | | | | | | | | | | | | | | | | | | | 19.69 | 7 | 14 | 452 | 52.0 | 9.10 |
| Q96LQ0 | 9.00 | 1 | 3 | 3 | 4 | | | | | | | | | | | | | | | | | | | | | | | | | | | | 9.00 | 3 | 4 | 422 | 49.4 | 9.36 |
| Q96LU5 | 10.84 | 1 | 2 | 2 | 2 | | | | | | | | | | | | | | | | | | | | | | | | | | | | 10.84 | 2 | 2 | 166 | 18.5 | 8.09 |
| Q96LZ3 | 39.41 | 1 | 5 | 5 | 7 | 10.59 | 1 | 2 | 15.29 | 2 | 2 | | | | | | | | | | | | | | | | | | | | | | 22.94 | 3 | 3 | 170 | 19.5 | 4.89 |
| Q96M20 | 7.47 | 1 | 3 | 3 | 4 | | | | | | | | | | | | | | | | | | | | | | | | | | | | 7.47 | 3 | 4 | 576 | 67.5 | 9.26 |
| Q96M29 | 88.66 | 1 | 52 | 52 | 416 | 24.54 | 11 | 12 | 27.63 | 12 | 12 | 7.42 | 3 | 4 | 53.81 | 23 | 31 | 74.23 | 31 | 46 | | | | 21.24 | 10 | 12 | 36.49 | 17 | 20 | 46.19 | 22 | 30 | 79.18 | 38 | 249 | 485 | 56.3 | 7.14 |
| Q96M32 | 43.15 | 1 | 29 | 29 | 114 | 4.84 | 2 | 2 | 3.60 | 2 | 2 | 2.35 | 1 | 1 | 14.66 | 8 | 10 | 22.41 | 13 | 14 | | | | | | | 2.63 | 1 | 1 | | | | 40.80 | 24 | 84 | 723 | 82.6 | 4.74 |
| Q96M34 | 12.13 | 1 | 6 | 6 | 12 | 5.60 | 2 | 2 | 2.99 | 1 | 1 | 2.61 | 1 | 1 | 2.05 | 1 | 1 | 6.53 | 4 | 4 | | | | | | | 1.68 | 1 | 1 | | | | 3.73 | 2 | 2 | 536 | 60.1 | 5.80 |
| Q96JB5 | 3.95 | 2 | 1 | 2 | 5 | | | | 1.19 | 1 | 1 | | | | 1.19 | 1 | 1 | | | | | | | | | | | | | | | | 3.95 | 2 | 3 | 506 | 56.9 | 4.75 |
| Q96M63 | 9.25 | 1 | 5 | 5 | 8 | | | | | | | | | | | | | | | | | | | | | | | | | | | | 9.25 | 5 | 8 | 670 | 75.0 | 6.28 |
| Q96M69 | 17.45 | 1 | 12 | 12 | 24 | | | | | | | | | | | | | | | | | | | | | | | | | | | | 17.45 | 12 | 24 | 825 | 93.6 | 6.43 |
| Q96M83 | 5.63 | 1 | 7 | 7 | 13 | | | | | | | | | | | | | 0.79 | 1 | 1 | | | | | | | | | | | | | 5.63 | 7 | 12 | 1385 | 157.4 | 7.58 |
| Q96M86 | 0.78 | 1 | 3 | 3 | 4 | | | | | | | | | | | | | | | | | | | | | | | | | | | | 0.78 | 3 | 4 | 4753 | 533.3 | 6.71 |
| Q96M91 | 65.37 | 1 | 45 | 45 | 117 | 6.03 | 3 | 3 | 1.36 | 1 | 1 | | | | 35.80 | 18 | 21 | 49.42 | 26 | 37 | | | | | | | 0.00 | | | 1.75 | 1 | 1 | 50.97 | 28 | 54 | 514 | 61.8 | 8.90 |
| O75674 | 6.30 | 1 | 1 | 2 | 5 | | | | 1.26 | 1 | 1 | | | | | | | | | | | | | | | | | | | 5.04 | 1 | 1 | 1.26 | 1 | 3 | 476 | 53.0 | 5.24 |
| Q96M95 | 54.43 | 1 | 15 | 15 | 33 | | | | | | | | | | 9.18 | 2 | 3 | 13.29 | 3 | 3 | | | | | | | | | | | | | 54.43 | 15 | 27 | 316 | 38.0 | 9.16 |
| Q96M98 | 63.18 | 1 | 17 | 17 | 79 | 13.51 | 3 | 4 | 21.28 | 5 | 7 | 4.39 | 1 | 1 | 29.39 | 6 | 9 | 38.85 | 9 | 12 | | | | | | | | | | | | | 47.64 | 12 | 46 | 296 | 33.3 | 8.21 |
| Q96MA6 | 55.95 | 1 | 27 | 27 | 96 | 3.13 | 1 | 1 | 10.23 | 4 | 5 | | | | 31.11 | 14 | 16 | 10.65 | 4 | 5 | | | | 3.55 | 2 | 2 | | | | 5.01 | 2 | 2 | 52.61 | 24 | 65 | 479 | 54.9 | 6.15 |
| Q96MC2 | 51.76 | 1 | 40 | 40 | 102 | | | | | | | | | | 12.84 | 10 | 10 | 26.22 | 17 | 20 | | | | | | | | | | | | | 46.08 | 32 | 72 | 740 | 87.1 | 5.38 |
| Q96MR6 | 52.56 | 1 | 76 | 76 | 278 | 3.92 | 4 | 4 | 2.88 | 3 | 3 | 6.48 | 7 | 7 | 23.60 | 28 | 38 | 23.60 | 30 | 35 | | | | | | | | | | | | | 48.32 | 63 | 191 | 1250 | 144.9 | 5.80 |
| Q96MT7 | 30.45 | 1 | 27 | 27 | 60 | | | | | | | 1.12 | 1 | 1 | 5.30 | 5 | 5 | 2.75 | 3 | 4 | | | | | | | | | | | | | 29.53 | 26 | 50 | 982 | 111.7 | 4.84 |
| Q96MW5 | 13.40 | 1 | 5 | 5 | 7 | | | | | | | | | | | | | | | | | | | | | | | | | | | | 13.40 | 5 | 7 | 612 | 68.4 | 5.20 |
| Q96N03 | 11.76 | 1 | 2 | 2 | 3 | | | | | | | | | | | | | | | | | | | | | | | | | | | | 11.76 | 2 | 3 | 204 | 22.3 | 8.57 |
| Q96N06 | 43.88 | 1 | 5 | 5 | 9 | 10.79 | 1 | 1 | 17.99 | 2 | 2 | | | | | | | | | | | | | | | | | | | | | | 33.09 | 4 | 6 | 139 | 15.5 | 9.22 |
| Q96N23 | 12.76 | 1 | 35 | 35 | 60 | | | | | | | | | | | | | | | | | | | | | | | | | | | | 12.76 | 35 | 60 | 3096 | 351.7 | 8.10 |
| Q9HC10 | 0.60 | 2 | 1 | 2 | 5 | | | | | | | 0.30 | 1 | 1 | 0.30 | 1 | 1 | | | | | | | 0.30 | 1 | 1 | | | | | | | 0.30 | 1 | 2 | 1997 | 226.6 | 5.69 |
| Q96N66 | 14.83 | 1 | 6 | 6 | 14 | | | | | | | | | | | | | | | | | | | | | | | | | | | | 14.83 | 6 | 14 | 472 | 52.7 | 8.97 |
| Q96ND0 | 14.71 | 1 | 3 | 3 | 3 | | | | | | | | | | | | | | | | | | | | | | | | | | | | 14.71 | 3 | 3 | 272 | 30.8 | 9.76 |
| Q96NG3 | 48.51 | 1 | 34 | 34 | 117 | 12.50 | 7 | 7 | 13.39 | 8 | 9 | 2.83 | 2 | 2 | 14.88 | 9 | 11 | 33.18 | 21 | 25 | | | | | | | | | | | | | 39.73 | 21 | 63 | 672 | 76.6 | 5.60 |
| Q96P26 | 43.93 | 2 | 24 | 24 | 77 | 7.54 | 4 | 4 | 5.90 | 3 | 3 | | | | 15.41 | 7 | 7 | 6.07 | 3 | 3 | | | | | | | | | | | | | 39.18 | 22 | 60 | 610 | 68.8 | 8.82 |
| Q96P56 | 14.34 | 1 | 6 | 6 | 12 | | | | | | | | | | | | | | | | | | | | | | | | | | | | 14.34 | 6 | 12 | 530 | 62.0 | 7.30 |
| Q96P70 | 12.49 | 1 | 11 | 11 | 21 | | | | | | | | | | | | | | | | | | | | | | | | | | | | 12.49 | 11 | 21 | 1041 | 115.9 | 4.81 |
| Q96PF2 | 37.15 | 1 | 11 | 11 | 33 | 5.87 | 2 | 2 | | | | 3.07 | 1 | 1 | 3.07 | 1 | 1 | 3.07 | 1 | 1 | | | | | | | | | | | | | 37.15 | 11 | 28 | 358 | 40.9 | 8.84 |
| Q96PK6 | 6.43 | 1 | 3 | 3 | 5 | | | | | | | | | | | | | | | | | | | | | | | | | | | | 6.43 | 3 | 5 | 669 | 69.4 | 9.67 |
| Q96PP4 | 8.73 | 1 | 2 | 2 | 3 | | | | | | | | | | | | | | | | | | | | | | | | | | | | 8.73 | 2 | 3 | 275 | 31.8 | 10.07 |
| Q96PU5 | 12.21 | 2 | 10 | 10 | 19 | | | | | | | | | | | | | | | | | | | | | | | | | | | | 12.21 | 10 | 19 | 975 | 111.9 | 5.82 |
| Q96PU9 | 79.13 | 1 | 20 | 20 | 184 | 28.74 | 6 | 6 | 27.95 | 6 | 9 | 13.78 | 3 | 3 | 64.96 | 14 | 18 | 72.44 | 16 | 29 | 5.51 | 1 | 1 | 35.04 | 8 | 9 | 31.10 | 7 | 10 | 55.51 | 13 | 17 | 77.95 | 19 | 82 | 254 | 27.7 | 9.89 |
| Q96QE4 | 43.29 | 1 | 28 | 31 | 274 | 24.71 | 16 | 21 | 24.29 | 17 | 25 | 15.63 | 11 | 13 | 5.91 | 4 | 5 | 9.50 | 7 | 8 | | | | | | | | | | | | | 38.01 | 26 | 202 | 947 | 105.5 | 4.91 |
| Q6ZU65 | 0.45 | 1 | 1 | 1 | 5 | | | | | | | | | | 0.45 | 1 | 1 | | | | | | | | | | | | | | | | 0.45 | 1 | 4 | 1347 | 146.0 | 9.19 |
| Q96QF0 | 10.71 | 1 | 3 | 3 | 7 | | | | 0.00 | | | | | | | | | | | | | | | | | | | | | | | | 10.71 | 3 | 7 | 476 | 53.0 | 5.97 |
| Q96QG7 | 4.74 | 1 | 3 | 3 | 3 | | | | | | | | | | | | | | | | | | | | | | | | | | | | 4.74 | 3 | 3 | 549 | 63.4 | 6.39 |
| Q96QH8 | 35.22 | 1 | 5 | 5 | 28 | 20.75 | 3 | 7 | 12.58 | 2 | 3 | 20.75 | 3 | 5 | 20.75 | 3 | 3 | 8.18 | 1 | 3 | | | | | | | | | | | | | 35.22 | 5 | 7 | 159 | 17.9 | 6.42 |
| Q96QK1 | 38.19 | 3 | 27 | 27 | 131 | 1.76 | 1 | 1 | 3.02 | 2 | 2 | 4.52 | 3 | 4 | 6.91 | 5 | 5 | 1.63 | 1 | 1 | | | | 2.39 | 1 | 2 | | | | 5.40 | 3 | 3 | 38.19 | 27 | 113 | 796 | 91.6 | 5.49 |
| Q96QR8 | 21.15 | 3 | 3 | 5 | 10 | | | | | | | | | | | | | | | | | | | | | | | | | 4.17 | 1 | 1 | 21.15 | 5 | 9 | 312 | 33.2 | 5.43 |
| Q96R05 | 32.09 | 1 | 4 | 4 | 4 | | | | | | | | | | | | | | | | | | | | | | | | | | | | 32.09 | 4 | 4 | 134 | 15.5 | 7.80 |
| Q96RG2 | 2.72 | 1 | 2 | 2 | 3 | | | | | | | | | | | | | | | | | | | | | | | | | | | | 2.72 | 2 | 3 | 1323 | 142.8 | 4.86 |
| P54252 | 5.22 | 1 | 1 | 2 | 5 | | | | | | | | | | | | | | | | | | | 3.30 | 1 | 1 | | | | | | | 5.22 | 2 | 4 | 364 | 41.8 | 4.91 |
| Q96RL7 | 20.48 | 1 | 56 | 56 | 102 | 0.72 | 2 | 2 | 3.53 | 9 | 9 | 2.08 | 5 | 5 | | | | | | | | | | | | | | | | | | | 19.03 | 52 | 86 | 3174 | 360.0 | 6.33 |
| Q96RN1 | 12.58 | 1 | 11 | 11 | 29 | 3.51 | 2 | 2 | 2.06 | 1 | 2 | 1.44 | 1 | 1 | | | | | | | | | | | | | | | | | | | 10.41 | 9 | 20 | 970 | 108.9 | 6.24 |
| Q96RQ1 | 38.46 | 1 | 10 | 10 | 31 | 7.96 | 3 | 3 | 11.41 | 4 | 5 | 5.84 | 2 | 2 | | | | | | | | | | | | | | | | | | | 38.46 | 10 | 21 | 377 | 42.5 | 6.77 |
| Q96RQ3 | 6.90 | 1 | 3 | 3 | 3 | | | | | | | | | | | | | | | | | | | | | | | | | | | | 6.90 | 3 | 3 | 725 | 80.4 | 7.78 |
| Q96RQ9 | 46.21 | 1 | 24 | 24 | 72 | 28.92 | 12 | 14 | 35.63 | 17 | 20 | | | | 2.65 | 1 | 1 | 1.41 | 1 | 1 | | | | | | | | | | | | | 28.22 | 14 | 36 | 567 | 62.8 | 8.68 |
| Q96RR4 | 3.40 | 1 | 2 | 2 | 2 | | | | | | | | | | | | | | | | | | | | | | | | | 1.53 | 1 | 1 | 1.87 | 1 | 1 | 588 | 64.7 | 6.68 |
| Q96RS6 | 5.32 | 1 | 3 | 3 | 5 | | | | | | | | | | | | | | | | | | | | | | | | | | | | 5.32 | 3 | 5 | 583 | 66.7 | 5.11 |
| Q96RT6 | 8.19 | 1 | 2 | 6 | 11 | 1.74 | 2 | 2 | 0.81 | 1 | 1 | 0.81 | 1 | 1 | | | | 1.61 | 1 | 1 | | | | | | | | | | | | | 4.83 | 3 | 6 | 745 | 85.2 | 6.16 |
| Q96RY7 | 15.46 | 1 | 19 | 19 | 26 | | | | | | | | | | | | | | | | | | | | | | | | | | | | 15.46 | 19 | 26 | 1462 | 165.1 | 6.04 |
| Q96S53 | 7.53 | 1 | 4 | 4 | 7 | | | | | | | | | | | | | | | | | | | | | | | | | | | | 7.53 | 4 | 7 | 571 | 63.6 | 7.06 |
| Q96S59 | 5.49 | 1 | 3 | 3 | 4 | | | | | | | | | | | | | | | | | | | | | | | | | | | | 5.49 | 3 | 4 | 729 | 77.8 | 6.79 |
| Q96S96 | 44.05 | 1 | 6 | 6 | 16 | 22.47 | 4 | 4 | 22.91 | 4 | 4 | | | | | | | 4.41 | 1 | 1 | | | | | | | | | | | | | 29.52 | 4 | 7 | 227 | 25.7 | 6.54 |
| Q96S97 | 19.57 | 1 | 5 | 5 | 23 | 10.25 | 2 | 2 | 7.14 | 1 | 1 | 7.14 | 1 | 2 | 7.14 | 1 | 1 | | | | | | | | | | | | | | | | 19.57 | 5 | 17 | 322 | 35.3 | 8.15 |
| Q96SB4 | 5.34 | 3 | 3 | 3 | 6 | | | | 1.37 | 1 | 1 | | | | | | | | | | | | | | | | | | | | | | 5.34 | 3 | 5 | 655 | 74.3 | 6.16 |
| Q96SF2 | 14.00 | 2 | 5 | 5 | 11 | | | | 3.05 | 1 | 2 | | | | | | | | | | | | | | | | | | | | | | 14.00 | 5 | 9 | 557 | 59.3 | 5.72 |
| Q96SK2 | 30.30 | 1 | 13 | 13 | 27 | 2.14 | 1 | 1 | 5.88 | 3 | 3 | | | | | | | | | | | | | | | | | | | | | | 30.30 | 13 | 23 | 561 | 62.9 | 8.63 |
| Q96SU4 | 3.94 | 1 | 2 | 3 | 3 | | | | | | | 0.00 | | | | | | | | | | | | | | | | | | | | | 3.94 | 3 | 3 | 736 | 83.1 | 6.18 |
| Q96T17 | 3.96 | 1 | 2 | 2 | 2 | | | | | | | | | | | | | | | | | | | | | | | | | | | | 3.96 | 2 | 2 | 732 | 81.9 | 8.84 |
| Q96T51 | 6.78 | 1 | 4 | 4 | 4 | | | | | | | 0.00 | | | 0.00 | | | | | | | | | | | | | | | | | | 6.78 | 4 | 4 | 708 | 79.8 | 5.74 |
| Q96T52 | 14.29 | 1 | 2 | 2 | 3 | | | | | | | | | | | | | | | | | | | | | | | | | | | | 14.29 | 2 | 3 | 175 | 19.7 | 9.36 |
| Q96T59 | 50.00 | 1 | 10 | 10 | 16 | 15.96 | 2 | 2 | 30.85 | 4 | 4 | | | | | | | | | | | | | | | | | | | | | | 50.00 | 8 | 10 | 188 | 20.6 | 8.05 |
| Q96T76 | 23.40 | 4 | 18 | 18 | 38 | | | | | | | | | | | | | | | | | | | | | | | | | | | | 23.40 | 18 | 38 | 1030 | 113.2 | 6.35 |
| Q96TA1 | 23.06 | 1 | 14 | 14 | 25 | | | | | | | | | | | | | | | | | | | | | | | | | | | | 23.06 | 14 | 25 | 746 | 84.1 | 6.19 |
| Q99417 | 73.79 | 1 | 8 | 8 | 31 | 56.31 | 5 | 5 | 50.49 | 5 | 8 | 30.10 | 2 | 2 | 66.02 | 5 | 5 | 31.07 | 4 | 4 | | | | | | | | | | | | | 65.05 | 6 | 7 | 103 | 12.0 | 5.91 |
| Q99426 | 49.59 | 1 | 11 | 11 | 41 | | | | 8.20 | 2 | 2 | | | | 3.69 | 1 | 1 | 23.36 | 4 | 7 | | | | 8.61 | 2 | 3 | 25.00 | 6 | 7 | 22.54 | 4 | 4 | 38.52 | 8 | 17 | 244 | 27.3 | 5.15 |
| Q99436 | 48.38 | 1 | 11 | 12 | 33 | 29.24 | 5 | 6 | 21.30 | 5 | 6 | 14.44 | 4 | 5 | | | | 3.25 | 1 | 1 | | | | | | | | | | | | | 47.29 | 11 | 15 | 277 | 29.9 | 7.68 |
| Q99442 | 13.78 | 1 | 7 | 7 | 16 | | | | 5.26 | 2 | 2 | 3.26 | 1 | 1 | | | | | | | | | | | | | | | | | | | 13.53 | 6 | 13 | 399 | 45.8 | 7.12 |
| Q99447 | 64.78 | 1 | 24 | 24 | 82 | 22.37 | 6 | 7 | 29.31 | 8 | 10 | 6.94 | 2 | 2 | 11.31 | 3 | 3 | 13.62 | 4 | 4 | | | | 10.03 | 3 | 3 | 19.54 | 6 | 8 | 19.02 | 6 | 6 | 51.93 | 19 | 39 | 389 | 43.8 | 6.92 |
| Q99460 | 56.24 | 1 | 41 | 41 | 223 | 11.96 | 7 | 9 | 10.18 | 7 | 9 | 13.43 | 8 | 8 | 8.39 | 7 | 7 | 14.38 | 9 | 9 | | | | | | | | | | | | | 56.24 | 40 | 181 | 953 | 105.8 | 5.39 |
| Q99471 | 26.62 | 1 | 3 | 3 | 5 | 11.69 | 1 | 1 | | | | | | | | | | | | | | | | | | | | | | | | | 26.62 | 3 | 4 | 154 | 17.3 | 6.33 |
| Q99487 | 28.06 | 1 | 7 | 7 | 8 | | | | | | | | | | | | | | | | | | | | | | | | | | | | 28.06 | 7 | 8 | 392 | 44.0 | 6.89 |
| Q99497 | 77.78 | 1 | 16 | 16 | 68 | 55.03 | 10 | 13 | 58.73 | 9 | 11 | 46.56 | 6 | 8 | | | | 12.17 | 2 | 2 | | | | | | | | | | 3.70 | 1 | 1 | 76.72 | 14 | 33 | 189 | 19.9 | 6.79 |
| Q99519 | 16.87 | 1 | 6 | 6 | 10 | | | | 2.41 | 1 | 1 | 3.37 | 1 | 1 | | | | | | | | | | | | | | | | | | | 14.46 | 5 | 8 | 415 | 45.4 | 5.88 |
| Q99523 | 16.85 | 1 | 12 | 12 | 19 | 1.20 | 1 | 1 | 1.20 | 1 | 1 | 9.03 | 5 | 5 | | | | | | | | | | | | | | | | | | | 13.36 | 8 | 12 | 831 | 92.0 | 5.74 |
| Q9UKX3 | 0.83 | 6 | 1 | 2 | 5 | | | | | | | | | | | | | | | | | | | | | | | | | | | | 0.83 | 2 | 5 | 1938 | 223.5 | 5.68 |
| P12883 | 0.78 | 7 | 1 | 2 | 5 | | | | | | | | | | | | | | | | | | | | | | | | | | | | 0.78 | 2 | 5 | 1935 | 223.0 | 5.80 |
| Q99536 | 56.23 | 1 | 18 | 18 | 65 | 19.59 | 6 | 8 | 22.14 | 6 | 7 | 18.32 | 5 | 5 | 8.91 | 3 | 3 | 4.83 | 2 | 2 | | | | | | | 2.29 | 1 | 1 | 7.38 | 2 | 2 | 55.22 | 17 | 37 | 393 | 41.9 | 6.29 |
| Q99549 | 1.51 | 1 | 2 | 2 | 3 | | | | | | | | | | | | | | | | | | | | | | | | | | | | 1.51 | 2 | 3 | 860 | 97.1 | 6.06 |
| Q99567 | 24.29 | 1 | 12 | 13 | 28 | | | | | | | | | | | | | 1.75 | 1 | 1 | | | | | | | | | | | | | 24.29 | 13 | 27 | 741 | 83.5 | 5.69 |
| Q99571 | 12.63 | 1 | 3 | 3 | 4 | | | | | | | | | | | | | | | | | | | | | | | | | | | | 12.63 | 3 | 4 | 388 | 43.3 | 7.99 |
| Q99598 | 50.69 | 1 | 10 | 10 | 19 | 3.10 | 1 | 1 | 15.17 | 3 | 3 | 8.97 | 2 | 2 | | | | | | | | | | | | | | | | | | | 50.69 | 10 | 13 | 290 | 33.1 | 6.55 |
| Q99613 | 18.29 | 2 | 15 | 15 | 36 | 2.85 | 2 | 2 | 1.10 | 1 | 1 | 2.85 | 2 | 2 | 1.86 | 2 | 2 | | | | | | | | | | | | | | | | 15.77 | 13 | 29 | 913 | 105.3 | 5.68 |
| Q99615 | 4.45 | 1 | 2 | 2 | 3 | | | | | | | | | | | | | | | | | | | | | | | | | 2.02 | 1 | 1 | 2.43 | 1 | 2 | 494 | 56.4 | 6.96 |
| Q99622 | 26.19 | 1 | 2 | 2 | 2 | | | | | | | | | | | | | | | | | | | | | | | | | | | | 26.19 | 2 | 2 | 126 | 13.2 | 5.14 |
| Q99623 | 59.53 | 1 | 16 | 16 | 43 | 16.72 | 4 | 4 | 27.42 | 7 | 7 | 19.40 | 4 | 4 | 4.01 | 1 | 1 | 4.01 | 1 | 1 | | | | | | | | | | 4.01 | 1 | 1 | 59.53 | 16 | 25 | 299 | 33.3 | 9.83 |
| Q99627 | 42.58 | 1 | 5 | 5 | 14 | | | | | | | | | | 7.18 | 1 | 1 | | | | | | | 6.70 | 1 | 1 | 6.70 | 1 | 1 | 13.40 | 2 | 2 | 35.41 | 4 | 9 | 209 | 23.2 | 5.38 |
| Q99643 | 21.89 | 1 | 3 | 3 | 5 | | | | | | | | | | | | | | | | | | | | | | | | | | | | 21.89 | 3 | 5 | 169 | 18.6 | 9.69 |
| Q99714 | 89.27 | 1 | 16 | 16 | 54 | 50.57 | 8 | 8 | 57.09 | 9 | 12 | 28.35 | 4 | 5 | | | | | | | | | | | | | 3.83 | 1 | 1 | 3.83 | 1 | 1 | 83.91 | 14 | 27 | 261 | 26.9 | 7.78 |
| Q99720 | 9.42 | 1 | 2 | 2 | 2 | | | | | | | | | | | | | | | | | | | | | | | | | | | | 9.42 | 2 | 2 | 223 | 25.1 | 5.96 |
| Q9P273 | 0.78 | 1 | 1 | 1 | 5 | | | | | | | | | | | | | | | | | | | | | | | | | | | | 0.78 | 1 | 5 | 2699 | 300.8 | 6.42 |
| Q99733 | 23.20 | 1 | 7 | 7 | 15 | 5.87 | 2 | 2 | | | | | | | | | | 2.93 | 1 | 1 | | | | 8.53 | 3 | 3 | 2.93 | 1 | 2 | 5.87 | 2 | 2 | 11.73 | 3 | 5 | 375 | 42.8 | 4.69 |
| Q99747 | 6.09 | 1 | 2 | 2 | 3 | | | | | | | | | | | | | | | | | | | | | | | | | 2.88 | 1 | 1 | 3.21 | 1 | 2 | 312 | 34.7 | 5.41 |
| Q99766 | 43.72 | 1 | 8 | 8 | 10 | | | | | | | | | | | | | | | | | | | | | | | | | | | | 43.72 | 8 | 10 | 215 | 24.8 | 7.56 |
| Q99798 | 56.03 | 1 | 42 | 42 | 268 | 46.03 | 24 | 31 | 42.18 | 27 | 42 | 49.74 | 31 | 48 | 11.41 | 6 | 7 | 26.15 | 13 | 14 | | | | 3.59 | 2 | 2 | | | | 5.64 | 3 | 3 | 51.79 | 37 | 121 | 780 | 85.4 | 7.61 |
| O95674 | 4.27 | 1 | 1 | 1 | 5 | | | | | | | | | | | | | | | | | | | | | | | | | | | | 4.27 | 1 | 5 | 445 | 51.4 | 7.09 |
| Q99805 | 8.75 | 1 | 4 | 4 | 22 | 4.37 | 2 | 2 | 2.11 | 1 | 1 | | | | | | | | | | | | | | | | | | | | | | 8.75 | 4 | 19 | 663 | 75.7 | 7.44 |
| Q99816 | 7.69 | 1 | 3 | 3 | 3 | 2.56 | 1 | 1 | | | | | | | | | | | | | | | | | | | | | | | | | 5.13 | 2 | 2 | 390 | 43.9 | 6.46 |
| Q99828 | 19.37 | 1 | 3 | 3 | 3 | | | | | | | | | | | | | | | | | | | | | | | | | | | | 19.37 | 3 | 3 | 191 | 21.7 | 4.78 |
| Q99829 | 14.34 | 1 | 5 | 5 | 15 | | | | | | | | | | | | | | | | | | | | | | | | | | | | 14.34 | 5 | 15 | 537 | 59.0 | 5.83 |
| Q99832 | 81.58 | 1 | 45 | 45 | 257 | 41.99 | 19 | 21 | 37.94 | 18 | 28 | 37.02 | 15 | 21 | 26.52 | 12 | 13 | 35.91 | 14 | 16 | | | | 3.50 | 2 | 2 | 1.29 | 1 | 1 | 7.73 | 4 | 4 | 79.37 | 40 | 151 | 543 | 59.3 | 7.65 |
| Q99836 | 6.08 | 1 | 2 | 2 | 2 | | | | 2.36 | 1 | 1 | | | | | | | | | | | | | | | | | | | | | | 3.72 | 1 | 1 | 296 | 33.2 | 6.15 |
| P50747 | 1.65 | 1 | 1 | 1 | 5 | | | | | | | | | | | | | | | | | | | | | | | | | | | | 1.65 | 1 | 5 | 726 | 80.7 | 5.62 |
| Q99873 | 18.28 | 1 | 6 | 6 | 8 | | | | | | | | | | | | | | | | | | | | | | | | | | | | 18.28 | 6 | 8 | 361 | 41.5 | 5.43 |
| Q99932 | 53.76 | 1 | 17 | 17 | 61 | | | | | | | | | | 21.13 | 6 | 7 | 34.51 | 12 | 12 | 3.05 | 1 | 1 | | | | | | | | | | 47.42 | 14 | 41 | 426 | 44.8 | 5.50 |
| Q99963 | 31.99 | 1 | 8 | 9 | 11 | | | | | | | | | | | | | | | | | | | | | | 6.05 | 2 | 2 | | | | 28.53 | 8 | 9 | 347 | 39.3 | 5.38 |
| Q96EK4 | 2.87 | 1 | 1 | 1 | 5 | | | | | | | | | | | | | | | | | | | | | | | | | | | | 2.87 | 1 | 5 | 314 | 34.4 | 8.98 |
| Q99965 | 4.22 | 1 | 3 | 3 | 3 | | | | | | | | | | | | | | | | | | | | | | | | | | | | 4.22 | 3 | 3 | 735 | 82.4 | 6.15 |
| Q99985 | 26.76 | 3 | 16 | 18 | 41 | 4.79 | 2 | 2 | 0.80 | 1 | 1 | 6.92 | 3 | 3 | | | | | | | | | | | | | 2.80 | 3 | 3 | 5.59 | 4 | 5 | 23.30 | 15 | 27 | 751 | 85.2 | 8.69 |
| Q99988 | 18.18 | 1 | 3 | 3 | 14 | 10.39 | 2 | 2 | | | | 10.39 | 2 | 3 | | | | | | | | | | | | | | | | | | | 18.18 | 3 | 9 | 308 | 34.1 | 9.66 |
| Q99996 | 0.69 | 2 | 2 | 4 | 9 | | | | 0.00 | | | | | | 0.00 | | | 0.15 | 1 | 1 | | | | | | | 0.15 | 1 | 1 | 0.31 | 2 | 2 | 0.54 | 3 | 5 | 3911 | 453.4 | 4.98 |
| Q9BPX5 | 16.34 | 1 | 2 | 2 | 2 | | | | | | | | | | | | | | | | | | | | | | | | | | | | 16.34 | 2 | 2 | 153 | 16.9 | 6.60 |
| Q9BPY3 | 24.50 | 1 | 7 | 7 | 13 | | | | | | | | | | | | | | | | | | | | | | | | | | | | 24.50 | 7 | 13 | 351 | 39.5 | 5.99 |
| Q9BQ16 | 8.94 | 1 | 3 | 3 | 4 | | | | | | | 3.44 | 1 | 2 | | | | | | | | | | | | | | | | | | | 5.50 | 2 | 2 | 436 | 49.4 | 5.00 |
| Q99755 | 4.27 | 2 | 1 | 2 | 5 | | | | | | | | | | | | | | | | | | | | | | | | | | | | 4.27 | 2 | 5 | 562 | 62.6 | 8.21 |
| Q9BQA1 | 28.36 | 1 | 7 | 7 | 33 | 8.48 | 2 | 2 | 10.53 | 3 | 3 | 24.85 | 6 | 8 | | | | 4.39 | 1 | 1 | | | | | | | 4.39 | 1 | 1 | 4.39 | 1 | 1 | 28.36 | 7 | 17 | 342 | 36.7 | 5.17 |
| Q9BQA9 | 8.56 | 1 | 2 | 2 | 2 | | | | | | | | | | | | | | | | | | | | | | | | | | | | 8.56 | 2 | 2 | 187 | 20.8 | 6.80 |

| | | | | | | | | | | | | | | | | | | | | | | | | | | | | | | | | | | | | | | |
|---|---|---|---|---|---|---|---|---|---|---|---|---|---|---|---|---|---|---|---|---|---|---|---|---|---|---|---|---|---|---|---|---|---|---|---|---|---|---|
| Q9BQE3 | 80.62 | 2 | 7 | 41 | 683 | 39.20 | 16 | 24 | 42.76 | 19 | 34 | 42.54 | 17 | 22 | 52.34 | 25 | 49 | 50.11 | 22 | 40 | 9.13 | 4 | 5 | 28.51 | 12 | 17 | 30.29 | 12 | 16 | 45.66 | 15 | 24 | 78.62 | 35 | 452 | 449 | 49.9 | 5.10 |
| Q9BQM9 | 16.99 | 1 | 2 | 2 | 2 |||||||||||||||||||||||||||| 16.99 | 2 | 2 | 153 | 17.2 | 11.47 |
| Q9BQP9 | 31.10 | 2 | 5 | 5 | 23 | 12.60 | 2 | 3 | 6.69 | 2 | 2 | 16.14 | 3 | 3 |||||||||||||||||||||| 31.10 | 5 | 15 | 254 | 28.4 | 6.65 |
| Q9BR76 | 39.26 | 2 | 16 | 16 | 79 |||| 1.64 | 1 | 1 | 8.18 | 4 | 4 | 14.93 | 7 | 7 | 4.50 | 2 | 2 |||| 4.09 | 2 | 2 | 5.32 | 3 | 3 | 25.97 | 9 | 10 | 29.45 | 10 | 46 | 489 | 54.2 | 5.88 |
| Q9P003 | 14.39 | 1 | 1 | 1 | 5 |||||||||||||||||||||||||||| 14.39 | 1 | 5 | 139 | 16.1 | 6.65 |
| O60331 | 3.59 | 2 | 1 | 2 | 5 |||||||||||||||||||||||||||| 3.59 | 2 | 5 | 668 | 73.2 | 5.29 |
| Q9BRA2 | 45.53 | 1 | 6 | 6 | 24 | 19.51 | 2 | 2 | 8.13 | 1 | 1 | 8.13 | 1 | 1 | 30.89 | 3 | 4 | 13.01 | 1 | 1 |||| 26.83 | 3 | 3 | 31.71 | 3 | 3 | 39.84 | 4 | 4 | 43.90 | 5 | 5 | 123 | 13.9 | 5.52 |
| Q9BRF8 | 12.10 | 1 | 2 | 2 | 2 |||||||||||||||||||||||||||| 12.10 | 2 | 2 | 314 | 35.5 | 6.20 |
| Q15847 | 18.42 | 1 | 1 | 1 | 5 |||||||||||||||||||||||||||| 18.42 | 1 | 5 | 76 | 7.8 | 5.31 |
| Q9BRG1 | 46.59 | 1 | 6 | 6 | 8 |||||||||||||||||||||||||||| 46.59 | 6 | 8 | 176 | 20.7 | 6.34 |
| Q9BRK5 | 40.33 | 1 | 13 | 13 | 40 | 11.60 | 4 | 4 | 32.04 | 10 | 10 | 6.91 | 2 | 2 | 6.91 | 2 | 2 | 6.63 | 2 | 2 |||| 20.17 | 6 | 7 | 12.43 | 4 | 4 | 8.84 | 3 | 4 | 9.12 | 4 | 5 | 362 | 41.8 | 4.86 |
| Q9BRQ6 | 51.06 | 1 | 10 | 10 | 16 | 5.11 | 1 | 1 | 8.51 | 2 | 2 | 5.11 | 1 | 1 |||||||||||||||||||||| 47.66 | 9 | 12 | 235 | 26.4 | 8.85 |
| Q9BRR6 | 14.89 | 1 | 5 | 5 | 7 |||| 2.62 | 1 | 1 |||||||||||||||||||||||| 12.27 | 4 | 6 | 497 | 54.1 | 6.20 |
| Q9BRX2 | 15.84 | 1 | 7 | 7 | 13 ||||||| 3.64 | 1 | 1 | 3.64 | 1 | 1 ||||||||||||| 4.16 | 1 | 1 | 15.32 | 6 | 10 | 385 | 43.3 | 6.34 |
| Q9BRX8 | 54.15 | 2 | 14 | 14 | 27 | 11.35 | 2 | 2 | 11.35 | 2 | 2 ||||||||||||||||||||||||| 54.15 | 14 | 23 | 229 | 25.7 | 8.84 |
| Q9BRY0 | 9.55 | 1 | 2 | 2 | 2 |||||||||||||||||||||||||||| 9.55 | 2 | 2 | 314 | 33.6 | 6.87 |
| Q9BS26 | 47.04 | 1 | 18 | 19 | 79 | 33.00 | 10 | 11 | 28.33 | 11 | 13 | 27.09 | 10 | 11 | 6.90 | 3 | 4 | 11.82 | 5 | 6 ||||||| 1.72 | 1 | 1 | 5.42 | 2 | 2 | 26.35 | 11 | 27 | 406 | 46.9 | 5.26 |
| Q9BS86 | 72.08 | 1 | 23 | 23 | 183 | 46.44 | 13 | 18 | 37.32 | 12 | 15 | 35.90 | 10 | 12 | 21.08 | 7 | 8 | 22.79 | 7 | 8 |||| 2.85 | 1 | 1 | 3.42 | 1 | 1 | 2.85 | 1 | 1 | 71.79 | 21 | 119 | 351 | 40.1 | 9.28 |
| Q9BSD7 | 41.58 | 1 | 6 | 6 | 7 |||||||||||||||||||||||||||| 41.58 | 6 | 7 | 190 | 20.7 | 9.54 |
| Q9BSF0 | 88.42 | 1 | 9 | 9 | 20 | 33.68 | 2 | 3 | 26.32 | 2 | 2 ||||||||||||||||||||||||| 88.42 | 9 | 15 | 95 | 11.0 | 4.87 |
| Q9BSU3 | 31.44 | 1 | 3 | 6 | 10 |||||||||||||||||||||| 6.55 | 1 | 1 | 6.55 | 1 | 1 | 31.44 | 6 | 8 | 229 | 26.0 | 5.17 |
| Q9BT09 | 27.34 | 1 | 6 | 6 | 8 | 11.15 | 2 | 2 | 4.68 | 1 | 1 | 10.43 | 2 | 2 |||||||||||||||||||||| 14.75 | 3 | 3 | 278 | 30.7 | 5.49 |
| Q9BT22 | 8.41 | 1 | 3 | 3 | 4 |||||||||||||||||||||||||||| 8.41 | 3 | 4 | 464 | 52.5 | 7.23 |
| Q9BT30 | 33.94 | 1 | 5 | 5 | 5 |||||||||||||||||||||||||||| 33.94 | 5 | 5 | 221 | 24.5 | 7.11 |
| Q9BT73 | 31.97 | 1 | 2 | 2 | 3 |||||||||||||||||||||||||||| 31.97 | 2 | 3 | 122 | 13.1 | 7.88 |
| Q9BT78 | 78.08 | 1 | 26 | 27 | 66 ||||||| 2.46 | 1 | 1 | 14.04 | 5 | 5 ||||||| 11.82 | 5 | 7 | 2.46 | 1 | 1 | 27.83 | 10 | 11 | 70.94 | 23 | 41 | 406 | 46.2 | 5.83 |
| Q9BTE1 | 32.42 | 1 | 6 | 6 | 11 | 5.49 | 1 | 1 |||| 6.04 | 1 | 1 |||||||||||||||||||||| 32.42 | 6 | 9 | 182 | 20.1 | 8.02 |
| Q9BTE6 | 67.48 | 1 | 29 | 29 | 162 | 14.08 | 5 | 5 | 13.35 | 4 | 5 | 10.44 | 3 | 3 | 37.14 | 12 | 14 | 20.39 | 7 | 8 |||| 22.33 | 8 | 8 | 26.46 | 11 | 12 | 28.64 | 11 | 13 | 64.81 | 24 | 94 | 412 | 45.5 | 6.42 |
| Q9BTV4 | 11.50 | 1 | 3 | 3 | 3 |||||||||||||||||||||||||||| 11.50 | 3 | 3 | 400 | 44.8 | 8.13 |
| Q9BTW9 | 30.20 | 1 | 27 | 27 | 57 |||| 2.10 | 2 | 2 |||||||||||||||||||||||| 30.20 | 27 | 55 | 1192 | 132.5 | 6.19 |
| Q9BTX1 | 30.56 | 1 | 14 | 14 | 31 | 2.52 | 1 | 1 | 3.12 | 2 | 2 ||||||||||||||||||||||||| 29.08 | 13 | 28 | 674 | 76.3 | 9.09 |
| Q9BU02 | 63.04 | 1 | 9 | 9 | 16 | 13.48 | 1 | 1 ||||||||||||||||||| 4.35 | 1 | 1 | 13.48 | 3 | 3 | 63.04 | 9 | 11 | 230 | 25.6 | 4.82 |
| Q9BU61 | 19.02 | 1 | 3 | 3 | 5 |||||||||||||||||||||||||||| 19.02 | 3 | 5 | 184 | 20.3 | 8.22 |
| Q9BU89 | 15.23 | 1 | 2 | 2 | 3 |||||||||||||||||||||||||||| 15.23 | 2 | 3 | 302 | 32.9 | 4.83 |
| Q9BUB7 | 8.85 | 1 | 2 | 2 | 3 |||||||||||||||||||||||||||| 8.85 | 2 | 3 | 260 | 29.0 | 8.91 |
| Q9BUD6 | 20.85 | 1 | 8 | 8 | 39 | 9.97 | 4 | 4 | 4.83 | 2 | 2 | 7.55 | 3 | 3 | 9.97 | 4 | 4 | 5.14 | 2 | 2 ||||||| 5.14 | 2 | 2 | 11.48 | 4 | 4 | 19.34 | 7 | 18 | 331 | 35.8 | 5.52 |
| O43286 | 2.06 | 3 | 1 | 1 | 4 | 2.06 | 1 | 1 | 2.06 | 1 | 1 | 0.00 |||||||||||||||||||||| 2.06 | 1 | 2 | 388 | 45.1 | 8.15 |
| Q9BUF5 | 31.17 | 1 | 3 | 14 | 291 | 16.82 | 7 | 15 | 16.82 | 7 | 19 | 9.87 | 5 | 11 | 15.70 | 8 | 20 | 18.83 | 8 | 22 |||| 8.52 | 3 | 5 | 7.40 | 3 | 6 | 7.40 | 3 | 6 | 29.15 | 12 | 187 | 446 | 49.8 | 4.88 |
| Q9BUK6 | 7.72 | 1 | 3 | 3 | 4 |||||||||||||||||||||||||||| 7.72 | 3 | 4 | 570 | 61.8 | 6.11 |
| Q9BUL8 | 21.70 | 1 | 3 | 3 | 3 |||||||||| 5.66 | 1 | 1 ||||||||||||||||||| 16.04 | 2 | 2 | 212 | 24.7 | 8.19 |
| Q04759 | 2.12 | 5 | 1 | 2 | 4 | 0.99 | 1 | 1 ||||||| 0.99 | 1 | 1 ||||||| 0.99 | 1 | 1 ||||||| 1.13 | 1 | 1 | 706 | 81.8 | 7.61 |
| Q9BUN1 | 60.70 | 1 | 13 | 13 | 70 | 43.70 | 9 | 12 | 34.02 | 7 | 8 | 13.20 | 3 | 3 | 7.92 | 2 | 2 | 10.56 | 3 | 3 |||||||||||||| 59.24 | 12 | 42 | 341 | 36.7 | 8.59 |
| P25815 | 13.68 | 1 | 1 | 1 | 4 | 13.68 | 1 | 1 | 13.68 | 1 | 1 | 13.68 | 1 | 1 | 13.68 | 1 | 1 |||||||||||||||||||| 95 | 10.4 | 4.88 |
| Q9BUN8 | 10.76 | 1 | 3 | 3 | 4 |||||||||||||||||||||||||||| 10.76 | 3 | 4 | 251 | 28.8 | 9.51 |
| Q08648 | 8.74 | 1 | 1 | 1 | 4 | 8.74 | 1 | 1 | 8.74 | 1 | 1 ||||||| 8.74 | 1 | 1 |||||||||||||| 8.74 | 1 | 1 | 103 | 11.4 | 11.49 |
| Q9BUP3 | 37.60 | 1 | 9 | 9 | 16 | 4.55 | 1 | 1 | 8.26 | 2 | 2 |||||||||||| 4.55 | 1 | 1 ||||||| 37.60 | 9 | 12 | 242 | 27.0 | 8.38 |
| Q9BUR5 | 44.44 | 1 | 6 | 6 | 17 | 5.56 | 1 | 1 | 13.13 | 2 | 2 ||||||||||||||||||||||||| 44.44 | 6 | 14 | 198 | 22.3 | 9.13 |
| Q9BUT1 | 19.59 | 1 | 3 | 3 | 7 |||||||||||||||||||||||||||| 19.59 | 3 | 7 | 245 | 26.7 | 7.65 |
| Q9BV20 | 23.85 | 1 | 8 | 8 | 9 |||||||||||||||||||||||||||| 23.85 | 8 | 9 | 369 | 39.1 | 6.30 |
| Q9BV36 | 12.17 | 1 | 6 | 6 | 7 |||||||||||||||||| 2.50 | 1 | 1 |||| 4.33 | 2 | 2 | 7.33 | 4 | 4 | 600 | 65.9 | 6.07 |
| Q9BV40 | 53.00 | 1 | 6 | 6 | 10 |||||||||||||||||||||||||||| 53.00 | 6 | 10 | 100 | 11.4 | 7.34 |
| Q9BV44 | 12.23 | 1 | 5 | 5 | 7 |||||||||||||||||||||||||||| 12.23 | 5 | 7 | 507 | 57.0 | 6.37 |
| Q9BV57 | 24.02 | 1 | 4 | 4 | 11 |||||||||||||||||| 7.82 | 1 | 1 |||| 12.85 | 2 | 3 | 18.99 | 3 | 7 | 179 | 21.5 | 5.68 |
| Q9BV68 | 6.75 | 1 | 2 | 2 | 2 |||||||||||||||||||||||||||| 6.75 | 2 | 2 | 326 | 35.6 | 5.72 |
| Q9BV86 | 31.84 | 1 | 9 | 9 | 19 |||||||||| 10.31 | 2 | 2 ||||||| 4.04 | 1 | 1 |||| 14.35 | 3 | 3 | 31.84 | 7 | 13 | 223 | 25.4 | 5.52 |
| Q9BVA1 | 87.19 | 2 | 2 | 39 | 990 | 45.39 | 18 | 44 | 52.58 | 20 | 59 | 46.07 | 19 | 41 | 53.71 | 25 | 70 | 53.48 | 27 | 93 | 6.97 | 2 | 3 | 27.42 | 11 | 20 | 29.21 | 13 | 27 | 29.89 | 14 | 28 | 80.45 | 32 | 605 | 445 | 49.9 | 4.89 |
| Q9BVC4 | 9.20 | 1 | 2 | 2 | 3 |||||||||||||||||||||||||||| 9.20 | 2 | 3 | 326 | 35.9 | 5.86 |
| Q9BVC6 | 9.05 | 1 | 3 | 3 | 3 |||||||||||||||||||||||||||| 9.05 | 3 | 3 | 243 | 26.2 | 10.48 |
| Q9BVG4 | 10.73 | 1 | 2 | 2 | 4 |||||||||||||||||| 4.29 | 1 | 1 |||| 4.29 | 1 | 1 | 6.44 | 1 | 2 | 233 | 26.0 | 4.79 |
| Q9BVG8 | 3.96 | 2 | 3 | 3 | 5 |||||||||||||||||||||||||||| 3.96 | 3 | 5 | 833 | 92.7 | 7.69 |
| Q9BVG9 | 17.86 | 1 | 8 | 8 | 20 | 4.52 | 1 | 1 | 11.70 | 4 | 4 ||||||||||||||||||||||||| 15.40 | 6 | 15 | 487 | 56.2 | 6.25 |
| Q9BVJ7 | 27.33 | 1 | 4 | 4 | 4 |||||||||||||||||||||||||||| 27.33 | 4 | 4 | 150 | 16.6 | 8.21 |
| P32321 | 3.93 | 1 | 1 | 1 | 4 |||| 0.00 ||| 3.93 | 1 | 1 |||||||||||||||||||||| 3.93 | 1 | 3 | 178 | 20.0 | 7.56 |
| Q9BVK6 | 32.77 | 1 | 8 | 9 | 39 | 20.00 | 4 | 4 | 12.77 | 3 | 3 | 20.00 | 4 | 4 | 8.51 | 2 | 2 ||||||||||||||||||| 32.77 | 9 | 26 | 235 | 27.3 | 8.02 |
| Q53G44 | 1.33 | 1 | 1 | 1 | 4 |||| 1.33 | 1 | 1 | 1.33 | 1 | 1 |||||||||||||||||||||| 1.33 | 1 | 2 | 452 | 51.3 | 6.79 |
| Q9BVL2 | 20.03 | 1 | 10 | 10 | 23 | 2.17 | 1 | 1 | 8.35 | 3 | 4 | 3.51 | 2 | 2 |||||||||||||||||||||| 17.86 | 9 | 16 | 599 | 60.9 | 9.33 |
| Q9BVM2 | 74.38 | 1 | 17 | 17 | 66 | 26.60 | 6 | 6 | 34.98 | 8 | 9 | 25.12 | 6 | 6 | 10.34 | 2 | 2 | 8.37 | 2 | 2 |||| 6.90 | 2 | 2 |||| 9.36 | 2 | 2 | 74.38 | 17 | 37 | 203 | 23.2 | 9.03 |
| Q9UPI3 | 2.28 | 1 | 1 | 1 | 4 |||| 2.28 | 1 | 1 | 2.28 | 1 | 1 |||||||||||||||||||||| 2.28 | 1 | 2 | 526 | 57.2 | 6.18 |
| Q9BVM4 | 36.60 | 1 | 4 | 4 | 5 |||||||||||||||||| 7.84 | 1 | 1 ||||||| 36.60 | 4 | 4 | 153 | 17.3 | 6.87 |
| Q9BW27 | 16.16 | 1 | 8 | 8 | 11 |||||||||||||||||||||||||||| 16.16 | 8 | 11 | 656 | 75.0 | 5.55 |
| P17844 | 3.75 | 2 | 1 | 2 | 4 ||||||| 1.79 | 1 | 1 | 1.79 | 1 | 1 |||||||||||||||||||| 1.95 | 1 | 2 | 614 | 69.1 | 8.92 |
| Q9BW83 | 56.99 | 1 | 7 | 7 | 14 |||||||||| 4.30 | 1 | 1 |||||||||| 4.30 | 1 | 1 | 4.30 | 1 | 1 | 52.69 | 6 | 11 | 186 | 20.5 | 5.41 |
| Q9BW91 | 8.57 | 1 | 3 | 3 | 4 ||||||||||||||||||||||||| 8.57 | 3 | 4 | 0.00 ||| 350 | 39.1 | 8.22 |
| Q5VT25 | 0.98 | 1 | 1 | 2 | 4 ||||||| 0.00 |||||||||||||||||||| 0.40 | 1 | 1 | 0.98 | 2 | 3 | 1732 | 197.2 | 6.58 |
| Q9BWD1 | 63.48 | 2 | 16 | 16 | 34 |||||||||| 10.08 | 2 | 2 |||||||||||||| 7.56 | 3 | 3 | 61.96 | 15 | 29 | 397 | 41.3 | 6.92 |
| P05166 | 2.97 | 1 | 1 | 1 | 4 ||||||| 2.97 | 1 | 1 |||||||||||||||||||||| 2.97 | 1 | 3 | 539 | 58.2 | 7.64 |
| Q9BWH2 | 67.20 | 2 | 16 | 16 | 45 | 15.87 | 2 | 2 | 16.93 | 3 | 3 | 11.11 | 2 | 2 |||| 5.82 | 1 | 1 ||||||| 5.82 | 1 | 1 |||| 67.20 | 15 | 36 | 189 | 20.7 | 9.73 |
| P0CG29 | 8.20 | 2 | 1 | 2 | 4 ||||||| 5.33 | 1 | 1 |||||||||||||||||||||| 2.87 | 1 | 3 | 244 | 27.5 | 7.02 |
| Q9BWM7 | 9.35 | 1 | 2 | 2 | 2 |||||||||||||||||||||||||||| 9.35 | 2 | 2 | 321 | 35.5 | 9.10 |
| Q9BWS9 | 45.80 | 1 | 15 | 15 | 38 | 16.03 | 4 | 5 | 18.58 | 5 | 7 | 19.34 | 6 | 7 |||||||||||||||||||||| 31.04 | 11 | 19 | 393 | 44.9 | 8.63 |
| Q9BX10 | 3.16 | 1 | 2 | 2 | 2 |||||||||||||||||||||||||||| 3.16 | 2 | 2 | 602 | 65.7 | 8.05 |
| Q9BX68 | 47.24 | 1 | 5 | 5 | 18 | 26.99 | 3 | 4 | 21.47 | 2 | 4 | 17.79 | 2 | 2 |||| 9.82 | 1 | 1 |||||||||||||| 47.24 | 5 | 7 | 163 | 17.2 | 9.16 |
| Q9BXA6 | 33.33 | 1 | 10 | 10 | 18 |||||||||| 5.86 | 2 | 2 | 2.93 | 1 | 1 ||||||||||||||| 33.33 | 10 | 15 | 273 | 30.3 | 9.11 |
| Q9BXA7 | 36.78 | 1 | 10 | 10 | 19 |||||||||||||||||||||||||||| 36.78 | 10 | 19 | 367 | 41.6 | 7.71 |
| Q9BXC9 | 2.64 | 1 | 2 | 2 | 2 |||||||||||||||||||||||||||| 2.64 | 2 | 2 | 721 | 79.8 | 6.15 |
| Q9BXF9 | 90.61 | 2 | 58 | 58 | 488 | 41.22 | 16 | 18 | 36.73 | 16 | 20 | 34.08 | 13 | 13 | 42.65 | 20 | 22 | 55.92 | 31 | 41 | 2.86 | 1 | 1 | 44.69 | 21 | 26 | 47.76 | 26 | 32 | 61.63 | 39 | 53 | 80.61 | 40 | 260 | 490 | 56.6 | 7.34 |
| P80192 | 0.54 | 1 | 1 | 1 | 4 |||||||||| 0.54 | 1 | 1 | 0.00 ||||||||| 0.54 | 1 | 1 | 0.00 ||| 0.54 | 1 | 2 | 1104 | 121.8 | 5.87 |
| Q9BXI2 | 18.60 | 1 | 4 | 4 | 5 |||||||||||||||||||||||||||| 18.60 | 4 | 5 | 301 | 32.6 | 9.09 |
| Q9BXJ8 | 15.16 | 1 | 6 | 6 | 14 | 2.92 | 1 | 1 | 2.92 | 1 | 1 | 3.21 | 1 | 1 |||||||||||||||||||||| 14.87 | 5 | 11 | 343 | 40.6 | 8.98 |
| Q9BXJ9 | 22.75 | 1 | 16 | 21 | 28 |||||||||||||||||||||||||||| 22.75 | 21 | 28 | 866 | 101.2 | 7.42 |
| P42285 | 0.67 | 1 | 1 | 1 | 4 ||||||||||||| 0.67 | 1 | 1 ||||||||||||||| 0.67 | 1 | 3 | 1042 | 117.7 | 6.52 |
| Q96PH6 | 8.94 | 1 | 1 | 1 | 4 ||||||||||||| 8.94 | 1 | 1 ||||||||||||||| 8.94 | 1 | 3 | 123 | 13.6 | 7.31 |
| Q8NAB2 | 1.15 | 1 | 1 | 1 | 4 |||||||||||||||||| 1.15 | 1 | 2 ||||||| 1.15 | 1 | 2 | 608 | 69.3 | 5.39 |
| Q9BXM0 | 1.71 | 1 | 3 | 3 | 3 |||||||||||||||||||||||||||| 1.71 | 3 | 3 | 1461 | 154.8 | 7.50 |
| Q9BXM9 | 17.55 | 1 | 8 | 8 | 13 |||||||||| 1.70 | 1 | 1 ||||||| 2.83 | 1 | 1 | 1.51 | 1 | 1 | 5.09 | 2 | 2 | 13.77 | 6 | 8 | 530 | 59.5 | 6.20 |
| Q9Y608 | 2.64 | 1 | 1 | 2 | 4 |||||||||||||||||| 1.66 | 1 | 1 | 1.66 | 1 | 1 |||| 2.64 | 2 | 3 | 721 | 82.1 | 6.95 |
| Q9BXN6 | 93.81 | 1 | 2 | 13 | 157 | 78.35 | 6 | 11 | 78.35 | 5 | 13 | 78.35 | 4 | 6 | 68.04 | 6 | 10 | 90.72 | 7 | 18 | 23.71 | 3 | 5 | 37.11 | 4 | 8 | 68.04 | 8 | 19 | 68.04 | 6 | 7 | 88.66 | 7 | 60 | 97 | 11.0 | 6.11 |
| Q9BXS5 | 22.46 | 1 | 6 | 8 | 12 |||||||||||||||||||||||||||| 22.46 | 8 | 12 | 423 | 48.6 | 7.30 |
| Q9BXT8 | 3.94 | 1 | 6 | 6 | 7 ||||||||||||| 0.68 | 1 | 1 ||||||||||||||| 3.94 | 6 | 6 | 1623 | 184.5 | 5.40 |
| Q9BXU1 | 7.36 | 1 | 6 | 6 | 14 |||||||||||||||||| 1.57 | 1 | 1 ||||||| 7.36 | 6 | 13 | 1019 | 115.6 | 5.14 |
| O60234 | 26.76 | 1 | 1 | 3 | 4 |||||||||||||||||||||||| 7.75 | 2 | 2 | 19.01 | 1 | 2 | 142 | 16.8 | 5.26 |
| P60983 | 36.62 | 1 | 1 | 3 | 4 |||||||||||||||||||||||| 6.34 | 1 | 1 | 30.28 | 2 | 3 | 142 | 16.7 | 5.29 |
| Q9BXW7 | 27.19 | 1 | 8 | 8 | 8 |||| 3.07 | 1 | 1 |||||||||||||||||||||||| 24.11 | 7 | 7 | 423 | 46.3 | 8.13 |
| Q9BY14 | 44.58 | 1 | 7 | 7 | 60 | 29.32 | 6 | 10 | 25.70 | 5 | 8 | 44.58 | 7 | 12 | 20.08 | 4 | 4 | 23.69 | 5 | 6 |||| 10.44 | 2 | 2 | 15.26 | 3 | 3 | 4.42 | 1 | 1 | 18.47 | 4 | 14 | 249 | 26.6 | 4.94 |
| Q9BY32 | 39.18 | 1 | 6 | 6 | 13 |||||||||||||||||||||||| 4.12 | 1 | 1 | 39.18 | 6 | 12 | 194 | 21.4 | 5.66 |
| P57737 | 1.41 | 1 | 1 | 1 | 4 |||||||||||||||||||||||| 1.41 | 1 | 1 | 1.41 | 1 | 3 | 925 | 100.5 | 5.80 |
| Q9BY43 | 10.36 | 1 | 2 | 2 | 4 |||||||||| 6.31 | 1 | 1 ||||||| 10.36 | 2 | 2 ||||||| 6.31 | 1 | 3 | 222 | 25.1 | 4.70 |
| Q9BY50 | 27.08 | 1 | 6 | 6 | 8 |||||||||||||||||||||||||||| 27.08 | 6 | 8 | 192 | 21.5 | 9.20 |
| Q9BY67 | 12.67 | 1 | 3 | 3 | 9 |||| 1.36 | 1 | 1 | 6.79 | 2 | 2 |||||||||||||||||||||| 12.67 | 3 | 6 | 442 | 48.5 | 5.07 |
| Q9BYC2 | 35.01 | 1 | 13 | 14 | 35 | 13.35 | 5 | 6 | 8.90 | 3 | 4 | 7.54 | 2 | 2 |||||||||||||||||||||| 35.01 | 14 | 23 | 517 | 56.1 | 7.14 |
| Q9BYC5 | 7.30 | 1 | 4 | 4 | 7 |||||||||| 0.00 ||| 0.00 ||||||||||||||| 7.30 | 4 | 7 | 575 | 66.5 | 7.66 |
| Q9BYD9 | 23.39 | 1 | 6 | 6 | 10 |||| 0.00 |||||||||||||||||||||||||| 23.39 | 6 | 10 | 372 | 41.0 | 5.82 |
| Q9BYT3 | 6.61 | 1 | 3 | 3 | 7 |||| 2.72 | 1 | 1 ||||||| 2.72 | 1 | 1 ||||||| 2.72 | 1 | 1 | 2.72 | 1 | 1 | 6.61 | 3 | 5 | 514 | 57.8 | 7.03 |
| Q9BYW3 | 24.32 | 1 | 5 | 5 | 7 |||||||||||||||||||||||||||| 24.32 | 5 | 7 | 111 | 12.2 | 9.31 |
| Q9BYZ2 | 46.98 | 1 | 20 | 22 | 119 | 28.08 | 10 | 18 | 31.76 | 11 | 17 | 28.87 | 10 | 11 | 17.32 | 5 | 6 | 5.77 | 2 | 3 |||||||||||||| 46.98 | 18 | 64 | 381 | 41.9 | 8.65 |
| Q9BZ23 | 3.51 | 1 | 2 | 2 | 2 |||||||||||||||||||||||||||| 3.51 | 2 | 2 | 570 | 62.6 | 9.28 |
| Q9BZH6 | 7.19 | 1 | 7 | 7 | 10 |||||||||||||||||||||||||||| 7.19 | 7 | 10 | 1224 | 136.6 | 6.92 |
| Q9BZQ2 | 3.98 | 1 | 2 | 2 | 3 ||||||||||||| 2.45 | 1 | 1 ||||||||||||||| 3.98 | 2 | 2 | 653 | 72.6 | 5.31 |
| Q9BZQ6 | 1.82 | 1 | 2 | 2 | 2 | 0.97 | 1 | 1 ||||||||||||||||||||||||| 0.86 | 1 | 1 | 932 | 104.6 | 4.93 |
| Q9BZQ8 | 24.14 | 1 | 19 | 19 | 87 | 2.80 | 2 | 3 | 1.29 | 1 | 1 | 2.59 | 1 | 1 | 2.26 | 2 | 2 ||||||| 1.40 | 1 | 1 |||| 5.71 | 5 | 5 | 22.95 | 17 | 74 | 928 | 103.1 | 4.78 |
| Q9BZV1 | 49.43 | 1 | 18 | 18 | 61 |||||||||| 2.27 | 1 | 1 ||||||| 4.08 | 2 | 2 | 1.81 | 1 | 1 | 11.56 | 4 | 6 | 44.90 | 15 | 51 | 441 | 49.7 | 6.89 |
| Q9BZW7 | 70.34 | 1 | 61 | 61 | 152 |||||||||| 2.87 | 2 | 2 | 1.43 | 1 | 1 |||| 20.06 | 12 | 14 | 10.03 | 6 | 6 | 47.85 | 31 | 35 | 59.03 | 43 | 94 | 698 | 81.4 | 5.97 |
| Q9BZX4 | 94.34 | 1 | 6 | 17 | 380 | 57.08 | 8 | 16 | 57.08 | 8 | 16 | 49.06 | 8 | 15 | 37.74 | 6 | 17 | 33.96 | 7 | 14 |||| 17.45 | 4 | 8 | 7.08 | 1 | 3 | 13.21 | 2 | 10 | 89.15 | 14 | 281 | 212 | 23.9 | 5.15 |
| Q8NCA5 | 2.89 | 1 | 1 | 1 | 4 |||||||||||||||||||||||||||| 2.89 | 1 | 4 | 519 | 55.4 | 9.03 |
| Q9C002 | 61.45 | 1 | 7 | 7 | 20 | 51.81 | 4 | 5 | 59.04 | 4 | 6 | 18.07 | 1 | 1 |||||||||||||||||||||| 61.45 | 5 | 8 | 83 | 9.6 | 9.47 |
| Q9C005 | 25.25 | 1 | 2 | 2 | 4 | 25.25 | 2 | 2 | 25.25 | 2 | 2 |||||||||||||||||||||||||||| 99 | 11.2 | 4.88 |
| Q9C093 | 20.47 | 2 | 36 | 36 | 77 | 1.04 | 2 | 2 | 1.70 | 3 | 3 ||||||| 1.59 | 2 | 2 |||||||||||||| 19.37 | 34 | 70 | 1822 | 209.7 | 5.54 |
| Q9C099 | 13.76 | 1 | 12 | 12 | 17 ||||||| 0.00 |||||||||||||||||||| 3.39 | 3 | 3 | 12.11 | 11 | 14 | 1032 | 119.5 | 5.88 |
| Q9C0B2 | 7.45 | 1 | 10 | 10 | 21 |||||||||||||||||||||||||||| 7.45 | 10 | 21 | 1584 | 178.5 | 6.42 |
| Q9C0C9 | 3.10 | 1 | 2 | 3 | 3 ||||||||||||||||||||| 0.00 |||||| 3.10 | 3 | 3 | 1292 | 141.2 | 5.12 |
| Q5TCQ9 | 0.53 | 1 | 1 | 1 | 4 |||||||||||||||||||||||||||| 0.53 | 1 | 4 | 1506 | 165.5 | 8.02 |
| Q9C0E2 | 10.25 | 1 | 8 | 9 | 10 |||||||||||||||||||||||||||| 10.25 | 9 | 10 | 1151 | 130.1 | 5.05 |
| Q9C0G6 | 40.02 | 2 | 149 | 152 | 362 | 0.53 | 2 | 2 | 0.94 | 3 | 3 | 0.51 | 2 | 2 | 6.93 | 26 | 28 | 3.05 | 10 | 10 |||||||||||||| 38.62 | 145 | 319 | 4158 | 475.7 | 6.00 |
| CON_007020243 | 19.67 | 6 | 1 | 1 | 4 |||||||||||||||||||||||||||| 19.67 | 1 | 4 | 61 | 5.9 | 7.96 |
| Q9GZN6 | 5.16 | 1 | 3 | 3 | 4 |||||||||||||||||||||||||||| 5.16 | 3 | 4 | 736 | 82.1 | 8.27 |
| Q9GZP4 | 34.60 | 1 | 6 | 6 | 10 |||||||||||||||||||||||||||| 34.60 | 6 | 10 | 211 | 24.2 | 5.74 |
| Q9GZS0 | 67.60 | 2 | 37 | 37 | 104 | 10.91 | 5 | 5 | 4.46 | 2 | 2 | 12.40 | 5 | 5 | 8.26 | 4 | 4 | 18.68 | 9 | 10 |||||||||||||| 61.32 | 31 | 78 | 605 | 68.8 | 4.74 |
| Q9GZS3 | 57.38 | 1 | 13 | 13 | 26 ||||||| 9.51 | 1 | 1 | 2.30 | 1 | 1 | 2.30 | 1 | 1 ||||||| 2.30 | 1 | 1 | 19.67 | 5 | 6 | 55.08 | 12 | 16 | 305 | 33.6 | 5.47 |
| Q9GZT6 | 22.83 | 1 | 6 | 6 | 6 |||||||||||||||||||||||||||| 22.83 | 6 | 6 | 254 | 29.5 | 7.55 |
| Q9GZT8 | 26.26 | 1 | 7 | 7 | 14 | 4.51 | 1 | 1 | 15.12 | 4 | 6 | 2.92 | 1 | 1 |||||||||||||||||||||| 14.06 | 4 | 6 | 377 | 41.9 | 6.65 |
| Q9GZY8 | 3.22 | 1 | 2 | 2 | 2 |||| 2.63 | 1 | 1 |||||||||||||||||||||||| 3.22 | 1 | 2 | 342 | 38.4 | 8.95 |
| Q9GZZ1 | 23.08 | 1 | 5 | 5 | 6 |||||||||||||||||||||||||||| 23.08 | 5 | 6 | 169 | 19.4 | 8.81 |
| P33527 | 0.52 | 1 | 1 | 1 | 4 |||||||||||||||||||||||||||| 0.52 | 1 | 4 | 1531 | 171.5 | 7.11 |
| Q9GZZ9 | 28.96 | 1 | 7 | 7 | 17 |||||||||||||||||||||||| 6.19 | 2 | 3 | 25.99 | 6 | 14 | 404 | 44.8 | 4.84 |
| Q9H008 | 18.52 | 1 | 2 | 2 | 4 | 10.74 | 1 | 1 | 7.78 | 1 | 1 ||||||||||||||||||||||||| 7.78 | 1 | 2 | 270 | 29.1 | 6.15 |
| Q16539 | 5.00 | 7 | 1 | 2 | 4 |||||||||||||||||||||||||||| 5.00 | 2 | 4 | 360 | 41.3 | 5.78 |
| Q9H061 | 50.26 | 1 | 7 | 7 | 14 |||| 9.23 | 2 | 2 |||||||||||||||||||||||| 50.26 | 7 | 12 | 195 | 21.5 | 9.26 |
| Q9H069 | 42.26 | 1 | 19 | 19 | 43 |||||||||| 18.55 | 7 | 9 | 14.15 | 6 | 7 ||||||||||||||| 37.09 | 16 | 27 | 523 | 61.0 | 4.78 |
| Q92504 | 2.99 | 1 | 1 | 1 | 4 |||||||||||||||||||||||||||| 2.99 | 1 | 4 | 469 | 50.1 | 6.87 |
| Q9H095 | 53.72 | 2 | 19 | 20 | 40 |||||||||| 5.19 | 2 | 2 | 14.90 | 6 | 7 ||||||||||||||| 48.31 | 15 | 31 | 443 | 51.9 | 6.52 |
| Q9H0A8 | 26.63 | 1 | 4 | 4 | 5 |||||||||||||||||||||||||||| 26.63 | 4 | 5 | 199 | 21.8 | 7.31 |
| Q9H0B3 | 21.53 | 1 | 20 | 20 | 56 | 3.22 | 3 | 3 | 2.03 | 2 | 2 |||| 0.93 | 1 | 1 | 1.10 | 1 | 1 |||||||||||||| 19.49 | 18 | 49 | 1180 | 127.6 | 10.23 |
| Q9H0B6 | 10.29 | 2 | 3 | 6 | 6 | 0.00 |||||||||||||||||||||| 0.00 ||| 5.79 | 4 | 4 | 4.50 | 2 | 2 | 622 | 68.9 | 7.15 |
| Q9H0B8 | 10.87 | 1 | 3 | 4 | 5 |||| 2.01 | 1 | 1 | 2.01 | 1 | 1 |||||||||||||||||||||| 8.85 | 3 | 3 | 497 | 55.9 | 8.02 |
| Q9H0C1 | 60.27 | 1 | 18 | 18 | 41 |||||||||||||||||||||||||||| 60.27 | 18 | 41 | 365 | 41.8 | 6.23 |
| Q9H0C2 | 68.25 | 1 | 22 | 25 | 126 | 32.06 | 10 | 11 | 53.97 | 19 | 21 | 34.29 | 11 | 13 | 13.65 | 4 | 4 | 13.02 | 4 | 4 |||||||||||||| 67.62 | 24 | 73 | 315 | 35.0 | 9.89 |
| Q9H0E2 | 32.12 | 1 | 7 | 7 | 15 | 5.11 | 1 | 1 | 5.11 | 1 | 1 | 5.11 | 1 | 1 | 4.74 | 1 | 1 ||||||||||| 2.92 | 1 | 1 | 2.92 | 1 | 1 | 29.20 | 6 | 9 | 274 | 30.3 | 5.97 |
| Q9H0F7 | 32.26 | 1 | 4 | 4 | 6 |||||||||||||||||||||||||||| 32.26 | 4 | 6 | 186 | 21.1 | 8.54 |
| Q9H0I3 | 33.95 | 1 | 15 | 15 | 28 |||| 2.65 | 1 | 1 |||| 13.53 | 5 | 5 | 11.94 | 5 | 5 |||||||||| 3.71 | 1 | 1 | 29.44 | 11 | 16 | 377 | 44.2 | 8.73 |
| Q9H0I9 | 29.55 | 1 | 11 | 13 | 38 |||| 4.79 | 2 | 2 | 2.24 | 2 | 2 | 4.31 | 2 | 2 | 2.40 | 1 | 1 |||| 2.40 | 1 | 1 | 4.79 | 2 | 2 | 4.79 | 2 | 2 | 26.36 | 11 | 26 | 626 | 67.8 | 6.33 |
| Q9H0J4 | 21.41 | 1 | 31 | 31 | 87 |||| 4.63 | 4 | 4 | 3.13 | 2 | 2 | 8.96 | 10 | 12 | 12.51 | 16 | 16 |||| 0.72 | 1 | 1 | 0.72 | 1 | 1 |||| 13.95 | 18 | 51 | 1663 | 180.7 | 6.73 |
| Q9H0K4 | 61.37 | 1 | 25 | 25 | 181 | 7.25 | 4 | 5 | 16.32 | 7 | 8 |||| 20.78 | 12 | 21 | 20.64 | 13 | 19 |||| 5.30 | 3 | 3 | 7.39 | 4 | 5 | 5.86 | 4 | 5 | 60.25 | 23 | 115 | 717 | 80.9 | 4.39 |
| Q9H0Q0 | 10.22 | 1 | 2 | 2 | 2 |||||||||||||||||||||||||||| 10.22 | 2 | 2 | 323 | 37.3 | 6.01 |
| Q9H0R4 | 18.53 | 1 | 4 | 4 | 9 | 3.09 | 1 | 1 |||||||||||||||||||||||||| 18.53 | 4 | 8 | 259 | 28.5 | 6.24 |
| Q9H0U3 | 11.94 | 1 | 5 | 5 | 11 |||||||||||||||||||||||||||| 11.94 | 5 | 11 | 335 | 38.0 | 9.63 |
| Q9H0U4 | 86.57 | 8 | 6 | 17 | 107 | 31.34 | 5 | 6 | 34.83 | 6 | 6 | 29.85 | 5 | 5 | 21.39 | 3 | 3 | 2.99 | 1 | 1 |||| 19.40 | 3 | 3 | 12.94 | 4 | 4 | 32.34 | 5 | 6 | 85.07 | 14 | 73 | 201 | 22.2 | 5.73 |
| Q9H0W9 | 28.25 | 1 | 8 | 8 | 25 | 6.35 | 2 | 2 | 7.94 | 2 | 2 ||||||||||||||||||||||||| 23.17 | 6 | 21 | 315 | 35.1 | 6.70 |
| Q9H0X4 | 3.99 | 1 | 2 | 2 | 2 |||||||||||||||||||||||||||| 3.99 | 2 | 2 | 552 | 59.6 | 6.28 |
| P36406 | 4.70 | 2 | 1 | 3 | 4 |||||||||||||||||||||||||||| 4.70 | 3 | 4 | 574 | 64.0 | 6.38 |
| Q9H173 | 21.91 | 1 | 9 | 9 | 21 | 2.39 | 1 | 1 | 3.04 | 1 | 1 | 2.60 | 1 | 1 |||||||||||||||||||||| 19.52 | 8 | 18 | 461 | 52.1 | 5.36 |
| Q9H1E5 | 33.52 | 1 | 9 | 9 | 40 | 15.47 | 4 | 5 | 16.62 | 4 | 6 | 15.47 | 4 | 6 | 3.72 | 1 | 1 ||||||||||||||||||| 28.65 | 8 | 22 | 349 | 38.9 | 4.37 |
| Q9H1H1 | 29.73 | 1 | 4 | 4 | 7 |||||||||||||||||||||||||||| 29.73 | 4 | 7 | 148 | 16.9 | 6.76 |
| Q9H1K1 | 44.91 | 1 | 7 | 7 | 10 | 11.38 | 1 | 1 | 5.39 | 1 | 1 ||||||||||||||||||||||||| 44.91 | 7 | 8 | 167 | 18.0 | 9.48 |
| Q9H1K4 | 7.62 | 2 | 3 | 3 | 3 |||| 2.54 | 1 | 1 |||||||||||||||||||||||| 5.08 | 2 | 2 | 315 | 33.8 | 9.25 |
| Q9H1M3 | 29.51 | 1 | 5 | 5 | 27 |||| 16.39 | 2 | 2 |||||||||||||||||||||||| 29.51 | 5 | 29 | 183 | 20.3 | 8.81 |
| Q9H1P6 | 47.45 | 1 | 6 | 6 | 9 |||||||||| 21.90 | 2 | 2 | 16.79 | 2 | 2 ||||||||||||||| 30.66 | 4 | 5 | 137 | 15.7 | 8.46 |
| Q9H1X1 | 86.96 | 2 | 26 | 26 | 117 | 13.77 | 4 | 4 | 38.04 | 9 | 9 | 7.61 | 2 | 2 | 40.58 | 16 | 20 | 39.49 | 11 | 15 |||| 7.61 | 2 | 2 | 14.49 | 3 | 5 | 19.57 | 4 | 4 | 83.33 | 23 | 56 | 276 | 31.3 | 5.43 |
| Q9H1Y0 | 6.18 | 1 | 2 | 2 | 2 |||||||||||||||||||||||||||| 6.18 | 2 | 2 | 275 | 32.4 | 5.77 |
| Q9H223 | 10.17 | 1 | 4 | 6 | 8 |||||||||||||||||||||||||||| 10.17 | 6 | 8 | 541 | 61.1 | 6.76 |
| Q9H269 | 7.15 | 1 | 4 | 4 | 5 |||||||||||||||||||||||||||| 7.15 | 4 | 5 | 839 | 94.6 | 6.77 |
| Q9H270 | 7.76 | 1 | 5 | 5 | 6 |||||||||||||||||||||||||||| 7.76 | 5 | 6 | 941 | 107.8 | 7.05 |
| Q9H299 | 36.56 | 1 | 3 | 3 | 9 ||||||||||||| 32.26 | 2 | 2 |||| 32.26 | 2 | 3 | 10.75 | 1 | 1 | 36.56 | 3 | 3 |||| 93 | 10.4 | 4.93 |
| Q9H2D1 | 16.83 | 1 | 2 | 2 | 3 |||||||||||||||||||||||||||| 16.83 | 2 | 3 | 315 | 35.4 | 9.45 |
| Q86TP1 | 4.64 | 1 | 1 | 1 | 4 |||||||||||||||||||||||||||| 4.64 | 1 | 4 | 453 | 50.2 | 5.50 |
| Q9H2G2 | 6.40 | 2 | 7 | 7 | 9 ||||||| 1.21 | 1 | 1 |||| 0.00 ||| 0.57 | 1 | 1 ||||||| 1.05 | 1 | 1 | 3.56 | 4 | 6 | 1235 | 142.6 | 5.15 |
| Q9H2H8 | 39.13 | 1 | 4 | 4 | 6 |||||||||||||||||| 7.45 | 1 | 1 | 7.45 | 1 | 1 |||| 31.68 | 3 | 4 | 161 | 18.1 | 6.79 |
| Q9H2K8 | 14.48 | 3 | 10 | 10 | 17 |||||||||||||||||||||||| 2.90 | 2 | 2 | 13.03 | 9 | 15 | 898 | 105.3 | 7.30 |
| Q9H2M9 | 4.95 | 1 | 5 | 5 | 7 |||||||||||||||||||||||||||| 4.95 | 5 | 7 | 1393 | 155.9 | 5.62 |
| Q9H2P9 | 9.12 | 1 | 2 | 2 | 2 |||||||||||||||||||||||||||| 9.12 | 2 | 2 | 285 | 31.6 | 5.31 |
| Q9H2T7 | 6.34 | 1 | 5 | 5 | 5 |||||||||||||||||||||||||||| 6.34 | 5 | 5 | 1088 | 124.3 | 6.44 |
| Q9H2U2 | 47.90 | 1 | 12 | 13 | 39 | 25.15 | 6 | 9 | 20.66 | 5 | 6 | 11.08 | 3 | 3 | 3.89 | 1 | 1 ||||||||||||||||||| 43.71 | 11 | 16 | 334 | 37.9 | 7.39 |
| P33908 | 1.68 | 1 | 1 | 1 | 4 |||||||||||||||||||||||||||| 1.68 | 1 | 4 | 653 | 72.9 | 6.47 |
| Q9H2U9 | 28.38 | 1 | 19 | 19 | 55 | 3.58 | 2 | 2 | 10.74 | 7 | 8 | 9.28 | 6 | 7 |||| 2.12 | 1 | 1 |||||||||||||| 27.32 | 17 | 37 | 754 | 85.6 | 6.52 |
| Q9H3G5 | 21.01 | 1 | 9 | 9 | 18 | 15.97 | 7 | 7 | 11.55 | 5 | 5 | 13.24 | 5 | 6 ||||||||||||||||||||||||| 476 | 54.1 | 5.62 |
| Q9H3H5 | 3.68 | 1 | 2 | 2 | 2 |||||||||||||||||||||||||||| 3.68 | 2 | 2 | 408 | 46.1 | 8.00 |
| Q9H3K2 | 21.45 | 1 | 8 | 8 | 29 | 6.38 | 2 | 2 | 6.38 | 2 | 2 | 2.90 | 1 | 1 |||||||||||||||||||||| 21.45 | 8 | 24 | 345 | 37.2 | 9.94 |
| Q9H3K6 | 53.49 | 1 | 5 | 5 | 14 |||||||||| 18.60 | 1 | 1 ||||||| 37.21 | 3 | 4 | 8.14 | 1 | 1 | 45.35 | 3 | 4 | 37.21 | 3 | 4 | 86 | 10.1 | 6.52 |
| Q9H3M9 | 26.76 | 1 | 6 | 7 | 11 |||||||||||||||||||||||||||| 26.76 | 7 | 11 | 355 | 40.7 | 4.96 |
| P43250 | 2.43 | 2 | 1 | 1 | 4 |||||||||||||||||||||||||||| 2.43 | 1 | 4 | 576 | 65.9 | 8.00 |
| Q9H3N1 | 8.57 | 1 | 3 | 3 | 8 | 7.50 | 2 | 2 |||| 4.29 | 1 | 1 |||||||||||||||||||||| 8.57 | 3 | 5 | 280 | 31.8 | 4.98 |
| Q9H3S7 | 2.69 | 1 | 3 | 3 | 5 | 0.00 ||||||| 0.00 |||||||||||||||||||||| 2.69 | 3 | 5 | 1636 | 178.9 | 6.92 |
| Q9H3U1 | 11.65 | 1 | 10 | 10 | 14 |||||||||||||||||||||||||||| 11.65 | 10 | 14 | 944 | 103.0 | 6.07 |
| Q9H3U7 | 17.71 | 1 | 7 | 7 | 16 | 6.28 | 2 | 2 | 2.91 | 1 | 2 ||||||||||||||||||||||||| 17.71 | 7 | 12 | 446 | 49.6 | 8.46 |
| Q9H3Z4 | 45.96 | 1 | 6 | 6 | 31 | 14.65 | 2 | 3 | 14.65 | 2 | 4 | 14.65 | 2 | 4 |||||||||||||||||||||| 45.96 | 5 | 20 | 198 | 22.1 | 5.07 |
| Q9H3Z7 | 5.33 | 1 | 2 | 2 | 2 |||||||||||||||||||||||||||| 5.33 | 2 | 2 | 469 | 52.5 | 8.57 |
| Q9H444 | 34.82 | 2 | 9 | 9 | 35 | 11.16 | 2 | 2 ||||||| 27.68 | 5 | 6 | 24.11 | 6 | 7 |||| 24.11 | 5 | 6 | 4.91 | 1 | 1 | 14.29 | 4 | 5 | 22.32 | 5 | 8 | 224 | 24.9 | 4.82 |
| Q9H492 | 17.36 | 3 | 2 | 2 | 2 |||||||||||||||||||||||||||| 17.36 | 2 | 2 | 121 | 14.3 | 8.68 |
| Q9H4A4 | 48.00 | 1 | 22 | 22 | 70 | 5.38 | 2 | 2 ||||||| 6.46 | 3 | 3 ||||||| 1.69 | 1 | 1 |||| 5.54 | 3 | 3 | 44.46 | 21 | 61 | 650 | 72.5 | 5.74 |
| Q9H4A6 | 15.77 | 1 | 3 | 3 | 6 |||||||||||||||||||||||||||| 15.77 | 3 | 6 | 298 | 33.8 | 6.44 |
| O95415 | 12.80 | 1 | 1 | 1 | 4 |||||||||||||||||||||||||||| 12.80 | 1 | 4 | 125 | 13.6 | 8.32 |
| Q9H4A9 | 11.93 | 1 | 3 | 4 | 4 |||| 4.32 | 2 | 2 |||||||||||||||||||||||| 7.61 | 2 | 2 | 486 | 53.3 | 6.40 |
| Q9H4B8 | 45.29 | 1 | 15 | 16 | 83 | 23.16 | 8 | 10 | 14.14 | 7 | 9 | 16.60 | 6 | 6 | 12.30 | 4 | 5 | 12.91 | 5 | 6 ||||||| 2.25 | 1 | 1 |||| 38.93 | 14 | 46 | 488 | 53.7 | 7.96 |
| Q9H4E7 | 3.65 | 1 | 2 | 3 | 4 |||||||||||||||||||||||| 1.43 | 1 | 1 | 2.22 | 2 | 3 | 631 | 73.9 | 6.05 |
| Q9H4F8 | 25.81 | 1 | 13 | 13 | 23 | 2.53 | 1 | 2 | 11.98 | 4 | 4 ||||||| 9.45 | 3 | 3 |||||||||||||| 21.66 | 8 | 14 | 434 | 48.1 | 8.22 |
| Q9NR96 | 0.68 | 1 | 1 | 1 | 4 |||||||||||||||||||||||||||| 0.68 | 1 | 4 | 1032 | 115.8 | 8.19 |
| Q8N7R7 | 11.14 | 1 | 1 | 3 | 4 |||||||||||||||||||||||||||| 11.14 | 3 | 4 | 359 | 40.7 | 6.07 |
| Q9H4G4 | 42.21 | 2 | 4 | 4 | 11 | 8.44 | 1 | 1 | 8.44 | 1 | 1 | 8.44 | 1 | 1 | 16.88 | 2 | 2 |||||||||| 8.44 | 1 | 1 | 8.44 | 1 | 1 | 33.77 | 3 | 4 | 154 | 17.2 | 9.41 |
| Q9H2A7 | 5.51 | 1 | 1 | 1 | 4 |||||||||||||||||||||||||||| 5.51 | 1 | 4 | 254 | 27.6 | 8.78 |
| Q9H4K1 | 63.11 | 1 | 26 | 26 | 72 |||||||||| 27.51 | 8 | 11 | 57.93 | 21 | 28 ||||||||||||||| 56.63 | 19 | 33 | 309 | 37.0 | 9.54 |
| Q9H4M9 | 35.02 | 2 | 12 | 15 | 22 | 5.06 | 2 | 2 | 2.25 | 1 | 1 |||| 4.31 | 2 | 2 ||||||||||||||||||| 30.15 | 12 | 17 | 534 | 60.6 | 6.83 |
| Q9H4Y5 | 23.46 | 1 | 5 | 5 | 7 |||||||||||||||||||||||||||| 23.46 | 5 | 7 | 243 | 28.2 | 7.56 |
| Q9H5S3 | 14.90 | 1 | 5 | 6 | 7 |||||||||||||||||||||||||||| 14.90 | 6 | 7 | 416 | 47.1 | 7.05 |
| Q9H579 | 20.70 | 1 | 9 | 9 | 14 |||| 2.28 | 1 | 1 |||||||||||||||||||||||| 20.70 | 9 | 13 | 483 | 54.8 | 6.40 |
| Q9H596 | 47.37 | 1 | 7 | 7 | 7 |||||||||||||||||||||||||||| 47.37 | 7 | 7 | 190 | 21.5 | 8.82 |
| O75843 | 2.55 | 1 | 1 | 2 | 4 |||||||||||||||||||||||||||| 2.55 | 2 | 4 | 785 | 87.1 | 6.55 |
| Q9H5F2 | 58.00 | 1 | 8 | 8 | 22 |||||||||| 18.00 | 3 | 3 | 48.00 | 6 | 6 |||||||||| 7.33 | 1 | 1 | 53.33 | 7 | 12 | 150 | 17.8 | 8.37 |
| Q9UN76 | 2.96 | 1 | 1 | 1 | 4 |||||||||||||||||||||||||||| 2.96 | 1 | 4 | 642 | 72.1 | 8.19 |
| Q9H5N1 | 8.44 | 1 | 5 | 5 | 10 | 1.76 | 1 | 1 | 1.76 | 1 | 1 |||||||||||||| 6.68 | 4 | 4 |||| 3.51 | 2 | 2 | 3.87 | 2 | 2 | 569 | 63.5 | 4.78 |
| Q9H5X1 | 12.50 | 1 | 2 | 2 | 2 |||||||||||||||||||||||||||| 12.50 | 2 | 2 | 160 | 18.3 | 4.88 |
| Q9H6K4 | 8.94 | 1 | 2 | 2 | 2 |||||||||||||||||||||||||||| 8.94 | 2 | 2 | 179 | 20.0 | 8.91 |
| Q9H6S3 | 7.83 | 1 | 5 | 5 | 7 |||||||||||||||||| 1.82 | 1 | 1 |||| 1.82 | 1 | 2 | 6.01 | 4 | 4 | 715 | 80.6 | 6.84 |
| Q9H6U8 | 5.73 | 1 | 4 | 4 | 7 |||||||||||||||||||||||||||| 5.73 | 4 | 7 | 611 | 69.8 | 8.68 |
| O95214 | 9.92 | 1 | 1 | 1 | 4 |||||||||||||||||||||||||||| 9.92 | 1 | 4 | 131 | 14.4 | 5.90 |
| Q9H6V9 | 7.38 | 1 | 2 | 2 | 2 |||||||||||||||||||||||||||| 7.38 | 2 | 2 | 325 | 37.3 | 6.54 |
| Q9H7D0 | 2.51 | 1 | 3 | 4 | 5 |||||||||||||||||||||||||||| 2.51 | 4 | 5 | 1870 | 215.2 | 7.96 |
| Q9H7D7 | 5.75 | 1 | 3 | 3 | 6 |||||||||||||||||||||||||||| 5.75 | 3 | 6 | 661 | 72.1 | 6.16 |
| Q9H7T0 | 18.46 | 1 | 17 | 17 | 31 |||| 2.69 | 2 | 2 |||||||||||||||||||||||| 17.47 | 16 | 29 | 1116 | 126.8 | 7.17 |
| Q9H7X7 | 33.51 | 1 | 4 | 4 | 5 |||||||||||||||||||||||||||| 33.51 | 4 | 5 | 185 | 20.8 | 5.15 |
| Q9H7Z7 | 12.20 | 1 | 2 | 2 | 3 |||||||||||||||||||||||||||| 12.20 | 2 | 3 | 377 | 41.9 | 9.16 |

| | | | | | | | | | | | | | | |
|---|---|---|---|---|---|---|---|---|---|---|---|---|---|---|
| Q14129 | 14.55 | 1 | 1 | 4 | 4 | | | | | | | | | |
| Q9BY27 | 14.55 | 1 | 1 | 4 | 4 | | | | | | | | | |
| Q9H832 | 5.65 | 1 | 2 | 2 | 3 | | | | | | | | | |
| Q1HG43 | 3.21 | 1 | 1 | 1 | 4 | | | | | | | | | |
| Q9H845 | 43.00 | 1 | 24 | 24 | 39 | 1.93 | 1 | 1 | 12.88 | 7 | 7 | 8.86 | 5 | 5 |
| Q9H871 | 7.16 | 1 | 2 | 2 | 3 | | | | | | | | | |
| Q9H892 | 25.96 | 1 | 16 | 16 | 27 | | | | | | | 0.99 | 1 | 1 |
| Q9H8H3 | 18.44 | 1 | 4 | 4 | 16 | 4.92 | 1 | 1 | | | | 0.00 | | |
| Q9H8X9 | 7.04 | 2 | 2 | 2 | 3 | | | | | | | | | |
| Q9H8Y8 | 7.30 | 1 | 2 | 2 | 3 | | | | | | | | | |
| Q9H993 | 43.99 | 1 | 15 | 15 | 32 | | | | | | | | | |
| Q9H9A5 | 3.76 | 1 | 3 | 3 | 3 | | | | | | | | | |
| Q9H9B4 | 6.83 | 1 | 2 | 2 | 2 | | | | | | | | | |
| Q9H9E3 | 16.18 | 1 | 11 | 11 | 13 | | | | | | | | | |
| Q9H9P8 | 12.74 | 1 | 5 | 5 | 5 | | | | | | | 0.00 | | |
| Q8NGR8 | 2.27 | 1 | 1 | 1 | 4 | | | | | | | | | |
| Q9H9Q2 | 25.76 | 1 | 4 | 4 | 7 | | | | | | | | | |
| Q9H9S3 | 7.77 | 1 | 2 | 4 | 14 | | | | 1.89 | 1 | 1 | | | |
| Q9H9S4 | 16.91 | 1 | 3 | 4 | 12 | | | | | | | | | |
| Q9H9V9 | 6.05 | 1 | 2 | 2 | 2 | | | | | | | | | |
| Q9HA64 | 8.41 | 1 | 2 | 2 | 3 | | | | | | | | | |
| Q9HA65 | 9.57 | 1 | 6 | 6 | 7 | | | | | | | | | |
| Q9HAB8 | 32.15 | 1 | 11 | 11 | 14 | | | | | | | | | |
| Q9HAE3 | 48.34 | 1 | 9 | 9 | 18 | 10.90 | 2 | 2 | 10.90 | 2 | 2 | | | |
| Q9HAQ2 | 35.44 | 2 | 29 | 30 | 99 | 3.42 | 2 | 2 | | | | 1.90 | 1 | 1 |
| Q9HAS0 | 35.35 | 2 | 13 | 13 | 23 | 2.78 | 1 | 1 | 6.31 | 2 | 2 | | | |
| Q9HAT0 | 85.38 | 1 | 8 | 19 | 448 | 57.55 | 10 | 21 | 54.72 | 10 | 20 | 58.96 | 11 | 21 |
| Q9HAU5 | 3.07 | 1 | 3 | 3 | 4 | | | | 0.00 | | | 0.00 | | |
| [illegible] | | | | | | | | | | | | | | |

| | | | | | | | | | | | | | | | | | | | | | | | | | | | | | | | | | | | | | | |
|---|---|---|---|---|---|---|---|---|---|---|---|---|---|---|---|---|---|---|---|---|---|---|---|---|---|---|---|---|---|---|---|---|---|---|---|---|---|---|
| Q9NZT1 | 31.51 | 1 | 6 | 6 | 31 | | | | 9.59 | 1 | 1 | | | | 25.34 | 2 | 2 | 15.75 | 3 | 4 | 9.59 | 2 | 2 | 31.51 | 6 | 8 | 15.75 | 3 | 4 | 31.51 | 5 | 7 | 9.59 | 1 | 3 | 146 | 15.9 | 4.44 |
| Q9NZZ3 | 21.92 | 1 | 4 | 4 | 6 | 7.31 | 1 | 1 | | | | | | | 16.89 | 2 | 2 | 5.02 | 1 | 1 | | | | | | | | | | | | | 3.20 | 1 | 2 | 219 | 24.6 | 4.83 |
| Q9P000 | 30.30 | 1 | 4 | 4 | 4 | | | | | | | | | | | | | | | | | | | | | | | | | | | | 30.30 | 4 | 4 | 198 | 21.8 | 5.88 |
| Q8IWU5 | 0.57 | 1 | 1 | 1 | 3 | | | | | | | | | | | | | | | | | | | | | | | | | | | | 0.57 | 1 | 3 | 870 | 100.4 | 9.17 |
| Q9P032 | 26.29 | 1 | 4 | 4 | 4 | | | | | | | | | | | | | | | | | | | | | | | | | | | | 26.29 | 4 | 4 | 175 | 20.3 | 8.82 |
| Q12933 | 2.00 | 1 | 1 | 1 | 3 | | | | | | | | | | | | | | | | | | | | | | | | | | | | 2.00 | 1 | 3 | 501 | 55.8 | 7.53 |
| Q9P035 | 16.85 | 1 | 5 | 5 | 9 | | | | | | | | | | | | | | | | | | | | | | | | | | | | 16.85 | 5 | 9 | 362 | 43.1 | 8.94 |
| Q9P281 | 0.54 | 2 | 1 | 2 | 3 | | | | | | | | | | | | | | | | | | | | | | | | | | | | 0.54 | 2 | 3 | 2608 | 276.8 | 8.81 |
| Q9P0J0 | 55.56 | 1 | 9 | 9 | 14 | | | | | | | | | | | | | | | | | | | | | | | | | | | | 55.56 | 9 | 14 | 144 | 16.7 | 8.43 |
| Q9P0J1 | 7.26 | 1 | 2 | 2 | 4 | | | | | | | | | | | | | | | | | | | | | | | | | | | | 7.26 | 2 | 4 | 537 | 61.0 | 6.67 |
| Q9P0J7 | 16.54 | 1 | 5 | 5 | 6 | | | | | | | | | | | | | 3.41 | 1 | 1 | | | | | | | | | | | | | 16.01 | 4 | 5 | 381 | 41.9 | 5.66 |
| Q9P0L0 | 49.00 | 1 | 10 | 11 | 50 | 22.89 | 4 | 5 | 21.29 | 4 | 5 | 14.86 | 3 | 3 | | | | 4.42 | 1 | 2 | | | | | | | | | | | | | 40.96 | 10 | 35 | 249 | 27.9 | 8.62 |
| Q9P0V9 | 56.39 | 1 | 16 | 20 | 58 | 1.98 | 1 | 1 | 6.39 | 2 | 2 | 3.08 | 1 | 1 | | | | 28.85 | 11 | 13 | | | | | | | 7.49 | 3 | 3 | 0.00 | | | 47.80 | 15 | 36 | 454 | 52.6 | 6.80 |
| Q9P1F3 | 43.21 | 1 | 3 | 3 | 4 | | | | | | | 19.75 | 1 | 1 | | | | | | | | | | | | | | | | | | | 43.21 | 3 | 3 | 81 | 9.1 | 6.29 |
| Q9P1V8 | 5.79 | 1 | 5 | 5 | 11 | | | | | | | | | | 4.01 | 3 | 3 | 5.79 | 3 | 3 | | | | | | | | | | | | | 3.26 | 2 | 5 | 674 | 77.1 | 4.45 |
| Q9P1Z9 | 14.16 | 2 | 19 | 20 | 38 | | | | | | | 0.36 | 1 | 1 | | | | | | | | | | | | | | | | | | | 14.16 | 20 | 37 | 1646 | 191.0 | 6.04 |
| Q9P0S3 | 21.57 | 2 | 1 | 3 | 3 | | | | | | | | | | | | | | | | | | | | | | | | | | | | 21.57 | 3 | 3 | 153 | 17.4 | 9.64 |
| Q9P225 | 39.48 | 6 | 165 | 167 | 342 | 0.20 | 1 | 1 | 0.70 | 3 | 3 | 0.63 | 3 | 3 | 2.62 | 11 | 11 | 1.29 | 6 | 6 | | | | | | | | | | | | | 39.24 | 166 | 318 | 4427 | 507.4 | 6.37 |
| Q9P253 | 8.32 | 2 | 5 | 6 | 7 | 1.85 | 1 | 1 | 1.13 | 1 | 1 | | | | | | | | | | | | | | | | | | | | | | 5.34 | 4 | 5 | 973 | 110.1 | 6.07 |
| Q9P260 | 4.61 | 1 | 4 | 4 | 4 | | | | 1.32 | 1 | 1 | | | | | | | | | | | | | | | | | | | | | | 3.29 | 3 | 3 | 1216 | 134.5 | 5.45 |
| Q9P2D3 | 2.99 | 1 | 6 | 6 | 7 | | | | | | | | | | 0.00 | | | 0.00 | | | | | | | | | | | | 0.00 | | | 2.99 | 6 | 7 | 2071 | 224.2 | 7.17 |
| Q9P2D7 | 41.11 | 2 | 155 | 161 | 414 | 0.92 | 3 | 3 | 0.58 | 2 | 2 | 0.21 | 1 | 1 | 7.30 | 28 | 31 | 7.64 | 27 | 30 | | | | | | | | | | 0.00 | | | 39.01 | 148 | 347 | 4330 | 493.6 | 5.94 |
| Q9P2E9 | 9.15 | 1 | 10 | 10 | 16 | | | | | | | 3.48 | 3 | 3 | | | | | | | | | | 0.00 | | | | | | 0.00 | | | 7.87 | 9 | 13 | 1410 | 152.4 | 8.60 |
| Q9P2H3 | 8.88 | 1 | 5 | 5 | 6 | | | | | | | | | | | | | | | | | | | | | | | | | | | | 8.88 | 5 | 6 | 777 | 88.0 | 7.69 |
| Q9P2J5 | 34.69 | 1 | 34 | 34 | 90 | 0.60 | 1 | 1 | 7.74 | 6 | 7 | 0.94 | 1 | 1 | | | | | | | | | | | | | | | | 2.55 | 2 | 2 | 32.65 | 32 | 79 | 1176 | 134.4 | 7.30 |
| Q9P2K2 | 3.39 | 1 | 2 | 2 | 3 | | | | | | | | | | | | | | | | | | | | | | | | | | | | 3.39 | 2 | 3 | 825 | 93.5 | 5.21 |
| Q9P2L0 | 15.07 | 1 | 13 | 13 | 27 | | | | | | | 1.27 | 1 | 1 | | | | | | | | | | | | | | | | | | | 13.80 | 12 | 26 | 1181 | 133.5 | 6.38 |
| Q9P2M1 | 25.94 | 1 | 9 | 9 | 15 | | | | | | | | | | | | | 2.59 | 1 | 1 | | | | | | | | | | | | | 25.94 | 9 | 14 | 347 | 39.8 | 7.90 |
| Q9P2M7 | 1.92 | 3 | 3 | 3 | 4 | | | | 0.00 | | | | | | | | | 0.00 | | | | | | 0.50 | 1 | 1 | | | | | | | 1.42 | 2 | 3 | 1197 | 136.3 | 5.54 |
| Q9P2R3 | 17.02 | 1 | 15 | 15 | 31 | | | | | | | | | | | | | | | | | | | | | | | | | | | | 17.02 | 15 | 31 | 1169 | 128.3 | 6.10 |
| P10072 | 1.37 | 1 | 1 | 1 | 3 | | | | | | | | | | | | | | | | | | | | | | | | | | | | 1.37 | 1 | 3 | 659 | 75.1 | 9.25 |
| Q6L9W6 | 0.90 | 1 | 1 | 1 | 3 | | | | | | | | | | | | | | | | | | | | | | | | | | | | 0.90 | 1 | 3 | 998 | 114.9 | 7.49 |
| Q9P2R7 | 32.61 | 1 | 14 | 14 | 38 | 11.02 | 5 | 5 | 19.22 | 7 | 7 | 20.52 | 8 | 10 | 2.16 | 1 | 1 | 2.16 | 1 | 1 | | | | | | | | | | | | | 25.05 | 11 | 14 | 463 | 50.3 | 7.42 |
| Q9P2S6 | 8.82 | 1 | 6 | 6 | 10 | | | | | | | | | | | | | | | | | | | | | | | | | | | | 8.82 | 6 | 10 | 941 | 105.4 | 6.73 |
| Q9P2T0 | 22.43 | 2 | 9 | 9 | 24 | | | | | | | | | | | | | 1.85 | 1 | 1 | | | | | | | | | | | | | 22.43 | 9 | 23 | 379 | 43.4 | 9.04 |
| Q8N668 | 14.21 | 1 | 1 | 2 | 3 | | | | | | | | | | | | | | | | | | | | | | | | | | | | 14.21 | 2 | 3 | 190 | 21.2 | 6.20 |
| Q9P2T1 | 15.52 | 1 | 3 | 5 | 5 | | | | | | | | | | | | | | | | | | | | | | | | | | | | 15.52 | 5 | 5 | 348 | 37.9 | 7.23 |
| Q9P2W9 | 10.45 | 1 | 2 | 2 | 3 | | | | 3.58 | 1 | 1 | | | | | | | | | | | | | | | | | | | | | | 6.87 | 1 | 2 | 335 | 38.6 | 5.49 |
| Q96IW7 | 3.58 | 1 | 1 | 1 | 3 | | | | | | | | | | | | | | | | | | | | | | | | | | | | 3.58 | 1 | 3 | 307 | 34.9 | 8.24 |
| Q9P2X3 | 19.06 | 1 | 6 | 6 | 6 | | | | | | | | | | | | | | | | | | | | | | | | | 3.44 | 1 | 1 | 15.63 | 5 | 5 | 320 | 36.5 | 4.97 |
| Q92959 | 2.02 | 1 | 1 | 1 | 3 | | | | | | | | | | | | | | | | | | | | | | | | | | | | 2.02 | 1 | 3 | 643 | 70.0 | 8.84 |
| Q9UBB4 | 40.00 | 2 | 15 | 16 | 38 | | | | | | | 0.00 | | | 3.16 | 1 | 1 | | | | | | | | | | | | | | | | 40.00 | 16 | 37 | 475 | 53.5 | 5.25 |
| Q9UBF2 | 25.14 | 1 | 12 | 15 | 36 | 2.30 | 2 | 2 | 1.38 | 1 | 1 | 1.38 | 1 | 1 | | | | | | | | | | | | | 0.92 | 1 | 2 | | | | 24.23 | 14 | 30 | 871 | 97.6 | 5.81 |
| Q9UBG3 | 12.53 | 1 | 4 | 4 | 5 | | | | | | | 12.53 | 4 | 5 | | | | | | | | | | | | | | | | | | | | | | 495 | 53.5 | 6.10 |
| Q9UBI1 | 10.26 | 1 | 2 | 2 | 2 | | | | | | | | | | | | | | | | | | | | | | | | | | | | 10.26 | 2 | 2 | 195 | 22.1 | 5.99 |
| Q9UBI6 | 37.50 | 1 | 2 | 2 | 2 | | | | | | | | | | | | | | | | | | | | | | | | | | | | 37.50 | 2 | 2 | 72 | 8.0 | 8.97 |
| Q9UBK7 | 48.25 | 2 | 9 | 9 | 19 | | | | | | | | | | | | | | | | | | | | | | | | | 7.89 | 1 | 1 | 48.25 | 9 | 18 | 228 | 26.1 | 4.89 |
| Q9UBM7 | 6.32 | 1 | 3 | 3 | 5 | | | | | | | | | | | | | | | | | | | | | | | | | | | | 6.32 | 3 | 5 | 475 | 54.5 | 8.70 |
| Q9UBN7 | 7.49 | 1 | 6 | 6 | 13 | | | | | | | | | | | | | | | | | | | | | | | | | 0.91 | 1 | 1 | 7.49 | 6 | 12 | 1215 | 131.3 | 5.30 |
| Q9UBP9 | 6.91 | 1 | 2 | 2 | 6 | | | | | | | | | | | | | | | | | | | | | | | | | | | | 6.91 | 2 | 6 | 304 | 34.5 | 7.90 |
| Q9UBQ0 | 45.05 | 1 | 8 | 8 | 22 | | | | | | | | | | | | | | | | | | | 5.49 | 1 | 1 | | | | 12.64 | 1 | 1 | 45.05 | 7 | 20 | 182 | 20.5 | 6.79 |
| Q9UBQ5 | 18.35 | 1 | 3 | 3 | 5 | | | | | | | | | | | | | | | | | | | | | | | | | | | | 18.35 | 3 | 5 | 218 | 25.0 | 4.93 |
| Q9GZP9 | 12.55 | 1 | 1 | 1 | 3 | | | | | | | | | | | | | | | | | | | | | | | | | | | | 12.55 | 1 | 3 | 239 | 27.5 | 7.28 |
| Q9UBQ7 | 50.61 | 2 | 16 | 16 | 44 | 8.54 | 2 | 2 | 8.54 | 2 | 2 | 3.35 | 1 | 1 | 18.90 | 4 | 4 | 3.35 | 1 | 1 | | | | 10.67 | 2 | 2 | | | | 10.67 | 2 | 3 | 50.30 | 14 | 29 | 328 | 35.6 | 7.39 |
| Q9UBR2 | 18.48 | 2 | 5 | 5 | 5 | 7.59 | 2 | 2 | | | | 8.58 | 2 | 2 | | | | | | | | | | | | | 2.31 | 1 | 1 | | | | | | | 303 | 33.8 | 7.11 |
| Q9UBS3 | 43.50 | 1 | 9 | 9 | 33 | 24.22 | 4 | 4 | 21.97 | 4 | 4 | 43.50 | 7 | 10 | 6.28 | 1 | 1 | 6.28 | 1 | 1 | | | | | | | | | | | | | 43.05 | 8 | 13 | 223 | 25.5 | 8.27 |
| Q9UBS4 | 36.03 | 1 | 12 | 12 | 30 | 8.94 | 2 | 2 | 7.54 | 2 | 2 | 24.86 | 6 | 6 | 4.47 | 1 | 1 | 9.78 | 2 | 2 | | | | | | | 5.31 | 1 | 2 | 3.35 | 1 | 1 | 22.91 | 8 | 14 | 358 | 40.5 | 6.18 |
| Q9UBV2 | 26.32 | 1 | 12 | 12 | 23 | | | | | | | | | | | | | | | | | | | | | | | | | | | | 26.32 | 12 | 23 | 794 | 88.7 | 5.39 |
| P13866 | 2.26 | 1 | 1 | 1 | 3 | | | | | | | | | | | | | | | | | | | | | | | | | | | | 2.26 | 1 | 3 | 664 | 73.4 | 7.68 |
| Q9UBV8 | 12.32 | 1 | 4 | 4 | 7 | | | | | | | | | | | | | | | | | | | | | | | | | 3.17 | 1 | 1 | 12.32 | 4 | 6 | 284 | 30.4 | 6.54 |
| Q9UBW8 | 29.82 | 1 | 6 | 6 | 7 | | | | | | | | | | | | | | | | | | | | | | | | | | | | 29.82 | 6 | 7 | 275 | 30.3 | 8.22 |
| P00414 | 5.36 | 1 | 1 | 1 | 3 | | | | | | | | | | | | | | | | | | | | | | | | | | | | 5.36 | 1 | 3 | 261 | 29.9 | 7.31 |
| Q8N138 | 21.57 | 1 | 1 | 3 | 3 | | | | | | | | | | | | | | | | | | | | | | | | | | | | 21.57 | 3 | 3 | 153 | 17.5 | 9.64 |
| Q9UBX1 | 25.00 | 1 | 12 | 12 | 29 | 13.22 | 6 | 6 | 15.08 | 7 | 7 | 4.55 | 2 | 2 | | | | | | | | | | | | | | | | | | | 16.94 | 8 | 14 | 484 | 53.3 | 8.22 |
| Q9UBX3 | 54.36 | 1 | 13 | 13 | 42 | 14.98 | 3 | 3 | 30.66 | 7 | 7 | 13.59 | 4 | 4 | | | | | | | | | | | | | | | | | | | 51.22 | 12 | 28 | 287 | 31.3 | 9.54 |
| Q6NTF9 | 4.40 | 1 | 1 | 1 | 3 | | | | | | | | | | | | | | | | | | | | | | | | | | | | 4.40 | 1 | 3 | 364 | 39.2 | 9.32 |
| Q9UBX7 | 7.45 | 1 | 2 | 2 | 2 | 0.00 | | | | | | 3.55 | 1 | 1 | | | | | | | | | | | | | | | | | | | 3.90 | 1 | 1 | 282 | 31.0 | 8.94 |
| Q8TAA9 | 1.72 | 1 | 1 | 1 | 3 | | | | | | | | | | | | | | | | | | | | | | | | | | | | 1.72 | 1 | 3 | 524 | 59.9 | 8.81 |
| Q9UDR5 | 9.07 | 1 | 8 | 8 | 8 | | | | | | | | | | | | | | | | | | | | | | | | | | | | 9.07 | 8 | 8 | 926 | 102.1 | 6.64 |
| Q9UBM1 | 6.53 | 1 | 1 | 1 | 3 | | | | | | | | | | | | | | | | | | | | | | | | | | | | 6.53 | 1 | 3 | 199 | 22.1 | 8.76 |
| Q9GIY3 | 15.41 | 2 | 1 | 3 | 3 | | | | | | | | | | | | | | | | | | | | | | | | | | | | 15.41 | 3 | 3 | 266 | 30.1 | 8.05 |
| Q9UDY2 | 4.54 | 1 | 5 | 5 | 10 | | | | | | | | | | | | | | | | | | | | | | 0.67 | 1 | 1 | 2.86 | 3 | 3 | 1.01 | 1 | 6 | 1190 | 133.9 | 7.40 |
| Q9UDY8 | 5.70 | 1 | 4 | 4 | 4 | | | | | | | | | | | | | | | | | | | | | | | | | | | | 5.70 | 4 | 4 | 824 | 92.2 | 5.73 |
| Q8WW43 | 4.67 | 1 | 1 | 1 | 3 | | | | | | | | | | | | | | | | | | | | | | | | | | | | 4.67 | 1 | 3 | 257 | 28.4 | 9.06 |
| Q9UEU0 | 12.93 | 1 | 2 | 2 | 3 | | | | | | | | | | | | | | | | | | | | | | | | | | | | 12.93 | 2 | 3 | 232 | 26.7 | 9.04 |
| Q9UEW8 | 37.80 | 1 | 11 | 15 | 64 | 2.75 | 1 | 1 | 4.04 | 1 | 1 | 4.95 | 2 | 2 | 2.75 | 1 | 1 | 2.75 | 1 | 1 | | | | | | | 2.75 | 1 | 1 | 2.75 | 1 | 1 | 37.80 | 15 | 56 | 545 | 59.4 | 6.29 |
| Q9UF47 | 23.12 | 1 | 3 | 3 | 3 | | | | | | | 14.57 | 2 | 2 | | | | | | | | | | | | | | | | | | | 8.54 | 1 | 1 | 199 | 22.5 | 5.30 |
| Q3MIR4 | 6.27 | 1 | 1 | 2 | 3 | | | | | | | | | | | | | | | | | | | | | | | | | | | | 6.27 | 2 | 3 | 351 | 38.9 | 7.88 |
| Q9UFE4 | 54.30 | 1 | 65 | 65 | 199 | 2.55 | 1 | 1 | 1.06 | 1 | 1 | | | | 22.00 | 20 | 23 | 39.00 | 38 | 48 | | | | | | | | | | 1.38 | 1 | 1 | 44.31 | 48 | 125 | 941 | 109.8 | 6.44 |
| Q9UFH2 | 52.60 | 1 | 244 | 247 | 982 | 12.26 | 42 | 48 | 13.07 | 49 | 52 | 13.02 | 51 | 53 | 8.23 | 30 | 31 | 6.04 | 25 | 28 | | | | 0.56 | 2 | 2 | | | | 0.33 | 1 | 1 | 50.28 | 234 | 767 | 4485 | 511.5 | 5.77 |
| Q9UFN0 | 44.53 | 2 | 10 | 10 | 24 | 10.53 | 2 | 2 | 29.15 | 6 | 7 | 15.79 | 3 | 3 | | | | | | | | | | | | | | | | | | | 44.13 | 8 | 12 | 247 | 28.4 | 9.16 |
| Q9UG01 | 18.58 | 1 | 28 | 28 | 38 | | | | | | | | | | | | | | | | | | | | | | | | | | | | 18.58 | 28 | 38 | 1749 | 197.5 | 6.13 |
| Q9UGI8 | 20.90 | 1 | 6 | 6 | 12 | | | | | | | | | | | | | | | | | | | 2.85 | 1 | 1 | | | | | | | 20.90 | 6 | 11 | 421 | 48.0 | 7.68 |
| Q8N687 | 3.85 | 1 | 1 | 1 | 3 | | | | | | | | | | | | | | | | | | | | | | | | | | | | 3.85 | 1 | 3 | 156 | 17.5 | 5.49 |
| Q9UGM3 | 10.69 | 1 | 2 | 2 | 3 | | | | | | | 10.69 | 2 | 2 | | | | | | | | | | | | | | | | | | | 6.96 | 1 | 1 | 2413 | 260.6 | 5.44 |
| Q9UGM6 | 5.83 | 1 | 2 | 2 | 2 | | | | | | | | | | | | | | | | | | | | | | | | | | | | 5.83 | 2 | 2 | 360 | 40.1 | 9.28 |
| Q9UGP8 | 24.74 | 1 | 15 | 15 | 30 | | | | 4.34 | 3 | 3 | | | | | | | | | | | | | | | | | | | | | | 24.61 | 14 | 27 | 760 | 87.9 | 5.31 |
| Q9H2S1 | 1.55 | 1 | 1 | 1 | 3 | | | | | | | | | | | | | | | | | | | | | | | | | | | | 1.55 | 1 | 3 | 579 | 63.7 | 9.52 |
| Q9UGV2 | 49.07 | 1 | 11 | 12 | 49 | 4.00 | 1 | 1 | 10.67 | 3 | 3 | 6.67 | 2 | 2 | 6.67 | 2 | 2 | | | | | | | | | | | | | 6.40 | 2 | 2 | 49.07 | 12 | 39 | 375 | 41.4 | 5.31 |
| Q86V88 | 7.39 | 1 | 1 | 1 | 3 | | | | | | | | | | | | | | | | | | | | | | | | | | | | 7.39 | 1 | 3 | 176 | 20.1 | 6.39 |
| Q502X0 | 17.72 | 1 | 1 | 1 | 3 | | | | | | | | | | | | | | | | | | | | | | | | | | | | 17.72 | 1 | 3 | 79 | 8.9 | 5.67 |
| Q9UHA4 | 41.13 | 1 | 4 | 4 | 4 | | | | | | | | | | | | | | | | | | | | | | | | | 0.00 | | | 41.13 | 4 | 4 | 124 | 13.6 | 7.34 |
| Q9UHB9 | 14.19 | 1 | 5 | 5 | 10 | | | | | | | | | | | | | | | | | | | | | | | | | | | | 14.19 | 5 | 10 | 627 | 70.7 | 8.56 |
| Q9UHD1 | 26.81 | 1 | 7 | 7 | 18 | | | | | | | | | | 5.42 | 1 | 1 | 8.43 | 2 | 2 | | | | | | | 7.53 | 2 | 2 | 8.43 | 2 | 2 | 21.69 | 5 | 11 | 332 | 37.5 | 7.87 |
| Q9UHD2 | 24.14 | 2 | 13 | 13 | 28 | | | | | | | | | | | | | | | | | | | | | | | | | | | | 24.14 | 13 | 26 | 729 | 83.6 | 6.79 |
| Q3MIN7 | 0.85 | 1 | 1 | 1 | 3 | | | | | | | | | | | | | | | | | | | | | | | | | | | | 0.85 | 1 | 3 | 710 | 78.0 | 7.68 |
| Q9UHD8 | 10.58 | 1 | 5 | 5 | 7 | | | | | | | | | | | | | | | | | | | | | | | | | | | | 10.58 | 5 | 7 | 586 | 65.4 | 8.97 |
| Q9UHG3 | 40.00 | 1 | 16 | 16 | 47 | | | | 8.91 | 4 | 5 | 15.05 | 6 | 8 | | | | | | | | | | | | | | | | | | | 37.23 | 15 | 34 | 505 | 56.6 | 6.18 |
| K7EJ46 | 12.59 | 1 | 1 | 1 | 3 | | | | | | | | | | | | | | | | | | | | | | | | | | | | 12.59 | 1 | 3 | 135 | 14.6 | 6.52 |
| Q9UHI8 | 14.58 | 1 | 10 | 10 | 17 | | | | | | | 3.41 | 2 | 2 | | | | | | | | | | | | | | | | 1.03 | 1 | 1 | 14.58 | 10 | 14 | 967 | 105.3 | 6.83 |
| Q496A3 | 5.00 | 1 | 1 | 1 | 3 | | | | | | | | | | | | | | | | | | | | | | | | | | | | 5.00 | 1 | 3 | 300 | 33.7 | 7.46 |
| Q9P0N9 | 8.19 | 1 | 1 | 2 | 3 | | | | | | | | | | | | | | | | | | | | | | | | | | | | 8.19 | 2 | 3 | 293 | 33.9 | 7.94 |
| Q9UHJ6 | 5.65 | 1 | 2 | 2 | 2 | | | | | | | | | | | | | | | | | | | | | | | | | | | | 5.65 | 2 | 2 | 478 | 51.5 | 6.83 |
| Q9UHL4 | 25.81 | 1 | 10 | 10 | 17 | 13.82 | 5 | 5 | 3.66 | 2 | 2 | 6.10 | 2 | 2 | | | | | | | | | | | | | | | | | | | 16.67 | 6 | 8 | 492 | 54.3 | 6.32 |
| Q9UHP6 | 50.86 | 1 | 16 | 16 | 49 | | | | | | | | | | 18.10 | 6 | 7 | 8.91 | 2 | 2 | | | | | | | 0.00 | | | 3.74 | 1 | 1 | 50.57 | 15 | 39 | 348 | 38.6 | 6.90 |
| Q9UHQ4 | 14.11 | 1 | 4 | 4 | 4 | | | | | | | | | | | | | | | | | | | | | | | | | | | | 14.11 | 4 | 4 | 241 | 28.3 | 9.54 |
| Q9UHQ9 | 19.34 | 1 | 6 | 6 | 8 | | | | | | | | | | | | | | | | | | | | | | | | | | | | 19.34 | 6 | 8 | 305 | 34.1 | 9.38 |
| Q9UHR6 | 15.63 | 1 | 5 | 5 | 10 | | | | | | | | | | 2.98 | 1 | 1 | | | | | | | 2.73 | 1 | 1 | | | | 5.71 | 2 | 2 | 15.63 | 5 | 6 | 403 | 42.9 | 5.99 |
| Q9UHV9 | 22.73 | 1 | 3 | 3 | 6 | 14.94 | 2 | 2 | | | | | | | | | | | | | | | | 7.79 | 1 | 1 | | | | 13.64 | 2 | 2 | 7.79 | 1 | 1 | 154 | 16.6 | 6.58 |
| Q5TID7 | 1.38 | 1 | 1 | 1 | 2 | 0.00 | | | 0.00 | | | 0.00 | | | 0.00 | | | 0.00 | | | | | | | | | | | | | | | 1.38 | 1 | 2 | 509 | 60.1 | 5.91 |
| Q9UHW9 | 5.65 | 2 | 6 | 6 | 9 | | | | | | | | | | | | | | | | | | | | | | | | | | | | 5.65 | 6 | 9 | 1150 | 127.5 | 7.08 |
| Q86YV0 | 1.19 | 1 | 1 | 1 | 2 | 1.19 | 1 | 1 | 1.19 | 1 | 1 | | | | | | | | | | | | | | | | | | | | | | | | | 1011 | 111.8 | 8.75 |
| Q9UHY1 | 12.34 | 1 | 5 | 6 | 12 | | | | | | | | | | | | | | | | | | | 2.43 | 1 | 1 | | | | 2.43 | 1 | 1 | 12.15 | 5 | 10 | 535 | 59.8 | 5.08 |
| Q9P2G4 | 0.77 | 1 | 1 | 1 | 2 | 0.77 | 1 | 1 | | | | 0.77 | 1 | 1 | | | | | | | | | | | | | | | | | | | 0.00 | | | 905 | 100.3 | 7.17 |
| Q63HR2 | 0.64 | 1 | 1 | 1 | 2 | 0.00 | | | 0.64 | 1 | 1 | 0.64 | 1 | 1 | | | | | | | | | | | | | | | | | | | 0.00 | | | 1409 | 152.5 | 8.35 |
| CON_009062951 | 1.70 | 2 | 1 | 1 | 2 | 1.70 | 1 | 1 | | | | 1.70 | 1 | 1 | | | | | | | | | | | | | | | | | | | | | | 705 | 76.5 | 5.17 |
| Q9UHY7 | 21.84 | 1 | 4 | 4 | 5 | | | | | | | | | | | | | | | | | | | | | | | | | 4.60 | 1 | 1 | 17.24 | 3 | 4 | 261 | 28.9 | 4.78 |
| Q5T1M5 | 1.64 | 2 | 1 | 2 | 2 | 0.49 | 1 | 1 | | | | | | | 0.00 | | | 0.00 | | | | | | | | | | | | 1.15 | 1 | 1 | 0.00 | | | 1219 | 133.5 | 5.20 |
| Q9UI09 | 47.59 | 1 | 6 | 6 | 9 | | | | | | | | | | | | | | | | | | | | | | | | | | | | 47.59 | 6 | 9 | 145 | 17.1 | 9.63 |
| Q96CN9 | 1.68 | 1 | 1 | 2 | 2 | 0.77 | 1 | 1 | | | | | | | | | | 0.00 | | | | | | | | | | | | | | | 0.90 | 1 | 1 | 775 | 87.8 | 5.45 |
| O43504 | 15.38 | 1 | 1 | 1 | 2 | 15.38 | 1 | 1 | | | | 15.38 | 1 | 1 | | | | | | | | | | | | | | | | | | | | | | 91 | 9.6 | 4.87 |
| Q5VZK9 | 0.58 | 1 | 1 | 1 | 2 | 0.00 | | | 0.00 | | | 0.00 | | | | | | | | | | | | | | | | | | | | | 0.58 | 1 | 2 | 1371 | 151.5 | 7.85 |
| Q9UI10 | 16.63 | 1 | 6 | 6 | 18 | | | | | | | | | | | | | | | | | | | | | | | | | | | | 16.63 | 6 | 18 | 523 | 57.5 | 9.38 |
| Q9NRZ9 | 0.72 | 2 | 1 | 1 | 2 | 0.72 | 1 | 1 | 0.72 | 1 | 1 | 0.00 | | | 0.00 | | | 0.00 | | | | | | 0.00 | | | 0.00 | | | 0.00 | | | 0.00 | | | 838 | 97.0 | 7.93 |
| Q9UI12 | 30.64 | 1 | 11 | 11 | 31 | | | | 0.00 | | | 4.76 | 1 | 1 | 0.00 | | | 0.00 | | | | | | 0.00 | | | 0.00 | | | 2.90 | 1 | 1 | 25.67 | 9 | 29 | 483 | 55.8 | 6.48 |
| Q9UI14 | 18.92 | 1 | 3 | 3 | 6 | 7.57 | 1 | 1 | 7.57 | 1 | 1 | | | | | | | | | | | | | | | | | | | | | | 18.92 | 3 | 4 | 185 | 20.6 | 7.34 |
| Q9UMN6 | 0.44 | 1 | 1 | 1 | 2 | 0.00 | | | 0.00 | | | 0.00 | | | 0.00 | | | 0.00 | | | | | | 0.00 | | | | | | 0.00 | | | 0.44 | 1 | 2 | 2715 | 293.3 | 8.22 |
| Q9UI26 | 12.10 | 1 | 7 | 7 | 9 | | | | | | | | | | | | | 0.00 | | | | | | | | | | | | | | | 12.10 | 7 | 9 | 975 | 112.5 | 5.25 |
| Q8TAC9 | 4.68 | 1 | 1 | 1 | 2 | 4.68 | 1 | 1 | | | | | | | | | | | | | | | | | | | | | | | | | 4.68 | 1 | 1 | 235 | 26.1 | 8.54 |
| Q9UI30 | 32.00 | 1 | 3 | 3 | 4 | | | | | | | | | | | | | | | | | | | | | | | | | | | | 32.00 | 3 | 4 | 125 | 14.2 | 5.26 |
| Q9UI38 | 21.56 | 1 | 5 | 5 | 20 | 14.81 | 4 | 4 | 6.23 | 2 | 2 | | | | 2.34 | 1 | 1 | 3.38 | 1 | 1 | | | | | | | | | | | | | 18.18 | 4 | 12 | 385 | 43.1 | 8.21 |
| P56378 | 12.07 | 1 | 1 | 1 | 2 | 12.07 | 1 | 1 | 12.07 | 1 | 1 | 0.00 | | | 0.00 | | | | | | | | | | | | | | | | | | | | | 58 | 6.7 | 10.08 |
| P52179 | 0.47 | 1 | 1 | 1 | 2 | 0.47 | 1 | 1 | | | | 0.47 | 1 | 1 | | | | | | | | | | | | | | | | | | | | | | 1685 | 187.5 | 6.93 |
| Q9UI46 | 64.95 | 1 | 44 | 44 | 203 | 15.45 | 9 | 9 | 17.17 | 10 | 12 | 14.31 | 8 | 8 | 17.45 | 9 | 9 | 22.60 | 11 | 14 | | | | | | | 3.00 | 1 | 2 | | | | 58.37 | 38 | 149 | 699 | 79.2 | 6.87 |
| Q8TDC0 | 2.39 | 1 | 1 | 1 | 2 | 0.00 | | | 0.00 | | | 0.00 | | | 0.00 | | | | | | | | | 0.00 | | | 0.00 | | | 2.39 | 1 | 1 | 2.39 | 1 | 1 | 251 | 27.1 | 9.38 |
| Q9UIA9 | 49.59 | 1 | 46 | 46 | 171 | 6.62 | 6 | 6 | 7.36 | 7 | 7 | 1.29 | 1 | 1 | | | | | | | | | | | | | | | | | | | 49.49 | 45 | 157 | 1087 | 123.8 | 6.32 |
| Q9BSV6 | 4.84 | 2 | 1 | 2 | 2 | 1.94 | 1 | 1 | | | | | | | | | | | | | | | | | | | | | | 2.90 | 1 | 1 | 0.00 | | | 310 | 33.6 | 8.43 |
| Q9UIC8 | 19.76 | 1 | 7 | 7 | 9 | | | | 2.99 | 1 | 1 | | | | | | | | | | | | | | | | | | | | | | 19.76 | 7 | 8 | 334 | 38.4 | 6.04 |
| Q9UID3 | 10.49 | 1 | 7 | 7 | 10 | | | | | | | | | | | | | | | | | | | | | | | | | | | | 10.49 | 7 | 10 | 782 | 86.0 | 6.47 |
| Q9UIF3 | 89.53 | 1 | 54 | 54 | 365 | 39.30 | 13 | 15 | 28.14 | 10 | 12 | 6.74 | 2 | 2 | 39.30 | 17 | 22 | 59.77 | 28 | 33 | 2.09 | 1 | 1 | 48.37 | 26 | 38 | 45.58 | 21 | 27 | 60.47 | 36 | 46 | 76.74 | 34 | 169 | 430 | 49.6 | 5.55 |
| Q9UII2 | 44.34 | 1 | 9 | 9 | 25 | 16.98 | 2 | 3 | 34.91 | 4 | 6 | 16.98 | 2 | 4 | 10.38 | 2 | 2 | 34.91 | 7 | 8 | | | | | | | | | | | | | 7.55 | 2 | 2 | 106 | 12.2 | 9.35 |
| Q9UII6 | 30.81 | 1 | 4 | 4 | 5 | | | | | | | | | | | | | | | | | | | | | | | | | | | | 30.81 | 4 | 5 | 198 | 22.1 | 9.14 |
| P22894 | 3.43 | 1 | 1 | 1 | 2 | 3.43 | 1 | 2 | | | | | | | | | | | | | | | | | | | | | | | | | | | | 467 | 53.4 | 6.87 |
| P30273 | 12.79 | 1 | 1 | 1 | 2 | 12.79 | 1 | 1 | | | | | | | | | | | | | | | | | | | | | | | | | 12.79 | 1 | 1 | 86 | 9.7 | 7.12 |
| Q9UIK5 | 19.25 | 1 | 6 | 6 | 17 | | | | | | | 8.56 | 3 | 3 | | | | 3.21 | 1 | 1 | | | | | | | | | | | | | 19.25 | 6 | 13 | 374 | 41.4 | 5.15 |
| Q9UIY3 | 6.51 | 1 | 2 | 2 | 2 | | | | | | | | | | | | | | | | | | | | | | | | | | | | 6.51 | 2 | 2 | 292 | 33.9 | 6.47 |
| P05093 | 1.18 | 1 | 1 | 1 | 2 | | | | 0.00 | | | 0.00 | | | 0.00 | | | 0.00 | | | | | | | | | | | | 1.18 | 1 | 1 | 1.18 | 1 | 1 | 508 | 57.3 | 8.57 |
| Q9UJ68 | 31.91 | 1 | 5 | 5 | 7 | | | | 6.38 | 1 | 1 | | | | | | | | | | | | | | | | | | | | | | 31.91 | 5 | 6 | 235 | 26.1 | 8.09 |
| O14763 | 2.95 | 1 | 1 | 1 | 2 | | | | 2.95 | 1 | 2 | | | | | | | | | | | | | | | | | | | | | | | | | 440 | 47.8 | 5.55 |
| O14548 | 11.40 | 1 | 1 | 1 | 2 | | | | 11.40 | 1 | 1 | | | | | | | | | | | | | | | | | | | | | | 11.40 | 1 | 1 | 114 | 12.6 | 9.42 |
| Q9UJ70 | 57.27 | 1 | 17 | 17 | 55 | 9.59 | 2 | 2 | 12.50 | 3 | 3 | 9.59 | 2 | 2 | 3.78 | 1 | 1 | | | | | | | | | | | | | 2.03 | 1 | 1 | 57.27 | 17 | 46 | 344 | 37.4 | 6.24 |
| Q9UJ83 | 28.37 | 1 | 14 | 14 | 25 | 5.02 | 2 | 2 | 13.49 | 7 | 7 | 4.15 | 2 | 2 | | | | | | | | | | | | | | | | | | | 20.59 | 9 | 14 | 578 | 63.7 | 7.36 |
| Q9UJC3 | 39.56 | 1 | 26 | 26 | 63 | | | | | | | | | | 2.47 | 1 | 1 | 3.43 | 2 | 2 | | | | 14.29 | 9 | 9 | 3.85 | 2 | 2 | 14.70 | 8 | 8 | 33.38 | 20 | 41 | 728 | 84.6 | 5.15 |
| Q9UJC5 | 43.93 | 1 | 3 | 3 | 4 | | | | | | | | | | | | | | | | | | | | | | | | | | | | 43.93 | 3 | 4 | 107 | 12.3 | 6.71 |
| Q9UJS0 | 27.56 | 1 | 11 | 15 | 28 | 1.19 | 1 | 1 | 4.59 | 2 | 2 | | | | | | | | | | | | | | | | | | | | | | 25.63 | 14 | 25 | 675 | 74.1 | 8.62 |
| Q9UJT2 | 30.41 | 1 | 17 | 18 | 35 | | | | | | | | | | 2.36 | 1 | 1 | 2.36 | 1 | 1 | | | | 8.11 | 4 | 4 | 5.41 | 3 | 3 | 11.66 | 7 | 7 | 20.44 | 11 | 19 | 592 | 65.0 | 5.97 |
| A6H8Y1 | 0.34 | 1 | 1 | 1 | 2 | | | | 0.00 | | | | | | | | | | | | | | | | | | | | | | | | 0.34 | 1 | 2 | 2624 | 293.7 | 5.11 |
| O94927 | 1.90 | 1 | 1 | 1 | 2 | | | | 0.00 | | | | | | | | | | | | | | | | | | | | | | | | 1.90 | 1 | 2 | 633 | 71.6 | 8.51 |
| Q9UJU6 | 36.51 | 1 | 13 | 13 | 26 | 2.56 | 1 | 1 | | | | | | | 3.02 | 1 | 1 | | | | | | | 6.28 | 2 | 2 | 25.35 | 8 | 8 | 18.60 | 6 | 7 | 16.28 | 6 | 7 | 430 | 48.2 | 5.05 |
| Q9UJW0 | 11.09 | 1 | 4 | 4 | 9 | | | | 4.35 | 2 | 2 | | | | | | | | | | | | | | | | | | | | | | 11.09 | 4 | 7 | 460 | 52.3 | 7.34 |
| P01591 | 7.55 | 1 | 1 | 1 | 2 | | | | 7.55 | 1 | 1 | 7.55 | 1 | 1 | | | | | | | | | | | | | | | | | | | | | | 159 | 18.1 | 5.24 |
| Q9UJZ1 | 61.24 | 1 | 15 | 15 | 34 | 19.38 | 4 | 4 | 19.38 | 4 | 4 | | | | | | | | | | | | | | | | | | | | | | 61.24 | 15 | 26 | 356 | 38.5 | 7.39 |
| Q3ZCN5 | 0.30 | 1 | 1 | 1 | 2 | | | | 0.30 | 1 | 1 | | | | 0.30 | 1 | 1 | 0.00 | | | | | | | | | | | | | | | | | | 2332 | 261.9 | 5.16 |
| P51795 | 1.74 | 1 | 1 | 2 | 2 | | | | 0.80 | 1 | 1 | | | | | | | | | | | | | | | | | | | | | | 0.94 | 1 | 1 | 746 | 83.1 | 6.81 |
| Q9UK22 | 28.38 | 1 | 7 | 7 | 8 | | | | | | | | | | | | | | | | | | | | | | 4.05 | 1 | 1 | 5.07 | 1 | 1 | 23.31 | 5 | 6 | 296 | 33.3 | 4.37 |
| Q9UK41 | 31.67 | 1 | 4 | 4 | 7 | | | | | | | | | | | | | | | | | | | | | | | | | 2.71 | 1 | 1 | 31.67 | 4 | 6 | 221 | 25.4 | 5.54 |
| Q9UK85 | 8.68 | 1 | 2 | 2 | 2 | | | | 8.68 | 2 | 2 | | | | | | | | | | | | | | | | | | | | | | | | | 242 | 27.0 | 8.46 |
| Q9UKA8 | 29.05 | 2 | 5 | 5 | 13 | | | | | | | | | | | | | | | | | | | | | | | | | | | | 29.05 | 5 | 13 | 241 | 27.5 | 4.63 |
| Q6PDA7 | 7.32 | 1 | 1 | 1 | 2 | | | | 7.32 | 1 | 1 | | | | | | | 7.32 | 1 | 1 | | | | | | | | | | | | | | | | 123 | 13.8 | 10.54 |
| P06703 | 8.89 | 1 | 1 | 1 | 2 | | | | | | | 8.89 | 1 | 1 | 8.89 | 1 | 1 | | | | | | | | | | | | | | | | | | | 90 | 10.2 | 5.48 |
| P54725 | 4.96 | 1 | 1 | 2 | 2 | | | | | | | 2.20 | 1 | 1 | | | | | | | | | | | | | 2.75 | 1 | 1 | | | | | | | 363 | 39.6 | 4.58 |
| Q9UKF2 | 8.86 | 1 | 6 | 6 | 11 | 3.80 | 2 | 2 | 1.52 | 1 | 1 | | | | | | | | | | | | | | | | | | | | | | 8.86 | 5 | 8 | 790 | 88.9 | 7.61 |
| Q9UKF5 | 11.22 | 1 | 8 | 8 | 14 | | | | 4.39 | 3 | 3 | 1.83 | 1 | 1 | | | | | | | | | | | | | | | | | | | 7.93 | 6 | 10 | 820 | 92.7 | 7.40 |
| Q9UIG0 | 0.94 | 1 | 1 | 2 | 2 | | | | | | | 0.00 | | | | | | 0.00 | | | | | | | | | | | | | | | 0.94 | 2 | 2 | 1483 | 170.8 | 8.48 |
| Q9UKG1 | 2.96 | 1 | 2 | 2 | 3 | | | | | | | | | | | | | | | | | | | | | | | | | | | | 2.96 | 2 | 3 | 709 | 79.6 | 5.41 |
| P49006 | 6.67 | 1 | 1 | 1 | 2 | | | | | | | 6.67 | 1 | 1 | | | | | | | | | | | | | | | | | | | 6.67 | 1 | 1 | 195 | 19.5 | 4.67 |
| Q9UK38 | 2.91 | 1 | 2 | 2 | 3 | | | | | | | | | | | | | | | | | | | | | | | | | | | | 2.91 | 2 | 3 | 722 | 80.8 | 6.68 |
| O95260 | 2.12 | 1 | 1 | 1 | 2 | | | | | | | 2.12 | 1 | 1 | | | | 2.12 | 1 | 1 | | | | | | | | | | | | | | | | 518 | 59.1 | 7.93 |
| Q9UKK9 | 9.13 | 1 | 2 | 2 | 2 | | | | | | | | | | | | | | | | | | | | | | | | | | | | 9.13 | 2 | 2 | 219 | 24.3 | 4.94 |
| Q9UKM7 | 3.43 | 1 | 2 | 2 | 2 | | | | | | | | | | | | | | | | | | | | | | | | | | | | 3.43 | 2 | 2 | 699 | 79.5 | 7.72 |
| P14060 | 1.88 | 2 | 1 | 1 | 2 | | | | | | | 1.88 | 1 | 1 | 0.00 | | | | | | | | | | | | | | | | | | 1.88 | 1 | 1 | 373 | 42.2 | 8.70 |
| Q9UKR5 | 27.14 | 1 | 4 | 4 | 8 | | | | | | | 0.00 | | | | | | | | | | | | | | | | | | | | | 27.14 | 4 | 8 | 140 | 15.9 | 9.83 |
| Q14790 | 1.46 | 1 | 1 | 1 | 2 | | | | | | | 0.00 | | | | | | | | | | | | | | | | | | | | | 1.46 | 1 | 2 | 479 | 55.4 | 5.10 |
| Q99732 | 4.35 | 1 | 1 | 1 | 2 | | | | | | | 4.35 | 1 | 1 | | | | | | | | | | | | | | | | | | | 4.35 | 1 | 1 | 161 | 17.1 | 6.44 |
| Q9UKR8 | 21.22 | 1 | 6 | 6 | 36 | 15.51 | 2 | 2 | 8.57 | 2 | 2 | 15.92 | 3 | 3 | | | | | | | | | | | | | | | | | | | 20.00 | 4 | 29 | 245 | 26.2 | 8.27 |
| Q9UKU0 | 61.12 | 1 | 35 | 38 | 124 | 16.21 | 9 | 11 | 21.09 | 12 | 14 | 23.24 | 15 | 15 | 1.15 | 1 | 1 | 1.15 | 1 | 1 | | | | | | | | | | 1.29 | 2 | 2 | 59.25 | 34 | 80 | 697 | 77.7 | 7.43 |
| Q3V6T2 | 0.64 | 1 | 1 | 1 | 2 | | | | | | | | | | 0.00 | | | 0.00 | | | | | | | | | | | | | | | 0.64 | 1 | 2 | 1871 | 215.9 | 6.21 |
| Q8N3T1 | 1.25 | 1 | 1 | 1 | 2 | | | | | | | | | | 1.25 | 1 | 1 | | | | | | | | | | | | | | | | 1.25 | 1 | 1 | 639 | 73.0 | 6.90 |
| Q04323 | 4.04 | 1 | 1 | 1 | 2 | | | | | | | | | | 4.04 | 1 | 1 | | | | | | | | | | 4.04 | 1 | 1 | | | | | | | 297 | 33.3 | 5.25 |
| Q9NZ56 | 0.35 | 1 | 1 | 1 | 2 | | | | | | | | | | 0.35 | 1 | 1 | 0.00 | | | | | | 0.00 | | | | | | 0.00 | | | 0.35 | 1 | 1 | 1722 | 180.0 | 5.47 |
| Q96P48 | 0.48 | 1 | 1 | 1 | 2 | | | | | | | | | | 0.48 | 1 | 1 | | | | | | | 0.00 | | | | | | | | | 0.48 | 1 | 1 | 1450 | 162.1 | 6.23 |
| Q15773 | 3.63 | 1 | 1 | 1 | 2 | | | | | | | | | | 3.63 | 1 | 1 | | | | | | | | | | | | | 3.63 | 1 | 1 | | | | 248 | 28.1 | 6.90 |
| Q4ZJI4 | 3.88 | 4 | 1 | 2 | 2 | | | | | | | | | | 0.00 | | | | | | | | | | | | | | | | | | 3.88 | 2 | 2 | 515 | 56.0 | 8.02 |
| Q15643 | 0.71 | 3 | 1 | 2 | 2 | | | | | | | | | | | | | 0.00 | | | | | | | | | | | | | | | 0.71 | 2 | 2 | 1979 | 227.4 | 5.26 |
| Q9UBK8 | 0.97 | 1 | 1 | 1 | 2 | | | | | | | | | | | | | 0.97 | 1 | 1 | | | | 0.00 | | | 0.97 | 1 | 1 | | | | 0.00 | | | 725 | 80.4 | 6.49 |
| Q9UKU7 | 30.84 | 1 | 12 | 12 | 28 | 10.84 | 3 | 3 | 7.23 | 3 | 4 | 18.55 | 7 | 7 | | | | | | | | | | | | | | | | | | | 27.95 | 9 | 14 | 415 | 45.0 | 7.85 |
| Q86UL8 | 0.55 | 1 | 1 | 1 | 2 | | | | | | | | | | | | | 0.55 | 1 | 1 | | | | | | | 0.55 | 1 | 1 | | | | | | | 1455 | 158.7 | 6.35 |
| Q86TI0 | 0.60 | 1 | 1 | 1 | 2 | | | | | | | | | | | | | 0.60 | 1 | 1 | | | | | | | | | | | | | 0.60 | 1 | 1 | 1168 | 133.0 | 6.96 |
| Q9UIA0 | 2.79 | 1 | 1 | 1 | 2 | | | | | | | | | | | | | 2.79 | 1 | 1 | | | | | | | | | | | | | 2.79 | 1 | 1 | 394 | 45.6 | 6.47 |
| O95342 | 0.53 | 1 | 1 | 1 | 2 | | | | | | | | | | | | | 0.53 | 1 | 1 | | | | | | | | | | 0.53 | 1 | 1 | | | | 1321 | 146.3 | 6.57 |
| O14625 | 7.45 | 1 | 1 | 1 | 2 | | | | | | | | | | | | | 7.45 | 1 | 1 | | | | 7.45 | 1 | 1 | 0.00 | | | | | | | | | 94 | 10.4 | 9.89 |
| P28332 | 2.72 | 1 | 1 | 1 | 2 | | | | | | | | | | | | | 2.72 | 1 | 1 | | | | | | | | | | | | | 2.72 | 1 | 1 | 368 | 39.1 | 7.77 |
| Q9UL15 | 55.03 | 1 | 19 | 19 | 60 | 21.92 | 7 | 7 | 29.98 | 9 | 10 | 11.63 | 4 | 4 | 8.95 | 3 | 3 | 2.68 | 1 | 1 | | | | | | | 2.68 | 1 | 1 | 8.95 | 3 | 3 | 54.36 | 18 | 31 | 447 | 51.2 | 6.05 |
| Q9UL16 | 73.14 | 2 | 64 | 65 | 242 | 12.16 | 6 | 6 | 9.26 | 5 | 6 | 3.81 | 2 | 2 | 39.02 | 26 | 35 | 63.88 | 48 | 82 | | | | 3.81 | 2 | 2 | 3.81 | 2 | 2 | 3.81 | 2 | 2 | 59.17 | 39 | 105 | 551 | 65.7 | 8.90 |
| P35637 | 3.04 | 1 | 1 | 1 | 2 | | | | | | | | | | | | | | | | 3.04 | 1 | 2 | | | | | | | | | | | | | 526 | 53.4 | 9.36 |
| P17096 | 15.89 | 1 | 1 | 1 | 2 | | | | | | | | | | | | | | | | 15.89 | 1 | 2 | | | | | | | | | | | | | 107 | 11.7 | 10.32 |
| Q9UL25 | 24.00 | 1 | 4 | 4 | 6 | 4.89 | 1 | 1 | 6.67 | 1 | 1 | | | | | | | | | | | | | | | | | | | | | | 19.11 | 3 | 4 | 225 | 24.3 | 7.94 |
| Q15714 | 1.96 | 2 | 1 | 2 | 2 | | | | | | | | | | | | | | | | | | | 0.84 | 1 | 1 | | | | 1.12 | 1 | 1 | | | | 1073 | 109.6 | 5.64 |
| Q3MHD2 | 6.67 | 1 | 1 | 1 | 2 | | | | | | | | | | | | | | | | | | | 6.67 | 1 | 1 | 6.67 | 1 | 1 | | | | | | | 195 | 21.7 | 7.74 |
| P19387 | 3.27 | 1 | 1 | 1 | 2 | | | | | | | | | | | | | | | | | | | 3.27 | 1 | 1 | | | | 3.27 | 1 | 1 | | | | 275 | 31.4 | 4.92 |
| Q99576 | 13.43 | 2 | 1 | 2 | 2 | | | | | | | | | | | | | | | | | | | 6.72 | 1 | 1 | | | | 6.72 | 1 | 1 | | | | 134 | 14.8 | 4.54 |
| Q9UL26 | 24.74 | 2 | 3 | 3 | 3 | | | | | | | | | | | | | | | | | | | | | | | | | | | | 24.74 | 3 | 3 | 194 | 21.8 | 8.15 |
| Q14011 | 6.40 | 1 | 1 | 1 | 2 | | | | | | | | | | | | | | | | | | | | | | 6.40 | 1 | 1 | 6.40 | 1 | 1 | | | | 172 | 18.6 | 9.51 |
| O96033 | 21.59 | 1 | 1 | 1 | 2 | | | | | | | | | | | | | | | | | | | | | | 21.59 | 1 | 1 | 21.59 | 1 | 1 | | | | 88 | 9.7 | 4.72 |
| Q15517 | 3.40 | 2 | 1 | 1 | 2 | | | | | | | | | | | | | | | | | | | | | | 3.40 | 1 | 2 | | | | | | | 529 | 51.5 | 8.35 |
| Q9UL33 | 25.71 | 1 | 3 | 3 | 3 | | | | | | | | | | | | | | | | | | | | | | | | | | | | 25.71 | 3 | 3 | 140 | 16.1 | 6.77 |
| Q9UL45 | 24.42 | 1 | 2 | 2 | 3 | | | | | | | | | | | | | | | | | | | | | | | | | | | | 24.42 | 2 | 3 | 172 | 19.7 | 6.40 |
| B7ZBB8 | 3.35 | 1 | 1 | 1 | 2 | | | | | | | | | | | | | | | | | | | | | | 3.35 | 1 | 2 | | | | | | | 358 | 38.0 | 4.93 |
| Q9UL46 | 49.79 | 1 | 10 | 11 | 22 | 13.39 | 2 | 2 | | | | 17.99 | 3 | 3 | | | | | | | | | | | | | | | | | | | 47.28 | 10 | 17 | 239 | 27.4 | 5.73 |
| Q9ULA0 | 61.05 | 1 | 20 | 20 | 63 | 20.84 | 7 | 7 | 14.11 | 5 | 5 | 12.84 | 5 | 5 | | | | | | | | | | | | | | | | | | | 57.89 | 19 | 46 | 475 | 52.4 | 7.42 |
| Q9ULC3 | 30.38 | 1 | 5 | 5 | 9 | | | | | | | | | | | | | | | | | | | | | | | | | 4.22 | 1 | 1 | 30.38 | 5 | 8 | 237 | 26.6 | 6.60 |
| Q9ULC4 | 39.23 | 1 | 6 | 6 | 12 | | | | | | | | | | 4.42 | 1 | 1 | | | | | | | | | | | | | | | | 39.23 | 6 | 11 | 181 | 20.5 | 8.82 |
| Q9ULE4 | 1.32 | 1 | 2 | 2 | 2 | | | | | | | | | | | | | | | | | | | | | | | | | | | | 1.32 | 2 | 2 | 1060 | 121.0 | 6.18 |
| Q9ULG3 | 10.02 | 1 | 3 | 3 | 3 | | | | | | | | | | | | | | | | | | | | | | | | | | | | 10.02 | 3 | 3 | 409 | 46.7 | 8.06 |
| Q9ULP9 | 9.30 | 1 | 4 | 4 | 8 | | | | | | | | | | | | | | | | | | | | | | | | | | | | 9.30 | 4 | 8 | 559 | 62.9 | 7.36 |
| Q9ULR3 | 4.86 | 1 | 2 | 2 | 2 | | | | | | | | | | | | | | | | | | | | | | | | | | | | 4.86 | 2 | 2 | 514 | 56.4 | 6.60 |
| P31327 | 1.33 | 1 | 1 | 2 | 2 | | | | | | | | | | | | | | | | | | | | | | | | | | | | 1.33 | 2 | 2 | 1500 | 164.8 | 6.74 |
| Q9ULV0 | 8.60 | 1 | 8 | 13 | 29 | | | | | | | | | | | | | | | | | | | 0.76 | 1 | 1 | | | | 0.76 | 1 | 1 | 8.60 | 13 | 27 | 1848 | 213.5 | 7.20 |
| Q5T8D3 | 1.87 | 1 | 1 | 1 | 2 | | | | | | | | | | | | | | | | | | | | | | | | | | | | 1.87 | 1 | 2 | 534 | 60.1 | 5.33 |
| Q7Z333 | 0.45 | 2 | 1 | 2 | 2 | | | | | | | | | | | | | | | | | | | | | | | | | | | | 0.45 | 2 | 2 | 2677 | 302.7 | 7.17 |
| Q9H672 | 4.40 | 1 | 1 | 2 | 2 | | | | | | | | | | | | | | | | | | | | | | | | | | | | 4.40 | 2 | 2 | 318 | 36.0 | 8.29 |
| Q9ULV4 | 10.34 | 1 | 4 | 4 | 5 | | | | | | | 6.75 | 2 | 2 | | | | | | | | | | | | | | | | | | | 6.12 | 3 | 3 | 474 | 53.2 | 7.08 |
| Q9H074 | 2.51 | 1 | 1 | 1 | 2 | | | | | | | | | | | | | | | | | | | | | | | | | | | | 2.51 | 1 | 2 | 479 | 53.5 | 4.81 |
| Q9UM00 | 17.02 | 1 | 3 | 3 | 4 | | | | 9.04 | 2 | 2 | | | | | | | | | | | | | | | | | | | | | | 12.23 | 2 | 2 | 188 | 21.2 | 9.74 |
| Q9UM22 | 8.93 | 1 | 2 | 2 | 3 | | | | | | | | | | | | | | | | | | | | | | | | | | | | 8.93 | 2 | 3 | 224 | 25.4 | 6.60 |
| Q9BPX3 | 0.59 | 1 | 1 | 1 | 2 | | | | | | | | | | | | | | | | | | | | | | | | | | | | 0.59 | 1 | 2 | 1015 | 114.3 | 5.59 |
| Q9UM54 | 44.20 | 1 | 52 | 52 | 149 | 4.79 | 5 | 5 | 2.70 | 3 | 3 | 3.63 | 4 | 4 | 0.85 | 1 | 1 | | | | | | | 3.55 | 4 | 4 | | | | 4.87 | 4 | 4 | 41.81 | 48 | 128 | 1294 | 149.6 | 8.53 |
| Q49A26 | 1.27 | 1 | 1 | 1 | 2 | | | | | | | | | | | | | | | | | | | | | | | | | | | | 1.27 | 1 | 2 | 553 | 60.5 | 9.17 |
| Q9UNK0 | 7.63 | 1 | 1 | 1 | 2 | | | | | | | | | | | | | | | | | | | | | | | | | | | | 7.63 | 1 | 2 | 236 | 26.9 | 4.98 |

| | | | | | | | | | | | | | | | | | | | | | | | | | | | | | | | | | | | | | | |
|---|---|---|---|---|---|---|---|---|---|---|---|---|---|---|---|---|---|---|---|---|---|---|---|---|---|---|---|---|---|---|---|---|---|---|---|---|---|---|
| Q8NEA5 | 5.58 | 1 | 1 | 1 | 2 | | | | | | | | | | | | | | | | | | | | | | | | | | | | 5.58 | 1 | 2 | 215 | 24.1 | 8.95 |
| Q9UMQ6 | 11.77 | 1 | 6 | 6 | 8 | | | | | | | | | | | | | | | | | | | | | | | | | | | | 11.77 | 6 | 8 | 739 | 84.4 | 5.60 |
| Q9UMX2 | 25.53 | 1 | 4 | 4 | 13 | | | | | | | | | | 6.38 | 1 | 1 | 6.38 | 1 | 1 | | | | 6.38 | 1 | 2 | 6.38 | 1 | 1 | | | | 25.53 | 4 | 8 | 235 | 27.4 | 6.86 |
| P36952 | 4.53 | 1 | 1 | 1 | 2 | | | | | | | | | | | | | | | | | | | | | | | | | | | | 4.53 | 1 | 2 | 375 | 42.1 | 6.05 |
| Q9UMY4 | 23.26 | 1 | 3 | 4 | 5 | | | | | | | | | | | | | | | | | | | | | | | | | 11.05 | 2 | 2 | 12.21 | 2 | 3 | 172 | 19.7 | 7.87 |
| O00592 | 2.33 | 1 | 1 | 1 | 2 | | | | | | | | | | | | | | | | | | | | | | | | | | | | 2.33 | 1 | 2 | 558 | 58.6 | 5.49 |
| Q9UN30 | 10.94 | 1 | 2 | 2 | 2 | | | | | | | | | | | | | | | | | | | | | | | | | | | | 10.94 | 2 | 2 | 329 | 37.4 | 6.89 |
| Q96D46 | 3.18 | 1 | 1 | 1 | 2 | | | | | | | | | | | | | | | | | | | | | | | | | | | | 3.18 | 1 | 2 | 503 | 57.6 | 7.14 |
| Q9UN36 | 45.82 | 1 | 9 | 10 | 19 | | | | | | | | | | | | | | | | | | | | | | | | | | | | 45.82 | 10 | 19 | 371 | 40.8 | 5.21 |
| O00567 | 1.52 | 1 | 1 | 1 | 2 | | | | | | | | | | | | | | | | | | | | | | | | | | | | 1.52 | 1 | 2 | 594 | 66.0 | 9.19 |
| Q9UN37 | 18.54 | 2 | 5 | 7 | 9 | | | | | | | | | | | | | | | | | | | | | | | | | | | | 18.54 | 7 | 9 | 437 | 48.9 | 7.80 |
| Q9UNE7 | 20.46 | 1 | 7 | 7 | 8 | | | | | | | | | | | | | | | | | | | | | | | | | | | | 20.46 | 7 | 8 | 303 | 34.8 | 5.87 |
| Q9UNF0 | 4.12 | 1 | 2 | 2 | 2 | | | | | | | | | | | | | | | | | | | | | | | | | 2.26 | 1 | 1 | 1.85 | 1 | 1 | 486 | 55.7 | 5.20 |
| Q9P287 | 7.32 | 1 | 1 | 1 | 2 | | | | | | | | | | | | | | | | | | | | | | | | | | | | 7.32 | 1 | 2 | 314 | 36.0 | 4.61 |
| P61165 | 10.13 | 1 | 1 | 1 | 2 | | | | | | | | | | | | | | | | | | | | | | | | | | | | 10.13 | 1 | 2 | 79 | 9.1 | 5.83 |
| Q8WU68 | 5.45 | 2 | 1 | 1 | 2 | | | | | | | | | | | | | | | | | | | | | | | | | | | | 5.45 | 1 | 2 | 220 | 25.7 | 7.15 |
| Q9UNF1 | 4.46 | 2 | 3 | 3 | 4 | | | | | | | | | | | | | | | | | | | | | | | | | | | | 4.46 | 3 | 4 | 606 | 64.9 | 9.32 |
| Q9UNH7 | 31.28 | 2 | 11 | 12 | 21 | | | | | | | | | | | | | | | | | | | 4.43 | 2 | 2 | | | | 14.29 | 6 | 6 | 24.63 | 9 | 13 | 406 | 46.6 | 6.16 |
| P35354 | 1.66 | 1 | 1 | 1 | 2 | | | | | | | | | | | | | | | | | | | | | | | | | | | | 1.66 | 1 | 2 | 604 | 69.0 | 7.39 |
| Q9UNM6 | 68.62 | 1 | 25 | 25 | 85 | 25.53 | 7 | 7 | 16.76 | 6 | 6 | 16.49 | 6 | 6 | 29.26 | 11 | 12 | 2.66 | 1 | 1 | | | | | | | | | | 8.24 | 3 | 3 | 61.70 | 21 | 50 | 376 | 42.9 | 5.81 |
| Q9UNN5 | 17.69 | 1 | 11 | 11 | 18 | | | | 5.23 | 3 | 3 | 1.54 | 1 | 1 | | | | | | | | | | 1.69 | 1 | 1 | 1.85 | 1 | 1 | | | | 14.00 | 8 | 12 | 650 | 73.9 | 4.88 |
| Q5TDH0 | 2.26 | 1 | 1 | 1 | 2 | | | | | | | | | | | | | | | | | | | | | | | | | | | | 2.26 | 1 | 2 | 399 | 44.5 | 5.05 |
| Q9P265 | 1.08 | 1 | 1 | 2 | 2 | | | | | | | | | | | | | | | | | | | | | | | | | | | | 1.08 | 2 | 2 | 1576 | 171.4 | 8.09 |
| Q9UNQ0 | 7.48 | 1 | 5 | 5 | 9 | | | | | | | | | | | | | | | | | | | | | | | | | | | | 7.48 | 5 | 9 | 655 | 72.3 | 8.69 |
| Q9UNS2 | 48.23 | 1 | 15 | 15 | 38 | 6.86 | 2 | 2 | 4.02 | 1 | 1 | | | | 13.95 | 4 | 4 | | | | 4.02 | 1 | 1 | | | | | | | 16.78 | 5 | 6 | 44.44 | 14 | 24 | 423 | 47.8 | 6.65 |
| Q9BW60 | 4.66 | 1 | 1 | 1 | 2 | | | | | | | | | | | | | | | | | | | | | | | | | | | | 4.66 | 1 | 2 | 279 | 32.6 | 9.60 |
| P50402 | 3.94 | 1 | 1 | 1 | 2 | | | | | | | | | | | | | | | | | | | | | | | | | | | | 3.94 | 1 | 2 | 254 | 29.0 | 5.50 |
| P62314 | 10.92 | 1 | 1 | 1 | 2 | | | | | | | | | | | | | | | | | | | | | | | | | | | | 10.92 | 1 | 2 | 119 | 13.3 | 11.56 |
| Q15436 | 3.53 | 1 | 1 | 2 | 2 | | | | | | | | | | | | | | | | | | | | | | | | | | | | 3.53 | 2 | 2 | 765 | 86.1 | 7.08 |
| Q9UNW1 | 6.37 | 1 | 2 | 2 | 2 | | | | | | | | | | | | | | | | | | | | | | | | | | | | 6.37 | 2 | 2 | 487 | 55.0 | 7.81 |
| Q9UNZ2 | 40.00 | 1 | 12 | 12 | 33 | 17.03 | 5 | 5 | 12.97 | 4 | 4 | 7.03 | 2 | 2 | 7.84 | 2 | 2 | 2.70 | 1 | 1 | | | | 12.16 | 4 | 4 | 16.22 | 4 | 4 | 12.97 | 4 | 4 | 16.49 | 5 | 7 | 370 | 40.5 | 5.10 |
| Q6IN85 | 0.84 | 1 | 1 | 1 | 2 | | | | | | | | | | | | | | | | | | | | | | | | | | | | 0.84 | 1 | 2 | 833 | 95.3 | 4.94 |
| Q9UP83 | 12.40 | 1 | 8 | 8 | 22 | | | | | | | | | | | | | | | | | | | | | | | | | | | | 12.40 | 8 | 22 | 839 | 92.7 | 6.60 |
| Q9UPM8 | 3.61 | 1 | 3 | 3 | 5 | | | | | | | | | | | | | | | | | | | | | | | | | | | | 3.61 | 3 | 5 | 1137 | 127.2 | 5.99 |
| O95453 | 1.10 | 1 | 1 | 1 | 2 | | | | | | | | | | | | | | | | | | | | | | | | | | | | 1.10 | 1 | 2 | 639 | 73.4 | 6.20 |
| Q8N5M9 | 6.56 | 1 | 1 | 1 | 2 | | | | | | | | | | | | | | | | | | | | | | | | | | | | 6.56 | 1 | 2 | 183 | 21.1 | 9.73 |
| Q9UPM9 | 13.24 | 1 | 2 | 2 | 2 | | | | | | | | | | | | | | | | | | | | | | | | | | | | 13.24 | 2 | 2 | 204 | 22.8 | 7.28 |
| Q9UPN3 | 0.81 | 1 | 6 | 6 | 11 | 0.00 | | | | | | | | | | | | | | | | | | | | | | | | | | | 0.81 | 6 | 11 | 7388 | 837.8 | 5.39 |
| Q99735 | 9.52 | 1 | 1 | 1 | 2 | | | | | | | | | | | | | | | | | | | | | | | | | | | | 9.52 | 1 | 2 | 147 | 16.6 | 9.55 |
| Q9UPN7 | 15.21 | 1 | 10 | 10 | 20 | 2.04 | 1 | 1 | 2.04 | 1 | 1 | | | | 1.25 | 1 | 1 | 1.25 | 1 | 1 | | | | | | | | | | | | | 13.17 | 9 | 16 | 881 | 96.7 | 4.55 |
| Q9UPQ0 | 3.14 | 1 | 3 | 3 | 10 | | | | | | | | | | | | | | | | | | | | | | | | | 2.40 | 2 | 2 | 0.74 | 1 | 8 | 1083 | 121.8 | 6.47 |
| Q9UPT5 | 10.20 | 1 | 5 | 5 | 7 | | | | | | | | | | | | | | | | | | | | | | | | | | | | 10.20 | 5 | 7 | 735 | 83.3 | 6.79 |
| Q9UPU5 | 2.37 | 1 | 4 | 4 | 8 | | | | | | | | | | | | | | | | | | | | | | | | | | | | 2.37 | 4 | 8 | 2620 | 294.2 | 6.14 |
| P48651 | 2.96 | 1 | 1 | 1 | 2 | | | | | | | | | | | | | | | | | | | | | | | | | | | | 2.96 | 1 | 2 | 473 | 55.5 | 8.43 |
| Q6AI08 | 1.27 | 1 | 1 | 1 | 2 | | | | | | | | | | | | | | | | | | | | | | | | | | | | 1.27 | 1 | 2 | 1181 | 128.7 | 7.03 |
| Q9UQ49 | 9.35 | 1 | 3 | 3 | 5 | | | | | | | | | | | | | | | | | | | | | | | | | | | | 9.35 | 3 | 5 | 428 | 48.2 | 7.17 |
| A6NDU8 | 4.76 | 1 | 1 | 1 | 2 | | | | | | | | | | | | | | | | | | | | | | | | | | | | 4.76 | 1 | 2 | 294 | 33.6 | 5.26 |
| Q9UQ80 | 46.70 | 1 | 17 | 17 | 70 | 15.74 | 6 | 6 | 15.48 | 6 | 7 | 4.82 | 2 | 2 | 9.14 | 3 | 3 | 11.17 | 4 | 4 | | | | 16.24 | 6 | 6 | 13.20 | 5 | 5 | 14.47 | 5 | 7 | 37.06 | 13 | 30 | 394 | 43.8 | 6.55 |
| Q9UNL2 | 7.57 | 1 | 1 | 1 | 2 | | | | | | | | | | | | | | | | | | | | | | | | | | | | 7.57 | 1 | 2 | 185 | 21.1 | 9.61 |
| Q9UQB8 | 9.96 | 1 | 5 | 5 | 7 | | | | | | | | | | | | | | | | | | | | | | | | | | | | 9.96 | 5 | 7 | 552 | 60.8 | 8.90 |
| Q96S52 | 3.78 | 1 | 1 | 1 | 2 | | | | | | | | | | | | | | | | | | | | | | | | | | | | 3.78 | 1 | 2 | 555 | 61.6 | 6.49 |
| Q9UQE7 | 14.54 | 3 | 14 | 15 | 17 | | | | | | | | | | | | | | | | | | | | | | | | | | | | 14.54 | 15 | 17 | 1217 | 141.5 | 7.18 |
| Q9UQN3 | 12.68 | 1 | 3 | 3 | 7 | | | | | | | | | | 4.69 | 1 | 1 | 4.69 | 1 | 1 | | | | 4.69 | 1 | 1 | | | | 4.69 | 1 | 1 | 12.68 | 3 | 3 | 213 | 23.9 | 8.76 |
| Q9Y217 | 3.54 | 3 | 2 | 2 | 2 | | | | | | | | | | | | | | | | | | | | | | | | | | | | 3.54 | 2 | 2 | 621 | 71.9 | 7.66 |
| Q9Y223 | 17.73 | 1 | 10 | 10 | 20 | | | | | | | | | | | | | | | | | | | | | | | | | | | | 17.73 | 10 | 20 | 722 | 79.2 | 6.80 |
| Q9Y224 | 43.03 | 1 | 9 | 9 | 10 | | | | | | | | | | | | | | | | | | | | | | | | | 7.38 | 2 | 2 | 39.34 | 8 | 8 | 244 | 28.1 | 6.65 |
| Q9Y230 | 73.87 | 1 | 40 | 40 | 260 | 34.13 | 14 | 16 | 50.54 | 21 | 26 | 49.46 | 19 | 21 | 43.84 | 18 | 20 | 44.71 | 18 | 22 | | | | 27.86 | 11 | 12 | 34.99 | 15 | 18 | 32.83 | 12 | 14 | 67.17 | 30 | 111 | 463 | 51.1 | 5.64 |
| Q13158 | 12.50 | 1 | 1 | 1 | 2 | | | | | | | | | | | | | | | | | | | | | | | | | | | | 12.50 | 1 | 2 | 208 | 23.3 | 5.69 |
| Q9Y233 | 5.91 | 1 | 3 | 3 | 5 | | | | | | | | | | | | | | | | | | | | | | | | | | | | 5.91 | 3 | 5 | 779 | 88.4 | 6.60 |
| Q9Y3L5 | 10.93 | 3 | 1 | 2 | 2 | | | | | | | | | | | | | | | | | | | | | | | | | | | | 10.93 | 2 | 2 | 183 | 20.7 | 4.94 |
| Q2TAC2 | 0.66 | 2 | 1 | 1 | 2 | | | | | | | | | | | | | | | | | | | | | | | | | | | | 0.66 | 1 | 2 | 916 | 103.1 | 6.55 |
| O95436 | 1.59 | 1 | 1 | 1 | 2 | | | | | | | | | | | | | | | | | | | | | | | | | | | | 1.59 | 1 | 2 | 690 | 75.7 | 8.15 |
| O15111 | 2.15 | 1 | 1 | 1 | 2 | | | | | | | | | | | | | | | | | | | | | | | | | | | | 2.15 | 1 | 2 | 745 | 84.6 | 6.73 |
| Q99653 | 6.15 | 1 | 1 | 1 | 2 | | | | | | | | | | | | | | | | | | | | | | | | | | | | 6.15 | 1 | 2 | 195 | 22.4 | 5.10 |
| Q9Y238 | 11.34 | 1 | 16 | 16 | 26 | | | | | | | | | | | | | 0.74 | 1 | 1 | | | | | | | | | | | | | 11.34 | 16 | 25 | 1755 | 195.6 | 6.33 |
| Q9NZN4 | 3.50 | 1 | 1 | 2 | 2 | | | | | | | | | | | | | | | | | | | | | | | | | | | | 3.50 | 2 | 2 | 543 | 61.1 | 6.46 |
| Q9Y241 | 29.03 | 2 | 2 | 2 | 4 | | | | | | | | | | | | | | | | | | | | | | | | | | | | 29.03 | 2 | 4 | 93 | 10.1 | 9.79 |
| Q969L2 | 6.25 | 1 | 1 | 1 | 2 | | | | | | | | | | | | | | | | | | | | | | | | | | | | 6.25 | 1 | 2 | 176 | 19.1 | 6.24 |
| Q9Y259 | 11.14 | 1 | 3 | 3 | 3 | | | | | | | | | | | | | | | | | | | | | | | | | | | | 11.14 | 3 | 3 | 395 | 45.2 | 5.49 |
| Q9Y262 | 28.90 | 1 | 14 | 14 | 31 | | | | 1.77 | 1 | 1 | 1.60 | 1 | 1 | 1.60 | 1 | 1 | 1.60 | 1 | 1 | | | | | | | | | | | | | 27.13 | 13 | 27 | 564 | 66.7 | 6.34 |
| Q99962 | 5.40 | 2 | 1 | 2 | 2 | | | | | | | | | | | | | | | | | | | | | | | | | | | | 5.40 | 2 | 2 | 352 | 39.9 | 5.45 |
| Q9Y263 | 15.97 | 1 | 10 | 10 | 24 | | | | 2.14 | 1 | 1 | 2.14 | 1 | 1 | | | | | | | | | | | | | | | | 2.01 | 1 | 1 | 14.97 | 9 | 21 | 795 | 87.1 | 6.37 |
| Q96BW9 | 2.88 | 1 | 1 | 1 | 2 | | | | | | | | | | | | | | | | | | | | | | | | | | | | 2.88 | 1 | 2 | 452 | 51.0 | 7.94 |
| Q9Y265 | 70.61 | 1 | 28 | 28 | 142 | 39.04 | 11 | 12 | 44.30 | 15 | 19 | 39.25 | 14 | 17 | 24.34 | 10 | 11 | 33.55 | 13 | 15 | 5.70 | 2 | 2 | 7.24 | 3 | 3 | 17.32 | 6 | 7 | 7.68 | 2 | 2 | 56.36 | 21 | 54 | 456 | 50.2 | 6.42 |
| Q15800 | 4.78 | 1 | 1 | 1 | 2 | | | | | | | | | | | | | | | | | | | | | | | | | | | | 4.78 | 1 | 2 | 293 | 35.2 | 7.23 |
| Q9Y266 | 46.83 | 1 | 16 | 16 | 39 | 9.37 | 3 | 3 | 13.90 | 4 | 5 | 3.32 | 1 | 1 | | | | 7.85 | 2 | 2 | | | | 11.48 | 3 | 3 | 13.29 | 4 | 4 | 16.01 | 4 | 4 | 43.81 | 14 | 17 | 331 | 38.2 | 5.38 |
| Q9Y276 | 6.92 | 1 | 2 | 2 | 2 | | | | | | | | | | | | | | | | | | | | | | | | | | | | 6.92 | 2 | 2 | 419 | 47.5 | 8.50 |
| Q5JS54 | 20.33 | 1 | 1 | 1 | 2 | | | | | | | | | | | | | | | | | | | | | | | | | | | | 20.33 | 1 | 2 | 123 | 13.8 | 6.52 |
| Q7Z404 | 1.40 | 1 | 1 | 1 | 2 | | | | | | | | | | | | | | | | | | | | | | | | | | | | 1.40 | 1 | 2 | 712 | 79.2 | 8.94 |
| Q63HN1 | 2.52 | 2 | 1 | 1 | 2 | | | | | | | | | | | | | | | | | | | | | | | | | | | | 2.52 | 1 | 2 | 556 | 61.8 | 7.50 |
| Q9Y277 | 62.90 | 1 | 15 | 16 | 144 | 36.75 | 9 | 12 | 39.93 | 9 | 13 | 54.77 | 12 | 19 | 19.79 | 5 | 5 | 40.28 | 9 | 10 | | | | | | | | | | | | | 62.90 | 16 | 85 | 283 | 30.6 | 8.66 |
| P09914 | 2.51 | 1 | 1 | 1 | 2 | | | | | | | | | | | | | | | | | | | | | | | | | | | | 2.51 | 1 | 2 | 478 | 55.3 | 7.20 |
| Q9Y281 | 50.00 | 1 | 4 | 10 | 34 | 16.27 | 2 | 4 | 6.63 | 1 | 1 | 16.87 | 2 | 3 | 6.63 | 1 | 1 | 6.63 | 1 | 1 | | | | 26.51 | 4 | 4 | 15.66 | 3 | 3 | 10.24 | 2 | 2 | 40.36 | 5 | 15 | 166 | 18.7 | 7.88 |
| Q9Y282 | 30.03 | 1 | 13 | 13 | 30 | | | | 6.53 | 2 | 2 | 1.83 | 1 | 1 | | | | | | | | | | | | | | | | | | | 30.03 | 13 | 27 | 383 | 43.2 | 6.06 |
| Q9Y285 | 29.53 | 1 | 12 | 12 | 31 | 10.24 | 4 | 4 | 6.30 | 3 | 3 | | | | | | | | | | | | | | | | | | | 9.84 | 2 | 2 | 25.59 | 10 | 22 | 508 | 57.5 | 7.80 |
| Q9Y287 | 38.35 | 1 | 5 | 5 | 13 | 3.76 | 1 | 1 | 3.76 | 1 | 1 | | | | 8.65 | 2 | 2 | | | | | | | | | | | | | | | | 38.35 | 5 | 9 | 266 | 30.3 | 5.14 |
| P60520 | 15.38 | 2 | 1 | 2 | 2 | | | | | | | | | | | | | | | | | | | | | | | | | | | | 15.38 | 2 | 2 | 117 | 13.7 | 8.10 |
| Q86TZ1 | 2.12 | 1 | 1 | 1 | 2 | | | | | | | | | | | | | | | | | | | | | | | | | | | | 2.12 | 1 | 2 | 520 | 59.3 | 6.37 |
| O95271 | 0.60 | 1 | 1 | 1 | 2 | | | | | | | | | | | | | | | | | | | | | | | | | | | | 0.60 | 1 | 2 | 1327 | 142.0 | 7.05 |
| Q5W0A0 | 1.01 | 1 | 1 | 1 | 2 | | | | | | | | | | | | | | | | | | | | | | | | | | | | 1.01 | 1 | 2 | 696 | 81.6 | 4.75 |
| Q9Y295 | 9.81 | 1 | 3 | 3 | 3 | | | | | | | | | | | | | | | | | | | | | | | | | | | | 9.81 | 3 | 3 | 367 | 40.5 | 8.90 |
| Q9Y2A7 | 21.99 | 1 | 20 | 20 | 29 | | | | | | | | | | | | | | | | | | | | | | | | | | | | 21.99 | 20 | 29 | 1128 | 128.7 | 6.62 |
| Q96JH7 | 1.31 | 1 | 1 | 1 | 2 | | | | | | | | | | | | | | | | | | | | | | | | | | | | 1.31 | 1 | 2 | 1222 | 134.2 | 7.20 |
| Q9Y2B0 | 31.32 | 1 | 4 | 4 | 7 | 8.79 | 1 | 1 | 8.79 | 1 | 1 | 8.79 | 1 | 1 | | | | | | | | | | | | | | | | | | | 31.32 | 4 | 4 | 182 | 20.6 | 4.92 |
| Q96HJ9 | 10.62 | 1 | 1 | 1 | 2 | | | | | | | | | | | | | | | | | | | | | | | | | | | | 10.62 | 1 | 2 | 113 | 12.7 | 10.07 |
| Q9Y2B4 | 6.21 | 1 | 2 | 2 | 4 | | | | | | | | | | | | | | | | | | | | | | | | | | | | 6.21 | 2 | 4 | 290 | 34.0 | 10.55 |
| Q99707 | 1.03 | 1 | 1 | 1 | 2 | | | | | | | | | | | | | | | | | | | | | | | | | | | | 1.03 | 1 | 2 | 1265 | 140.4 | 5.58 |
| P36639 | 6.09 | 1 | 1 | 1 | 2 | | | | | | | | | | | | | | | | | | | | | | | | | | | | 6.09 | 1 | 2 | 197 | 22.5 | 5.27 |
| Q3B820 | 2.42 | 1 | 1 | 1 | 2 | | | | | | | | | | | | | | | | | | | | | | | | | | | | 2.42 | 1 | 2 | 660 | 76.7 | 8.03 |
| Q9Y2C4 | 14.40 | 1 | 3 | 3 | 3 | | | | | | | | | | | | | | | | | | | | | | | | | | | | 14.40 | 3 | 3 | 368 | 41.1 | 8.27 |
| Q9BQS8 | 1.15 | 5 | 1 | 2 | 2 | | | | | | | | | | | | | | | | | | | | | | | | | | | | 1.15 | 2 | 2 | 1478 | 166.9 | 4.92 |
| Q86VN1 | 2.07 | 1 | 1 | 1 | 2 | | | | | | | | | | | | | | | | | | | | | | | | | | | | 2.07 | 1 | 2 | 386 | 43.8 | 7.20 |
| Q9Y2I8 | 6.48 | 1 | 2 | 3 | 7 | | | | | | | | | | | | | | | | | | | 3.04 | 1 | 2 | | | | 3.04 | 1 | 1 | 6.48 | 3 | 4 | 494 | 54.6 | 7.23 |
| Q9BPW8 | 5.28 | 1 | 1 | 2 | 2 | | | | | | | | | | | | | | | | | | | | | | | | | | | | 5.28 | 2 | 2 | 284 | 33.3 | 9.31 |
| Q9Y2J2 | 2.94 | 1 | 2 | 2 | 2 | | | | | | | | | | | | | | | | | | | | | | 2.94 | 2 | 2 | | | | | | | 1087 | 120.6 | 5.19 |
| Q9Y2L5 | 2.23 | 1 | 3 | 3 | 4 | | | | | | | | | | | | | | | | | | | | | | | | | | | | 2.23 | 3 | 4 | 1435 | 160.9 | 6.87 |
| Q9Y2Q0 | 16.67 | 3 | 16 | 18 | 32 | | | | 1.03 | 1 | 1 | 3.09 | 3 | 3 | | | | | | | | | | | | | | | | | | | 14.86 | 16 | 28 | 1164 | 131.3 | 6.84 |
| Q9Y2Q3 | 17.26 | 1 | 3 | 3 | 7 | 10.18 | 2 | 2 | | | | | | | | | | | | | | | | | | | | | | 6.19 | 1 | 1 | 13.27 | 2 | 4 | 226 | 25.5 | 8.41 |
| Q9Y2Q5 | 31.20 | 1 | 3 | 3 | 5 | | | | | | | | | | | | | | | | | | | | | | | | | | | | 31.20 | 3 | 5 | 125 | 13.5 | 5.40 |
| Q9Y2R0 | 17.92 | 1 | 2 | 2 | 2 | | | | | | | | | | | | | | | | | | | | | | | | | | | | 17.92 | 2 | 2 | 106 | 11.7 | 9.60 |
| Q8IZD9 | 0.44 | 1 | 1 | 1 | 2 | | | | | | | | | | | | | | | | | | | | | | | | | | | | 0.44 | 1 | 2 | 2030 | 233.0 | 6.98 |
| Q5JV73 | 0.72 | 5 | 1 | 2 | 2 | | | | | | | | | | | | | | | | | | | | | | | | | | | | 0.72 | 2 | 2 | 1810 | 199.1 | 8.15 |
| Q9Y2S2 | 47.34 | 1 | 13 | 13 | 35 | | | | | | | | | | 6.27 | 2 | 2 | | | | | | | 6.27 | 2 | 2 | | | | 13.48 | 4 | 4 | 47.34 | 13 | 27 | 319 | 35.4 | 6.18 |
| P03891 | 2.59 | 1 | 1 | 1 | 2 | | | | | | | | | | | | | | | | | | | | | | | | | | | | 2.59 | 1 | 2 | 347 | 38.9 | 9.83 |
| Q9Y2T2 | 10.29 | 1 | 3 | 3 | 3 | | | | | | | | | | | | | | | | | | | | | | | | | | | | 10.29 | 3 | 3 | 418 | 46.9 | 6.93 |
| Q9H4M3 | 11.37 | 1 | 1 | 2 | 2 | | | | | | | | | | | | | | | | | | | | | | | | | | | | 11.37 | 2 | 2 | 255 | 29.7 | 5.82 |
| Q9Y2T7 | 23.90 | 1 | 5 | 5 | 6 | | | | | | | | | | | | | | | | | | | | | | | | | | | | 23.90 | 5 | 6 | 364 | 38.5 | 10.80 |
| Q9Y2V2 | 44.22 | 1 | 7 | 7 | 19 | | | | | | | | | | 10.88 | 1 | 1 | 18.37 | 2 | 2 | | | | 18.37 | 2 | 2 | 23.13 | 3 | 3 | 13.61 | 2 | 2 | 30.61 | 4 | 9 | 147 | 15.9 | 8.21 |
| Q9Y2V7 | 3.96 | 1 | 2 | 2 | 2 | | | | | | | | | | | | | | | | | | | | | | | | | | | | 3.96 | 2 | 2 | 657 | 73.2 | 5.76 |
| Q9Y2Z0 | 49.32 | 1 | 15 | 15 | 51 | 16.44 | 5 | 5 | 11.51 | 4 | 4 | 8.77 | 3 | 3 | 7.40 | 2 | 3 | 4.11 | 1 | 1 | | | | 10.41 | 3 | 3 | 14.25 | 4 | 4 | 10.14 | 3 | 3 | 41.10 | 11 | 25 | 365 | 41.0 | 5.16 |
| Q9Y305 | 4.56 | 1 | 2 | 2 | 2 | | | | | | | | | | | | | | | | | | | | | | | | | | | | 4.56 | 2 | 2 | 439 | 49.9 | 8.60 |
| Q9Y312 | 5.73 | 1 | 2 | 2 | 2 | | | | | | | 1.82 | 1 | 1 | | | | | | | | | | | | | | | | | | | 3.91 | 1 | 1 | 384 | 43.4 | 5.96 |
| Q99943 | 7.42 | 1 | 1 | 1 | 2 | | | | | | | | | | | | | | | | | | | | | | | | | | | | 7.42 | 1 | 2 | 283 | 31.7 | 9.38 |
| O95379 | 7.07 | 1 | 1 | 1 | 2 | | | | | | | | | | | | | | | | | | | | | | | | | | | | 7.07 | 1 | 2 | 198 | 23.0 | 7.93 |
| Q9Y315 | 27.04 | 1 | 6 | 6 | 8 | 3.77 | 1 | 1 | 3.77 | 1 | 1 | | | | | | | | | | | | | | | | | | | | | | 27.04 | 6 | 6 | 318 | 35.2 | 8.94 |
| Q9Y333 | 40.00 | 1 | 3 | 3 | 3 | 8.42 | 1 | 1 | 11.58 | 1 | 1 | | | | | | | | | | | | | | | | | | | | | | 20.00 | 1 | 1 | 95 | 10.8 | 6.52 |
| Q9Y365 | 47.08 | 1 | 9 | 9 | 24 | | | | | | | | | | | | | | | | | | | | | | | | | 11.68 | 2 | 2 | 40.89 | 8 | 22 | 291 | 33.0 | 7.12 |
| Q96M93 | 4.34 | 1 | 1 | 1 | 2 | | | | | | | | | | | | | | | | | | | | | | | | | | | | 4.34 | 1 | 2 | 576 | 64.4 | 9.01 |
| Q8TEQ8 | 0.92 | 1 | 1 | 1 | 2 | | | | | | | | | | | | | | | | | | | | | | | | | | | | 0.92 | 1 | 2 | 1089 | 118.6 | 8.12 |
| Q9Y366 | 18.08 | 1 | 5 | 5 | 7 | | | | | | | | | | | | | | | | | | | | | | | | | | | | 18.08 | 5 | 7 | 437 | 49.7 | 5.26 |
| Q9Y371 | 32.88 | 1 | 12 | 13 | 57 | 20.00 | 7 | 7 | 14.25 | 6 | 6 | 16.44 | 6 | 6 | 4.38 | 2 | 2 | 6.58 | 3 | 3 | | | | 7.95 | 3 | 3 | 3.56 | 1 | 1 | 10.96 | 3 | 3 | 27.67 | 12 | 26 | 365 | 40.8 | 6.04 |
| Q9Y376 | 22.58 | 1 | 6 | 7 | 10 | | | | | | | | | | | | | | | | | | | | | | | | | | | | 22.58 | 7 | 10 | 341 | 39.8 | 6.89 |
| Q9Y394 | 48.97 | 1 | 15 | 15 | 45 | 14.75 | 4 | 4 | 8.26 | 2 | 2 | 8.26 | 2 | 3 | 5.60 | 2 | 2 | | | | | | | | | | | | | | | | 48.97 | 15 | 34 | 339 | 38.3 | 8.32 |
| Q6NXP6 | 5.01 | 1 | 1 | 1 | 2 | | | | | | | | | | | | | | | | | | | | | | | | | | | | 5.01 | 1 | 2 | 359 | 39.9 | 6.67 |
| Q9Y3A3 | 12.44 | 1 | 2 | 2 | 2 | | | | | | | | | | | | | | | | | | | | | | | | | | | | 12.44 | 2 | 2 | 225 | 26.0 | 5.78 |
| Q5XXA6 | 1.42 | 1 | 1 | 1 | 2 | | | | | | | | | | | | | | | | | | | | | | | | | | | | 1.42 | 1 | 2 | 986 | 114.0 | 8.54 |
| Q9Y3A5 | 66.40 | 1 | 16 | 16 | 33 | | | | | | | | | | | | | | | | | | | 7.20 | 2 | 2 | | | | | | | 66.40 | 15 | 31 | 250 | 28.7 | 8.75 |
| Q5T0D9 | 2.94 | 1 | 1 | 1 | 2 | | | | | | | | | | | | | | | | | | | | | | | | | | | | 2.94 | 1 | 2 | 272 | 30.2 | 7.34 |
| Q9Y3A6 | 10.04 | 1 | 2 | 2 | 3 | 10.04 | 1 | 1 | | | | | | | | | | | | | | | | | | | | | | | | | 5.24 | 1 | 2 | 229 | 26.0 | 4.84 |
| Q9Y2H1 | 2.16 | 1 | 1 | 1 | 2 | | | | | | | | | | | | | | | | | | | | | | | | | | | | 2.16 | 1 | 2 | 464 | 54.0 | 6.81 |
| Q9BXF6 | 2.76 | 1 | 1 | 1 | 2 | | | | | | | | | | | | | | | | | | | | | | | | | | | | 2.76 | 1 | 2 | 653 | 70.4 | 9.23 |
| O43323 | 2.78 | 1 | 1 | 1 | 2 | | | | | | | | | | | | | | | | | | | | | | | | | | | | 2.78 | 1 | 2 | 396 | 43.5 | 9.28 |
| Q9Y3B3 | 33.48 | 1 | 7 | 7 | 33 | 9.82 | 2 | 3 | 9.82 | 2 | 2 | 24.11 | 4 | 5 | 4.46 | 1 | 1 | | | | | | | | | | | | | | | | 33.48 | 7 | 22 | 224 | 25.2 | 6.89 |
| Q68EM7 | 1.02 | 1 | 1 | 1 | 2 | | | | | | | | | | | | | | | | | | | | | | | | | | | | 1.02 | 1 | 2 | 881 | 95.4 | 7.62 |
| Q8N122 | 1.42 | 1 | 1 | 1 | 2 | | | | | | | | | | | | | | | | | | | | | | | | | | | | 1.42 | 1 | 2 | 1335 | 148.9 | 6.89 |
| Q9Y3B8 | 21.52 | 1 | 4 | 4 | 6 | | | | | | | | | | | | | | | | | | | | | | | | | | | | 21.52 | 4 | 6 | 237 | 26.8 | 6.87 |
| Q9Y3C4 | 30.86 | 1 | 4 | 4 | 6 | | | | | | | | | | | | | | | | | | | | | | | | | | | | 30.86 | 4 | 6 | 175 | 19.6 | 6.79 |
| Q9Y3C6 | 19.28 | 1 | 3 | 3 | 4 | | | | | | | | | | | | | | | | | | | | | | | | | | | | 19.28 | 3 | 4 | 166 | 18.2 | 7.99 |
| Q32NB8 | 2.52 | 1 | 1 | 1 | 2 | | | | | | | | | | | | | | | | | | | | | | | | | | | | 2.52 | 1 | 2 | 556 | 62.7 | 8.90 |
| Q9Y3C8 | 20.36 | 1 | 4 | 4 | 5 | | | | | | | | | | | | | | | | | | | | | | | | | | | | 20.36 | 4 | 5 | 167 | 19.4 | 7.40 |
| Q9Y3D0 | 39.26 | 1 | 3 | 3 | 3 | | | | | | | | | | | | | | | | | | | 13.50 | 1 | 1 | | | | | | | 25.77 | 2 | 2 | 163 | 17.7 | 5.19 |
| P62341 | 10.77 | 1 | 1 | 1 | 2 | | | | | | | | | | | | | | | | | | | | | | | | | | | | 10.77 | 1 | 2 | 195 | 22.3 | 8.60 |
| Q9Y3D6 | 20.39 | 1 | 3 | 3 | 5 | 7.24 | 1 | 1 | 7.24 | 1 | 1 | | | | | | | | | | | | | | | | | | | | | | 20.39 | 3 | 3 | 152 | 16.9 | 8.79 |
| Q15398 | 2.01 | 2 | 1 | 2 | 2 | | | | | | | | | | | | | | | | | | | | | | | | | | | | 2.01 | 2 | 2 | 846 | 95.1 | 9.00 |
| Q9Y3E0 | 26.09 | 1 | 3 | 3 | 9 | | | | | | | | | | | | | | | | | | | | | | | | | | | | 26.09 | 3 | 9 | 138 | 15.4 | 10.36 |
| Q9Y3E5 | 31.84 | 1 | 3 | 3 | 3 | | | | | | | | | | | | | | | | | | | | | | | | | | | | 31.84 | 3 | 3 | 179 | 19.2 | 8.73 |
| Q9UMF0 | 0.76 | 1 | 1 | 1 | 2 | | | | | | | | | | | | | | | | | | | | | | | | | | | | 0.76 | 1 | 2 | 924 | 97.1 | 5.95 |
| Q9NW68 | 2.79 | 1 | 1 | 1 | 2 | | | | | | | | | | | | | | | | | | | | | | | | | | | | 2.79 | 1 | 2 | 430 | 47.1 | 4.49 |
| Q9Y3E7 | 12.16 | 1 | 3 | 3 | 3 | | | | | | | | | | | | | | | | | | | | | | | | | | | | 12.16 | 3 | 3 | 222 | 25.1 | 5.12 |
| Q3MJ40 | 0.97 | 2 | 1 | 1 | 2 | | | | | | | | | | | | | | | | | | | | | | | | | | | | 0.97 | 1 | 2 | 725 | 82.9 | 5.01 |
| Q9Y3F4 | 44.57 | 1 | 13 | 13 | 29 | 5.14 | 1 | 1 | | | | | | | 11.14 | 3 | 3 | | | | | | | 3.71 | 1 | 1 | 12.86 | 3 | 3 | 23.43 | 6 | 6 | 37.71 | 10 | 15 | 350 | 38.4 | 5.12 |
| Q5T5C0 | 1.04 | 1 | 1 | 1 | 2 | | | | | | | | | | | | | | | | | | | | | | | | | | | | 1.04 | 1 | 2 | 1151 | 127.5 | 7.28 |
| P35226 | 3.37 | 1 | 1 | 1 | 2 | | | | | | | | | | | | | | | | | | | | | | | | | | | | 3.37 | 1 | 2 | 326 | 36.9 | 8.63 |
| P42331 | 1.86 | 1 | 1 | 1 | 2 | | | | | | | | | | | | | | | | | | | | | | | | | | | | 1.86 | 1 | 2 | 645 | 73.4 | 6.37 |
| P42574 | 6.86 | 1 | 1 | 2 | 2 | | | | | | | | | | | | | | | | | | | | | | | | | | | | 6.86 | 2 | 2 | 277 | 31.6 | 6.54 |
| Q9Y3I0 | 28.51 | 1 | 14 | 14 | 22 | 1.39 | 1 | 1 | | | | 2.18 | 1 | 1 | 1.39 | 1 | 1 | | | | | | | | | | 2.18 | 1 | 2 | 10.30 | 5 | 5 | 20.40 | 9 | 12 | 505 | 55.2 | 7.23 |
| Q8NEE8 | 1.72 | 1 | 1 | 1 | 2 | | | | | | | | | | | | | | | | | | | | | | | | | | | | 1.72 | 1 | 2 | 873 | 98.2 | 9.01 |
| Q9Y3P9 | 5.52 | 1 | 5 | 5 | 7 | | | | | | | | | | | | | | | | | | | | | | | | | | | | 5.52 | 5 | 7 | 1069 | 121.7 | 5.25 |
| Q15431 | 0.61 | 1 | 1 | 1 | 2 | | | | | | | | | | | | | | | | | | | | | | | | | | | | 0.61 | 1 | 2 | 976 | 114.1 | 5.96 |
| Q86VW1 | 2.08 | 1 | 1 | 1 | 2 | | | | | | | | | | | | | | | | | | | | | | | | | | | | 2.08 | 1 | 2 | 577 | 64.6 | 8.15 |
| Q9NUY8 | 2.43 | 1 | 1 | 1 | 2 | | | | | | | | | | | | | | | | | | | | | | | | | | | | 2.43 | 1 | 2 | 699 | 78.3 | 5.41 |
| Q9Y3Q0 | 10.41 | 1 | 5 | 7 | 10 | | | | | | | 0.81 | 1 | 1 | | | | | | | | | | | | | | | | | | | 10.41 | 7 | 9 | 740 | 83.5 | 8.50 |
| Q9Y3Q3 | 14.29 | 1 | 3 | 3 | 6 | | | | | | | | | | | | | | | | | | | | | | | | | | | | 14.29 | 3 | 6 | 217 | 24.8 | 5.60 |
| P0C869 | 0.90 | 1 | 1 | 1 | 2 | | | | | | | | | | | | | | | | | | | | | | | | | | | | 0.90 | 1 | 2 | 781 | 87.9 | 5.96 |
| Q01968 | 1.11 | 1 | 1 | 1 | 2 | | | | | | | | | | | | | | | | | | | | | | | | | | | | 1.11 | 1 | 2 | 901 | 104.1 | 6.55 |
| P48059 | 3.69 | 1 | 1 | 1 | 2 | | | | | | | | | | | | | | | | | | | | | | | | | | | | 3.69 | 1 | 2 | 325 | 37.2 | 8.05 |
| Q8TCY9 | 1.07 | 1 | 1 | 1 | 2 | | | | | | | | | | | | | | | | | | | | | | | | | | | | 1.07 | 1 | 2 | 931 | 104.9 | 6.42 |
| Q5JWR5 | 0.69 | 1 | 1 | 2 | 2 | | | | | | | | | | | | | | | | | | | | | | | | | | | | 0.69 | 2 | 2 | 2465 | 277.2 | 6.29 |
| Q5GH77 | 3.49 | 1 | 1 | 1 | 2 | | | | | | | | | | | | | | | | | | | | | | | | | | | | 3.49 | 1 | 2 | 459 | 53.4 | 8.95 |
| Q9NXE8 | 1.65 | 2 | 1 | 2 | 2 | | | | | | | | | | | | | | | | | | | | | | | | | | | | 1.65 | 2 | 2 | 425 | 49.6 | 10.18 |
| Q6ZVX9 | 3.71 | 1 | 1 | 1 | 2 | | | | | | | | | | | | | | | | | | | | | | | | | | | | 3.71 | 1 | 2 | 377 | 42.7 | 8.59 |
| P98155 | 1.03 | 1 | 1 | 1 | 2 | | | | | | | | | | | | | | | | | | | | | | | | | | | | 1.03 | 1 | 2 | 873 | 96.0 | 4.79 |
| Q14689 | 1.08 | 1 | 1 | 2 | 2 | | | | | | | | | | | | | | | | | | | | | | | | | | | | 1.08 | 2 | 2 | 1571 | 170.3 | 8.03 |
| Q9Y3R5 | 9.44 | 3 | 16 | 17 | 33 | | | | 0.00 | | | 0.61 | 1 | 1 | | | | | | | | | | | | | | | | | | | 9.44 | 17 | 32 | 2298 | 258.1 | 6.29 |
| Q9Y625 | 5.77 | 1 | 1 | 2 | 2 | | | | | | | | | | | | | | | | | | | | | | | | | | | | 5.77 | 2 | 2 | 555 | 62.7 | 5.41 |
| Q8J025 | 4.67 | 1 | 1 | 1 | 2 | | | | | | | | | | | | | | | | | | | | | | | | | | | | 4.67 | 1 | 2 | 514 | 58.8 | 8.15 |
| Q96J92 | 0.80 | 1 | 1 | 1 | 2 | | | | | | | | | | | | | | | | | | | | | | | | | | | | 0.80 | 1 | 2 | 1243 | 134.7 | 5.48 |
| P31629 | 0.25 | 1 | 1 | 1 | 2 | | | | | | | | | | | | | | | | | | | | | | | | | | | | 0.25 | 1 | 2 | 2446 | 268.9 | 6.96 |
| Q9Y2G3 | 1.44 | 1 | 1 | 2 | 2 | | | | | | | | | | | | | | | | | | | | | | | | | | | | 1.44 | 2 | 2 | 1177 | 134.1 | 6.96 |
| Q16760 | 0.58 | 1 | 1 | 1 | 2 | | | | | | | | | | | | | | | | | | | | | | | | | | | | 0.58 | 1 | 2 | 1214 | 134.4 | 7.56 |
| Q7L099 | 2.35 | 1 | 1 | 1 | 2 | | | | | | | | | | | | | | | | | | | | | | | | | | | | 2.35 | 1 | 2 | 469 | 52.9 | 5.49 |
| Q9Y3U8 | 34.29 | 1 | 3 | 4 | 5 | | | | | | | | | | | | | | | | | | | | | | | | | | | | 34.29 | 4 | 5 | 105 | 12.2 | 11.59 |
| Q9Y448 | 11.39 | 1 | 3 | 3 | 7 | | | | | | | | | | | | | | | | | | | | | | 2.85 | 1 | 1 | 2.85 | 1 | 1 | 11.39 | 3 | 5 | 316 | 35.4 | 6.30 |
| Q9Y450 | 4.82 | 1 | 3 | 3 | 4 | | | | | | | | | | | | | | | | | | | | | | | | | | | | 4.82 | 3 | 4 | 684 | 75.4 | 6.61 |
| Q9Y485 | 10.41 | 2 | 24 | 24 | 36 | | | | | | | | | | | | | | | | | | | | | | | | | | | | 10.41 | 24 | 36 | 3027 | 337.6 | 6.34 |
| Q9NUQ2 | 3.30 | 1 | 1 | 1 | 2 | | | | | | | | | | | | | | | | | | | | | | | | | | | | 3.30 | 1 | 2 | 364 | 42.0 | 9.10 |
| Q71RG4 | 6.23 | 1 | 1 | 2 | 2 | | | | | | | | | | | | | | | | | | | | | | | | | | | | 6.23 | 2 | 2 | 321 | 33.8 | 5.03 |
| P26572 | 4.04 | 1 | 1 | 1 | 2 | | | | | | | | | | | | | | | | | | | | | | | | | | | | 4.04 | 1 | 2 | 445 | 50.8 | 9.16 |
| Q9Y490 | 33.77 | 2 | 52 | 58 | 154 | 1.65 | 3 | 3 | | | | 2.40 | 4 | 4 | 0.55 | 1 | 1 | | | | | | | 2.16 | 4 | 4 | 0.51 | 1 | 1 | 4.76 | 9 | 9 | 31.84 | 53 | 132 | 2541 | 269.6 | 6.07 |
| Q9UIG8 | 1.55 | 1 | 1 | 1 | 2 | | | | | | | | | | | | | | | | | | | | | | | | | | | | 1.55 | 1 | 2 | 710 | 76.5 | 6.73 |
| Q9Y4B6 | 1.86 | 1 | 2 | 2 | 4 | | | | | | | | | | | | | | | | | | | | | | | | | | | | 1.86 | 2 | 4 | 1507 | 168.9 | 5.06 |
| Q9Y4D8 | 0.95 | 2 | 3 | 4 | 6 | | | | | | | | | | 0.23 | 1 | 1 | | | | | | | | | | | | | | | | 0.95 | 4 | 5 | 3996 | 439.1 | 6.19 |
| Q9H6X4 | 6.15 | 1 | 1 | 1 | 2 | | | | | | | | | | | | | | | | | | | | | | | | | | | | 6.15 | 1 | 2 | 195 | 21.6 | 6.54 |
| Q9NPR2 | 2.40 | 2 | 1 | 2 | 2 | | | | | | | | | | | | | | | | | | | | | | | | | | | | 2.40 | 2 | 2 | 832 | 92.1 | 6.95 |
| Q9Y4E6 | 3.02 | 1 | 3 | 3 | 5 | | | | | | | | | | | | | | | | | | | | | | | | | | | | 3.02 | 3 | 5 | 1490 | 163.7 | 6.92 |
| Q9Y4I1 | 7.44 | 1 | 8 | 13 | 26 | | | | | | | | | | | | | | | | | | | 0.75 | 1 | 1 | | | | 0.75 | 1 | 1 | 7.44 | 13 | 24 | 1855 | 215.3 | 8.48 |
| Q8NEC5 | 2.95 | 1 | 1 | 1 | 2 | | | | | | | | | | | | | | | | | | | | | | | | | | | | 2.95 | 1 | 2 | 780 | 90.0 | 7.71 |
| Q9Y4L1 | 56.26 | 1 | 56 | 57 | 393 | 38.84 | 29 | 38 | 42.94 | 34 | 62 | 45.05 | 35 | 53 | 11.61 | 10 | 11 | 18.42 | 17 | 21 | | | | 8.21 | 9 | 9 | 14.51 | 13 | 16 | 14.71 | 13 | 16 | 48.05 | 42 | 167 | 999 | 111.3 | 5.22 |
| Q8N9H8 | 0.80 | 1 | 1 | 1 | 2 | | | | | | | | | | | | | | | | | | | | | | | | | | | | 0.80 | 1 | 2 | 876 | 96.5 | 8.22 |
| Q9Y4P1 | 37.40 | 1 | 9 | 9 | 17 | | | | | | | | | | | | | | | | | | | | | | | | | | | | 37.40 | 9 | 17 | 393 | 44.3 | 5.07 |
| Q9Y4P3 | 53.24 | 1 | 20 | 20 | 47 | 5.59 | 2 | 2 | 2.91 | 1 | 1 | 15.66 | 5 | 5 | 2.91 | 1 | 1 | 2.91 | 1 | 1 | | | | 2.91 | 1 | 1 | | | | 2.91 | 1 | 1 | 53.24 | 20 | 35 | 447 | 49.8 | 9.44 |
| P50613 | 3.18 | 1 | 1 | 1 | 2 | | | | | | | | | | | | | | | | | | | | | | | | | | | | 3.18 | 1 | 2 | 346 | 39.0 | 8.47 |
| Q9UFC0 | 2.16 | 1 | 1 | 1 | 2 | | | | | | | | | | | | | | | | | | | | | | | | | | | | 2.16 | 1 | 2 | 647 | 70.8 | 7.21 |
| Q9Y4R8 | 5.26 | 1 | 4 | 4 | 4 | | | | | | | | | | | | | | | | | | | | | | | | | | | | 5.26 | 4 | 4 | 837 | 91.7 | 5.76 |
| Q9Y4W6 | 40.65 | 3 | 28 | 28 | 39 | 4.52 | 3 | 3 | 5.02 | 3 | 3 | | | | | | | | | | | | | | | | | | | | | | 38.27 | 26 | 33 | 797 | 88.5 | 8.66 |
| Q9Y4X5 | 8.98 | 1 | 3 | 3 | 9 | | | | | | | | | | | | | | | | | | | | | | | | | | | | 8.98 | 3 | 9 | 557 | 64.1 | 5.08 |
| Q9Y4Z0 | 15.11 | 1 | 3 | 3 | 3 | 0.00 | | | | | | | | | 0.00 | | | | | | | | | | | | | | | | | | 15.11 | 3 | 3 | 139 | 15.3 | 9.99 |
| Q8N6L1 | 12.50 | 1 | 1 | 1 | 2 | | | | | | | | | | | | | | | | | | | | | | | | | | | | 12.50 | 1 | 2 | 136 | 14.7 | 9.61 |
| Q92845 | 2.53 | 1 | 1 | 1 | 2 | | | | | | | | | | | | | | | | | | | | | | | | | | | | 2.53 | 1 | 2 | 792 | 91.1 | 5.08 |
| Q9Y508 | 14.47 | 1 | 3 | 3 | 6 | | | | | | | | | | | | | | | | | | | | | | | | | | | | 14.47 | 3 | 6 | 228 | 25.7 | 7.25 |
| Q9Y512 | 33.26 | 1 | 11 | 11 | 16 | | | | 2.35 | 1 | 1 | | | | | | | 0.00 | | | | | | | | | | | | | | | 33.26 | 11 | 15 | 469 | 51.9 | 6.90 |
| Q9Y530 | 46.71 | 1 | 6 | 6 | 7 | | | | | | | | | | | | | | | | | | | | | | 5.92 | 1 | 1 | | | | 46.71 | 6 | 6 | 152 | 17.0 | 8.31 |
| Q07889 | 0.53 | 1 | 1 | 1 | 2 | | | | | | | | | | | | | | | | | | | | | | | | | | | | 0.53 | 1 | 2 | 1333 | 152.4 | 6.84 |
| Q9Y547 | 36.11 | 1 | 3 | 3 | 14 | | | | | | | | | | | | | | | | | | | 17.36 | 1 | 3 | 27.78 | 2 | 2 | 27.78 | 2 | 2 | 36.11 | 3 | 7 | 144 | 16.3 | 5.03 |
| Q9H5U6 | 1.36 | 1 | 1 | 1 | 2 | | | | | | | | | | | | | | | | | | | | | | | | | | | | 1.36 | 1 | 2 | 513 | 59.0 | 8.75 |
| Q9Y570 | 19.43 | 1 | 7 | 7 | 9 | | | | | | | | | | | | | | | | | | | | | | | | | | | | 19.43 | 7 | 9 | 386 | 42.3 | 5.97 |
| Q9Y587 | 14.58 | 1 | 2 | 2 | 3 | | | | | | | | | | | | | | | | | | | | | | | | | | | | 14.58 | 2 | 3 | 144 | 17.0 | 5.19 |
| Q9Y5A7 | 2.76 | 1 | 2 | 2 | 2 | | | | | | | | | | | | | | | | | | | | | | | | | | | | 2.76 | 2 | 2 | 615 | 70.5 | 5.96 |
| Q9Y5B8 | 76.86 | 1 | 26 | 26 | 95 | 8.51 | 2 | 2 | 15.43 | 5 | 5 | | | | 46.01 | 16 | 19 | 32.71 | 11 | 13 | | | | | | | | | | | | | 72.34 | 22 | 56 | 376 | 42.5 | 6.47 |
| P30408 | 4.46 | 1 | 1 | 1 | 2 | | | | | | | | | | | | | | | | | | | | | | | | | | | | 4.46 | 1 | 2 | 202 | 21.6 | 4.98 |
| Q9Y5K5 | 50.76 | 2 | 11 | 12 | 26 | | | | 3.34 | 1 | 1 | | | | | | | | | | | | | | | | 3.34 | 1 | 1 | 3.34 | 1 | 1 | 47.42 | 11 | 23 | 329 | 37.6 | 5.33 |
| P78356 | 3.37 | 1 | 1 | 1 | 2 | | | | | | | | | | | | | | | | | | | | | | | | | | | | 3.37 | 1 | 2 | 416 | 47.3 | 7.33 |
| P15151 | 3.36 | 1 | 1 | 1 | 2 | | | | | | | | | | | | | | | | | | | | | | | | | | | | 3.36 | 1 | 2 | 417 | 45.3 | 6.52 |
| Q7Z4R8 | 14.14 | 1 | 1 | 1 | 2 | | | | | | | | | | | | | | | | | | | | | | | | | | | | 14.14 | 1 | 2 | 191 | 20.8 | 4.84 |
| Q9Y5K6 | 5.32 | 1 | 3 | 3 | 5 | 1.56 | 1 | 1 | | | | | | | | | | | | | | | | | | | 2.19 | 1 | 2 | 3.76 | 2 | 2 | | | | 639 | 71.4 | 6.40 |
| C9JQI7 | 2.13 | 1 | 1 | 1 | 2 | | | | | | | | | | | | | | | | | | | | | | | | | | | | 2.13 | 1 | 2 | 657 | 76.4 | 8.06 |
| Q9Y5K8 | 31.58 | 1 | 6 | 6 | 7 | | | | | | | | | | | | | | | | | | | | | | | | | | | | 31.58 | 6 | 7 | 247 | 28.2 | 9.36 |
| Q9Y5L0 | 18.96 | 1 | 11 | 11 | 22 | | | | | | | | | | | | | | | | | | | | | | | | | | | | 18.96 | 11 | 22 | 923 | 104.1 | 5.57 |
| O00258 | 12.07 | 1 | 1 | 1 | 2 | | | | | | | | | | | | | | | | | | | | | | | | | | | | 12.07 | 1 | 2 | 174 | 19.8 | 9.67 |

| | | | | | | | | | | | | | | | | | | | | | | | | | | | | | | | | | | | | | | |
|---|---|---|---|---|---|---|---|---|---|---|---|---|---|---|---|---|---|---|---|---|---|---|---|---|---|---|---|---|---|---|---|---|---|---|---|---|---|---|
| Q9Y5M8 | 35.42 | 1 | 8 | 8 | 12 | | | | 3.69 | 1 | 1 | 2.95 | 1 | 1 | | | | | | | | | | | | | | | | | | | 28.78 | 6 | 10 | 271 | 29.7 | 9.04 |
| Q9Y5P4 | 8.65 | 1 | 4 | 4 | 8 | | | | | | | | | | | | | | | | | | | | | | | | | | | | 8.65 | 4 | 8 | 624 | 70.8 | 5.48 |
| Q8NEE6 | 0.82 | 1 | 1 | 1 | 2 | | | | | | | | | | | | | | | | | | | | | | | | | | | | 0.82 | 1 | 2 | 735 | 83.9 | 8.72 |
| Q13394 | 3.62 | 1 | 1 | 1 | 2 | | | | | | | | | | | | | | | | | | | | | | | | | | | | 3.62 | 1 | 2 | 359 | 40.9 | 8.66 |
| Q9Y5P6 | 33.89 | 1 | 9 | 9 | 19 | 5.00 | 1 | 1 | | | | 5.00 | 1 | 1 | 5.00 | 1 | 1 | | | | | | | | | | | | | | | | 33.89 | 9 | 16 | 360 | 39.8 | 6.61 |
| Q9Y5S2 | 5.38 | 1 | 6 | 8 | 13 | | | | | | | | | | | | | | | | | | | | | | | | | 0.94 | 1 | 1 | 4.44 | 7 | 12 | 1711 | 194.2 | 6.37 |
| Q9Y5U8 | 52.29 | 1 | 5 | 5 | 10 | | | | | | | | | | | | | | | | | | | | | | | | | | | | 52.29 | 5 | 10 | 109 | 12.3 | 9.61 |
| P18827 | 5.48 | 1 | 1 | 1 | 2 | | | | | | | | | | | | | | | | | | | | | | | | | | | | 5.48 | 1 | 2 | 310 | 32.4 | 4.63 |
| Q9Y5X1 | 17.82 | 1 | 7 | 7 | 23 | | | | | | | | | | | | | | | | | | | | | | | | | | | | 17.82 | 7 | 23 | 595 | 66.5 | 5.58 |
| Q9Y5X2 | 4.52 | 1 | 2 | 2 | 2 | | | | | | | | | | | | | | | | | | | | | | | | | | | | 4.52 | 2 | 2 | 465 | 52.5 | 7.39 |
| Q8NE01 | 1.13 | 2 | 1 | 1 | 2 | | | | | | | | | | | | | | | | | | | | | | | | | | | | 1.13 | 1 | 2 | 707 | 76.1 | 6.09 |
| O95807 | 5.10 | 1 | 1 | 1 | 2 | | | | | | | | | | | | | | | | | | | | | | | | | | | | 5.10 | 1 | 2 | 157 | 17.4 | 5.87 |
| Q9Y5R8 | 11.03 | 1 | 1 | 1 | 2 | | | | | | | | | | | | | | | | | | | | | | | | | | | | 11.03 | 1 | 2 | 145 | 16.8 | 9.16 |
| O60603 | 1.02 | 1 | 1 | 1 | 2 | | | | | | | | | | | | | | | | | | | | | | | | | | | | 1.02 | 1 | 2 | 784 | 89.8 | 6.61 |
| O75426 | 1.55 | 1 | 1 | 1 | 2 | | | | | | | | | | | | | | | | | | | | | | | | | | | | 1.55 | 1 | 2 | 580 | 64.8 | 8.69 |
| Q9BVT8 | 4.07 | 1 | 1 | 1 | 2 | | | | | | | | | | | | | | | | | | | | | | | | | | | | 4.07 | 1 | 2 | 246 | 26.2 | 5.72 |
| Q9Y5X3 | 24.75 | 1 | 8 | 10 | 15 | | | | 1.49 | 1 | 1 | | | | | | | | | | | | | | | | | | | 5.20 | 2 | 2 | 20.54 | 8 | 12 | 404 | 46.8 | 6.76 |
| Q9Y5X9 | 19.60 | 1 | 7 | 8 | 20 | 3.60 | 1 | 1 | | | | | | | | | | | | | | | | | | | | 2.20 | 1 | 1 | 5.80 | 2 | 2 | 19.40 | 7 | 16 | 500 | 56.8 | 7.93 |
| Q9BV10 | 2.05 | 1 | 1 | 1 | 2 | | | | | | | | | | | | | | | | | | | | | | | | | | | | 2.05 | 1 | 2 | 488 | 54.6 | 9.58 |
| Q9Y5Y2 | 54.61 | 1 | 8 | 8 | 20 | | | | | | | | | | | | | | | | | | | | | | | | | | | | 54.61 | 8 | 20 | 271 | 28.8 | 5.83 |
| Q9NX04 | 11.33 | 1 | 1 | 1 | 2 | | | | | | | | | | | | | | | | | | | | | | | | | | | | 11.33 | 1 | 2 | 203 | 23.4 | 5.67 |
| Q3KRA6 | 13.49 | 1 | 1 | 1 | 2 | | | | | | | | | | | | | | | | | | | | | | | | | | | | 13.49 | 1 | 2 | 126 | 14.6 | 6.93 |
| Q9Y5Y5 | 19.05 | 1 | 6 | 6 | 9 | | | | | | | | | | | | | | | | | | | | | | | | | | | | 19.05 | 6 | 9 | 336 | 38.6 | 9.85 |
| Q9Y5Z4 | 41.95 | 1 | 10 | 10 | 33 | 19.51 | 3 | 3 | | | | | | | 9.76 | 2 | 2 | | | | | | | 30.24 | 6 | 7 | 11.22 | 3 | 3 | 32.20 | 6 | 9 | 27.32 | 5 | 9 | 205 | 22.9 | 4.63 |
| O95406 | 20.14 | 1 | 1 | 1 | 2 | | | | | | | | | | | | | | | | | | | | | | | | | | | | 20.14 | 1 | 2 | 144 | 16.7 | 5.71 |
| Q9H693 | 10.13 | 1 | 1 | 1 | 2 | | | | | | | | | | | | | | | | | | | | | | | | | | | | 10.13 | 1 | 2 | 158 | 16.8 | 9.63 |
| Q9BUF7 | 14.17 | 1 | 1 | 1 | 2 | | | | | | | | | | | | | | | | | | | | | | | | | | | | 14.17 | 1 | 2 | 120 | 12.8 | 9.96 |
| Q9Y614 | 29.40 | 1 | 7 | 7 | 11 | | | | | | | 1.93 | 1 | 1 | | | | | | | | | | | | | | | | | | | 29.40 | 7 | 10 | 415 | 45.2 | 5.49 |
| Q6ICB0 | 14.88 | 1 | 1 | 1 | 2 | | | | | | | | | | | | | | | | | | | | | | | | | | | | 14.88 | 1 | 2 | 168 | 18.3 | 4.94 |
| P13762 | 5.64 | 1 | 1 | 2 | 2 | | | | | | | | | | | | | | | | | | | | | | | | | | | | 5.64 | 2 | 2 | 266 | 29.9 | 7.33 |
| Q9H479 | 3.24 | 1 | 1 | 1 | 2 | | | | | | | | | | | | | | | | | | | | | | | | | | | | 3.24 | 1 | 2 | 309 | 35.1 | 7.55 |
| Q9Y615 | 16.55 | 1 | 6 | 6 | 9 | | | | | | | | | | | | | | | | | | | | | | | | | | | | 16.55 | 6 | 9 | 435 | 48.6 | 7.09 |
| Q75N03 | 2.85 | 1 | 1 | 1 | 2 | | | | | | | | | | | | | | | | | | | | | | | | | | | | 2.85 | 1 | 2 | 491 | 54.5 | 8.29 |
| Q9Y617 | 19.19 | 1 | 7 | 7 | 13 | | | | | | | | | | 2.70 | 1 | 1 | | | | | | | | | | | | | 2.70 | 1 | 1 | 19.19 | 7 | 11 | 370 | 40.4 | 7.66 |
| P0C7M6 | 7.14 | 1 | 1 | 1 | 2 | | | | | | | | | | | | | | | | | | | | | | | | | | | | 7.14 | 1 | 2 | 154 | 18.2 | 10.48 |
| Q9NQX7 | 3.75 | 1 | 1 | 1 | 2 | | | | | | | | | | | | | | | | | | | | | | | | | | | | 3.75 | 1 | 2 | 267 | 30.2 | 8.00 |
| Q9Y619 | 8.97 | 1 | 3 | 3 | 3 | | | | | | | | | | | | | | | | | | | | | | | | | | | | 8.97 | 3 | 3 | 301 | 32.7 | 9.13 |
| Q9Y639 | 16.83 | 1 | 6 | 6 | 8 | | | | | | | 2.76 | 1 | 1 | | | | 4.02 | 1 | 1 | | | | | | | | | | | | | 12.81 | 4 | 6 | 398 | 44.4 | 7.99 |
| O75121 | 5.62 | 1 | 1 | 1 | 2 | | | | | | | | | | | | | | | | | | | | | | | | | | | | 5.62 | 1 | 2 | 409 | 45.4 | 5.22 |
| Q9Y678 | 54.46 | 1 | 30 | 33 | 130 | 1.37 | 1 | 1 | 1.37 | 1 | 1 | 1.37 | 1 | 1 | | | | | | | | | | | | | 0.92 | 1 | 2 | 3.32 | 2 | 2 | 54.46 | 33 | 123 | 874 | 97.7 | 5.47 |
| Q9Y696 | 61.66 | 4 | 9 | 11 | 46 | 2.37 | 1 | 1 | 2.37 | 1 | 1 | 10.28 | 3 | 3 | 4.74 | 2 | 2 | | | | | | | 7.11 | 2 | 2 | | | | 9.49 | 3 | 3 | 61.66 | 11 | 34 | 253 | 28.8 | 5.59 |
| Q9Y697 | 45.73 | 1 | 19 | 19 | 33 | 13.35 | 4 | 6 | 22.54 | 8 | 9 | 8.32 | 3 | 3 | | | | | | | | | | | | | | | | | | | 33.70 | 14 | 15 | 457 | 50.2 | 8.31 |
| Q9Y6A4 | 74.61 | 1 | 15 | 15 | 49 | 6.22 | 1 | 1 | 19.69 | 3 | 3 | | | | 18.13 | 3 | 5 | 9.84 | 2 | 2 | | | | | | | | | | | | | 73.58 | 14 | 38 | 193 | 22.8 | 9.76 |
| Q9NSY0 | 4.79 | 1 | 1 | 2 | 2 | | | | | | | | | | | | | | | | | | | | | | | | | | | | 4.79 | 2 | 2 | 501 | 57.8 | 6.49 |
| Q86UD1 | 9.89 | 1 | 1 | 1 | 2 | | | | | | | | | | | | | | | | | | | | | | | | | | | | 9.89 | 1 | 2 | 273 | 30.7 | 6.84 |
| Q9Y6A9 | 31.37 | 1 | 3 | 3 | 14 | 11.76 | 1 | 1 | 26.47 | 2 | 3 | 11.76 | 1 | 1 | | | | | | | | | | | | | | | | | | | 31.37 | 3 | 9 | 102 | 11.8 | 9.31 |
| Q9UHJ9 | 4.33 | 1 | 1 | 1 | 2 | | | | | | | | | | | | | | | | | | | | | | | | | | | | 4.33 | 1 | 2 | 254 | 29.4 | 8.15 |
| Q8WWB5 | 2.22 | 1 | 1 | 1 | 2 | | | | | | | | | | | | | | | | | | | | | | | | | | | | 2.22 | 1 | 2 | 315 | 35.9 | 6.37 |
| P48378 | 1.24 | 1 | 1 | 1 | 2 | | | | | | | | | | | | | | | | | | | | | | | | | | | | 1.24 | 1 | 2 | 723 | 79.9 | 6.79 |
| Q3KRB8 | 2.25 | 1 | 1 | 1 | 2 | | | | | | | | | | | | | | | | | | | | | | | | | | | | 2.25 | 1 | 2 | 267 | 30.2 | 9.14 |
| Q86VV4 | 1.29 | 1 | 1 | 1 | 2 | | | | | | | | | | | | | | | | | | | | | | | | | | | | 1.29 | 1 | 2 | 465 | 52.2 | 8.95 |
| Q9Y6B6 | 50.51 | 1 | 5 | 10 | 21 | 5.56 | 1 | 1 | 21.21 | 4 | 4 | | | | 10.10 | 2 | 2 | | | | | | | 5.56 | 1 | 1 | | | | | | | 50.00 | 8 | 13 | 198 | 22.4 | 6.11 |
| Q9Y6B7 | 7.31 | 1 | 3 | 3 | 3 | | | | | | | | | | | | | | | | | | | | | | | | | | | | 7.31 | 3 | 3 | 739 | 83.2 | 5.86 |
| Q9Y6C9 | 59.74 | 1 | 15 | 15 | 99 | 34.98 | 8 | 9 | 28.05 | 7 | 12 | 30.36 | 8 | 9 | 14.85 | 3 | 3 | 6.93 | 1 | 2 | | | | | | | | | | | | | 59.74 | 15 | 64 | 303 | 33.3 | 7.97 |
| Q9Y6D5 | 27.23 | 1 | 28 | 41 | 71 | 2.30 | 3 | 3 | 0.73 | 1 | 1 | | | | 0.56 | 1 | 1 | | | | | | | | | | | | | | | | 25.77 | 39 | 66 | 1785 | 201.9 | 6.33 |
| Q9Y6D6 | 12.98 | 1 | 8 | 21 | 34 | | | | | | | | | | | | | | | | | | | | | | | | | | | | 12.98 | 21 | 34 | 1849 | 208.6 | 5.85 |
| Q9Y6E0 | 17.83 | 1 | 4 | 6 | 9 | | | | | | | | | | | | | | | | | | | 2.93 | 1 | 1 | | | | 1.58 | 1 | 1 | 17.83 | 6 | 7 | 443 | 49.3 | 5.69 |
| Q99727 | 5.80 | 1 | 1 | 1 | 2 | | | | | | | | | | | | | | | | | | | | | | | | | | | | 5.80 | 1 | 2 | 224 | 25.5 | 7.80 |
| Q8TB96 | 1.47 | 1 | 1 | 1 | 2 | | | | | | | | | | | | | | | | | | | | | | | | | | | | 1.47 | 1 | 2 | 612 | 68.1 | 5.39 |
| Q9Y6G9 | 18.36 | 1 | 8 | 8 | 19 | 1.72 | 1 | 1 | | | | 1.72 | 1 | 1 | 1.72 | 1 | 1 | | | | | | | 4.59 | 2 | 2 | | | | 4.02 | 2 | 2 | 16.63 | 7 | 13 | 523 | 56.5 | 6.42 |
| E9PQX1 | 6.90 | 1 | 1 | 1 | 2 | | | | | | | | | | | | | | | | | | | | | | | | | | | | 6.90 | 1 | 2 | 116 | 13.7 | 6.62 |
| Q9Y6J8 | 56.55 | 1 | 12 | 12 | 32 | | | | | | | | | | 4.15 | 1 | 1 | | | | | | | | | | | | | | | | 56.55 | 12 | 31 | 313 | 35.8 | 6.14 |
| Q96PN7 | 0.50 | 1 | 1 | 1 | 2 | | | | | | | | | | | | | | | | | | | | | | | | | | | | 0.50 | 1 | 2 | 1200 | 132.2 | 6.74 |
| Q96RU8 | 2.69 | 1 | 1 | 1 | 2 | | | | | | | | | | | | | | | | | | | | | | | | | | | | 2.69 | 1 | 2 | 372 | 41.0 | 7.28 |
| O95164 | 15.38 | 1 | 1 | 2 | 2 | | | | | | | | | | | | | | | | | | | | | | | | | | | | 15.38 | 2 | 2 | 117 | 13.1 | 6.77 |
| Q9Y6M0 | 25.48 | 1 | 6 | 6 | 15 | 2.87 | 1 | 1 | 7.01 | 2 | 2 | | | | | | | | | | | | | | | | | | | | | | 22.61 | 5 | 12 | 314 | 34.9 | 7.62 |
| Q9Y6M9 | 33.52 | 1 | 5 | 5 | 8 | 7.82 | 1 | 1 | | | | | | | | | | | | | | | | | | | | | | | | | 33.52 | 5 | 7 | 179 | 21.8 | 8.38 |
| Q8WWF5 | 4.43 | 1 | 1 | 1 | 2 | | | | | | | | | | | | | | | | | | | | | | | | | | | | 4.43 | 1 | 2 | 429 | 46.9 | 7.99 |
| Q9Y6N5 | 15.11 | 1 | 6 | 6 | 7 | 3.11 | 1 | 1 | | | | | | | | | | | | | | | | | | | | | | | | | 12.00 | 5 | 6 | 450 | 49.9 | 9.11 |
| H3BQJ8 | 10.87 | 1 | 1 | 1 | 2 | | | | | | | | | | | | | | | | | | | | | | | | | | | | 10.87 | 1 | 2 | 138 | 15.3 | 8.65 |
| Q9Y6Q5 | 20.57 | 1 | 6 | 8 | 14 | | | | | | | | | | | | | | | | | | | | | | | | | | | | 20.57 | 8 | 14 | 423 | 48.1 | 8.22 |
| Q9Y6R7 | 6.38 | 1 | 13 | 13 | 13 | | | | | | | 5.61 | 11 | 11 | | | | | | | | | | | | | | | | | | | 0.78 | 2 | 2 | 5405 | 571.6 | 5.34 |
| Q9Y6W5 | 7.43 | 1 | 3 | 3 | 6 | 2.01 | 1 | 1 | | | | | | | | | | 1.61 | 1 | 1 | | | | | | | | | | | | | 3.82 | 1 | 4 | 498 | 54.3 | 5.53 |
| Q9Y2Z9 | 1.92 | 1 | 1 | 1 | 1 | 0.00 | | | | | | | | | | | | | | | | | | | | | | | | | | | 1.92 | 1 | 1 | 468 | 50.8 | 7.30 |
| Q9H9F9 | 1.48 | 1 | 1 | 1 | 1 | 1.48 | 1 | 1 | 0.00 | | | | | | | | | | | | | | | | | | | | | | | | 0.00 | | | 607 | 68.3 | 5.27 |
| Q8IV36 | 1.40 | 1 | 1 | 1 | 1 | 0.00 | | | 0.00 | | | 0.00 | | | 0.00 | | | | | | | | | | | | | | | | | | 1.40 | 1 | 1 | 788 | 88.7 | 6.06 |
| O43147 | 0.70 | 1 | 1 | 1 | 1 | 0.00 | | | 0.00 | | | | | | | | | 0.00 | | | | | | | | | | | | | | | 0.70 | 1 | 1 | 1006 | 113.2 | 6.67 |
| Q9H0K1 | 0.76 | 1 | 1 | 1 | 1 | 0.76 | 1 | 1 | 0.00 | | | 0.00 | | | | | | | | | | | | | | | | | | | | | 0.00 | | | 926 | 103.8 | 6.02 |
| Q9Y2X3 | 1.89 | 1 | 1 | 1 | 1 | 1.89 | 1 | 1 | | | | | | | | | | | | | | | | | | | | | | | | | 0.00 | | | 529 | 59.5 | 8.92 |
| Q86U86 | 0.36 | 1 | 1 | 1 | 1 | 0.36 | 1 | 1 | | | | | | | | | | | | | | | | | | | | | | | | | 0.00 | | | 1689 | 192.8 | 6.89 |
| Q6P2E9 | 1.64 | 1 | 1 | 1 | 1 | 0.00 | | | | | | | | | | | | | | | | | | | | | | | | | | | 1.64 | 1 | 1 | 1401 | 151.6 | 5.86 |
| Q9P2I0 | 1.66 | 1 | 1 | 1 | 1 | 0.00 | | | | | | | | | | | | | | | | | | | | | | | | | | | 1.66 | 1 | 1 | 782 | 88.4 | 5.11 |
| Q8WYS4 | 1.96 | 1 | 1 | 1 | 1 | 0.00 | | | | | | | | | | | | | | | | | | | | | | | | | | | 1.96 | 1 | 1 | 764 | 84.9 | 5.03 |
| Q96KK3 | 1.52 | 1 | 1 | 1 | 1 | 1.52 | 1 | 1 | | | | | | | | | | | | | | | | | | | | | | | | | | | | 526 | 58.3 | 7.15 |
| Q9NXG0 | 0.43 | 1 | 1 | 1 | 1 | 0.00 | | | | | | | | | | | | | | | | | | | | | | | | | | | 0.43 | 1 | 1 | 1405 | 161.5 | 8.15 |
| Q5UIP0 | 0.32 | 1 | 1 | 1 | 1 | 0.00 | | | | | | | | | | | | | | | | | | | | | | | | | | | 0.32 | 1 | 1 | 2472 | 274.3 | 5.52 |
| Q92839 | 1.56 | 1 | 1 | 1 | 1 | 0.00 | | | 0.00 | | | 0.00 | | | 0.00 | | | 0.00 | | | | | | 0.00 | | | 0.00 | | | 0.00 | | | 1.56 | 1 | 1 | 578 | 64.8 | 9.19 |
| Q9NWV8 | 3.34 | 1 | 1 | 1 | 1 | 3.34 | 1 | 1 | | | | | | | | | | | | | | | | | | | | | | | | | | | | 329 | 36.5 | 4.64 |
| Q9UH65 | 2.05 | 1 | 1 | 1 | 1 | 0.00 | | | 0.00 | | | | | | | | | | | | | | | | | | | | | | | | 2.05 | 1 | 1 | 585 | 69.0 | 5.87 |
| Q99661 | 1.79 | 1 | 1 | 1 | 1 | 0.00 | | | | | | 0.00 | | | | | | 0.00 | | | | | | | | | 0.00 | | | 0.00 | | | 1.79 | 1 | 1 | 725 | 81.3 | 7.83 |
| Q5H913 | 2.41 | 1 | 1 | 1 | 1 | 0.00 | | | | | | | | | | | | | | | | | | 0.00 | | | 0.00 | | | | | | 2.41 | 1 | 1 | 290 | 33.0 | 9.03 |
| Q15750 | 2.58 | 1 | 1 | 1 | 1 | 0.00 | | | | | | | | | | | | | | | | | | | | | | | | 2.58 | 1 | 1 | | | | 504 | 54.6 | 5.52 |
| Q9UH99 | 2.65 | 1 | 1 | 1 | 1 | 2.65 | 1 | 1 | | | | | | | | | | | | | | | | | | | | | | | | | | | | 717 | 80.3 | 6.73 |
| Q8IVL1 | 0.32 | 1 | 1 | 1 | 1 | 0.00 | | | | | | 0.00 | | | | | | | | | | | | | | | 0.00 | | | | | | 0.32 | 1 | 1 | 2488 | 268.0 | 9.03 |
| Q9UL12 | 0.87 | 1 | 1 | 1 | 1 | 0.00 | | | 0.00 | | | 0.00 | | | | | | | | | | | | | | | | | | | | | 0.87 | 1 | 1 | 918 | 101.0 | 7.25 |
| P35610 | 2.91 | 1 | 1 | 1 | 1 | 2.91 | 1 | 1 | | | | | | | | | | | | | | | | | | | | | | | | | | | | 550 | 64.7 | 8.94 |
| Q96PV6 | 0.77 | 1 | 1 | 1 | 1 | 0.77 | 1 | 1 | | | | | | | | | | | | | | | | | | | | | | | | | | | | 779 | 86.1 | 9.35 |
| Q9NUW8 | 1.32 | 1 | 1 | 1 | 1 | 0.00 | | | | | | | | | | | | | | | | | | | | | | | | | | | 1.32 | 1 | 1 | 608 | 68.4 | 7.65 |
| Q9UHB7 | 0.52 | 1 | 1 | 1 | 1 | 0.52 | 1 | 1 | | | | | | | | | | | | | | | | | | | | | | | | | 0.00 | | | 1163 | 127.4 | 9.31 |
| Q13009 | 0.38 | 1 | 1 | 1 | 1 | 0.38 | 1 | 1 | | | | | | | | | | | | | | | | | | | | | | | | | 0.00 | | | 1591 | 177.4 | 6.61 |
| Q96AP4 | 1.73 | 1 | 1 | 1 | 1 | 1.73 | 1 | 1 | | | | | | | | | | | | | | | | | | | | | | | | | | | | 578 | 65.9 | 6.49 |
| P51003 | 1.74 | 1 | 1 | 1 | 1 | 1.74 | 1 | 1 | | | | | | | | | | | | | | | | | | | | | | | | | | | | 745 | 82.8 | 7.37 |
| Q15058 | 0.73 | 1 | 1 | 1 | 1 | 0.00 | | | | | | | | | | | | | | | | | | | | | 0.73 | 1 | 1 | | | | | | | 1648 | 186.4 | 7.91 |
| Q53ET0 | 1.01 | 1 | 1 | 1 | 1 | 1.01 | 1 | 1 | | | | | | | | | | | | | | | | | | | | | | | | | | | | 693 | 73.3 | 7.11 |
| P08637 | 4.33 | 1 | 1 | 1 | 1 | 4.33 | 1 | 1 | | | | | | | | | | | | | | | | | | | | | | | | | | | | 254 | 29.1 | 8.07 |
| P08571 | 5.07 | 1 | 1 | 1 | 1 | 5.07 | 1 | 1 | | | | | | | | | | | | | | | | | | | | | | | | | | | | 375 | 40.1 | 6.23 |
| Q6P087 | 1.99 | 1 | 1 | 1 | 1 | 1.99 | 1 | 1 | | | | | | | | | | | | | | | | | | | | | | | | | 0.00 | | | 351 | 38.4 | 10.32 |
| Q8WUD4 | 4.22 | 1 | 1 | 1 | 1 | 0.00 | | | | | | | | | | | | | | | | | | 4.22 | 1 | 1 | | | | | | | 0.00 | | | 166 | 19.2 | 7.34 |
| Q969T9 | 5.36 | 1 | 1 | 1 | 1 | 5.36 | 1 | 1 | | | | | | | | | | | | | | | | | | | | | | | | | | | | 261 | 28.1 | 5.91 |
| Q9BPW9 | 3.76 | 1 | 1 | 1 | 1 | 3.76 | 1 | 1 | | | | | | | | | | | | | | | | | | | | | | | | | 0.00 | | | 319 | 35.2 | 8.60 |
| P24386 | 1.07 | 2 | 1 | 1 | 1 | 0.00 | | | | | | | | | | | | | | | | | | | | | | | | | | | 1.07 | 1 | 1 | 653 | 73.4 | 4.75 |
| Q6ZMV7 | 1.80 | 1 | 1 | 1 | 1 | 1.80 | 1 | 1 | | | | 0.00 | | | | | | | | | | | | | | | | | | | | | 0.00 | | | 388 | 45.1 | 6.68 |
| P24557 | 1.88 | 1 | 1 | 1 | 1 | 1.88 | 1 | 1 | | | | | | | | | | | | | | | | | | | | | | | | | | | | 533 | 60.5 | 7.62 |
| Q53EZ4 | 3.66 | 1 | 1 | 1 | 1 | 0.00 | | | 0.00 | | | | | | | | | 0.00 | | | | | | | | | | | | | | | 3.66 | 1 | 1 | 464 | 54.1 | 7.01 |
| Q96DD0 | 1.79 | 1 | 1 | 1 | 1 | 1.79 | 1 | 1 | | | | | | | | | | | | | | | | | | | | | | | | | 0.00 | | | 335 | 38.8 | 6.34 |
| Q96IZ5 | 2.91 | 1 | 1 | 1 | 1 | 2.91 | 1 | 1 | | | | | | | | | | | | | | | | | | | | | | | | | 0.00 | | | 413 | 47.1 | 8.70 |
| Q9NPP4 | 0.68 | 1 | 1 | 1 | 1 | 0.68 | 1 | 1 | | | | | | | | | | | | | | | | | | | | | | | | | | | | 1024 | 116.1 | 6.77 |
| Q96BQ1 | 3.57 | 1 | 1 | 1 | 1 | 3.57 | 1 | 1 | | | | | | | | | | | | | | | | | | | | | | | | | | | | 224 | 24.9 | 9.33 |
| Q10588 | 2.83 | 1 | 1 | 1 | 1 | 2.83 | 1 | 1 | | | | | | | | | | | | | | | | | | | | | | | | | | | | 318 | 35.7 | 7.80 |
| B4DYI2 | 0.79 | 7 | 1 | 1 | 1 | 0.00 | | | | | | | | | | | | | | | | | | | | | | | | 0.79 | 1 | 1 | | | | 1134 | 124.4 | 9.29 |
| Q96A32 | 4.14 | 2 | 1 | 1 | 1 | 4.14 | 1 | 1 | | | | | | | | | | | | | | | | | | | | | | | | | | | | 169 | 19.0 | 5.01 |
| Q0VD83 | 1.29 | 1 | 1 | 1 | 1 | 1.29 | 1 | 1 | | | | | | | | | | | | | | | | | | | | | | | | | | | | 1088 | 114.8 | 4.40 |
| P36222 | 3.66 | 1 | 1 | 1 | 1 | 3.66 | 1 | 1 | | | | | | | | | | | | | | | | | | | | | | | | | | | | 383 | 42.6 | 8.46 |
| Q8NGA8 | 2.95 | 3 | 1 | 1 | 1 | 2.95 | 1 | 1 | | | | | | | | | | | | | | | | | | | | | | | | | | | | 305 | 34.2 | 8.38 |
| Q5JRM2 | 2.77 | 1 | 1 | 1 | 1 | 2.77 | 1 | 1 | | | | | | | | | | | | | | | | | | | | | | | | | | | | 361 | 39.9 | 9.54 |
| A6NF34 | 3.65 | 1 | 1 | 1 | 1 | 3.65 | 1 | 1 | | | | | | | | | | | | | | | | | | | | | | | | | | | | 631 | 70.6 | 8.37 |
| Q8N104 | 15.38 | 1 | 1 | 1 | 1 | 15.38 | 1 | 1 | | | | | | | | | | | | | | | | | | | | | | | | | | | | 65 | 7.4 | 8.60 |
| Q9HBL8 | 4.01 | 1 | 1 | 1 | 1 | | | | 0.00 | | | | | | 0.00 | | | | | | | | | | | | | | | | | | 4.01 | 1 | 1 | 299 | 33.3 | 7.52 |
| P11532 | 0.33 | 1 | 1 | 1 | 1 | | | | 0.00 | | | | | | 0.33 | 1 | 1 | | | | | | | | | | 0.00 | | | 0.00 | | | 0.00 | | | 3685 | 426.5 | 5.88 |
| Q9UIF8 | 0.32 | 1 | 1 | 1 | 1 | | | | 0.00 | | | | | | 0.00 | | | 0.00 | | | | | | | | | | | | | | | 0.32 | 1 | 1 | 2168 | 240.3 | 6.54 |
| Q99490 | 0.59 | 1 | 1 | 1 | 1 | | | | 0.00 | | | 0.00 | | | | | | | | | | | | | | | | | | | | | 0.59 | 1 | 1 | 1192 | 124.6 | 9.89 |
| Q92974 | 0.81 | 1 | 1 | 1 | 1 | | | | 0.00 | | | | | | | | | | | | | | | | | | | | | 0.00 | | | 0.81 | 1 | 1 | 986 | 111.5 | 7.27 |
| Q16740 | 5.42 | 1 | 1 | 1 | 1 | | | | 5.42 | 1 | 1 | | | | | | | | | | | | | | | | | | | | | | | | | 277 | 30.2 | 8.09 |
| Q9UKX7 | 3.21 | 1 | 1 | 1 | 1 | | | | 3.21 | 1 | 1 | | | | | | | | | | | | | | | | | | | | | | | | | 468 | 50.1 | 7.06 |
| Q6ZSJ9 | 1.40 | 1 | 1 | 1 | 1 | | | | 1.40 | 1 | 1 | 0.00 | | | | | | | | | | | | | | | | | | | | | 0.00 | | | 500 | 55.7 | 9.35 |
| P19878 | 3.61 | 1 | 1 | 1 | 1 | | | | 0.00 | | | | | | 0.00 | | | 0.00 | | | | | | | | | | | | | | | 3.61 | 1 | 1 | 526 | 59.7 | 6.16 |
| P56381 | 15.69 | 2 | 1 | 1 | 1 | | | | 15.69 | 1 | 1 | | | | | | | | | | | | | | | | | | | | | | | | | 51 | 5.8 | 9.92 |
| Q8NDH3 | 2.29 | 1 | 1 | 1 | 1 | | | | 0.00 | | | | | | | | | | | | | | | | | | | | | | | | 2.29 | 1 | 1 | 523 | 55.8 | 6.87 |
| Q8IWJ2 | 0.48 | 1 | 1 | 1 | 1 | | | | 0.48 | 1 | 1 | | | | | | | | | | | | | | | | | | | | | | | | | 1684 | 195.8 | 5.14 |
| O95359 | 0.47 | 1 | 1 | 1 | 1 | | | | 0.00 | | | | | | | | | | | | | | | | | | | | | 0.47 | 1 | 1 | | | | 2948 | 309.2 | 4.79 |
| Q5FWF5 | 1.07 | 1 | 1 | 1 | 1 | | | | 1.07 | 1 | 1 | | | | | | | | | | | | | | | | | | | | | | | | | 840 | 94.9 | 9.16 |
| Q96PQ6 | 1.68 | 1 | 1 | 1 | 1 | | | | 0.00 | | | 1.68 | 1 | 1 | | | | | | | | | | | | | | | | | | | | | | 595 | 67.9 | 8.98 |
| A8K855 | 1.27 | 1 | 1 | 1 | 1 | | | | 1.27 | 1 | 1 | | | | | | | | | | | | | | | | | | | | | | | | | 629 | 71.9 | 6.38 |
| Q9H1K6 | 3.31 | 1 | 1 | 1 | 1 | | | | 3.31 | 1 | 1 | | | | | | | | | | | | | | | | | | | | | | | | | 362 | 37.7 | 8.15 |
| Q9UL63 | 1.09 | 1 | 1 | 1 | 1 | | | | 1.09 | 1 | 1 | | | | | | | | | | | | | | | | | | | | | | | | | 735 | 84.7 | 6.34 |
| Q11203 | 1.87 | 1 | 1 | 1 | 1 | | | | 1.87 | 1 | 1 | 0.00 | | | | | | | | | | | | | | | | | | | | | | | | 375 | 42.1 | 8.98 |
| A4UGR9 | 0.30 | 1 | 1 | 1 | 1 | | | | 0.00 | | | 0.00 | | | 0.00 | | | 0.00 | | | | | | | | | 0.00 | | | 0.00 | | | 0.30 | 1 | 1 | 3374 | 382.1 | 6.38 |
| Q460N5 | 0.39 | 1 | 1 | 1 | 1 | | | | 0.39 | 1 | 1 | | | | | | | | | | | | | | | | | | | | | | | | | 1801 | 202.7 | 7.18 |
| Q9BRR9 | 1.33 | 1 | 1 | 1 | 1 | | | | 0.00 | | | | | | | | | | | | | | | | | | | | | | | | 1.33 | 1 | 1 | 750 | 83.2 | 8.34 |
| Q96C45 | 0.55 | 1 | 1 | 1 | 1 | | | | 0.00 | | | 0.00 | | | | | | | | | | | | | | | | | | | | | 0.55 | 1 | 1 | 1275 | 142.4 | 6.29 |
| Q96RS0 | 0.82 | 1 | 1 | 1 | 1 | | | | 0.00 | | | | | | | | | | | | | | | | | | | | | | | | 0.82 | 1 | 1 | 853 | 96.6 | 4.94 |
| Q9UPP5 | 0.50 | 1 | 1 | 1 | 1 | | | | 0.50 | 1 | 1 | | | | | | | | | | | | | | | | | | | | | | | | | 1409 | 155.6 | 6.19 |
| Q8N0Z2 | 2.89 | 1 | 1 | 1 | 1 | | | | 2.89 | 1 | 1 | | | | | | | | | | | | | | | | | | | | | | | | | 381 | 43.1 | 8.54 |
| O43897 | 0.79 | 1 | 1 | 1 | 1 | | | | 0.79 | 1 | 1 | | | | | | | | | | | | | | | | | | | | | | | | | 1013 | 114.6 | 6.02 |
| Q96GW7 | 0.66 | 1 | 1 | 1 | 1 | | | | 0.66 | 1 | 1 | | | | | | | | | | | | | | | | | | | | | | | | | 911 | 99.1 | 4.64 |
| Q16595 | 5.24 | 1 | 1 | 1 | 1 | | | | 5.24 | 1 | 1 | | | | | | | | | | | | | | | | | | | | | | | | | 210 | 23.1 | 8.69 |
| Q96BQ5 | 2.69 | 2 | 1 | 1 | 1 | | | | 0.00 | | | | | | | | | | | | | | | | | | | | | | | | 2.69 | 1 | 1 | 260 | 30.8 | 9.20 |
| Q9NTU4 | 5.00 | 1 | 1 | 1 | 1 | | | | 5.00 | 1 | 1 | | | | | | | | | | | | | | | | | | | | | | 0.00 | | | 200 | 22.8 | 6.16 |
| Q9P0U3 | 1.09 | 1 | 1 | 1 | 1 | | | | 1.09 | 1 | 1 | | | | | | | | | | | | | | | | | | | | | | | | | 644 | 73.4 | 8.47 |
| Q96BK5 | 5.49 | 1 | 1 | 1 | 1 | | | | 5.49 | 1 | 1 | | | | | | | | | | | | | | | | | | | | | | | | | 328 | 37.0 | 9.60 |
| Q49SW5 | 2.03 | 1 | 1 | 1 | 1 | | | | 2.03 | 1 | 1 | | | | | | | | | | | | | | | | | | | | | | 0.00 | | | 492 | 55.8 | 5.94 |
| Q9UBB9 | 0.84 | 1 | 1 | 1 | 1 | | | | 0.84 | 1 | 1 | | | | | | | | | | | | | | | | | | | | | | | | | 837 | 96.8 | 5.67 |
| Q6ZNG9 | 1.63 | 1 | 1 | 1 | 1 | | | | 1.63 | 1 | 1 | | | | | | | | | | | | | | | | | | | | | | | | | 492 | 56.2 | 5.97 |
| Q5RHP9 | 0.78 | 1 | 1 | 1 | 1 | | | | 0.78 | 1 | 1 | | | | | | | | | | | | | | | | | | | | | | | | | 1530 | 168.4 | 4.88 |
| Q96HP0 | 0.34 | 2 | 1 | 1 | 1 | | | | 0.00 | | | | | | | | | | | | | | | | | | | | | | | | 0.34 | 1 | 1 | 2047 | 229.4 | 6.74 |
| Q86YT6 | 0.60 | 1 | 1 | 1 | 1 | | | | 0.00 | | | | | | | | | | | | | | | | | | | | | | | | 0.60 | 1 | 1 | 1006 | 110.1 | 6.92 |
| P19793 | 2.16 | 1 | 1 | 1 | 1 | | | | 2.16 | 1 | 1 | | | | | | | | | | | | | | | | | | | | | | | | | 462 | 50.8 | 7.80 |
| Q9NXF7 | 6.48 | 1 | 1 | 1 | 1 | | | | 6.48 | 1 | 1 | | | | | | | | | | | | | | | | | | | | | | | | | 216 | 24.2 | 6.04 |
| Q9Y581 | 3.76 | 1 | 1 | 1 | 1 | | | | 3.76 | 1 | 1 | | | | | | | | | | | | | | | | | | | | | | | | | 213 | 24.8 | 9.67 |
| E9PKD4 | 3.14 | 5 | 1 | 1 | 1 | | | | 3.14 | 1 | 1 | | | | | | | | | | | | | | | | | | | | | | | | | 350 | 40.1 | 9.36 |
| Q3MJ13 | 0.54 | 1 | 1 | 1 | 1 | | | | | | | 0.00 | | | | | | | | | | | | 0.00 | | | | | | | | | 0.54 | 1 | 1 | 1102 | 123.3 | 6.67 |
| P12270 | 0.38 | 1 | 1 | 1 | 1 | | | | | | | 0.00 | | | 0.00 | | | | | | | | | | | | | | | 0.00 | | | 0.38 | 1 | 1 | 2363 | 267.1 | 5.02 |
| Q01130 | 3.62 | 2 | 1 | 1 | 1 | | | | | | | 3.62 | 1 | 1 | | | | | | | | | | | | | | | | | | | | | | 221 | 25.5 | 11.85 |
| CON_008274691 | 7.29 | 1 | 1 | 1 | 1 | | | | | | | 7.29 | 1 | 1 | | | | | | | | | | | | | | | | | | | | | | 192 | 20.9 | 5.02 |
| P13995 | 2.00 | 1 | 1 | 1 | 1 | | | | | | | 2.00 | 1 | 1 | | | | | | | | | | | | | | | | | | | | | | 350 | 37.9 | 8.73 |
| CON_007078841 | 2.61 | 2 | 1 | 1 | 1 | | | | | | | 2.61 | 1 | 1 | | | | | | | | | | | | | | | | | | | | | | 345 | 38.2 | 8.15 |
| Q9NY91 | 0.91 | 1 | 1 | 1 | 1 | | | | | | | 0.91 | 1 | 1 | | | | | | | | | | | | | | | | | | | 0.00 | | | 659 | 72.4 | 6.15 |
| P20701 | 1.62 | 1 | 1 | 1 | 1 | | | | | | | 1.62 | 1 | 1 | | | | | | | | | | | | | | | | | | | | | | 1170 | 128.7 | 5.63 |
| Q9BXT2 | 2.69 | 1 | 1 | 1 | 1 | | | | | | | 0.00 | | | | | | | | | | | | | | | | | | | | | 2.69 | 1 | 1 | 260 | 28.1 | 9.17 |
| Q9NYP7 | 4.35 | 1 | 1 | 1 | 1 | | | | | | | 4.35 | 1 | 1 | | | | | | | | | | | | | | | | | | | | | | 299 | 35.3 | 9.42 |
| O95613 | 0.18 | 1 | 1 | 1 | 1 | | | | | | | 0.00 | | | | | | 0.00 | | | | | | | | | | | | | | | 0.18 | 1 | 1 | 3336 | 377.8 | 5.55 |
| Q9NVU7 | 1.02 | 1 | 1 | 1 | 1 | | | | | | | 1.02 | 1 | 1 | | | | | | | | | | | | | | | | | | | | | | 687 | 79.8 | 9.25 |
| Q92692 | 1.86 | 1 | 1 | 1 | 1 | | | | | | | 1.86 | 1 | 1 | | | | | | | | | | | | | | | | | | | 0.00 | | | 538 | 57.7 | 4.82 |
| Q6NUS8 | 1.91 | 1 | 1 | 1 | 1 | | | | | | | 1.91 | 1 | 1 | | | | | | | | | | | | | | | | | | | | | | 523 | 59.1 | 8.16 |
| Q9Y336 | 3.89 | 1 | 1 | 1 | 1 | | | | | | | 3.89 | 1 | 1 | | | | | | | | | | | | | | | | | | | | | | 463 | 50.1 | 7.25 |
| Q05823 | 1.08 | 1 | 1 | 1 | 1 | | | | | | | 1.08 | 1 | 1 | | | | | | | | | | | | | | | | | | | | | | 741 | 83.5 | 6.65 |
| Q99797 | 1.12 | 1 | 1 | 1 | 1 | | | | | | | 0.00 | | | 0.00 | | | 0.00 | | | | | | 0.00 | | | 0.00 | | | 0.00 | | | 1.12 | 1 | 1 | 713 | 80.6 | 7.05 |
| Q9C0E8 | 2.57 | 1 | 1 | 1 | 1 | | | | | | | 2.57 | 1 | 1 | | | | | | | | | | | | | | | | | | | | | | 428 | 47.7 | 5.11 |
| Q9NSI2 | 4.35 | 1 | 1 | 1 | 1 | | | | | | | 4.35 | 1 | 1 | | | | | | | | | | | | | | | | | | | | | | 230 | 25.4 | 11.08 |
| P00751 | 1.83 | 1 | 1 | 1 | 1 | | | | | | | 1.83 | 1 | 1 | | | | | | | | | | | | | | | | | | | | | | 764 | 85.5 | 7.06 |
| Q96JN8 | 0.70 | 1 | 1 | 1 | 1 | | | | | | | 0.00 | | | | | | | | | | | | | | | | | | | | | 0.70 | 1 | 1 | 1562 | 166.8 | 5.86 |
| Q8N5F7 | 1.69 | 1 | 1 | 1 | 1 | | | | | | | 0.00 | | | 1.69 | 1 | 1 | | | | | | | 0.00 | | | | | | | | | 0.00 | | | 415 | 47.1 | 10.11 |
| Q6YHU6 | 0.51 | 1 | 1 | 1 | 1 | | | | | | | 0.00 | | | | | | | | | | | | | | | | | | | | | 0.51 | 1 | 1 | 1953 | 219.5 | 6.06 |
| Q9BUA6 | 3.10 | 1 | 1 | 1 | 1 | | | | | | | 3.10 | 1 | 1 | | | | | | | | | | | | | | | | | | | | | | 226 | 25.3 | 5.73 |
| Q8TDW0 | 1.00 | 1 | 1 | 1 | 1 | | | | | | | 1.00 | 1 | 1 | | | | | | | | | | | | | | | | | | | | | | 803 | 92.4 | 7.62 |
| P28336 | 3.08 | 1 | 1 | 1 | 1 | | | | | | | 3.08 | 1 | 1 | | | | | | | | | | | | | | | | | | | | | | 390 | 43.4 | 8.66 |
| P23141 | 1.94 | 1 | 1 | 1 | 1 | | | | | | | 1.94 | 1 | 1 | | | | | | | | | | | | | | | | | | | | | | 567 | 62.5 | 6.60 |
| O75015 | 6.44 | 1 | 1 | 1 | 1 | | | | | | | 6.44 | 1 | 1 | | | | | | | | | | | | | | | | | | | | | | 233 | 26.2 | 6.71 |
| Q9BY44 | 3.42 | 1 | 1 | 1 | 1 | | | | | | | | | | 0.00 | | | 0.00 | | | | | | | | | | | | | | | 3.42 | 1 | 1 | 585 | 64.9 | 8.87 |
| Q8N6R0 | 1.14 | 1 | 1 | 1 | 1 | | | | | | | | | | 0.00 | | | | | | | | | | | | | | | | | | 1.14 | 1 | 1 | 699 | 78.7 | 6.73 |
| CON_007089593 | 1.85 | 1 | 1 | 1 | 1 | | | | | | | | | | 1.85 | 1 | 1 | | | | | | | | | | | | | | | | | | | 433 | 49.3 | 6.60 |
| O75506 | 14.47 | 1 | 1 | 1 | 1 | | | | | | | | | | 14.47 | 1 | 1 | | | | | | | | | | | | | | | | | | | 76 | 8.5 | 4.36 |
| CON_007152872 | 1.59 | 2 | 1 | 1 | 1 | | | | | | | | | | 1.59 | 1 | 1 | | | | | | | | | | | | | | | | | | | 946 | 106.1 | 7.94 |
| Q08345 | 0.88 | 1 | 1 | 1 | 1 | | | | | | | | | | 0.88 | 1 | 1 | | | | | | | | | | | | | | | | 0.00 | | | 913 | 101.1 | 6.83 |
| Q8TD16 | 1.09 | 1 | 1 | 1 | 1 | | | | | | | | | | 1.09 | 1 | 1 | | | | | | | | | | | | | | | | 0.00 | | | 824 | 93.5 | 5.44 |
| Q15063 | 0.96 | 1 | 1 | 1 | 1 | | | | | | | | | | 0.96 | 1 | 1 | | | | | | | | | | | | | | | | | | | 836 | 93.3 | 7.53 |
| Q7Z2Y8 | 0.33 | 1 | 1 | 1 | 1 | | | | | | | | | | 0.00 | | | | | | | | | | | | | | | | | | 0.33 | 1 | 1 | 2422 | 278.9 | 6.55 |
| O75943 | 1.03 | 1 | 1 | 1 | 1 | | | | | | | | | | 1.03 | 1 | 1 | | | | | | | | | | | | | | | | | | | 681 | 77.0 | 7.06 |
| Q13506 | 1.44 | 1 | 1 | 1 | 1 | | | | | | | | | | 1.44 | 1 | 1 | | | | | | | | | | | | | | | | | | | 487 | 54.4 | 6.70 |
| Q9BW92 | 1.11 | 1 | 1 | 1 | 1 | | | | | | | | | | 0.00 | | | | | | | | | | | | | | | 0.00 | | | 1.11 | 1 | 1 | 718 | 81.0 | 7.30 |
| Q96A37 | 2.95 | 1 | 1 | 1 | 1 | | | | | | | | | | 2.95 | 1 | 1 | | | | | | | | | | | | | | | | | | | 237 | 26.1 | 8.06 |
| O95140 | 0.92 | 1 | 1 | 1 | 1 | | | | | | | | | | 0.92 | 1 | 1 | | | | | | | | | | | | | | | | | | | 757 | 86.3 | 6.98 |
| O00472 | 1.09 | 1 | 1 | 1 | 1 | | | | | | | | | | 0.00 | | | | | | | | | | | | | | | | | | 1.09 | 1 | 1 | 640 | 72.3 | 9.00 |
| O95996 | 1.35 | 1 | 1 | 1 | 1 | | | | | | | | | | 0.00 | | | 0.00 | | | | | | | | | | | | | | | 1.35 | 1 | 1 | 2303 | 243.8 | 8.82 |
| O94916 | 0.72 | 1 | 1 | 1 | 1 | | | | | | | | | | 0.00 | | | | | | | | | | | | | | | | | | 0.72 | 1 | 1 | 1531 | 165.7 | 5.24 |
| Q96HU1 | 1.47 | 1 | 1 | 1 | 1 | | | | | | | | | | 0.00 | | | | | | | | | | | | | | | | | | 1.47 | 1 | 1 | 749 | 85.3 | 6.00 |
| Q5T2T1 | 1.39 | 1 | 1 | 1 | 1 | | | | | | | | | | 1.39 | 1 | 1 | | | | | | | | | | | | | | | | | | | 576 | 65.5 | 7.14 |
| Q96JB8 | 1.10 | 1 | 1 | 1 | 1 | | | | | | | | | | 1.10 | 1 | 1 | | | | | | | | | | | | | | | | 0.00 | | | 637 | 72.7 | 5.62 |
| Q86SQ4 | 0.49 | 1 | 1 | 1 | 1 | | | | | | | | | | 0.49 | 1 | 1 | | | | | | | | | | | | | | | | | | | 1221 | 136.6 | 7.87 |
| P55017 | 0.88 | 1 | 1 | 1 | 1 | | | | | | | | | | 0.88 | 1 | 1 | | | | | | | | | | | | | | | | | | | 1021 | 113.1 | 7.88 |
| Q8NGJ9 | 2.45 | 1 | 1 | 1 | 1 | | | | | | | | | | 2.45 | 1 | 1 | | | | | | | | | | | | | | | | | | | 327 | 37.0 | 9.29 |
| P49915 | 1.30 | 1 | 1 | 1 | 1 | | | | | | | | | | | | | 0.00 | | | | | | | | | | | | | | | 1.30 | 1 | 1 | 693 | 76.7 | 6.87 |
| Q9NU22 | 0.16 | 1 | 1 | 1 | 1 | | | | | | | | | | | | | 0.00 | | | | | | | | | | | | 0.16 | 1 | 1 | | | | 5596 | 632.4 | 5.68 |
| Q8WWI1 | 0.48 | 1 | 1 | 1 | 1 | | | | | | | | | | | | | 0.00 | | | | | | | | | | | | | | | 0.48 | 1 | 1 | 1683 | 192.6 | 8.09 |
| Q7Z460 | 0.72 | 1 | 1 | 1 | 1 | | | | | | | | | | | | | 0.00 | | | | | | | | | | | | | | | 0.72 | 1 | 1 | 1538 | 169.3 | 9.03 |
| Q9H6N6 | 0.55 | 3 | 1 | 1 | 1 | | | | | | | | | | | | | 0.00 | | | | | | | | | 0.55 | 1 | 1 | 0.00 | | | 0.00 | | | 1097 | 128.2 | 5.49 |
| A4D2H0 | 1.29 | 5 | 1 | 1 | 1 | | | | | | | | | | | | | 1.29 | 1 | 1 | | | | | | | | | | | | | | | | 777 | 87.8 | 5.24 |
| Q6PKH6 | 3.04 | 1 | 1 | 1 | 1 | | | | | | | | | | | | | 3.04 | 1 | 1 | | | | | | | | | | | | | | | | 230 | 24.6 | 10.11 |
| Q6WKZ4 | 0.55 | 1 | 1 | 1 | 1 | | | | | | | | | | | | | 0.55 | 1 | 1 | | | | | | | | | | | | | | | | 1283 | 137.1 | 5.43 |
| Q08AF8 | 1.40 | 3 | 1 | 1 | 1 | | | | | | | | | | | | | 0.00 | | | | | | 0.00 | | | 0.00 | | | 0.00 | | | 1.40 | 1 | 1 | 430 | 48.4 | 5.57 |
| P26022 | 2.62 | 1 | 1 | 1 | 1 | | | | | | | | | | | | | 2.62 | 1 | 1 | | | | | | | | | | | | | | | | 381 | 41.9 | 5.01 |
| Q2VPJ9 | 2.22 | 1 | 1 | 1 | 1 | | | | | | | | | | | | | 2.22 | 1 | 1 | | | | | | | | | | | | | | | | 315 | 34.7 | 9.31 |
| Q9NWZ3 | 1.52 | 1 | 1 | 1 | 1 | | | | | | | | | | | | | 0.00 | | | | | | 0.00 | | | | | | 0.00 | | | 1.52 | 1 | 1 | 460 | 51.5 | 5.41 |
| O94915 | 0.27 | 1 | 1 | 1 | 1 | | | | | | | | | | | | | 0.00 | | | | | | | | | | | | 0.00 | | | 0.27 | 1 | 1 | 3013 | 339.4 | 5.58 |
| Q9P1Y5 | 0.56 | 1 | 1 | 1 | 1 | | | | | | | | | | | | | 0.00 | | | | | | | | | | | | | | | 0.56 | 1 | 1 | 1249 | 134.7 | 8.35 |
| Q9H5V9 | 4.95 | 1 | 1 | 1 | 1 | | | | | | | | | | | | | 4.95 | 1 | 1 | | | | | | | | | | | | | | | | 222 | 25.6 | 8.73 |
| Q8IZX4 | 0.38 | 1 | 1 | 1 | 1 | | | | | | | | | | | | | 0.00 | | | | | | 0.00 | | | 0.00 | | | | | | 0.38 | 1 | 1 | 1826 | 207.2 | 5.40 |
| Q6NXP0 | 1.75 | 1 | 1 | 1 | 1 | | | | | | | | | | | | | 1.75 | 1 | 1 | | | | | | | | | | | | | | | | 572 | 66.5 | 9.47 |
| Q8IVF7 | 0.58 | 2 | 1 | 1 | 1 | | | | | | | | | | | | | 0.00 | | | | | | | | | | | | | | | 0.58 | 1 | 1 | 1028 | 117.1 | 6.65 |

| | | | | | | | | | | | | | | | | | | | | | | | | | | |
|---|---|---|---|---|---|---|---|---|---|---|---|---|---|---|---|---|---|---|---|---|---|---|---|---|---|---|
| Q8NGC5 | 2.02 | 1 | 1 | 1 | 1 | 2.02 | 1 | 1 | | | | | | | | | | | | | | | | 347 | 38.7 | 8.47 |
| Q9P2Q2 | 0.58 | 1 | 1 | 1 | 1 | 0.58 | 1 | 1 | | | | | | | | | | | | | 0.00 | | | 1039 | 115.4 | 8.87 |
| P63136 | 0.84 | 5 | 1 | 1 | 1 | 0.84 | 1 | 1 | | | | | | | | | | | | | | | | 954 | 107.4 | 9.00 |
| Q86UB2 | 6.16 | 1 | 1 | 1 | 1 | 6.16 | 1 | 1 | | | | | | | | | | | | | | | | 503 | 56.7 | 8.98 |
| Q86XI6 | 2.46 | 1 | 1 | 1 | 1 | 2.46 | 1 | 1 | | | | | | | | | | | | | | | | 285 | 32.7 | 6.14 |
| Q86V48 | 0.84 | 1 | 1 | 1 | 1 | | | | 0.00 | | | | | | | | | | | | 0.84 | 1 | 1 | 1076 | 120.2 | 8.50 |
| Q9NQ50 | 3.40 | 1 | 1 | 1 | 1 | | | | 3.40 | 1 | 1 | | | | | | | | | | | | | 206 | 24.5 | 9.63 |
| Q03518 | 1.36 | 1 | 1 | 1 | 1 | | | | 0.00 | | | | | | | | | | | | 1.36 | 1 | 1 | 808 | 87.2 | 8.02 |
| Q9H788 | 3.74 | 1 | 1 | 1 | 1 | | | | 0.00 | | | | | | | | | | | | 3.74 | 1 | 1 | 454 | 52.7 | 8.06 |
| Q6V0I7 | 0.26 | 1 | 1 | 1 | 1 | | | | 0.26 | 1 | 1 | | | | | | | | | | 0.00 | | | 4981 | 542.4 | 4.94 |
| Q7Z5M8 | 1.66 | 1 | 1 | 1 | 1 | | | | 1.66 | 1 | 1 | | | | | | | | | | | | | 362 | 40.7 | 8.28 |
| Q5VX52 | 1.83 | 1 | 1 | 1 | 1 | | | | | | | 0.00 | | | 0.00 | | | | | | 1.83 | 1 | 1 | 437 | 50.3 | 8.43 |
| Q9NZN1 | 1.01 | 1 | 1 | 1 | 1 | | | | | | | 0.00 | | | 1.01 | 1 | 1 | | | | | | | 696 | 79.9 | 6.23 |
| Q96MW1 | 4.46 | 1 | 1 | 1 | 1 | | | | | | | 4.46 | 1 | 1 | | | | | | | | | | 224 | 25.2 | 4.92 |
| Q9Y6R1 | 0.65 | 1 | 1 | 1 | 1 | | | | | | | 0.65 | 1 | 1 | | | | | | | | | | 1079 | 121.4 | 6.80 |
| P08F94 | 0.27 | 1 | 1 | 1 | 1 | | | | | | | 0.27 | 1 | 1 | | | | | | | | | | 4074 | 446.4 | 6.57 |
| Q8N302 | 0.84 | 1 | 1 | 1 | 1 | | | | | | | 0.00 | | | | | | | | | 0.84 | 1 | 1 | 714 | 80.9 | 5.44 |
| P07476 | 3.42 | 1 | 1 | 1 | 1 | | | | | | | 3.42 | 1 | 1 | | | | | | | | | | 585 | 68.4 | 4.61 |
| Q9Y6G3 | 4.93 | 1 | 1 | 1 | 1 | | | | | | | 4.93 | 1 | 1 | | | | | | | | | | 142 | 16.7 | 8.35 |
| P49790 | 0.88 | 1 | 1 | 1 | 1 | | | | | | | | | | 0.88 | 1 | 1 | | | | 0.00 | | | 1475 | 153.8 | 8.73 |
| O14979 | 3.81 | 1 | 1 | 1 | 1 | | | | | | | | | | 3.81 | 1 | 1 | | | | 0.00 | | | 420 | 46.4 | 9.57 |
| Q9ULD6 | 0.74 | 1 | 1 | 1 | 1 | | | | | | | | | | 0.74 | 1 | 1 | | | | | | | 942 | 105.6 | 5.90 |
| Q96LI9 | 1.81 | 1 | 1 | 1 | 1 | | | | | | | | | | 1.81 | 1 | 1 | | | | | | | 332 | 38.9 | 10.18 |
| Q9NRL2 | 0.51 | 1 | 1 | 1 | 1 | | | | | | | | | | 0.00 | | | 0.00 | | | 0.51 | 1 | 1 | 1556 | 178.6 | 6.60 |
| Q15406 | 1.25 | 2 | 1 | 1 | 1 | | | | | | | | | | 1.25 | 1 | 1 | | | | | | | 480 | 54.3 | 6.25 |
| P08100 | 4.02 | 1 | 1 | 1 | 1 | | | | | | | | | | 4.02 | 1 | 1 | | | | | | | 348 | 38.9 | 6.65 |
| O15061 | 0.64 | 1 | 1 | 1 | 1 | | | | | | | | | | 0.00 | | | | | | 0.64 | 1 | 1 | 1565 | 172.7 | 5.16 |
| Q9P219 | 0.54 | 1 | 1 | 1 | 1 | | | | | | | | | | 0.00 | | | | | | 0.54 | 1 | 1 | 2028 | 228.1 | 6.23 |
| Q96PM5 | 5.36 | 1 | 1 | 1 | 1 | | | | | | | | | | 5.36 | 1 | 1 | | | | | | | 261 | 30.1 | 6.71 |
| P21217 | 2.49 | 2 | 1 | 1 | 1 | | | | | | | | | | 2.49 | 1 | 1 | | | | | | | 361 | 42.1 | 9.01 |
| Q9BYV6 | 2.01 | 1 | 1 | 1 | 1 | | | | | | | | | | 2.01 | 1 | 1 | | | | | | | 548 | 60.4 | 4.77 |
| P13010 | 1.91 | 1 | 1 | 1 | 1 | | | | | | | | | | | | | 1.91 | 1 | 1 | | | | 732 | 82.7 | 5.81 |
| P15884 | 1.05 | 1 | 1 | 1 | 1 | | | | | | | | | | | | | 1.05 | 1 | 1 | 0.00 | | | 667 | 71.3 | 7.01 |
| O14907 | 13.71 | 1 | 1 | 1 | 1 | | | | | | | | | | | | | 13.71 | 1 | 1 | | | | 124 | 13.7 | 8.48 |
| Q8IZP0 | 1.77 | 2 | 1 | 1 | 1 | | | | | | | | | | | | | 1.77 | 1 | 1 | | | | 508 | 55.0 | 7.06 |
| Q9NZJ4 | 0.24 | 1 | 1 | 1 | 1 | | | | | | | | | | | | | 0.24 | 1 | 1 | 0.00 | | | 4579 | 520.8 | 7.05 |
| Q96RT7 | 0.38 | 1 | 1 | 1 | 1 | | | | | | | | | | | | | 0.38 | 1 | 1 | | | | 1819 | 200.4 | 6.32 |
| Q9Y6Y8 | 1.30 | 1 | 1 | 1 | 1 | | | | | | | | | | | | | 1.30 | 1 | 1 | | | | 1000 | 111.0 | 5.54 |
| Q96FV9 | 0.91 | 2 | 1 | 1 | 1 | | | | | | | | | | | | | 0.91 | 1 | 1 | | | | 657 | 75.6 | 4.98 |
| O14746 | 0.53 | 1 | 1 | 1 | 1 | | | | | | | | | | | | | 0.53 | 1 | 1 | 0.00 | | | 1132 | 126.9 | 10.52 |
| Q8WZA9 | 2.25 | 1 | 1 | 1 | 1 | | | | | | | | | | | | | 2.25 | 1 | 1 | | | | 623 | 62.7 | 4.88 |
| O60925 | 9.02 | 1 | 1 | 1 | 1 | | | | | | | | | | | | | 9.02 | 1 | 1 | | | | 122 | 14.2 | 6.81 |
| Q12830 | 0.23 | 1 | 1 | 1 | 1 | | | | | | | | | | | | | 0.23 | 1 | 1 | 0.00 | | | 3046 | 338.1 | 6.54 |
| Q8N7Z5 | 0.59 | 1 | 1 | 1 | 1 | | | | | | | | | | | | | 0.59 | 1 | 1 | 0.00 | | | 1873 | 210.7 | 6.20 |
| Q6V1X1 | 1.34 | 1 | 1 | 1 | 1 | | | | | | | | | | | | | 1.34 | 1 | 1 | 0.00 | | | 898 | 103.3 | 5.80 |
| Q70CQ1 | 1.02 | 1 | 1 | 1 | 1 | | | | | | | | | | | | | 1.02 | 1 | 1 | 0.00 | | | 688 | 79.1 | 9.19 |
| Q7Z4V0 | 0.97 | 1 | 1 | 1 | 1 | | | | | | | | | | | | | 0.97 | 1 | 1 | | | | 828 | 91.8 | 9.39 |
| Q2M1Z3 | 0.48 | 1 | 1 | 1 | 1 | | | | | | | | | | | | | 0.48 | 1 | 1 | | | | 1444 | 156.9 | 5.76 |
| Q2M3G4 | 2.00 | 1 | 1 | 1 | 1 | | | | | | | | | | | | | 2.00 | 1 | 1 | | | | 852 | 90.7 | 6.15 |
| Q6IQ55 | 0.72 | 1 | 1 | 1 | 1 | | | | | | | | | | | | | 0.00 | | | 0.72 | 1 | 1 | 1244 | 137.3 | 7.01 |
| Q6P1X5 | 1.92 | 1 | 1 | 1 | 1 | | | | | | | | | | | | | 1.92 | 1 | 1 | 0.00 | | | 1199 | 136.9 | 8.19 |
| Q8NFH8 | 1.52 | 2 | 1 | 1 | 1 | | | | | | | | | | | | | 1.52 | 1 | 1 | | | | 660 | 71.5 | 7.65 |
| Q8N4S9 | 1.79 | 1 | 1 | 1 | 1 | | | | | | | | | | | | | 1.79 | 1 | 1 | | | | 558 | 64.1 | 7.53 |
| P51948 | 3.24 | 1 | 1 | 1 | 1 | | | | | | | | | | | | | 3.24 | 1 | 1 | | | | 309 | 35.8 | 6.09 |
| Q4V348 | 2.20 | 2 | 1 | 1 | 1 | | | | | | | | | | | | | 2.20 | 1 | 1 | | | | 819 | 94.3 | 8.56 |
| O43663 | 1.77 | 1 | 1 | 1 | 1 | | | | | | | | | | | | | | | | 1.77 | 1 | 1 | 620 | 71.6 | 6.68 |
| Q9ULL1 | 0.43 | 1 | 1 | 1 | 1 | | | | | | | | | | | | | | | | 0.43 | 1 | 1 | 1385 | 155.3 | 6.24 |
| P04114 | 0.24 | 1 | 1 | 1 | 1 | | | | | | | | | | | | | | | | 0.24 | 1 | 1 | 4563 | 515.3 | 7.05 |
| Q8TDB6 | 2.16 | 1 | 1 | 1 | 1 | | | | | | | | | | | | | | | | 2.16 | 1 | 1 | 740 | 83.5 | 8.06 |
| Q92830 | 0.72 | 1 | 1 | 1 | 1 | | | | | | | | | | | | | | | | 0.72 | 1 | 1 | 837 | 93.9 | 9.04 |
| Q13573 | 1.49 | 1 | 1 | 1 | 1 | | | | | | | | | | | | | | | | 1.49 | 1 | 1 | 536 | 61.5 | 9.52 |
| O95571 | 3.54 | 1 | 1 | 1 | 1 | | | | | | | | | | | | | | | | 3.54 | 1 | 1 | 254 | 27.9 | 6.83 |
| Q6VY07 | 0.93 | 1 | 1 | 1 | 1 | | | | | | | | | | | | | | | | 0.93 | 1 | 1 | 963 | 104.8 | 7.74 |
| P12004 | 2.68 | 1 | 1 | 1 | 1 | | | | | | | | | | | | | | | | 2.68 | 1 | 1 | 261 | 28.8 | 4.69 |
| Q96J66 | 1.01 | 1 | 1 | 1 | 1 | | | | | | | | | | | | | | | | 1.01 | 1 | 1 | 1382 | 154.2 | 8.09 |
| Q8NBJ5 | 1.61 | 1 | 1 | 1 | 1 | | | | | | | | | | | | | | | | 1.61 | 1 | 1 | 622 | 71.6 | 7.31 |
| Q6P2Q9 | 0.64 | 1 | 1 | 1 | 1 | | | | | | | | | | | | | | | | 0.64 | 1 | 1 | 2335 | 273.4 | 8.84 |
| Q9UJX2 | 1.68 | 1 | 1 | 1 | 1 | | | | | | | | | | | | | | | | 1.68 | 1 | 1 | 597 | 68.8 | 7.02 |
| P01024 | 0.42 | 1 | 1 | 1 | 1 | | | | | | | | | | | | | | | | 0.42 | 1 | 1 | 1663 | 187.0 | 6.40 |
| Q8NC51 | 3.92 | 1 | 1 | 1 | 1 | | | | | | | | | | | | | | | | 3.92 | 1 | 1 | 408 | 44.9 | 8.65 |
| P08579 | 4.89 | 1 | 1 | 1 | 1 | | | | | | | | | | | | | | | | 4.89 | 1 | 1 | 225 | 25.5 | 9.72 |
| Q9UNL4 | 2.81 | 1 | 1 | 1 | 1 | | | | | | | | | | | | | | | | 2.81 | 1 | 1 | 249 | 28.5 | 7.62 |
| Q8NB16 | 1.49 | 1 | 1 | 1 | 1 | | | | | | | | | | | | | | | | 1.49 | 1 | 1 | 471 | 54.4 | 8.82 |
| Q99543 | 2.09 | 1 | 1 | 1 | 1 | | | | | | | | | | | | | | | | 2.09 | 1 | 1 | 621 | 72.0 | 8.70 |
| P35659 | 1.87 | 1 | 1 | 1 | 1 | | | | | | | | | | | | | | | | 1.87 | 1 | 1 | 375 | 42.6 | 8.56 |
| Q14683 | 0.57 | 1 | 1 | 1 | 1 | | | | | | | | | | | | | | | | 0.57 | 1 | 1 | 1233 | 143.1 | 7.64 |
| Q6XQN6 | 1.86 | 1 | 1 | 1 | 1 | | | | | | | | | | | | | | | | 1.86 | 1 | 1 | 538 | 57.5 | 5.68 |
| Q5T5U3 | 0.36 | 1 | 1 | 1 | 1 | | | | | | | | | | | | | | | | 0.36 | 1 | 1 | 1957 | 217.2 | 7.80 |
| P49959 | 0.85 | 1 | 1 | 1 | 1 | | | | | | | | | | | | | | | | 0.85 | 1 | 1 | 708 | 80.5 | 5.90 |
| Q15650 | 1.55 | 1 | 1 | 1 | 1 | | | | | | | | | | | | | | | | 1.55 | 1 | 1 | 581 | 66.1 | 7.85 |
| Q12955 | 0.16 | 1 | 1 | 1 | 1 | | | | | | | | | | | | | | | | 0.16 | 1 | 1 | 4377 | 480.1 | 6.49 |
| Q9UKK3 | 0.58 | 1 | 1 | 1 | 1 | | | | | | | | | | | | | | | | 0.58 | 1 | 1 | 1724 | 192.5 | 5.66 |
| Q0VF96 | 0.54 | 1 | 1 | 1 | 1 | | | | | | | | | | | | | | | | 0.54 | 1 | 1 | 1302 | 149.0 | 5.67 |
| Q8IYM0 | 0.90 | 1 | 1 | 1 | 1 | | | | | | | | | | | | | | | | 0.90 | 1 | 1 | 893 | 103.7 | 9.01 |
| Q16204 | 2.95 | 1 | 1 | 1 | 1 | | | | | | | | | | | | | | | | 2.95 | 1 | 1 | 474 | 53.3 | 7.34 |
| Q5TGZ0 | 10.26 | 1 | 1 | 1 | 1 | | | | | | | | | | | | | | | | 10.26 | 1 | 1 | 78 | 8.8 | 8.50 |
| Q96C01 | 7.25 | 1 | 1 | 1 | 1 | | | | | | | | | | | | | | | | 7.25 | 1 | 1 | 138 | 15.6 | 7.61 |
| O95347 | 0.84 | 1 | 1 | 1 | 1 | | | | | | | | | | | | | | | | 0.84 | 1 | 1 | 1197 | 135.6 | 8.43 |
| Q9HAV4 | 1.66 | 1 | 1 | 1 | 1 | | | | | | | | | | | | | | | | 1.66 | 1 | 1 | 1204 | 136.2 | 5.80 |
| Q14318 | 3.40 | 1 | 1 | 1 | 1 | | | | | | | | | | | | | | | | 3.40 | 1 | 1 | 412 | 44.5 | 4.84 |
| P04181 | 3.64 | 1 | 1 | 1 | 1 | | | | | | | | | | | | | | | | 3.64 | 1 | 1 | 439 | 48.5 | 7.03 |
| O75526 | 3.32 | 1 | 1 | 1 | 1 | | | | | | | | | | | | | | | | 3.32 | 1 | 1 | 392 | 42.8 | 10.32 |
| Q9BZX2 | 3.07 | 2 | 1 | 1 | 1 | | | | | | | | | | | | | | | | 3.07 | 1 | 1 | 261 | 29.3 | 6.70 |
| O00139 | 1.56 | 1 | 1 | 1 | 1 | | | | | | | | | | | | | | | | 1.56 | 1 | 1 | 706 | 79.9 | 6.68 |
| Q16719 | 3.44 | 1 | 1 | 1 | 1 | | | | | | | | | | | | | | | | 3.44 | 1 | 1 | 465 | 52.3 | 7.03 |
| Q9BWU0 | 1.63 | 1 | 1 | 1 | 1 | | | | | | | | | | | | | | | | 1.63 | 1 | 1 | 796 | 88.8 | 5.19 |
| Q12797 | 1.85 | 1 | 1 | 1 | 1 | | | | | | | | | | | | | | | | 1.85 | 1 | 1 | 758 | 85.8 | 5.01 |
| Q7Z478 | 1.24 | 1 | 1 | 1 | 1 | | | | | | | | | | | | | | | | 1.24 | 1 | 1 | 1369 | 155.1 | 8.09 |
| CON_007100251 | 1.50 | 1 | 1 | 1 | 1 | | | | | | | | | | | | | | | | 1.50 | 1 | 1 | 468 | 51.7 | 6.67 |
| O43447 | 7.91 | 1 | 1 | 1 | 1 | | | | | | | | | | | | | | | | 7.91 | 1 | 1 | 177 | 19.2 | 8.07 |
| O43325 | 6.56 | 1 | 1 | 1 | 1 | | | | | | | | | | | | | | | | 6.56 | 1 | 1 | 122 | 14.3 | 9.94 |
| Q12905 | 5.13 | 1 | 1 | 1 | 1 | | | | | | | | | | | | | | | | 5.13 | 1 | 1 | 390 | 43.0 | 5.26 |
| P62306 | 15.12 | 1 | 1 | 1 | 1 | | | | | | | | | | | | | | | | 15.12 | 1 | 1 | 86 | 9.7 | 4.67 |
| Q5BJF2 | 4.55 | 1 | 1 | 1 | 1 | | | | | | | | | | | | | | | | 4.55 | 1 | 1 | 176 | 20.8 | 9.38 |
| Q9UPY8 | 2.85 | 1 | 1 | 1 | 1 | | | | | | | | | | | | | | | | 2.85 | 1 | 1 | 281 | 32.0 | 5.54 |
| Q8N999 | 3.08 | 1 | 1 | 1 | 1 | | | | | | | | | | | | | | | | 3.08 | 1 | 1 | 325 | 37.5 | 7.06 |
| O00625 | 4.14 | 1 | 1 | 1 | 1 | | | | | | | | | | | | | | | | 4.14 | 1 | 1 | 290 | 32.1 | 6.92 |
| Q96KR1 | 0.65 | 1 | 1 | 1 | 1 | | | | | | | | | | | | | | | | 0.65 | 1 | 1 | 1074 | 116.9 | 9.04 |
| Q9GZT3 | 9.17 | 1 | 1 | 1 | 1 | | | | | | | | | | | | | | | | 9.17 | 1 | 1 | 109 | 12.3 | 10.24 |
| Q8WUY1 | 3.85 | 1 | 1 | 1 | 1 | | | | | | | | | | | | | | | | 3.85 | 1 | 1 | 208 | 23.8 | 9.55 |
| O15355 | 5.31 | 1 | 1 | 1 | 1 | | | | | | | | | | | | | | | | 5.31 | 1 | 1 | 546 | 59.2 | 4.36 |
| Q9Y2Q9 | 5.35 | 1 | 1 | 1 | 1 | | | | | | | | | | | | | | | | 5.35 | 1 | 1 | 187 | 20.8 | 9.10 |
| Q8NBN7 | 7.25 | 1 | 1 | 1 | 1 | | | | | | | | | | | | | | | | 7.25 | 1 | 1 | 331 | 35.9 | 8.10 |
| O75027 | 1.33 | 1 | 1 | 1 | 1 | | | | | | | | | | | | | | | | 1.33 | 1 | 1 | 752 | 82.6 | 9.33 |
| P61011 | 2.78 | 1 | 1 | 1 | 1 | | | | | | | | | | | | | | | | 2.78 | 1 | 1 | 504 | 55.7 | 8.75 |
| Q9BZD4 | 1.51 | 1 | 1 | 1 | 1 | | | | | | | | | | | | | | | | 1.51 | 1 | 1 | 464 | 54.3 | 8.27 |
| Q9BQE5 | 3.26 | 1 | 1 | 1 | 1 | | | | | | | | | | | | | | | | 3.26 | 1 | 1 | 337 | 37.1 | 6.74 |
| Q92685 | 2.74 | 1 | 1 | 1 | 1 | | | | | | | | | | | | | | | | 2.74 | 1 | 1 | 438 | 50.1 | 9.44 |
| Q712K3 | 4.62 | 1 | 1 | 1 | 1 | | | | | | | | | | | | | | | | 4.62 | 1 | 1 | 238 | 27.1 | 4.42 |
| O75368 | 8.77 | 1 | 1 | 1 | 1 | | | | | | | | | | | | | | | | 8.77 | 1 | 1 | 114 | 12.8 | 5.25 |
| Q14624 | 0.86 | 1 | 1 | 1 | 1 | | | | | | | | | | | | | | | | 0.86 | 1 | 1 | 930 | 103.3 | 6.98 |
| Q4V328 | 0.71 | 1 | 1 | 1 | 1 | | | | | | | | | | | | | | | | 0.71 | 1 | 1 | 841 | 95.9 | 5.11 |
| Q8N5I2 | 2.77 | 1 | 1 | 1 | 1 | | | | | | | | | | | | | | | | 2.77 | 1 | 1 | 433 | 46.0 | 7.02 |
| Q9NY27 | 1.92 | 1 | 1 | 1 | 1 | | | | | | | | | | | | | | | | 1.92 | 1 | 1 | 417 | 46.9 | 4.54 |
| P04062 | 4.29 | 1 | 1 | 1 | 1 | | | | | | | | | | | | | | | | 4.29 | 1 | 1 | 536 | 59.7 | 7.61 |
| P20810 | 1.84 | 1 | 1 | 1 | 1 | | | | | | | | | | | | | | | | 1.84 | 1 | 1 | 708 | 76.5 | 5.07 |
| O00273 | 4.83 | 1 | 1 | 1 | 1 | | | | | | | | | | | | | | | | 4.83 | 1 | 1 | 331 | 36.5 | 4.79 |
| P84157 | 4.90 | 1 | 1 | 1 | 1 | | | | | | | | | | | | | | | | 4.90 | 1 | 1 | 204 | 21.5 | 4.26 |
| Q6IQ49 | 1.55 | 1 | 1 | 1 | 1 | | | | | | | | | | | | | | | | 1.55 | 1 | 1 | 451 | 49.7 | 6.05 |
| Q9BRT9 | 2.69 | 1 | 1 | 1 | 1 | | | | | | | | | | | | | | | | 2.69 | 1 | 1 | 223 | 26.0 | 4.98 |
| Q9BQE4 | 6.35 | 1 | 1 | 1 | 1 | | | | | | | | | | | | | | | | 6.35 | 1 | 1 | 189 | 21.2 | 9.70 |
| Q9Y2S0 | 6.02 | 1 | 1 | 1 | 1 | | | | | | | | | | | | | | | | 6.02 | 1 | 1 | 133 | 15.2 | 5.80 |
| Q96FQ6 | 10.68 | 1 | 1 | 1 | 1 | | | | | | | | | | | | | | | | 10.68 | 1 | 1 | 103 | 11.8 | 6.79 |
| P53992 | 1.19 | 1 | 1 | 1 | 1 | | | | | | | | | | | | | | | | 1.19 | 1 | 1 | 1094 | 118.2 | 7.06 |
| P10619 | 2.71 | 1 | 1 | 1 | 1 | | | | | | | | | | | | | | | | 2.71 | 1 | 1 | 480 | 54.4 | 6.61 |
| Q5TAP6 | 1.04 | 1 | 1 | 1 | 1 | | | | | | | | | | | | | | | | 1.04 | 1 | 1 | 766 | 87.1 | 7.15 |
| Q9UBK9 | 7.01 | 1 | 1 | 1 | 1 | | | | | | | | | | | | | | | | 7.01 | 1 | 1 | 157 | 18.2 | 7.59 |
| P82933 | 3.03 | 1 | 1 | 1 | 1 | | | | | | | | | | | | | | | | 3.03 | 1 | 1 | 396 | 45.8 | 9.51 |
| Q6NUM9 | 1.97 | 1 | 1 | 1 | 1 | | | | | | | | | | | | | | | | 1.97 | 1 | 1 | 610 | 66.8 | 8.28 |
| P51398 | 1.76 | 1 | 1 | 1 | 1 | | | | | | | | | | | | | | | | 1.76 | 1 | 1 | 398 | 45.5 | 8.88 |
| Q03169 | 2.75 | 1 | 1 | 1 | 1 | | | | | | | | | | | | | | | | 2.75 | 1 | 1 | 654 | 72.6 | 6.46 |
| Q15796 | 2.36 | 1 | 1 | 1 | 1 | | | | | | | | | | | | | | | | 2.36 | 1 | 1 | 467 | 52.3 | 6.58 |
| P82663 | 7.51 | 1 | 1 | 1 | 1 | | | | | | | | | | | | | | | | 7.51 | 1 | 1 | 173 | 20.1 | 8.82 |
| P41743 | 2.35 | 1 | 1 | 1 | 1 | | | | | | | | | | | | | | | | 2.35 | 1 | 1 | 596 | 68.2 | 5.85 |
| Q9BQ67 | 3.59 | 1 | 1 | 1 | 1 | | | | | | | | | | | | | | | | 3.59 | 1 | 1 | 446 | 49.4 | 4.92 |
| Q01085 | 2.93 | 1 | 1 | 1 | 1 | | | | | | | | | | | | | | | | 2.93 | 1 | 1 | 375 | 41.6 | 7.74 |
| Q5T1C6 | 5.00 | 1 | 1 | 1 | 1 | | | | | | | | | | | | | | | | 5.00 | 1 | 1 | 240 | 27.1 | 8.28 |
| Q12802 | 0.57 | 1 | 1 | 1 | 1 | | | | | | | | | | | | | | | | 0.57 | 1 | 1 | 2813 | 307.4 | 5.24 |
| P28799 | 3.04 | 1 | 1 | 1 | 1 | | | | | | | | | | | | | | | | 3.04 | 1 | 1 | 593 | 63.5 | 6.83 |
| P13164 | 12.80 | 3 | 1 | 1 | 1 | | | | | | | | | | | | | | | | 12.80 | 1 | 1 | 125 | 14.0 | 7.93 |
| P61960 | 9.41 | 1 | 1 | 1 | 1 | | | | | | | | | | | | | | | | 9.41 | 1 | 1 | 85 | 9.1 | 9.31 |
| O60551 | 2.61 | 1 | 1 | 1 | 1 | | | | | | | | | | | | | | | | 2.61 | 1 | 1 | 498 | 56.9 | 7.58 |
| Q9H3P7 | 2.46 | 1 | 1 | 1 | 1 | | | | | | | | | | | | | | | | 2.46 | 1 | 1 | 528 | 60.6 | 5.06 |
| Q16850 | 2.78 | 1 | 1 | 1 | 1 | | | | | | | | | | | | | | | | 2.78 | 1 | 1 | 503 | 56.8 | 8.53 |
| O75717 | 1.06 | 1 | 1 | 1 | 1 | | | | | | | | | | | | | | | | 1.06 | 1 | 1 | 1129 | 125.9 | 5.62 |
| O15269 | 2.96 | 1 | 1 | 1 | 1 | | | | | | | | | | | | | | | | 2.96 | 1 | 1 | 473 | 52.7 | 6.01 |
| Q5BKZ1 | 1.03 | 1 | 1 | 1 | 1 | | | | | | | | | | | | | | | | 1.03 | 1 | 1 | 582 | 65.6 | 5.15 |
| O14974 | 1.17 | 1 | 1 | 1 | 1 | | | | | | | | | | | | | | | | 1.17 | 1 | 1 | 1030 | 115.2 | 5.40 |
| Q9BX95 | 1.36 | 1 | 1 | 1 | 1 | | | | | | | | | | | | | | | | 1.36 | 1 | 1 | 441 | 49.1 | 8.82 |
| Q7Z4Q2 | 0.88 | 1 | 1 | 1 | 1 | | | | | | | | | | | | | | | | 0.88 | 1 | 1 | 680 | 74.5 | 5.11 |
| O43809 | 7.93 | 1 | 1 | 1 | 1 | | | | | | | | | | | | | | | | 7.93 | 1 | 1 | 227 | 26.2 | 8.82 |
| Q9NPF4 | 5.07 | 1 | 1 | 1 | 1 | | | | | | | | | | | | | | | | 5.07 | 1 | 1 | 335 | 36.4 | 6.35 |
| Q9C0D9 | 4.03 | 1 | 1 | 1 | 1 | | | | | | | | | | | | | | | | 4.03 | 1 | 1 | 397 | 45.2 | 6.60 |
| Q96B26 | 9.78 | 1 | 1 | 1 | 1 | | | | | | | | | | | | | | | | 9.78 | 1 | 1 | 276 | 30.0 | 5.30 |
| Q15165 | 2.54 | 1 | 1 | 1 | 1 | | | | | | | | | | | | | | | | 2.54 | 1 | 1 | 354 | 39.4 | 5.60 |
| Q06203 | 3.68 | 1 | 1 | 1 | 1 | | | | | | | | | | | | | | | | 3.68 | 1 | 1 | 517 | 57.4 | 6.76 |
| O60658 | 1.57 | 1 | 1 | 1 | 1 | | | | | | | | | | | | | | | | 1.57 | 1 | 1 | 829 | 93.2 | 6.11 |
| Q9UHI6 | 1.82 | 1 | 1 | 1 | 1 | | | | | | | | | | | | | | | | 1.82 | 1 | 1 | 824 | 92.2 | 6.95 |
| Q12849 | 4.58 | 1 | 1 | 1 | 1 | | | | | | | | | | | | | | | | 4.58 | 1 | 1 | 480 | 53.1 | 6.19 |
| A6NHR9 | 0.65 | 1 | 1 | 1 | 1 | | | | | | | | | | | | | | | | 0.65 | 1 | 1 | 2005 | 226.2 | 7.30 |
| Q9NTX5 | 3.58 | 1 | 1 | 1 | 1 | | | | | | | | | | | | | | | | 3.58 | 1 | 1 | 307 | 33.7 | 8.21 |
| Q8IWI9 | 0.26 | 1 | 1 | 1 | 1 | | | | | | | | | | | | | | | | 0.26 | 1 | 1 | 3026 | 331.6 | 6.79 |
| Q9Y2Z4 | 4.40 | 1 | 1 | 1 | 1 | | | | | | | | | | | | | | | | 4.40 | 1 | 1 | 477 | 53.2 | 8.98 |
| Q9Y3D7 | 18.40 | 1 | 1 | 1 | 1 | | | | | | | | | | | | | | | | 18.40 | 1 | 1 | 125 | 13.8 | 9.70 |
| P52594 | 3.02 | 1 | 1 | 1 | 1 | | | | | | | | | | | | | | | | 3.02 | 1 | 1 | 562 | 58.2 | 8.63 |
| Q9BQ69 | 4.31 | 1 | 1 | 1 | 1 | | | | | | | | | | | | | | | | 4.31 | 1 | 1 | 325 | 35.5 | 9.51 |
| Q06124 | 3.02 | 1 | 1 | 1 | 1 | | | | | | | | | | | | | | | | 3.02 | 1 | 1 | 597 | 68.4 | 7.30 |
| Q9ULT0 | 1.28 | 1 | 1 | 1 | 1 | | | | | | | | | | | | | | | | 1.28 | 1 | 1 | 858 | 96.1 | 6.46 |
| Q8IVS2 | 2.82 | 1 | 1 | 1 | 1 | | | | | | | | | | | | | | | | 2.82 | 1 | 1 | 390 | 42.9 | 8.72 |
| Q7Z6I6 | 0.91 | 1 | 1 | 1 | 1 | | | | | | | | | | | | | | | | 0.91 | 1 | 1 | 1101 | 118.5 | 4.81 |
| Q9ULJ6 | 0.56 | 1 | 1 | 1 | 1 | | | | | | | | | | | | | | | | 0.56 | 1 | 1 | 1067 | 115.4 | 7.46 |
| Q8N4H5 | 13.73 | 1 | 1 | 1 | 1 | | | | | | | | | | | | | | | | 13.73 | 1 | 1 | 51 | 6.0 | 9.70 |
| Q96T58 | 0.22 | 1 | 1 | 1 | 1 | | | | | | | | | | | | | | | | 0.22 | 1 | 1 | 3664 | 402.0 | 7.64 |
| Q9ULM0 | 1.39 | 1 | 1 | 1 | 1 | | | | | | | | | | | | | | | | 1.39 | 1 | 1 | 1364 | 151.1 | 7.94 |
| Q12796 | 2.14 | 1 | 1 | 1 | 1 | | | | | | | | | | | | | | | | 2.14 | 1 | 1 | 327 | 35.2 | 10.51 |
| Q8NEZ2 | 2.02 | 1 | 1 | 1 | 1 | | | | | | | | | | | | | | | | 2.02 | 1 | 1 | 397 | 44.3 | 5.57 |
| Q07954 | 0.24 | 1 | 1 | 1 | 1 | | | | | | | | | | | | | | | | 0.24 | 1 | 1 | 4544 | 504.3 | 5.39 |
| Q96AX1 | 3.52 | 1 | 1 | 1 | 1 | | | | | | | | | | | | | | | | 3.52 | 1 | 1 | 596 | 67.6 | 6.96 |
| Q9BZC7 | 0.25 | 1 | 1 | 1 | 1 | | | | | | | | | | | | | | | | 0.25 | 1 | 1 | 2435 | 269.7 | 6.81 |
| Q9BS40 | 8.11 | 1 | 1 | 1 | 1 | | | | | | | | | | | | | | | | 8.11 | 1 | 1 | 222 | 25.7 | 5.78 |
| Q9BYJ9 | 2.33 | 2 | 1 | 1 | 1 | | | | | | | | | | | | | | | | 2.33 | 1 | 1 | 559 | 60.8 | 8.79 |
| Q8N1G2 | 0.84 | 1 | 1 | 1 | 1 | | | | | | | | | | | | | | | | 0.84 | 1 | 1 | 835 | 95.3 | 7.05 |
| Q8N2G8 | 1.70 | 1 | 1 | 1 | 1 | | | | | | | | | | | | | | | | 1.70 | 1 | 1 | 530 | 57.5 | 7.88 |
| Q9BZ29 | 1.35 | 1 | 1 | 1 | 1 | | | | | | | | | | | | | | | | 1.35 | 1 | 1 | 2069 | 236.3 | 7.49 |
| P47736 | 1.96 | 1 | 1 | 1 | 1 | | | | | | | | | | | | | | | | 1.96 | 1 | 1 | 663 | 73.3 | 5.82 |
| Q15042 | 1.94 | 1 | 1 | 1 | 1 | | | | | | | | | | | | | | | | 1.94 | 1 | 1 | 981 | 110.5 | 5.55 |
| Q8IYU8 | 3.92 | 1 | 1 | 1 | 1 | | | | | | | | | | | | | | | | 3.92 | 1 | 1 | 434 | 49.6 | 9.09 |
| Q9NQP4 | 11.19 | 1 | 1 | 1 | 1 | | | | | | | | | | | | | | | | 11.19 | 1 | 1 | 134 | 15.3 | 4.53 |
| Q96L92 | 5.36 | 1 | 1 | 1 | 1 | | | | | | | | | | | | | | | | 5.36 | 1 | 1 | 541 | 61.2 | 6.49 |
| P46734 | 3.17 | 1 | 1 | 1 | 1 | | | | | | | | | | | | | | | | 3.17 | 1 | 1 | 347 | 39.3 | 7.43 |
| O60783 | 11.72 | 1 | 1 | 1 | 1 | | | | | | | | | | | | | | | | 11.72 | 1 | 1 | 128 | 15.1 | 11.41 |
| Q9NPL8 | 3.16 | 1 | 1 | 1 | 1 | | | | | | | | | | | | | | | | 3.16 | 1 | 1 | 285 | 32.2 | 8.50 |
| P63098 | 5.88 | 1 | 1 | 1 | 1 | | | | | | | | | | | | | | | | 5.88 | 1 | 1 | 170 | 19.3 | 4.81 |
| Q8N490 | 2.08 | 1 | 1 | 1 | 1 | | | | | | | | | | | | | | | | 2.08 | 1 | 1 | 385 | 42.8 | 9.09 |
| Q9NX47 | 5.76 | 1 | 1 | 1 | 1 | | | | | | | | | | | | | | | | 5.76 | 1 | 1 | 278 | 31.2 | 8.70 |
| Q5TIE3 | 0.66 | 1 | 1 | 1 | 1 | | | | | | | | | | | | | | | | 0.66 | 1 | 1 | 1220 | 133.7 | 8.15 |
| Q9Y6I3 | 1.74 | 2 | 1 | 1 | 1 | | | | | | | | | | | | | | | | 1.74 | 1 | 1 | 576 | 60.3 | 4.83 |
| Q9Y320 | 5.07 | 1 | 1 | 1 | 1 | | | | | | | | | | | | | | | | 5.07 | 1 | 1 | 296 | 34.0 | 8.69 |
| P36954 | 18.40 | 1 | 1 | 1 | 1 | | | | | | | | | | | | | | | | 18.40 | 1 | 1 | 125 | 14.5 | 5.14 |
| Q8NEZ4 | 0.14 | 1 | 1 | 1 | 1 | | | | | | | | | | | | | | | | 0.14 | 1 | 1 | 4911 | 541.0 | 6.49 |
| Q5TAX3 | 0.43 | 1 | 1 | 1 | 1 | | | | | | | | | | | | | | | | 0.43 | 1 | 1 | 1644 | 185.0 | 7.97 |
| Q92481 | 1.74 | 1 | 1 | 1 | 1 | | | | | | | | | | | | | | | | 1.74 | 1 | 1 | 460 | 50.4 | 8.24 |
| O95714 | 0.52 | 1 | 1 | 1 | 1 | | | | | | | | | | | | | | | | 0.52 | 1 | 1 | 4834 | 526.9 | 6.28 |
| Q38SD2 | 0.40 | 1 | 1 | 1 | 1 | | | | | | | | | | | | | | | | 0.40 | 1 | 1 | 2015 | 225.2 | 6.68 |
| P20929 | 0.12 | 1 | 1 | 1 | 1 | | | | | | | | | | | | | | | | 0.12 | 1 | 1 | 6669 | 772.4 | 9.07 |
| P08069 | 0.51 | 1 | 1 | 1 | 1 | | | | | | | | | | | | | | | | 0.51 | 1 | 1 | 1367 | 154.7 | 5.80 |
| Q3YEC7 | 1.51 | 1 | 1 | 1 | 1 | | | | | | | | | | | | | | | | 1.51 | 1 | 1 | 729 | 79.5 | 5.22 |
| Q9NX02 | 1.69 | 1 | 1 | 1 | 1 | | | | | | | | | | | | | | | | 1.69 | 1 | 1 | 1062 | 120.4 | 6.07 |
| Q969I3 | 1.99 | 1 | 1 | 1 | 1 | | | | | | | | | | | | | | | | 1.99 | 1 | 1 | 302 | 35.1 | 6.87 |
| O75192 | 4.05 | 1 | 1 | 1 | 1 | | | | | | | | | | | | | | | | 4.05 | 1 | 1 | 247 | 28.3 | 9.64 |
| Q8WY91 | 1.56 | 1 | 1 | 1 | 1 | | | | | | | | | | | | | | | | 1.56 | 1 | 1 | 577 | 62.9 | 9.28 |
| Q68D91 | 3.58 | 1 | 1 | 1 | 1 | | | | | | | | | | | | | | | | 3.58 | 1 | 1 | 279 | 31.4 | 6.92 |
| Q6UXV4 | 4.48 | 1 | 1 | 1 | 1 | | | | | | | | | | | | | | | | 4.48 | 1 | 1 | 268 | 29.1 | 9.52 |
| Q9NRL3 | 0.93 | 1 | 1 | 1 | 1 | | | | | | | | | | | | | | | | 0.93 | 1 | 1 | 753 | 80.5 | 5.40 |
| Q9HB90 | 3.01 | 2 | 1 | 1 | 1 | | | | | | | | | | | | | | | | 3.01 | 1 | 1 | 399 | 44.2 | 5.10 |
| Q9H4A3 | 0.55 | 1 | 1 | 1 | 1 | | | | | | | | | | | | | | | | 0.55 | 1 | 1 | 2382 | 250.6 | 6.34 |
| Q9BW04 | 2.50 | 1 | 1 | 1 | 1 | | | | | | | | | | | | | | | | 2.50 | 1 | 1 | 601 | 63.9 | 8.62 |
| P08910 | 2.59 | 1 | 1 | 1 | 1 | | | | | | | | | | | | | | | | 2.59 | 1 | 1 | 425 | 48.3 | 6.70 |
| Q96EC8 | 5.08 | 1 | 1 | 1 | 1 | | | | | | | | | | | | | | | | 5.08 | 1 | 1 | 236 | 26.2 | 5.64 |
| Q07617 | 0.76 | 1 | 1 | 1 | 1 | | | | | | | | | | | | | | | | 0.76 | 1 | 1 | 926 | 103.6 | 6.86 |
| P49593 | 4.41 | 1 | 1 | 1 | 1 | | | | | | | | | | | | | | | | 4.41 | 1 | 1 | 454 | 49.8 | 5.10 |
| O60522 | 0.52 | 1 | 1 | 1 | 1 | | | | | | | | | | | | | | | | 0.52 | 1 | 1 | 2096 | 236.4 | 5.25 |
| Q92556 | 1.38 | 1 | 1 | 1 | 1 | | | | | | | | | | | | | | | | 1.38 | 1 | 1 | 727 | 83.8 | 6.28 |
| Q9HC96 | 1.34 | 1 | 1 | 1 | 1 | | | | | | | | | | | | | | | | 1.34 | 1 | 1 | 672 | 74.9 | 7.74 |
| Q8TB72 | 0.94 | 2 | 1 | 1 | 1 | | | | | | | | | | | | | | | | 0.94 | 1 | 1 | 1066 | 114.1 | 7.08 |
| A6NJZ7 | 0.61 | 3 | 1 | 1 | 1 | | | | | | | | | | | | | | | | 0.61 | 1 | 1 | 1639 | 180.8 | 6.77 |
| Q8N4P3 | 8.94 | 1 | 1 | 1 | 1 | | | | | | | | | | | | | | | | 8.94 | 1 | 1 | 179 | 20.3 | 6.74 |
| Q6DN90 | 1.14 | 1 | 1 | 1 | 1 | | | | | | | | | | | | | | | | 1.14 | 1 | 1 | 963 | 108.2 | 6.93 |
| Q8TD31 | 0.90 | 1 | 1 | 1 | 1 | | | | | | | | | | | | | | | | 0.90 | 1 | 1 | 782 | 88.6 | 5.95 |

| | | | | | | | | | | | |
|---|---|---|---|---|---|---|---|---|---|---|---|
| O94817 | 7.14 | 1 | 1 | 1 | 1 | 7.14 | 1 | 1 | 140 | 15.1 | 5.10 |
| Q6JVE6 | 5.35 | 1 | 1 | 1 | 1 | 5.35 | 1 | 1 | 187 | 20.7 | 10.35 |
| Q13835 | 1.61 | 1 | 1 | 1 | 1 | 1.61 | 1 | 1 | 747 | 82.8 | 9.13 |
| O15534 | 0.54 | 1 | 1 | 1 | 1 | 0.54 | 1 | 1 | 1290 | 136.1 | 6.09 |
| O00635 | 2.58 | 1 | 1 | 1 | 1 | 2.58 | 1 | 1 | 465 | 53.4 | 7.02 |
| Q6P995 | 1.09 | 1 | 1 | 1 | 1 | 1.09 | 1 | 1 | 826 | 92.1 | 8.65 |
| P51798 | 1.12 | 1 | 1 | 1 | 1 | 1.12 | 1 | 1 | 805 | 88.6 | 8.43 |
| Q8TBQ9 | 12.50 | 1 | 1 | 1 | 1 | 12.50 | 1 | 1 | 72 | 8.1 | 8.95 |
| Q9BSL1 | 3.21 | 1 | 1 | 1 | 1 | 3.21 | 1 | 1 | 405 | 45.3 | 4.92 |
| Q9Y5Z9 | 2.96 | 1 | 1 | 1 | 1 | 2.96 | 1 | 1 | 338 | 36.8 | 8.15 |
| Q9H799 | 0.25 | 1 | 1 | 1 | 1 | 0.25 | 1 | 1 | 3197 | 361.5 | 6.99 |
| A6NKD2 | 5.84 | 5 | 1 | 1 | 1 | 5.84 | 1 | 1 | 308 | 35.1 | 5.39 |
| Q15388 | 4.83 | 1 | 1 | 1 | 1 | 4.83 | 1 | 1 | 145 | 16.3 | 8.60 |
| Q9P0I2 | 5.36 | 1 | 1 | 1 | 1 | 5.36 | 1 | 1 | 261 | 29.9 | 6.81 |
| P62256 | 8.20 | 1 | 1 | 1 | 1 | 8.20 | 1 | 1 | 183 | 20.6 | 4.67 |
| Q7Z7N9 | 7.31 | 1 | 1 | 1 | 1 | 7.31 | 1 | 1 | 219 | 23.5 | 7.85 |
| Q8N319 | 3.31 | 1 | 1 | 1 | 1 | 3.31 | 1 | 1 | 242 | 26.1 | 12.19 |
| Q86YV5 | 0.78 | 1 | 1 | 1 | 1 | 0.78 | 1 | 1 | 1406 | 149.5 | 7.20 |
| Q5S007 | 0.24 | 1 | 1 | 1 | 1 | 0.24 | 1 | 1 | 2527 | 285.9 | 6.80 |
| Q9BVL4 | 2.99 | 1 | 1 | 1 | 1 | 2.99 | 1 | 1 | 669 | 73.4 | 5.97 |
| Q9BXI6 | 2.17 | 1 | 1 | 1 | 1 | 2.17 | 1 | 1 | 508 | 57.1 | 8.44 |
| Q9UKN7 | 0.28 | 1 | 1 | 1 | 1 | 0.28 | 1 | 1 | 3530 | 395.0 | 9.17 |
| Q7L8A9 | 6.30 | 1 | 1 | 1 | 1 | 6.30 | 1 | 1 | 365 | 40.9 | 9.48 |
| Q8TDX7 | 2.65 | 2 | 1 | 1 | 1 | 2.65 | 1 | 1 | 302 | 34.5 | 8.25 |
| Q8IZA0 | 1.62 | 1 | 1 | 1 | 1 | 1.62 | 1 | 1 | 1049 | 115.6 | 6.05 |
| Q15118 | 2.52 | 1 | 1 | 1 | 1 | 2.52 | 1 | 1 | 436 | 49.2 | 8.81 |
| Q96KA5 | 2.04 | 1 | 1 | 1 | 1 | 2.04 | 1 | 1 | 538 | 62.2 | 8.56 |
| Q9BZF1 | 1.69 | 1 | 1 | 1 | 1 | 1.69 | 1 | 1 | 889 | 101.1 | 6.96 |
| Q9NPI6 | 1.03 | 1 | 1 | 1 | 1 | 1.03 | 1 | 1 | 582 | 63.3 | 6.25 |
| Q13190 | 7.89 | 1 | 1 | 1 | 1 | 7.89 | 1 | 1 | 355 | 39.6 | 9.16 |
| Q9UGK3 | 2.73 | 1 | 1 | 1 | 1 | 2.73 | 1 | 1 | 403 | 44.9 | 8.16 |
| Q9UQB9 | 2.27 | 1 | 1 | 1 | 1 | 2.27 | 1 | 1 | 309 | 35.6 | 8.81 |
| Q9BQG2 | 1.52 | 1 | 1 | 1 | 1 | 1.52 | 1 | 1 | 462 | 52.0 | 6.83 |
| Q0VDG4 | 2.59 | 1 | 1 | 1 | 1 | 2.59 | 1 | 1 | 424 | 48.5 | 5.55 |
| Q9UIV1 | 6.67 | 1 | 1 | 1 | 1 | 6.67 | 1 | 1 | 285 | 32.7 | 4.84 |
| Q15051 | 2.01 | 1 | 1 | 1 | 1 | 2.01 | 1 | 1 | 598 | 68.9 | 9.10 |
| Q02083 | 2.23 | 1 | 1 | 1 | 1 | 2.23 | 1 | 1 | 359 | 40.0 | 8.75 |
| Q96MX6 | 3.36 | 1 | 1 | 1 | 1 | 3.36 | 1 | 1 | 357 | 39.7 | 8.09 |
| Q5T0N5 | 1.49 | 1 | 1 | 1 | 1 | 1.49 | 1 | 1 | 605 | 70.0 | 6.64 |
| P78560 | 8.04 | 1 | 1 | 1 | 1 | 8.04 | 1 | 1 | 199 | 22.7 | 6.80 |
| Q15404 | 2.53 | 1 | 1 | 1 | 1 | 2.53 | 1 | 1 | 277 | 31.5 | 8.65 |
| Q13488 | 0.96 | 1 | 1 | 1 | 1 | 0.96 | 1 | 1 | 830 | 92.9 | 7.12 |
| Q9HCI6 | 0.89 | 1 | 1 | 1 | 1 | 0.89 | 1 | 1 | 787 | 89.6 | 7.74 |
| O43299 | 0.87 | 1 | 1 | 1 | 1 | 0.87 | 1 | 1 | 807 | 88.5 | 7.01 |
| Q9NZ42 | 14.85 | 1 | 1 | 1 | 1 | 14.85 | 1 | 1 | 101 | 12.0 | 9.19 |
| O15013 | 0.51 | 1 | 1 | 1 | 1 | 0.51 | 1 | 1 | 1369 | 151.5 | 5.68 |
| Q96CN5 | 0.90 | 1 | 1 | 1 | 1 | 0.90 | 1 | 1 | 670 | 75.9 | 6.23 |
| Q9NZB8 | 1.26 | 1 | 1 | 1 | 1 | 1.26 | 1 | 1 | 636 | 70.1 | 9.26 |
| Q8IZY2 | 0.23 | 1 | 1 | 1 | 1 | 0.23 | 1 | 1 | 2146 | 234.2 | 7.24 |
| Q16890 | 6.86 | 1 | 1 | 1 | 1 | 6.86 | 1 | 1 | 204 | 22.4 | 5.62 |
| P14635 | 2.54 | 1 | 1 | 1 | 1 | 2.54 | 1 | 1 | 433 | 48.3 | 7.47 |
| Q6UB98 | 0.39 | 1 | 1 | 1 | 1 | 0.39 | 1 | 1 | 2062 | 235.5 | 7.01 |
| Q8IXW5 | 1.47 | 1 | 1 | 1 | 1 | 1.47 | 1 | 1 | 612 | 69.5 | 7.78 |
| Q9H7C9 | 9.02 | 1 | 1 | 1 | 1 | 9.02 | 1 | 1 | 122 | 13.3 | 8.46 |
| Q9NXZ2 | 1.23 | 1 | 1 | 1 | 1 | 1.23 | 1 | 1 | 648 | 72.8 | 8.75 |
| O95907 | 1.39 | 1 | 1 | 1 | 1 | 1.39 | 1 | 1 | 504 | 52.3 | 5.67 |
| Q14191 | 0.49 | 1 | 1 | 1 | 1 | 0.49 | 1 | 1 | 1432 | 162.4 | 6.34 |
| Q96M89 | 1.50 | 1 | 1 | 1 | 1 | 1.50 | 1 | 1 | 665 | 76.2 | 8.53 |
| P28039 | 1.22 | 1 | 1 | 1 | 1 | 1.22 | 1 | 1 | 575 | 65.1 | 8.16 |
| O60500 | 0.56 | 1 | 1 | 1 | 1 | 0.56 | 1 | 1 | 1241 | 134.7 | 5.64 |
| Q7Z392 | 1.24 | 1 | 1 | 1 | 1 | 1.24 | 1 | 1 | 1133 | 128.8 | 7.14 |
| Q96Q35 | 1.80 | 1 | 1 | 1 | 1 | 1.80 | 1 | 1 | 445 | 52.4 | 6.71 |
| Q969G6 | 7.10 | 1 | 1 | 1 | 1 | 7.10 | 1 | 1 | 155 | 17.6 | 8.13 |
| P40818 | 1.16 | 1 | 1 | 1 | 1 | 1.16 | 1 | 1 | 1118 | 127.4 | 8.51 |
| F5H4B4 | 3.33 | 1 | 1 | 1 | 1 | 3.33 | 1 | 1 | 570 | 66.1 | 9.31 |
| O43687 | 11.54 | 2 | 1 | 1 | 1 | 11.54 | 1 | 1 | 104 | 11.5 | 4.94 |
| Q67FW5 | 1.94 | 1 | 1 | 1 | 1 | 1.94 | 1 | 1 | 361 | 40.7 | 7.40 |
| Q16513 | 0.91 | 1 | 1 | 1 | 1 | 0.91 | 1 | 1 | 984 | 112.0 | 6.30 |
| Q9H300 | 2.37 | 1 | 1 | 1 | 1 | 2.37 | 1 | 1 | 379 | 42.2 | 9.79 |
| Q8TBB5 | 2.12 | 1 | 1 | 1 | 1 | 2.12 | 1 | 1 | 520 | 57.9 | 5.72 |
| Q9NXL2 | 1.29 | 1 | 1 | 1 | 1 | 1.29 | 1 | 1 | 777 | 89.0 | 6.25 |
| Q05209 | 1.28 | 1 | 1 | 1 | 1 | 1.28 | 1 | 1 | 780 | 88.1 | 5.62 |
| Q96EX3 | 1.31 | 1 | 1 | 1 | 1 | 1.31 | 1 | 1 | 536 | 57.8 | 6.64 |
| Q8TCT1 | 4.12 | 1 | 1 | 1 | 1 | 4.12 | 1 | 1 | 267 | 29.7 | 7.71 |
| Q96PZ2 | 1.47 | 1 | 1 | 1 | 1 | 1.47 | 1 | 1 | 611 | 70.2 | 8.53 |
| Q32P41 | 1.38 | 1 | 1 | 1 | 1 | 1.38 | 1 | 1 | 509 | 58.2 | 8.62 |
| O15055 | 0.72 | 1 | 1 | 1 | 1 | 0.72 | 1 | 1 | 1255 | 136.5 | 6.47 |
| O14578 | 0.49 | 1 | 1 | 1 | 1 | 0.49 | 1 | 1 | 2027 | 231.3 | 6.57 |
| Q96AJ9 | 4.61 | 1 | 1 | 1 | 1 | 4.61 | 1 | 1 | 217 | 25.2 | 6.40 |
| Q8NBJ4 | 3.49 | 1 | 1 | 1 | 1 | 3.49 | 1 | 1 | 401 | 45.3 | 4.97 |
| Q5TGL8 | 3.90 | 1 | 1 | 1 | 1 | 3.90 | 1 | 1 | 231 | 26.5 | 5.03 |
| Q96J88 | 3.14 | 1 | 1 | 1 | 1 | 3.14 | 1 | 1 | 318 | 36.8 | 9.89 |
| Q5THJ4 | 0.18 | 1 | 1 | 1 | 1 | 0.18 | 1 | 1 | 4388 | 491.6 | 6.58 |
| O75716 | 2.95 | 1 | 1 | 1 | 1 | 2.95 | 1 | 1 | 305 | 34.6 | 6.90 |
| Q96C11 | 2.00 | 1 | 1 | 1 | 1 | 2.00 | 1 | 1 | 551 | 60.0 | 6.35 |
| A8MTZ7 | 2.60 | 1 | 1 | 1 | 1 | 2.60 | 1 | 1 | 269 | 30.3 | 5.40 |
| Q6ZRI0 | 0.34 | 1 | 1 | 1 | 1 | 0.34 | 1 | 1 | 2925 | 314.6 | 5.91 |
| Q9BRT3 | 9.57 | 1 | 1 | 1 | 1 | 9.57 | 1 | 1 | 115 | 12.4 | 4.37 |
| Q9HD26 | 1.95 | 1 | 1 | 1 | 1 | 1.95 | 1 | 1 | 462 | 50.5 | 5.92 |
| Q7Z3E5 | 1.84 | 1 | 1 | 1 | 1 | 1.84 | 1 | 1 | 817 | 91.7 | 6.20 |
| Q6UWE0 | 0.83 | 3 | 1 | 1 | 1 | 0.83 | 1 | 1 | 723 | 83.5 | 5.94 |
| Q7Z6G3 | 1.81 | 1 | 1 | 1 | 1 | 1.81 | 1 | 1 | 386 | 43.2 | 5.50 |
| Q58EX2 | 0.32 | 1 | 1 | 1 | 1 | 0.32 | 1 | 1 | 2172 | 239.2 | 7.06 |
| Q5T3U5 | 0.80 | 1 | 1 | 1 | 1 | 0.80 | 1 | 1 | 1492 | 161.5 | 7.08 |
| O95876 | 1.74 | 1 | 1 | 1 | 1 | 1.74 | 1 | 1 | 746 | 85.0 | 6.39 |
| Q6ZSS3 | 1.82 | 1 | 1 | 1 | 1 | 1.82 | 1 | 1 | 439 | 49.2 | 8.94 |
| Q8NEN9 | 0.69 | 1 | 1 | 1 | 1 | 0.69 | 1 | 1 | 1154 | 128.5 | 6.09 |
| O60239 | 1.32 | 1 | 1 | 1 | 1 | 1.32 | 1 | 1 | 455 | 50.4 | 4.97 |
| Q8N8E3 | 0.63 | 1 | 1 | 1 | 1 | 0.63 | 1 | 1 | 955 | 112.7 | 6.68 |
| Q7L0X0 | 0.86 | 1 | 1 | 1 | 1 | 0.86 | 1 | 1 | 811 | 88.7 | 9.60 |
| Q99457 | 1.38 | 1 | 1 | 1 | 1 | 1.38 | 1 | 1 | 506 | 57.6 | 5.08 |
| P12107 | 0.50 | 1 | 1 | 1 | 1 | 0.50 | 1 | 1 | 1806 | 181.0 | 5.17 |
| Q8N608 | 0.88 | 1 | 1 | 1 | 1 | 0.88 | 1 | 1 | 796 | 90.8 | 6.57 |
| Q8N6K0 | 11.92 | 1 | 1 | 1 | 1 | 11.92 | 1 | 1 | 151 | 16.6 | 5.07 |
| Q92730 | 3.88 | 1 | 1 | 1 | 1 | 3.88 | 1 | 1 | 232 | 26.0 | 7.94 |
| O95167 | 8.33 | 1 | 1 | 1 | 1 | 8.33 | 1 | 1 | 84 | 9.3 | 8.46 |
| Q9NZR1 | 1.99 | 1 | 1 | 1 | 1 | 1.99 | 1 | 1 | 351 | 39.6 | 5.27 |
| P11117 | 1.65 | 1 | 1 | 1 | 1 | 1.65 | 1 | 1 | 423 | 48.3 | 6.74 |
| Q9BTT6 | 1.91 | 2 | 1 | 1 | 1 | 1.91 | 1 | 1 | 524 | 59.2 | 5.02 |
| Q9UKG9 | 2.78 | 1 | 1 | 1 | 1 | 2.78 | 1 | 1 | 612 | 70.1 | 7.08 |
| Q8TE59 | 0.83 | 1 | 1 | 1 | 1 | 0.83 | 1 | 1 | 1207 | 134.0 | 7.56 |
| Q5VWQ0 | 0.87 | 1 | 1 | 1 | 1 | 0.87 | 1 | 1 | 802 | 90.0 | 8.60 |
| Q96IW2 | 2.35 | 1 | 1 | 1 | 1 | 2.35 | 1 | 1 | 340 | 38.2 | 5.01 |
| Q96AZ6 | 4.42 | 1 | 1 | 1 | 1 | 4.42 | 1 | 1 | 181 | 20.4 | 8.92 |
| Q9H172 | 1.70 | 1 | 1 | 1 | 1 | 1.70 | 1 | 1 | 646 | 71.8 | 8.16 |
| Q96JK4 | 0.77 | 1 | 1 | 1 | 1 | 0.77 | 1 | 1 | 782 | 86.7 | 7.58 |
| P34820 | 2.49 | 2 | 1 | 1 | 1 | 2.49 | 1 | 1 | 402 | 44.7 | 8.41 |
| P05106 | 1.27 | 1 | 1 | 1 | 1 | 1.27 | 1 | 1 | 788 | 87.0 | 5.24 |
| Q8IUG5 | 0.23 | 1 | 1 | 1 | 1 | 0.23 | 1 | 1 | 2567 | 285.0 | 6.90 |
| Q9BRP4 | 2.55 | 1 | 1 | 1 | 1 | 2.55 | 1 | 1 | 392 | 42.2 | 6.32 |
| Q8N9M1 | 2.84 | 1 | 1 | 1 | 1 | 2.84 | 1 | 1 | 422 | 44.7 | 10.11 |
| O75503 | 3.07 | 1 | 1 | 1 | 1 | 3.07 | 1 | 1 | 358 | 41.5 | 7.40 |
| Q12981 | 3.95 | 1 | 1 | 1 | 1 | 3.95 | 1 | 1 | 228 | 26.1 | 8.95 |
| A6ZKI3 | 12.39 | 2 | 1 | 1 | 1 | 12.39 | 1 | 1 | 113 | 13.2 | 5.07 |
| Q6DKK2 | 2.63 | 1 | 1 | 1 | 1 | 2.63 | 1 | 1 | 380 | 42.4 | 5.77 |
| Q7Z3D6 | 1.95 | 1 | 1 | 1 | 1 | 1.95 | 1 | 1 | 616 | 66.4 | 6.79 |
| Q9H7B4 | 2.34 | 1 | 1 | 1 | 1 | 2.34 | 1 | 1 | 428 | 49.1 | 7.25 |
| Q9Y4F9 | 0.94 | 1 | 1 | 1 | 1 | 0.94 | 1 | 1 | 1068 | 118.4 | 5.41 |
| Q9NVG8 | 3.00 | 1 | 1 | 1 | 1 | 3.00 | 1 | 1 | 400 | 46.5 | 5.24 |
| Q9HB19 | 2.59 | 1 | 1 | 1 | 1 | 2.59 | 1 | 1 | 425 | 47.2 | 8.66 |
| Q9NSB4 | 1.95 | 1 | 1 | 1 | 1 | 1.95 | 1 | 1 | 513 | 56.6 | 6.74 |
| Q6GMV2 | 2.87 | 1 | 1 | 1 | 1 | 2.87 | 1 | 1 | 418 | 47.3 | 5.05 |
| Q9BPZ7 | 2.68 | 1 | 1 | 1 | 1 | 2.68 | 1 | 1 | 522 | 59.1 | 7.55 |
| Q9UBS9 | 0.64 | 1 | 1 | 1 | 1 | 0.64 | 1 | 1 | 1254 | 139.3 | 5.01 |
| P49407 | 5.74 | 1 | 1 | 1 | 1 | 5.74 | 1 | 1 | 418 | 47.0 | 6.20 |
| Q8IVF5 | 0.47 | 1 | 1 | 1 | 1 | 0.47 | 1 | 1 | 1701 | 190.0 | 7.21 |
| Q96PN6 | 0.93 | 1 | 1 | 1 | 1 | 0.93 | 1 | 1 | 1610 | 187.0 | 7.31 |
| Q9H1U4 | 2.49 | 1 | 1 | 1 | 1 | 2.49 | 1 | 1 | 602 | 62.9 | 5.67 |
| O43402 | 4.29 | 1 | 1 | 1 | 1 | 4.29 | 1 | 1 | 210 | 23.8 | 6.40 |
| Q86YW0 | 3.62 | 1 | 1 | 1 | 1 | 3.62 | 1 | 1 | 608 | 70.4 | 9.04 |
| Q496Y0 | 0.79 | 1 | 1 | 1 | 1 | 0.79 | 1 | 1 | 759 | 84.4 | 7.25 |
| Q8TD20 | 1.94 | 1 | 1 | 1 | 1 | 1.94 | 1 | 1 | 617 | 66.9 | 8.37 |
| Q96BW1 | 3.56 | 1 | 1 | 1 | 1 | 3.56 | 1 | 1 | 309 | 33.8 | 6.04 |
| Q9H7F0 | 1.14 | 1 | 1 | 1 | 1 | 1.14 | 1 | 1 | 1226 | 138.0 | 6.64 |
| Q9BV19 | 8.04 | 1 | 1 | 1 | 1 | 8.04 | 1 | 1 | 199 | 21.9 | 5.88 |
| Q9BZZ2 | 0.35 | 1 | 1 | 1 | 1 | 0.35 | 1 | 1 | 1709 | 182.5 | 6.62 |
| Q86VQ0 | 0.86 | 2 | 1 | 1 | 1 | 0.86 | 1 | 1 | 697 | 80.5 | 7.68 |
| P14778 | 1.76 | 1 | 1 | 1 | 1 | 1.76 | 1 | 1 | 569 | 65.4 | 7.83 |
| Q9Y296 | 4.11 | 1 | 1 | 1 | 1 | 4.11 | 1 | 1 | 219 | 24.3 | 6.21 |
| P59044 | 0.78 | 1 | 1 | 1 | 1 | 0.78 | 1 | 1 | 892 | 98.7 | 8.07 |
| Q9ULL5 | 0.66 | 1 | 1 | 1 | 1 | 0.66 | 1 | 1 | 1215 | 129.9 | 8.00 |
| Q9NVF7 | 2.17 | 1 | 1 | 1 | 1 | 2.17 | 1 | 1 | 368 | 41.1 | 9.55 |
| P51957 | 0.71 | 1 | 1 | 1 | 1 | 0.71 | 1 | 1 | 841 | 94.5 | 7.90 |
| Q86TI2 | 1.27 | 1 | 1 | 1 | 1 | 1.27 | 1 | 1 | 863 | 98.2 | 6.46 |
| Q6NSW5 | 2.52 | 2 | 1 | 1 | 1 | 2.52 | 1 | 1 | 357 | 40.5 | 6.54 |
| Q96EB1 | 2.36 | 1 | 1 | 1 | 1 | 2.36 | 1 | 1 | 424 | 46.6 | 8.51 |
| O94952 | 2.39 | 1 | 1 | 1 | 1 | 2.39 | 1 | 1 | 628 | 72.2 | 6.09 |
| Q9HCE7 | 1.32 | 1 | 1 | 1 | 1 | 1.32 | 1 | 1 | 757 | 86.1 | 7.02 |
| Q09013 | 1.11 | 1 | 1 | 1 | 1 | 1.11 | 1 | 1 | 629 | 69.3 | 5.02 |
| Q9UG56 | 1.96 | 1 | 1 | 1 | 1 | 1.96 | 1 | 1 | 409 | 46.6 | 9.42 |
| P41226 | 2.96 | 1 | 1 | 1 | 1 | 2.96 | 1 | 1 | 1012 | 111.6 | 6.04 |
| Q3B7J2 | 2.60 | 1 | 1 | 1 | 1 | 2.60 | 1 | 1 | 385 | 42.2 | 6.44 |
| Q17RC7 | 1.11 | 1 | 1 | 1 | 1 | 1.11 | 1 | 1 | 722 | 79.8 | 6.32 |
| P47972 | 4.41 | 1 | 1 | 1 | 1 | 4.41 | 1 | 1 | 431 | 47.0 | 5.63 |
| Q07973 | 1.17 | 1 | 1 | 1 | 1 | 1.17 | 1 | 1 | 514 | 58.8 | 8.75 |
| P22531 | 25.00 | 6 | 1 | 1 | 1 | 25.00 | 1 | 1 | 72 | 7.8 | 8.31 |
| P35711 | 0.92 | 1 | 1 | 1 | 1 | 0.92 | 1 | 1 | 763 | 84.0 | 6.60 |
| Q8NHQ1 | 1.51 | 1 | 1 | 1 | 1 | 1.51 | 1 | 1 | 597 | 69.7 | 5.71 |
| P22897 | 0.82 | 1 | 1 | 1 | 1 | 0.82 | 1 | 1 | 1456 | 165.9 | 6.54 |
| Q5U649 | 4.49 | 1 | 1 | 1 | 1 | 4.49 | 1 | 1 | 245 | 27.6 | 7.96 |
| Q8IY21 | 0.35 | 1 | 1 | 1 | 1 | 0.35 | 1 | 1 | 1712 | 197.7 | 7.59 |
| Q8NC74 | 1.51 | 1 | 1 | 1 | 1 | 1.51 | 1 | 1 | 664 | 71.4 | 6.51 |
| Q12766 | 0.39 | 1 | 1 | 1 | 1 | 0.39 | 1 | 1 | 1538 | 168.2 | 8.05 |
| P48553 | 0.87 | 1 | 1 | 1 | 1 | 0.87 | 1 | 1 | 1259 | 142.1 | 6.04 |
| Q7L1V2 | 1.65 | 1 | 1 | 1 | 1 | 1.65 | 1 | 1 | 547 | 59.2 | 6.29 |
| Q9NWU2 | 9.65 | 1 | 1 | 1 | 1 | 9.65 | 1 | 1 | 228 | 26.7 | 4.97 |
| Q9Y6Q2 | 1.50 | 1 | 1 | 1 | 1 | 1.50 | 1 | 1 | 735 | 83.1 | 6.20 |
| Q8NB91 | 0.70 | 1 | 1 | 1 | 1 | 0.70 | 1 | 1 | 859 | 97.7 | 7.61 |
| Q9UHL9 | 0.63 | 1 | 1 | 1 | 1 | 0.63 | 1 | 1 | 959 | 106.0 | 6.87 |
| Q86UW6 | 0.45 | 1 | 1 | 1 | 1 | 0.45 | 1 | 1 | 1770 | 198.7 | 5.21 |
| Q9H0F6 | 3.36 | 1 | 1 | 1 | 1 | 3.36 | 1 | 1 | 387 | 39.9 | 5.80 |
| Q8IYU4 | 3.58 | 1 | 1 | 1 | 1 | 3.58 | 1 | 1 | 475 | 52.9 | 6.07 |
| Q14669 | 0.75 | 1 | 1 | 1 | 1 | 0.75 | 1 | 1 | 1992 | 220.3 | 8.48 |
| Q8NEG2 | 2.71 | 1 | 1 | 1 | 1 | 2.71 | 1 | 1 | 295 | 32.8 | 8.06 |
| P15498 | 0.83 | 1 | 1 | 1 | 1 | 0.83 | 1 | 1 | 845 | 98.3 | 6.62 |
| Q9H4P4 | 2.84 | 1 | 1 | 1 | 1 | 2.84 | 1 | 1 | 317 | 35.9 | 6.14 |
| Q5SRE7 | 4.12 | 1 | 1 | 1 | 1 | 4.12 | 1 | 1 | 291 | 32.4 | 6.32 |
| Q96QA5 | 3.37 | 1 | 1 | 1 | 1 | 3.37 | 1 | 1 | 445 | 49.3 | 5.29 |
| Q8IVH8 | 0.89 | 1 | 1 | 1 | 1 | 0.89 | 1 | 1 | 894 | 101.3 | 7.56 |
| Q9Y4H2 | 0.45 | 1 | 1 | 1 | 1 | 0.45 | 1 | 1 | 1338 | 137.2 | 8.65 |
| Q07890 | 0.75 | 1 | 1 | 1 | 1 | 0.75 | 1 | 1 | 1332 | 152.9 | 6.84 |
| Q53S58 | 2.25 | 1 | 1 | 1 | 1 | 2.25 | 1 | 1 | 311 | 33.7 | 9.61 |
| O14681 | 1.76 | 1 | 1 | 1 | 1 | 1.76 | 1 | 1 | 340 | 38.9 | 9.72 |
| Q13470 | 0.90 | 1 | 1 | 1 | 1 | 0.90 | 1 | 1 | 666 | 72.4 | 9.48 |
| O14879 | 3.47 | 1 | 1 | 1 | 1 | 3.47 | 1 | 1 | 490 | 55.9 | 5.20 |
| P55286 | 1.25 | 1 | 1 | 1 | 1 | 1.25 | 1 | 1 | 799 | 88.2 | 4.70 |
| Q5VSL9 | 1.67 | 1 | 1 | 1 | 1 | 1.67 | 1 | 1 | 837 | 95.5 | 6.29 |
| O00308 | 1.49 | 1 | 1 | 1 | 1 | 1.49 | 1 | 1 | 870 | 98.9 | 7.12 |
| Q9H1M4 | 13.13 | 1 | 1 | 1 | 1 | 13.13 | 1 | 1 | 99 | 11.3 | 8.75 |
| Q92903 | 3.90 | 1 | 1 | 1 | 1 | 3.90 | 1 | 1 | 461 | 53.3 | 8.09 |
| Q96PZ7 | 0.20 | 1 | 1 | 1 | 1 | 0.20 | 1 | 1 | 3565 | 388.6 | 6.07 |
| Q8NCE2 | 1.38 | 1 | 1 | 1 | 1 | 1.38 | 1 | 1 | 650 | 72.2 | 6.24 |
| Q96RK4 | 2.12 | 1 | 1 | 1 | 1 | 2.12 | 1 | 1 | 519 | 58.2 | 7.31 |
| O95427 | 1.29 | 1 | 1 | 1 | 1 | 1.29 | 1 | 1 | 931 | 105.7 | 8.65 |
| Q12879 | 1.71 | 1 | 1 | 1 | 1 | 1.71 | 1 | 1 | 1464 | 165.2 | 7.11 |
| Q6ZSS7 | 1.77 | 1 | 1 | 1 | 1 | 1.77 | 1 | 1 | 791 | 88.0 | 5.53 |
| Q15274 | 13.47 | 1 | 1 | 1 | 1 | 13.47 | 1 | 1 | 297 | 30.8 | 6.21 |
| P48546 | 1.50 | 2 | 1 | 1 | 1 | 1.50 | 1 | 1 | 466 | 53.1 | 8.78 |
| Q5TC82 | 0.53 | 1 | 1 | 1 | 1 | 0.53 | 1 | 1 | 1133 | 125.7 | 7.27 |
| Q9NPC7 | 1.15 | 1 | 1 | 1 | 1 | 1.15 | 1 | 1 | 610 | 68.6 | 8.25 |
| Q5T890 | 0.70 | 1 | 1 | 1 | 1 | 0.70 | 1 | 1 | 1561 | 177.0 | 8.60 |
| Q9Y4P8 | 2.20 | 1 | 1 | 1 | 1 | 2.20 | 1 | 1 | 454 | 49.4 | 6.46 |
| Q6ZVD7 | 1.01 | 1 | 1 | 1 | 1 | 1.01 | 1 | 1 | 989 | 110.9 | 7.78 |
| P43378 | 1.85 | 1 | 1 | 1 | 1 | 1.85 | 1 | 1 | 593 | 68.0 | 8.02 |
| Q9Y240 | 8.67 | 1 | 1 | 1 | 1 | 8.67 | 1 | 1 | 323 | 35.7 | 5.16 |
| A1KZ92 | 0.62 | 1 | 1 | 1 | 1 | 0.62 | 1 | 1 | 1463 | 163.6 | 7.43 |
| Q68DN1 | 0.30 | 1 | 1 | 1 | 1 | 0.30 | 1 | 1 | 1984 | 224.2 | 10.08 |
| Q96HY7 | 0.76 | 1 | 1 | 1 | 1 | 0.76 | 1 | 1 | 919 | 103.0 | 6.93 |
| Q8N5C6 | 0.80 | 1 | 1 | 1 | 1 | 0.80 | 1 | 1 | 995 | 111.7 | 8.72 |
| Q5KU26 | 1.48 | 1 | 1 | 1 | 1 | 1.48 | 1 | 1 | 742 | 81.5 | 5.69 |
| O60927 | 12.70 | 1 | 1 | 1 | 1 | 12.70 | 1 | 1 | 126 | 13.9 | 7.01 |
| O75354 | 2.48 | 1 | 1 | 1 | 1 | 2.48 | 1 | 1 | 484 | 53.2 | 9.26 |
| Q8IYA6 | 1.61 | 1 | 1 | 1 | 1 | 1.61 | 1 | 1 | 745 | 83.5 | 9.83 |
| Q96NT5 | 2.61 | 1 | 1 | 1 | 1 | 2.61 | 1 | 1 | 459 | 49.7 | 8.81 |
| Q9UBS8 | 2.11 | 1 | 1 | 1 | 1 | 2.11 | 1 | 1 | 474 | 53.8 | 4.75 |
| Q8NAT1 | 5.17 | 1 | 1 | 1 | 1 | 5.17 | 1 | 1 | 580 | 66.6 | 8.57 |
| Q8N2C7 | 0.18 | 1 | 1 | 1 | 1 | 0.18 | 1 | 1 | 3258 | 363.2 | 6.86 |
| Q14155 | 2.49 | 1 | 1 | 1 | 1 | 2.49 | 1 | 1 | 803 | 90.0 | 7.09 |
| Q08357 | 1.84 | 1 | 1 | 1 | 1 | 1.84 | 1 | 1 | 652 | 70.3 | 6.54 |
| Q9BQT8 | 4.01 | 1 | 1 | 1 | 1 | 4.01 | 1 | 1 | 299 | 33.3 | 9.51 |
| Q008S8 | 1.11 | 1 | 1 | 1 | 1 | 1.11 | 1 | 1 | 904 | 104.8 | 8.40 |
| Q9H330 | 1.43 | 1 | 1 | 1 | 1 | 1.43 | 1 | 1 | 911 | 100.9 | 8.87 |
| Q13042 | 1.45 | 1 | 1 | 1 | 1 | 1.45 | 1 | 1 | 620 | 71.6 | 5.85 |
| Q9BWC9 | 2.50 | 1 | 1 | 1 | 1 | 2.50 | 1 | 1 | 280 | 32.0 | 9.42 |
| Q53GL7 | 1.27 | 1 | 1 | 1 | 1 | 1.27 | 1 | 1 | 1025 | 109.9 | 4.97 |
| Q17R55 | 2.71 | 1 | 1 | 1 | 1 | 2.71 | 1 | 1 | 369 | 42.4 | 6.90 |
| A2RUB6 | 0.84 | 1 | 1 | 1 | 1 | 0.84 | 1 | 1 | 948 | 109.3 | 8.25 |
| Q8TB03 | 2.51 | 1 | 1 | 1 | 1 | 2.51 | 1 | 1 | 319 | 36.6 | 6.30 |
| P16383 | 0.90 | 1 | 1 | 1 | 1 | 0.90 | 1 | 1 | 781 | 89.3 | 5.99 |

| | | | | | | | | | | | |
|---|---|---|---|---|---|---|---|---|---|---|---|
| Q96AQ8 | 3.34 | 1 | 1 | 1 | 1 | 3.34 | 1 | 1 | 359 | 39.7 | 9.63 |
| P35475 | 1.07 | 1 | 1 | 1 | 1 | 1.07 | 1 | 1 | 653 | 72.6 | 9.14 |
| Q9H0R3 | 6.73 | 1 | 1 | 1 | 1 | 6.73 | 1 | 1 | 208 | 23.2 | 6.51 |
| Q86VR2 | 1.93 | 1 | 1 | 1 | 1 | 1.93 | 1 | 1 | 466 | 51.4 | 4.97 |
| Q9NQC7 | 0.94 | 1 | 1 | 1 | 1 | 0.94 | 1 | 1 | 956 | 107.2 | 5.59 |
| O95999 | 4.29 | 1 | 1 | 1 | 1 | 4.29 | 1 | 1 | 233 | 26.2 | 5.74 |
| Q9UHL0 | 3.52 | 1 | 1 | 1 | 1 | 3.52 | 1 | 1 | 483 | 54.7 | 6.28 |
| Q96M60 | 3.35 | 1 | 1 | 1 | 1 | 3.35 | 1 | 1 | 508 | 59.9 | 8.90 |
| O43314 | 0.56 | 1 | 1 | 1 | 1 | 0.56 | 1 | 1 | 1243 | 140.3 | 8.22 |
| P0C7P0 | 5.51 | 1 | 1 | 1 | 1 | 5.51 | 1 | 1 | 127 | 14.2 | 10.55 |
| Q9BZE7 | 3.23 | 1 | 1 | 1 | 1 | 3.23 | 1 | 1 | 217 | 24.9 | 9.80 |
| P61927 | 6.19 | 1 | 1 | 1 | 1 | 6.19 | 1 | 1 | 97 | 11.1 | 11.74 |
| P51512 | 1.15 | 1 | 1 | 1 | 1 | 1.15 | 1 | 1 | 607 | 69.5 | 8.54 |
| Q8N2U0 | 24.78 | 1 | 1 | 1 | 1 | 24.78 | 1 | 1 | 113 | 11.7 | 8.94 |
| Q9BSJ2 | 1.22 | 1 | 1 | 1 | 1 | 1.22 | 1 | 1 | 902 | 102.5 | 6.84 |
| P31431 | 6.06 | 1 | 1 | 1 | 1 | 6.06 | 1 | 1 | 198 | 21.6 | 4.50 |
| P07711 | 3.60 | 1 | 1 | 1 | 1 | 3.60 | 1 | 1 | 333 | 37.5 | 5.45 |
| O15533 | 1.79 | 1 | 1 | 1 | 1 | 1.79 | 1 | 1 | 448 | 47.6 | 7.15 |
| Q8N4N8 | 1.78 | 1 | 1 | 1 | 1 | 1.78 | 1 | 1 | 673 | 76.2 | 8.65 |
| F5GYI3 | 2.36 | 1 | 1 | 1 | 1 | 2.36 | 1 | 1 | 381 | 40.6 | 5.54 |
| Q96AQ1 | 5.03 | 2 | 1 | 1 | 1 | 5.03 | 1 | 1 | 378 | 41.6 | 10.52 |
| Q6ZR85 | 5.26 | 1 | 1 | 1 | 1 | 5.26 | 1 | 1 | 190 | 19.9 | 7.46 |
| Q5VWP3 | 1.75 | 1 | 1 | 1 | 1 | 1.75 | 1 | 1 | 458 | 50.4 | 6.38 |
| Q8NBL1 | 3.57 | 1 | 1 | 1 | 1 | 3.57 | 1 | 1 | 392 | 46.2 | 8.72 |
| Q9H1Z4 | 2.68 | 1 | 1 | 1 | 1 | 2.68 | 1 | 1 | 485 | 53.7 | 9.14 |
| P03897 | 13.04 | 1 | 1 | 1 | 1 | 13.04 | 1 | 1 | 115 | 13.2 | 4.44 |
| Q86TE4 | 1.73 | 3 | 1 | 1 | 1 | 1.73 | 1 | 1 | 346 | 38.9 | 8.73 |
| O00445 | 1.81 | 1 | 1 | 1 | 1 | 1.81 | 1 | 1 | 386 | 42.9 | 9.17 |
| Q9Y680 | 2.32 | 1 | 1 | 1 | 1 | 2.32 | 1 | 1 | 259 | 30.0 | 6.54 |
| P51178 | 1.46 | 1 | 1 | 1 | 1 | 1.46 | 1 | 1 | 756 | 85.6 | 6.70 |
| Q9P2F6 | 0.59 | 1 | 1 | 1 | 1 | 0.59 | 1 | 1 | 1191 | 132.5 | 7.99 |
| P10915 | 3.39 | 1 | 1 | 1 | 1 | 3.39 | 1 | 1 | 354 | 40.1 | 7.42 |
| Q14832 | 0.68 | 1 | 1 | 1 | 1 | 0.68 | 1 | 1 | 879 | 98.8 | 7.66 |
| Q14117 | 2.31 | 1 | 1 | 1 | 1 | 2.31 | 1 | 1 | 519 | 56.6 | 7.27 |
| Q96CN4 | 1.26 | 1 | 1 | 1 | 1 | 1.26 | 1 | 1 | 794 | 91.3 | 5.34 |
| Q00872 | 0.53 | 1 | 1 | 1 | 1 | 0.53 | 1 | 1 | 1141 | 128.2 | 6.04 |
| P21246 | 4.17 | 1 | 1 | 1 | 1 | 4.17 | 1 | 1 | 168 | 18.9 | 9.60 |
| O95185 | 0.75 | 1 | 1 | 1 | 1 | 0.75 | 1 | 1 | 931 | 103.1 | 6.10 |
| C9JBD0 | 4.69 | 1 | 1 | 1 | 1 | 4.69 | 1 | 1 | 128 | 14.9 | 9.36 |
| P09455 | 8.89 | 1 | 1 | 1 | 1 | 8.89 | 1 | 1 | 135 | 15.8 | 5.11 |
| Q8ND61 | 1.55 | 1 | 1 | 1 | 1 | 1.55 | 1 | 1 | 904 | 101.2 | 9.13 |
| Q969Q5 | 5.42 | 1 | 1 | 1 | 1 | 5.42 | 1 | 1 | 203 | 23.1 | 6.23 |
| Q14159 | 0.77 | 1 | 1 | 1 | 1 | 0.77 | 1 | 1 | 915 | 100.3 | 6.62 |
| O00221 | 2.80 | 1 | 1 | 1 | 1 | 2.80 | 1 | 1 | 500 | 52.8 | 6.68 |
| Q96KC2 | 6.15 | 2 | 1 | 1 | 1 | 6.15 | 1 | 1 | 179 | 20.4 | 6.52 |
| Q9H496 | 10.69 | 1 | 1 | 1 | 1 | 10.69 | 1 | 1 | 131 | 15.3 | 7.31 |
| Q14767 | 1.15 | 1 | 1 | 1 | 1 | 1.15 | 1 | 1 | 1821 | 194.9 | 5.19 |
| P52746 | 0.71 | 1 | 1 | 1 | 1 | 0.71 | 1 | 1 | 1687 | 187.8 | 7.91 |
| Q08426 | 2.49 | 1 | 1 | 1 | 1 | 2.49 | 1 | 1 | 723 | 79.4 | 9.14 |
| P0C2W1 | 3.85 | 1 | 1 | 1 | 1 | 3.85 | 1 | 1 | 286 | 30.6 | 7.83 |
| P51617 | 1.69 | 1 | 1 | 1 | 1 | 1.69 | 1 | 1 | 712 | 76.5 | 6.62 |
| Q8TEA1 | 1.49 | 1 | 1 | 1 | 1 | 1.49 | 1 | 1 | 469 | 51.7 | 7.96 |
| O14657 | 1.79 | 2 | 1 | 1 | 1 | 1.79 | 1 | 1 | 336 | 38.0 | 8.54 |
| Q5T160 | 2.08 | 1 | 1 | 1 | 1 | 2.08 | 1 | 1 | 578 | 65.5 | 8.21 |
| Q9UHK6 | 3.14 | 1 | 1 | 1 | 1 | 3.14 | 1 | 1 | 382 | 42.4 | 6.44 |
| Q9Y3D2 | 4.95 | 1 | 1 | 1 | 1 | 4.95 | 1 | 1 | 182 | 19.5 | 8.63 |
| Q9ULK2 | 1.74 | 1 | 1 | 1 | 1 | 1.74 | 1 | 1 | 861 | 91.5 | 9.79 |
| Q9NUT2 | 1.09 | 1 | 1 | 1 | 1 | 1.09 | 1 | 1 | 735 | 79.9 | 8.98 |
| Q96GX2 | 10.31 | 1 | 1 | 1 | 1 | 10.31 | 1 | 1 | 97 | 10.8 | 4.36 |
| Q9Y478 | 11.11 | 1 | 1 | 1 | 1 | 11.11 | 1 | 1 | 270 | 30.4 | 6.40 |
| Q8IUX4 | 2.41 | 1 | 1 | 1 | 1 | 2.41 | 1 | 1 | 373 | 45.0 | 7.23 |
| Q9Y3Q7 | 1.08 | 1 | 1 | 1 | 1 | 1.08 | 1 | 1 | 739 | 82.8 | 7.28 |
| Q01432 | 1.43 | 1 | 1 | 1 | 1 | 1.43 | 1 | 1 | 767 | 88.8 | 6.98 |
| P01019 | 3.51 | 1 | 1 | 1 | 1 | 3.51 | 1 | 1 | 485 | 53.1 | 6.32 |
| Q8WXJ9 | 8.14 | 1 | 1 | 1 | 1 | 8.14 | 1 | 1 | 295 | 34.3 | 8.79 |
| Q9H503 | 14.44 | 1 | 1 | 1 | 1 | 14.44 | 1 | 1 | 90 | 10.3 | 5.76 |
| Q16520 | 5.60 | 1 | 1 | 1 | 1 | 5.60 | 1 | 1 | 125 | 14.1 | 8.75 |
| P13497 | 0.91 | 1 | 1 | 1 | 1 | 0.91 | 1 | 1 | 986 | 111.2 | 6.90 |
| O14569 | 5.41 | 1 | 1 | 1 | 1 | 5.41 | 1 | 1 | 222 | 24.0 | 9.89 |
| Q9BXY5 | 1.26 | 1 | 1 | 1 | 1 | 1.26 | 1 | 1 | 557 | 63.8 | 8.66 |
| A6NN90 | 2.93 | 1 | 1 | 1 | 1 | 2.93 | 1 | 1 | 581 | 63.0 | 5.62 |
| Q8N5N4 | 6.38 | 1 | 1 | 1 | 1 | 6.38 | 1 | 1 | 141 | 15.7 | 9.85 |
| Q96KX1 | 14.53 | 1 | 1 | 1 | 1 | 14.53 | 1 | 1 | 117 | 13.3 | 9.47 |
| Q76N32 | 1.32 | 1 | 1 | 1 | 1 | 1.32 | 1 | 1 | 757 | 81.1 | 5.16 |
| Q6NUI6 | 1.05 | 1 | 1 | 1 | 1 | 1.05 | 1 | 1 | 762 | 82.3 | 8.84 |
| Q96D05 | 10.74 | 1 | 1 | 1 | 1 | 10.74 | 1 | 1 | 121 | 13.2 | 11.58 |
| Q9BRQ4 | 5.62 | 1 | 1 | 1 | 1 | 5.62 | 1 | 1 | 267 | 30.8 | 6.86 |
| O95476 | 4.51 | 1 | 1 | 1 | 1 | 4.51 | 1 | 1 | 244 | 28.4 | 9.76 |
| Q9GZY4 | 7.53 | 1 | 1 | 1 | 1 | 7.53 | 1 | 1 | 146 | 16.7 | 8.81 |
| Q86VX2 | 6.00 | 1 | 1 | 1 | 1 | 6.00 | 1 | 1 | 200 | 22.5 | 5.92 |
| Q6V0L0 | 2.11 | 1 | 1 | 1 | 1 | 2.11 | 1 | 1 | 522 | 57.1 | 9.00 |
| Q0P670 | 3.19 | 1 | 1 | 1 | 1 | 3.19 | 1 | 1 | 501 | 57.1 | 9.61 |
| Q9H4D0 | 0.63 | 1 | 1 | 1 | 1 | 0.63 | 1 | 1 | 955 | 106.9 | 5.39 |
| Q9NUD7 | 5.79 | 1 | 1 | 1 | 1 | 5.79 | 1 | 1 | 363 | 42.8 | 8.68 |
| Q8TF08 | 12.35 | 1 | 1 | 1 | 1 | 12.35 | 1 | 1 | 81 | 9.1 | 9.72 |
| Q96B18 | 1.11 | 1 | 1 | 1 | 1 | 1.11 | 1 | 1 | 629 | 64.9 | 10.36 |
| Q30KQ8 | 7.96 | 1 | 1 | 1 | 1 | 7.96 | 1 | 1 | 113 | 13.0 | 8.48 |
| Q92564 | 2.74 | 1 | 1 | 1 | 1 | 2.74 | 1 | 1 | 292 | 34.0 | 5.88 |
| Q6ZN92 | 5.67 | 1 | 1 | 1 | 1 | 5.67 | 1 | 1 | 141 | 15.5 | 7.64 |
| P78545 | 2.16 | 1 | 1 | 1 | 1 | 2.16 | 1 | 1 | 371 | 41.4 | 5.77 |
| Q9Y5L3 | 3.03 | 1 | 1 | 1 | 1 | 3.03 | 1 | 1 | 495 | 53.6 | 8.27 |
| Q7L0X2 | 2.26 | 1 | 1 | 1 | 1 | 2.26 | 1 | 1 | 663 | 75.2 | 4.93 |
| Q9NWS6 | 5.04 | 1 | 1 | 1 | 1 | 5.04 | 1 | 1 | 357 | 40.2 | 6.09 |
| Q6IPT2 | 4.86 | 1 | 1 | 1 | 1 | 4.86 | 1 | 1 | 247 | 27.6 | 8.65 |
| P23610 | 6.20 | 1 | 1 | 1 | 1 | 6.20 | 1 | 1 | 371 | 39.1 | 6.84 |
| A8MYZ6 | 1.22 | 1 | 1 | 1 | 1 | 1.22 | 1 | 1 | 492 | 50.6 | 5.45 |
| Q9BZS1 | 1.86 | 1 | 1 | 1 | 1 | 1.86 | 1 | 1 | 431 | 47.2 | 9.42 |
| Q9H469 | 9.33 | 1 | 1 | 1 | 1 | 9.33 | 1 | 1 | 300 | 33.0 | 7.44 |
| Q9NQ84 | 3.63 | 1 | 1 | 1 | 1 | 3.63 | 1 | 1 | 441 | 48.2 | 8.43 |
| P78333 | 2.45 | 1 | 1 | 1 | 1 | 2.45 | 1 | 1 | 572 | 63.7 | 6.81 |
| P86397 | 16.67 | 1 | 1 | 1 | 1 | 16.67 | 1 | 1 | 168 | 18.5 | 8.92 |
| Q9NQC8 | 4.61 | 1 | 1 | 1 | 1 | 4.61 | 1 | 1 | 304 | 34.3 | 4.54 |
| P01854 | 1.87 | 2 | 1 | 1 | 1 | 1.87 | 1 | 1 | 428 | 47.0 | 8.06 |
| Q9HBE4 | 5.81 | 1 | 1 | 1 | 1 | 5.81 | 1 | 1 | 155 | 17.9 | 9.31 |
| A8MTL0 | 7.43 | 1 | 1 | 1 | 1 | 7.43 | 1 | 1 | 148 | 18.0 | 10.01 |
| Q8WWA0 | 6.07 | 1 | 1 | 1 | 1 | 6.07 | 1 | 1 | 313 | 34.9 | 6.01 |
| O76095 | 8.90 | 1 | 1 | 1 | 1 | 8.90 | 1 | 1 | 146 | 16.3 | 8.28 |
| A6NE52 | 0.49 | 1 | 1 | 1 | 1 | 0.49 | 1 | 1 | 1622 | 180.2 | 7.61 |
| Q9BQ13 | 4.71 | 1 | 1 | 1 | 1 | 4.71 | 1 | 1 | 255 | 29.6 | 8.59 |
| Q4G0X4 | 5.38 | 1 | 1 | 1 | 1 | 5.38 | 1 | 1 | 260 | 29.6 | 6.54 |
| Q9Y4X4 | 3.73 | 1 | 1 | 1 | 1 | 3.73 | 1 | 1 | 402 | 44.2 | 9.70 |
| Q99612 | 3.53 | 1 | 1 | 1 | 1 | 3.53 | 1 | 1 | 283 | 31.8 | 6.92 |
| Q96S06 | 2.82 | 1 | 1 | 1 | 1 | 2.82 | 1 | 1 | 567 | 64.8 | 9.31 |
| Q9H1C0 | 2.69 | 1 | 1 | 1 | 1 | 2.69 | 1 | 1 | 372 | 41.3 | 10.18 |
| Q6ZNQ3 | 8.93 | 1 | 1 | 1 | 1 | 8.93 | 1 | 1 | 347 | 39.6 | 8.57 |
| Q6P1Q0 | 3.33 | 1 | 1 | 1 | 1 | 3.33 | 1 | 1 | 360 | 41.8 | 10.32 |
| A6BM72 | 0.77 | 1 | 1 | 1 | 1 | 0.77 | 1 | 1 | 1044 | 110.8 | 6.21 |
| Q5TF39 | 1.54 | 1 | 1 | 1 | 1 | 1.54 | 1 | 1 | 518 | 56.2 | 5.43 |
| Q86W26 | 1.22 | 1 | 1 | 1 | 1 | 1.22 | 1 | 1 | 655 | 75.0 | 7.20 |
| Q93015 | 4.20 | 1 | 1 | 1 | 1 | 4.20 | 1 | 1 | 286 | 31.4 | 7.85 |
| Q16649 | 1.52 | 1 | 1 | 1 | 1 | 1.52 | 1 | 1 | 462 | 51.4 | 7.18 |
| P48745 | 4.20 | 1 | 1 | 1 | 1 | 4.20 | 1 | 1 | 357 | 39.1 | 7.72 |
| Q9NQX5 | 2.77 | 1 | 1 | 1 | 1 | 2.77 | 1 | 1 | 325 | 34.5 | 7.01 |
| Q8NH19 | 2.33 | 1 | 1 | 1 | 1 | 2.33 | 1 | 1 | 301 | 34.1 | 8.78 |
| P30954 | 2.19 | 1 | 1 | 1 | 1 | 2.19 | 1 | 1 | 320 | 35.9 | 9.07 |
| Q5BKU9 | 6.80 | 1 | 1 | 1 | 1 | 6.80 | 1 | 1 | 147 | 15.8 | 8.22 |
| A6NNC1 | 2.01 | 1 | 1 | 1 | 1 | 2.01 | 1 | 1 | 897 | 94.0 | 10.26 |
| Q15198 | 1.87 | 1 | 1 | 1 | 1 | 1.87 | 1 | 1 | 375 | 41.8 | 8.50 |
| O75594 | 11.73 | 1 | 1 | 1 | 1 | 11.73 | 1 | 1 | 196 | 21.7 | 8.59 |
| Q7Z6Z6 | 2.10 | 1 | 1 | 1 | 1 | 2.10 | 1 | 1 | 429 | 47.9 | 6.77 |
| Q9Y2Y8 | 4.89 | 1 | 1 | 1 | 1 | 4.89 | 1 | 1 | 225 | 25.4 | 4.81 |
| Q9UKY0 | 5.11 | 1 | 1 | 1 | 1 | 5.11 | 1 | 1 | 176 | 20.3 | 9.03 |
| O43490 | 2.20 | 1 | 1 | 1 | 1 | 2.20 | 1 | 1 | 865 | 97.1 | 7.27 |
| A8MZF0 | 2.72 | 1 | 1 | 1 | 1 | 2.72 | 1 | 1 | 331 | 35.2 | 8.35 |
| Q9Y6I8 | 2.83 | 1 | 1 | 1 | 1 | 2.83 | 1 | 1 | 212 | 24.2 | 10.07 |
| Q16769 | 3.88 | 1 | 1 | 1 | 1 | 3.88 | 1 | 1 | 361 | 40.9 | 6.61 |
| A4D1S5 | 5.07 | 1 | 1 | 1 | 1 | 5.07 | 1 | 1 | 217 | 24.4 | 6.52 |
| Q7Z5H3 | 1.29 | 1 | 1 | 1 | 1 | 1.29 | 1 | 1 | 698 | 76.7 | 8.15 |
| A6NNX1 | 28.26 | 1 | 1 | 1 | 1 | 28.26 | 1 | 1 | 92 | 10.8 | 9.57 |
| Q9H920 | 2.75 | 1 | 1 | 1 | 1 | 2.75 | 1 | 1 | 327 | 37.9 | 7.06 |
| Q8WU17 | 1.96 | 1 | 1 | 1 | 1 | 1.96 | 1 | 1 | 664 | 75.9 | 6.44 |
| Q9Y3C5 | 7.79 | 1 | 1 | 1 | 1 | 7.79 | 1 | 1 | 154 | 17.4 | 4.78 |
| Q9BPW5 | 2.82 | 1 | 1 | 1 | 1 | 2.82 | 1 | 1 | 248 | 27.5 | 9.11 |
| Q9HAS3 | 2.17 | 1 | 1 | 1 | 1 | 2.17 | 1 | 1 | 691 | 76.9 | 7.62 |
| Q969S0 | 3.02 | 1 | 1 | 1 | 1 | 3.02 | 1 | 1 | 331 | 37.4 | 9.11 |
| Q9Y6D0 | 12.77 | 1 | 1 | 1 | 1 | 12.77 | 1 | 1 | 94 | 10.6 | 10.86 |
| Q86VE9 | 2.13 | 1 | 1 | 1 | 1 | 2.13 | 1 | 1 | 423 | 47.0 | 7.62 |
| P58005 | 3.46 | 1 | 1 | 1 | 1 | 3.46 | 1 | 1 | 492 | 57.3 | 6.25 |
| Q8IWL1 | 5.24 | 2 | 1 | 1 | 1 | 5.24 | 1 | 1 | 248 | 26.2 | 5.17 |
| Q8WV19 | 16.98 | 1 | 1 | 1 | 1 | 16.98 | 1 | 1 | 159 | 17.8 | 8.65 |
| O95562 | 6.25 | 1 | 1 | 1 | 1 | 6.25 | 1 | 1 | 160 | 17.8 | 9.13 |
| A6NKC9 | 1.55 | 1 | 1 | 1 | 1 | 1.55 | 1 | 1 | 451 | 49.8 | 6.42 |
| Q08ET2 | 2.27 | 2 | 1 | 1 | 1 | 2.27 | 1 | 1 | 396 | 43.9 | 7.96 |
| Q9BVQ7 | 2.26 | 1 | 1 | 1 | 1 | 2.26 | 1 | 1 | 753 | 80.7 | 8.09 |
| Q6RVD6 | 10.48 | 1 | 1 | 1 | 1 | 10.48 | 1 | 1 | 105 | 11.7 | 10.02 |
| Q96GX1 | 1.58 | 1 | 1 | 1 | 1 | 1.58 | 1 | 1 | 697 | 76.8 | 5.24 |
| Q6PEX7 | 8.25 | 1 | 1 | 1 | 1 | 8.25 | 1 | 1 | 206 | 23.3 | 6.27 |
| P04216 | 9.32 | 1 | 1 | 1 | 1 | 9.32 | 1 | 1 | 161 | 17.9 | 8.73 |
| Q99595 | 12.87 | 1 | 1 | 1 | 1 | 12.87 | 1 | 1 | 171 | 18.0 | 7.87 |
| Q8N9M5 | 2.17 | 1 | 1 | 1 | 1 | 2.17 | 1 | 1 | 508 | 54.1 | 5.91 |
| Q5U3C3 | 6.06 | 1 | 1 | 1 | 1 | 6.06 | 1 | 1 | 297 | 33.5 | 8.16 |
| Q9H6L2 | 2.22 | 1 | 1 | 1 | 1 | 2.22 | 1 | 1 | 316 | 36.0 | 7.94 |
| Q8TBM7 | 8.94 | 1 | 1 | 1 | 1 | 8.94 | 1 | 1 | 123 | 14.2 | 9.67 |
| Q5JXX7 | 19.64 | 1 | 1 | 1 | 1 | 19.64 | 1 | 1 | 168 | 19.8 | 6.54 |
| Q6UXN7 | 6.58 | 1 | 1 | 1 | 1 | 6.58 | 1 | 1 | 152 | 17.7 | 8.59 |
| Q96J77 | 6.43 | 1 | 1 | 1 | 1 | 6.43 | 1 | 1 | 140 | 15.5 | 9.95 |
| Q68CL5 | 5.00 | 1 | 1 | 1 | 1 | 5.00 | 1 | 1 | 300 | 33.3 | 6.73 |
| O94972 | 0.62 | 1 | 1 | 1 | 1 | 0.62 | 1 | 1 | 964 | 107.8 | 5.15 |
| Q9NRE2 | 0.58 | 1 | 1 | 1 | 1 | 0.58 | 1 | 1 | 1034 | 114.9 | 7.83 |
| P35442 | 1.02 | 1 | 1 | 1 | 1 | 1.02 | 1 | 1 | 1172 | 129.9 | 4.83 |
| O14604 | 27.27 | 1 | 1 | 1 | 1 | 27.27 | 1 | 1 | 44 | 5.0 | 5.36 |
| Q13432 | 3.75 | 2 | 1 | 1 | 1 | 3.75 | 1 | 1 | 240 | 26.9 | 6.42 |
| Q8IX04 | 2.55 | 1 | 1 | 1 | 1 | 2.55 | 1 | 1 | 471 | 52.2 | 7.09 |
| Q8IV63 | 2.32 | 1 | 1 | 1 | 1 | 2.32 | 1 | 1 | 474 | 52.8 | 9.04 |
| Q8NEX6 | 16.09 | 1 | 1 | 1 | 1 | 16.09 | 1 | 1 | 87 | 10.3 | 8.40 |
| Q9Y484 | 2.78 | 1 | 1 | 1 | 1 | 2.78 | 1 | 1 | 360 | 39.8 | 7.14 |
| O96014 | 1.98 | 1 | 1 | 1 | 1 | 1.98 | 1 | 1 | 354 | 39.2 | 8.69 |
| Q9H1B5 | 1.16 | 1 | 1 | 1 | 1 | 1.16 | 1 | 1 | 865 | 96.7 | 8.22 |
| P58557 | 6.59 | 1 | 1 | 1 | 1 | 6.59 | 1 | 1 | 167 | 19.3 | 7.55 |
| A6XGL0 | 4.35 | 1 | 1 | 1 | 1 | 4.35 | 1 | 1 | 299 | 32.6 | 6.09 |
| Q9HBF4 | 1.16 | 1 | 1 | 1 | 1 | 1.16 | 1 | 1 | 777 | 87.1 | 7.42 |
| Q14590 | 1.49 | 1 | 1 | 1 | 1 | 1.49 | 1 | 1 | 738 | 83.9 | 8.56 |
| P51522 | 1.16 | 1 | 1 | 1 | 1 | 1.16 | 1 | 1 | 516 | 59.7 | 9.22 |
| Q5SWL8 | 2.09 | 1 | 1 | 1 | 1 | 2.09 | 1 | 1 | 479 | 55.2 | 7.71 |

SUPPMENTARY TABLE 2, SHEET 1

| Accession | Description | ΣCoverage | Σ# Proteins | Unique Pepti | Σ# Peptides | Σ# PSMs | Exp1_01 Coverage A4 |
|---|---|---|---|---|---|---|---|
| Q9NQR4 | Omega-amidase NIT2 OS=Ho | 18.12 | 1 | 4 | 4 | 5 | 0 |
| Q8TCU4 | Alstrom syndrome protein 1 O | 1.3 | 1 | 3 | 3 | 6 | 0 |
| Q5HYA8 | Meckelin OS=Homo sapiens G | 2.31 | 1 | 2 | 2 | 2 | 0 |
| Q9H7X7 | Intraflagellar transport protein | 33.51 | 1 | 4 | 4 | 5 | 0 |
| Q14980 | Nuclear mitotic apparatus prot | 2.17 | 1 | 4 | 4 | 4 | 0 |
| Q5VYK3 | Proteasome-associated protei | 21.46 | 1 | 32 | 32 | 48 | 0.65 |
| Q9UPT5 | Exocyst complex component 7 | 10.2 | 1 | 5 | 5 | 7 | 0 |
| P42858 | Huntingtin OS=Homo sapiens | 1.34 | 1 | 3 | 3 | 4 | 0 |
| Q9NRY4 | Rho GTPase-activating protei | 1.33 | 1 | 2 | 2 | 2 | 0 |
| O60733 | 85/88 kDa calcium-independe | 32.51 | 2 | 18 | 18 | 29 | 0 |
| Q14008 | Cytoskeleton-associated prote | 1.87 | 1 | 3 | 3 | 5 | 0 |
| Q9NTK5 | Obg-like ATPase 1 OS=Homo | 14.9 | 1 | 5 | 5 | 16 | 3.79 |
| O75800 | Zinc finger MYND domain-con | 25.68 | 1 | 9 | 9 | 10 | 0 |
| Q9Y3P9 | Rab GTPase-activating protei | 5.52 | 1 | 5 | 5 | 7 | 0 |
| P22694 | cAMP-dependent protein kina | 38.46 | 1 | 2 | 15 | 53 | 13.68 |
| Q8IY31 | Intraflagellar transport protein | 40.91 | 1 | 4 | 4 | 6 | 10.61 |
| Q8TD19 | Serine/threonine-protein kinas | 5.62 | 1 | 4 | 4 | 7 | 0 |
| Q8TF46 | DIS3-like exonuclease 1 OS=H | 1.8 | 1 | 2 | 2 | 2 | 0 |
| Q8WW35 | Tctex1 domain-containing pro | 38.73 | 1 | 3 | 3 | 4 | 0 |
| Q8IVU9 | Uncharacterized protein C10o | 26.44 | 1 | 5 | 5 | 6 | 0 |
| Q8WYA0 | Intraflagellar transport protein | 15.68 | 1 | 11 | 11 | 11 | 0 |
| Q13485 | Mothers against decapentaple | 5.43 | 1 | 2 | 2 | 2 | 0 |
| Q02241 | Kinesin-like protein KIF23 OS= | 2.71 | 1 | 2 | 2 | 3 | 0 |
| Q15172 | Serine/threonine-protein phos | 25.93 | 1 | 9 | 11 | 20 | 0 |
| Q15691 | Microtubule-associated protei | 23.13 | 1 | 4 | 4 | 4 | 0 |
| Q5SW79 | Centrosomal protein of 170 kD | 1.77 | 2 | 3 | 3 | 3 | 0 |
| Q13099 | Intraflagellar transport protein | 2.04 | 1 | 2 | 2 | 2 | 0 |
| A1XBS5 | Protein FAM92A1 OS=Homo s | 14.53 | 1 | 4 | 4 | 4 | 0 |
| Q9UN37 | Vacuolar protein sorting-assoc | 18.54 | 2 | 5 | 7 | 9 | 0 |
| Q14145 | Kelch-like ECH-associated pro | 3.04 | 1 | 2 | 2 | 2 | 0 |
| Q15018 | BRISC complex subunit Abro1 | 16.63 | 1 | 6 | 6 | 10 | 2.65 |
| A0AVF1 | Intraflagellar transport protein | 9.75 | 1 | 4 | 4 | 5 | 0 |
| Q96AJ1 | Clusterin-associated protein 1 | 20.58 | 2 | 8 | 9 | 15 | 0 |
| Q8WXH0 | Nesprin-2 OS=Homo sapiens | 0.52 | 1 | 3 | 4 | 4 | 0 |
| Q66GS9 | Centrosomal protein of 135 kD | 6.93 | 4 | 6 | 8 | 11 | 0 |
| Q9BW83 | Intraflagellar transport protein | 56.99 | 1 | 7 | 7 | 14 | 0 |
| P49768 | Presenilin-1 OS=Homo sapien | 18.42 | 2 | 6 | 6 | 12 | 0 |
| P23258 | Tubulin gamma-1 chain OS=H | 39.02 | 2 | 9 | 9 | 18 | 0 |
| O75695 | Protein XRP2 OS=Homo sapi | 5.14 | 1 | 2 | 2 | 3 | 0 |
| Q8IYX3 | Coiled-coil domain-containing | 30.68 | 1 | 12 | 12 | 47 | 0 |
| O75116 | Rho-associated protein kinase | 9.01 | 2 | 6 | 10 | 10 | 0 |
| Q8TC44 | POC1 centriolar protein homo | 12.76 | 1 | 5 | 5 | 6 | 0 |
| Q3SYG4 | Protein PTHB1 OS=Homo sap | 4.06 | 1 | 2 | 2 | 3 | 0 |
| Q99996 | A-kinase anchor protein 9 OS | 0.69 | 2 | 2 | 4 | 9 | 0 |
| Q9UPM9 | B9 domain-containing protein | 13.24 | 1 | 2 | 2 | 2 | 0 |
| Q9Y366 | Intraflagellar transport protein | 18.08 | 1 | 5 | 5 | 7 | 0 |
| P42025 | Beta-centractin OS=Homo sap | 48.94 | 1 | 6 | 14 | 33 | 11.7 |
| O43237 | Cytoplasmic dynein 1 light inte | 27.85 | 1 | 8 | 8 | 12 | 0 |
| Q9NWB7 | Intraflagellar transport protein | 22.61 | 1 | 7 | 7 | 13 | 0 |
| P60510 | Serine/threonine-protein phos | 31.27 | 1 | 6 | 8 | 17 | 0 |
| Q9ULC3 | Ras-related protein Rab-23 O | 30.38 | 1 | 5 | 5 | 9 | 0 |
| A6NC98 | Coiled-coil domain-containing | 1.96 | 3 | 2 | 3 | 3 | 0 |
| P11388 | DNA topoisomerase 2-alpha C | 7.51 | 1 | 10 | 10 | 11 | 0 |
| Q9Y5Y2 | Cytosolic Fe-S cluster assemb | 54.61 | 1 | 8 | 8 | 20 | 0 |
| P49841 | Glycogen synthase kinase-3 b | 32.38 | 1 | 6 | 10 | 13 | 0 |
| O75604 | Ubiquitin carboxyl-terminal hyd | 24.79 | 1 | 9 | 9 | 11 | 0 |
| Q9P2H3 | Intraflagellar transport protein | 8.88 | 1 | 5 | 5 | 6 | 0 |
| O15078 | Centrosomal protein of 290 kD | 1.53 | 4 | 4 | 5 | 6 | 0 |
| Q92834 | X-linked retinitis pigmentosa G | 7.35 | 1 | 5 | 5 | 5 | 0 |
| Q502W7 | Coiled-coil domain-containing | 11.55 | 1 | 5 | 6 | 7 | 0 |
| Q15154 | Pericentriolar material 1 protei | 3.9 | 1 | 6 | 6 | 6 | 0 |
| Q8TCT7 | Signal peptide peptidase-like | 8.28 | 1 | 4 | 4 | 8 | 0 |
| Q9P2L0 | WD repeat-containing protein | 15.07 | 1 | 13 | 13 | 27 | 0 |
| Q9Y5K8 | V-type proton ATPase subuni | 31.58 | 1 | 6 | 6 | 7 | 0 |
| Q9BVG8 | Kinesin-like protein KIFC3 OS | 3.96 | 2 | 2 | 3 | 5 | 0 |
| Q08211 | ATP-dependent RNA helicase | 8.35 | 1 | 8 | 8 | 19 | 0 |
| P36404 | ADP-ribosylation factor-like pro | 50 | 1 | 7 | 7 | 10 | 0 |
| Q99828 | Calcium and integrin-binding p | 19.37 | 1 | 3 | 3 | 3 | 0 |
| Q76KD6 | Speriolin OS=Homo sapiens G | 8.46 | 1 | 4 | 4 | 5 | 0 |
| P53990 | IST1 homolog OS=Homo sapi | 18.41 | 1 | 7 | 7 | 20 | 1.92 |
| Q8TCI5 | Protein pitchfork OS=Homo sa | 26.18 | 1 | 4 | 4 | 10 | 0 |
| Q8IZT6 | Abnormal spindle-like microcep | 0.4 | 1 | 3 | 3 | 3 | 0 |
| Q96RY7 | Intraflagellar transport protein | 15.46 | 1 | 19 | 19 | 26 | 0 |
| Q8IWZ6 | Bardet-Biedl syndrome 7 prote | 14.13 | 1 | 8 | 8 | 8 | 0 |
| Q8TDR0 | TRAF3-interacting protein 1 O | 2.75 | 1 | 2 | 2 | 2 | 0 |
| O75665 | Oral-facial-digital syndrome 1 p | 3.66 | 1 | 3 | 3 | 7 | 1.28 |
| Q8TAP6 | Centrosomal protein of 76 kDa | 8.5 | 1 | 5 | 5 | 10 | 0 |
| Q96C92 | Serologically defined colon ca | 9.89 | 1 | 4 | 4 | 5 | 0 |
| O94886 | CSC1-like protein 1 OS=Homo | 5.33 | 2 | 5 | 5 | 11 | 0 |
| P30622 | CAP-Gly domain-containing lin | 5.84 | 1 | 7 | 7 | 7 | 0 |
| Q9Y6D5 | Brefeldin A-inhibited guanine | 27.23 | 1 | 28 | 41 | 71 | 2.3 |
| Q9UNZ2 | NSFL1 cofactor p47 OS=Hom | 40 | 1 | 12 | 12 | 33 | 17.03 |
| P17987 | T-complex protein 1 subunit a | 78.78 | 1 | 37 | 37 | 295 | 44.96 |
| P37198 | Nuclear pore glycoprotein p62 | 23.95 | 1 | 10 | 10 | 50 | 12.07 |
| P25786 | Proteasome subunit alpha typ | 74.14 | 1 | 19 | 19 | 54 | 65.4 |
| Q9Y6A4 | UPF0468 protein C16orf80 O | 74.61 | 1 | 15 | 15 | 49 | 6.22 |
| Q02750 | Dual specificity mitogen-activa | 34.35 | 1 | 7 | 13 | 30 | 0 |
| P60953 | Cell division control protein 42 | 62.83 | 3 | 8 | 9 | 37 | 14.14 |
| O75935 | Dynactin subunit 3 OS=Homo | 50.54 | 1 | 8 | 8 | 11 | 0 |
| Q9Y6G9 | Cytoplasmic dynein 1 light inte | 18.36 | 1 | 8 | 8 | 19 | 1.72 |
| Q8N865 | Uncharacterized protein C7orf | 61.36 | 3 | 31 | 31 | 118 | 0 |
| Q13561 | Dynactin subunit 2 OS=Homo | 46.63 | 1 | 17 | 17 | 68 | 27.43 |
| P63167 | Dynein light chain 1, cytoplasr | 65.17 | 1 | 4 | 7 | 108 | 59.55 |
| Q9UJT2 | Testis-specific serine kinase su | 30.41 | 1 | 17 | 18 | 35 | 0 |
| P36405 | ADP-ribosylation factor-like pro | 71.98 | 1 | 12 | 12 | 27 | 6.04 |
| Q9Y230 | RuvB-like 2 OS=Homo sapien | 73.87 | 1 | 40 | 40 | 260 | 34.13 |
| Q8IYX7 | Protein FAM154A OS=Homo s | 87.34 | 1 | 44 | 44 | 279 | 20.46 |
| A5D8V7 | Coiled-coil domain-containing | 46.55 | 2 | 25 | 26 | 49 | 0 |
| O43379 | WD repeat-containing protein | 8.04 | 1 | 9 | 9 | 20 | 0 |
| P29966 | Myristoylated alanine-rich C-ki | 33.73 | 2 | 6 | 6 | 23 | 20.78 |
| P61006 | Ras-related protein Rab-8A O | 48.79 | 7 | 4 | 12 | 86 | 24.64 |
| O43805 | Ras-related protein Rab-8A O | 36.97 | 1 | 5 | 5 | 12 | 20.17 |
| P35222 | Catenin beta-1 OS=Homo sap | 17.8 | 1 | 7 | 11 | 32 | 0 |
| P48729 | Casein kinase I isoform alpha | 36.8 | 1 | 4 | 10 | 19 | 2.97 |
| P28074 | Proteasome subunit beta type | 61.22 | 2 | 15 | 15 | 50 | 32.32 |
| Q14203 | Dynactin subunit 1 OS=Homo | 50.47 | 2 | 57 | 58 | 161 | 10.09 |
| O75955 | Flotillin-1 OS=Homo sapiens G | 61.12 | 1 | 22 | 22 | 50 | 17.33 |
| P17612 | cAMP-dependent protein kina | 55.27 | 1 | 6 | 20 | 76 | 30.77 |
| Q14204 | Cytoplasmic dynein 1 heavy c | 47.27 | 3 | 200 | 201 | 618 | 0.65 |
| Q9BZV1 | UBX domain-containing protei | 49.43 | 1 | 18 | 18 | 61 | 0 |
| P50990 | T-complex protein 1 subunit th | 72.26 | 1 | 45 | 45 | 322 | 45.26 |
| O60271 | C-Jun-amino-terminal kinase-ir | 5.98 | 2 | 6 | 6 | 11 | 0 |
| P50991 | T-complex protein 1 subunit d | 73.65 | 1 | 38 | 39 | 263 | 31.91 |
| Q13464 | Rho-associated protein kinase | 12.33 | 1 | 14 | 17 | 19 | 0 |
| Q12798 | Centrin-1 OS=Homo sapiens G | 52.33 | 2 | 10 | 10 | 21 | 5.23 |
| Q9Y265 | RuvB-like 1 OS=Homo sapien | 70.61 | 1 | 28 | 28 | 142 | 39.04 |
| Q9UJW0 | Dynactin subunit 4 OS=Homo | 11.09 | 1 | 4 | 4 | 9 | 0 |
| Q9NRG9 | Aladin OS=Homo sapiens GN | 42.86 | 1 | 15 | 15 | 28 | 0 |
| P43034 | Platelet-activating factor acety | 41.95 | 1 | 16 | 17 | 69 | 1.71 |
| Q9BTW9 | Tubulin-specific chaperone D | 30.2 | 1 | 27 | 27 | 57 | 0 |
| Q9HD42 | Charged multivesicular body p | 12.76 | 1 | 3 | 3 | 4 | 0 |
| P62826 | GTP-binding nuclear protein R | 51.39 | 1 | 12 | 12 | 72 | 9.26 |
| P04899 | Guanine nucleotide-binding p | 56.06 | 4 | 10 | 17 | 92 | 23.38 |
| P15311 | Ezrin OS=Homo sapiens GN= | 54.95 | 2 | 25 | 38 | 182 | 18.26 |
| P13861 | cAMP-dependent protein kina | 78.96 | 2 | 33 | 33 | 242 | 57.43 |
| Q5VTH2 | UPF0740 protein C1orf192 O | 59.89 | 1 | 10 | 10 | 48 | 0 |
| Q9Y224 | UPF0568 protein C14orf166 C | 43.03 | 1 | 9 | 9 | 10 | 0 |
| P05783 | Keratin, type I cytoskeletal 18 | 43.49 | 13 | 17 | 19 | 57 | 9.53 |
| Q5T4S7 | E3 ubiquitin-protein ligase UB | 20.66 | 3 | 79 | 80 | 204 | 0.68 |
| P62081 | 40S ribosomal protein S7 OS= | 60.82 | 1 | 10 | 10 | 24 | 5.67 |
| P43487 | Ran-specific GTPase-activatin | 43.28 | 2 | 7 | 7 | 22 | 5.47 |
| P06748 | Nucleophosmin OS=Homo sap | 12.93 | 2 | 5 | 5 | 10 | 0 |
| P40121 | Macrophage-capping protein C | 37.36 | 1 | 13 | 13 | 38 | 6.9 |
| Q9Y547 | Heat shock protein beta-11 O | 36.11 | 1 | 3 | 3 | 14 | 0 |
| P61421 | V-type proton ATPase subuni | 43.59 | 2 | 11 | 11 | 24 | 13.11 |
| P50570 | Dynamin-2 OS=Homo sapiens | 23.45 | 3 | 16 | 16 | 31 | 0 |
| P53384 | Cytosolic Fe-S cluster assemb | 24.69 | 1 | 5 | 5 | 11 | 0 |
| P53985 | Monocarboxylate transporter 1 | 17.8 | 1 | 8 | 8 | 37 | 2.4 |
| P33176 | Kinesin-1 heavy chain OS=Ho | 46.42 | 1 | 29 | 41 | 111 | 2.28 |
| Q96PF2 | Testis-specific serine/threonin | 37.15 | 1 | 11 | 11 | 33 | 5.87 |
| Q9H892 | Tetratricopeptide repeat prote | 25.96 | 1 | 16 | 16 | 27 | 0 |
| Q9H0I3 | Coiled-coil domain-containing | 33.95 | 1 | 15 | 15 | 28 | 0 |
| Q8IYE0 | Coiled-coil domain-containing | 30.99 | 1 | 29 | 29 | 51 | 0 |
| Q9C099 | Leucine-rich repeat and coiled | 13.76 | 1 | 12 | 12 | 17 | 0 |
| Q5BJF6 | Outer dense fiber protein 2 O | 65.86 | 3 | 97 | 98 | 1166 | 34.50 |
| P48643 | T-complex protein 1 subunit e | 71.72 | 1 | 42 | 43 | 346 | 53.42 |
| P61163 | Alpha-centractin OS=Homo sa | 61.17 | 1 | 10 | 18 | 69 | 34.31 |
| Q9UJC3 | Protein Hook homolog 1 OS=H | 39.56 | 1 | 26 | 26 | 63 | 0 |
| Q5JVL4 | EF-hand domain-containing p | 70.16 | 1 | 50 | 50 | 267 | 15.78 |
| Q92597 | Protein NDRG1 OS=Homo sap | 45.18 | 1 | 12 | 12 | 160 | 19.8 |
| Q2M329 | Coiled-coil domain-containing | 38.38 | 1 | 25 | 25 | 38 | 0 |
| Q00839 | Heterogeneous nuclear ribonu | 13.94 | 1 | 9 | 9 | 18 | 1.45 |
| Q13045 | Protein flightless-1 homolog O | 14.58 | 1 | 15 | 15 | 22 | 0 |
| Q9Y5B8 | Nucleoside diphosphate kinas | 76.86 | 1 | 26 | 26 | 95 | 8.51 |
| Q9Y696 | Chloride intracellular channel p | 61.66 | 4 | 9 | 11 | 46 | 2.37 |
| P08754 | Guanine nucleotide-binding p | 38.98 | 4 | 6 | 12 | 65 | 7.34 |
| A7E2U8 | UPF0602 protein C4orf47 OS | 64.08 | 1 | 15 | 15 | 51 | 0 |
| Q8WVJ2 | NudC domain-containing prote | 52.87 | 1 | 6 | 6 | 16 | 0 |
| Q96LB3 | Intraflagellar transport protein | 15.33 | 1 | 8 | 8 | 15 | 0 |
| Q8ND07 | Coiled-coil domain-containing | 30.81 | 1 | 15 | 15 | 18 | 0 |
| Q6ZN84 | Coiled-coil domain-containing | 50.61 | 1 | 37 | 37 | 59 | 0 |
| Q8WUM4 | Programmed cell death 6-inter | 56.91 | 1 | 48 | 48 | 208 | 13.36 |
| P08107;P0DMV8 | Heat shock 70 kDa protein 1A | 69.11 | 3 | 31 | 49 | 566 | 44.31 |
| P15531 | Nucleoside diphosphate kinas | 65.79 | 2 | 2 | 9 | 39 | 48.03 |
| Q9BZW7 | Testis-specific gene 10 protein | 70.34 | 1 | 61 | 61 | 152 | |
| O15066 | Kinesin-like protein KIF3B OS | 4.42 | 3 | 2 | 3 | 5 | 0.00 |
| O95721 | Synaptosomal-associated pro | 8.53 | 1 | 2 | 2 | 2 | 0.00 |
| Q6NXR4 | TELO2-interacting protein 2 O | 5.31 | 1 | 2 | 2 | 2 | |
| Q9H5N1 | Rab GTPase-binding effector | 8.44 | 1 | 5 | 5 | 10 | 1.76 |
| Q86WT1 | Tetratricopeptide repeat prote | 32.03 | 2 | 17 | 17 | 20 | |

| Accession | Exp1_01 Coverage A4 | # Peptides A4 | # PSM A4 | Exp02_01 Coverage B4 | # Peptides B4 | # PSM B4 | Exp03_01 Coverage C4 | # Peptides C4 | # PSM C4 | Exp01_02 Coverage D4 | # Peptides D4 | # PSM D4 | Exp02_02 Coverage E4 | # Peptides E4 | # PSM E4 | Exp03_02 Coverage F4 | # Peptides F4 | # PSM F4 | Exp01_03 Coverage G4 | # Peptides G4 | # PSM G4 | Exp02_03 Coverage H4 | # Peptides H4 | # PSM H4 | Exp03_03 Coverage I4 | # Peptides I4 | # PSM I4 | Exp04 Coverage J4 | # Peptides J4 | # PSM J4 | # AAs | MW [kDa] | calc. pI |
|---|---|---|---|---|---|---|---|---|---|---|---|---|---|---|---|---|---|---|---|---|---|---|---|---|---|---|---|---|---|---|---|---|---|
| Q9NQR4 | 0 | 0 | 0 | 0 | 0 | 0 | 0 | 0 | 0 | 0 | 0 | 0 | 0 | 0 | 0 | 0 | 0 | 0 | 0 | 0 | 0 | 0 | 0 | 0 | 0 | 0 | 0 | 18.12 | 4 | 5 | 276 | [illegible] | [illegible] |
| Q8TCU4 | 0 | 0 | 0 | 0 | 0 | 0 | 0 | 0 | 0 | 0 | 0 | 0 | 0 | 0 | 0 | 0 | 0 | 0 | 0 | 0 | 0 | 0 | 0 | 0 | 0 | 0 | 0 | 1.3 | 3 | 6 | 4167 | [illegible] | [illegible] |
| Q5HYA8 | 0 | 0 | 0 | 0 | 0 | 0 | 0 | 0 | 0 | 0 | 0 | 0 | 0 | 0 | 0 | 0 | 0 | 0 | 0 | 0 | 0 | 0 | 0 | 0 | 0 | 0 | 0 | 2.31 | 2 | 2 | 995 | [illegible] | [illegible] |
| Q9H7X7 | 0 | 0 | 0 | 0 | 0 | 0 | 0 | 0 | 0 | 0 | 0 | 0 | 0 | 0 | 0 | 0 | 0 | 0 | 0 | 0 | 0 | 0 | 0 | 0 | 0 | 0 | 0 | 33.51 | 4 | 5 | 185 | [illegible] | [illegible] |
| Q14980 | 0 | 0 | 0 | 0 | 0 | 0 | 0 | 0 | 0 | 0 | 0 | 0 | 0 | 0 | 0 | 0 | 0 | 0 | 0 | 0 | 0 | 0 | 0 | 0 | 0 | 0 | 0 | 2.17 | 4 | 4 | 2115 | [illegible] | [illegible] |
| Q5VYK3 | 0.65 | 1 | 1 | 0.65 | 1 | 1 | 0 | 0 | 0 | 0 | 0 | 0 | 0 | 0 | 0 | 0 | 0 | 0 | 0 | 0 | 0 | 0 | 0 | 0 | 0 | 0 | 0 | 20.81 | 31 | 46 | 1845 | [illegible] | [illegible] |
| Q9UPT5 | 0 | 0 | 0 | 0 | 0 | 0 | 0 | 0 | 0 | 0 | 0 | 0 | 0 | 0 | 0 | 0 | 0 | 0 | 0 | 0 | 0 | 0 | 0 | 0 | 0 | 0 | 0 | 10.2 | 5 | 7 | 735 | [illegible] | [illegible] |
| P42858 | 0 | 0 | 0 | 0 | 0 | 0 | 0 | 0 | 0 | 0 | 0 | 0 | 0 | 0 | 0 | 0 | 0 | 0 | 0 | 0 | 0 | 0 | 0 | 0 | 0 | 0 | 0 | 1.34 | 3 | 4 | 3142 | [illegible] | [illegible] |
| Q9NRY4 | 0 | 0 | 0 | 0 | 0 | 0 | 0 | 0 | 0 | 0 | 0 | 0 | 0 | 0 | 0 | 0 | 0 | 0 | 0 | 0 | 0 | 0 | 0 | 0 | 0 | 0 | 0 | 1.33 | 2 | 2 | 1499 | [illegible] | [illegible] |
| O60733 | 0 | 0 | 0 | 2.11 | 1 | 1 | 0 | 0 | 0 | 0 | 0 | 0 | 0 | 0 | 0 | 0 | 0 | 0 | 0 | 0 | 0 | 0 | 0 | 0 | 0 | 0 | 0 | 32.51 | 18 | 28 | 806 | [illegible] | [illegible] |
| Q14008 | 0 | 0 | 0 | 0 | 0 | 0 | 0 | 0 | 0 | 0 | 0 | 0 | 0 | 0 | 0 | 0 | 0 | 0 | 0 | 0 | 0 | 0 | 0 | 0 | 0 | 0 | 0 | 1.87 | 3 | 5 | 2032 | [illegible] | [illegible] |
| Q9NTK5 | 3.79 | 1 | 1 | 0 | 0 | 0 | 0 | 0 | 0 | 3.79 | 1 | 1 | 0 | 0 | 0 | 0 | 0 | 0 | 0 | 0 | 0 | 0 | 0 | 0 | 0 | 0 | 0 | 14.9 | 5 | 14 | 396 | [illegible] | [illegible] |
| O75800 | 0 | 0 | 0 | 0 | 0 | 0 | 0 | 0 | 0 | 0 | 0 | 0 | 0 | 0 | 0 | 0 | 0 | 0 | 0 | 0 | 0 | 0 | 0 | 0 | 0 | 0 | 0 | 25.68 | 9 | 10 | 440 | [illegible] | [illegible] |
| Q9Y3P9 | 0 | 0 | 0 | 0 | 0 | 0 | 0 | 0 | 0 | 0 | 0 | 0 | 0 | 0 | 0 | 0 | 0 | 0 | 0 | 0 | 0 | 0 | 0 | 0 | 0 | 0 | 0 | 5.52 | 5 | 7 | 1069 | [illegible] | [illegible] |
| P22694 | 13.68 | 4 | 4 | 13.68 | 4 | 5 | 7.41 | 3 | 3 | 0 | 0 | 0 | 0 | 0 | 0 | 0 | 0 | 0 | 0 | 0 | 0 | 0 | 0 | 0 | 2.56 | 1 | 1 | 38.46 | 15 | 40 | 351 | [illegible] | [illegible] |

*New PCM proteins identified in this work

- Uniprot
- Protein atlas
- Uniprot + Protein atlas
- GO term

**SUPPMENTARY TABLE 2, SHEET 2**

| Accession | Description | ΣCoverage | Σ# Proteins | Unique Peptid | Σ# Peptides | Σ# PSMs | Exp1_01 Coverage A4 | # Peptides A4 | # PSM A4 | Exp02_01 Coverage B4 | # Peptides B4 | # PSM B4 | Exp03_01 Coverage C4 | # Peptides C4 | # PSM C4 | Exp01_02 Coverage D4 | # Peptides D4 | # PSM D4 | Exp02_02 Coverage E4 | # Peptides E4 | # PSM E4 | Exp03_02 Coverage F4 | # Peptides F4 | # PSM F4 | Exp01_03 Coverage G4 | # Peptides G4 | # PSM G4 | Exp02_03 Coverage H4 | # Peptides H4 | # PSM H4 | Exp03_03 Coverage I4 | # Peptides I4 | # PSM I4 | Exp04 Coverage J4 | # Peptides J4 | # PSM J4 | # AAs | MW [kDa] | calc. pI |
|---|---|---|---|---|---|---|---|---|---|---|---|---|---|---|---|---|---|---|---|---|---|---|---|---|---|---|---|---|---|---|---|---|---|---|---|---|---|---|---|
| Q8TD16 | Protein bicaudal D homolog 2 OS=Homo sapiens GN=BICD2 PE=1 SV=1 - [BICD2_HUMAN] | 1.09 | 1 | 1 | 1 | 1 | | | | | | | | | | 1.09 | 1 | 1 | | | | | | | | | | | | | | | | 0.00 | | | 824 | 93.5 | 5.44 |
| Q5T890 | DNA excision repair protein ERCC-6-like 2 OS=Homo sapiens GN=ERCC6L2 PE=1 SV=2 - [ER6L2_ | 0.70 | 1 | 1 | 1 | 1 | | | | | | | | | | | | | | | | | | | | | | | | | | | | 0.70 | 1 | 1 | 1561 | 177.0 | 8.60 |
| Q9P1Y5 | Calmodulin-regulated spectrin-associated protein 3 OS=Homo sapiens GN=CAMSAP3 PE=1 SV=2 - | 0.56 | 1 | 1 | 1 | 1 | | | | | | | | | | | | | 0.00 | | | | | | | | | | | | | | | 0.56 | 1 | 1 | 1249 | 134.7 | 8.35 |
| Q8N4N8 | Kinesin-like protein KIF2B OS=Homo sapiens GN=KIF2B PE=1 SV=3 - [KIF2B_HUMAN] | 1.78 | 1 | 1 | 1 | 1 | | | | | | | | | | | | | | | | | | | | | | | | | | | | 1.78 | 1 | 1 | 673 | 76.2 | 8.65 |
| Q86YT6 | E3 ubiquitin-protein ligase MIB1 OS=Homo sapiens GN=MIB1 PE=1 SV=1 - [MIB1_HUMAN] | 0.60 | 1 | 1 | 1 | 1 | | | | 0.00 | | | | | | | | | | | | | | | | | | | | | | | | 0.60 | 1 | 1 | 1006 | 110.1 | 6.92 |
| O95613 | Pericentrin OS=Homo sapiens GN=PCNT PE=1 SV=4 - [PCNT_HUMAN] | 0.18 | 1 | 1 | 1 | 1 | | | | | | | 0.00 | | | | | | 0.00 | | | | | | | | | | | | | | | 0.18 | 1 | 1 | 3336 | 377.8 | 5.55 |
| Q9P219 | Protein Daple OS=Homo sapiens GN=CCDC88C PE=1 SV=3 - [DAPLE_HUMAN] | 0.54 | 1 | 1 | 1 | 1 | | | | | | | | | | | | | | | | | | | | | | 0.00 | | | | | | 0.54 | 1 | 1 | 2028 | 228.1 | 6.23 |
| Q15051 | IQ calmodulin-binding motif-containing protein 1 OS=Homo sapiens GN=IQCB1 PE=1 SV=1 - [IQCB | 2.01 | 1 | 1 | 1 | 1 | | | | | | | | | | | | | | | | | | | | | | | | | | | | 2.01 | 1 | 1 | 598 | 68.9 | 9.10 |
| Q92845 | Kinesin-associated protein 3 OS=Homo sapiens GN=KIFAP3 PE=1 SV=2 - [KIFA3_HUMAN] | 2.53 | 1 | 1 | 1 | 2 | | | | | | | | | | | | | | | | | | | | | | | | | | | | 2.53 | 1 | 2 | 792 | 91.1 | 5.08 |
| Q92830 | Histone acetyltransferase KAT2A OS=Homo sapiens GN=KAT2A PE=1 SV=3 - [KAT2A_HUMAN] | 0.72 | 1 | 1 | 1 | 1 | | | | | | | | | | | | | | | | | | | | | | | | | | | | 0.72 | 1 | 1 | 837 | 93.9 | 9.04 |
| Q9P2G4 | Microtubule-associated protein 10 OS=Homo sapiens GN=MAP10 PE=1 SV=2 - [MAP10_HUMAN] | 0.77 | 1 | 1 | 1 | 2 | 0.77 | 1 | 1 | | | | 0.77 | 1 | 1 | | | | | | | | | | | | | | | | | | | 0.00 | | | 905 | 100.3 | 7.17 |
| Q3V6T2 | Girdin OS=Homo sapiens GN=CCDC88A PE=1 SV=2 - [GRDN_HUMAN] | 0.64 | 1 | 1 | 1 | 2 | | | | | | | | | | 0.00 | | | 0.00 | | | | | | | | | | | | | | | 0.64 | 1 | 2 | 1871 | 215.9 | 6.21 |
| Q6DKK2 | Tetratricopeptide repeat protein 19, mitochondrial OS=Homo sapiens GN=TTC19 PE=1 SV=4 - [TTC | 2.63 | 1 | 1 | 1 | 1 | | | | | | | | | | | | | | | | | | | | | | | | | | | | 2.63 | 1 | 1 | 380 | 42.4 | 5.77 |
| P63096 | Guanine nucleotide-binding protein G(i) subunit alpha-1 OS=Homo sapiens GN=GNAI1 PE=1 SV=2 | 25.14 | 4 | 1 | 8 | 52 | 7.06 | 2 | 2 | | | | 12.43 | 4 | 5 | 12.43 | 4 | 4 | 3.11 | 1 | 1 | | | | | | | 3.11 | 1 | 1 | 14.41 | 5 | 7 | 21.19 | 7 | 32 | 354 | 40.3 | 5.97 |
| Q9NQC8 | Intraflagellar transport protein 46 homolog OS=Homo sapiens GN=IFT46 PE=1 SV=1 - [IFT46_HUM | 4.61 | 1 | 1 | 1 | 1 | | | | | | | | | | | | | | | | | | | | | | | | | | | | 4.61 | 1 | 1 | 304 | 34.3 | 4.54 |
| Q13432 | Protein unc-119 homolog A OS=Homo sapiens GN=UNC119 PE=1 SV=1 - [U119A_HUMAN] | 3.75 | 2 | 1 | 1 | 1 | | | | | | | | | | | | | | | | | | | | | | | | | | | | 3.75 | 1 | 1 | 240 | 26.9 | 6.42 |
| Q99661 | Kinesin-like protein KIF2C OS=Homo sapiens GN=KIF2C PE=1 SV=2 - [KIF2C_HUMAN] | 1.79 | 1 | 1 | 1 | 1 | 0.00 | | | | | | 0.00 | | | | | | 0.00 | | | | | | | | | 0.00 | | | 0.00 | | | 1.79 | 1 | 1 | 725 | 81.3 | 7.83 |
| Q14191 | Werner syndrome ATP-dependent helicase OS=Homo sapiens GN=WRN PE=1 SV=2 - [WRN_HUMA | 0.49 | 1 | 1 | 1 | 1 | | | | | | | | | | | | | | | | | | | | | | | | | | | | 0.49 | 1 | 1 | 1432 | 162.4 | 6.34 |
| Q9UQB9 | Aurora kinase C OS=Homo sapiens GN=AURKC PE=1 SV=1 - [AURKC_HUMAN] | 2.27 | 1 | 1 | 1 | 1 | | | | | | | | | | | | | | | | | | | | | | | | | | | | 2.27 | 1 | 1 | 309 | 35.6 | 8.81 |
| Q8IYA6 | Cytoskeleton-associated protein 2-like OS=Homo sapiens GN=CKAP2L PE=1 SV=4 - [CKP2L_HUM | 1.61 | 1 | 1 | 1 | 1 | | | | | | | | | | | | | | | | | | | | | | | | | | | | 1.61 | 1 | 1 | 745 | 83.5 | 9.83 |
| P14635 | G2/mitotic-specific cyclin-B1 OS=Homo sapiens GN=CCNB1 PE=1 SV=1 - [CCNB1_HUMAN] | 2.54 | 1 | 1 | 1 | 1 | | | | | | | | | | | | | | | | | | | | | | | | | | | | 2.54 | 1 | 1 | 433 | 48.3 | 7.47 |
| O14974 | Protein phosphatase 1 regulatory subunit 12A OS=Homo sapiens GN=PPP1R12A PE=1 SV=1 - [M | 1.17 | 1 | 1 | 1 | 1 | | | | | | | | | | | | | | | | | | | | | | | | | | | | 1.17 | 1 | 1 | 1030 | 115.2 | 5.40 |
| Q9P287 | BRCA2 and CDKN1A-interacting protein OS=Homo sapiens GN=BCCIP PE=1 SV=1 - [BCCIP_HUMA | 7.32 | 1 | 1 | 1 | 2 | | | | | | | | | | | | | | | | | | | | | | | | | | | | 7.32 | 1 | 2 | 314 | 36.0 | 4.61 |
| Q96RK4 | Bardet-Biedl syndrome 4 protein OS=Homo sapiens GN=BBS4 PE=1 SV=2 - [BBS4_HUMAN] | 2.12 | 1 | 1 | 1 | 1 | | | | | | | | | | | | | | | | | | | | | | | | | | | | 2.12 | 1 | 1 | 519 | 58.2 | 7.31 |
| Q7Z460 | CLIP-associating protein 1 OS=Homo sapiens GN=CLASP1 PE=1 SV=1 - [CLAP1_HUMAN] | 0.72 | 1 | 1 | 1 | 1 | | | | | | | | | | | | | 0.00 | | | | | | | | | | | | | | | 0.72 | 1 | 1 | 1538 | 169.3 | 9.03 |
| P12004 | Proliferating cell nuclear antigen OS=Homo sapiens GN=PCNA PE=1 SV=1 - [PCNA_HUMAN] | 2.68 | 1 | 1 | 1 | 1 | | | | | | | | | | | | | | | | | | | | | | | | | | | | 2.68 | 1 | 1 | 261 | 28.8 | 4.69 |
| Q3B820 | Protein FAM161A OS=Homo sapiens GN=FAM161A PE=1 SV=2 - [F161A_HUMAN] | 2.42 | 1 | 1 | 1 | 2 | | | | | | | | | | | | | | | | | | | | | | | | | | | | 2.42 | 1 | 2 | 660 | 76.7 | 8.03 |
| O76095 | Protein JTB OS=Homo sapiens GN=JTB PE=1 SV=1 - [JTB_HUMAN] | 8.90 | 1 | 1 | 1 | 1 | | | | | | | | | | | | | | | | | | | | | | | | | | | | 8.90 | 1 | 1 | 146 | 16.3 | 8.28 |
| Q53EZ4 | Centrosomal protein of 55 kDa OS=Homo sapiens GN=CEP55 PE=1 SV=3 - [CEP55_HUMAN] | 3.66 | 1 | 1 | 1 | 1 | 0.00 | | | 0.00 | | | | | | | | | 0.00 | | | | | | | | | | | | | | | 3.66 | 1 | 1 | 464 | 54.1 | 7.01 |
| Q9NY27 | Serine/threonine-protein phosphatase 4 regulatory subunit 2 OS=Homo sapiens GN=PPP4R2 PE=1 | 1.92 | 1 | 1 | 1 | 1 | | | | | | | | | | | | | | | | | | | | | | | | | | | | 1.92 | 1 | 1 | 417 | 46.9 | 4.54 |
| Q15650 | Activating signal cointegrator 1 OS=Homo sapiens GN=TRIP4 PE=1 SV=4 - [TRIP4_HUMAN] | 1.55 | 1 | 1 | 1 | 1 | | | | | | | | | | | | | | | | | | | | | | | | | | | | 1.55 | 1 | 1 | 581 | 66.1 | 7.85 |
| Q16513 | Serine/threonine-protein kinase N2 OS=Homo sapiens GN=PKN2 PE=1 SV=1 - [PKN2_HUMAN] | 0.91 | 1 | 1 | 1 | 1 | | | | | | | | | | | | | | | | | | | | | | | | | | | | 0.91 | 1 | 1 | 984 | 112.0 | 6.30 |
| Q13042 | Cell division cycle protein 16 homolog OS=Homo sapiens GN=CDC16 PE=1 SV=2 - [CDC16_HUMAN | 1.45 | 1 | 1 | 1 | 1 | | | | | | | | | | | | | | | | | | | | | | | | | | | | 1.45 | 1 | 1 | 620 | 71.6 | 5.85 |
| O95271 | Tankyrase-1 OS=Homo sapiens GN=TNKS PE=1 SV=2 - [TNKS1_HUMAN] | 0.60 | 1 | 1 | 1 | 2 | | | | | | | | | | | | | | | | | | | | | | | | | | | | 0.60 | 1 | 2 | 1327 | 142.0 | 7.05 |
| Q96CW5 | Gamma-tubulin complex component 3 OS=Homo sapiens GN=TUBGCP3 PE=1 SV=2 - [GCP3_HUM | 2.43 | 1 | 1 | 1 | 3 | | | | | | | | | | | | | | | | | | | | | | | | | | | | 2.43 | 1 | 3 | 907 | 103.5 | 8.12 |
| Q9UBK9 | Protein UXT OS=Homo sapiens GN=UXT PE=1 SV=1 - [UXT_HUMAN] | 7.01 | 1 | 1 | 1 | 1 | | | | | | | | | | | | | | | | | | | | | | | | | | | | 7.01 | 1 | 1 | 157 | 18.2 | 7.59 |
| P05062 | Fructose-bisphosphate aldolase B OS=Homo sapiens GN=ALDOB PE=1 SV=2 - [ALDOB_HUMAN] | 3.85 | 2 | 1 | 2 | 13 | 1.92 | 1 | 1 | 0.00 | | | 1.92 | 1 | 1 | | | | 1.92 | 1 | 1 | | | | 1.92 | 1 | 1 | 1.92 | 1 | 1 | 1.92 | 1 | 1 | 1.92 | 1 | 7 | 364 | 39.4 | 7.87 |
| Q9NQC7 | Ubiquitin carboxyl-terminal hydrolase CYLD OS=Homo sapiens GN=CYLD PE=1 SV=1 - [CYLD_HUM | 0.94 | 1 | 1 | 1 | 1 | | | | | | | | | | | | | | | | | | | | | | | | | | | | 0.94 | 1 | 1 | 956 | 107.2 | 5.59 |
| Q96RT7 | Gamma-tubulin complex component 6 OS=Homo sapiens GN=TUBGCP6 PE=1 SV=3 - [GCP6_HUM | 0.38 | 1 | 1 | 1 | 1 | | | | | | | | | | | | | | | | | | | | | | | | | 0.38 | 1 | 1 | | | | 1819 | 200.4 | 6.32 |
| Q76N32 | Centrosomal protein of 68 kDa OS=Homo sapiens GN=CEP68 PE=1 SV=2 - [CEP68_HUMAN] | 1.32 | 1 | 1 | 1 | 1 | | | | | | | | | | | | | | | | | | | | | | | | | | | | 1.32 | 1 | 1 | 757 | 81.1 | 5.16 |
| Q15398 | Disks large-associated protein 5 OS=Homo sapiens GN=DLGAP5 PE=1 SV=2 - [DLGP5_HUMAN] | 2.01 | 2 | 1 | 2 | 2 | | | | | | | | | | | | | | | | | | | | | | | | | | | | 2.01 | 2 | 2 | 846 | 95.1 | 9.00 |
| Q6ZVD7 | Storkhead-box protein 1 OS=Homo sapiens GN=STOX1 PE=1 SV=2 - [STOX1_HUMAN] | 1.01 | 1 | 1 | 1 | 1 | | | | | | | | | | | | | | | | | | | | | | | | | | | | 1.01 | 1 | 1 | 989 | 110.9 | 7.78 |
| Q9H0N0 | Ras-related protein Rab-6C OS=Homo sapiens GN=RAB6C PE=1 SV=2 - [RAB6C_HUMAN] | 11.02 | 1 | 1 | 2 | 10 | | | | 2.36 | 1 | 1 | | | | | | | | | | | | | | | | | | | | | | 11.02 | 2 | 9 | 254 | 28.3 | 7.71 |
| Q8N8E3 | Centrosomal protein of 112 kDa OS=Homo sapiens GN=CEP112 PE=1 SV=2 - [CE112_HUMAN] | 0.63 | 1 | 1 | 1 | 1 | | | | | | | | | | | | | | | | | | | | | | | | | | | | 0.63 | 1 | 1 | 955 | 112.7 | 6.68 |
| O75351 | Vacuolar protein sorting-associated protein 4B OS=Homo sapiens GN=VPS4B PE=1 SV=2 - [VPS4I | 8.78 | 2 | 1 | 3 | 7 | | | | | | | | | | | | | | | | | | | | | | | | | | | | 8.78 | 3 | 7 | 444 | 49.3 | 7.23 |
| A2RUB6 | Coiled-coil domain-containing protein 66 OS=Homo sapiens GN=CCDC66 PE=1 SV=4 - [CCD66_HU | 0.84 | 1 | 1 | 1 | 1 | | | | | | | | | | | | | | | | | | | | | | | | | | | | 0.84 | 1 | 1 | 948 | 109.3 | 8.25 |
| O15013 | Rho guanine nucleotide exchange factor 10 OS=Homo sapiens GN=ARHGEF10 PE=1 SV=4 - [ARH | 0.51 | 1 | 1 | 1 | 1 | | | | | | | | | | | | | | | | | | | | | | | | | | | | 0.51 | 1 | 1 | 1369 | 151.5 | 5.68 |
| O00139 | Kinesin-like protein KIF2A OS=Homo sapiens GN=KIF2A PE=1 SV=3 - [KIF2A_HUMAN] | 1.56 | 1 | 1 | 1 | 1 | | | | | | | | | | | | | | | | | | | | | | | | | | | | 1.56 | 1 | 1 | 706 | 79.9 | 6.68 |
| Q3YEC7 | Rab-like protein 6 OS=Homo sapiens GN=RABL6 PE=1 SV=2 - [RABL6_HUMAN] | 1.51 | 1 | 1 | 1 | 1 | | | | | | | | | | | | | | | | | | | | | | | | | | | | 1.51 | 1 | 1 | 729 | 79.5 | 5.22 |
| Q8NEZ2 | Vacuolar protein sorting-associated protein 37A OS=Homo sapiens GN=VPS37A PE=1 SV=1 - [VP3 | 2.02 | 1 | 1 | 1 | 1 | | | | | | | | | | | | | | | | | | | | | | | | | | | | 2.02 | 1 | 1 | 397 | 44.3 | 5.57 |
| Q96JB5 | CDK5 regulatory subunit-associated protein 3 OS=Homo sapiens GN=CDK5RAP3 PE=1 SV=2 - [CK | 3.95 | 2 | 1 | 2 | 5 | | | | 1.19 | 1 | 1 | | | | 1.19 | 1 | 1 | | | | | | | | | | | | | | | | 3.95 | 2 | 3 | 506 | 56.9 | 4.75 |
| Q8NHQ1 | Centrosomal protein of 70 kDa OS=Homo sapiens GN=CEP70 PE=1 SV=2 - [CEP70_HUMAN] | 1.51 | 1 | 1 | 1 | 1 | | | | | | | | | | | | | | | | | | | | | | | | | | | | 1.51 | 1 | 1 | 597 | 69.7 | 5.71 |
| Q9NXG0 | Centlein OS=Homo sapiens GN=CNTLN PE=2 SV=5 - [CNTLN_HUMAN] | 0.43 | 1 | 1 | 1 | 1 | 0.00 | | | | | | | | | | | | | | | | | | | | | | | | | | | 0.43 | 1 | 1 | 1405 | 161.5 | 8.15 |
| P62491 | Ras-related protein Rab-11A OS=Homo sapiens GN=RAB11A PE=1 SV=3 - [RB11A_HUMAN] | 62.50 | 1 | 1 | 14 | 71 | 29.17 | 6 | 7 | 29.17 | 5 | 6 | 20.83 | 5 | 5 | 29.63 | 6 | 6 | 5.09 | 1 | 1 | | | | 9.26 | 2 | 2 | 5.09 | 1 | 1 | 25.00 | 5 | 6 | 59.72 | 12 | 37 | 216 | 24.4 | 6.57 |
| Q9BSJ2 | Gamma-tubulin complex component 2 OS=Homo sapiens GN=TUBGCP2 PE=1 SV=2 - [GCP2_HUM | 1.22 | 1 | 1 | 1 | 1 | | | | | | | | | | | | | | | | | | | | | | | | | | | | 1.22 | 1 | 1 | 902 | 102.5 | 6.84 |
| O43809 | Cleavage and polyadenylation specificity factor subunit 5 OS=Homo sapiens GN=NUDT21 PE=1 SV | 7.93 | 1 | 1 | 1 | 1 | | | | | | | | | | | | | | | | | | | | | | | | | | | | 7.93 | 1 | 1 | 227 | 26.2 | 8.82 |
| P06493 | Cyclin-dependent kinase 1 OS=Homo sapiens GN=CDK1 PE=1 SV=3 - [CDK1_HUMAN] | 6.06 | 17 | 1 | 2 | 6 | 0.00 | | | 0.00 | | | 2.69 | 1 | 1 | | | | | | | | | | | | | | | | | | | 6.06 | 2 | 5 | 297 | 34.1 | 8.40 |
| P08F94 | Fibrocystin OS=Homo sapiens GN=PKHD1 PE=1 SV=1 - [PKHD1_HUMAN] | 0.27 | 1 | 1 | 1 | 1 | | | | | | | | | | | | | | | | | | | 0.27 | 1 | 1 | | | | | | | | | | 4074 | 446.4 | 6.57 |
| Q96CN5 | Leucine-rich repeat-containing protein 45 OS=Homo sapiens GN=LRRC45 PE=1 SV=1 - [LRC45_HU | 0.90 | 1 | 1 | 1 | 1 | | | | | | | | | | | | | | | | | | | | | | | | | | | | 0.90 | 1 | 1 | 670 | 75.9 | 6.23 |
| O94927 | HAUS augmin-like complex subunit 5 OS=Homo sapiens GN=HAUS5 PE=1 SV=2 - [HAUS5_HUMAN | 1.90 | 1 | 1 | 1 | 2 | | | | 0.00 | | | | | | | | | | | | | | | | | | | | | | | | 1.90 | 1 | 2 | 633 | 71.6 | 8.51 |
| Q9BVT8 | Transmembrane and ubiquitin-like domain-containing protein 1 OS=Homo sapiens GN=TMUB1 PE= | 4.07 | 1 | 1 | 1 | 2 | | | | | | | | | | | | | | | | | | | | | | | | | | | | 4.07 | 1 | 2 | 246 | 26.2 | 5.72 |
| O95714 | E3 ubiquitin-protein ligase HERC2 OS=Homo sapiens GN=HERC2 PE=1 SV=2 - [HERC2_HUMAN] | 0.52 | 1 | 1 | 1 | 1 | | | | | | | | | | | | | | | | | | | | | | | | | | | | 0.52 | 1 | 1 | 4834 | 526.9 | 6.28 |
| Q9BXF6 | Rab11 family-interacting protein 5 OS=Homo sapiens GN=RAB11FIP5 PE=1 SV=1 - [RFIP5_HUMAN | 2.76 | 1 | 1 | 1 | 2 | | | | | | | | | | | | | | | | | | | | | | | | | | | | 2.76 | 1 | 2 | 653 | 70.4 | 9.23 |
| Q9H1Z4 | WD repeat-containing protein 13 OS=Homo sapiens GN=WDR13 PE=1 SV=2 - [WDR13_HUMAN] | 2.68 | 1 | 1 | 1 | 1 | | | | | | | | | | | | | | | | | | | | | | | | | | | | 2.68 | 1 | 1 | 485 | 53.7 | 9.14 |
| Q8TD31 | Coiled-coil alpha-helical rod protein 1 OS=Homo sapiens GN=CCHCR1 PE=1 SV=2 - [CCHCR_HUMA | 0.90 | 1 | 1 | 1 | 1 | | | | | | | | | | | | | | | | | | | | | | | | | | | | 0.90 | 1 | 1 | 782 | 88.6 | 5.95 |
| Q6IQ55 | Tau-tubulin kinase 2 OS=Homo sapiens GN=TTBK2 PE=1 SV=2 - [TTBK2_HUMAN] | 0.72 | 1 | 1 | 1 | 1 | | | | | | | | | | | | | | | | | | | | | | | | | 0.00 | | | 0.72 | 1 | 1 | 1244 | 137.3 | 7.01 |
| O95359 | Transforming acidic coiled-coil-containing protein 2 OS=Homo sapiens GN=TACC2 PE=1 SV=3 - [TA | 0.47 | 1 | 1 | 1 | 1 | | | | 0.00 | | | | | | | | | | | | | | | | | | | | | 0.47 | 1 | 1 | | | | 2948 | 309.2 | 4.79 |
| Q9UFC0 | Leucine-rich repeat and WD repeat-containing protein 1 OS=Homo sapiens GN=LRWD1 PE=1 SV=2 | 2.16 | 1 | 1 | 1 | 2 | | | | | | | | | | | | | | | | | | | | | | | | | | | | 2.16 | 1 | 2 | 647 | 70.8 | 7.21 |
| Q9UPY8 | Microtubule-associated protein RP/EB family member 3 OS=Homo sapiens GN=MAPRE3 PE=1 SV= | 2.85 | 1 | 1 | 1 | 1 | | | | | | | | | | | | | | | | | | | | | | | | | | | | 2.85 | 1 | 1 | 281 | 32.0 | 5.54 |
| Q04759 | Protein kinase C theta type OS=Homo sapiens GN=PRKCQ PE=1 SV=3 - [KPCT_HUMAN] | 2.12 | 5 | 1 | 2 | 4 | 0.99 | 1 | 1 | | | | | | | 0.99 | 1 | 1 | | | | | | | 0.99 | 1 | 1 | | | | | | | 1.13 | 1 | 1 | 706 | 81.8 | 7.61 |
| Q86VQ0 | Lebercilin OS=Homo sapiens GN=LCA5 PE=1 SV=2 - [LCA5_HUMAN] | 0.86 | 2 | 1 | 1 | 1 | | | | | | | | | | | | | | | | | | | | | | | | | | | | 0.86 | 1 | 1 | 697 | 80.5 | 7.68 |
| O75923 | Dysferlin OS=Homo sapiens GN=DYSF PE=1 SV=1 - [DYSF_HUMAN] | 2.12 | 1 | 1 | 4 | 5 | 0.77 | 1 | 1 | | | | | | | | | | | | | | | | | | | | | | | | | 1.35 | 3 | 4 | 2080 | 237.1 | 5.64 |
| Q03518 | Antigen peptide transporter 1 OS=Homo sapiens GN=TAP1 PE=1 SV=2 - [TAP1_HUMAN] | 1.36 | 1 | 1 | 1 | 1 | | | | | | | | | | | | | | | | 0.00 | | | | | | | | | | | | 1.36 | 1 | 1 | 808 | 87.2 | 8.02 |
| O00592 | Podocalyxin OS=Homo sapiens GN=PODXL PE=1 SV=2 - [PODXL_HUMAN] | 2.33 | 1 | 1 | 1 | 2 | | | | | | | | | | | | | | | | | | | | | | | | | | | | 2.33 | 1 | 2 | 558 | 58.6 | 5.49 |
| Q9P2P6 | StAR-related lipid transfer protein 9 OS=Homo sapiens GN=STARD9 PE=1 SV=3 - [STAR9_HUMAN | 0.51 | 3 | 1 | 3 | 5 | 0.00 | | | 0.00 | | | 0.00 | | | | | | | | | | | | | | | | | | | | | 0.51 | 3 | 5 | 4700 | 516.0 | 6.32 |
| Q96EX3 | WD repeat-containing protein 34 OS=Homo sapiens GN=WDR34 PE=1 SV=2 - [WDR34_HUMAN] | 1.31 | 1 | 1 | 1 | 1 | | | | | | | | | | | | | | | | | | | | | | | | | | | | 1.31 | 1 | 1 | 536 | 57.8 | 6.64 |
| Q6IN85 | Serine/threonine-protein phosphatase 4 regulatory subunit 3A OS=Homo sapiens GN=SMEK1 PE= | 0.84 | 1 | 1 | 1 | 2 | | | | | | | | | | | | | | | | | | | | | | | | | | | | 0.84 | 1 | 2 | 833 | 95.3 | 4.94 |
| Q96JN8 | Neuralized-like protein 4 OS=Homo sapiens GN=NEURL4 PE=1 SV=2 - [NEUL4_HUMAN] | 0.70 | 1 | 1 | 1 | 1 | | | | | | | 0.00 | | | | | | | | | | | | | | | | | | | | | 0.70 | 1 | 1 | 1562 | 166.8 | 5.86 |
| Q7Z3E5 | LisH domain-containing protein ARMC9 OS=Homo sapiens GN=ARMC9 PE=1 SV=2 - [ARMC9_HUM | 1.84 | 1 | 1 | 1 | 1 | | | | | | | | | | | | | | | | | | | | | | | | | | | | 1.84 | 1 | 1 | 817 | 91.7 | 6.20 |
| Q8TDX7 | Serine/threonine-protein kinase Nek7 OS=Homo sapiens GN=NEK7 PE=1 SV=1 - [NEK7_HUMAN] | 2.65 | 2 | 1 | 1 | 1 | | | | | | | | | | | | | | | | | | | | | | | | | | | | 2.65 | 1 | 1 | 302 | 34.5 | 8.25 |
| P50402 | Emerin OS=Homo sapiens GN=EMD PE=1 SV=1 - [EMD_HUMAN] | 3.94 | 1 | 1 | 1 | 2 | | | | | | | | | | | | | | | | | | | | | | | | | | | | 3.94 | 1 | 2 | 254 | 29.0 | 5.50 |

*New PCM proteins identified in this work

Uniprot

Protein atlas

Uniprot + Protein atlas

**SUPPMENTARY TABLE 2, SHEET 3**

| 2 unique peptides: | | | |
|---|---|---|---|
| Accession | Description | Uniprot description | References |
| Q8TCU4 | Alstrom syndrome protein 1 OS=Homo sapiens GN=ALMS1 PE=1 SV=3 - [ALMS1_HUMAN] | Involved in PCM1-dependent intracellular transport. Required, directly or indirectly, for the **localization of NCAPD2 to the proximal ends of centrioles**. Required for proper formation and/or maintenance of primary cilia (PC), microtubule-based structures that protrude from the surface of epithelial cells. | 10.1038/nature02166. Proteomic characterization of the human centrosome by protein correlation profiling |
| Q5HYA8 | Meckelin OS=Homo sapiens GN=TMEM67 PE=1 SV=2 - [MKS3_HUMAN] | Required for ciliary structure and function. Part of the tectonic-like complex which is required for tissue-specific ciliogenesis and may regulate ciliary membrane composition (By similarity). Involved in centrosome migration to the apical cell surface during early ciliogenesis. Involved in the **regulation of cilia length and appropriate number through the control of centrosome duplication**. Required for cell branching morphology. Essential for endoplasmic reticulum-associated degradation (ERAD) of surfactant protein C (SFTPC). | 10.1093/hmg/ddl459. The Meckel-Gruber Syndrome Proteins MKS1 and Meckelin Interact and Are Required for Primary Cilium Formation. 10.1093/hmg/ddp272. Ciliary and Centrosomal Defects Associated With Mutation and Depletion of the Meckel Syndrome Genes MKS1 and MKS3. 10.1074/jbc.M109.034371. Meckel-Gruber Syndrome Protein MKS3 Is Required for Endoplasmic Reticulum-Associated Degradation of Surfactant Protein C. 10.1242/jcs.043794. Nesprin-2 Interacts With Meckelin and Mediates Ciliogenesis via Remodelling of the Actin Cytoskeleton. |
| Q14980 | Nuclear mitotic apparatus protein 1 OS=Homo sapiens GN=NUMA1 PE=1 SV=2 - [NUMA1_HUMAN] | Microtubule (MT)-binding protein that plays a role in the formation and maintenance of the spindle poles and the alignement and the segregation of chromosomes during mitotic cell division (PubMed:7769006, PubMed:17172455, PubMed:19255246, PubMed:24996901, PubMed:26195665, PubMed:27462074). Functions to tether the minus ends of MTs at the spindle poles, which is critical for the establishment and maintenance of the spindle poles (PubMed:12445386, PubMed:11956313). Plays a role in the establishment of the mitotic spindle orientation during metaphase and elongation during anaphase in a dynein-dynactin-dependent manner (PubMed:23870127, PubMed:24109598, PubMed:24996901, PubMed:26765568). In metaphase, part of a ternary complex composed of GPSM2 and G(i) alpha proteins, that regulates the recruitment and anchorage of the dynein-dynactin complex in the mitotic cell cortex regions situated above the two spindle poles, and hence regulates the correct oritentation of the mitotic spindle (PubMed:23027904, PubMed:22327364, PubMed:23921553). During anaphase, mediates the recruitment and accumulation of the dynein-dynactin complex at the cell membrane of the polar cortical region through direct association with phosphatidylinositol 4,5-bisphosphate (PI(4,5)P2), and hence participates in the regulation of the spindle elongation and chromosome segregation (PubMed:22327364, PubMed:23921553, PubMed:24996901, PubMed:24371089). Binds also to other polyanionic phosphoinositides, such as phosphatidylinositol 3-phosphate (PIP), lysophosphatidic acid (LPA) and phosphatidylinositol triphosphate (PIP3), in vitro (PubMed:24996901, PubMed:24371089). Also required for proper orientation of the mitotic spindle during asymmetric cell divisions (PubMed:21816348). Plays a role in mitotic MT aster assembly (PubMed:11163243, PubMed:11229403, PubMed:12445386). Involved in anastral spindle assembly (PubMed:25657325). Positively regulates TNKS protein localization to spindle poles in mitosis (PubMed:16076287). Highly abundant component of the nuclear matrix where it may serve a non-mitotic structural role, occupies the majority of the nuclear volume (PubMed:10075938). Required for epidermal differentiation and hair follicle morphogenesis | 10.1083/jcb.116.6.1303. NuMA: An Unusually Long Coiled-Coil Related Protein in the Mammalian Nucleus. 10.1083/jcb.116.6.1395. Primary Structure of NuMA, an Intranuclear Protein That Defines a Novel Pathway for Segregation of Proteins at Mitosis. 10.1083/jcb.149.4.851. Formation of Spindle Poles by Dynein/Dynactin-Dependent Transport of NuMA. 10.1091/mbc.E14-09-1366. CENP-32 Is Required to Maintain **Centrosomal** Dominance in Bipolar Spindle Assembly. 10.1371/journal.pone.0142798. Polyglutamylated Tubulin Binding Protein C1orf96/CSAP Is Involved in Microtubule Stabilization in Mitotic Spindles. 10.7554/eLife.12504. NuMA-microtubule Interactions Are Critical for Spindle Orientation and the Morphogenesis of Diverse Epidermal Structures. |
| Q14008 | Cytoskeleton-associated protein 5 OS=Homo sapiens GN=CKAP5 PE=1 SV=3 - [CKAP5_HUMAN]. Alternative name: Colonic and hepatic tumor overexpressed gene protein Ch-TOG | Binds to the plus end of microtubules and regulates microtubule dynamics and microtubule organization. Acts as processive microtubule polymerase. Promotes cytoplasmic microtubule nucleation and elongation. Plays a major role in organizing spindle poles. In spindle formation protects kinetochore microtubules from depolymerization by KIF2C and has an essential role in centrosomal microtubule assembly independently of KIF2C activity. **Contributes to centrosome integrity.** Acts as component of the TACC3/ch-TOG/clathrin complex proposed to contribute to stabilization of kinetochore fibers of the mitotic spindle by acting as inter-microtubule bridge. The TACC3/ch-TOG/clathrin complex is required for the maintenance of kinetochore fiber tension (PubMed:23532825). Enhances the strength of NDC80 complex-mediated kinetochore-tip microtubule attachments (PubMed:27156448). | 10.1083/jcb.201209107. The UBXN-2/p37/p47 Adaptors of CDC-48/p97 Regulate Mitosis by Limiting the Centrosomal Recruitment of Aurora A. The TOGp protein is a new human microtubule-associated protein homologous to the Xenopus XMAP215 |
| Q9NTK5 | Obg-like ATPase 1 OS=Homo sapiens GN=OLA1 PE=1 SV=2 - [OLA1_HUMAN] | Hydrolyzes ATP, and can also hydrolyze GTP with lower efficiency. Has lower affinity for GTP. **Binds to γ-tubulin and localized to centrosomes** in interphase and to the spindle pole in mitotic phase. OLA1 knockdown results in centrosome amplification and the activation of microtubule aster formation. | 10.1016/j.molcel.2013.10.028. The BRCA1/BARD1-interacting Protein OLA1 Functions in Centrosome Regulation |
| Q9Y3P9 | Rab GTPase-activating protein 1 OS=Homo sapiens GN=RABGAP1 PE=1 SV=3 - [RBGP1_HUMAN] | May act as a GTPase-activating protein of RAB6A. May play a role in microtubule nucleation by centrosome. May participate in a RAB6A-mediated pathway involved in the metaphase-anaphase transition. | 10.1093/emboj/18.7.1772. Characterization of GAPCenA, a GTPase Activating Protein for Rab6, **Part of Which Associates With the Centrosome** |
| Q8IY31 | Intraflagellar transport protein 20 homolog OS=Homo sapiens GN=IFT20 PE=1 SV=1 - [IFT20_HUMAN] | Part of intraflagellar transport (IFT) particles involved in ciliary process assembly. May play a role in the trafficking of ciliary membrane proteins from the Golgi complex to the cilium (PubMed:16775004). Regulates the platelet-derived growth factor receptor-alpha (PDGFRA) signaling pathway. Required for protein stability of E3 ubiquitin ligases CBL and CBLB that mediate ubiquitination and internalization of PDGFRA for proper feedback inhibition of PDGFRA signaling (PubMed:29237719). Essential for male fertility. Plays an important role in spermatogenesis, particularly spermiogenesis, when germ cells form flagella. May play a role in the transport of flagellar proteins ODF2 and SPAG16 to build sperm flagella and in the removal of redundant sperm cytoplasm (By similarity). Also involved in autophagy since it is required for trafficking of ATG16L and the expansion of the autophagic compartment. Present at the centrosomes during the cell cycle and **associated with the proximal portion of the mother centriole and the lateral aspect of the daughter centriole.** | 10.1083/jcb.200405023. Pericentrin Forms a Complex With Intraflagellar Transport Proteins and polycystin-2 and Is Required for Primary Cilia Assembly |
| Q8TD19 | Serine/threonine-protein kinase Nek9 OS=Homo sapiens GN=NEK9 PE=1 SV=2 - [NEK9_HUMAN] | Pleiotropic regulator of mitotic progression, participating in the control of spindle dynamics and chromosome separation. Phosphorylates different histones, myelin basic protein, beta-casein, and BICD2. Phosphorylates histone H3 on serine and threonine residues and beta-casein on serine residues. Important for G1/S transition and S phase progression. Phosphorylates NEK6 and NEK7 and stimulates their activity by releasing the autoinhibitory functions of Tyr-108 and Tyr-97 respectively | 10.1016/j.cub.2017.11.046. Nek9 Phosphorylation Defines a New Role for TPX2 in Eg5-Dependent Centrosome Separation Before Nuclear Envelope Breakdown. 10.1016/j.cub.2012.06.027. Nek9 Phosphorylation of NEDD1/GCP-WD Contributes to Plk1 Control of γ-Tubulin Recruitment to the Mitotic Centrosome. 10.1038/emboj.2011.179. **Nek9 Is a Plk1-activated Kinase That Controls Early Centrosome Separation Through Nek6/7** and Eg5 |
| Q8IVU9 | Uncharacterized protein C10orf107 OS=Homo sapiens GN=C10orf107 PE=2 SV=1 - [CJ107_HUMAN] | Calcium-binding protein. May be involved in the control of sperm flagellar movement. Colocalized **with pericentrin at centrosome of spermatocytes and round spermatids**. | 10.1002/mrd.22639. CABCOCO1, a Novel Coiled-Coil Protein With Calcium-Binding Activity, Is Localized in the Sperm Flagellum |
| Q15172 | Serine/threonine-protein phosphatase 2A 56 kDa regulatory subunit alpha isoform OS=Homo sapiens GN=PPP2R5A PE=1 SV=1 - [2A5A_HUMAN] | The B regulatory subunit might modulate substrate selectivity and catalytic activity, and also might direct the localization of the catalytic enzyme to a particular subcellular compartment. **Centrosome localization** | 10.1074/jbc.M109.093294. Nuclear Export and Centrosome Targeting of the Protein Phosphatase 2A Subunit B56alpha: Role of B56alpha in Nuclear Export of the Catalytic Subunit |
| Q15691 | Microtubule-associated protein RP/EB family member 1 OS=Homo sapiens GN=MAPRE1 PE=1 SV=3 - [MARE1_HUMAN] | Plus-end tracking protein (+TIP) that binds to the plus-end of microtubules and regulates the dynamics of the microtubule cytoskeleton (PubMed:12388762, PubMed:16109370, PubMed:19632184, PubMed:21646404, PubMed:28726242, PubMed:28814570). Promotes cytoplasmic microtubule nucleation and elongation (PubMed:12388762, PubMed:16109370, PubMed:19632184, PubMed:21646404, PubMed:28726242, PubMed:28814570). May be involved in spindle function by stabilizing **microtubules and anchoring them at centrosomes** (PubMed:12388762). Also acts as a regulator of minus-end microtubule organization: interacts with the complex formed by AKAP9 and PDE4DIP, leading to recruit CAMSAP2 to the Golgi apparatus, thereby tethering non-centrosomal minus-end microtubules to the Golgi, an important step for polarized cell movement (PubMed:28814570). Promotes elongation of CAMSAP2-decorated microtubule stretches on the minus-end of microtubules (PubMed:28814570). Acts as a regulator of autophagosome transport via interaction with CAMSAP2 (PubMed:28726242). May play a role in cell migration (By similarity). | 10.1091/mbc.e02-01-0061. Evidence That an Interaction Between EB1 and p150(Glued) Is Required for the Formation and Maintenance of a Radial Microtubule Array Anchored at the Centrosome. 10.1038/nature02166. Proteomic Characterization of the Human Centrosome by Protein Correlation Profiling |
| Q5SW79 | Centrosomal protein of 170 kDa OS=Homo sapiens GN=CEP170 PE=1 SV=1 - [CE170_HUMAN] | Plays a role in microtubule organization (PubMed:15616186). Required for **centriole subdistal appendage assembly** (PubMed:28422092). | 10.1091/mbc.e04-10-0939. The Forkhead-Associated Domain Protein Cep170 Interacts With Polo-like Kinase 1 and Serves as a Marker for Mature Centrioles. 10.1038/ncomms15057. Hierarchical Assembly of Centriole Subdistal Appendages via Centrosome Binding Proteins CCDC120 and CCDC68. |
| Q13099 | Intraflagellar transport protein 88 homolog OS=Homo sapiens GN=IFT88 PE=2 SV=2 - [IFT88_HUMAN] | Involved in primary cilium biogenesis. Also involved in autophagy since it is required for trafficking of ATG16L and the expansion of the autophagic compartment. **IFT88 is tightly associated with the centrosome** throughout the cell cycle in a microtubule- and dynein-independent manner. | 10.1242/jcs.03366. The intraflagellar transport component IFT88/polaris is a centrosomal protein regulating G1-S transition in non-ciliated cells |
| A1XBS5 | Protein FAM92A1 OS=Homo sapiens GN=FAM92A1 PE=1 SV=2 - [F92A1_HUMAN] | Acts as a positive regulator of ciliary hedgehog signaling (By similarity). Probable regulator of ciliogenesis involved in limb morphogenesis. In cooperation with CBY1 it is involved in the recruitment and fusion of endosomal vesicles at distal appendages during early stages of ciliogenesis. **Localizes at the distal end of mother centrioles** (PubMed:27528616). Extensive colocalization with CBY1 at mother centrioles (PubMed:27528616). | 10.1128/MCB.00160-16. BAR Domain-Containing FAM92 Proteins Interact With Chibby1 To Facilitate Ciliogenesis |
| Q9UN37 | Vacuolar protein sorting-associated protein 4A OS=Homo sapiens GN=VPS4A PE=1 SV=1 - [VPS4A_HUMAN] | Involved in late steps of the endosomal multivesicular bodies (MVB) pathway. Recognizes membrane-associated ESCRT-III assemblies and catalyzes their disassembly, possibly in combination with membrane fission. Redistributes the ESCRT-III components to the cytoplasm for further rounds of MVB sorting. MVBs contain intraluminal vesicles (ILVs) that are generated by invagination and scission from the limiting membrane of the endosome and mostly are delivered to lysosomes enabling degradation of membrane proteins, such as stimulated growth factor receptors, lysosomal enzymes and lipids. In conjunction with the ESCRT machinery also appears to function in topologically equivalent membrane fission events, such as the terminal stages of cytokinesis and enveloped virus budding (HIV-1 and other lentiviruses). Involved in cytokinesis: retained at the midbody by ZFYVE19/ANCHR and CHMP4C until abscission checkpoint signaling is terminated at late cytokinesis. It is then released following dephosphorylation of CHMP4C, leading to abscission (PubMed:24814515). VPS4A/B are required for the exosomal release of SDCBP, CD63 and syndecan (PubMed:22660413). VPS4 is a **dynamic component of both mother and daughter centrioles.** | 10.1038/s41598-018-21491-x. VPS4 Is a **Dynamic Component of the Centrosome That Regulates Centrosome Localization of γ-Tubulin, Centriolar Satellite Stability and Ciliogenesis.** 10.1073/pnas.1005938107. Human ESCRT-III and VPS4 Proteins Are Required for Centrosome and Spindle Maintenance. |
| Q8WXH0 | Nesprin-2 OS=Homo sapiens GN=SYNE2 PE=1 SV=3 - [SYNE2_HUMAN] | Multi-isomeric modular protein which forms a linking network between organelles and the actin cytoskeleton to maintain the subcellular spatial organization. As a component of the LINC (LInker of Nucleoskeleton and Cytoskeleton) complex involved in the connection between the nuclear lamina and the cytoskeleton. The nucleocytoplasmic interactions established by the LINC complex play an important role in the transmission of mechanical forces across the nuclear envelope and in nuclear movement and positioning. Specifically, SYNE2 and SUN2 assemble in arrays of transmembrane actin-associated nuclear (TAN) lines which are bound to F-actin cables and couple the nucleus to retrograde actin flow during actin-dependent nuclear movement. **May be involved in nucleus-centrosome attachment.** During interkinetic nuclear migration (INM) at G2 phase and nuclear migration in neural progenitors its LINC complex association with SUN1/2 and probable association with cytoplasmic dynein-dynactin motor complexes functions to pull the nucleus toward the centrosome; SYNE1 and SYNE2 may act redundantly. During INM at G1 phase mediates respective LINC complex association with kinesin to push the nucleus away from the centrosome. Involved in nuclear migration in retinal photoreceptor progenitors. Required for centrosome migration to the apical cell surface during early ciliogenesis. | 10.1007/s00018-010-0535-z. Molecular Mechanisms of Centrosome and Cytoskeleton Anchorage at the Nuclear Envelope. 10.3390/cells8101238.Lack of a Retinal Phenotype in a Syne-2/Nesprin-2 Knockout Mouse Model. |
| Q66GS9 | Centrosomal protein of 135 kDa OS=Homo sapiens GN=CEP135 PE=1 SV=2 - [CP135_HUMAN] | Involved in early **centriole assembly/duplication/biogenesis/formation/**. Required for centriole elongation. Required for the recruitment of CEP295 to the proximal end of new-born centrioles at the centriolar microtubule wall during early S phase in a PLK4-dependent manner. Centrosomal protein involved in centriole biogenesis. Acts as a scaffolding protein during early centriole biogenesis. Required for the targeting of centriole satellite proteins to centrosomes such as of PCM1, SSX2IP and CEP290 and recruitment of WRAP73 to centrioles. Also required for centriole-centriole cohesion during interphase by acting as a platform protein for CEP250 at the centriole. Required for the recruitment of CEP295 to the proximal end of new-born centrioles at the centriolar microtubule wall during early S phase in a PLK4-dependent manner. Required for the recruitment of CEP295 to the proximal end of new-born centrioles at the centriolar microtubule wall during early S phase in a PLK4-dependent manner | 10.1038/ncb3274. Conserved Molecular Interactions in Centriole-To-Centrosome Conversion. 10.1016/j.cub.2015.08.039. A Short CEP135 Splice Isoform Controls Centriole Duplication. 10.1242/jcs.113506. Drosophila Cep135/Bld10 Maintains Proper Centriole Structure but Is Dispensable for Cartwheel Formation |
| P23258 | Tubulin gamma-1 chain OS=Homo sapiens GN=TUBG1 PE=1 SV=2 - [TBG1_HUMAN] | Tubulin is the major constituent of microtubules. The **gamma chain is found at microtubule organizing centers (MTOC) such as the spindle poles or the centrosome**. Pericentriolar matrix component that regulates alpha/beta chain minus-end nucleation, centrosome duplication and spindle formation. | 10.1083/jcb.141.3.663. The Mammalian Gamma-Tubulin Complex Contains Homologues of the Yeast Spindle Pole Body Components spc97p and spc98p. 10.1038/nature02166. Proteomic Characterization of the Human Centrosome by Protein Correlation Profiling. 10.1083/jcb.141.3.689. Characterization of the Human Homologue of the Yeast spc98p and Its Association With Gamma-Tubulin |
| Q8IYX3 | Coiled-coil domain-containing protein 116 OS=Homo sapiens GN=CCDC116 PE=1 SV=2 - [CC116_HUMAN] | **Identified as a centrosomal protein.** | 10.1242/jcs.157008. Proteomic Analysis of Mammalian Sperm Cells Identifies New Components of the Centrosome. |
| O75116 | Rho-associated protein kinase 2 OS=Homo sapiens GN=ROCK2 PE=1 SV=4 - [ROCK2_HUMAN] | Protein kinase which is a key regulator of actin cytoskeleton and cell polarity. Involved in regulation of smooth muscle contraction, actin cytoskeleton organization, stress fiber and focal adhesion formation, neurite retraction, cell adhesion and motility via phosphorylation of ADD1, BRCA2, CNN1, EZR, DPYSL2, EP300, MSN, MYL9/MLC2, NPM1, RDX, PPP1R12A and VIM. Phosphorylates SORL1 and IRF4. Acts as a negative regulator of VEGF-induced angiogenic endothelial cell activation. Positively regulates the activation of p42/MAPK1-p44/MAPK3 and of p90RSK/RPS6KA1 during myogenic differentiation. **Plays an important role in the timely initiation of centrosome duplication.** Inhibits keratinocyte terminal differentiation. May regulate closure of the eyelids and ventral body wall through organization of actomyosin bundles. Plays a critical role in the regulation of spine and synaptic properties in the hippocampus. Plays an important role in generating the circadian rhythm of the aortic myofilament $Ca^{2+}$ sensitivity and vascular contractility by modulating the myosin light chain phosphorylation. | 10.4161/15384101.2014.989121. Functional Relationship Among PLK2, PLK4 and ROCK2 to Induce Centrosome Amplification. 10.1128/MCB.01383-06. Interaction Between ROCK II and nucleophosmin/B23 in the Regulation of Centrosome Duplication. 10.1158/0008-5472.CAN-10-0030. BRCA2 and Nucleophosmin Coregulate Centrosome Amplification and Form a Complex With the Rho Effector Kinase ROCK2. |
| Q8TC44 | POC1 centriolar protein homolog B OS=Homo sapiens GN=POC1B PE=1 SV=1 - [POC1B_HUMAN] | Plays an important role in **centriole assembly and/or stability and ciliogenesis** (PubMed:20008567). Involved in early steps of centriole duplication, as well as in the later steps of centriole length control (PubMed:19109428). Acts in concert with POC1A to ensure centriole integrity and proper mitotic spindle formation. Required for primary cilia formation, ciliary length and also cell proliferation (PubMed:23015594). Required for retinal integrity (PubMed:25044745). | 10.1083/jcb.200908019. Basal Body Stability and Ciliogenesis Requires the Conserved Component Poc1. 10.1091/mbc.e08-06-0619. Molecular Architecture of the Centriole Proteome: The Conserved WD40 Domain Protein POC1 Is Required for Centriole Duplication and Length Control. 10.1242/jcs.111203.. Poc1A and Poc1B Act Together in Human Cells to Ensure Centriole Integrity. |
| Q99996 | A-kinase anchor protein 9 OS=Homo sapiens GN=AKAP9 PE=1 SV=3 - [AKAP9_HUMAN] | **Scaffolding protein that assembles several protein kinases and phosphatases on the centrosome** and Golgi apparatus. Required to maintain the integrity of the Golgi apparatus (PubMed:10202149, PubMed:15047863). Required for microtubule nucleation at the cis-side of the Golgi apparatus (PubMed:15047863, PubMed:19242490). Required for association of the centrosomes with the poles of the bipolar mitotic spindle during metaphase (PubMed:25657325). In complex with PDE4DIP isoform 13/MMG8/SMYLE, recruits CAMSAP2 to the Golgi apparatus and tethers non-centrosomal minus-end microtubules to the Golgi, an important step for polarized cell movement (PubMed:27666745, PubMed:28814570). In complex with PDE4DIP isoform 13/MMG8/SMYLE, EB1/MAPRE1 and CDK5RAP2, contributes to microtubules nucleation and extension also from the centrosome to the cell periphery (PubMed:29162697). | 10.1091/mbc.E14-09-1366. CENP-32 is required to maintain centrosomal dominance in bipolar spindle assembly. 10.1073/pnas.1705682114. EB1-binding-myomegalin protein complex promotes centrosomal microtubules functions. 10.1093/emboj/18.7.1858. Cloning and characterization of a cDNA encoding an A-kinase anchoring protein located in the centrosome, AKAP450. |
| P42025 | Beta-centractin OS=Homo sapiens GN=ACTR1B PE=1 SV=1 - [ACTY_HUMAN] | Component of a multi-subunit complex involved in microtubule based vesicle motility. It is **associated with the centrosome** | 10.1038/359246a0. Centractin is an actin homologue associated with the centrosome |
| P60510 | Serine/threonine-protein phosphatase 4 catalytic subunit OS=Homo sapiens GN=PPP4C PE=1 SV=1 - [PP4C_HUMAN] | Protein phosphatase that is involved in many processes such as **microtubule organization at centrosomes**, maturation of spliceosomal snRNPs, apoptosis, DNA repair, tumor necrosis factor (TNF)-alpha signaling, activation of c-Jun N-terminal kinase MAPK8, regulation of histone acetylation, DNA damage checkpoint signaling, NF-kappa-B activation and cell migration. The PPP4C-PPP4R1 PP4 complex may play a role in dephosphorylation and regulation of HDAC3. The PPP4C-PPP4R2-PPP4R3A PP4 complex specifically dephosphorylates H2AX phosphorylated on Ser-140 (gamma-H2AX) generated during DNA replication and required for DNA double strand break repair. Dephosphorylates NDEL1 at CDK1 phosphorylation sites and negatively regulates CDK1 activity in interphase (By similarity). In response to DNA damage, catalyzes RPA2 dephosphorylation, an essential step for DNA repair since it allows the efficient RPA2-mediated recruitment of RAD51 to chromatin. | 10.1016/j.biocel.2008.03.021. Depletion of protein phosphatase 4 in human cells reveals essential roles in centrosome maturation, cell migration and the regulation of Rho GTPases. 10.1083/jcb.200705148. Protein phosphatase 4 catalytic subunit regulates Cdk1 activity and microtubule organization via NDEL1 dephosphorylation |
| Q9Y5Y2 | Cytosolic Fe-S cluster assembly factor NUBP2 OS=Homo sapiens GN=NUBP2 PE=1 SV=1 - [NUBP2_HUMAN] | Enriched at the centrosomes during mitosis. **Enriched in centrioles** of microtubule asters during prophase, prometaphase and telophase stages of mitosis (By similarity). Localized at centrioles and in the nucleus at interphase (By similarity). Colocalizes with nubp-1 at prometaphase (By similarity). | Similarity with the mouse protein. 0.1242/jcs.02922. Motor protein KIFC5A interacts with Nubp1 and Nubp2, and is implicated in the regulation of centrosome duplication |
| O15078 | Centrosomal protein of 290 kDa OS=Homo sapiens GN=CEP290 PE=1 SV=2 - [CE290_HUMAN] | Involved in early and late steps in cilia formation. Its association with CCP110 is required for inhibition of primary cilia formation by CCP110 (PubMed:18694559). May play a role in early ciliogenesis in the **disappearance of centriolar satellites** and in the transition of primary ciliar vesicles (PCVs) to capped ciliary vesicles (CCVs). Required for the centrosomal recruitment of RAB8A and for the targeting of centriole satellite proteins to centrosomes such as of PCM1 (PubMed:24421332). Required for the correct localization of ciliary and phototransduction proteins in retinal photoreceptor cells; may play a role in ciliary transport processes (By similarity). Required for efficient recruitment of RAB8A to primary cilium (PubMed:17705300). In the ciliary transition zone is part of the tectonic-like complex which is required for tissue-specific ciliogenesis and may regulate ciliary membrane composition (By similarity). Involved in regulation of the BBSome complex integrity, specifically for presence of BBS2, BBS5 and BBS8/TTC8 in the complex, and in ciliary targeting of selected BBSome cargos. May play a role in controlling entry of the BBSome complex to cilia possibly implicating IQCB1/NPHP5 (PubMed:25552655). Activates ATF4-mediated transcription | 10.1002/humu.20614. Mutations of the CEP290 gene encoding a centrosomal protein cause Meckel-Gruber syndrome. 10.1083/jcb.201304153.The CP110-interacting proteins Talpid3 and Cep290 play overlapping and distinct roles in cilia assembly |
| Q502W7 | Coiled-coil domain-containing protein 38 OS=Homo sapiens GN=CCDC38 PE=2 SV=1 - [CCD38_HUMAN] | **Identified as a centrosomal protein.** | 10.1242/jcs.157008. Proteomic analysis of mammalian sperm cells identifies new components of the centrosome |

| | | | |
|---|---|---|---|
| Q15154 | Pericentriolar material 1 protein OS=Homo sapiens GN=PCM1 PE=1 SV=4 - [PCM1_HUMAN] | **Required for centrosome assembly and function** (PubMed:12403812, PubMed:15659651, PubMed:16943179). Essential for the correct localization of several centrosomal proteins including CEP250, CETN3, PCNT and NEK2 (PubMed:12403812, PubMed:15659651). Required to anchor microtubules to the centrosome (PubMed:12403812, PubMed:15659651). Involved in the biogenesis of cilia (PubMed:20551181, PubMed:24121310). | 10.1083/jcb.200204023. Assembly of centrosomal proteins and microtubule organization depends on PCM-1. 10.1091/mbc.e04-08-0688. Dynamic recruitment of Nek2 kinase to the centrosome involves microtubules, PCM-1, and localized proteasomal degradation. 10.1083/jcb.200606051. Inhibition of centrosome protein assembly leads to p53-dependent exit from the cell cycle. 10.1083/jcb.200204023. Assembly of centrosomal proteins and microtubule organization depends on PCM-1. 10.1091/mbc.e04-08-0688. Dynamic recruitment of Nek2 kinase to the centrosome involves microtubules, PCM-1, and localized proteasomal degradation. 10.1083/jcb.200204023. Assembly of centrosomal proteins and microtubule organization depends on PCM-1. 10.1091/mbc.e04-08-0688. Dynamic recruitment of Nek2 kinase to the centrosome involves microtubules, PCM-1, and localized proteasomal degradation. |
| Q9P2L0 | WD repeat-containing protein 35 OS=Homo sapiens GN=WDR35 PE=1 SV=3 - [WDR35_HUMAN] | Similarity to the mouse protein. **Centrosome localization** during embryo development. | 10.1016/j.ajhg.2011.03.015. Human and mouse mutations in WDR35 cause short-rib polydactyly syndromes due to abnormal ciliogenesis |
| P36404 | ADP-ribosylation factor-like protein 2 OS=Homo sapiens GN=ARL2 PE=1 SV=4 - [ARL2_HUMAN] | Small GTP-binding protein which cycles between an inactive GDP-bound and an active GTP-bound form, and the rate of cycling is regulated by guanine nucleotide exchange factors (GEF) and GTPase-activating proteins (GAP). GTP-binding protein that does not act as an allosteric activator of the cholera toxin catalytic subunit. Regulates formation of new microtubules and **centrosome integrity**. Prevents the TBCD-induced microtubule destruction. Participates in association with TBCD, in the disassembly of the apical junction complexes. Antagonizes the effect of TBCD on epithelial cell detachment and tight and adherens junctions disassembly. Together with ARL2, plays a role in the nuclear translocation, retention and transcriptional activity of STAT3. Component of a regulated secretory pathway involved in $Ca^{2+}$-dependent release of acetylcholine. Required for normal progress through the cell cycle | 10.1091/mbc.e05-10-0929. Arl2 and Arl3 regulate different microtubule-dependent processes. 10.1083/jcb.201503047. Arl2- and Msps-dependent microtubule growth governs asymmetric division |
| Q76KD6 | Speriolin OS=Homo sapiens GN=SPATC1 PE=2 SV=2 - [SPERI_HUMAN] | Similarity to the mouse protein. **Novel sperm centrosomal protein** | 10.1093/humrep/deq138. Speriolin is a novel human and mouse sperm centrosome protein. |
| P53990 | IST1 homolog OS=Homo sapiens GN=IST1 PE=1 SV=1 - [IST1_HUMAN] | **Localizes to centrosomes during mitosis** | 10.1091/mbc.E09-10-0879. SPG20 protein spartin is recruited to midbodies by ESCRT-III protein Ist1 and participates in cytokinesis |
| Q8IZT6 | Abnormal spindle-like microcephaly-associated protein OS=Homo sapiens GN=ASPM PE=1 SV=2 - [ASPM_HUMAN] | Involved in mitotic spindle regulation and coordination of mitotic processes. The function in regulating microtubule dynamics at spindle poles including spindle orientation, astral microtubule density and poleward microtubule flux seems to depend on the association with the katanin complex formed by KATNA1 and KATNB1. Enhances the microtubule lattice severing activity of KATNA1 by recruiting the katanin complex to microtubules. Can block microtubule minus-end growth and reversely this function can be enhanced by the katanin complex (PubMed:28436967). **Aspm localize to the proximal end of the mother centriole** | 10.1016/j.neuron.2016.09.056. Microcephaly Proteins Wdr62 and Aspm Define a Mother Centriole Complex Regulating Centriole Biogenesis, Apical Complex, and Cell Fate. 10.4161/cc.6.13.4356. ASPM and citron kinase co-localize to the midbody ring during cytokinesis. 10.4161/cc.4.9.2029. The abnormal spindle-like, microcephaly-associated (ASPM) gene encodes a centrosomal protein |
| O75665 | Oral-facial-digital syndrome 1 protein OS=Homo sapiens GN=OFD1 PE=1 SV=1 - [OFD1_HUMAN] | **Component of the centrioles controlling mother and daughter centrioles length**. Recruits to the centriole IFT88 and centriole distal appendage-specific proteins including CEP164. Involved in the biogenesis of the cilium, a centriole-associated function. The cilium is a cell surface projection found in many vertebrate cells required to transduce signals important for development and tissue homeostasis. Plays an important role in development by regulating Wnt signaling and the specification of the left-right axis. Only OFD1 localized at the centriolar satellites is removed by autophagy, which is an important step in the ciliogenesis regulation (By similarity). | 10.1097/01.asn.0000054497.48394.d2. OFD1, the gene mutated in oral-facial-digital syndrome type 1, is expressed in the metanephros and in human embryonic renal mesenchymal cells. 10.1038/nature02166. Proteomic characterization of the human centrosome by protein correlation profiling. 10.1016/j.devcel.2009.12.022. Ofd1, a human disease gene, regulates the length and distal structure of centrioles. 10.1093/hmg/ddv488. OFIP/KIAA0753 forms a complex with OFD1 and FOR20 at pericentriolar satellites and centrosomes and is mutated in one individual with oral-facial-digital syndrome. |
| Q8TAP6 | Centrosomal protein of 76 kDa OS=Homo sapiens GN=CEP76 PE=1 SV=1 - [CEP76_HUMAN] | Centrosomal protein involved in **regulation of centriole duplication**. Required to limit centriole duplication to once per cell cycle by preventing centriole reduplication. | 10.1016/j.devcel.2009.03.004. Cep76, a centrosomal protein that specifically restrains centriole reduplication |
| Q96C92 | Serologically defined colon cancer antigen 3 OS=Homo sapiens GN=SDCCAG3 PE=1 SV=3 - [SDCG3_HUMAN] | GIT1 and SDCCAG3 partially colocalize at the centrosomal area. **SDCCAG3 and IFT88 both localize at centrosomes and cilia**. Endosome-associated protein that plays a role in membrane receptor sorting, cytokinesis and ciliogenesis (PubMed:23108400, PubMed:25278552, PubMed:27767179). Involved in the endosome-to-plasma membrane trafficking and recycling of SNX27-retromer-dependent cargo proteins, such as GLUT1 (PubMed:25278552). Involved in the regulation of cytokinesis; the function may involve PTPN13 and GIT1 (PubMed:23108400). Plays a role in the formation of cilia (PubMed:27767179). Involved in cargo protein localization, such as PKD2, at primary cilia (PubMed:27767179). Involved in the presentation of the tumor necrosis factor (TNF) receptor TNFRSF1A on the cell surface, and hence in the modulation of the TNF-induced apoptosis (By similarity). | 10.1038/onc.2012.485. The serologically defined colon cancer antigen-3 interacts with the protein tyrosine phosphatase PTPN13 and is involved in the regulation of cytokinesis. 10.1038/srep35399. The serologically defined colon cancer antigen-3 (SDCCAG3) is involved in the regulation of ciliogenesis. |
| Q9Y6D5 | Brefeldin A-inhibited guanine nucleotide-exchange protein 2 OS=Homo sapiens GN=ARFGEF2 PE=1 SV=3 - [BIG2_HUMAN] | By immunofluorescence microscopy, endogenous BIG2 in HepG2 cells was visualized at Golgi membranes and apparently at the microtubule-organizing center (MTOC).**BIG2 was identified in purified centrosomes. Immunoreactive BIG2 partially or completely overlapped with gamma-tubulin at the MTOC in cells** inspected by confocal microscopy. In HepG2 cells BIG2 interacts in trans-Golgi network and centrosomes, as well as in exocyst structures or complexes that move along microtubules to the plasma membrane, consistent with a functional association in early and late stages of vesicular trafficking. | 10.1073/pnas.0409871102. Interaction of BIG2, a brefeldin A-inhibited guanine nucleotide-exchange protein, with exocyst protein Exo70 |
| Q9UNZ2 | NSFL1 cofactor p47 OS=Homo sapiens GN=NSFL1C PE=1 SV=2 - [NSF1C_HUMAN] | Reduces the ATPase activity of VCP (By similarity). Necessary for the fragmentation of Golgi stacks during mitosis and for VCP-mediated reassembly of Golgi stacks after mitosis (By similarity). May play a role in VCP-mediated formation of transitional endoplasmic reticulum (tER) (By similarity). Inhibits the activity of CTSL (in vitro) (PubMed:15498563). Together with UBXN2B/p37, **regulates the centrosomal levels of kinase AURKA/Aurora A during mitotic progression by promoting AURKA removal from centrosomes in prophase** (PubMed:23649807). Also, regulates spindle orientation during mitosis (PubMed:23649807). | 10.1083/jcb.201209107. The UBXN-2/p37/p47 Adaptors of CDC-48/p97 Regulate Mitosis by Limiting the Centrosomal Recruitment of Aurora A |
| P37198 | Nuclear pore glycoprotein p62 OS=Homo sapiens GN=NUP62 PE=1 SV=3 - [NUP62_HUMAN] | Essential component of the nuclear pore complex (PubMed:1915414). The N-terminal is probably involved in nucleocytoplasmic transport (PubMed:1915414). The C-terminal is involved in protein-protein interaction probably via coiled-coil formation, promotes its association with centrosomes and may function in anchorage of p62 to the pore complex (PubMed:1915414, PubMed:24107630). Plays a role in mitotic cell cycle progression by **regulating centrosome segregation, centriole maturation and spindle orientation** (PubMed:24107630). It might be involved in protein recruitment to the centrosome after nuclear breakdown (PubMed:24107630) | 10.4161/cc.26671. Nucleoporin Nup62 maintains centrosome homeostasis |
| Q02750 | Dual specificity mitogen-activated protein kinase kinase 1 OS=Homo sapiens GN=MAP2K1 PE=1 SV=2 - [MP2K1_HUMAN] | **Localizes at centrosomes** during prometaphase, midzone during anaphase and midbody during telophase/cytokinesis (PubMed:14737111). Membrane localization is probably regulated by its interaction with KSR1 (PubMed:10409742). Dual specificity protein kinase which acts as an essential component of the MAP kinase signal transduction pathway. Binding of extracellular ligands such as growth factors, cytokines and hormones to their cell-surface receptors activates RAS and this initiates RAF1 activation. RAF1 then further activates the dual-specificity protein kinases MAP2K1/MEK1 and MAP2K2/MEK2. Both MAP2K1/MEK1 and MAP2K2/MEK2 function specifically in the MAPK/ERK cascade, and catalyze the concomitant phosphorylation of a threonine and a tyrosine residue in a Thr-Glu-Tyr sequence located in the extracellular signal-regulated kinases MAPK3/ERK1 and MAPK1/ERK2, leading to their activation and further transduction of the signal within the MAPK/ERK cascade. Activates BRAF in a KSR1 or KSR2-dependent manner; by binding to KSR1 or KSR2 releases the inhibitory intramolecular interaction between KSR1 or KSR2 protein kinase and N-terminal domains which promotes KSR1 or KSR2-BRAF dimerization and BRAF activation (PubMed:29433126). Depending on the cellular context, this pathway mediates diverse biological functions such as cell growth, adhesion, survival and differentiation, predominantly through the regulation of transcription, metabolism and cytoskeletal rearrangements. One target of the MAPK/ERK cascade is peroxisome proliferator-activated receptor gamma (PPARG), a nuclear receptor that promotes differentiation and apoptosis. MAP2K1/MEK1 has been shown to export PPARG from the nucleus. The MAPK/ERK cascade is also involved in the regulation of endosomal dynamics, including lysosome processing and endosome cycling through the perinuclear recycling compartment (PNRC), as well as in the fragmentation of the Golgi apparatus during mitosis. | 10.1038/sj.onc.1207188. The MAP kinase pathway is required for entry into mitosis and cell survival. |
| P60953 | Cell division control protein 42 homolog OS=Homo sapiens GN=CDC42 PE=1 SV=2 - [CDC42_HUMAN] | **Colocalizes with NEK6 in the centrosome.** Plasma membrane-associated small GTPase which cycles between an active GTP-bound and an inactive GDP-bound state. In active state binds to a variety of effector proteins to regulate cellular responses. Involved in epithelial cell polarization processes. Regulates the bipolar attachment of spindle microtubules to kinetochores before chromosome congression in metaphase (PubMed:15642749). Regulates cell migration (PubMed:17038317). In neurons, plays a role in the extension and maintenance of the formation of filopodia, thin and actin-rich surface projections (PubMed:14978216). Required for DOCK10-mediated spine formation in Purkinje cells and hippocampal neurons. Facilitates filopodia formation upon DOCK11-activation (By similarity). Upon activation by CaMKII, modulates dendritic spine structural plasticity by relaying CaMKII transient activation to synapse-specific, long-term signaling (By similarity). Also plays a role in phagocytosis through organization of the F-actin cytoskeleton associated with forming phagocytic cups (PubMed:26465210). | 10.1021/pr100562w. Characterization of hNek6 interactome reveals an important role for its short N-terminal domain and colocalization with proteins at the centrosome |
| O75935 | Dynactin subunit 3 OS=Homo sapiens GN=DCTN3 PE=1 SV=1 - [DCTN3_HUMAN] | Together with dynein may be involved in spindle assembly and cytokinesis. **Localizes to the centrosome.** | 10.1083/jcb.142.4.1023. Characterization of the p22 subunit of dynactin reveals the localization of cytoplasmic dynein and dynactin to the midbody of dividing cells |
| Q8N865 | Uncharacterized protein C7orf31 OS=Homo sapiens GN=C7orf31 PE=2 SV=3 - [CG031_HUMAN] | **Localizes to the sperm centrosome** | 10.1242/jcs.157008. Proteomic analysis of mammalian sperm cells identifies new components of the centrosome. |
| Q13561 | Dynactin subunit 2 OS=Homo sapiens GN=DCTN2 PE=1 SV=4 - [DCTN2_HUMAN] | Modulates cytoplasmic dynein binding to an organelle, and plays a role in prometaphase chromosome alignment and spindle organization during mitosis. **Involved in anchoring microtubules to centrosomes**. May play a role in synapse formation during brain development. | 10.1083/jcb.132.4.617. Molecular characterization of the 50-kD subunit of dynactin reveals function for the complex in chromosome alignment and spindle organization during mitosis. |
| P63167 | Dynein light chain 1, cytoplasmic OS=Homo sapiens GN=DYNLL1 PE=1 SV=1 - [DYL1_HUMAN] | **Centrosome localization.** Acts as one of several non-catalytic accessory components of the cytoplasmic dynein 1 complex that are thought to be involved in linking dynein to cargos and to adapter proteins that regulate dynein function. Cytoplasmic dynein 1 acts as a motor for the intracellular retrograde motility of vesicles and organelles along microtubules. May play a role in changing or maintaining the spatial distribution of cytoskeletal structures.<br>Binds and inhibits the catalytic activity of neuronal nitric oxide synthase.<br>Promotes transactivation functions of ESR1 and plays a role in the nuclear localization of ESR1.<br>Regulates apoptotic activities of BCL2L11 by sequestering it to microtubules. Upon apoptotic stimuli the BCL2L11-DYNLL1 complex dissociates from cytoplasmic dynein and translocates to mitochondria and sequesters BCL2 thus neutralizing its antiapoptotic activity. | 10.1038/nature02166. Proteomic characterization of the human centrosome by protein correlation profiling |
| Q9UJT2 | Testis-specific serine kinase substrate OS=Homo sapiens GN=TSKS PE=1 SV=3 - [TSKS_HUMAN] | Localization in the **spermatid centrioles**. May play a role in testicular physiology, most probably in the process of spermatogenesis or spermatid development. | 10.1016/j.ydbio.2008.03.043. TSKS concentrates in spermatid centrioles during flagellogenesis |
| P36405 | ADP-ribosylation factor-like protein 3 OS=Homo sapiens GN=ARL3 PE=1 SV=2 - [ARL3_HUMAN] | **Centrosome localization**. Small GTP-binding protein which cycles between an inactive GDP-bound and an active GTP-bound form, and the rate of cycling is regulated by guanine nucleotide exchange factors (GEF) and GTPase-activating proteins (GAP) (PubMed:16525022, PubMed:18588884). Required for normal cytokinesis and cilia signaling (PubMed:22085962). Requires assistance from GTPase-activating proteins (GAPs) like RP2 and PDE6D, in order to cycle between inactive GDP-bound and active GTP-bound forms. Required for targeting proteins to the cilium, including myristoylated NPHP3 and prenylated INPP5E (PubMed:30269812). Targets NPHP3 to the ciliary membrane by releasing myristoylated NPHP3 from UNC119B cargo adapter into the cilium (PubMed:22085962). Required for PKD1:PKD2 complex targeting from the trans-Golgi network to the cilium (By similarity). | 10.1091/mbc.e05-10-0929. Arl2 and Arl3 regulate different microtubule-dependent processes |
| Q8IYX7 | Protein FAM154A OS=Homo sapiens GN=FAM154A PE=1 SV=2 - [F154A_HUMAN] | In multi-ciliated cells, **localizes to the basal bodies and in non-ciliated cells, to the centrosome.** In spermatozoa, colocalizes with microtubules along the length of the axoneme from its proximal end to its distal tip and with tubulin at the distal end of the flagellum and at **the proximal centriole**. May play a role in the regulation of cilium length. Stabilizes microtubules at low temperature. | 10.1242/jcs.155143. Human FAM154A (SAXO1) is a microtubule-stabilizing protein specific to cilia and related structures. |
| A5D8V7 | Coiled-coil domain-containing protein 151 OS=Homo sapiens GN=CCDC151 PE=1 SV=1 - [CC151_HUMAN] | In dividing IMCD3 cells, CCDC151 is enriched at the **centrioles**. In ciliated cells, CCDC151 localizes to the primary cilia and basal bodies. | 10.1093/hmg/ddt445. The coiled-coil domain containing protein CCDC151 is required for the function of IFT-dependent motile cilia in animals |
| O43379 | WD repeat-containing protein 62 OS=Homo sapiens GN=WDR62 PE=1 SV=4 - [WDR62_HUMAN] | Required for cerebral cortical development. Plays a role in neuronal proliferation and migration (PubMed:20890278, PubMed:20729831). Plays a role in **mother-centriole-dependent centriole duplication**; the function seems also to involve CEP152, CDK5RAP2 and CEP63 through a stepwise assembled complex at the centrosome that recruits CDK2 required for centriole duplication (PubMed:26297806). | 10.1038/ng.683. Mutations in WDR62, encoding a centrosome-associated protein, cause microcephaly with simplified gyri and abnormal cortical architecture. 10.7554/eLife.07519. Centriolar satellites assemble centrosomal microcephaly proteins to recruit CDK2 and promote centriole duplication. 10.1111/j.1399-0004.2011.01686.x. Mutations in WDR62, encoding a centrosomal and nuclear protein, in Indian primary microcephaly families with cortical malformations. |
| O43805 | Sjoegren syndrome nuclear autoantigen 1 OS=Homo sapiens GN=SSNA1 PE=1 SV=2 - [SSNA1_HUMAN] | **Novel centrosomal protein.** | 10.1038/nature02166. Proteomic characterization of the human centrosome by protein correlation profiling |
| P35222 | Catenin beta-1 OS=Homo sapiens GN=CTNNB1 PE=1 SV=1 - [CTNB1_HUMAN] | Key downstream component of the canonical Wnt signaling pathway (PubMed:17524503, PubMed:18077326, PubMed:18086858, PubMed:18957423, PubMed:21262353, PubMed:22155184, PubMed:22647378, PubMed:22699938). In the absence of Wnt, forms a complex with AXIN1, AXIN2, APC, CSNK1A1 and GSK3B that promotes phosphorylation on N-terminal Ser and Thr residues and ubiquitination of CTNNB1 via BTRC and its subsequent degradation by the proteasome (PubMed:17524503, PubMed:18077326, PubMed:18086858, PubMed:18957423, PubMed:21262353, PubMed:22155184, PubMed:22647378, PubMed:22699938). In the presence of Wnt ligand, CTNNB1 is not ubiquitinated and accumulates in the nucleus, where it acts as a coactivator for transcription factors of the TCF/LEF family, leading to activate Wnt responsive genes (PubMed:17524503, PubMed:18077326, PubMed:18086858, PubMed:18957423, PubMed:21262353, PubMed:22155184, PubMed:22647378, PubMed:22699938). Involved in the regulation of cell adhesion, as component of an E-cadherin:catenin adhesion complex (By similarity). **Acts as a negative regulator of centrosome cohesion** (PubMed:18086858). Involved in the CDK2/PTPN6/CTNNB1/CEACAM1 pathway of insulin internalization (PubMed:21262353). Blocks anoikis of malignant kidney and intestinal epithelial cells and promotes their anchorage-independent growth by down-regulating DAPK2 (PubMed:18957423). Disrupts PML function and PML-NB formation by inhibiting RANBP2-mediated sumoylation of PML (PubMed:22155184). Promotes neurogenesis by maintaining sympathetic neuroblasts within the cell cycle (By similarity). Involved in chondrocyte differentiation via interaction with SOX9: SOX9-binding competes with the binding sites of TCF/LEF within CTNNB1, thereby inhibiting the Wnt signaling (By similarity). | 10.1101/gad.1596308. beta-Catenin is a Nek2 substrate involved in centrosome separation. |
| P48729 | Casein kinase I isoform alpha OS=Homo sapiens GN=CSNK1A1 PE=1 SV=2 - [KC1A_HUMAN] | **Localizes to the centrosome in interphase cells**, and to kinetochore fibers during mitosis. Also recruited to the keratin cytoskeleton. Casein kinases are operationally defined by their preferential utilization of acidic proteins such as caseins as substrates. It can phosphorylate a large number of proteins. Participates in Wnt signaling. Phosphorylates CTNNB1 at 'Ser-45'. May phosphorylate PER1 and PER2. May play a role in segregating chromosomes during mitosis (PubMed:11955436, PubMed:1409656, PubMed:18305108). May play a role in keratin cytoskeleton disassembly and thereby, it may regulate epithelial cell migration (PubMed:23902688) | 10.1073/pnas.89.20.9454. Cell cycle-dependent localization of casein kinase I to mitotic spindles |
| Q14203 | Dynactin subunit 1 OS=Homo sapiens GN=DCTN1 PE=1 SV=3 - [DCTN1_HUMAN] | Plays a key role in dynein-mediated retrograde transport of vesicles and organelles along microtubules by recruiting and tethering dynein to microtubules. Binds to both dynein and microtubules providing a link between specific cargos, microtubules and dynein. Essential for targeting dynein to microtubule plus ends, recruiting dynein to membranous cargos and enhancing dynein processivity (the ability to move along a microtubule for a long distance without falling off the track). Can also act as a brake to slow the dynein motor during motility along the microtubule (PubMed:25185702). Can regulate microtubule stability by promoting microtubule formation, nucleation and polymerization and by inhibiting microtubule catastrophe in neurons. Inhibits microtubule catastrophe by binding both to microtubules and to tubulin, leading to enhanced microtubule stability along the axon (PubMed:23874158). Plays a role in metaphase spindle orientation (PubMed:22327364). **Plays a role in centriole cohesion and subdistal appendage organization and function.** Its recruitment to the centriole in a KIF3A-dependent manner is essential for the maintenance of centriole cohesion and the formation of subdistal appendage. Also required for microtubule anchoring at the mother centriole (PubMed:23386061). Plays a role in primary cilia formation (PubMed:25774020). | 10.1038/emboj.2013.3. Kif3a interacts with Dynactin subunit p150 Glued to organize centriole subdistal appendages. 10.1091/mbc.E10-05-0430. Par6 alpha interacts with the dynactin subunit p150 Glued and is a critical regulator of centrosomal protein recruitment. 10.1091/mbc.E13-03-0137. Ste20-like protein kinase SLK (LOSK) regulates microtubule organization by targeting dynactin to the centrosome. 10.1002/jcb.25160. Cell Cycle-Dependent Localization of Dynactin Subunit p150 glued at Centrosome. |
| Q9BZV1 | UBX domain-containing protein 6 OS=Homo sapiens GN=UBXN6 PE=1 SV=1 - [UBXN6_HUMAN] | Ubxd1 is a stable protein that localizes to the cytoplasm and nucleus and is **highly enriched in centrosomes**. May negatively regulate the ATPase activity of VCP, an ATP-driven segregase that associates with different cofactors to control a wide variety of cellular processes (PubMed:26475856). As a cofactor of VCP, it may play a role in the transport of CAV1 to lysosomes for degradation (PubMed:21822278, PubMed:23335559). It may also play a role in endoplasmic reticulum-associated degradation (ERAD) of misfolded proteins (PubMed:19275885). Together with VCP and other cofactors, it may play a role in macroautophagy, regulating for instance the clearance of damaged lysosomes (PubMed:27753622). | 10.1016/j.biocel.2008.06.008. Ubxd1 is a novel co-factor of the human p97 ATPase. |

| | | | |
|---|---|---|---|
| Q13464 | Rho-associated protein kinase 1 OS=Homo sapiens GN=ROCK1 PE=1 SV=1 - [ROCK1_HUMAN] | Protein kinase which is a key regulator of actin cytoskeleton and cell polarity (PubMed:10436159, PubMed:10652353, PubMed:11018042, PubMed:11283607, PubMed:17158456, PubMed:18573880, PubMed:19131646, PubMed:8617235, PubMed:9722579). Involved in regulation of smooth muscle contraction, actin cytoskeleton organization, stress fiber and focal adhesion formation, neurite retraction, cell adhesion and motility via phosphorylation of DAPK3, GFAP, LIMK1, LIMK2, MYL9/MLC2, TPPP, PFN1 and PPP1R12A (PubMed:10436159, PubMed:10652353, PubMed:11018042, PubMed:11283607, PubMed:17158456, PubMed:18573880, PubMed:19131646, PubMed:8617235, PubMed:9722579, PubMed:23093407, PubMed:23355470). Phosphorylates FHOD1 and acts synergistically with it to promote SRC-dependent non-apoptotic plasma membrane blebbing (PubMed:18694941). Phosphorylates JIP3 and regulates the recruitment of JNK to JIP3 upon UVB-induced stress (PubMed:19036714). Acts as a suppressor of inflammatory cell migration by regulating PTEN phosphorylation and stability (By similarity). Acts as a negative regulator of VEGF-induced angiogenic endothelial cell activation (PubMed:19181962). **Required for centrosome positioning and centrosome-dependent exit from mitosis** (By similarity). Plays a role in terminal erythroid differentiation (PubMed:21072057). May regulate closure of the eyelids and ventral body wall by inducing the assembly of actomyosin bundles (By similarity). Promotes keratinocyte terminal differentiation (PubMed:19997641). Involved in osteoblast compaction through the fibronectin fibrillogenesis cell-mediated matrix assembly process, essential for osteoblast mineralization (By similarity). | 10.1083/jcb.200203034. The Rho-associated protein kinase p160ROCK is required for centrosome positioning |
| Q12798 | Centrin-1 OS=Homo sapiens GN=CETN1 PE=1 SV=1 - [CETN1_HUMAN] | Plays a fundamental **role in microtubule-organizing center structure and function** (PubMed:8175926). Plays a role in sperm cilia formation (By similarity). | PMID: 8175926. Cloning of a cDNA encoding human centrin, an EF-hand protein of centrosomes and mitotic spindle poles |
| Q9Y265 | RuvB-like 1 OS=Homo sapiens GN=RUVBL1 PE=1 SV=1 - [RUVB1_HUMAN] | Mainly localized in the nucleus, associated with nuclear matrix or in the nuclear cytosol, although it is also present in the cytoplasm and associated with the cell membranes. In prophase and prometaphase it is **located at the centrosome** and the branching microtubule spindles. After mitotic nuclear membrane disintigration it accumulates at the centrosome and sites of tubulin polymerization. As cells pass through metaphase and into telophase it is located close to the centrosome at the early phase of tubulin polymerization. In anaphase it accumulates at the zone of tubule interdigitation. In telophase it is found at polar tubule overlap, and it reappears at the site of chromosomal decondensation in the daughter cells. Possesses single-stranded DNA-stimulated ATPase and ATP-dependent DNA helicase (3' to 5') activity; hexamerization is thought to be critical for ATP hydrolysis and adjacent subunits in the ring-like structure contribute to the ATPase activity (PubMed:17157868). Component of the NuA4 histone acetyltransferase complex which is involved in transcriptional activation of select genes principally by acetylation of nucleosomal histones H4 and H2A (PubMed:14966270). This modification may both alter nucleosome-DNA interactions and promote interaction of the modified histones with other proteins which positively regulate transcription (PubMed:14966270). This complex may be required for the activation of transcriptional programs associated with oncogene and proto-oncogene mediated growth induction, tumor suppressor mediated growth arrest and replicative senescence, apoptosis, and DNA repair (PubMed:14966270). The NuA4 complex ATPase and helicase activities seem to be, at least in part, contributed by the association of RUVBL1 and RUVBL2 with EP400. NuA4 may also play a direct role in DNA repair when recruited to sites of DNA damage (PubMed:14966270). Component of a SWR1-like complex that specifically mediates the removal of histone H2A.Z/H2AZ1 from the nucleosome (PubMed:24463511). Proposed core component of the chromatin remodeling INO80 complex which exhibits DNA- and nucleosome-activated ATPase activity and catalyzes ATP-dependent nucleosome sliding (PubMed:16230350, PubMed:21303910). Plays an essential role in oncogenic transformation by MYC and also modulates transcriptional activation by the LEF1/TCF1-CTNNB1 complex (PubMed:10882073, PubMed:16014379). Essential for cell proliferation (PubMed:14506706). May be able to bind plasminogen at cell surface and enhance plasminogen activation (PubMed:11027681). | 10.1002/cm.10136. The ATP-dependent helicase RUVBL1/TIP49a associates with tubulin during mitosis |
| Q9UJW0 | Dynactin subunit 4 OS=Homo sapiens GN=DCTN4 PE=1 SV=1 - [DCTN4_HUMAN] | Could have a dual role in dynein targeting and in ACTR1A/Arp1 subunit of dynactin pointed-end capping. Could be involved in ACTR1A pointed-end binding and in additional roles in linking dynein and dynactin to the cortical cytoskeleton. **Centrosome localization during the cell cycle** | 10.1074/jbc.275.7.4834. A Dynactin Subunit with a Highly Conserved Cysteine-rich Motif Interacts Directly with Arp1 |
| Q9NRG9 | Aladin OS=Homo sapiens GN=AAAS PE=1 SV=1 - [AAAS_HUMAN] | Plays a role in the normal development of the peripheral and central nervous system (PubMed:11062474, PubMed:11159947, PubMed:16022285). Required for the correct localization of aurora kinase AURKA and the microtubule minus end-binding protein NUMA1 as well as a subset of AURKA targets which ensures proper spindle formation and timely chromosome alignment (PubMed:26246606). **Without ALADIN, Aurora A spreads from centrosomes** onto spindle microtubules, which affects the distribution of a subset of microtubule regulators and slows spindle assembly and chromosome alignment. | 10.1091/mbc.E15-02-0113. The nucleoporin ALADIN regulates Aurora A localization to ensure robust mitotic spindle formation |
| P43034 | Platelet-activating factor acetylhydrolase IB subunit alpha OS=Homo sapiens GN=PAFAH1B1 PE=1 SV=2 - [LIS1_HUMAN] | Required for proper activation of Rho GTPases and actin polymerization at the leading edge of locomoting cerebellar neurons and postmigratory hippocampal neurons in response to calcium influx triggered via NMDA receptors. Non-catalytic subunit of an acetylhydrolase complex which inactivates platelet-activating factor (PAF) by removing the acetyl group at the SN-2 position (By similarity). Positively regulates the activity of the minus-end directed microtubule motor protein dynein. May enhance dynein-mediated microtubule sliding by targeting dynein to the microtubule plus end. Required for several dynein- and microtubule-dependent processes such as the maintenance of Golgi integrity, the peripheral transport of microtubule fragments and the **coupling of the nucleus and centrosome**. Required during brain development for the proliferation of neuronal precursors and the migration of newly formed neurons from the ventricular/subventricular zone toward the cortical plate. Neuronal migration involves a process called nucleokinesis, whereby migrating cells extend an anterior process into which the nucleus subsequently translocates. During nucleokinesis dynein at the nuclear surface may translocate the nucleus towards the centrosome by exerting force on centrosomal microtubules. May also play a role in other forms of cell locomotion including the migration of fibroblasts during wound healing. Required for dynein recruitment to microtubule plus ends and BICD2-bound cargos (PubMed:22956769). | 10.1038/nature02166. Proteomic characterization of the human centrosome by protein correlation profiling. 10.1083/jcb.200309025. Lis1 and doublecortin function with dynein to mediate coupling of the nucleus to the centrosome in neuronal migration. |
| Q9BTW9 | Tubulin-specific chaperone D OS=Homo sapiens GN=TBCD PE=1 SV=2 - [TBCD_HUMAN] | **TBCD localizes at the centrosome**, the major microtubule-organizing center. Tubulin-folding protein implicated in the first step of the tubulin folding pathway and required for tubulin complex assembly. Involved in the regulation of microtubule polymerization or depolymerization, it modulates microtubule dynamics by capturing GTP-bound beta-tubulin (TUBB). Its ability to interact with beta tubulin is regulated via its interaction with ARL2. Acts as a GTPase-activating protein (GAP) for ARL2. Induces microtubule disruption in absence of ARL2. Increases degradation of beta tubulin, when overexpressed in polarized cells. Promotes epithelial cell detachment, a process antagonized by ARL2. Induces tight adherens and tight junctions disassembly at the lateral cell membrane (PubMed:10722852, PubMed:10831612, PubMed:11847227, PubMed:20740604, PubMed:27666370, PubMed:28158450). Required for correct assembly and maintenance of the mitotic spindle, and proper progression of mitosis (PubMed:27666370). Involved in neuron morphogenesis (PubMed:27666374). | 10.1016/j.ajhg.2016.08.003. Biallelic Mutations in TBCD, Encoding the Tubulin Folding Cofactor D, Perturb Microtubule Dynamics and Cause Early-Onset Encephalopathy. |
| P04899 | Guanine nucleotide-binding protein G(i) subunit alpha-2 OS=Homo sapiens GN=GNAI2 PE=1 SV=3 - [GNAI2_HUMAN] | **Localization of G(ialpha1), G(ialpha2), and G(ialpha3) in the centrosomes** and at the midbody. Guanine nucleotide-binding proteins (G proteins) are involved as modulators or transducers in various transmembrane signaling systems. The G(i) proteins are involved in hormonal regulation of adenylate cyclase: they inhibit the cyclase in response to beta-adrenergic stimuli. May play a role in cell division. | 10.1083/jcb.200604114. Localization of Gi alpha proteins in the centrosomes and at the midbody: implication for their role in cell division. |
| Q9Y224 | UPF0568 protein C14orf166 OS=Homo sapiens GN=C14orf166 PE=1 SV=1 - [CN166_HUMAN] | **Interacts with ninein affecting its centrosomal localization and ninein function.** RNA-binding protein involved in modulation of mRNA transcription by Polymerase II (PubMed:16950395). Component of the tRNA-splicing ligase complex and is required for tRNA ligation (PubMed:24870230). May be required for RNA transport (PubMed:24608264). | 10.1016/j.febslet.2004.04.024. A novel ninein-interaction protein, CGI-99, blocks ninein phosphorylation by GSK3beta and is highly expressed in brain tumors |
| P62081 | 40S ribosomal protein S7 OS=Homo sapiens GN=RPS7 PE=1 SV=1 - [RS7_HUMAN] | **Colocalizes with NEK6 in the centrosome**. Required for rRNA maturation. | 10.1021/pr100562w. Characterization of hNek6 interactome reveals an important role for its short N-terminal domain and colocalization with proteins at the centrosome. |
| P06748 | Nucleophosmin OS=Homo sapiens GN=NPM1 PE=1 SV=2 - [NPM_HUMAN] | Involved in diverse cellular processes such as ribosome biogenesis, **centrosome** duplication, protein chaperoning, histone assembly, cell proliferation, and regulation of tumor suppressors p53/TP53 and ARF. Binds ribosome presumably to drive ribosome nuclear export. Associated with nucleolar ribonucleoprotein structures and bind single-stranded nucleic acids. Acts as a chaperonin for the core histones H3, H2B and H4. Stimulates APEX1 endonuclease activity on apurinic/apyrimidinic (AP) double-stranded DNA but inhibits APEX1 endonuclease activity on AP single-stranded RNA. May exert a control of APEX1 endonuclease activity within nucleoli devoted to repair AP on rDNA and the removal of oxidized rRNA molecules. In concert with BRCA2, **regulates centrosome duplication. Regulates centriole duplication**: phosphorylation by PLK2 is able to trigger centriole replication. Negatively regulates the activation of EIF2AK2/PKR and suppresses apoptosis through inhibition of EIF2AK2/PKR autophosphorylation. Antagonizes the inhibitory effect of ATF5 on cell proliferation and relieves ATF5-induced G2/M blockade (PubMed:22528486). In complex with MYC enhances the transcription of MYC target genes (PubMed:25956029). | 10.1128/MCB.01383-06. Interaction between ROCK II and nucleophosmin/B23 in the regulation of centrosome duplication. 10.1371/journal.pone.0009849. Polo-like kinase 2-dependent phosphorylation of NPM/B23 on serine 4 triggers centriole duplication. 10.1158/0008-5472.CAN-10-0030. BRCA2 and nucleophosmin coregulate centrosome amplification and form a complex with the Rho effector kinase ROCK2. |
| P53384 | Cytosolic Fe-S cluster assembly factor NUBP1 OS=Homo sapiens GN=NUBP1 PE=1 SV=2 - [NUBP1_HUMAN] | Component of the cytosolic iron-sulfur (Fe/S) protein assembly (CIA) machinery (PubMed:18573874). Required for maturation of extramitochondrial Fe-S proteins (PubMed:18573874). The NUBP1-NUBP2 heterotetramer forms a Fe-S scaffold complex, mediating the de novo assembly of an Fe-S cluster and its transfer to target apoproteins (PubMed:18573874). Implicated in the **regulation of centrosome duplication** (By similarity). Negatively regulates cilium formation and structure (By similarity) | 10.1242/jcs.02922. Motor protein KIFC5A interacts with Nubp1 and Nubp2, and is implicated in the regulation of centrosome duplication. 10.1007/s00018-013-1401-6. The nucleotide-binding proteins Nubp1 and Nubp2 are negative regulators of ciliogenesis. |
| P33176 | Kinesin-1 heavy chain OS=Homo sapiens GN=KIF5B PE=1 SV=1 - [KINH_HUMAN] | Microtubule-dependent motor required for normal distribution of mitochondria and lysosomes. Can induce formation of neurite-like membrane protrusions in non-neuronal cells in a ZFYVE27-dependent manner (By similarity). **Regulates centrosome and nuclear positioning** during mitotic entry. During the G2 phase of the cell cycle in a BICD2-dependent manner, antagonizes dynein function and drives the separation of nuclei and centrosomes (PubMed:20386726). Required for anterograde axonal transportation of MAPK8IP3/JIP3 which is essential for MAPK8IP3/JIP3 function in axon elongation (By similarity). | 10.1371/journal.pbio.1000350. Bicaudal D2, dynein, and kinesin-1 associate with nuclear pore complexes and regulate centrosome and nuclear positioning during mitotic entry. |
| Q96PF2 | Testis-specific serine/threonine-protein kinase 2 OS=Homo sapiens GN=TSSK2 PE=1 SV=2 - [TSSK2_HUMAN] | TSSK2/TSKS is the **first kinase/substrate pair localized to the centrioles of spermatids and spermatozoa.** Testis-specific serine/threonine-protein kinase required during spermatid development. Phosphorylates TSKS at 'Ser-288' and SPAG16. Involved in the late stages of spermatogenesis, during the reconstruction of the cytoplasm. During spermatogenesis, required for the transformation of a ring-shaped structure around the base of the flagellum originating from the chromatoid body. | 10.1016/j.ydbio.2008.03.043. TSKS concentrates in spermatid centrioles during flagellogenesis |
| Q9H892 | Tetratricopeptide repeat protein 12 OS=Homo sapiens GN=TTC12 PE=1 SV=2 - [TTC12_HUMAN] | **TTC12 localizes to the centrosome** as defined by gamma-tubulin labeling. | 10.1371/journal.pone.0052166. Transcriptional program of ciliated epithelial cells reveals new cilium and centrosome components and links to human disease. |
| Q9H0I3 | Coiled-coil domain-containing protein 113 OS=Homo sapiens GN=CCDC113 PE=1 SV=1 - [CC113_HUMAN] | **Component of centriolar satellites** contributing to primary cilium formation. | 10.1242/jcs.157008. Proteomic analysis of mammalian sperm cells identifies new components of the centrosome |
| Q8IYE0 | Coiled-coil domain-containing protein 146 OS=Homo sapiens GN=CCDC146 PE=2 SV=2 - [CC146_HUMAN] | Localized to only one centriole identified as the **mother centriole** by co-staining with CEP164 | 10.1242/jcs.157008. Proteomic analysis of mammalian sperm cells identifies new components of the centrosome |
| Q9C099 | Leucine-rich repeat and coiled-coil domain-containing protein 1 OS=Homo sapiens GN=LRRCC1 PE=1 SV=2 - [LRCC1_HUMAN] | Required for the organization of the mitotic spindle. **Maintains the structural integrity of centrosomes** during mitosis. | 10.4161/cc.7.17.6591. An evolutionarily conserved leucine-rich repeat protein CLERC is a centrosomal protein required for spindle pole integrity |
| Q5BJF6 | Outer dense fiber protein 2 OS=Homo sapiens GN=ODF2 PE=1 SV=1 - [ODFP2_HUMAN] | Seems to be a major component of sperm tail outer dense fibers (ODF). ODFs are filamentous structures located on the outside of the axoneme in the midpiece and principal piece of the mammalian sperm tail and may help to maintain the passive elastic structures and elastic recoil of the sperm tail. May have a modulating influence on sperm motility. Functions as a general scaffold protein that is **specifically localized at the distal/subdistal appendages of mother centrioles**. Component of the centrosome matrix required for the localization of PLK1 and NIN to the centrosomes. Required for the formation and/or maintenance of normal CETN1 assembly. | 10.1128/MCB.00671-06. Requirement of hCenexin for proper mitotic functions of polo-like kinase 1 at the centrosomes. |
| P61163 | Alpha-centractin OS=Homo sapiens GN=ACTR1A PE=1 SV=1 - [ACTZ_HUMAN] | Component of a multi-subunit complex involved in microtubule based vesicle motility. It is **associated with the centrosome**. | 10.1038/nature02166. Proteomic characterization of the human centrosome by protein correlation profiling |
| Q5JVL4 | EF-hand domain-containing protein 1 OS=Homo sapiens GN=EFHC1 PE=1 SV=1 - [EFHC1_HUMAN] | EFHC1 is a microtubule-associated protein (MAP) involved in cell division and radial migration during cerebral corticogenesis. **EFHC1 colocalizes with the centrosome** and the mitotic spindle. | 10.1093/hmg/dds356. Mutations of EFHC1, linked to juvenile myoclonic epilepsy, disrupt radial and tangential migrations during brain development |
| Q92597 | Protein NDRG1 OS=Homo sapiens GN=NDRG1 PE=1 SV=1 - [NDRG1_HUMAN] | **Centrosome localization**. NDRG1 mediates centrosome homeostasis, maintaining genome stability. | 10.1038/s41598-020-68629-4. Analysis of the "centrosome-ome" identifies MCPH1 deletion as a cause of centrosome amplification in human cancer. 10.1073/pnas.1503683112. NDRG1 links p53 with proliferation-mediated centrosome homeostasis and genome stability. 10.1186/s12917-019-1872-2. Cell and context-dependent sorting of neuropathy-associated protein NDRG1 - insights from canine tissues and primary Schwann cell cultures |
| Q2M329 | Coiled-coil domain-containing protein 96 OS=Homo sapiens GN=CCDC96 PE=2 SV=2 - [CCD96_HUMAN] | **Identified as a centrosomal protein.** | 10.1242/jcs.157008. Proteomic analysis of mammalian sperm cells identifies new components of the centrosome |
| Q00839 | Heterogeneous nuclear ribonucleoprotein U OS=Homo sapiens GN=HNRNPU PE=1 SV=6 - [HNRPU_HUMAN] | The Ser-59 phosphorylated form **localizes to centrosomes** during prophase and metaphase, to mitotic spindles in anaphase and to the midbody during cytokinesis. | 10.1038/nature02166. Proteomic characterization of the human centrosome by protein correlation profiling. 10.1083/jcb.200309025. Lis1 and doublecortin function with dynein to mediate coupling of the nucleus to the centrosome in neuronal migration. 10.1128/MCB.01312-14. Phosphorylation of SAF-A/hnRNP-U Serine 59 by Polo-Like Kinase 1 Is Required for Mitosis |
| Q9Y696 | Chloride intracellular channel protein 4 OS=Homo sapiens GN=CLIC4 PE=1 SV=4 - [CLIC4_HUMAN] | It **localizes at the centrosomes** with AKAP350 in cultured mammalian cells. | 10.1002/cm.10141. CLIC4 is enriched at cell-cell junctions and colocalizes with AKAP350 at the centrosome and midbody of cultured mammalian cells |
| P08754 | Guanine nucleotide-binding protein G(k) subunit alpha OS=Homo sapiens GN=GNAI3 PE=1 SV=3 - [GNAI3_HUMAN] | **Centrosome localization.** | 10.1083/jcb.200604114. Localization of Gi alpha proteins in the centrosomes and at the midbody: implication for their role in cell division |
| A7E2U8 | UPF0602 protein C4orf47 OS=Homo sapiens GN=C4orf47 PE=2 SV=1 - [CD047_HUMAN] | **Identified as a centrosomal protein.** | 10.1242/jcs.157008. Proteomic analysis of mammalian sperm cells identifies new components of the centrosome |
| Q8WVJ2 | NudC domain-containing protein 2 OS=Homo sapiens GN=NUDCD2 PE=1 SV=1 - [NUDC2_HUMAN] | It **associates with centrosomes** in interphase and to spindle poles and kinetochores during mitosis. | 10.1073/pnas.0914307107. NudC-like protein 2 regulates the LIS1/dynein pathway by stabilizing LIS1 with Hsp90. |
| Q6ZN84 | Coiled-coil domain-containing protein 81 OS=Homo sapiens GN=CCDC81 PE=2 SV=2 - [CCD81_HUMAN] | Identified as a **centrosomal protein.** | 10.1242/jcs.157008. Proteomic analysis of mammalian sperm cells identifies new components of the centrosome |
| Q8WUM4 | Programmed cell death 6-interacting protein OS=Homo sapiens GN=PDCD6IP PE=1 SV=1 - [PDC6I_HUMAN] | **Colocalizes with CEP55 at centrosomes** of non-dividing cells. Multifunctional protein involved in endocytosis, multivesicular body biogenesis, membrane repair, cytokinesis, apoptosis and maintenance of tight junction integrity. Class E VPS protein involved in concentration and sorting of cargo proteins of the multivesicular body (MVB) for incorporation into intralumenal vesicles (ILVs) that are generated by invagination and scission from the limiting membrane of the endosome. Binds to the phospholipid lysobisphosphatidic acid (LBPA) which is abundant in MVBs internal membranes. The MVB pathway requires the sequential function of ESCRT-O, -I,-II and -III complexes (PubMed:14739459). The ESCRT machinery also functions in topologically equivalent membrane fission events, such as the terminal stages of cytokinesis (PubMed:17853893, PubMed:17556548). Adapter for a subset of ESCRT-III proteins, such as CHMP4, to function at distinct membranes. Required for completion of cytokinesis (PubMed:17853893, PubMed:17556548, PubMed:18641129). May play a role in the regulation of both apoptosis and cell proliferation. Regulates exosome biogenesis in concert with SDC1/4 and SDCBP (PubMed:22660413). By interacting with F-actin, PARD3 and TJP1 secures the proper assembly and positioning of actomyosin-tight junction complex at the apical sides of adjacent epithelial cells that defines a spatial membrane domain essential for the maintenance of epithelial cell polarity and barrier (By similarity). | 10.1038/sj.emboj.7601850. Human ESCRT and ALIX proteins interact with proteins of the midbody and function in cytokinesis. |

| | | | |
|---|---|---|---|
| P08107;P0DMV8 | Heat shock 70 kDa protein 1A/1B OS=Homo sapiens GN=HSPA1A PE=1 SV=5 - [HSP71_HUMAN] | Molecular chaperone implicated in a wide variety of cellular processes, including protection of the proteome from stress, folding and transport of newly synthesized polypeptides, activation of proteolysis of misfolded proteins and the formation and dissociation of protein complexes. Plays a pivotal role in the protein quality control system, ensuring the correct folding of proteins, the re-folding of misfolded proteins and controlling the targeting of proteins for subsequent degradation. This is achieved through cycles of ATP binding, ATP hydrolysis and ADP release, mediated by co-chaperones. The co-chaperones have been shown to not only regulate different steps of the ATPase cycle, but they also have an individual specificity such that one co-chaperone may promote folding of a substrate while another may promote degradation. The affinity for polypeptides is regulated by its nucleotide bound state. In the ATP-bound form, it has a low affinity for substrate proteins. However, upon hydrolysis of the ATP to ADP, it undergoes a conformational change that increases its affinity for substrate proteins. It goes through repeated cycles of ATP hydrolysis and nucleotide exchange, which permits cycles of substrate binding and release. The co-chaperones are of three types: J-domain co-chaperones such as HSP40s (stimulate ATPase hydrolysis by HSP70), the nucleotide exchange factors (NEF) such as BAG1/2/3 (facilitate conversion of HSP70 from the ADP-bound to the ATP-bound state thereby promoting substrate release), and the TPR domain chaperones such as HOPX and STUB1 (PubMed:24012426, PubMed:26865365, PubMed:24318877). Maintains protein homeostasis during cellular stress through two opposing mechanisms: protein refolding and degradation. Its acetylation/deacetylation state determines whether it functions in protein refolding or protein degradation by controlling the competitive binding of co-chaperones HOPX and STUB1. During the early stress response, the acetylated form binds to HOPX which assists in chaperone-mediated protein refolding, thereafter, it is deacetylated and binds to ubiquitin ligase STUB1 that promotes ubiquitin-mediated protein degradation (PubMed:27708256). **Regulates centrosome integrity** during mitosis, and is required for the **maintenance of a functional mitotic centrosome** that supports the assembly of a bipolar mitotic spindle (PubMed:27137183). Enhances STUB1-mediated SMAD3 ubiquitination and degradation and facilitates STUB1-mediated inhibition of TGF-beta signaling (PubMed:24613385). Essential for STUB1-mediated ubiquitination and degradation of FOXP3 in regulatory T-cells (Treg) during inflammation (PubMed:23973223). Negatively regulates heat shock-induced HSF1 transcriptional activity during the attenuation and recovery phase period of the heat shock response | 10.1007/s00018-016-2236-8. HSP70 regulates the function of mitotic centrosomes |
| Q9BZW7 | Testis-specific gene 10 protein OS=Homo sapiens GN=TSGA10 PE=1 SV=1 - [TSG10_HUMAN] | In mature spermatozoa, **localizes to the centriole and midpiece** (PubMed:28905369). The 27-kDa peptide associates with the fibrous sheath in mature spermatozoa and localizes to the principal piece of sperm tail, while the 55-kDa peptide localizes to the midpiece (By similarity). Detected in the cytoplasm of almost all spermatogonial cells within the seminiferous tubules (By similarity). Plays a role in spermatogenesis (PubMed:28905369). When overexpressed, prevents nuclear localization of HIF1A (By similarity). | 10.1111/cge.13140. TSGA10 is a novel candidate gene associated with acephalic spermatozoa |
| Q9H5N1 | Rab GTPase-binding effector protein 2 OS=Homo sapiens GN=RABEP2 PE=1 SV=2 - [RABE2_HUMAN] | Plays a role in membrane trafficking and in homotypic early endosome fusion (PubMed:9524116). Participates in arteriogenesis by regulating vascular endothelial growth factor receptor 2/VEGFR2 cell surface expression and endosomal trafficking (PubMed:29425100). By interacting with SDCCAG8, **localizes to centrosomes** and plays a critical role in ciliogenesis | 10.1371/journal.pone.0156081. SDCCAG8 Interacts with RAB Effector Proteins RABEP2 and ERC1 and Is Required for Hedgehog Signaling |
| Q5VYK3 | Proteasome-associated protein ECM29 homolog OS=Homo sapiens GN=ECM29 PE=1 SV=2 - [ECM29_HUMAN] | **Localizes to centrosomes.** Proteasome-associated protein ECM29 homolog; Adapter/scaffolding protein that binds to the 26S proteasome, motor proteins and other compartment specific proteins. May couple the proteasome to different compartments including endosome, endoplasmic reticulum and **centrosome**. May play a role in ERAD and other enhanced proteolyis *[a.k.a. ECM29, ECM29_HUMAN, R4GMY1, Q5VYK3]* | 10.1074/jbc.M410444200. Characterization of mammalian Ecm29, a 26 S proteasome-associated protein that localizes to the nucleus and membrane vesicles |
| Q8WW35 | Tctex1 domain-containing protein 2 OS=Homo sapiens GN=TCTEX1D2 PE=1 SV=2 - [TC1D2_HUMAN] | Tctex1 domain-containing protein 2; Required for proper retrograde ciliary transport; Belongs to the dynein light chain Tctex-type family. **Localizes at the centrosome.** *[a.k.a. Tctex1 domain-containing protein 2, ENST00000545438, NP_689986, Q8WW35]* | 10.4161/15384101.2014.985066. Tctex1d2 associates with short-rib polydactyly syndrome proteins and is required for ciliogenesis |
| O43237 | Cytoplasmic dynein 1 light intermediate chain 2 OS=Homo sapiens GN=DYNC1LI2 PE=1 SV=1 - [DC1L2_HUMAN] | Cytoplasmic dynein 1 light intermediate chain 2; Acts as one of several non-catalytic accessory components of the cytoplasmic dynein 1 complex that are thought to be involved in linking dynein to cargos and to adapter proteins that regulate dynein function. Cytoplasmic dynein 1 acts as a motor for the intracellular retrograde motility of vesicles and organelles along microtubules. May play a role in binding dynein to membranous organelles or chromosomes; Belongs to the dynein light intermediate chain family *[a.k.a. LIC2, DNCLI2, NM_006141, O43237]*. **Maintains centriole cohesion during mitosis.** | 10.1083/jcb.201408025. Dynein light intermediate chains maintain spindle bipolarity by functioning in centriole cohesion |
| Q8TCI5 | Protein pitchfork OS=Homo sapiens GN=PIFO PE=1 SV=2 - [PIFO_HUMAN] | Protein pitchfork; During primary cilia disassembly, involved in cilia disassembly. Required specifically to control cilia retraction as well as the liberation and duplication of the **basal body/centrosome.** May act by stimulating AURKA activity at the basal body in a cell cycle-dependent manner *[a.k.a. C1orf88, FLJ23853, ENST00000369738, Q8TCI5]* | 10.1016/j.devcel.2010.06.005. Pitchfork regulates primary cilia disassembly and left-right asymmetry |
| Q9Y6G9 | Cytoplasmic dynein 1 light intermediate chain 1 OS=Homo sapiens GN=DYNC1LI1 PE=1 SV=3 - [DC1L1_HUMAN] | Cytoplasmic dynein 1 light intermediate chain 1; Acts as one of several non-catalytic accessory components of the cytoplasmic dynein 1 complex that are thought to be involved in linking dynein to cargos and to adapter proteins that regulate dynein function. Cytoplasmic dynein 1 acts as a motor for the intracellular retrograde motility of vesicles and organelles along microtubules. May play a role in binding dynein to membranous organelles or chromosomes. Probably involved in the microtubule-dependent transport of pericentrin. Is required for progress through the spindle assembly checkp [...]. **Maintains centriole cohesion during mitosis. Interacts with pericentrin**. *[a.k.a. DNCLI1, AC104306, OTTHUMT00000342171, Q9Y6G9]* | 10.1083/jcb.201408025. Dynein light intermediate chains maintain spindle bipolarity by functioning in centriole cohesion. 10.1074/jbc.M001536200. Light intermediate chain 1 defines a functional subfraction of cytoplasmic dynein which binds to pericentrin. |
| Q9Y230 | RuvB-like 2 OS=Homo sapiens GN=RUVBL2 PE=1 SV=3 - [RUVB2_HUMAN] | RuvB-like 2; Involved in the endoplasmic reticulum (ER)-associated degradation (ERAD) pathway where it negatively regulates expression of ER stress response genes; Belongs to the RuvB family. **During anaphase, some TIP48 was detected in the centrosome colocalizing with tubulin** *[a.k.a. CGI-46, INO80J, TIP49B, Q9Y230]* | 10.1016/j.yexcr.2005.07.030. Relocalization of human chromatin remodeling cofactor TIP48 in mitosis |
| Q14204 | Cytoplasmic dynein 1 heavy chain 1 OS=Homo sapiens GN=DYNC1H1 PE=1 SV=5 - [DYHC1_HUMAN] | **Localizes at the centrosome.** Cytoplasmic dynein 1 acts as a motor for the intracellular retrograde motility of vesicles and organelles along microtubules. Dynein has ATPase activity; the force-producing power stroke is thought to occur on release of ADP. Plays a role in mitotic spindle assembly and metaphase plate congression; Belongs to the dynein heavy chain family *[a.k.a. DYHC, KIAA0325, DNCH1, Q14204]* | 10.1016/j.neuron.2020.01.030. Posterior Neocortex-Specific Regulation of Neuronal Migration by CEP85L Identifies Maternal Centriole-Dependent Activation of CDK5 |
| Q5VTH2 | UPF0740 protein C1orf192 OS=Homo sapiens GN=C1orf192 PE=2 SV=1 - [CA192_HUMAN] | Protein Flattop; Acts as a regulator of cilium basal body docking and positioning in mono- and multiciliated cells. Regulates basal body docking and cilia formation in multiciliated lung cells. Regulates kinocilium positioning and stereocilia bundle morphogenesis in the inner ear. **Flattop functions to position basal bodies** [a.k.a. FLTP, CFAP126, Chromosome 1 open reading frame 192, Q5VTH2] | 10.7554/eLife.03842. Flattop regulates basal body docking and positioning in mono- and multiciliated cell |
| Q8ND07 | Coiled-coil domain-containing protein 176 OS=Homo sapiens GN=CCDC176 PE=2 SV=3 - [CC176_HUMAN] | Basal body-orientation factor 1; Basal body protein required in multiciliate cells to align and maintain cilia orientation in response to flow. May act by mediating a maturation step that stabilizes and aligns cilia orientation. Not required to respond to planar cell polarity (PCP) or flow-based orientation cues (By similarity). **Transiently associated with basal bodies** *[a.k.a. BBOF1, C14orf45, NP_079333.2, Q8ND07]* | 10.1242/dev.096727. Bbof1 is required to maintain cilia orientation |

1 unique peptide:

| Accession | Description | Uniprot description | References |
|---|---|---|---|
| Q5T890 | DNA excision repair protein ERCC-6-like 2 OS=Homo sapiens GN=ERCC6L2 PE=1 SV=2 - [ER6L2_HUMAN] | **Colocalizes with NEK6 in the centrosome**. May be involved in early DNA damage response | 10.1021/pr100562w. Characterization of hNek6 interactome reveals an important role for its short N-terminal domain and colocalization with proteins at the centrosome. |
| Q8N4N8 | Kinesin-like protein KIF2B OS=Homo sapiens GN=KIF2B PE=1 SV=3 - [KIF2B_HUMAN] | Kif2b is expressed at very low levels in cultured cells and that **GFP-Kif2b localizes predominately to centrosomes** and midbodies, but also to spindle microtubules and transiently to kinetochores. Kif2b-deficient. cells assemble monopolar or disorganized spindles. Plus end-directed microtubule-dependent motor required for spindle assembly and chromosome movement. Has microtubule depolymerization activity (PubMed:17538014). Plays a role in chromosome congression (PubMed:23891108). | 10.1091/mbc.e07-02-0110. The kinesin-13 proteins Kif2a, Kif2b, and Kif2c/MCAK have distinct roles during mitosis in human cells |
| Q86YT6 | E3 ubiquitin-protein ligase MIB1 OS=Homo sapiens GN=MIB1 PE=1 SV=1 - [MIB1_HUMAN] | **Identified as a centrosomal protein**. E3 ubiquitin-protein ligase that mediates ubiquitination of Delta receptors, which act as ligands of Notch proteins. Positively regulates the Delta-mediated Notch signaling by ubiquitinating the intracellular domain of Delta, leading to endocytosis of Delta receptors. Probably mediates ubiquitination and subsequent proteasomal degradation of DAPK1, thereby antagonizing anti-apoptotic effects of DAPK1 to promote TNF-induced apoptosis (By similarity). Involved in ubiquitination of centriolar satellite CEP131, CEP290 and PCM1 proteins and hence inhibits primary cilium formation in proliferating cells. Mediates 'Lys-63'-linked polyubiquitination of TBK1, which probably participates in kinase activation. | 10.1038/emboj.2011.63. Novel asymmetrically localizing components of human centrosomes identified by complementary proteomics methods. |
| O95613 | Pericentrin OS=Homo sapiens GN=PCNT PE=1 SV=4 - [PCNT_HUMAN] | **Integral component of the filamentous matrix of the centrosome** involved in the initial establishment of organized microtubule arrays in both mitosis and meiosis. Plays a role, together with DISC1, in the microtubule network formation. Is an integral component of the pericentriolar material (PCM). May play an important role in preventing premature centrosome splitting during interphase by inhibiting NEK2 kinase activity at the centrosome. | PMID: 11171385. Kendrin/pericentrin-B, a centrosome protein with homology to pericentrin that complexes with PCM-1. 10.1073/pnas.97.11.5919. Identification of a human centrosomal calmodulin-binding protein that shares homology with pericentrin. 10.1016/j.bbrc.2008.10.100. DISC1-kendrin interaction is involved in centrosomal microtubule network formation. 10.1016/j.bbrc.2010.06.063. Involvement of a centrosomal protein kendrin in the maintenance of centrosome cohesion by modulating Nek2A kinase activity |
| Q15051 | IQ calmodulin-binding motif-containing protein 1 OS=Homo sapiens GN=IQCB1 PE=1 SV=1 - [IQCB1_HUMAN] | **Localization to the centrosome** depends on the interaction with CEP290/NPHP6. Involved in ciliogenesis. The function in an early step in cilia formation depends on its association with CEP290/NPHP6 (PubMed:21565611, PubMed:23446637). Involved in regulation of the BBSome complex integrity, specifically for presence of BBS2 and BBS5 in the complex, and in ciliary targeting of selected BBSome cargos. May play a role in controlling entry of the BBSome complex to cilia possibly implicating CEP290/NPHP6 (PubMed:25552655). | 10.1016/j.cell.2011.04.019. Mapping the NPHP-JBTS-MKS protein network reveals ciliopathy disease genes and pathways. |
| Q92830 | Histone acetyltransferase KAT2A OS=Homo sapiens GN=KAT2A PE=1 SV=3 - [KAT2A_HUMAN] | Acts as a negative regulator of centrosome amplification by mediating acetylation of PLK4. **Localizes to centrosomes in late G1 and around the G1/S transition, coinciding with the onset of centriole formation** (PubMed:27796307). Protein lysine acyltransferase that can act as a acetyltransferase, glutaryltransferase or succinyltransferase, depending on the context (PubMed:29211711). Acts as a histone lysine succinyltransferase: catalyzes succinylation of histone H3 on 'Lys-79' (H3K79succ), with a maximum frequency around the transcription start sites of genes (PubMed:29211711). Succinylation of histones gives a specific tag for epigenetic transcription activation (PubMed:29211711). Association with the 2-oxoglutarate dehydrogenase complex, which provides succinyl-CoA, is required for histone succinylation (PubMed:29211711). In different complexes, functions either as an acetyltransferase (HAT) or as a succinyltransferase: in the SAGA and ATAC complexes, acts as a histone acetyltransferase (PubMed:17301242, PubMed:19103755, PubMed:29211711). Has significant histone acetyltransferase activity with core histones, but not with nucleosome core particles (PubMed:17301242, PubMed:19103755). Acetylation of histones gives a specific tag for epigenetic transcription activation (PubMed:17301242, PubMed:19103755, PubMed:29211711). Recruited by the XPC complex at promoters, where it specifically mediates acetylation of histone variant H2A.Z.1/H2A.Z, thereby promoting expression of target genes (PubMed:29973595, PubMed:31527837). Involved in long-term memory consolidation and synaptic plasticity: acts by promoting expression of a hippocampal gene expression network linked to neuroactive receptor signaling (By similarity). Acts as a positive regulator of T-cell activation: upon TCR stimulation, recruited to the IL2 promoter following interaction with NFATC2 and catalyzes acetylation of histone H3 at 'Lys-9' (H3K9ac), leading to promote IL2 expression (By similarity). Required for growth and differentiation of craniofacial cartilage and bone by regulating acetylation of histone H3 at 'Lys-9' (H3K9ac) (By similarity). Regulates embryonic stem cell (ESC) pluripotency and differentiation (By similarity). Also acetylates non-histone proteins, such as CEBPB, PLK4 and TBX5 (PubMed:17301242, PubMed:27796307, PubMed:29174768). Involved in heart and limb development by mediating acetylation of TBX5, acetylation regulating nucleocytoplasmic shuttling of TBX5 (PubMed:29174768). | 10.1038/ncomms13227. KAT2A/KAT2B-targeted acetylome reveals a role for PLK4 acetylation in preventing centrosome amplification |
| Q6DKK2 | Tetratricopeptide repeat protein 19, mitochondrial OS=Homo sapiens GN=TTC19 PE=1 SV=4 - [TTC19_HUMAN] | **Localizes to the centrosome** during all stages of the cell cycle. Required for the preservation of the structural and functional integrity of mitochondrial respiratory complex III by allowing the physiological turnover of the Rieske protein UQCRFS1 (PubMed:21278747, PubMed:28673544). Involved in the clearance of UQCRFS1 N-terminal fragments, which are produced upon incorporation of UQCRFS1 into the complex III and whose presence is detrimental for its catalytic activity (PubMed:28673544). | 10.1038/ncb2036. PtdIns(3)P controls cytokinesis through KIF13A-mediated recruitment of FYVE-CENT to the midbody. |
| P63096 | Guanine nucleotide-binding protein G(i) subunit alpha-1 OS=Homo sapiens GN=GNAI1 PE=1 SV=2 - [GNAI1_HUMAN] | Guanine nucleotide-binding proteins (G proteins) function as transducers downstream of G protein-coupled receptors (GPCRs) in numerous signaling cascades. The alpha chain contains the guanine nucleotide binding site and alternates between an active, GTP-bound state and an inactive, GDP-bound state. Signaling by an activated GPCR promotes GDP release and GTP binding. The alpha subunit has a low GTPase activity that converts bound GTP to GDP, thereby terminating the signal. Both GDP release and GTP hydrolysis are modulated by numerous regulatory proteins (PubMed:8774883, PubMed:18434541). Signaling is mediated via effector proteins, such as adenylate cyclase. Inhibits adenylate cyclase activity, leading to decreased intracellular cAMP levels (By similarity). The inactive GDP-bound form prevents the association of RGS14 with centrosomes and is required for the translocation of RGS14 from the cytoplasm to the plasma membrane. Required for normal cytokinesis during mitosis (PubMed:17635935). Required for cortical dynein-dynactin complex recruitment during metaphase (PubMed:22327364). | 10.1083/jcb.200604114. **Localization of Gi alpha proteins in the centrosomes** and at the midbody: implication for their role in cell division. |
| Q13432 | Protein unc-119 homolog A OS=Homo sapiens GN=UNC119 PE=1 SV=1 - [U119A_HUMAN] | **Localizes to the centrosome in interphase** cells and begins to translocate from the spindle pole to the spindle midzone after the onset of mitosis; it then localizes to the intercellular bridge in telophase cells and to the midbody in cytokinetic cells. | 10.4161/cc.24404. UNC119a bridges the transmission of Fyn signals to Rab11, leading to the completion of cytokinesis |
| Q8IYA6 | Cytoskeleton-associated protein 2-like OS=Homo sapiens GN=CKAP2L PE=1 SV=4 - [CKP2L_HUMAN] | Uniformly distributed along each microtubule bundle of spindles in addition to **centrioles** during mitosis, expression promptly diminishes at interphase. Microtubule-associated protein required for mitotic spindle formation and cell-cycle progression in neural progenitor cells. | 10.1016/j.ajhg.2014.10.008. Mutations in CKAP2L, the human homolog of the mouse Radmis gene, cause Filippi syndrome |
| Q9P287 | BRCA2 and CDKN1A-interacting protein OS=Homo sapiens GN=BCCIP PE=1 SV=1 - [BCCIP_HUMAN] | In interphase, preferential localizes to the **mother centriole**. Recruited to the spindle pole matrix and centrosome by microtubules and dynein/dynactin activity. During interphase, required for microtubule organizing and anchoring activities. During mitosis, required for the organization and stabilization of the spindle pole (PubMed:28394342). Isoform 2/alpha is particularly important for the regulation of microtubule anchoring, microtubule stability, spindle architecture and spindle orientation, compared to isoform 1/beta (PubMed:28394342). May promote cell cycle arrest by enhancing the inhibition of CDK2 activity by CDKN1A. May be required for repair of DNA damage by homologous recombination in conjunction with BRCA2. May not be involved in non-homologous end joining (NHEJ). | 10.1038/onc.2017.92. Regulation of spindle integrity and mitotic fidelity by BCCIP. |
| Q7Z460 | CLIP-associating protein 1 OS=Homo sapiens GN=CLASP1 PE=1 SV=1 - [CLAP1_HUMAN] | Localizes to microtubule plus ends. **Localizes to centrosomes**, kinetochores and the mitotic spindle from prometaphase. Subsequently localizes to the spindle midzone from anaphase and to the midbody from telophase. In migrating cells localizes to the plus ends of microtubules within the cell body and to the entire microtubule lattice within the lamella. Localizes to the cell cortex and this requires ERC1 and PHLDB2. Microtubule plus-end tracking protein that promotes the stabilization of dynamic microtubules. Involved in the nucleation of noncentrosomal microtubules originating from the trans-Golgi network (TGN). Required for the polarization of the cytoplasmic microtubule arrays in migrating cells towards the leading edge of the cell. May act at the cell cortex to enhance the frequency of rescue of depolymerizing microtubules by attaching their plus-ends to cortical platforms composed of ERC1 and PHLDB2. This cortical microtubule stabilizing activity is regulated at least in part by phosphatidylinositol 3-kinase signaling. Also performs a similar stabilizing function at the kinetochore which is essential for the bipolar alignment of chromosomes on the mitotic spindle. | 10.1038/nature02166. Proteomic characterization of the human centrosome by protein correlation profiling |
| Q3B820 | Protein FAM161A OS=Homo sapiens GN=FAM161A PE=1 SV=2 - [F161A_HUMAN] | Localize to the scaffold structure along the inner wall of the **centriole microtubules**. | 10.1007/978-1-4614-3209-8_24. FAM161A, a novel centrosomal-ciliary protein implicated in autosomal recessive retinitis pigmentosa. 10.1093/hmg/dds268. The retinitis pigmentosa 28 protein FAM161A is a novel ciliary protein involved in intermolecular protein interaction and microtubule association. 10.1126/sciadv.aaz4137. A helical inner scaffold provides a structural basis for centriole cohesion |
| O76095 | Protein JTB OS=Homo sapiens GN=JTB PE=1 SV=1 - [JTB_HUMAN] | **Detected at the centrosome** and along spindle fibers during prophase and metaphase. Detected at the midbody during telophase. Required for normal cytokinesis during mitosis. Plays a role in the regulation of cell proliferation. May be a component of the chromosomal passenger complex (CPC), a complex that acts as a key regulator of mitosis. The CPC complex has essential functions at the centromere in ensuring correct chromosome alignment and segregation and is required for chromatin-induced microtubule stabilization and spindle assembly. Increases AURKB activity. Inhibits apoptosis induced by TGFB1 (By similarity). Overexpression induces swelling of mitochondria and reduces mitochondrial membrane potential (By similarity). | 10.3892/ijo.2011.900. PAR, a protein involved in the cell cycle, is functionally related to chromosomal passenger proteins. |
| Q53EZ4 | Centrosomal protein of 55 kDa OS=Homo sapiens GN=CEP55 PE=1 SV=3 - [CEP55_HUMAN] | Present at the centrosomes at interphase. A small portion is associated preferentially with the **mother centriole**, whereas the majority localizes to the pericentriolar material. During mitosis, loses affinity for the centrosome at the onset of prophase and diffuses throughout the cell. This dissociation from the centrosome is phosphorylation-dependent. May remain localized at the centrosome during mitosis in certain cell types. Appears at the cleavage furrow in late anaphase and in the midbody in cytokinesis. Plays a role in mitotic exit and cytokinesis (PubMed:16198290, PubMed:17853893). Recruits PDCD6IP and TSG101 to midbody during cytokinesis. Required for successful completion of cytokinesis (PubMed:17853893). Not required for microtubule nucleation (PubMed:16198290). Plays a role in the development of the brain and kidney (PubMed:28264986). | 10.1016/j.devcel.2005.09.003. Cdk1/Erk2- and Plk1-dependent phosphorylation of a centrosome protein, Cep55, is required for its recruitment to midbody and cytokinesis |
| Q9NY27 | Serine/threonine-protein phosphatase 4 regulatory subunit 2 OS=Homo sapiens GN=PPP4R2 PE=1 SV=3 - [PP4R2_HUMAN] | Regulatory subunit of serine/threonine-protein phosphatase 4 (PP4). May regulate the activity of PPP4C at centrosomal microtubule organizing centers. Its interaction with the SMN complex leads to enhance the temporal localization of snRNPs, suggesting a role of PPP4C in maturation of spliceosomal snRNPs. The PPP4C-PPP4R2-PPP4R3A PP4 complex specifically dephosphorylates H2AX phosphorylated on 'Ser-140' (gamma-H2AX) generated during DNA replication and required for DNA double strand break repair. Mediates RPA2 dephosphorylation by recruiting PPP4C to RPA2 in a DNA damage-dependent manner. RPA2 dephosphorylation is required for the efficient RPA2-mediated recruitment of RAD51 to chromatin following double strand breaks, an essential step for DNA repair. | PMID: 10769191. **A novel 50 kDa protein forms complexes with protein phosphatase 4 and is located at centrosomal microtubule organizing centres** |
| Q15650 | Activating signal cointegrator 1 OS=Homo sapiens GN=TRIP4 PE=1 SV=4 - [TRIP4_HUMAN] | **Colocalizes with NEK6 in the centrosome.** May be involved in early DNA damage response. Transcription coactivator which associates with nuclear receptors, transcriptional coactivators including EP300, CREBBP and NCOA1, and basal transcription factors like TBP and TFIIA to facilitate nuclear receptors-mediated transcription. May thereby play an important role in establishing distinct coactivator complexes under different cellular conditions. Plays a role in thyroid hormone receptor and estrogen receptor transactivation (PubMed:10454579, PubMed:25219498). Also involved in androgen receptor transactivation (By similarity). Plays a pivotal role in the transactivation of NF-kappa-B, SRF and AP1. Acts as a mediator of transrepression between nuclear receptor and either AP1 or NF-kappa-B (PubMed:12077347). May play a role in the development of neuromuscular junction (PubMed:26924529). May play a role in late myogenic differentiation (By similarity). | 10.1021/pr100562w. Characterization of hNek6 interactome reveals an important role for its short N-terminal domain and colocalization with proteins at the centrosome. |

| | | | |
|---|---|---|---|
| Q13042 | Cell division cycle protein 16 homolog OS=Homo sapiens GN=CDC16 PE=1 SV=2 - [CDC16_HUMAN] | **Colocalizes with CDC27 to the centrosome** at all stages of the cell cycle and to the mitotic spindle. Component of the anaphase promoting complex/cyclosome (APC/C), a cell cycle-regulated E3 ubiquitin ligase that controls progression through mitosis and the G1 phase of the cell cycle. The APC/C complex acts by mediating ubiquitination and subsequent degradation of target proteins: it mainly mediates the formation of 'Lys-11'-linked polyubiquitin chains and, to a lower extent, the formation of 'Lys-48'- and 'Lys-63'-linked polyubiquitin chains. | 10.1016/0092-8674(95)90336-4. CDC27Hs colocalizes with CDC16Hs to the centrosome and mitotic spindle and is essential for the metaphase to anaphase transition. |
| O95271 | Tankyrase-1 OS=Homo sapiens GN=TNKS PE=1 SV=2 - [TNKS1_HUMAN] | Poly-ADP-ribosyltransferase involved in various processes such as Wnt signaling pathway, telomere length and vesicle trafficking (PubMed:10988299, PubMed:11739745, PubMed:16076287, PubMed:19759537, PubMed:21478859, PubMed:22864114, PubMed:23622245, PubMed:25043379). Acts as an activator of the Wnt signaling pathway by mediating poly-ADP-ribosylation (PARsylation) of AXIN1 and AXIN2, 2 key components of the beta-catenin destruction complex: poly-ADP-ribosylated target proteins are recognized by RNF146, which mediates their ubiquitination and subsequent degradation (PubMed:19759537, PubMed:21478859). Also mediates PARsylation of BLZF1 and CASC3, followed by recruitment of RNF146 and subsequent ubiquitination (PubMed:21478859). Mediates PARsylation of TERF1, thereby contributing to the regulation of telomere length (PubMed:11739745). Involved in centrosome maturation during prometaphase by mediating PARsylation of HEPACAM2/MIKI (PubMed:22864114). May also regulate vesicle trafficking and modulate the subcellular distribution of SLC2A4/GLUT4-vesicles (PubMed:10988299). May be involved in spindle pole assembly through PARsylation of NUMA1 (PubMed:16076287). Stimulates 26S proteasome activity (PubMed:23622245). | 10.1016/j.molcel.2012.06.033. **Poly-ADP ribosylation of Miki by tankyrase-1 promotes centrosome maturation.** PMID: 10523501. Cell cycle dependent localization of the telomeric PARP, tankyrase, to nuclear pore complexes and centrosomes. 10.1371/journal.pone.0022595. Ubiquitin ligase RNF146 regulates tankyrase and Axin to promote Wnt signaling. |
| Q96CW5 | Gamma-tubulin complex component 3 OS=Homo sapiens GN=TUBGCP3 PE=1 SV=2 - [GCP3_HUMAN] | Gamma-tubulin complex is necessary for **microtubule nucleation at the centrosome.** | 10.1038/nature02166. Proteomic characterization of the human centrosome by protein correlation profiling. |
| Q9UBK9 | Protein UXT OS=Homo sapiens GN=UXT PE=1 SV=1 - [UXT_HUMAN] | UXT is overexpressed in a number of human tumor tissues but not in the matching normal tissues. **UXT is located in human centrosomes** and is associated with gamma-tubulin. Overexpression of UXT disrupts centrosome structure. | 10.1091/mbc.e05-08-0705. UXT is a novel centrosomal protein essential for cell viability |
| Q9NQC7 | Ubiquitin carboxyl-terminal hydrolase CYLD OS=Homo sapiens GN=CYLD PE=1 SV=1 - [CYLD_HUMAN] | **CYLD localizes at centrosomes and basal bodies** via interaction with the centrosomal protein CAP350. CYLD must be both at the centrosome and catalytically active to promote ciliogenesis independently of NF-κB. Detected at the microtubule cytoskeleton during interphase. Detected at the midbody during telophase. During metaphase, it remains localized to the centrosome but is also present along the spindle. Deubiquitinase that specifically cleaves 'Lys-63'- and linear 'Met-1'-linked polyubiquitin chains and is involved in NF-kappa-B activation and TNF-alpha-induced necroptosis (PubMed:18636086, PubMed:26670046, PubMed:27458237, PubMed:26997266, PubMed:27591049, PubMed:29291351, PubMed:18313383). Plays an important role in the regulation of pathways leading to NF-kappa-B activation (PubMed:12917689, PubMed:12917691). Contributes to the regulation of cell survival, proliferation and differentiation via its effects on NF-kappa-B activation (PubMed:12917690). Negative regulator of Wnt signaling (PubMed:20227366). Inhibits HDAC6 and thereby promotes acetylation of alpha-tubulin and stabilization of microtubules (PubMed:19893491). Plays a role in the regulation of microtubule dynamics, and thereby contributes to the regulation of cell proliferation, cell polarization, cell migration, and angiogenesis (PubMed:18222923, PubMed:20194890). Required for normal cell cycle progress and normal cytokinesis (PubMed:17495026, PubMed:19893491). Inhibits nuclear translocation of NF-kappa-B (PubMed:18636086). Plays a role in the regulation of inflammation and the innate immune response, via its effects on NF-kappa-B activation (PubMed:18636086). Dispensable for the maturation of intrathymic natural killer cells, but required for the continued survival of immature natural killer cells (By similarity). Negatively regulates TNFRSF11A signaling and osteoclastogenesis (By similarity). Involved in the regulation of ciliogenesis, allowing ciliary basal bodies to migrate and dock to the plasma membrane; this process does not depend on NF-kappa-B activation (By similarity). Ability to remove linear ('Met-1'-linked) polyubiquitin chains regulates innate immunity and TNF-alpha-induced necroptosis: recruited to the LUBAC complex via interaction with SPATA2 and restricts linear polyubiquitin formation on target proteins (PubMed:26997266, PubMed:26670046, PubMed:27458237, PubMed:27591049). Regulates innate immunity by restricting linear polyubiquitin formation on RIPK2 in response to NOD2 stimulation (PubMed:26997266). Involved in TNF-alpha-induced necroptosis by removing linear ('Met-1'-linked) polyubiquitin chains from RIPK1, thereby regulating the kinase activity of RIPK1 (By similarity). Removes 'Lys-63' linked polyubiquitin chain of MAP3K7, which inhibits phosphorylation | 10.1038/ncomms5585. The deubiquitinating enzyme CYLD controls apical docking of basal bodies in ciliated epithelial cells |
| Q96RT7 | Gamma-tubulin complex component 6 OS=Homo sapiens GN=TUBGCP6 PE=1 SV=3 - [GCP6_HUMAN] | Gamma-tubulin complex is necessary for **microtubule nucleation at the centrosome.** | 10.1091/mbc.12.11.3340. GCP5 and GCP6: two new members of the human gamma-tubulin complex |
| Q76N32 | Centrosomal protein of 68 kDa OS=Homo sapiens GN=CEP68 PE=1 SV=2 - [CEP68_HUMAN] | **Involved in maintenance of centrosome cohesion**, probably as part of a linker structure which prevents centrosome splitting (PubMed:18042621). Required for localization of CDK5RAP2 to the centrosome during interphase | 10.1242/jcs.020248. Cep68 and Cep215 (Cdk5rap2) are required for centrosome cohesion. 10.1242/jcs.139451. Centlein mediates an interaction between C-Nap1 and Cep68 to maintain centrosome cohesion. 10.1038/ncb3076. Degradation of Cep68 and PCNT cleavage mediate Cep215 removal from the PCM to allow **centriole separation, disengagement and licensing**. |
| Q6ZVD7 | Storkhead-box protein 1 OS=Homo sapiens GN=STOX1 PE=1 SV=2 - [STOX1_HUMAN] | **Concentrated at centrosomes** during metaphase. Involved in regulating the levels of reactive oxidative species and reactive nitrogen species and in mitochondrial homeostasis in the placenta (PubMed:24738702). Required for regulation of inner ear epithelial cell proliferation via the AKT signaling pathway | 10.1371/journal.pone.0029769. Transcription factor STOX1A promotes mitotic entry by binding to the CCNB1 promotor |
| Q9H0N0 | Ras-related protein Rab-6C OS=Homo sapiens GN=RAB6C PE=1 SV=2 - [RAB6C_HUMAN] | May be involved in the **regulation of centrosome duplication** and cell cycle progression. | 10.1016/j.jmb.2010.01.009. RAB6C is a retrogene that encodes a centrosomal protein involved in cell cycle progression |
| Q8N8E3 | Centrosomal protein of 112 kDa OS=Homo sapiens GN=CEP112 PE=1 SV=2 - [CE112_HUMAN] | **Identified as a centrosomal protein.** | 10.1038/emboj.2011.63. Novel asymmetrically localizing components of human centrosomes identified by complementary proteomics methods |
| A2RUB6 | Coiled-coil domain-containing protein 66 OS=Homo sapiens GN=CCDC66 PE=1 SV=4 - [CCD66_HUMAN] | Microtubule-binding protein required for ciliogenesis (PubMed:28235840). May function in ciliogenesis by mediating the transport of proteins like BBS4 to the cilium, but also through the organization of the **centriolar satellites** (PubMed:28235840). Plays a role in retina morphogenesis and/or homeostasis | 10.1242/jcs.196832. The centriolar satellite protein CCDC66 interacts with CEP290 and functions in cilium formation and trafficking. |
| O15013 | Rho guanine nucleotide exchange factor 10 OS=Homo sapiens GN=ARHGEF10 PE=1 SV=4 - [ARHGA_HUMAN] | **Localize to centrosomes in HeLa cells** | 10.1186/1471-2121-10-56. Regulation of mitotic spindle formation by the RhoA guanine nucleotide exchange factor ARHGEF10 |
| Q96JB5 | CDK5 regulatory subunit-associated protein 3 OS=Homo sapiens GN=CDK5RAP3 PE=1 SV=2 - [CK5P3_HUMAN] | A **portion of C53 protein is localized at the centrosome**, and centrosome-targeting C53 potently promotes local Cdk1 activation. Substrate adapter for ufmylation, the covalent attachment of the ubiquitin-like modifier UFM1 to substrate proteins, in response to endoplasmic reticulum stress (PubMed:23152784, PubMed:30635284). Negatively regulates NF-kappa-B-mediated gene transcription through the control of RELA phosphorylation (PubMed:17785205, PubMed:20228063). Probable tumor suppressor initially identified as a CDK5R1 interactor controlling cell proliferation (PubMed:12054757, PubMed:12737517). Also regulates mitotic G2/M transition checkpoint and mitotic G2 DNA damage checkpoint (PubMed:15790566, PubMed:19223857). Through its interaction with CDKN2A/ARF and MDM2 may induce MDM2-dependent p53/TP53 ubiquitination, stabilization and activation in the nucleus, thereby promoting G1 cell cycle arrest and inhibition of cell proliferation (PubMed:16173922). May also play a role in the rupture of the nuclear envelope during apoptosis (PubMed:23478299). May regulate MAPK14 activity by regulating its dephosphorylation by PPM1D/WIP1 (PubMed:21283629). Required for liver development. | 10.1038/cr.2009.14. Tumor suppressor protein C53 antagonizes checkpoint kinases to promote cyclin-dependent kinase 1 activation |
| Q8NHQ1 | Centrosomal protein of 70 kDa OS=Homo sapiens GN=CEP70 PE=1 SV=2 - [CEP70_HUMAN] | Plays a role in the organization of both preexisting and nascent microtubules in interphase cells. During mitosis, required for the organization and orientation of the mitotic spindle. | 10.1038/nature02166. Proteomic characterization of the human centrosome by protein correlation profiling. 10.1074/jbc.M111.252262. **CEP70 protein interacts with γ-tubulin to localize at the centrosome** and is critical for mitotic spindle assembly |
| Q9NXG0 | Centlein OS=Homo sapiens GN=CNTLN PE=2 SV=5 - [CNTLN_HUMAN] | Required for **centrosome cohesion and recruitment of CEP68 to centrosomes**. | 10.1242/jcs.139451. Centlein mediates an interaction between C-Nap1 and Cep68 to maintain centrosome cohesion |
| Q9BSJ2 | Gamma-tubulin complex component 2 OS=Homo sapiens GN=TUBGCP2 PE=1 SV=2 - [GCP2_HUMAN] | Gamma-tubulin complex is necessary for **microtubule nucleation at the centrosome**. | 10.1038/nature02166. Proteomic characterization of the human centrosome by protein correlation profiling |
| P06493 | Cyclin-dependent kinase 1 OS=Homo sapiens GN=CDK1 PE=1 SV=3 - [CDK1_HUMAN] | Plays a key role in the control of the eukaryotic cell cycle by **modulating the centrosome cycle** as well as mitotic onset; promotes G2-M transition, and regulates G1 progress and G1-S transition via association with multiple interphase cyclins. | 10.1038/nature02166. Proteomic characterization of the human centrosome by protein correlation profiling |
| P08F94 | Fibrocystin OS=Homo sapiens GN=PKHD1 PE=1 SV=1 - [PKHD1_HUMAN] | May be required for correct bipolar cell division through the **regulation of centrosome duplication** and mitotic spindle assembly. May be a receptor protein that acts in collecting-duct and biliary differentiation. | 10.1093/hmg/ddq233. Polycystic kidney disease protein fibrocystin localizes to the mitotic spindle and regulates spindle bipolarity. 10.1097/01.asn.0000113793.12558.1d. The autosomal recessive polycystic kidney disease protein is localized to primary cilia, with concentration in the basal body area |
| Q96CN5 | Leucine-rich repeat-containing protein 45 OS=Homo sapiens GN=LRRC45 PE=1 SV=1 - [LRC45_HUMAN] | Component of the proteinaceous fiber-like linker between **two centrioles**, required for centrosome cohesion. | 10.1016/j.celrep.2013.08.005. LRRC45 is a centrosome linker component required for centrosome cohesion |
| O94927 | HAUS augmin-like complex subunit 5 OS=Homo sapiens GN=HAUS5 PE=1 SV=2 - [HAUS5_HUMAN] | Contributes to mitotic spindle assembly, maintenance of centrosome integrity and completion of cytokinesis as part of the HAUS augmin-like complex. | 10.1016/j.cub.2009.04.033. **HAUS, the 8-subunit human Augmin complex, regulates centrosome** and spindle integrity. 10.1073/pnas.0901587106. The augmin complex plays a critical role in spindle microtubule generation for mitotic progression and cytokinesis in human cells |
| Q9BVT8 | Transmembrane and ubiquitin-like domain-containing protein 1 OS=Homo sapiens GN=TMUB1 PE=1 SV=1 - [TMUB1_HUMAN] | Involved in sterol-regulated ubiquitination and degradation of HMG-CoA reductase HMGCR (By similarity). Involved in positive regulation of AMPA-selective glutamate receptor GRIA2 recycling to the cell surface (By similarity). Acts as negative regulator of hepatocyte growth during regeneration. When clevelad into iHOPS, it is involved in centrosome assembly | 10.4161/cc.7.10.5882. **HOPS is an essential constituent of centrosome assembly** |
| Q6IQ55 | Tau-tubulin kinase 2 OS=Homo sapiens GN=TTBK2 PE=1 SV=2 - [TTBK2_HUMAN] | Serine/threonine kinase that acts as a key regulator of ciliogenesis: controls the initiation of ciliogenesis by **binding to the distal end of the basal body** and promoting the removal of CCP110, which caps the mother centriole, leading to the recruitment of IFT proteins, which build the ciliary axoneme. Has some substrate preference for proteins that are already phosphorylated on a Tyr residue at the +2 position relative to the phosphorylation site. Able to phosphorylate tau on serines in vitro. | 10.1016/j.cell.2012.10.010. The spinocerebellar ataxia-associated gene Tau tubulin kinase 2 controls the initiation of ciliogenesis. 10.1042/BJ20110276. TTBK2 kinase substrate specificity and the impact of spinocerebellar-ataxia-causing mutations on expression, activity, localization and development |
| O95359 | Transforming acidic coiled-coil-containing protein 2 OS=Homo sapiens GN=TACC2 PE=1 SV=3 - [TACC2_HUMAN] | Plays a role in the microtubule-dependent **coupling of the nucleus and the centrosome**. Involved in the processes that regulate centrosome-mediated interkinetic nuclear migration (INM) of neural progenitors (By similarity). May play a role in organizing centrosomal microtubules. May act as a tumor suppressor protein. May represent a tumor progression marker. | 10.1016/j.febslet.2004.06.092. TTK kinase is essential for the centrosomal localization of TACC2 |
| Q9UFC0 | Leucine-rich repeat and WD repeat-containing protein 1 OS=Homo sapiens GN=LRWD1 PE=1 SV=2 - [LRWD1_HUMAN] | **Localizes to the centrosome and interacts with gamma-tubulin** in mouse spermatozoa. Required for G1/S transition. Recruits and stabilizes the origin recognition complex (ORC) onto chromatin during G1 to establish pre-replication complex (preRC) and to heterochromatic sites in post-replicated cells. Binds a combination of DNA and histone methylation repressive marks on heterochromatin. Binds histone H3 and H4 trimethylation marks H3K9me3, H3K27me3 and H4K20me3 in a cooperative manner with DNA methylation (By similarity). Required for silencing of major satellite repeats. May be important ORC2, ORC3 and ORC4 stability. | 10.1111/j.1365-2605.2009.01038.x. Expression of lrwd1 in mouse testis and its centrosomal localization |
| Q86VQ0 | Lebercilin OS=Homo sapiens GN=LCA5 PE=1 SV=2 - [LCA5_HUMAN] | In non- ciliated cells, **localizes to the centrosome** and its associated microtubule array. Involved in intraflagellar protein **(IFT) transport** in photoreceptor cilia. | 10.1038/ng2066. Mutations in LCA5, encoding the ciliary protein lebercilin, cause Leber congenital amaurosis. |
| Q9P2P6 | StAR-related lipid transfer protein 9 OS=Homo sapiens GN=STARD9 PE=1 SV=3 - [STAR9_HUMAN] | Microtubule-dependent motor protein required for spindle pole assembly during mitosis. Required to stabilize the pericentriolar material (PCM). Localizes throughout the cytoplasm and nucleus during interphase. **Localizes to the daughter centriole** during mitosis. Disappears in cytokinesis. | 10.1016/j.cell.2011.11.020. The STARD9/Kif16a kinesin associates with mitotic microtubules and regulates spindle pole assembly. |
| Q6IN85 | Serine/threonine-protein phosphatase 4 regulatory subunit 3A OS=Homo sapiens GN=SMEK1 PE=1 SV=1 - [P4R3A_HUMAN] | In interphase localized in the cytoplasm and in the nucleus (with higher levels). During metaphase located in pericentriolar regions. Regulatory subunit of serine/threonine-protein phosphatase 4. **May regulate the activity of PPP4C at centrosomal microtubule organizing centers**. The PPP4C-PPP4R2-PPP4R3A PP4 complex specifically dephosphorylates H2AX phosphorylated on 'Ser-140' (gamma-H2AX) generated during DNA replication and required for DNA DSB repair. | 10.1016/j.biocel.2008.03.021. Depletion of protein phosphatase 4 in human cells reveals essential roles in centrosome maturation, cell migration and the regulation of Rho GTPases. |
| Q96JN8 | Neuralized-like protein 4 OS=Homo sapiens GN=NEURL4 PE=1 SV=2 - [NEUL4_HUMAN] | Promotes CCP110 ubiquitination and proteasome-dependent degradation. By counteracting accumulation of CP110, maintains normal centriolar homeostasis and preventing formation of ectopic microtubular organizing centers. **Localizes to procentriole and daughter centriole** in growing and quiescent cells (PubMed:22441691). May loose association with centrosomes during mitosis (PubMed:22261722). | 10.1038/embor.2012.40. Neurl4, a novel daughter centriole protein, prevents formation of ectopic microtubule organizing centres. 10.1074/mcp.M111.014233. Interaction proteomics identify NEURL4 and the HECT E3 ligase HERC2 as novel modulators of centrosome architecture. |
| Q7Z3E5 | LisH domain-containing protein ARMC9 OS=Homo sapiens GN=ARMC9 PE=1 SV=2 - [ARMC9_HUMAN] | Localized to the proximal region in cilia. Stimulation of Hh signaling leads to redistribution of ARMC9 toward the ciliary tip within 6 hours, follow by a gradual return to its original proximal location (By similarity). **Localizes to the daughter centriole of the primary cilium** in RPE1 cells. Acts as a positive regulator of hedgehog (Hh)signaling (By similarity). Involved in ciliogenesis (By similarity). May participate in the trafficking and/or retention of GLI2 and GLI3 proteins at the ciliary tip (By similarity). | 10.1016/j.ajhg.2017.05.010. Mutations in ARMC9, which Encodes a Basal Body Protein, Cause Joubert Syndrome in Humans and Ciliopathy Phenotypes in Zebrafish |
| Q8TDX7 | Serine/threonine-protein kinase Nek7 OS=Homo sapiens GN=NEK7 PE=1 SV=1 - [NEK7_HUMAN] | Protein kinase which plays an important role in mitotic cell cycle progression. Required for **microtubule nucleation activity of the centrosome**, robust mitotic spindle formation and cytokinesis. Phosphorylates RPS6KB1 | 10.1016/j.febslet.2006.10.069. **Nek7 kinase is enriched at the centrosome**, and is required for proper spindle assembly and mitotic progression. 10.1016/j.bbrc.2007.05.206. NEK7 is a centrosomal kinase critical for microtubule nucleation. |
| P50402 | Emerin OS=Homo sapiens GN=EMD PE=1 SV=1 - [EMD_HUMAN] | Stabilizes and promotes the formation of a nuclear actin cortical network. Stimulates actin polymerization in vitro by binding and stabilizing the pointed end of growing filaments. Inhibits beta-catenin activity by preventing its accumulation in the nucleus. Acts by influencing the nuclear accumulation of beta-catenin through a CRM1-dependent export pathway. Links centrosomes to the nuclear envelope via a microtubule association. EMD and BAF are cooperative cofactors of HIV-1 infection. Association of EMD with the viral DNA requires the presence of BAF and viral integrase. The association of viral DNA with chromatin requires the presence of BAF and EMD. Required for proper localization of non-farnesylated prelamin-A/C. | 10.1083/jcb.200702026. **A novel role for the nuclear membrane protein emerin in association of the centrosome** to the outer nuclear membrane. |
| Q8TD16 | Protein bicaudal D homolog 2 OS=Homo sapiens GN=BICD2 PE=1 SV=1 - [BICD2_HUMAN] | Acts as an adapter protein linking the dynein motor complex to various cargos and converts dynein from a non- processive to a highly processive motor in the presence of dynactin. Facilitates and stabilizes the interaction between dynein and dynactin and activates dynein processivity (the ability to move along a microtubule for a long distance without falling off the track) (By similarity). Facilitates the binding of RAB6A to the Golgi by stabilizing its GTP-bound form. Regulates coat complex coatomer protein I (COPI)-independent Golgi-endoplasmic reticulum. **Plays a role in regulating centrosome localization at G2** [...] *[a.k.a. KIAA0699, AAH73970.1, Q9H561, Q8TD16]* | 10.1371/journal.pbio.1000350. Bicaudal D2, dynein, and kinesin-1 associate with nuclear pore complexes and regulate centrosome and nuclear positioning during mitotic entry. 10.1091/mbc.E15-07-0482. Lamins position the nuclear pores and centrosomes by modulating dynein. |
| Q9P1Y5 | Calmodulin-regulated spectrin-associated protein 3 OS=Homo sapiens GN=CAMSAP3 PE=1 SV=2 - [CAMP3_HUMAN] | **Localizes at the PCM. Releases microtubules from the centrosome.** Key microtubule-organizing protein that specifically binds the minus-end of non-centrosomal microtubules and regulates their dynamics and organization. Specifically recognizes growing microtubule minus-ends and autonomously decorates and stabilizes microtubule lattice formed by microtubule minus-end polymerization. Acts on free microtubule minus-ends that are not capped by microtubule-nucleating proteins or other factors and protects microtubule minus-ends from depolymerization. In addition, it also reduces the velocity of microtubule [...] *[a.k.a. KIAA1543, PPP1R80, AF289580, Q9P1Y5]* | 10.1242/jcs.198010. CAMSAP3 accumulates in the pericentrosomal area and accompanies microtubule release from the centrosome via katanin. |
| Q99661 | Kinesin-like protein KIF2C OS=Homo sapiens GN=KIF2C PE=1 SV=2 - [KIF2C_HUMAN] | In complex with KIF18B, constitutes the major microtubule plus-end depolymerizing activity in mitotic cells. Regulates the turnover of microtubules at the kinetochore and functions in chromosome segregation during mitosis. Plays a role in chromosome congression and is required for the lateral to end-on conversion of the chromosome- microtubule attachment, and **centrosome separation.** Belongs to the TRAFAC class myosin-kinesin ATPase superfamily. Kinesin family. MCAK/KIF2 subfamily [a.k.a. KNSL6, Kinesin-like protein KIF2C, Q5JR89, Q99661] | 10.1083/jcb.201902109. Cell polarity-dependent centrosome separation in the C. elegans embryo. 10.1007/s00412-016-0607-4. Aurora A, MCAK, and Kif18b promote Eg5-independent spindle formation. |
| O00139 | Kinesin-like protein KIF2A OS=Homo sapiens GN=KIF2A PE=1 SV=3 - [KIF2A_HUMAN] | Plus end-directed microtubule-dependent motor required for normal brain development. May regulate microtubule dynamics during axonal growth. Required for normal progression through mitosis. Required for normal congress of chromosomes at the metaphase plate. Required for normal spindle dynamics during mitosis. Promotes spindle turnover. **Implicated in formation of bipolar mitotic spindles. Interacts with CEP85L and localizes to the centriole. It also localizes to the cilium basal body.** Has microtubule depolymerization activity; Kinesins [a.k.a. KIF2, KNS2, CDCBM3, O00139] | 10.1016/j.neuron.2020.01.030. Posterior Neocortex-Specific Regulation of Neuronal Migration by CEP85L Identifies Maternal Centriole-Dependent Activation of CDK5. 10.1038/s41467-019-10497-2. Modeling microcephaly with cerebral organoids reveals a WDR62-CEP170-KIF2A pathway promoting cilium disassembly in neural progenitors. |

**Centriole**
Centrosome

#### SUPPMENTARY TABLE 2, SHEET 4

| 2 unique peptides: | |
|---|---|
| Accession | Description |
| Q9NQR4 | Omega-amidase NIT2 OS=Homo sapiens GN=NIT2 PE=1 SV=1 - [NIT2_HUMAN] |
| Q8TCU4 | Alstrom syndrome protein 1 OS=Homo sapiens GN=ALMS1 PE=1 SV=3 - [ALMS1_HUMAN] |
| Q5HYA8 | Meckelin OS=Homo sapiens GN=TMEM67 PE=1 SV=2 - [MKS3_HUMAN] |
| Q5VYK3 | Proteasome-associated protein ECM29 homolog OS=Homo sapiens GN=ECM29 PE=1 SV=2 - [ECM29_HUMAN] |
| O60733 | 85/88 kDa calcium-independent phospholipase A2 OS=Homo sapiens GN=PLA2G6 PE=1 SV=2 - [PLPL9_HUMAN] |
| Q14008 | Cytoskeleton-associated protein 5 OS=Homo sapiens GN=CKAP5 PE=1 SV=3 - [CKAP5_HUMAN]. |
| Q9NTK5 | Obg-like ATPase 1 OS=Homo sapiens GN=OLA1 PE=1 SV=2 - [OLA1_HUMAN] |
| Q9Y3P9 | Rab GTPase-activating protein 1 OS=Homo sapiens GN=RABGAP1 PE=1 SV=3 - [RBGP1_HUMAN] |
| Q8TF46 | DIS3-like exonuclease 1 OS=Homo sapiens GN=DIS3L PE=1 SV=2 - [DI3L1_HUMAN] |
| Q8IVU9 | Uncharacterized protein C10orf107 OS=Homo sapiens GN=C10orf107 PE=2 SV=1 - [CJ107_HUMAN] |
| Q15691 | Microtubule-associated protein RP/EB family member 1 OS=Homo sapiens GN=MAPRE1 PE=1 SV=3 - [MARE1_HUMAN] |
| Q5SW79 | Centrosomal protein of 170 kDa OS=Homo sapiens GN=CEP170 PE=1 SV=1 - [CE170_HUMAN] |
| Q8WXH0 | Nesprin-2 OS=Homo sapiens GN=SYNE2 PE=1 SV=3 - [SYNE2_HUMAN] |
| Q66GS9 | Centrosomal protein of 135 kDa OS=Homo sapiens GN=CEP135 PE=1 SV=2 - [CP135_HUMAN] |
| P23258 | Tubulin gamma-1 chain OS=Homo sapiens GN=TUBG1 PE=1 SV=2 - [TBG1_HUMAN] |
| O75116 | Rho-associated protein kinase 2 OS=Homo sapiens GN=ROCK2 PE=1 SV=4 - [ROCK2_HUMAN] |
| Q8TC44 | POC1 centriolar protein homolog B OS=Homo sapiens GN=POC1B PE=1 SV=1 - [POC1B_HUMAN] |
| Q99996 | A-kinase anchor protein 9 OS=Homo sapiens GN=AKAP9 PE=1 SV=3 - [AKAP9_HUMAN] |
| P42025 | Beta-centractin OS=Homo sapiens GN=ACTR1B PE=1 SV=1 - [ACTY_HUMAN] |
| O43237 | Cytoplasmic dynein 1 light intermediate chain 2 OS=Homo sapiens GN=DYNC1LI2 PE=1 SV=1 - [DC1L2_HUMAN] |
| P60510 | Serine/threonine-protein phosphatase 4 catalytic subunit OS=Homo sapiens GN=PPP4C PE=1 SV=1 - [PP4C_HUMAN] |
| A6NC98 | Coiled-coil domain-containing protein 88B OS=Homo sapiens GN=CCDC88B PE=1 SV=1 - [CC88B_HUMAN] |
| O15078 | Centrosomal protein of 290 kDa OS=Homo sapiens GN=CEP290 PE=1 SV=2 - [CE290_HUMAN] |
| Q15154 | Pericentriolar material 1 protein OS=Homo sapiens GN=PCM1 PE=1 SV=4 - [PCM1_HUMAN] |
| Q8TCT7 | Signal peptide peptidase-like 2B OS=Homo sapiens GN=SPPL2B PE=1 SV=2 - [SPP2B_HUMAN] |
| P36404 | ADP-ribosylation factor-like protein 2 OS=Homo sapiens GN=ARL2 PE=1 SV=4 - [ARL2_HUMAN] |
| Q8TCI5 | Protein pitchfork OS=Homo sapiens GN=PIFO PE=1 SV=2 - [PIFO_HUMAN] |
| O75665 | Oral-facial-digital syndrome 1 protein OS=Homo sapiens GN=OFD1 PE=1 SV=1 - [OFD1_HUMAN] |
| Q8TAP6 | Centrosomal protein of 76 kDa OS=Homo sapiens GN=CEP76 PE=1 SV=1 - [CEP76_HUMAN] |
| O94886 | CSC1-like protein 1 OS=Homo sapiens GN=TMEM63A PE=1 SV=3 - [CSCL1_HUMAN] |
| Q9UNZ2 | NSFL1 cofactor p47 OS=Homo sapiens GN=NSFL1C PE=1 SV=2 - [NSF1C_HUMAN] |
| P37198 | Nuclear pore glycoprotein p62 OS=Homo sapiens GN=NUP62 PE=1 SV=3 - [NUP62_HUMAN] |
| Q9Y6G9 | Cytoplasmic dynein 1 light intermediate chain 1 OS=Homo sapiens GN=DYNC1LI1 PE=1 SV=3 - [DC1L1_HUMAN] |
| Q13561 | Dynactin subunit 2 OS=Homo sapiens GN=DCTN2 PE=1 SV=4 - [DCTN2_HUMAN] |
| P63167 | Dynein light chain 1, cytoplasmic OS=Homo sapiens GN=DYNLL1 PE=1 SV=1 - [DYL1_HUMAN] |
| P36405 | ADP-ribosylation factor-like protein 3 OS=Homo sapiens GN=ARL3 PE=1 SV=2 - [ARL3_HUMAN] |
| Q9Y230 | RuvB-like 2 OS=Homo sapiens GN=RUVBL2 PE=1 SV=3 - [RUVB2_HUMAN] |
| O43379 | WD repeat-containing protein 62 OS=Homo sapiens GN=WDR62 PE=1 SV=4 - [WDR62_HUMAN] |
| P35222 | Catenin beta-1 OS=Homo sapiens GN=CTNNB1 PE=1 SV=1 - [CTNB1_HUMAN] |
| P28074 | Proteasome subunit beta type-5 OS=Homo sapiens GN=PSMB5 PE=1 SV=3 - [PSB5_HUMAN] |
| Q14203 | Dynactin subunit 1 OS=Homo sapiens GN=DCTN1 PE=1 SV=3 - [DCTN1_HUMAN] |
| Q14204 | Cytoplasmic dynein 1 heavy chain 1 OS=Homo sapiens GN=DYNC1H1 PE=1 SV=5 - [DYHC1_HUMAN] |
| Q9BZV1 | UBX domain-containing protein 6 OS=Homo sapiens GN=UBXN6 PE=1 SV=1 - [UBXN6_HUMAN] |
| O60271 | C-Jun-amino-terminal kinase-interacting protein 4 OS=Homo sapiens GN=SPAG9 PE=1 SV=4 - [JIP4_HUMAN] |
| Q13464 | Rho-associated protein kinase 1 OS=Homo sapiens GN=ROCK1 PE=1 SV=1 - [ROCK1_HUMAN] |
| Q12798 | Centrin-1 OS=Homo sapiens GN=CETN1 PE=1 SV=1 - [CETN1_HUMAN] |
| Q9UJW0 | Dynactin subunit 4 OS=Homo sapiens GN=DCTN4 PE=1 SV=1 - [DCTN4_HUMAN] |
| Q9NRG9 | Aladin OS=Homo sapiens GN=AAAS PE=1 SV=1 - [AAAS_HUMAN] |
| P43034 | Platelet-activating factor acetylhydrolase IB subunit alpha OS=Homo sapiens GN=PAFAH1B1 PE=1 SV=2 - [LIS1_HUMAN] |
| Q9BTW9 | Tubulin-specific chaperone D OS=Homo sapiens GN=TBCD PE=1 SV=2 - [TBCD_HUMAN] |
| Q5T4S7 | E3 ubiquitin-protein ligase UBR4 OS=Homo sapiens GN=UBR4 PE=1 SV=1 - [UBR4_HUMAN] |
| P06748 | Nucleophosmin OS=Homo sapiens GN=NPM1 PE=1 SV=2 - [NPM_HUMAN] |
| P53384 | Cytosolic Fe-S cluster assembly factor NUBP1 OS=Homo sapiens GN=NUBP1 PE=1 SV=2 - [NUBP1_HUMAN] |
| P33176 | Kinesin-1 heavy chain OS=Homo sapiens GN=KIF5B PE=1 SV=1 - [KINH_HUMAN] |
| Q9H0I3 | Coiled-coil domain-containing protein 113 OS=Homo sapiens GN=CCDC113 PE=1 SV=1 - [CC113_HUMAN] |
| Q9C099 | Leucine-rich repeat and coiled-coil domain-containing protein 1 OS=Homo sapiens GN=LRRCC1 PE=1 SV=2 - [LRCC1_HUMAN] |
| Q5BJF6 | Outer dense fiber protein 2 OS=Homo sapiens GN=ODF2 PE=1 SV=1 - [ODFP2_HUMAN] |
| P61163 | Alpha-centractin OS=Homo sapiens GN=ACTR1A PE=1 SV=1 - [ACTZ_HUMAN] |
| Q13045 | Protein flightless-1 homolog OS=Homo sapiens GN=FLII PE=1 SV=2 - [FLII_HUMAN] |

| | |
|---|---|
| Q9Y5B8 | Nucleoside diphosphate kinase 7 OS=Homo sapiens GN=NME7 PE=1 SV=1 - [NDK7_HUMAN] |
| P08754 | Guanine nucleotide-binding protein G(k) subunit alpha OS=Homo sapiens GN=GNAI3 PE=1 SV=3 - [GNAI3_HUMAN] |
| Q8ND07 | Coiled-coil domain-containing protein 176 OS=Homo sapiens GN=CCDC176 PE=2 SV=3 - [CC176_HUMAN] |
| P08107;P0DI | Heat shock 70 kDa protein 1A/1B OS=Homo sapiens GN=HSPA1A PE=1 SV=5 - [HSP71_HUMAN] |
| Q9H5N1 | Rab GTPase-binding effector protein 2 OS=Homo sapiens GN=RABEP2 PE=1 SV=2 - [RABE2_HUMAN] |
| Q86WT1 | Tetratricopeptide repeat protein 30A OS=Homo sapiens GN=TTC30A PE=2 SV=3 - [TT30A_HUMAN] |
| Q8WYA0 | Intraflagellar transport protein 81 homolog OS=Homo sapiens GN=IFT81 PE=1 SV=1 - [IFT81_HUMAN] |
| Q13485 | Mothers against decapentaplegic homolog 4 OS=Homo sapiens GN=SMAD4 PE=1 SV=1 - [SMAD4_HUMAN] |
| Q14145 | Kelch-like ECH-associated protein 1 OS=Homo sapiens GN=KEAP1 PE=1 SV=2 - [KEAP1_HUMAN] |

#### 1 unique peptide:

| Accession | Description |
|---|---|
| Q8TD16 | Protein bicaudal D homolog 2 OS=Homo sapiens GN=BICD2 PE=1 SV=1 - [BICD2_HUMAN] |
| Q9P1Y5 | Calmodulin-regulated spectrin-associated protein 3 OS=Homo sapiens GN=CAMSAP3 PE=1 SV=2 - [CAMP3_HUMAN] |
| Q86YT6 | E3 ubiquitin-protein ligase MIB1 OS=Homo sapiens GN=MIB1 PE=1 SV=1 - [MIB1_HUMAN] |
| O95613 | Pericentrin OS=Homo sapiens GN=PCNT PE=1 SV=4 - [PCNT_HUMAN] |
| Q92830 | Histone acetyltransferase KAT2A OS=Homo sapiens GN=KAT2A PE=1 SV=3 - [KAT2A_HUMAN] |
| P63096 | Guanine nucleotide-binding protein G(i) subunit alpha-1 OS=Homo sapiens GN=GNAI1 PE=1 SV=2 - [GNAI1_HUMAN] |
| Q99661 | Kinesin-like protein KIF2C OS=Homo sapiens GN=KIF2C PE=1 SV=2 - [KIF2C_HUMAN] |
| Q96RK4 | Bardet-Biedl syndrome 4 protein OS=Homo sapiens GN=BBS4 PE=1 SV=2 - [BBS4_HUMAN] |
| Q53EZ4 | Centrosomal protein of 55 kDa OS=Homo sapiens GN=CEP55 PE=1 SV=3 - [CEP55_HUMAN] |
| Q9NY27 | Serine/threonine-protein phosphatase 4 regulatory subunit 2 OS=Homo sapiens GN=PPP4R2 PE=1 SV=3 - [PP4R2_HUMAN] |
| Q16513 | Serine/threonine-protein kinase N2 OS=Homo sapiens GN=PKN2 PE=1 SV=1 - [PKN2_HUMAN] |
| O95271 | Tankyrase-1 OS=Homo sapiens GN=TNKS PE=1 SV=2 - [TNKS1_HUMAN] |
| Q96CW5 | Gamma-tubulin complex component 3 OS=Homo sapiens GN=TUBGCP3 PE=1 SV=2 - [GCP3_HUMAN] |
| Q9UBK9 | Protein UXT OS=Homo sapiens GN=UXT PE=1 SV=1 - [UXT_HUMAN] |
| Q9NQC7 | Ubiquitin carboxyl-terminal hydrolase CYLD OS=Homo sapiens GN=CYLD PE=1 SV=1 - [CYLD_HUMAN] |
| Q96RT7 | Gamma-tubulin complex component 6 OS=Homo sapiens GN=TUBGCP6 PE=1 SV=3 - [GCP6_HUMAN] |
| Q76N32 | Centrosomal protein of 68 kDa OS=Homo sapiens GN=CEP68 PE=1 SV=2 - [CEP68_HUMAN] |
| Q15398 | Disks large-associated protein 5 OS=Homo sapiens GN=DLGAP5 PE=1 SV=2 - [DLGP5_HUMAN] |
| Q9H0N0 | Ras-related protein Rab-6C OS=Homo sapiens GN=RAB6C PE=1 SV=2 - [RAB6C_HUMAN] |
| Q8N8E3 | Centrosomal protein of 112 kDa OS=Homo sapiens GN=CEP112 PE=1 SV=2 - [CE112_HUMAN] |
| A2RUB6 | Coiled-coil domain-containing protein 66 OS=Homo sapiens GN=CCDC66 PE=1 SV=4 - [CCD66_HUMAN] |
| O00139 | Kinesin-like protein KIF2A OS=Homo sapiens GN=KIF2A PE=1 SV=3 - [KIF2A_HUMAN] |
| Q3YEC7 | Rab-like protein 6 OS=Homo sapiens GN=RABL6 PE=1 SV=2 - [RABL6_HUMAN] |
| Q8NEZ2 | Vacuolar protein sorting-associated protein 37A OS=Homo sapiens GN=VPS37A PE=1 SV=1 - [VP37A_HUMAN] |
| Q8NHQ1 | Centrosomal protein of 70 kDa OS=Homo sapiens GN=CEP70 PE=1 SV=2 - [CEP70_HUMAN] |
| Q9NXG0 | Centlein OS=Homo sapiens GN=CNTLN PE=2 SV=5 - [CNTLN_HUMAN] |
| P62491 | Ras-related protein Rab-11A OS=Homo sapiens GN=RAB11A PE=1 SV=3 - [RB11A_HUMAN] |
| Q9BSJ2 | Gamma-tubulin complex component 2 OS=Homo sapiens GN=TUBGCP2 PE=1 SV=2 - [GCP2_HUMAN] |
| O43809 | Cleavage and polyadenylation specificity factor subunit 5 OS=Homo sapiens GN=NUDT21 PE=1 SV=1 - [CPSF5_HUMAN] |
| P06493 | Cyclin-dependent kinase 1 OS=Homo sapiens GN=CDK1 PE=1 SV=3 - [CDK1_HUMAN] |
| P08F94 | Fibrocystin OS=Homo sapiens GN=PKHD1 PE=1 SV=1 - [PKHD1_HUMAN] |
| Q96CN5 | Leucine-rich repeat-containing protein 45 OS=Homo sapiens GN=LRRC45 PE=1 SV=1 - [LRC45_HUMAN] |
| O94927 | HAUS augmin-like complex subunit 5 OS=Homo sapiens GN=HAUS5 PE=1 SV=2 - [HAUS5_HUMAN] |
| Q9BXF6 | Rab11 family-interacting protein 5 OS=Homo sapiens GN=RAB11FIP5 PE=1 SV=1 - [RFIP5_HUMAN] |
| Q9H1Z4 | WD repeat-containing protein 13 OS=Homo sapiens GN=WDR13 PE=1 SV=2 - [WDR13_HUMAN] |
| Q6IQ55 | Tau-tubulin kinase 2 OS=Homo sapiens GN=TTBK2 PE=1 SV=2 - [TTBK2_HUMAN] |
| O95359 | Transforming acidic coiled-coil-containing protein 2 OS=Homo sapiens GN=TACC2 PE=1 SV=3 - [TACC2_HUMAN] |
| Q9UPY8 | Microtubule-associated protein RP/EB family member 3 OS=Homo sapiens GN=MAPRE3 PE=1 SV=1 - [MARE3_HUMAN] |
| Q04759 | Protein kinase C theta type OS=Homo sapiens GN=PRKCQ PE=1 SV=3 - [KPCT_HUMAN] |
| Q86VQ0 | Lebercilin OS=Homo sapiens GN=LCA5 PE=1 SV=2 - [LCA5_HUMAN] |
| O75923 | Dysferlin OS=Homo sapiens GN=DYSF PE=1 SV=1 - [DYSF_HUMAN] |
| Q03518 | Antigen peptide transporter 1 OS=Homo sapiens GN=TAP1 PE=1 SV=2 - [TAP1_HUMAN] |
| O00592 | Podocalyxin OS=Homo sapiens GN=PODXL PE=1 SV=2 - [PODXL_HUMAN] |
| Q96EX3 | WD repeat-containing protein 34 OS=Homo sapiens GN=WDR34 PE=1 SV=2 - [WDR34_HUMAN] |
| Q6IN85 | Serine/threonine-protein phosphatase 4 regulatory subunit 3A OS=Homo sapiens GN=SMEK1 PE=1 SV=1 - [P4R3A_HUMAN] |
| Q96JN8 | Neuralized-like protein 4 OS=Homo sapiens GN=NEURL4 PE=1 SV=2 - [NEUL4_HUMAN] |
| Q8TDX7 | Serine/threonine-protein kinase Nek7 OS=Homo sapiens GN=NEK7 PE=1 SV=1 - [NEK7_HUMAN] |
| P50402 | Emerin OS=Homo sapiens GN=EMD PE=1 SV=1 - [EMD_HUMAN] |

#### SUPPMENTARY TABLE 2, SHEET 5

| 2 unique peptides: | |
|---|---|
| Accession | Description |
| Q9H7X7 | Intraflagellar transport protein 22 homolog OS=Homo sapiens GN=IFT22 PE=2 SV=1 - [IFT22_HUMAN] |
| Q9UPT5 | Exocyst complex component 7 OS=Homo sapiens GN=EXOC7 PE=1 SV=3 - [EXOC7_HUMAN] |
| P42858 | Huntingtin OS=Homo sapiens GN=HTT PE=1 SV=2 - [HD_HUMAN] |
| P22694 | cAMP-dependent protein kinase catalytic subunit beta OS=Homo sapiens GN=PRKACB PE=1 SV=2 - [KAPCB_HUMAN] |
| Q02241 | Kinesin-like protein KIF23 OS=Homo sapiens GN=KIF23 PE=1 SV=3 - [KIF23_HUMAN] |
| Q15018 | BRISC complex subunit Abro1 OS=Homo sapiens GN=FAM175B PE=1 SV=2 - [F175B_HUMAN] |
| A0AVF1 | Intraflagellar transport protein 56 OS=Homo sapiens GN=TTC26 PE=2 SV=1 - [IFT56_HUMAN] |
| Q96AJ1 | Clusterin-associated protein 1 OS=Homo sapiens GN=CLUAP1 PE=1 SV=4 - [CLUA1_HUMAN] |
| Q9BW83 | Intraflagellar transport protein 27 homolog OS=Homo sapiens GN=IFT27 PE=1 SV=1 - [IFT27_HUMAN] |
| P49768 | Presenilin-1 OS=Homo sapiens GN=PSEN1 PE=1 SV=1 - [PSN1_HUMAN] |
| Q3SYG4 | Protein PTHB1 OS=Homo sapiens GN=BBS9 PE=1 SV=1 - [PTHB1_HUMAN] |
| Q9UPM9 | B9 domain-containing protein 1 OS=Homo sapiens GN=B9D1 PE=2 SV=1 - [B9D1_HUMAN] |
| Q9Y366 | Intraflagellar transport protein 52 homolog OS=Homo sapiens GN=IFT52 PE=2 SV=3 - [IFT52_HUMAN] |
| Q9NWB7 | Intraflagellar transport protein 57 homolog OS=Homo sapiens GN=IFT57 PE=1 SV=1 - [IFT57_HUMAN] |
| Q9ULC3 | Ras-related protein Rab-23 OS=Homo sapiens GN=RAB23 PE=1 SV=1 - [RAB23_HUMAN] |
| P11388 | DNA topoisomerase 2-alpha OS=Homo sapiens GN=TOP2A PE=1 SV=3 - [TOP2A_HUMAN] |
| P49841 | Glycogen synthase kinase-3 beta OS=Homo sapiens GN=GSK3B PE=1 SV=2 - [GSK3B_HUMAN] |
| O75604 | Ubiquitin carboxyl-terminal hydrolase 2 OS=Homo sapiens GN=USP2 PE=1 SV=2 - [UBP2_HUMAN] |
| Q9P2H3 | Intraflagellar transport protein 80 homolog OS=Homo sapiens GN=IFT80 PE=1 SV=3 - [IFT80_HUMAN] |
| Q92834 | X-linked retinitis pigmentosa GTPase regulator OS=Homo sapiens GN=RPGR PE=1 SV=2 - [RPGR_HUMAN] |
| Q9Y5K8 | V-type proton ATPase subunit D OS=Homo sapiens GN=ATP6V1D PE=1 SV=1 - [VATD_HUMAN] |
| Q9BVG8 | Kinesin-like protein KIFC3 OS=Homo sapiens GN=KIFC3 PE=1 SV=4 - [KIFC3_HUMAN] |
| Q08211 | ATP-dependent RNA helicase A OS=Homo sapiens GN=DHX9 PE=1 SV=4 - [DHX9_HUMAN] |
| Q99828 | Calcium and integrin-binding protein 1 OS=Homo sapiens GN=CIB1 PE=1 SV=4 - [CIB1_HUMAN] |
| Q96RY7 | Intraflagellar transport protein 140 homolog OS=Homo sapiens GN=IFT140 PE=1 SV=1 - [IF140_HUMAN] |
| Q8IWZ6 | Bardet-Biedl syndrome 7 protein OS=Homo sapiens GN=BBS7 PE=1 SV=2 - [BBS7_HUMAN] |
| Q8TDR0 | TRAF3-interacting protein 1 OS=Homo sapiens GN=TRAF3IP1 PE=1 SV=1 - [MIPT3_HUMAN] |
| P30622 | CAP-Gly domain-containing linker protein 1 OS=Homo sapiens GN=CLIP1 PE=1 SV=2 - [CLIP1_HUMAN] |
| P17987 | T-complex protein 1 subunit alpha OS=Homo sapiens GN=TCP1 PE=1 SV=1 - [TCPA_HUMAN] |
| P25786 | Proteasome subunit alpha type-1 OS=Homo sapiens GN=PSMA1 PE=1 SV=1 - [PSA1_HUMAN] |
| Q9Y6A4 | UPF0468 protein C16orf80 OS=Homo sapiens GN=C16orf80 PE=1 SV=1 - [CP080_HUMAN] |
| P29966 | Myristoylated alanine-rich C-kinase substrate OS=Homo sapiens GN=MARCKS PE=1 SV=4 - [MARCS_HUMAN] |
| P61006 | Ras-related protein Rab-8A OS=Homo sapiens GN=RAB8A PE=1 SV=1 - [RAB8A_HUMAN] |
| O75955 | Flotillin-1 OS=Homo sapiens GN=FLOT1 PE=1 SV=3 - [FLOT1_HUMAN] |

| | |
|---|---|
| P17612 | cAMP-dependent protein kinase catalytic subunit alpha OS=Homo sapiens GN=PRKACA PE=1 SV=2 - [KAPCA_HUMAN] |
| P50990 | T-complex protein 1 subunit theta OS=Homo sapiens GN=CCT8 PE=1 SV=4 - [TCPQ_HUMAN] |
| P50991 | T-complex protein 1 subunit delta OS=Homo sapiens GN=CCT4 PE=1 SV=4 - [TCPD_HUMAN] |
| Q9HD42 | Charged multivesicular body protein 1a OS=Homo sapiens GN=CHMP1A PE=1 SV=1 - [CHM1A_HUMAN] |
| P62826 | GTP-binding nuclear protein Ran OS=Homo sapiens GN=RAN PE=1 SV=3 - [RAN_HUMAN] |
| P15311 | Ezrin OS=Homo sapiens GN=EZR PE=1 SV=4 - [EZRI_HUMAN] |
| P13861 | cAMP-dependent protein kinase type II-alpha regulatory subunit OS=Homo sapiens GN=PRKAR2A PE=1 SV=2 - [KAP2_HUMAN] |
| P05783 | Keratin, type I cytoskeletal 18 OS=Homo sapiens GN=KRT18 PE=1 SV=2 - [K1C18_HUMAN] |
| P43487 | Ran-specific GTPase-activating protein OS=Homo sapiens GN=RANBP1 PE=1 SV=1 - [RANG_HUMAN] |
| P40121 | Macrophage-capping protein OS=Homo sapiens GN=CAPG PE=1 SV=2 - [CAPG_HUMAN] |
| Q9Y547 | Heat shock protein beta-11 OS=Homo sapiens GN=HSPB11 PE=1 SV=1 - [HSB11_HUMAN] |
| P61421 | V-type proton ATPase subunit d 1 OS=Homo sapiens GN=ATP6V0D1 PE=1 SV=1 - [VA0D1_HUMAN] |
| P50570 | Dynamin-2 OS=Homo sapiens GN=DNM2 PE=1 SV=2 - [DYN2_HUMAN] |
| P53985 | Monocarboxylate transporter 1 OS=Homo sapiens GN=SLC16A1 PE=1 SV=3 - [MOT1_HUMAN] |
| P48643 | T-complex protein 1 subunit epsilon OS=Homo sapiens GN=CCT5 PE=1 SV=1 - [TCPE_HUMAN] |
| Q9UJC3 | Protein Hook homolog 1 OS=Homo sapiens GN=HOOK1 PE=1 SV=2 - [HOOK1_HUMAN] |
| Q96LB3 | Intraflagellar transport protein 74 homolog OS=Homo sapiens GN=IFT74 PE=1 SV=1 - [IFT74_HUMAN] |
| P15531 | Nucleoside diphosphate kinase A OS=Homo sapiens GN=NME1 PE=1 SV=1 - [NDKA_HUMAN] |
| O15066 | Kinesin-like protein KIF3B OS=Homo sapiens GN=KIF3B PE=1 SV=1 - [KIF3B_HUMAN] |
| O95721 | Synaptosomal-associated protein 29 OS=Homo sapiens GN=SNAP29 PE=1 SV=1 - [SNP29_HUMAN] |
| Q6NXR4 | TELO2-interacting protein 2 OS=Homo sapiens GN=TTI2 PE=1 SV=1 - [TTI2_HUMAN] |

#### 1 unique peptide:

| Accession | Description |
|---|---|
| Q9P219 | Protein Daple OS=Homo sapiens GN=CCDC88C PE=1 SV=3 - [DAPLE_HUMAN] |
| Q92845 | Kinesin-associated protein 3 OS=Homo sapiens GN=KIFAP3 PE=1 SV=2 - [KIFA3_HUMAN] |
| Q9P2G4 | Microtubule-associated protein 10 OS=Homo sapiens GN=MAP10 PE=1 SV=2 - [MAP10_HUMAN] |
| Q3V6T2 | Girdin OS=Homo sapiens GN=CCDC88A PE=1 SV=2 - [GRDN_HUMAN] |
| Q9NQC8 | Intraflagellar transport protein 46 homolog OS=Homo sapiens GN=IFT46 PE=1 SV=1 - [IFT46_HUMAN] |
| Q14191 | Werner syndrome ATP-dependent helicase OS=Homo sapiens GN=WRN PE=1 SV=2 - [WRN_HUMAN] |
| Q9UQB9 | Aurora kinase C OS=Homo sapiens GN=AURKC PE=1 SV=1 - [AURKC_HUMAN] |
| P14635 | G2/mitotic-specific cyclin-B1 OS=Homo sapiens GN=CCNB1 PE=1 SV=1 - [CCNB1_HUMAN] |
| O14974 | Protein phosphatase 1 regulatory subunit 12A OS=Homo sapiens GN=PPP1R12A PE=1 SV=1 - [MYPT1_HUMAN] |
| P12004 | Proliferating cell nuclear antigen OS=Homo sapiens GN=PCNA PE=1 SV=1 - [PCNA_HUMAN] |
| P05062 | Fructose-bisphosphate aldolase B OS=Homo sapiens GN=ALDOB PE=1 SV=2 - [ALDOB_HUMAN] |
| O75351 | Vacuolar protein sorting-associated protein 4B OS=Homo sapiens GN=VPS4B PE=1 SV=2 - [VPS4B_HUMAN] |
| O95714 | E3 ubiquitin-protein ligase HERC2 OS=Homo sapiens GN=HERC2 PE=1 SV=2 - [HERC2_HUMAN] |
| Q8TD31 | Coiled-coil alpha-helical rod protein 1 OS=Homo sapiens GN=CCHCR1 PE=1 SV=2 - [CCHCR_HUMAN] |

#### SUPPLEMENTARY TABLE 3

| Uniprot code | Name | Uniprot information (localization) |
|---|---|---|
| H3BRN8 | C15orf65 | - |
| Q53QW1 | TEX44 | Cytoplasm |
| Q5JU67 | CFAP157 | Cilium basal body |
| Q5T5A4 | C1orf194 | - |
| **Q5T681** | **C10orf62** | - |
| Q5TEZ5 | C6orf163 | - |
| Q5VTT2 | C9orf135 | Cell membrane and cytoplasm |
| **Q5VU69** | **C1orf189** | - |
| Q6P656 | CFAP161 | - |
| Q6V702 | C4orf22 | - |
| Q6ZQR2 | CFAP77 | Cilium localization |
| Q8IZ16 | C7orf61 | Nucleus |
| Q8N1D5 | C1orf158 | - |
| Q8N801 | STPG4 | Nucleus and cytoplasm |
| **Q8N865** | **C7orf31** | Centrosome in bovine sperm |
| Q8NA69 | TEX45 | - |
| **Q8NEP4** | **C17orf47** | - |
| **Q8WW14** | **C10orf82** | - |
| Q96LM5 | C4orf45 | - |
| **Q96M34** | **C3orf30** | - |
| **Q9H0B3** | **KIAA1683** | Mitochondirion and Nucleus |
| **Q9H1P6** | **C20orf85** | - |
| A4D263 | C7orf72 | - |
| **A4QMS7** | **C5orf49** | - |
| A6NCJ1 | C19orf71 | - |
| **B2RV13** | **C17orf105** | - |

**SUPPLEMENTARY TABLE 4**

| Accession | Description | ΣCoverage | Σ# Proteins | Unique Peptic | Σ# Peptides | Σ# PSMs | Exp1_01 | | | Exp02_01 | | | Exp03_01 | | |
|---|---|---|---|---|---|---|---|---|---|---|---|---|---|---|---|
| | | | | | | | Coverage A4 | # Peptides A4 | # PSM A4 | Coverage B4 | # Peptides B4 | # PSM B4 | Coverage C4 | # Peptides C4 | # PSM C4 |
| Q9NRG9 | Aladin OS=Hc | 42.86 | 1 | 15 | 15 | 28 | 0 | 0 | 0 | 9.89 | 4 | 6 | 0 | 0 | 0 |
| P61163 | Alpha-centrac | 61.17 | 1 | 10 | 18 | 69 | 34.31 | 9 | 11 | 31.65 | 8 | 11 | 34.31 | 7 | 8 |
| P42025 | Beta-centracti | 48.94 | 1 | 6 | 14 | 33 | 11.7 | 4 | 4 | 9.04 | 3 | 3 | 11.7 | 3 | 3 |
| P36404 | ADP-ribosylati | 50 | 1 | 7 | 7 | 10 | 0 | 0 | 0 | 5.98 | 1 | 1 | 0 | 0 | 0 |
| P36405 | ADP-ribosylati | 71.98 | 1 | 12 | 12 | 27 | 6.04 | 1 | 1 | 6.04 | 1 | 1 | 0 | 0 | 0 |
| P61421 | V-type proton | 43.59 | 2 | 11 | 11 | 24 | 13.11 | 3 | 3 | 0 | 0 | 0 | 3.7 | 1 | 1 |
| Q9Y5K8 | V-type proton | 31.58 | 1 | 6 | 6 | 7 | 0 | 0 | 0 | 0 | 0 | 0 | 0 | 0 | 0 |
| P50991 | T-complex prc | 73.65 | 1 | 38 | 39 | 263 | 31.91 | 12 | 15 | 39.33 | 18 | 22 | 30.98 | 15 | 17 |
| P48643 | T-complex prc | 71.72 | 1 | 42 | 43 | 346 | 53.42 | 23 | 31 | 53.23 | 26 | 46 | 40.11 | 19 | 25 |
| P50990 | T-complex prc | 72.26 | 1 | 45 | 45 | 322 | 45.26 | 21 | 24 | 51.46 | 26 | 32 | 49.27 | 26 | 32 |
| P60953 | Cell division c | 62.83 | 3 | 8 | 9 | 37 | 14.14 | 2 | 3 | 0 | 0 | 0 | 36.65 | 5 | 5 |
| Q14008 | Cytoskeleton- | 1.87 | 1 | 3 | 3 | 5 | 0 | 0 | 0 | 0 | 0 | 0 | 0 | 0 | 0 |
| P35222 | Catenin beta- | 17.8 | 1 | 7 | 11 | 32 | 0 | 0 | 0 | 0 | 0 | 0 | 0 | 0 | 0 |
| Q14203 | Dynactin subu | 50.47 | 2 | 57 | 58 | 161 | 10.09 | 10 | 12 | 10.72 | 11 | 12 | 5.01 | 5 | 5 |
| Q13561 | Dynactin subu | 46.63 | 1 | 17 | 17 | 68 | 27.43 | 8 | 8 | 31.42 | 9 | 11 | 17.46 | 5 | 5 |
| Q14204 | Cytoplasmic d | 47.27 | 3 | 200 | 201 | 618 | 0.65 | 3 | 3 | | | | 5.47 | 21 | 21 |
| Q9Y6G9 | Cytoplasmic d | 18.36 | 1 | 8 | 8 | 19 | 1.72 | 1 | 1 | 0 | 0 | 0 | 1.72 | 1 | 1 |
| P63167 | Dynein light cl | 65.17 | 1 | 4 | 7 | 108 | 59.55 | 6 | 8 | 59.55 | 4 | 6 | 58.43 | 5 | 6 |
| P15311 | Ezrin OS=Hor | 54.95 | 2 | 25 | 38 | 182 | 18.26 | 11 | 13 | 21.5 | 13 | 16 | 31.06 | 17 | 17 |
| P04899 | Guanine nucl | 56.06 | 4 | 10 | 17 | 92 | 23.38 | 6 | 7 | 0 | 0 | 0 | 34.37 | 9 | 11 |
| P08754 | Guanine nucl | 38.98 | 4 | 6 | 12 | 65 | 7.34 | 2 | 2 | 0 | 0 | 0 | 12.43 | 4 | 5 |
| Q9UJC3 | Protein Hook | 39.56 | 1 | 26 | 26 | 63 | 0 | 0 | 0 | 0 | 0 | 0 | 0 | 0 | 0 |
| P33176 | Kinesin-1 hea | 46.42 | 1 | 29 | 41 | 111 | 2.28 | 2 | 2 | 4.26 | 3 | 3 | 4.05 | 3 | 3 |
| Q15691 | Microtubule-a | 23.13 | 1 | 4 | 4 | 4 | 0 | 0 | 0 | 0 | 0 | 0 | 0 | 0 | 0 |
| P29966 | Myristoylated | 33.73 | 2 | 6 | 6 | 23 | 20.78 | 4 | 4 | 9.64 | 2 | 2 | 15.06 | 3 | 3 |
| Q9NQR4 | Omega-amida | 18.12 | 1 | 4 | 4 | 5 | 0 | 0 | 0 | 0 | 0 | 0 | 0 | 0 | 0 |
| P06748 | Nucleophosm | 12.93 | 2 | 5 | 5 | 10 | 0 | 0 | 0 | 6.8 | 2 | 3 | 0 | 0 | 0 |
| Q9UNZ2 | NSFL1 cofact | 40 | 1 | 12 | 12 | 33 | 17.03 | 5 | 5 | 12.97 | 4 | 4 | 7.03 | 2 | 2 |
| P43034 | Platelet-activa | 41.95 | 1 | 16 | 17 | 69 | 1.71 | 1 | 1 | 10.98 | 4 | 4 | 24.88 | 8 | 8 |
| Q8WUM4 | Programmed ( | 56.91 | 1 | 48 | 48 | 208 | 13.36 | 9 | 9 | 8.06 | 5 | 5 | 12.44 | 10 | 10 |
| P13861 | cAMP-depend | 78.96 | 2 | 33 | 33 | 242 | 57.43 | 19 | 23 | 50.74 | 18 | 22 | 41.34 | 13 | 18 |
| P28074 | Proteasome s | 61.22 | 2 | 15 | 15 | 50 | 32.32 | 7 | 7 | 26.62 | 7 | 7 | 33.46 | 7 | 9 |
| P61006 | Ras-related p | 48.79 | 7 | 4 | 12 | 86 | 24.64 | 4 | 5 | 30.43 | 5 | 6 | 27.54 | 5 | 5 |
| P62826 | GTP-binding | 51.39 | 1 | 12 | 12 | 72 | 9.26 | 2 | 2 | 18.98 | 4 | 4 | 18.98 | 4 | 4 |
| P43487 | Ran-specific ( | 43.28 | 2 | 7 | 7 | 22 | 5.47 | 1 | 1 | 5.47 | 1 | 1 | 8.96 | 2 | 2 |
| P62081 | 40S ribosoma | 60.82 | 1 | 10 | 10 | 24 | 5.67 | 1 | 1 | 10.31 | 2 | 2 | 0 | 0 | 0 |
| Q9Y265 | RuvB-like 1 O | 70.61 | 1 | 28 | 28 | 142 | 39.04 | 11 | 12 | 44.3 | 15 | 19 | 39.25 | 14 | 17 |
| Q9Y230 | RuvB-like 2 O | 73.87 | 1 | 40 | 40 | 260 | 34.13 | 14 | 16 | 50.54 | 21 | 26 | 49.46 | 19 | 21 |
| P53985 | Monocarboxyl | 17.8 | 1 | 8 | 8 | 37 | 2.4 | 1 | 1 | 11 | 6 | 9 | 9 | 3 | 3 |
| Q9BTW9 | Tubulin-specil | 30.2 | 1 | 27 | 27 | 57 | 0 | 0 | 0 | 2.1 | 2 | 2 | 0 | 0 | 0 |
| P17987 | T-complex prc | 78.78 | 1 | 37 | 37 | 295 | 44.96 | 17 | 22 | 53.42 | 22 | 26 | 42.45 | 17 | 21 |
| P23258 | Tubulin gamn | 39.02 | 2 | 9 | 9 | 18 | 0 | 0 | 0 | 0 | 0 | 0 | 0 | 0 | 0 |

| Accession | Exp01_02 | | | Exp02_02 | | | Exp03_02 | | |
|---|---|---|---|---|---|---|---|---|---|
| | Coverage D4 | # Peptides D4 | # PSM D4 | Coverage E4 | # Peptides E4 | # PSM E4 | Coverage F4 | # Peptides F4 | # PSM F4 |
| Q9NRG9 | 0 | 0 | 0 | 0 | 0 | 0 | 0 | 0 | 0 |
| P61163 | 12.5 | 3 | 3 | 0 | 0 | 0 | 0 | 0 | 0 |
| P42025 | 6.65 | 2 | 2 | 0 | 0 | 0 | 0 | 0 | 0 |
| P36404 | 0 | 0 | 0 | 0 | 0 | 0 | 0 | 0 | 0 |
| P36405 | 0 | 0 | 0 | 0 | 0 | 0 | 0 | 0 | 0 |
| P61421 | 5.41 | 2 | 2 | 0 | 0 | 0 | 0 | 0 | 0 |
| Q9Y5K8 | 0 | 0 | 0 | 0 | 0 | 0 | 0 | 0 | 0 |
| P50991 | 25.60 | 11 | 14 | 26.16 | 12 | 15 | 5.19 | 3 | 3 |
| P48643 | 44.73 | 23 | 30 | 31.79 | 18 | 23 | | | |
| P50990 | 61.86 | 32 | 38 | 42.70 | 24 | 26 | | | |
| P60953 | 16.75 | 3 | 3 | 10.99 | 2 | 2 | 0 | 0 | 0 |
| Q14008 | 0 | 0 | 0 | 0 | 0 | 0 | 0 | 0 | 0 |
| P35222 | 0 | 0 | 0 | 0.9 | 1 | 1 | 0 | 0 | 0 |
| Q14203 | 4.46 | 5 | 6 | 4.3 | 5 | 5 | 0 | 0 | 0 |
| Q13561 | 19.2 | 6 | 6 | 20.2 | 7 | 10 | 0 | 0 | 0 |
| Q14204 | 4.20 | 18 | 18 | 0.24 | 1 | 1 | | | |
| Q9Y6G9 | 1.72 | 1 | 1 | 0 | 0 | 0 | 0 | 0 | 0 |
| P63167 | 59.55 | 6 | 11 | 59.55 | 5 | 10 | 0 | 0 | 0 |
| P15311 | 14.51 | 8 | 8 | 12.12 | 7 | 8 | 0 | 0 | 0 |
| P04899 | 20.28 | 6 | 6 | 3.1 | 1 | 1 | 0 | 0 | 0 |
| P08754 | 15.25 | 5 | 5 | 3.11 | 1 | 1 | 0 | 0 | 0 |
| Q9UJC3 | 2.47 | 1 | 1 | 3.43 | 2 | 2 | 0 | 0 | 0 |
| P33176 | 4.26 | 3 | 3 | 2.91 | 2 | 2 | 0 | 0 | 0 |
| Q15691 | 0 | 0 | 0 | 0 | 0 | 0 | 0 | 0 | 0 |
| P29966 | 0 | 0 | 0 | 0 | 0 | 0 | 0 | 0 | 0 |
| Q9NQR4 | 0 | 0 | 0 | 0 | 0 | 0 | 0 | 0 | 0 |
| P06748 | 0 | 0 | 0 | 12.93 | 4 | 4 | 0 | 0 | 0 |
| Q9UNZ2 | 7.84 | 2 | 2 | 2.7 | 1 | 1 | 0 | 0 | 0 |
| P43034 | 11.95 | 4 | 4 | 9.27 | 3 | 3 | 0 | 0 | 0 |
| Q8WUM4 | 12.10 | 9 | 9 | 1.27 | 1 | 1 | | | |
| P13861 | 28.22 | 9 | 10 | 56.19 | 20 | 26 | | | |
| P28074 | 4.56 | 1 | 1 | 4.56 | 1 | 1 | 0 | 0 | 0 |
| P61006 | 26.09 | 5 | 5 | 0 | 0 | 0 | 0 | 0 | 0 |
| P62826 | 14.81 | 3 | 3 | 13.89 | 3 | 3 | 0 | 0 | 0 |
| P43487 | 0 | 0 | 0 | 9.95 | 2 | 2 | 0 | 0 | 0 |
| P62081 | 0 | 0 | 0 | 0 | 0 | 0 | 0 | 0 | 0 |
| Q9Y265 | 24.34 | 10 | 11 | 33.55 | 13 | 15 | 5.7 | 2 | 2 |
| Q9Y230 | 43.84 | 18 | 20 | 44.71 | 18 | 22 | 0 | 0 | 0 |
| P53985 | 0 | 0 | 0 | 0 | 0 | 0 | 0 | 0 | 0 |
| Q9BTW9 | 0 | 0 | 0 | 0 | 0 | 0 | 0 | 0 | 0 |
| P17987 | 43.88 | 19 | 20 | 33.27 | 14 | 19 | | | |
| P23258 | 0 | 0 | 0 | 0 | 0 | 0 | 0 | 0 | 0 |

| Accession | Exp01_03 | | | Exp02_03 | | | Exp03_03 | | |
|---|---|---|---|---|---|---|---|---|---|
| | Coverage G4 | # Peptides G4 | # PSM G4 | Coverage H4 | # Peptides H4 | # PSM H4 | Coverage I4 | # Peptides I4 | # PSM I4 |
| Q9NRG9 | 0 | 0 | 0 | 0 | 0 | 0 | 0 | 0 | 0 |
| P61163 | 0 | 0 | 0 | 0 | 0 | 0 | 0 | 0 | 0 |
| P42025 | 0 | 0 | 0 | 0 | 0 | 0 | 0 | 0 | 0 |
| P36404 | 0 | 0 | 0 | 0 | 0 | 0 | 0 | 0 | 0 |
| P36405 | 0 | 0 | 0 | 4.4 | 1 | 1 | 22.53 | 3 | 3 |
| P61421 | 0 | 0 | 0 | 0 | 0 | 0 | 0 | 0 | 0 |
| Q9Y5K8 | 0 | 0 | 0 | 0 | 0 | 0 | 0 | 0 | 0 |
| P50991 | 6.31 | 3 | 3 | 3.90 | 2 | 3 | 15.03 | 7 | 8 |
| P48643 | 15.53 | 9 | 9 | 16.45 | 9 | 11 | 32.53 | 17 | 23 |
| P50990 | 13.69 | 6 | 7 | 12.41 | 6 | 7 | 32.66 | 16 | 19 |
| P60953 | 5.76 | 1 | 1 | 5.24 | 1 | 1 | 33.51 | 5 | 5 |
| Q14008 | 0 | 0 | 0 | 0 | 0 | 0 | 0 | 0 | 0 |
| P35222 | 0.9 | 1 | 1 | 0 | 0 | 0 | 2.43 | 2 | 2 |
| Q14203 | 3.6 | 4 | 4 | 6.26 | 7 | 9 | 12.52 | 13 | 14 |
| Q13561 | 0 | 0 | 0 | 6.23 | 2 | 2 | 17.96 | 5 | 5 |
| Q14204 | 0.60 | 3 | 3 | 0.30 | 1 | 1 | 4.52 | 18 | 20 |
| Q9Y6G9 | 4.59 | 2 | 2 | 0 | 0 | 0 | 4.02 | 2 | 2 |
| P63167 | 46.07 | 4 | 8 | 59.55 | 4 | 7 | 59.55 | 4 | 10 |
| P15311 | 3.75 | 2 | 2 | 1.54 | 1 | 1 | 6.31 | 4 | 4 |
| P04899 | 0 | 0 | 0 | 3.1 | 1 | 1 | 18.59 | 6 | 8 |
| P08754 | 0 | 0 | 0 | 3.11 | 1 | 1 | 14.41 | 5 | 7 |
| Q9UJC3 | 14.29 | 9 | 9 | 3.85 | 2 | 2 | 14.7 | 8 | 8 |
| P33176 | 3.53 | 3 | 3 | 0 | 0 | 0 | 16.72 | 14 | 16 |
| Q15691 | 0 | 0 | 0 | 0 | 0 | 0 | 0 | 0 | 0 |
| P29966 | 0 | 0 | 0 | 0 | 0 | 0 | 0 | 0 | 0 |
| Q9NQR4 | 0 | 0 | 0 | 0 | 0 | 0 | 0 | 0 | 0 |
| P06748 | 0 | 0 | 0 | 0 | 0 | 0 | 0 | 0 | 0 |
| Q9UNZ2 | 12.16 | 4 | 4 | 16.22 | 4 | 4 | 12.97 | 4 | 4 |
| P43034 | 3.9 | 1 | 1 | 17.07 | 5 | 5 | 18.29 | 5 | 5 |
| Q8WUM4 | 12.21 | 10 | 11 | 3.57 | 3 | 3 | 25.23 | 18 | 22 |
| P13861 | 17.82 | 7 | 8 | 25.25 | 8 | 10 | 30.20 | 10 | 11 |
| P28074 | 0 | 0 | 0 | 4.56 | 1 | 1 | 0 | 0 | 0 |
| P61006 | 18.36 | 4 | 4 | 13.53 | 4 | 4 | 26.57 | 5 | 5 |
| P62826 | 4.63 | 1 | 1 | 10.19 | 2 | 2 | 15.74 | 3 | 3 |
| P43487 | 0 | 0 | 0 | 4.48 | 1 | 1 | 15.42 | 3 | 3 |
| P62081 | 0 | 0 | 0 | 0 | 0 | 0 | 0 | 0 | 0 |
| Q9Y265 | 7.24 | 3 | 3 | 17.32 | 6 | 7 | 7.68 | 2 | 2 |
| Q9Y230 | 27.86 | 11 | 12 | 34.99 | 15 | 18 | 32.83 | 12 | 14 |
| P53985 | 0 | 0 | 0 | 0 | 0 | 0 | 0 | 0 | 0 |
| Q9BTW9 | 0 | 0 | 0 | 0 | 0 | 0 | 0 | 0 | 0 |
| P17987 | 8.45 | 5 | 5 | 12.41 | 6 | 7 | 19.60 | 10 | 10 |
| P23258 | 0 | 0 | 0 | 0 | 0 | 0 | 0 | 0 | 0 |

| Accession | Exp04 | | | | | |
|---|---|---|---|---|---|---|
| | Coverage J4 | # Peptides J4 | # PSM J4 | # AAs | MW [kDa] | calc. pI |
| Q9NRG9 | 35.16 | 12 | 22 | 546 | 59.5363286 | 7.50341797 |
| P61163 | 47.34 | 14 | 36 | 376 | 42.5869075 | 6.63916016 |
| P42025 | 43.09 | 12 | 21 | 376 | 42.266726 | 6.40478516 |
| P36404 | 50 | 7 | 9 | 184 | 20.8648543 | 6.34130859 |
| P36405 | 71.43 | 11 | 21 | 182 | 20.44276 | 7.23974609 |
| P61421 | 43.59 | 11 | 18 | 351 | 40.3032185 | 4.99560547 |
| Q9Y5K8 | 31.58 | 6 | 7 | 247 | 28.245322 | 9.36376953 |
| P50991 | 66.05 | 33 | 163 | 539 | 57.9 | 7.83 |
| P48643 | 62.85 | 32 | 148 | 541 | 59.6 | 5.66 |
| P50990 | 66.24 | 38 | 137 | 548 | 59.6 | 5.60 |
| P60953 | 56.02 | 8 | 17 | 191 | 21.2450182 | 6.55126953 |
| Q14008 | 1.87 | 3 | 5 | 2032 | 225.351785 | 7.79638672 |
| P35222 | 15.36 | 9 | 28 | 781 | 85.4422164 | 5.85888672 |
| Q14203 | 45.54 | 47 | 94 | 1278 | 141.60708 | 5.80810547 |
| Q13561 | 35.66 | 11 | 21 | 401 | 44.2038581 | 5.21142578 |
| Q14204 | 45.16 | 190 | 551 | 4646 | 532.1 | 6.40 |
| Q9Y6G9 | 16.63 | 7 | 12 | 523 | 56.544003 | 6.41748047 |
| P63167 | 65.17 | 7 | 42 | 89 | 10.3591084 | 7.40087891 |
| P15311 | 44.88 | 28 | 113 | 586 | 69.369741 | 6.26513672 |
| P04899 | 51.27 | 14 | 58 | 355 | 40.4250561 | 5.54150391 |
| P08754 | 38.98 | 12 | 44 | 354 | 40.5062694 | 5.69384766 |
| Q9UJC3 | 33.38 | 20 | 41 | 728 | 84.5948379 | 5.14794922 |
| P33176 | 39.15 | 32 | 79 | 963 | 109.616994 | 6.50732422 |
| Q15691 | 23.13 | 4 | 4 | 268 | 29.9801912 | 5.13525391 |
| P29966 | 29.22 | 5 | 14 | 332 | 31.5358666 | 4.44970703 |
| Q9NQR4 | 18.12 | 4 | 5 | 276 | 30.5885573 | 7.21044922 |
| P06748 | 6.8 | 2 | 3 | 294 | 32.5548433 | 4.77978516 |
| Q9UNZ2 | 16.49 | 5 | 7 | 370 | 40.5483107 | 5.09716797 |
| P43034 | 36.83 | 15 | 38 | 410 | 46.6081598 | 7.37158203 |
| Q8WUM4 | 54.84 | 43 | 138 | 868 | 96.0 | 6.52 |
| P13861 | 73.27 | 25 | 114 | 404 | 45.5 | 5.07 |
| P28074 | 60.08 | 13 | 24 | 263 | 28.462211 | 6.91748047 |
| P61006 | 47.83 | 11 | 52 | 207 | 23.6531897 | 9.07080078 |
| P62826 | 50.93 | 11 | 50 | 216 | 24.4076189 | 7.48876953 |
| P43487 | 37.81 | 6 | 12 | 201 | 23.2955715 | 5.28759766 |
| P62081 | 60.82 | 10 | 21 | 194 | 22.113259 | 10.0961914 |
| Q9Y265 | 56.36 | 21 | 54 | 456 | 50.1963165 | 6.41748047 |
| Q9Y230 | 67.17 | 30 | 111 | 463 | 51.1245568 | 5.64306641 |
| P53985 | 15.8 | 6 | 24 | 500 | 53.9088535 | 8.66064453 |
| Q9BTW9 | 30.2 | 27 | 55 | 1192 | 132.515292 | 6.18896484 |
| P17987 | 71.58 | 31 | 165 | 556 | 60.3 | 6.11 |
| P23258 | 39.02 | 9 | 18 | 451 | 51.1377609 | 6.13818359 |

### SUPPLEMENTARY TABLE 4 SHEET 2

| | | | | | | | Exp1_01 | | | Exp02_01 | | | Exp03_01 | | | Exp01_02 | | | Exp02_02 | | | Exp03_02 | | | Exp01_03 | | | Exp02_03 | | | Exp03_03 | | | Exp04 | | | | | |
|---|---|---|---|---|---|---|---|---|---|---|---|---|---|---|---|---|---|---|---|---|---|---|---|---|---|---|---|---|---|---|---|---|---|---|---|---|---|---|---|
| Accession | Description | ΣCoverage | Σ# Proteins | Unique Pepti | Σ# Peptides | Σ# PSMs | Coverage A4 | # Peptides A4 | # PSM A4 | Coverage B4 | # Peptides B4 | # PSM B4 | Coverage C4 | # Peptides C4 | # PSM C4 | Coverage D4 | # Peptides D4 | # PSM D4 | Coverage E4 | # Peptides E4 | # PSM E4 | Coverage F4 | # Peptides F4 | # PSM F4 | Coverage G4 | # Peptides G4 | # PSM G4 | Coverage H4 | # Peptides H4 | # PSM H4 | Coverage I4 | # Peptides I4 | # PSM I4 | Coverage J4 | # Peptides J4 | # PSM J4 | # AAs | MW [kDa] | calc. pI |
| Q15398 | Disks large-as | 2.01 | 2 | 1 | 2 | 2 | | | | | | | | | | | | | | | | | | | | | | | | | | | | 2.01 | 2 | 2 | 846 | 95.1 | 9.00 |
| P63096 | Guanine nucl | 25.14 | 4 | 1 | 8 | 52 | 7.06 | 2 | 2 | | | | 12.43 | 4 | 5 | 12.43 | 4 | 4 | 3.11 | 1 | 1 | | | | | | | 3.11 | 1 | 1 | 14.41 | 5 | 7 | 21.19 | 7 | 32 | 354 | 40.3 | 5.97 |
| Q99661 | Kinesin-like p | 1.79 | 1 | 1 | 1 | 1 | 0.00 | | | | | | 0.00 | | | | | | 0.00 | | | | | | | | | 0.00 | | | 0.00 | | | 1.79 | 1 | 1 | 725 | 81.3 | 7.83 |
| P12004 | Proliferating c | 2.68 | 1 | 1 | 1 | 1 | | | | | | | | | | | | | | | | | | | | | | | | | | | | 2.68 | 1 | 1 | 261 | 28.8 | 4.69 |
| Q9BSJ2 | Gamma-tubul | 1.22 | 1 | 1 | 1 | 1 | | | | | | | | | | | | | | | | | | | | | | | | | | | | 1.22 | 1 | 1 | 902 | 102.5 | 6.84 |
| Q96CW5 | Gamma-tubul | 2.43 | 1 | 1 | 1 | 3 | | | | | | | | | | | | | | | | | | | | | | | | | | | | 2.43 | 1 | 3 | 907 | 103.5 | 8.12 |

#### SUPPLEMENTARY TABLE 5

| Sample | Pronucleus formation | Pronuclear Break Down | 2 cells | 3 cells | 4 cells | 5 cells | Compaction | Early Blastocyst | Expanded Blastocyst | IF | MTOC Signal |
|---|---|---|---|---|---|---|---|---|---|---|---|
| EXPERIMENT 1 | | | | | | | | | | | |
| Control_1 | no data | 65 | 71 | 104 | no data | 135 | 214 | | | YES | NO |
| Control_2 | no data | 62 | 71 | | | | | | | YES | NO |
| Control_3 | no data | 61 | 75 | 81 | | | | | | YES | NO |
| Control_4 | no data | 71 | 81 | 114 | 116 | 161 | | | | YES | NO |
| Control_5 | no data | 73 | | | | | | | | NO | |
| Injected_1 | 28 | 57 | 65 | 97 | 106 | 142 | | | | YES | NO |
| Injected_2 | 17 | 36 | 44 | | | | | | | YES | YES |
| Injected_3 | 14 | 33 | 43 | 69 | 71 | 96 | 232 | | | YES | YES |
| Injected_4 | 16 | 40 | 48 | 73 | 80 | 100 | 218 | 253 | 261 | YES | YES |
| Injected_5 | 15 | 39 | 47 | 51 | 57 | 89 | | | | YES | NO |
| Injected_6 | 18 | 42 | 49 | 80 | 115 | 188 | 219 | | | YES | YES |
| Injected_7 | no data | no data | 116 | 123 | | | | | | YES | NO |
| Injected_8 | 13 | 49 | 57 | 79 | 96 | 118 | 182 | 325 | 369 | YES | YES |
| EXPERIMENT 2 | | | | | | | | | | | |
| Control_1 | 18 | 45 | 54 | 68 | | | | | | NO | |
| Control_2 | 25 | 43 | 50 | 80 | 107 | 109 | 213 | 307 | 316 | YES | YES |
| Control_3 | 24 | 44 | 54 | 71 | 86 | 109 | 259 | 299 | | YES | YES |
| Control_4 | 16 | 40 | 46 | 78 | 83 | 106 | 223 | 282 | 362 | YES | YES |
| Control_5 | 19 | no data | 49 | 110 | 110 | 120 | | | | NO | |
| Injected_1 | 23 | 55 | 64 | 75 | 91 | 115 | 173 | | | YES | NO |
| Injected_2 | 20 | 50 | 56 | 88 | no data | 136 | 225 | 320 | | YES | YES |
| Injected_3 | 33 | 46 | 53 | | | | | | | YES | NO |
| Injected_4 | 28 | 44 | 58 | 85 | 88 | 125 | 176 | | | YES | NO |
| Injected_5 | no data | no data | 24 | 39 | | | | | | YES | NO |
| Injected_6 | 21 | 46 | 52 | 80 | 87 | 109 | 197 | 278 | 301 | YES | YES |
| Injected_7 | 21 | 61 | 67 | 72 | 98 | 105 | 210 | 316 | 338 | YES | YES |

#### SUPPLEMENTARY TABLE 6

| Sample | Pronucleus formation | Pronuclear Break Down | 2 cells | 3 cells | 4 cells | 5 cells | Compaction | Early Blastocyst | Expanded Blastocyst | IF | MTOC Signal |
|---|---|---|---|---|---|---|---|---|---|---|---|
| FIXED AT D+3 | | | | | | | | | | | |
| Control_1 | no data | no data | 35 | 51 | 54 | 54 | | | | NO | |
| Control_2 | 12 | 21 | 23 | | | | | | | YES | NO |
| Control_3 | 17 | 19 | 22 | 34 | 36 | 48 | | | | YES | NO |
| Control_4 | 9 | 17 | 20 | 36 | 39 | 57 | | | | YES | NO |
| Control_5 | 7 | 28 | 31 | | | | | | | YES | NO |
| Control_6 | 9 | 22 | 25 | 24 | | | | | | YES | NO |
| FIXED AT D+5 | | | | | | | | | | | |
| Control_1 | no data | no data | 21 | 22 | 46 | 53 | 63 | 93 | 102 | YES | YES |
| Control_2 | 10 | 26 | 31 | 44 | 44 | 56 | 83 | | | NO | |
| Control_3 | 7 | 19 | 22 | | | | | | | YES | NO |
| Control_4 | 11 | 23 | 25 | 28 | 43 | 63 | | | | YES | NO |
| Control_5 | no data | 22 | 24 | | | | | | | NO | |
